# Egg MVBs elicit an antimicrobial pathway to degrade paternal mitochondria after fertilization

**DOI:** 10.1101/2023.11.05.565307

**Authors:** Sharon Ben-Hur, Sara Afar, Yoav Politi, Liron Gal, Ofra Golani, Ehud Sivan, Rebecca Haffner-Krausz, Elina Maizenberg, Sima Peretz, Zvi Roth, Dorit Kalo, Nili Dezorella, David Morgenstern, Shmuel Pietrokovski, Keren Yacobi-Sharon, Eli Arama

## Abstract

Mitochondria are maternally inherited, but the mechanisms underlying paternal mitochondrial elimination (PME) after fertilization are far less clear. Using *Drosophila*, we show that special egg-derived multivesicular bodies (MVBs) promote PME by activating LC3-associated phagocytosis (LAP), a cellular defense pathway commonly employed against invading microbes. Upon fertilization, the egg MVBs engage and densely coat the sperm flagellum, forming extended flagellum vesicular sheaths (FVSs), within which the paternal mitochondria degrade. Inactivation of multiple LAP pathway components, such as Rubicon, a LAP-specific class III PI(3)K complex protein, significantly attenuates PME. Furthermore, recruitment of Atg8/LC3 to the FVS requires both Rubicon and the Atg8/LC3 conjugation machinery. Other LAP pathway events, such as production of the phospholipid PtdIns(3)P and reactive oxygen species (ROS), also unfold during PME. Finally, we provide evidence that a similar pathway might also mediate PME in mammals, highlighting the notion that eggs may regard paternal mitochondria as potentially dangerous trespassers.

## Introduction

Organisms as diverse as plants, fungi, protists, and animals differ greatly in external morphology, internal structure, and physiology, but when it comes to sexual mode of reproduction, they share multiple similar cellular and molecular patterns, such as meiosis, merger of the gametes, and a uniparental mode of mitochondrial inheritance ^1–9^. In mammals, the union of the gametes during fertilization occurs between a pair of highly distinct cells, the elongated male sperm, and the spherical female egg, each contributing half of the parental genomic material to the zygote ^10–12^. In contrast, only one parent, generally the mother, passes on the mitochondria to the offspring ^8,13,14^. Although this uniparental mode of inheritance is ubiquitous among eukaryotes, there are still abundant gaps in our knowledge concerning the molecular mechanisms underlying maternal mitochondrial inheritance, the fate of the paternal mitochondria (PM) after fertilization, and the rationale behind this highly conserved phenomenon ^8,9,13–15^. This ambiguity may be attributed to various factors, as well as to the widely received misconception that upon fertilization, sperm cells discard both the flagellum and the mitochondria, and that only the sperm head enters the egg^16^. In addition, it is difficult to exclude technical artefacts and false positive or negative findings due to assay sensitivity limits, such as assays for detection of scarce paternal mitochondrial (mt)DNA copies among the vast copy number of maternal mtDNA molecules ^17^. Moreover, such assays may be highly misleading, as in some organisms, and perhaps also in humans, elimination of the paternal mtDNA has been reported to be uncoupled from, and already occurring prior to, degradation of the paternal vacuolar mitochondria (i.e., mtDNA-free mitochondria) ^18–23^. The anatomical divergence in male gametes and their mitochondria among different organisms is another impeding factor, as this often hinders cross-study comparisons. Finally, although it is widely believed that the main purpose of paternal mitochondrial elimination (PME) is to prevent mtDNA heteroplasmy (i.e., a state of having more than one type of mtDNA in a cell), this is still a contentious issue, as the heteroplasmic state has been shown to be very unstable, and documented negative effects of paternal mtDNA leakage on the well-being of the organism are rare ^8,17,24–35.^

Two general theories have emerged to explain the mechanisms underlying PME after fertilization: A passive model of simple dilution of the PM by excess copy number of the egg mitochondria; and an active model, in which egg-derived mechanisms specifically target the PM for degradation. Both models have their drawbacks. The passive model is primarily based on two studies using mainly, albeit not exclusively, interspecific mice cross ^21,36^, an experimental setup which could account for the observed PM leakage in the progeny, as has been previously documented with murine and bovine hybrids ^37–39^. Another reason to be cautious about this model is that proteins which may imply involvement of active mechanisms in mediating PME, such as ubiquitin, P62, Atg8 or its mammalian ortholog LC3, all known to be involved in recycling of damaged mitochondria by selective autophagy (mitophagy), were shown to decorate the PM after fertilization in mice and other mammals ^21,40–42^. Moreover, genetic studies in the roundworm *Caenorhabditis elegans*, which constitute the basis for the active model, suggest that PME is mediated by mechanisms reminiscent of mitophagy ^18,41,43–48^. However, as opposed to flagellated sperm in mammals, in which the mitochondria undergo extensive structural organization and become part of the flagellum midpiece ^6,49^, the *C. elegans* amoeboid sperm lacks a flagellum and contains mitochondria of simple morphology, which can be accommodated by mitophagy ^50^.

Unlike *C. elegans*, *Drosophila melanogaster* flies produce unusually long sperm, consisting of an approximately 10 µm long needle-shaped nucleus and a ∼1.8 mm long flagellum (∼35 times the length of human sperm). The flagellum is mainly occupied by two major organelles: a single cylindrical mitochondrion (for convenience herein referred to as PM) which extends from the nucleus posterior edge along the entire length of the flagellum, and an intimately associated axoneme, the microtubule cytoskeleton which generates the propulsive force for sperm cell movement. The extraordinary size of the PM in *Drosophila*, that completely penetrates the egg, facilitates high spatiotemporal resolution of the PME process, with possible implications for PME in mammals ^5,51,52^. Indeed, we previously demonstrated that in *Drosophila*, PME is an active process mediated by two types of egg-derived vesicles: multivesicular bodies (MVBs) of the endocytic pathway, which are the first to engage and target large segments of the PM for degradation; and autophagosomes of the autophagy pathway, that recycle remnants of the PM ^53^. However, the pathway by which the MVBs mediate PME and whether a similar pathway also operates in mammals remain unknown.

Although originating in the endocytic pathway, the MVBs that are associated with the PM also display Atg8, a ubiquitin-like protein of the autophagy system, on their limiting membrane ^53^. In recent years, several related noncanonical autophagy pathways, which mediate <u>c</u>onjugation of <u>A</u>tg8 to <u>s</u>ingle <u>m</u>embranes (CASM), have emerged. As opposed to the double membrane autophagosomes that mediate classical autophagy, CASM pathways commonly utilize a subset of the autophagy machinery for Atg8 lipidation onto single-membrane vesicles ^54^. The most extensively investigated CASM pathway is LC3-associated phagocytosis (LAP), which is involved in clearance of different extracellular cargos by phagocytosis, including a variety of pathogenic microbes, as well as dying cells and axon debris, soluble ligands and protein aggregates, in different tissues and cell types ^55–65^. LAP is activated upon engagement of extracellular cargos with specific surface receptors, mainly but not exclusively Toll-like receptors (TLRs), triggering inward invagination of the plasma membrane and engulfment of the cargo into a single-membrane phagosome (also termed LAPosome). A LAP-specific class III PI(3)K complex (PI3KC3), composed of at least four proteins (Rubicon, Uvrag, Pi3k/Vps34 and Atg6/Beclin1), is then recruited to the LAPosome membrane, where it generates the phospholipid phosphatidylinositol-3-phosphate (PtdIns(3)P). Consequently, in a yet unknown mechanism, PtdIns(3)P recruits the Atg8/LC3 conjugating system, which in turn promotes recruitment of Atg8/LC3 to the LAPosomal membrane in a mechanism that requires the NADPH oxidase-2 (Nox2)-derived reactive oxygen species (ROS). Atg8/LC3 lipidation then facilitates fusion of the LAPosomes with lysosomes, promoting cargo digestion ^58,59,64,66,67^.

Here, to uncover the PM cellular degradative pathway mediated by the MVBs, we isolated these vesicles from early fertilized *Drosophila* eggs, using a subcellular fractionation procedure. Mass spectrometry (MS) analysis of the MVB-enriched fractions, followed by pathway enrichment analysis of the protein composition data, identified an innate immunity pathway with prevailing elements of phagocytosis. This, and the observation that Atg8 is expressed on the limiting membrane of the MVBs that associate with the PM, prompted us to explore the idea that the MVBs might mediate PME by activating a LAP-like CASM pathway. We demonstrate that mutants and RNAi knockdown of the LAP-specific PI3KC3 component, Rubicon, as well as downregulation of the other three main components of the complex, all significantly attenuate degradation of the PM. Maternal expression of a *rubicon-eGFP* rescue transgene and of tdTomato-tagged endogenous Rubicon in the early fertilized egg, revealed the presence of multiple MVBs expressing Rubicon mainly on their limiting membrane. We then show that immediately after fertilization, many of the Rubicon positive MVBs engage the sperm flagellum, creating extended vesicular sheaths around large flagellar segments, which display both PtdIns(3)P and ROS. Subsequently, Atg8 is recruited to the flagellum <u>v</u>esicular <u>s</u>heaths (FVSs) in a Rubicon- and Atg7-dependent manner, ultimately leading to degradation of the PM, but not the axoneme, within large Atg8 positive vesicles that bud off from the FVSs. These findings indicate that the egg activates a LAP-like cellular defense pathway against invading microbes to target the PM for degradation. Interestingly, we found that Rubicon is readily expressed on the flagellum of both mouse and bull sperm cells. Given previous observations of MVBs in the vicinity of the PM in both bovine and hamster fertilized eggs ^38,68^, these findings suggest that a similar pathway might also mediate PME in mammals.

## Results

### Egg-derived MVBs contain components of innate immunity with elements of phagocytosis

Degradation of the paternal mitochondria (PM) after fertilization in *Drosophila* is an efficient and rapid process, such that within only 1-hour after egg laying (AEL), the entire ∼1.8 mm long PM is eliminated (Supplementary Video 1). Note that time zero AEL is not equivalent to time zero after fertilization, as the latter occurs prior to egg laying, inside the female reproductive tract. A key step in the process of paternal mitochondrial elimination (PME) is the association of multiple egg-derived MVBs with the sperm flagellum, an event that gradually and asynchronously culminates in the degradation of large PM segments (illustrated in Fig. 1a) ^53^. Accordingly, inactivation of genes required for proper biogenesis of the intraluminal vesicles (ILVs), the functional units of the MVBs, strongly attenuate PME ^53^. Reasoning that the MVBs trigger a cellular degradation pathway to eliminate the PM, we set out to isolate the MVBs from early fertilized eggs and determine their proteome by MS. For this, we first generated two transgenic fly lines, to co-label the MVBs and the PM. The first fly line contains an eGFP-tagged human CD63 (hCD63) gene, a member of the tetraspanin superfamily and a general marker of MVBs in human and *Drosophila*, which mainly localizes to the ILV compartment of these vesicles ^69^. This transgene is expressed under the control of the female germline *UASp* promoter, which allows efficient maternal expression of transgenes by the maternal driver *matα-GAL4*. Note that all transgenes and RNAi lines used in this study are under the control of the maternal promoters *UASp* or *UASz* ^70,71^. The second transgenic line was designed to produce sperm with red fluorescently labeled PM (red-PM). This transgene consists of the red fluorescent protein tdTomato fused to a mitochondrial targeting sequence (MTS) and expressed under the regulatory elements of *don juan* (*dj*), a late spermatogenesis expressed gene ^72^. Eggs laid by females maternally expressing the *eGFP-hCD63* transgene and fertilized by males producing red-PM sperm, displayed multiple eGFP-hCD63 labeled vesicles, including vesicles associated with large segments of the PM (Fig. 1b).

**Fig. 1:**
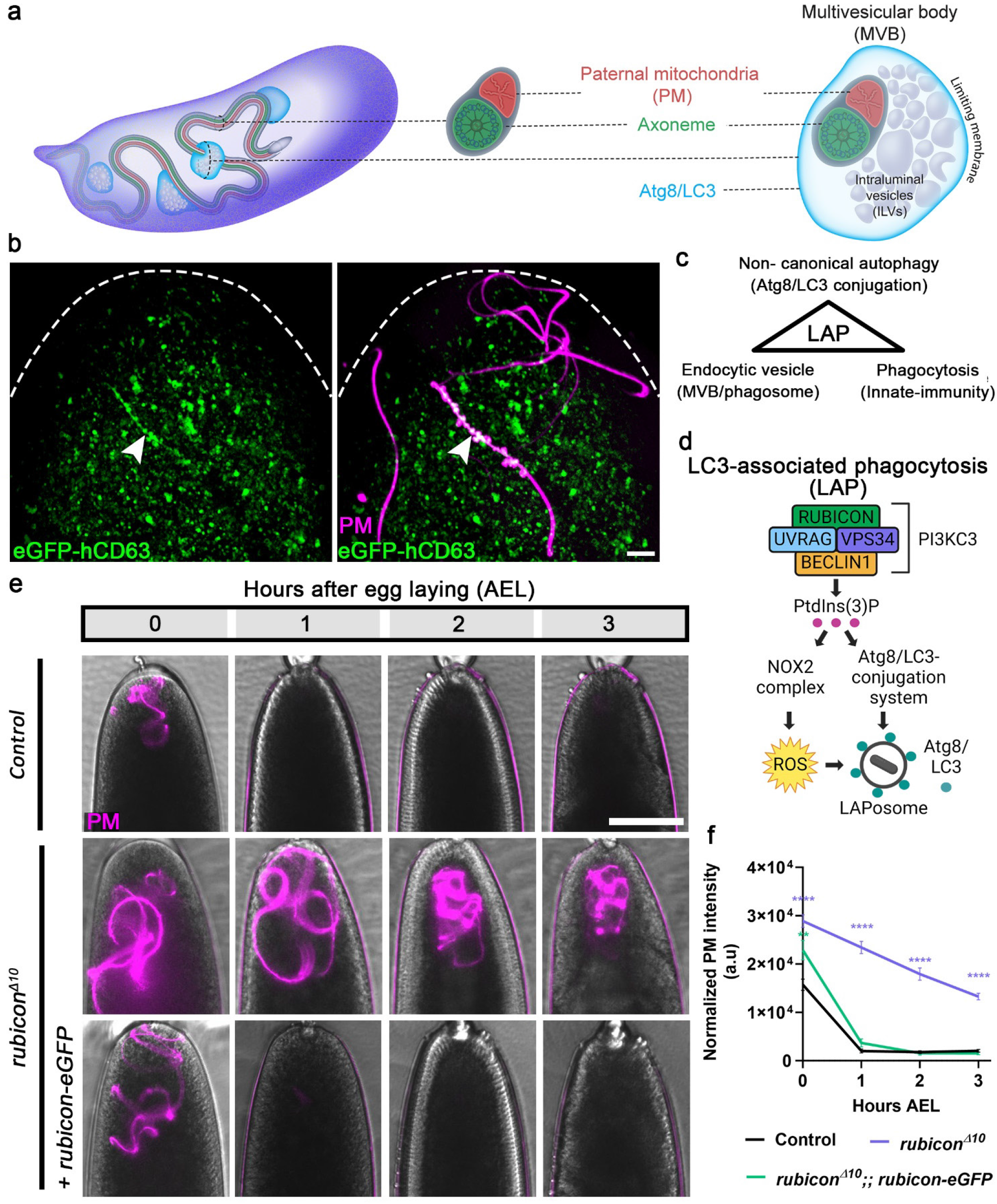
Rubicon is required for PME after fertilization. **a**, Illustration of the onset of PME in *Drosophila* based on ^53^. Left, an early fertilized egg. Egg-derived MVBs associate with the 1.8 mm long sperm flagellum. Middle, the flagellum mainly consists of a single cylindrical mitochondrial derivative (PM; red) and an intimately associated axoneme (green), shown in cross-section, both extending along the entire length of the flagellum. Right, enlargement of an MVB (shown in a cross-section displaying the ILVs and the encapsulating limiting membrane) that associates with the flagellum. Atg8/LC3 (cyan) is loaded on the MVB limiting membrane. **b,** A representative confocal image of the anterior (top) region of an egg maternally expressing the MVB marker hCD63-eGFP (green), and fertilized by a coiled red-PM sperm cell (magenta). While free MVBs are colored green in the merged panel, MVBs that coat large flagellar segments (arrowhead) are white. Scale bar, 10 μm. **c,** Schematic view of three main features shared by the egg MVBs and the LAP pathway. **d,** Schematic view of the LAP pathway as defined in mammalian cells. **e,** Live imaging of the anterior regions of early developing embryos laid by females of the indicated genotypes and fertilized by red-PM sperm. Embryos carrying a single copy of the *rubicon* gRNA served as control. Representative time-lapse images of each embryo were taken at 0, 1, 2, and 3 hours AEL. Brightfield and fluorescence channel image overlay reveals the egg (grey) and the red-PM sperm flagellum (magenta), respectively. The PM signal is lost within one hour in the control embryos, but persists in the *rubicon^Δ^*^10^ mutant embryos, a phenotype rescued by *rubicon-eGFP*. Scale bar, 100 μm. **f,** Quantifications of normalized red-PM fluorescence intensities (arbitrary units [a.u.]) in embryos represented in (**e**). Error bars indicate standard error of the mean (SEM). The respective numbers of scored embryos (n) laid by females of the control, *rubicon^Δ10^*, and *rubicon^Δ10^*;; *rubicon-eGFP* genotypes are 61, 31, and 34. \*\**p <* 0.01 and \*\*\*\**p <* 0.0001. Two-way repeated measures ANOVA, followed by Dunnett’s multiple comparisons test.

To isolate the MVBs, extracts prepared from eGFP-hCD63-expressing early fertilized eggs were subjected to a modified OptiPrep ^TM^ density gradient procedure of subcellular fractionation, and the MVB-enriched fractions were detected using Western blotting with anti-GFP antibodies (Extended Data Fig. 1a). The enrichment of largely intact MVBs and their ILVs in the peak fractions was confirmed by transmission electron microscopy (TEM; Extended Data Fig. 1b), and the composition of the proteins in these fractions was revealed by MS (Table S1). Pathway enrichment analysis of the data revealed six major pathways, five of which were somewhat trivial or unrelated directly to cellular degradation processes (i.e., mitochondria, metabolism, endomembrane system, glycosylation, and transport), but one pathway, innate immunity with elements of phagocytosis, seemed particularly intriguing, as it implied possible involvement of a targeted cellular destruction pathway in PME (Extended Data Fig. 1c).

### Components of the LAP-specific PI3KC3 are required for PME

In the past decade, an unconventional phagocytosis pathway termed <u>L</u>C3-<u>a</u>ssociated <u>p</u>hagocytosis (LAP) has emerged as an important defense pathway against invading microbes. LAP combines the pathways of endocytosis and autophagy, and mediates the delivery of pathogens and other extracellular elements to lysosomes by phagocytosis ^58,59,66,67^. The finding that the egg MVBs, which originate in the endocytic pathway, contain elements of phagocytosis, as well as the observed association of Atg8/LC3 with the limiting membrane of the MVBs that engage the flagellum ^53^, raised the hypothesis that the MVBs mediate PME by activating a LAP-like pathway (Fig. 1c). Conventional autophagy and LAP are functionally and mechanistically distinct, as the autophagosome is a double-membrane structure, while the LAPosome, the characteristic vesicle mediating LAP phagocytosis, is surrounded by a single membrane ^56^. Furthermore, although LAP involves a subset of the autophagic machinery, it does not require the pre-initiation complex. Moreover, the Rubicon protein, a component of the class III PI(3)K complex (PI3KC3) mediating LAP, is essential for LAP but dispensable for autophagy (Fig. 1d)^73^. Significantly, MVBs are also vesicles with a single limiting membrane, and as in LAP, the autophagy pre-initiation complex is also dispensable for PME ^53,74^.

In order to explore the possibility that a LAP-like pathway underlies PME in fertilized *Drosophila* eggs, we set out to examine whether Rubicon, the main protein that distinguishes between the LAP-specific and the autophagy-specific PI3KC3s, is involved in this process ^65,73^. Using CRISPR/Cas9, we generated four *rubicon* mutant alleles, all containing small frameshifting deletions at the beginning of the second exon, leading to premature stop codons and putative truncated proteins that lack the highly conserved RUN domain and the Rubicon homology (RH) domain (Extended Data Fig. 2a). Flies homozygous for the *rubicon* mutant alleles are viable and fertile. To test for possible effects of loss of Rubicon on PME, females homozygous for each of these alleles were crossed with males producing red-PM sperm. Critically, *rubicon* mutant eggs displayed significant attenuation in the degradation of the PM, as revealed by the persistence of the red-PM in the developing mutant embryos, even at 3 hours AEL (Fig. 1e,f and Extended Data Fig. 2b-e). Furthermore, we generated a *rubicon-eGFP* transgene, which upon maternal expression fully restores degradation of the PM in *rubicon* mutant eggs (Fig. 1e,f and Extended Data Fig. 2b,c). In addition to Rubicon, the LAP-specific PI3KC3 protein complex also contains Uvrag, Pi3k/Vps34, and Atg6/Beclin1 (Fig. 1d). Knocking down these components by maternal expression of specific shRNA transgenes (ShR) significantly attenuated PME, although the effect of *atg6* knockdown was less pronounced than that of the *uvrag*, *pi3k* and *rubicon* knockdowns (Fig. 2 and Extended Data Fig. 3a-d). Taken together, these findings suggest that a LAP-specific PI3KC3 is assembled in fertilized eggs and mediates degradation of the PM. It is important to note that to validate that the effects on PME are not a consequence of failed embryonic development, we monitored normal progression in development of the manipulated embryos through the process of blastoderm cellularization, and omitted from our analysis the embryos that failed to develop.

**Fig. 2:**
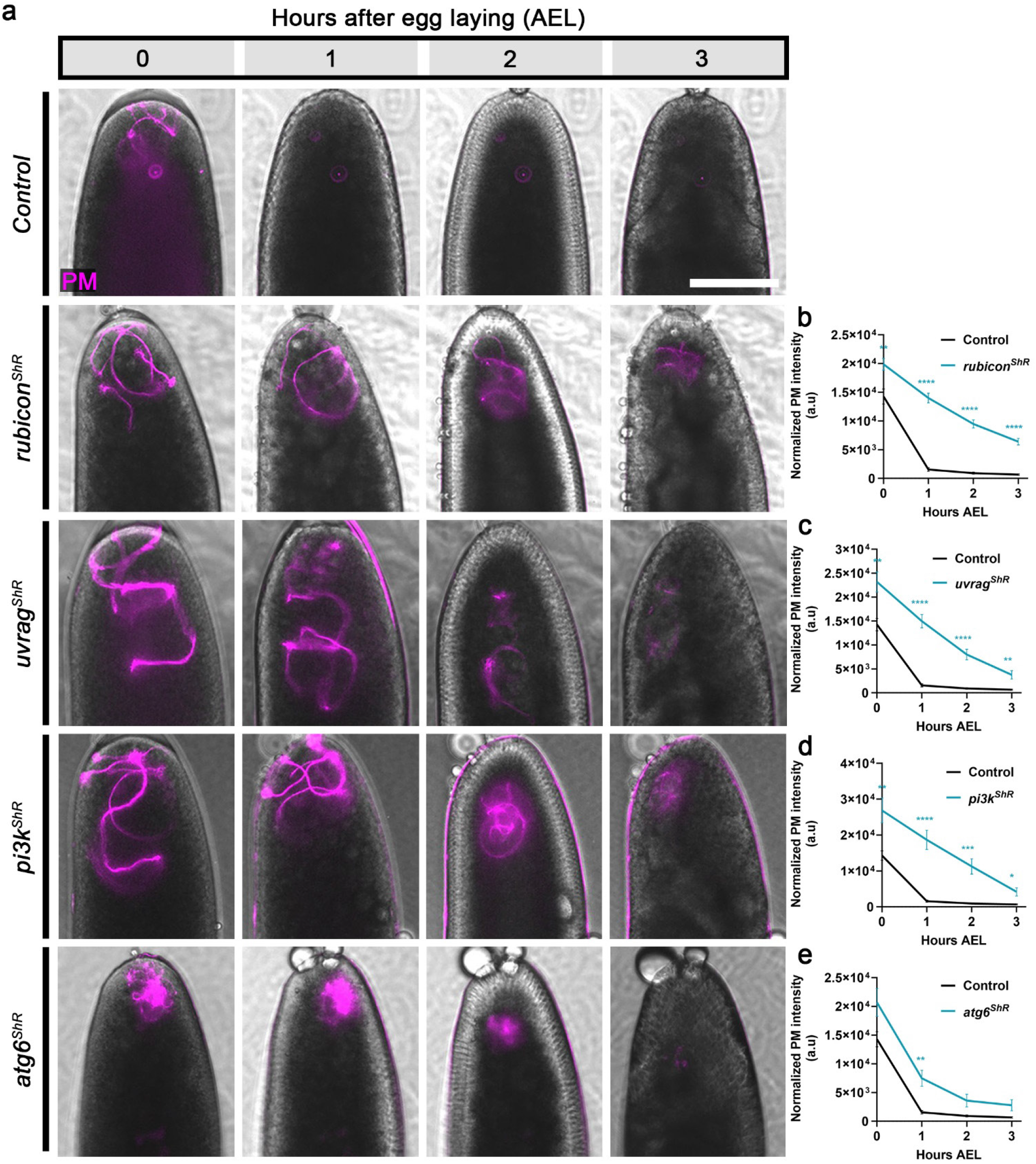
The main members of the LAP-specific PI3KC3 complex are required for PME. **a**, shRNA (ShR) mediated maternal knockdown of *rubicon*, *uvrag*, *pi3k*, and *atg6* significantly attenuates PME. A live imaging assay performed and presented as in Fig. 1e. Embryos expressing the maternal driver alone served as control. Scale bar, 100 μm. **b-e,** Quantifications of normalized red-PM fluorescence intensities (arbitrary units [a.u.]) in embryos represented in (**a**). Error bars indicate SEM. The respective numbers of scored embryos (n) laid by females of the control, *rubicon^ShR^*, *uvrag^ShR^*, *pi3k^ShR^*, and *atg6^ShR^*, are 58, 60, 24, 22, and 24. \**p <* 0.05, \*\**p <* 0.01, \*\*\**p <* 0.001, and \*\*\*\**p <* 0.0001. Two-way repeated measures ANOVA, followed by Šídák’s multiple comparisons test.

### Rubicon coated egg MVBs associate with the sperm flagellum and degrade the PM upon fertilization

To examine whether the LAP-specific PI3KC3 assembles on egg MVBs, live fertilized eggs, laid by females maternally expressing the *rubicon-eGFP* rescuing transgene, and fertilized by males producing red-PM sperm, were collected 0–15 minutes (min) AEL and imaged for 45 min. Remarkably, Rubicon-eGFP specifically labeled numerous vesicles in the egg, including vesicles which enwrap large flagellar segments, implying that Rubicon resides on the MVBs that mediate PME (Fig. 3a and Supplementary Video 2). Closer examination of the coated flagellar segments revealed two types of vesicles, distinguished by their size: Numerous Rubicon-eGFP positive small vesicles densely arranged one next to the other on the flagellum creating extended flagellum vesicular sheaths (FVSs), such that the coated flagellar segments can be detected merely by virtue of Rubicon-eGFP expression (Fig. 3b); and larger vesicles, 1-1.5 µm in diameter on average, budding off from and containing PM pieces and debris (Fig. 3b,c). Interestingly, Rubicon-eGFP primarily localizes to the limiting membrane of these vesicles and the FVS, reminiscent of the assembly of the LAP-specific PI3KC3 on the LAPosome limiting membrane (Fig. 3c). It is noteworthy that, consistent with the requirement of an intact PI3KC3 for PME, the appearance of Rubicon positive FVSs always precedes the elimination of the PM fluorescent signal (Supplementary Video 2).

**Fig. 3:**
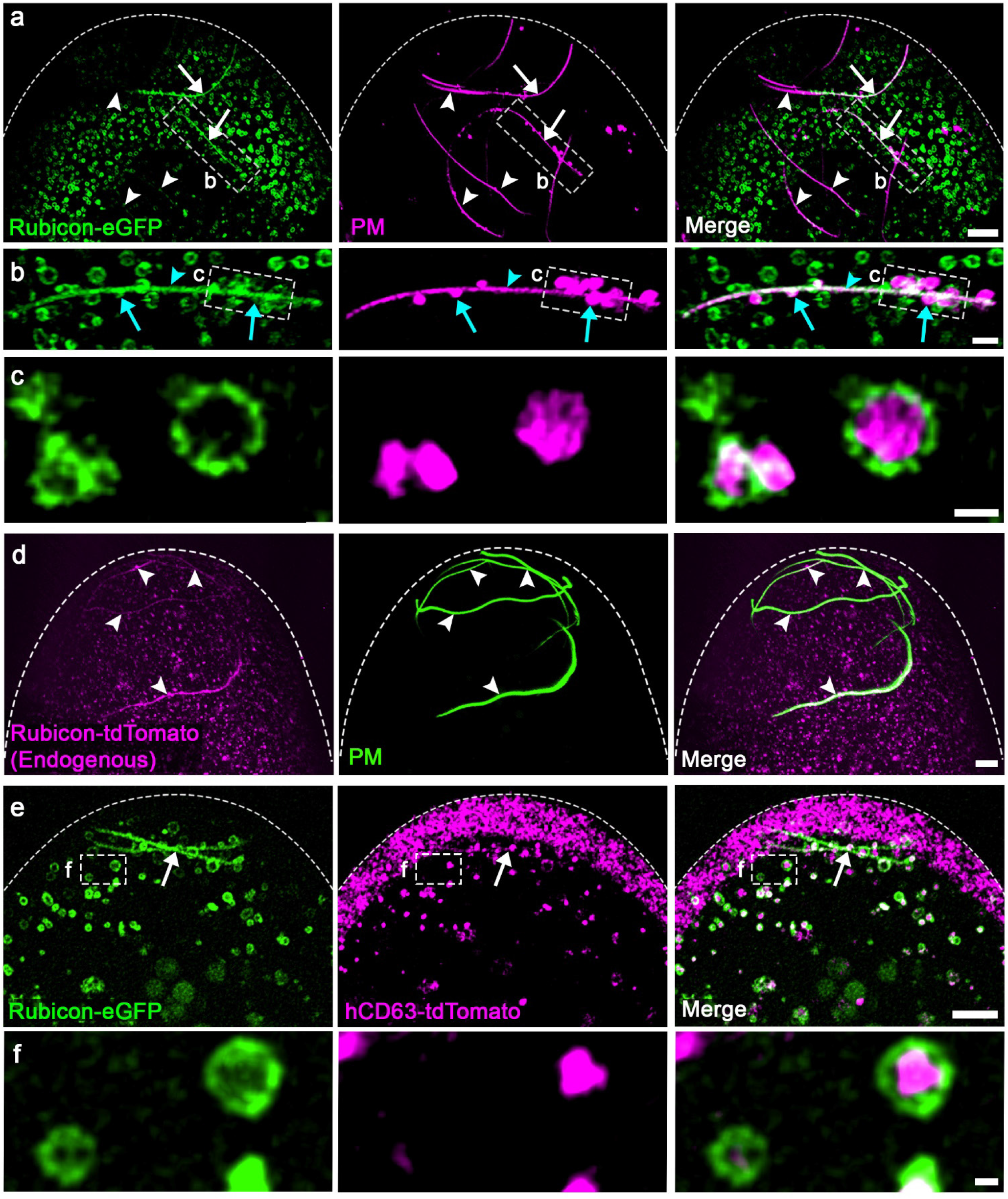
Rubicon is localized to the limiting membrane of multiple egg MVBs which associate with the sperm flagellum. **a,d,e** Representative confocal images of the anterior (top) region of early fertilized eggs. **a,** An egg maternally expressing the *rubicon-eGFP* transgene (green) and fertilized by a red-PM sperm cell (magenta). Numerous egg vesicles display Rubicon-eGFP, many of which associate with and coat large segments of the sperm flagellum (arrows), whereas other flagellar segments show only a few vesicles or are still free of vesicles (arrowheads), due to the asynchronous manner of the PME process along the sperm flagellum. Scale bar, 10 μm. **b,** Magnification of the area outlined by a dashed rectangle in (**a**). Rubicon positive vesicles densely coat flagellar segments creating extended FVS segments (arrowhead). Large Rubicon positive vesicles (green) bud off from the FVS, encapsulating PM pieces and debris (magenta; arrows). Scale bar, 2 μm. **c,** Magnification of a projection of only a few z-sections in the area outlined by a dashed rectangle in (**b**). Rubicon (green) remains localized on the limiting membrane of the vesicles that bud off from the FVS. Scale bar, 1 μm. **d,** An egg expressing tdTomato-tagged endogenous Rubicon (magenta) and fertilized by green-PM sperm (green). Rubicon is localized on multiple egg vesicles, many of which associate with and coat large flagellar segments (arrowheads). Scale bar, 10 μm. **e,** An egg maternally expressing both a *rubicon-eGFP* transgene (green) and the MVB transgenic marker *hCD63-tdTomato* (magenta), and fertilized by a WT (non-fluorescent) sperm cell. Essentially all the Rubicon-eGFP positive vesicles also display the MVB marker. Note that the FVS also exhibits both transgenes (arrow). Scale bar, 5 μm. **f,** Magnification of the area outlined by a dashed rectangle in (**e**). Note that Rubicon is localized on the MVB limiting membrane, while hCD63 resides in the ILV compartments. Scale bar, 0.5 μm.

To confirm that the expression dynamics and localization pattern of the *rubicon-eGFP* transgene in early fertilized eggs reliably reflect those of the endogenous Rubicon locus and protein, we used CRISPR/Cas9 to tag the endogenous *rubicon* gene with tdTomato. Live imaging of eggs laid by Rubicon-tdTomato females and fertilized by males producing green-PM sperm (flies expressing the DJ-GFP transgenic protein ^53,72^), revealed that endogenous Rubicon is expressed in essentially identical spatiotemporal patterns as those of the *rubicon-eGFP* transgene, displaying numerous Rubicon-tdTomato positive vesicles, of which multiple vesicles densely associate with large flagellar segments (Fig. 3d).

To validate that the Rubicon positive vesicles are MVBs, we generated flies maternally co-expressing the Rubicon-eGFP and the MVB marker protein hCD63 fused to tdTomato (i.e., *UASz-hCD63-tdTomato*). Eggs laid by these females and fertilized by wild-type (WT) males displayed numerous vesicles labeled by either Rubicon-eGFP, hCD63-tdTomato, or both (Fig. 3e). Whereas multiple vesicles labeled only by hCD63-tdTomato and located in the egg periphery were not associated with the PM, essentially all the Rubicon-eGFP positive vesicles were also positive for hCD63-tdTomato, including large vesicles associated with the flagellum (Fig. 3e; note that the large flagellar segments can be detected merely by virtue of being enwrapped by multiple small Rubicon-eGFP vesicles). Closer examination of the dually labeled vesicles reveled that the two transgenes occupy different compartments in these vesicles: Rubicon-eGFP is primarily located at the limiting membrane of the MVBs, while hCD63-tdTomato resides in the ILV compartment (Fig. 3f). Interestingly, traces of hCD63-tdTomato are detected along the flagellar segments within the Rubicon-eGFP FVSs (Fig. 3e), which is consistent with our previous ultrastructural studies suggesting that upon engagement with the flagellum, the MVBs release their ILVs in the vicinity of the PM ^53^. Collectively, these observations indicate that the egg contains numerous MVBs pre-loaded with Rubicon, which associate with the flagellum and mediate PME.

### Rubicon positive MVBs and the FVSs exhibit PI3KC3 activity

Active Rubicon PI3KC3 facilitates sustained PtdIns(3)P production by Pi3k/Vps34, which is required for recruitment of the Atg8/LC3 conjugating system to LAPosomes and subsequent Atg8/LC3 lipidation (Fig. 1d). We therefore tested whether the Rubicon positive MVBs are also associated with PtdIns(3)P. For this, we first maternally co-expressed the *rubicon-eGFP* transgene together with TagRFPt-2xFYVE, a genetically encoded reporter for PtdIns(3)P based on the PtdIns(3)P-binding peptide motif FYVE, fused to a red fluorescent protein ^75,76^. Multiple PtdIns(3)P positive vesicles were detected by live imaging of eggs laid by these females and fertilized with WT sperm, while a subset of these vesicles were also positive for Rubicon-eGFP (Fig. 4a). Significantly, sustained presence of PtdIns(3)P on the FVS is detected in eggs maternally expressing the PtdIns(3)P reporter and fertilized by green-PM sperm, indicating that the Rubicon/PtdIns(3)P double-labeled vesicles are the MVBs that engage and densely coat the flagellum (Fig. 4b and Supplementary Video 3). Note that at least some of the vesicles labeled by PtdIns(3)P alone may correspond to the reported PtdIns(3)P expressing intermediate endosomes that accumulate yolk proteins ^75^.

**Fig 4:**
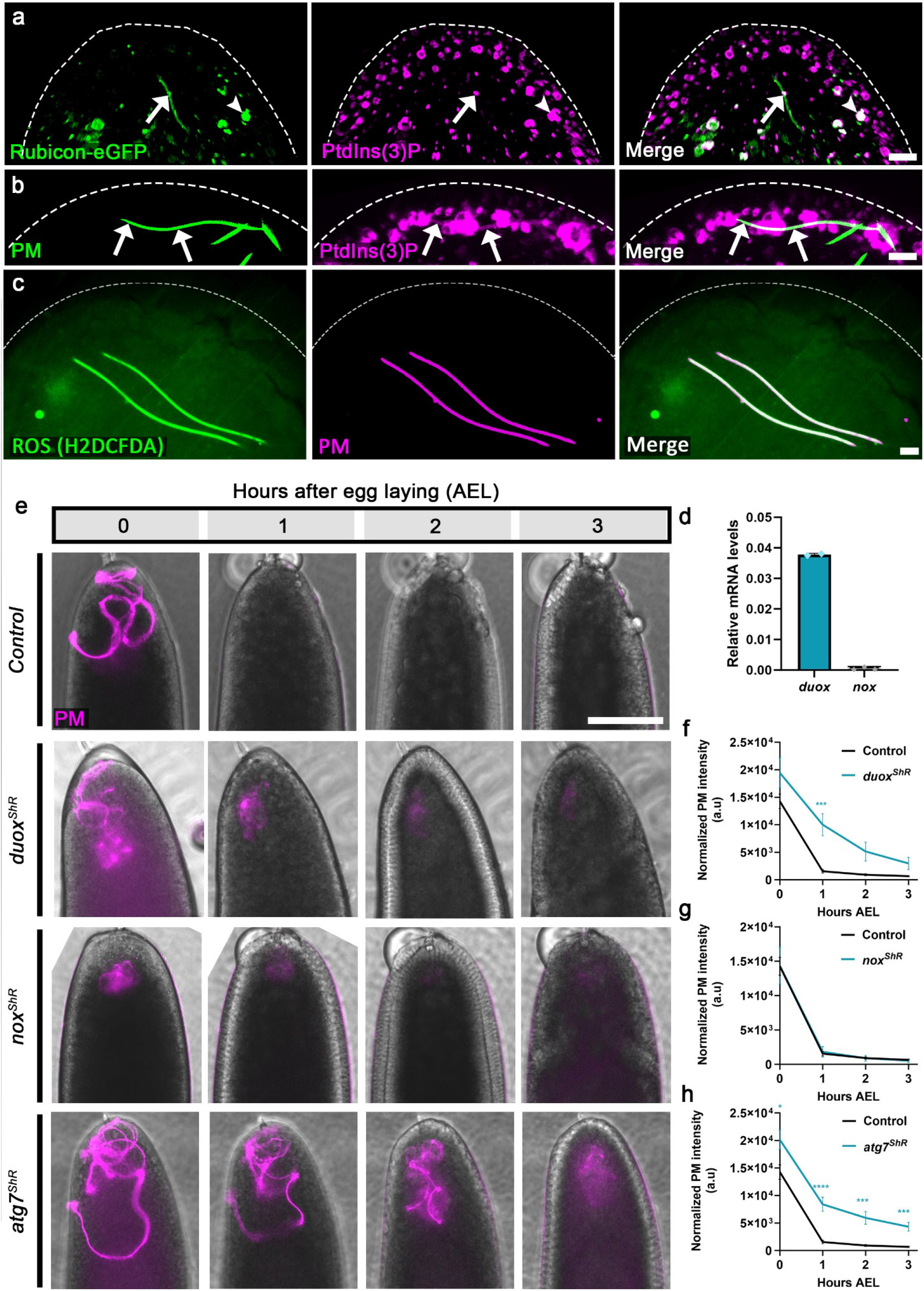
The molecular events associated with PME conform with the LAP pathway cascade. **a-c,** Representative confocal images of the anterior (top) region of early fertilized eggs. **a,** An egg maternally expressing both the *rubicon-eGFP* transgene (green) and the PtdIns(3)P reporter TagRFPt-2xFYVE (magenta), and fertilized by a WT (non-fluorescent) sperm cell. Essentially all the Rubicon-eGFP positive vesicles (MVBs) display PtdIns(3)P as well (arrowhead), including vesicles that generate the FVS (arrow). Scale bar, 6 μm. **b,** An egg maternally expressing the TagRFPt-2xFYVE reporter (magenta) and fertilized by green-PM sperm (green). Note that in addition to the PtdIns(3)P positive vesicles that engage the flagellum (arrowhead), the FVS is also readily displaying PtdIns(3)P (arrows). Scale bar, 10 μm. **c,** Large PM segments display high ROS levels. Shown is a WT egg fertilized by red-PM sperm (magenta) and stained with the fluorescent ROS indicator H2DCFDA (green). Scale bar, 10 μm. **d,** Duox is essentially the sole maternally expressed egg NADPH oxidase. The histogram depicts RT-qPCR analysis of relative mRNA levels of *duox* and *nox* in early fertilized WT embryos. Error bars indicate standard deviation (SD). **e,** shRNA (ShR) mediated maternal knockdowns of *duox* and *atg7*, but not of *nox*, significantly attenuate PME. A live imaging assay performed and presented as in Fig. 1e. Scale bar, 100 μm. **f-h,** Quantifications of normalized red-PM fluorescence intensities (arbitrary units [a.u.]) in embryos represented in (**e**). Error bars indicate SEM. The respective numbers of scored embryos (n) laid by females of the control, *nox^ShR^*, *duox^ShR^*, and *atg7^ShR^*, are 58, 19, 30, and 33. \*\*\**p <* 0.001 and \*\*\*\**p <* 0.0001. Two-way repeated measures ANOVA, followed by Šídák’s multiple comparisons test.

### ROS are involved in PME

In addition to PtdIns(3)P, recruitment of the Atg8/LC3 conjugating system to the maturing LAPosome also requires ROS production, which in phagocytes is generated by the NOX2 multiprotein complex ^58,67,77,78^. WT eggs fertilized by red-PM sperm and labeled with the cell-permeant, fluorescent ROS indicator H2DCFDA, revealed high levels of ROS on the PM (Fig. 4c). Two NADPH oxidase (NOX) homologs, Nox and Duox, are present in *Drosophila* ^79^. Quantitative reverse transcription PCR (RT-qPCR) analyses of early fertilized egg transcripts indicated that of the two, *duox* is essentially the sole maternally expressed gene (Fig. 4d), in agreement with omics data in two public databases ^80,81^. Duox is also enriched in the egg MVB proteome (Table S1). Accordingly, knockdown of *duox*, but not *nox*, in eggs fertilized by red-PM sperm, moderately attenuated PME, suggesting that Duox is responsible for at least a portion of the ROS required for PME (Fig. 4e-g and Extended Data Fig. 3e,f). Note that we cannot exclude the possibility that the PM itself also produces ROS. Taken together, we conclude that the two established mechanistic requirements for recruitment of the Atg8/LC3 conjugating system to the LAPosome, production of PtdIns(3)P and ROS, are also fulfilled upon engagement of the MVBs with the sperm flagellum in early fertilized *Drosophila* eggs.

### Conjugation of Atg8/LC3 on the FVS is a late onset event

Atg7 is an E1 ligase-like enzyme, that plays a central role in autophagy by activating two different ubiquitin-like proteins, Atg12 and Atg8/LC3 ^82,83^. Atg12 then undergoes conjugation to Atg5, and the Atg12-Atg5 complex interacts with Atg16L1 to form a large multimeric complex that facilitates efficient Atg8/LC3 lipidation ^84^. Likewise, in LAP, the Atg12-Atg5-Atg16L1 complex is required for Atg8/LC3 conjugation to the LAPosome membrane ^56,85–87^. We previously showed that in *atg7* mutant eggs, PME is attenuated and the MVBs are deformed due to improper biogenesis of the ILVs ^53^. Consistently, mild knockdown of maternal *atg7* moderately attenuated PME, as revealed by persistence of the red-PM in these eggs (Fig. 4e,h and Extended Data Fig. 3g). Unexpectedly however, efficient maternal knockdowns of *Drosophila atg5*, *atg12*, and *atg16* homologs caused only weak PME attenuation if any (Extended Data Fig. 4a-d and Extended Data Fig. 3h-j). Moreover, maternal knockdown of *atg8a*, the major fly *atg8* gene, had no significant effect on PME as revealed by this assay (Extended Data Fig. 4a,e and Extended Data Fig. 3k). While *Drosophila* has another closely related but distinct *atg8* gene, *atg8b*, its expression is mainly restricted to the testis and is barely detected in other tissues ^88–90^. Although these findings may suggest that conjugation of Atg8 on the FVS is dispensable for PME, pronounced Atg8a expression can still be detected on the FVSs that coat large flagellar segments during advanced PM degradation stages, which may imply an important role for Atg8 conjugation in PME (Fig. 5a). It is possible that for some technical reasons, the assay which we use to determine PME kinetics, i.e., monitoring the elimination of red-PM fluorescence, is inappropriate for assessment of impaired Atg8 conjugation effects. For example, since Atg8/LC3 conjugation is the most downstream event in the LAP pathway, it may occur only after elimination of the red-PM fluorescence, while the destruction of the PM is still in progress and requires Atg8 conjugation.

**Fig. 5:**
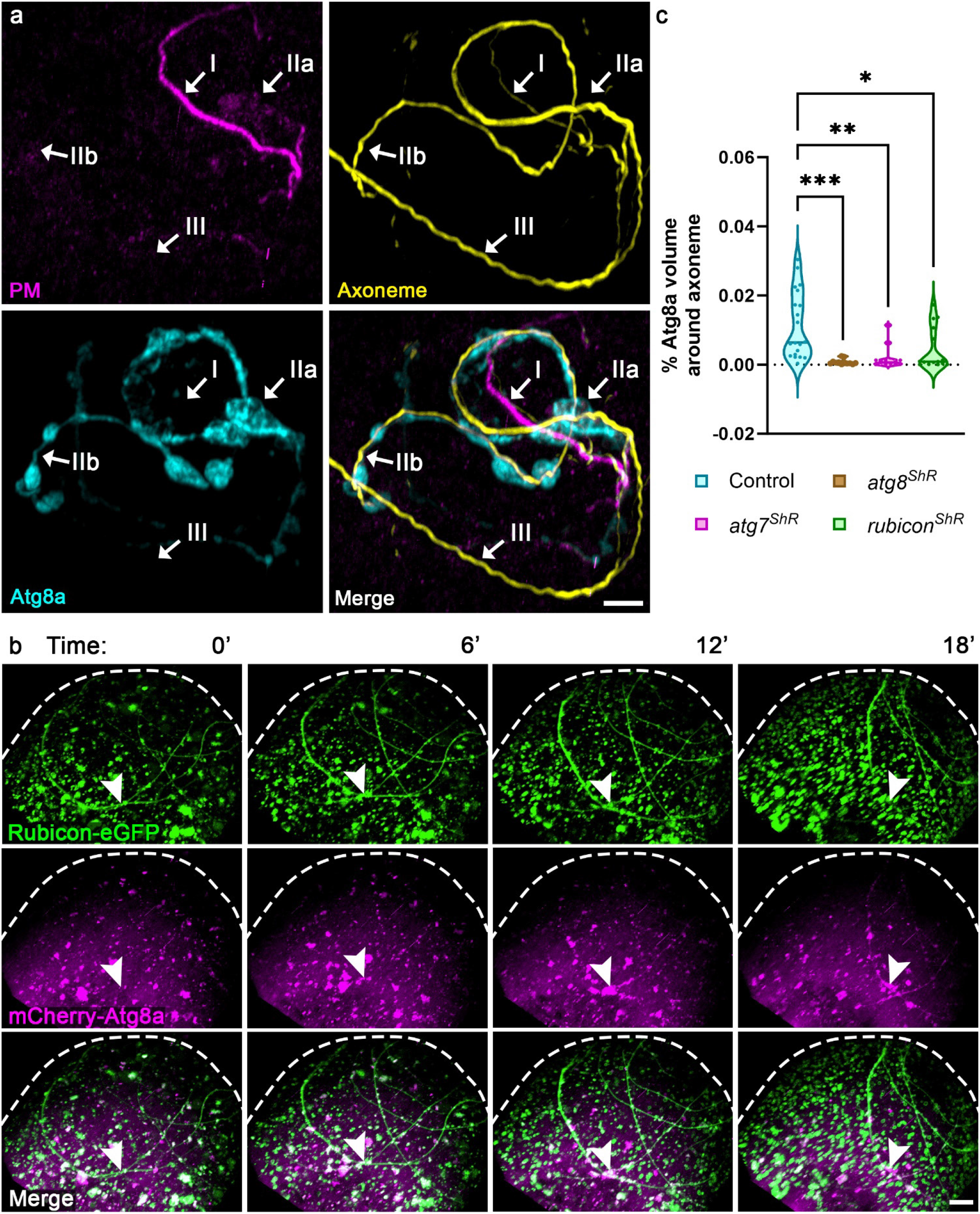
Atg8a recruitment to the FVS coincides with advanced PM degradation stages and requires prior expression of Rubicon. **a**, A representative confocal image of a WT egg fertilized by green-PM sperm (magenta; stained with anti-GFP antibodies) and also immunostained to visualize the axoneme (yellow) and Atg8a (cyan). Shown is a sperm flagellum which exhibits at least three defined consecutive PME stages (I; IIa,b; III) detailed in the main text. Scale bar, 4 μm. **b,** Live confocal imaging of the anterior region of an early developing embryo laid by a female maternally expressing both *rubicon-eGFP* (green) and *mCherry-atg8a* (magenta) transgenes. Shown are representative images from Supplementary Video 4 at the indicated times (min AEL). Note the 6-12 min lag between the formation of Rubicon-eGFP positive FVS and the subsequent recruitment of mCherry-Atg8a (an arrowhead points to the same position on the FVS). Scale bar, 5 μm. **c,** Downregulation of *rubicon* and *atg7* abrogates Atg8a recruitment to the FVS. The violin plots depict the staining volumes of Atg8a in a 0-1 μm radius area around the axoneme, measured in 0–1-hour AEL fertilized eggs maternally expressing shRNA (*ShR*) transgenes against *rubicon*, *atg7*, and *atg8a*, and immunostained to reveal Atg8a and the axoneme. Summary of the statistics reported in the violin plots: the center line represents the median of the frequency distribution of the data. Quartiles are represented by a dashed black line. Each dot corresponds to the percentage of Atg8a volume in a single embryo to reflect n number, where n indicates the number of examined embryos. Corresponding plots measuring Atg8a staining volumes at further distances from the axoneme, as well as representative images are presented in Extended Data Fig. 5. The respective numbers of scored embryos (n) laid by females of the control (WT), *atg8a^ShR^*, *atg7^ShR^*, and *rubicon^ShR^*, are 20, 12, 11, and 15. One-way ANOVA, followed by Dunnett’s multiple comparisons test. *p* values for *atg8a^ShR^*, *atg7^ShR^*, and *rubicon^ShR^* are 0.0002, 0.0016, and 0.0119, respectively.

To explore this idea further, we first sought to determine the timing of Atg8a conjugation to the FVS during the PME sequence of events. For this, ∼50 WT eggs fertilized by green-PM sperm were collected at 0–1-hour AEL, fixed, and immunostained to visualize the PM (anti-GFP antibody), Atg8a (anti-Atg8a antibody), and the axoneme (anti-polyglycylated Tubulin antibody ^53,91,92^). Three consecutive and distinct PME phases can be clearly detected, often within a single fertilized egg: 1) Intact PM segments that are Atg8a negative and usually still reside in the vicinity of parallel axonemal segments (Fig. 5a, I); 2) FVSs that become positive for Atg8a, which often display large Atg8a positive vesicles that bud off from the FVSs and encapsulate degrading PM pieces (Fig. 5a, IIa), resulting in PM-free axonemal segments (Fig. 5a, IIb); 3) PM-free axonemal segments displaying neither PM traces nor Atg8a positive FVS, indicative of completion of the PME process (Fig. 5a, III). Taken together, these observations suggest that conjugation of Atg8a on the FVS occurs when most, if not all, of the PM transgenic fluorescent signal is lost. This indicates that the PM degradation process already begins with the coating of the flagellum by the Rubicon MVBs, and therefore, the decaying PM fluorescent signal cannot be reliably used to explore the effects of improper *atg8a* conjugation on PME.

To evaluate the time elapsed between coating of the flagellum by the Rubicon MVBs and conjugation of Atg8a on the FVS, we maternally expressed *rubicon-eGFP* and *mCherry-atg8a* transgenes, and used live imaging to monitor their relative spatiotemporal patterns of expression. Large Rubicon-eGFP positive FVSs that are mCherry-Atg8a negative were readily detected at time 0 AEL, while conjugation of mCherry-Atg8a on the FVS occurred 6–12 min later (Fig. 5b and Supplementary Video 4). Altogether, these findings are consistent with the established LAP pathway sequence of events, in which loading of the Rubicon PI3KC3 on the FVS precedes conjugation of Atg8/LC3.

### Rubicon is required for Atg8/LC3 conjugation to the FVS

To examine whether early loading of Rubicon PI3KC3 is required for Atg8a conjugation to the FVS, we next monitored the staining volume of Atg8a in 0–1-hour AEL fertilized eggs maternally expressing an shRNA transgene against *rubicon*. Maternal expression of an *atg7* shRNA transgene served as positive control, whereas down regulation of *atg8a* was used to validate the specificity of the anti-Atg8a antibody. To avoid possible unrelated Atg8a positive vesicles, such as unrelated autophagosomes, we restricted the measurements to the area surrounding the axoneme. In addition, to assure that the developmental stages of the embryos are highly synchronized, we monitored syncytial embryos up to nuclear division cycle six. The Atg8a staining volume in a 0-1 µm radius around the axoneme, corresponding to the FVS-associated Atg8a, was significantly reduced upon down regulation of *rubicon*, *atg7*, and *atg8a*, as compared with corresponding WT eggs, indicating that Atg8a conjugation to the FVS requires both Rubicon and Atg7 (Fig. 5c). Furthermore, this trend persisted even up to a 10 µm radius around the axoneme, demonstrating that vesicles which bud off from the FVSs and/or MVBs that were not associated with the flagellum, both require Rubicon and Atg7 for Atg8a conjugation (Extended Data Fig. 5a and corresponding representative images in Extended Data Fig. 5b-e).

Because PM degradation is significantly delayed upon down regulation of *rubicon* (Fig. 2b), we asked whether the kinetics of Atg8a conjugation to the FVS is also affected in this setup. For this, eggs maternally expressing the *mCherry-atg8a* transgene and fertilized by green-PM sperm were monitored by live imaging. In an otherwise WT background, most of the flagellar segments underwent Atg8a conjugation and PM degradation within 10-20 min AEL, such that only a very few short PM segments are still detected at 50 min AEL, by virtue of their mCherry-Atg8a positive FVSs (Extended Data Fig. 6a and Supplementary Video 5). In contrast, eggs maternally expressing the *rubicon* shRNA transgene displayed a delayed appearance of mCherry-Atg8a positive FVS (40–50 min AEL), coating only a few short flagellar segments (Extended Data Fig. 6b and Supplementary Video 6). Interestingly, despite the significant delay in elimination of the fluorescent PM signal, it eventually disappeared, implying that either some residual Rubicon protein may still promote low efficiency level of PM degradation, or that other mechanisms may at least partially degrade the PM (Extended Data Fig. 6b and Supplementary Video 6). Collectively, we conclude that a complete LAP-like pathway mediated by the egg MVBs, is required for efficient elimination of the PM after fertilization, and that interfering with this pathway leads to a highly inefficient PME process.

### A LAP-like pathway may mediate PME in mammals

Although the involvement of classical autophagy was proposed to be dispensable for PME in mice ^21^, autophagy-related proteins, including ubiquitin, P62 and Atg8/LC3, were shown to decorate the mammalian PM just prior to or immediately after fertilization ^21,40,41^. We therefore asked whether it is possible that a LAP-like pathway might also mediate PME in mammals. To examine the feasibility of this idea, we immunostained spermatozoa isolated from mouse cauda epididymis and from bull semen with two anti-Rubicon antibodies, raised against different human RUBICON epitopes. To detect the PM, sperm cells were pre-incubated with MitoTracker. Significantly, as revealed by both antibodies, Rubicon readily decorates the entire flagellum of both mouse and bull sperm cells, including the midpiece region where the mitochondria are localized (Extended Data Fig. 7). These observations imply that a LAP-like pathway might also operate in mammals to eliminate the PM after fertilization, and that the initial steps of the pathway might commence prior to fertilization.

## Discussion

In this work, we uncovered the pathway by which the paternal mitochondria (PM) are eliminated immediately after fertilization in *Drosophila*. We show that special MVBs, which are abundant in the early fertilized egg, engage the sperm flagellum immediately after fertilization, while triggering a LAP-like pathway to specifically target the PM for degradation. This highly efficient process takes about 20 min, as revealed by the disappearance of transgenic fluorescent signals that label the PM, with an average PM degradation rate of 100 nm/min. According to the sequence of events we uncovered, numerous egg MVBs that are loaded with the Rubicon-containing PI3K complex (PI3KC3) on their limiting membrane, approach and densely coat large segments of the penetrating sperm flagellum in an asynchronous manner, forming flagellum vesicular sheaths (FVSs) within which the corresponding PM segments are degraded (Fig. 6, I, II). At the molecular level, activated PI3KC3 generates PtdIns(3)P, which in turn recruits the Atg8 conjugating system to the FVS (Fig. 6, III). Consequently, Atg8 is conjugated on the FVS, presumably with the assistance of ROS generated by Duox in the egg and/or the PM itself (Fig. 6, IV). Eventually, the Atg8 positive FVSs bud off large vesicles that contain degrading PM pieces and debris, leaving behind the axoneme (Fig. 6, V). Ultimately, this process assures inheritance of the mitochondria from the maternal lineage only. The mechanism by which the MVBs specifically target the PM while the axoneme and the maternal mitochondria are spared is uncertain, but may involve specific ubiquitination of PM surface proteins, as we previously demonstrated ^53^.

**Fig. 6:**
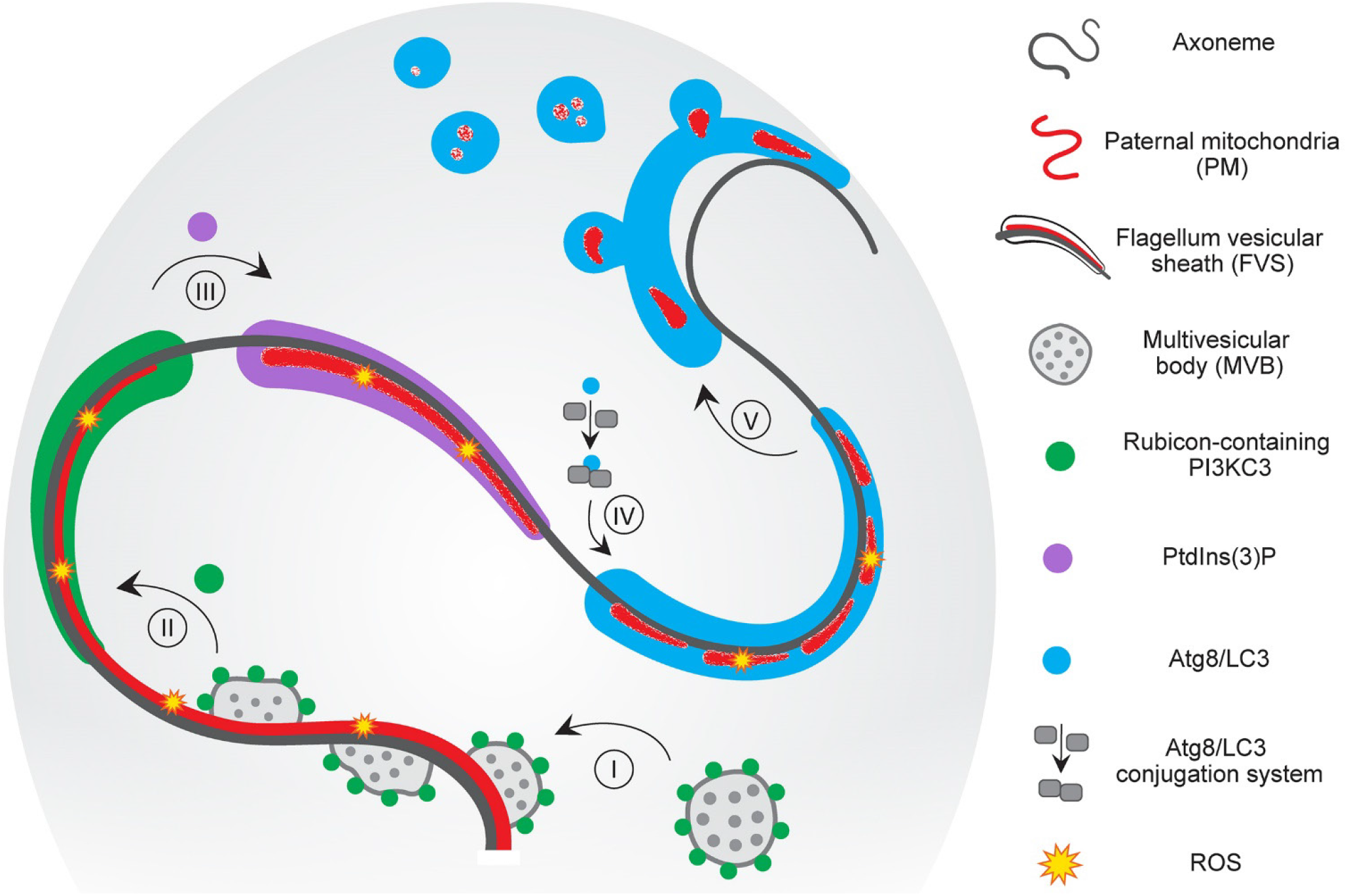
An integrated model of PME by a LAP-like pathway in *Drosophila*. An illustration of a sperm flagellum inside the anterior part of an early fertilized egg. Relevant flagellar and LAP pathway components are indicated by different signs and colors as defined on the right-hand side. Five PME/LAP pathway steps are indicated by Roman numerals. (I) Egg-derived Rubicon positive MVBs engage the sperm flagellum in an asynchronous manner immediately after fertilization, (II) forming FVSs around large flagellar segments. (III) Active Rubicon PI3KC3 generates PtdIns(3)P, that promotes the recruitment of the Atg8/LC3 conjugation machinery, (IV) which in turn conjugates Atg8/LC3 on the FVS, presumably with the help of ROS (produced by the egg and perhaps also by the PM). (V) Consequently, large vesicles containing degraded PM material bud off from the FVS, leaving behind a PM-free axoneme.

Whereas LAP is known to target extracellular pathogenic microbes by phagocytosis, targeting of the PM occurs after the sperm enters the egg ooplasm. Nevertheless, these two processes are not only molecularly reminiscent of each other but also anatomically similar. In LAP, a single membrane LAPosomal vesicle encapsulates the cargo by inward invagination of the plasma membrane, while during PME, single membrane MVBs form extended FVSs that encapsulate large flagellar segments. Interestingly, in addition to LAP, other pathways, such as LC3-associated endocytosis (LANDO) and LC3-dependent extracellular vesicle loading and secretion (LDELS), were reported to involve conjugation of Atg8/LC3 to the limiting membrane of endosomes and MVBs, respectively, indicating that convergence of the endocytosis and the autophagy pathways may be more common than has been previously recognized. Furthermore, similar to *Drosophila* PME, induction of LDELS does not require surface receptor activation ^66,87,93–95^.

A general belief is that elimination of the PM is important to prevent mtDNA heteroplasmy, partially because sperm mtDNA was reported to have a higher mutation rate than egg mtDNA ^96^. Furthermore, deliberate major mtDNA heteroplasmy in mice was reported to lead to reduced animal activity, increased anxiety, reduced feeding, and impaired spatial learning ^31^. However, prevention of mtDNA heteroplasmy may not be the entire story, as several reports suggest that in some cases, including in *Drosophila*, the sperm mtDNA is degraded before the observed degradation of the vacuolar PM ^18–22^, implying that the latter degradation is a second line of defense against paternal mtDNA transmission, or that other PM factors are deleterious to the organism. An alternative theory is that, similar to the role of macroautophagy in providing nutrients during starvation, recycling of the PM may support proper embryonic development, by providing developmental cues or important substances to the fertilized egg, such as iron, copper, folates, etc. ^97–99^. Our findings that the egg triggers a conserved cellular defense pathway against invading microbes to target the PM for degradation, support the idea that the PM deliver deleterious factors to the embryos, as they imply that the egg regards the PM as a potentially dangerous trespasser, reminiscent of the manner by which other cells react against intrusive bacteria.

Paternal mitochondrial elimination and maternal mitochondrial inheritance are highly ubiquitous across the animal kingdom. Scattered evidence from different research groups suggests that at least some elements of the autophagy pathway are expressed on the flagellum of mammalian sperm, either before and/or immediately after fertilization ^21,40–42^. The current work provides evidence for Rubicon presence on the sperm flagellum of bulls and mice prior to fertilization. Evidence from electron microscopy studies showed the presence of MVBs in the vicinity of the PM in some mammalian early fertilized eggs ^38,68^. Collectively, it is attractive to propose that a LAP-like pathway similar to the pathway underlying PME in *Drosophila* might also mediate the process in mammals and perhaps also in other organisms that produce flagellated sperm. Such a mechanism may well operate in other instances where non-conventional mitochondria and other organelles, in terms of unusual size, anatomy and structure, need to be efficiently cleared. The importance of understanding the mechanisms underlying PME is not only of academic interest, as the use of medically assisted reproduction (MAR) treatments is constantly increasing, leading to diversion of research efforts from molecular studies to improvement of MAR technologies. Consequently, much less attention has been given to possible negative impacts of some of these treatments, which inevitably involve PM leakage, on embryo development, implantation, and clinical and obstetric outcomes ^100–102^. Furthermore, the novel reproductive technologies known as mitochondrial donation or mitochondrial replacement therapy (MRT), raise challenging ethical and safety issues, as only a few successful MRT case reports are available. Furthermore, since we still lack long-term studies of MRT outcomes, the potential risks of both major mtDNA heteroplasmy and the mixing of genetic materials (nuclear and mitochondrial genomes) from three parents are under much debate ^103–107^. Therefore, delineating the molecular pathway underlying PME and the (possible negative) effects of PM persistence on the developing embryo, will allow for deeper understanding of the importance of this highly conserved phenomenon for embryo development, and may ultimately lead to the development and/or modification of MAR technologies that guard against these effects.

## Methods

### Fly strains used in this study

Fly stocks were maintained on standard yeast/molasses medium at 25°C. Unless indicated otherwise, *yellow white (yw)* flies were used as WT control. Fly strains, including mutant and transgenic flies used in this study are listed in the following table:

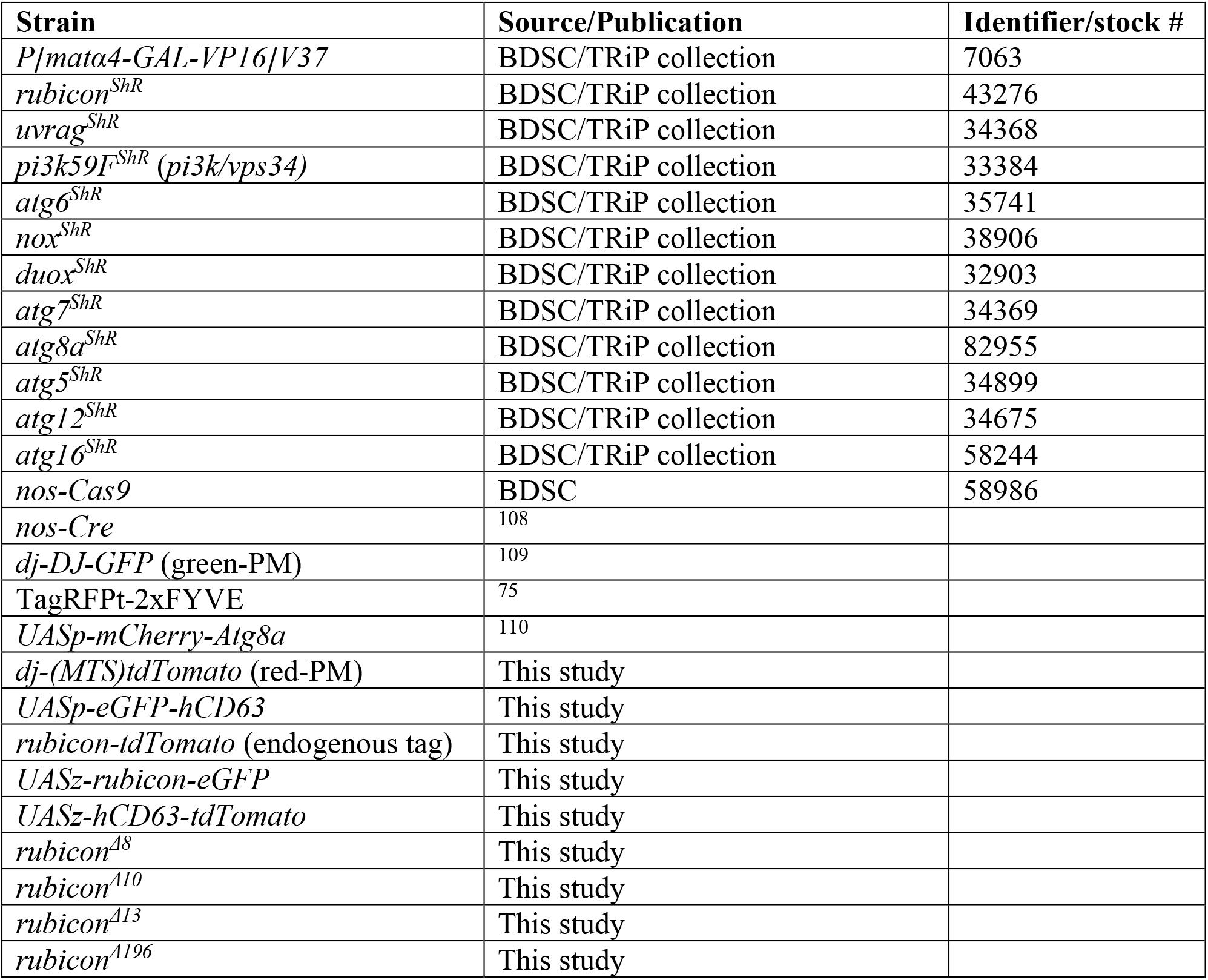

### Generation of the *rubicon* mutant alleles

The *rubicon^Δ8^*, *rubicon^Δ^*^10^, *rubicon^Δ^*^13^ and *rubicon^Δ^*^196^ mutant alleles were generated using CRISPR/Cas9-mediated mutagenesis. Briefly, *nos-Cas9* expressing flies were crossed to flies ubiquitously expressing a single guide RNA (gRNA) against *rubicon* (BDSC #81781). F1 females carrying both the *nos-Cas9* and the *rubicon* gRNA, and hence subjected to potential germline mutagenesis in the *rubicon* gene, were crossed with males carrying an X chromosome balancer (FM7), as the *rubicon* gene is located on the X chromosome. F2 females were then individually crossed again with FM7 males to establish stable fly lines with potential mutations in *rubicon*. Genomic DNA was extracted from each line, the *rubicon* gene was PCR amplified using forward (F) primer: CCGAGACAGTGCGTCAATG and reverse (R) primer: CTTCCTCGCTGGGATAGTACA, and sequenced to identify *rubicon* mutant lines.

### Generation of DNA constructs and transgenic lines

#### *dj-(MTS)tdTomato* (red-PM)

The tdTomato coding sequence (CDS) was PCR amplified from the *pRSETB-tdTomato* vector ^111^ using F primer: CATTCGTTGGGGGATCCACCGGTCGCCACCATGGTGAGCAAGGGCGAGGAGG and R primer: ATCTTGTTGTTTCGCAGGCGATTGCGGCCGTTACTTGTACAGCTCGTCCATGC.

Applying the restriction free technique (RF) ^112^, the DsRed CDS in the *dj-(MTS)DsRed* plasmid ^53^ was replaced with the tdTomato CDS. Note that for expression during late-stage spermatogenesis, the *(MTS)tdTomato* is placed under the regulatory elements of the *dj* gene (i.e., promoter and 5’ UTR) ^72,113^, as well as the 3’UTR of the male germ cell-specific gene *cyt-c-d* ^114^. Embryo injection to introduce the transgene into the attP2 site was carried out by BestGene Inc.

#### UASp-eGFP-hCD63

The eGFP-hCD63 CDS was PCR amplified from a *pUASt-eGFP-hCD63* plasmid (kindly provided by Suzanne Eaton, the Max Plank Institute of Molecular Cell Biology and Genetics, Germany) using F primer: CCGCGCGGCCGCATGGTGAGCAAGGGCGAGGAGCTG and R primer: GCGGACTAGTCTACATCACCTCGTAGCCAC. The eGFP-hCD63 CDS was subcloned into the *pUASp2* vector ^70^ using the restriction enzymes NotI and SpeI for the insert, and NotI and XbaI for the vector. Embryo injection to introduce the transgene into the 2^nd^-chromosome through P element-mediated transformation was carried out by BestGene Inc.

#### UASz-rubicon-eGFP

This line was generated using the *In-fusion* cloning technique (Takara) in two steps as follows: First, the *eGFP* DNA sequence was PCR amplified using F primer: GTACCGCCTCTCTAGGTGAGCAAGGGCGAGGAG and R primer: GAATTCACACTCTAGTTACTTGTACAGCTCGTCCATGCC. This amplicon was inserted into the *pUASz1.1* plasmid ^71^, obtained from the *Drosophila* Genomics Resource Center (DGRC; stock #1433), following digestion of the plasmid with XbaI to obtain a *pUASz-eGFP* vector. Next, the full-length *rubicon* CDS was PCR amplified from the cDNA clone *RH61467* (DGRC, Stock #11152) using F primer: CAAAGGATCCCTCGAATGACCACGCCCCCG and R primer: AGGCGGTACCCTCGAGCTGGCACGGCTTTGA. The amplicon was inserted into the *pUASz-eGFP* vector following digestion with XhoI to generate the *pUASz-Rubicon-eGFP* vector. Embryo injections to introduce the transgene into the attP2 and attP40 sites were carried out by FlyORF.

#### UASz-hCD63-tdTomato

The *tdTomato* gene was first isolated from a plasmid by digestion with BamHI and ligated into BamHI linearized *pUASz1.1* plasmid to obtain the *pUASz-tdTomato* vector. *hCD63* CDS was PCR amplified from the *UASp-eGFP-hCD63* vector using F primer: CAAAGGATCCACCGGATGGCGGTGGAAGGAGGAAT and R primer: CATGGTGGCGACCGGCATCACCTCGTAGCCACTTCTGA. The amplicon was then inserted into the *pUASz-tdTomato* vector using the *In-fusion* cloning technique (Takara) following digestion with AgeI. Embryo injection to introduce the transgene into the attP2 site was carried out by FlyORF.

#### Tagging the endogenous rubicon gene with tdTomato

A single gRNA was designed to target Cas9 mediated excision around the stop codon of the *rubicon* gene (TTCTTTGATCGGATGTTAGC in the *pCFD5* plasmid). For homology-directed repair, we generated a donor plasmid in several steps as follows: We first inserted the *tdTomato* gene into the *pHD-DsRed-attP* vector, which contains a loxP flanked 3XP3-DsRed cassette that produces eye-specific expression of the DsRed ^115^. Then, using *In-fusion* cloning (Takara), we subcloned into the *pHD-tdTomato-DsRed-attP* vector the two homology arms (HAs) which contain sequences flanking the Cas9 cleavage site. The HAs were PCR amplified from *yw Drosophila* genomic DNA using F primer: CTGGGCCTTTCGCCCCCAGTCTGACCAGAGGCACC and R primer: CCCCATAATTGGCCCTTGCTGGCACGGCTTTGAAGG, for the 5’ HA, and F primer: ATAGAAGAGCACTAGCATCCGATCAAAGAAAATCGAAGGG and R primer: GGAGATCTTTACTAGACGCGTTCGGCAAAATACC, for the 3’ HA. The *pHD-tdTomato-DsRed-attP* vector was digested with SmaI to first insert the 5’ HA and subsequently with SpeI to insert the 3’ HA, obtaining the *pHD-5’HA-tdTomato-3’HA-DsRed-attP* donor plasmid. Embryo injections to obtain transgenic lines were carried out by FlyORF. F1 progeny was screened for insertion events using the eye DsRed expression. Positive lines were validated by sequencing and then crossed to *nos*-Cre flies to remove the 3XP3-DsRed cassette.

### Purification of the egg-derived MVBs

To isolate the egg-derived MVBs, we devised a procedure that is based on two protocols, one for mitochondria isolation from *Drosophila* embryos and the second for isolation of exosomes from cultured cells ^116,117^ .To adjust the procedure, MVB isolation trials were first performed on small-scale eGFP-hCD63 expressing embryos (i.e., embryos laid by females carrying the maternal driver and the *UASp-eGFP-hCD63* transgene and fertilized by WT males). Once adjusted, the procedure was then scaled-up to produce a large amount of protein that is needed for MS analysis. To avoid slow embryo collection due to repeated crossings, the large-scale MVB isolation was performed with WT (*yw*) embryos. Collection, preparation, and MVB extraction from the eGFP-hCD63 expressing embryos was performed as follows: Flies were placed in a population cage, allowing to lay eggs on juice agar plates with a dollop of thick yeast paste for one hour. Embryos were then collected in mesh baskets, dechorionated in undiluted 6% bleach, washed with water, shock frozen in liquid nitrogen, and stored at −80°C until the day of procedure. All further steps were performed on ice and using a cooled centrifuge. 60 mg of embryos were thawed on ice and transferred to a Dounce homogenizer containing 6-times volume of homogenization buffer (HB; 0.25 M sucrose, 1 mM EDTA, 0.03 M Tris pH 7.4, supplemented with protease inhibitor cocktail (Sigma-Aldrich, P8340)). The homogenate was centrifuged three times at 500 x g for 15 min to pellet cell debris. A fraction of the supernatant was removed and served as cell lysate control, while the remaining supernatant was centrifuged at 12,000 x g for 20 min. The pellet was then resuspended in HB and layered onto a 5-50% OptiPrep™ Density Gradient Medium (Sigma-Aldrich, D1556), and ultracentrifuged at 50,000 x g for 3 hours in a SW41Ti rotor (Beckman). 12 fractions of 1 ml each were manually separated, diluted 1:4 in OptiPrep™ buffer (OB, 0.07 M EDTA, 0.03 M Tris pH 7.4) and centrifuged at 30,000 x g for 15 min. Fractions pellet was resuspended in SDS sample buffer and analyzed by Western blotting (WB) using anti-GFP antibody (Abcam, ab 290, 1:1,000). GFP positive fractions (5 and 6) were subjected to TEM analysis to assess the level of MVB purification. The scaled-up procedure was applied to 2 gm of *yw* embryos, and the homogenization buffer volume was accordingly adjusted (all other steps were not modified). Fractions 5 and 6 were combined and subjected to proteomic analysis.

### Transmission electron microscopy (TEM)

Samples were fixed in a solution containing 4% paraformaldehyde (Electron Microscopy Sciences, EMS) and 2% glutaraldehyde (EMS) in 0.1 M cacodylate buffer containing 5 mM CaCl2 (pH 7.4). Fixed samples were drawn into cellulose capillary tubes with an inner diameter of 200 µm ^118^. Samples were postfixed in 1% osmium tetroxide (EMS) supplemented with 0.5% potassium hexacyanoferrate tryhidrate (BDH chemicals) and potassium dichromate (BDH chemicals) in 0.1 M cacodylate for one hour, stained with 2% uranyl acetate (EMS) in double distilled water for one hour, dehydrated in graded ethanol solutions and embedded in epoxy resin (Agar scientific Ltd.). Ultrathin sections (70 nm) were obtained with a Leica EMUC7 ultramicrotome. Sections were transferred to 200 mesh copper transmission electron microscopy grids (SPI). Grids were stained with lead citrate (Merk) and examined with a Tecnai T12 transmission electron microscope (Thermo Fisher Scientific). Digital electron micrographs were acquired with a bottom-mounted TVIPS TemCam-XF416 4k × 4k CMOS camera.

### Mass spectrometry

MVB-containing fractions and cell lysate control were resuspended in lysis buffer (5% SDS, 50 mM Tris pH 7.4) and centrifuged at 14,000 x g for 5 min. Pellets were discarded, and the supernatant was collected. Samples were reduced in 6 mM dithiothreitol and alkylated with 12 mM iodoacetamide in the dark. Each sample was loaded onto S-Trap minicolumns (Protifi, USA) according to the manufacturer’s instructions. In brief, after loading, samples were washed with a solution of 90% methanol and 10% 50 mM Triethylammonium bicarbonate. Samples were then digested with trypsin (Promega) at 1:50 trypsin/protein ratio for 90 min at 47°C. The digested peptides were eluted using 50 mM ammonium bicarbonate; trypsin was added to this fraction and incubated overnight at 37°C. Two more elutions were made using 0.2% formic acid and 0.2% formic acid in 50% acetonitrile. The three elutions were pooled together and vacuum-centrifuged to dry.

UPLC/MS grade solvents were used for all chromatographic steps. Each sample was loaded using splitless nanoUltra Performance Liquid Chromatography (10k psi nanoAcquity; Waters, Milford, MA, USA). The mobile phase was: A) H2O + 0.1% formic acid and B) acetonitrile + 0.1% formic acid. The samples were desalted online using a reversed-phase Symmetry C18 trapping column (180 µm internal diameter, 20 mm length, 5 µm particle size; Waters). The peptides were then separated using a T3 HSS nano-column (75 µm internal diameter, 250 mm length, 1.8 µm particle size; Waters) at 0.35 µL/min. Peptides were eluted from the column into the mass spectrometer using the following buffer B (99.9% Acetonitrile, 0.1% formic acid) gradient in three steps: (1) 4% to 25% of buffer B for 150 min, (2) 25% to 90% of buffer B for 5 min, (III) 5 min incubation in 90% buffer B, and then back to initial conditions. Samples were run in random order.

A nanoESI emitter (20 µm tip, Fossilion Tech, Madrid, Spain) was used on a FlexIon source (Thermo Fisher), mounted on a Tribrid orbitrap mass spectrometer (Fusion Lumos, Thermo Fisher). Data were acquired in data-dependent acquisition (DDA) mode, using a Top Speed method for 3 sec. MS1 resolution was set to 120,000@200 m/z, mass range of 380-1650 m/z, AGC at Standard and maximum injection time was set to 50 msec. Precursors selected for MS2 were limited to charge states of 2-8 with a minimum intensity of 50000. MS2 resolution was set to 15,000@200 m/z, quadrupole isolation 1 m/z, first mass at 130 m/z, AGC set at Standard. Dynamic exclusion of 30 sec with +/-10 ppm window and maximum injection time set to 100 ms. Ions were fragmented using HCD at 30 NCE.

The data analysis was performed using Byonic (v3.3.11) search engine (Protein Metrics) ^119^ against the *Drosophila melanogaster* proteome database (UniProt Nov 2018). Data were searched against specific C-terminal cleavage at K and R, allowing one missed cleavage at a tolerance of 10 ppm in MS1 and 20 ppm in MS2, using HCD settings for precursors up to 5500 Da. Modifications allowed were fixed Carbamidomethylation on C and variable modifications on the following amino acids: oxidation on M, deamidation on NQ and protein N-terminal acetylation. Data was filtered for 1% FDR. Quantification was performed by FlashLFQ ^120^ without match between runs or normalization since the samples were very different from one another. Protein annotations obtained from MS analysis were scored by the number of peptide sequences that are unique to a protein group (unique peptides value), and the number of peptide spectrum matches (PSM value), which is the total number of identified peptide spectra matched for the protein. Low abundance proteins (PSM<10) were disregarded, and Log2 fold change (log2FC) ratios were calculated to obtain a list of MVB-enriched proteins (log2FC>1). Pathway enrichment analysis was performed using Flymine ^121^.

### Western blotting

Samples were run in SDS-PAGE and transferred to nitroglycerin membrane. After blocking in 5% milk solution (dry milk in PBTw), the membrane was incubated with anti-GFP (Abcam, ab290 1:1000), followed by incubation with anti-rabbit HRP secondary antibody (1:10,000) for one hour at room temperature (Jackson Immuno-Research). After washes in PBTw, images were obtained using the ImageQuant LAS 4000 mini.

### Live imaging

#### PM elimination kinetic assay

Female flies of the appropriate genotype were crossed with males producing red-PM sperm in population cages, and were allowed to lay eggs on juice agar plates with a dollop of thick yeast paste for 15 min. Embryos were collected in mesh baskets, dechorionated in bleach and washed with water. Embryos were then aligned on an agar pad, picked up on an embryo glue (Permanent double-sided tape, Scotch^®^, 665)-coated coverslip and covered with halocarbon oil 27 and 700 (1:1 mixture, Sigma-Aldrich H8773 and H8898, respectively). Embryos were then imaged under a Nikon Eclipse fluorescence microscope. 2D images were acquired for each embryo at 1-hour intervals for the duration of 3 hours AEL.

### High resolution 3D live imaging

Embryos at 0–15 min AEL of the appropriate genotype were collected and mounted for imaging as described above. Imaging was performed using a Dragonfly Spinning disc confocal system (Andor Technology PLC) mounted on an inverted Leica Dmi8 microscope (Leica GMBH). When needed, images were deconvolved with the internal Andor Fusion deconvolution application. MIP (Max-Intensity-Projection) processing was performed using Imaris 9.5.0 (Bitplane, http://www.bitplane.com/, RRID:SCR_007370).

### ROS staining

WT embryos at 0–15 min AEL were collected and dechorionated as described above. The embryos were then permeabilized by 1 min incubation in EPS solution (90% D[+] Limonene [Thermo Fisher Scientific, FL/1860/07], 5% ethoxylated alcohol [Bio-Soft N1-7], 5% cocamide DEA [kind gift from Sano International; https://www.sano-international.com/], diluted 1:10 in pre-warmed [37°C] PBS) ^122^. Embryos were washed four times in PBS and incubated in 10 μM 2,7-Dichlorodihydrofluorescein diacetate solution (H2DCFDA; Cayman chemical, 85155) for 3 min. Embryos were then washed twice in PBS and mounted and imaged as described above.

### Embryo fixation and immunostaining

Embryos at 0–1-hour AEL of the appropriate genotype were collected and dechorionated as described above. Embryos were placed in boiling Triton/Salt solution (0.7% NaCl, 0.04% TritonX-100) for 5 sec, and immediately moved to cool Triton/Salt solution on ice for 15 min. Vitelline membranes were removed by manual shaking in 50% methanol: 50% heptane for 1 min. Embryos were rehydrated by passing through a series of solutions with decreasing methanol concentrations, washed with 1× PBS + 0.1% TritonX-100 (PBTx), and blocked in 5% normal goat serum in PBTx for one hour. Then, the embryos were incubated with the relevant primary antibody overnight at 4°C, washed in PBTx, and incubated with the secondary antibody at room temperature for 2 hours. Embryos were washed in PBTx and then mounted onto slides in Fluoromount medium (SouthernBiotech, Birmingham, AL, USA). Primary antibodies used in this study are biotin Anti-GFP antibody (1:100, Abcam, ab6658), anti-*Drosophila melanogaster* Atg8a polyclonal antibody (1:100, Creative-diagnostics,CABT-L1690), anti-pan polyglycylated Tubulin antibody, clone AXO 49 (1:5000, Sigma-Aldrich, MABS276). All secondary antibodies (Jackson Immuno-Research) were used at a dilution of 1:250. All images were acquired using the Andor Dragonfly Spinning disc confocal imaging system (Andor Technology PLC) mounted on an inverted Leica DMi8 microscope (Leica GMBH). Images in Extended Data Fig. 7a depicting immunostaining of mouse sperm with the anti-Rubicon antibody from Abcam were acquired using a Zeiss LSM900 confocal microscope.

### Image analysis

#### PM elimination

To quantify PM intensity, two channel 2D images were acquired for each embryo: A bright field (BF) channel and a fluorescent channel of the (MTS)tdTomato (red-PM). Embryos that failed to complete cellularization were omitted from the analysis. Each image includes one or two embryos. We used the BF channel to segment the whole embryo using Ilastik AutoContext pixel classifier ^123^. We trained the Ilastik classifier on multiple images from multiple different conditions. To avoid artifacts on the edge of the embryo, we eroded the embryo segments by 15 pixels. Small non-embryo segments were discarded from further analysis. We then identified positive PM regions by applying background subtraction with Rolling ball (sigma = 35 pixels) to the fluorescent channel and selecting all the pixels above a fixed selected value (1200). For each embryo, we measured the total and average signal in the whole embryo, PM-positive regions and PM-negative regions, and calculated the difference between the average PM-positive signal to the average PM-negative signal. We saved segmented regions on top of the BF channel and the fluorescent channel for quality control. The regions-of-interest of all embryos were saved into a file, to enable manual correction in case of incorrect embryo segmentation. Normalized PM intensity values were calculated for each embryo, by subtracting the mean intensity value of the background (PM-negative) from the mean PM intensity value (PM-positive), and multiplying by the fraction area of intensity above threshold, to take into account the PM size. The above workflow was implemented as a Fiji 125 macro ^124^, which also enables the above measurement from manually modified regions instead of automatic regions, to overcome segmentation problems.

#### Atg8a recruitment

To determine Atg8a recruitment levels to the sperm flagellum, we used an Arivis Vision4D pipeline to analyze the relative volume of Atg8a as a function of the distance from the axoneme, in 0–1-hour AEL embryos immunostained to reveal the axoneme and Atg8a. First, the axoneme was identified using a random forest pixel classifier. False-positive axoneme objects were manually removed. Then, the Atg8a signal was identified using a random forest pixel classifier. To save computation time and avoid background signal, the smallest 40% Atg8a objects were filtered out from each file, such that only 60% of the total Atg8a volume was further analyzed. We verified on a subset of the data that we get the same spatial distribution of the Atg8a objects when analyzing all the objects or just the bigger ones as described. Next, total Atg8a object volume was measured in 10 envelopes of 1 µm thick around the identified axoneme (the first envelope captures Atg8a objects with less than 1 µm distance from the axoneme, the second envelope captures Atg8a objects residing between 1 and 2 µm distance and so on). Then, the normalized Atg8a volume was obtained by dividing the total Atg8a volume within an envelope by the corresponding envelope volume.

### Deamination of relative RNA expression levels by RT-qPCR

Maternal gene knockdowns were validated by comparing the levels of the examined genes in control embryos (expressing the *matα-GAL4* driver alone) and embryos expressing corresponding *UAS-ShR* transgenes. Total RNA was isolated from 0–30 min AEL embryos, using the Quick-RNA Microprep Kit (Zymo Research, R1051). cDNA was then synthesized using the High-Capacity cDNA Reverse Transcription Kit (Applied Biosystems™, 4368814). Gene expression levels were determined by RT-qPCR using the StepOnePlus™ Real-Time PCR System (Applied Biosystems™, 4376600) with the KAPA SYBR® FAST qPCR Master Mix Kit (KAPA Biosystems, KR0389_S-v2.17). Expression values were normalized to the *αTub84B* gene (CG1913). Fold change in the expression levels in the knockdown validation experiments were analyzed using the ΔΔCt method ^125^, while relative mRNA levels in Fig. 4d are represented by ΔCt values. Primer sets were designed using the FlyPrimerBank online database ^126^ and are listed herein:

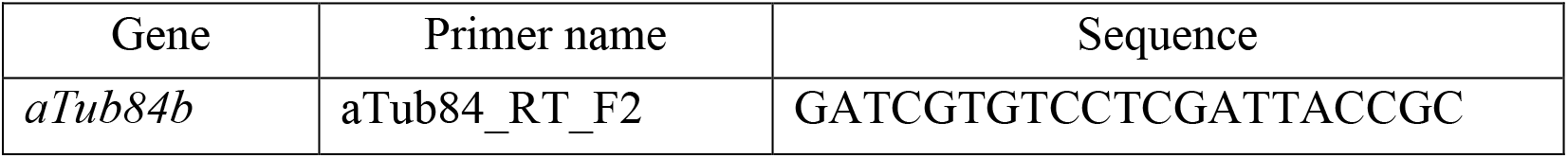
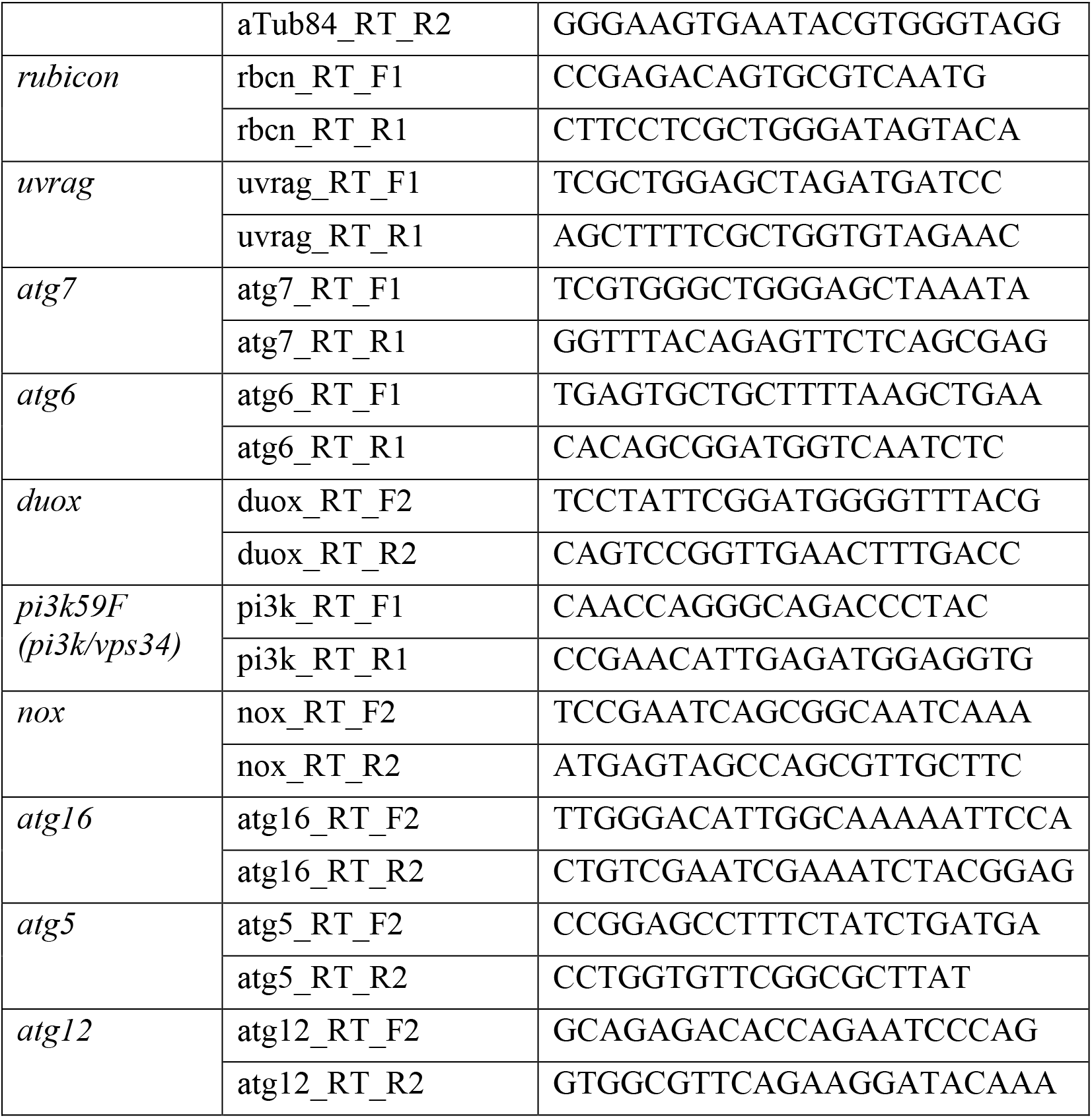

### Immunostaining of mammalian sperm

#### Mice

C57Bl/6JOlaHsd mice were purchased from Envigo (https://www.envigo.com/model/c57bl-6jolahsd). For sperm collection, male sperm donor mice were sacrificed, and the cauda epididymis were dissected and cleaned of the adipose and vascular tissue. Cauda were transferred into a 0.2 μM MitoTracker™ Green FM (Thermo Fisher Scientific, M7514), diluted in TYH medium^127^. Several cuts were scored in the cauda to allow the sperm to swim out. Cauda were incubated in 5% CO2, at 37°C for 20 min. 20 μl of labeled sperm were collected from the edge of the drop, washed in TYH and centrifuged at 300 x g for 1 min. Sperm were fixed in 4% paraformaldehyde for 30 min, permeabilized in 0.2% triton for 20 min, and blocked in 2% BSA, 0.05% Tween 20 for one hour. Sperm were then incubated with the primary antibody at 4°C overnight, washed in 0.05% Tween 20 and incubated with the secondary antibody at room temperature for two hours. 0.1 % BSA was added to all solutions to prevent stickiness. Primary antibodies used: anti-Rubicon antibody (1:100, Biorbyt, orb186000), Anti-Rubicon/Baron antibody (1:100, ab156052), anti-pan polyglycylated Tubulin antibody, clone AXO 49 (1:5000, Sigma-Aldrich, MABS276). All secondary antibodies were used at a dilution of 1:250 (Jackson Immuno-Research). Sperm were washed in 0.05% Tween 20, mounted onto slides in Fluoromount medium (SouthernBiotech, Birmingham, AL, USA) and visualized using a Dragonfly spinning disc microscope (Andor). All mouse experiments are approved by the Weizmann Institute’s IACUC committee and are carried out in accordance with their approved guidelines (https://www.weizmann.ac.il/vet/iacuc/introduction; IACUC approval 08101022-1).

#### Bovine

This protocol was adopted from ^128^ with minor modifications. Briefly, the experiments were conducted using frozen-thawed semen from the same mature working bull (SION; Israeli Company for Artificial Insemination and Breeding LtD), as previously described ^129^. Two straws, composing a single sample, were thawed at 38.5°C (cow normal core body temperature) in a pre-warmed bath for 1 min. The samples were then subjected to the swim-up procedure to ensure that further analysis will be performed on viable motile spermatozoa. Samples were washed once in prewarmed (38.5°C) Sperm-Tyrode lactate buffer (SP-TL; 0.09 M NaCl, 3.03 mM KCl, 24.9 mM NaHCO3, 3.47 µM NaH2PO4, 0.01 M HEPES, 2.03 mM CaCl2·2H2O, 1.13 µM MgCl2·6H2O and 0.19 % [v/v] Na-lactate adjusted to pH 7.4). The samples then were centrifuged at 600 x g for 10 min at room temperature, followed by 20 min at 38.5°C, allowing the viable motile sperm to swim up. After the incubation, a 10 µl of 0.01 mM MitoTracker™ Green FM (Thermo Fisher Scientific, M7514) was added into a 310 µL of washed swim-up cells. The samples were then incubated at 38.5°C for additional 10 min, vortexed and centrifuged at 600 x g for 5 min, and washed in SP-TL buffer. Samples were then fixed in 2% paraformaldehyde for 15 min, washed in PBS, and blocked in 0.1 M glycine, 0.15 mM BSA, 5% goat serum, 1 % milk in PBS for 45 min. Samples were washed in PBS and incubated with a primary antibody for 30 min, washed in PBS, and incubated with a secondary antibody for 30 min. Primary and secondary antibody staining, as well as further procedures, were performed as described for the mouse sperm.

### Data visualization, statistics, and reproducibility

Illustrations in Fig. 1d, Extended Data Fig. 1a, and Extended Data Fig. 7 were generated using BioRender.com. The pathway enrichment analysis graph was generated using R (R Core Team, 2020 https://www.r-project.org/, RRID:SCR_001905). Illustrations in Fig. 1a and Fig. 6 were generated using Adobe Illustrator (Adobe Inc. [2019]. Retrieved from https://adobe.com/products/illustrator). All graphs and statistical analyses in this manuscript were generated using the GraphPad Prism software version 9.5.1 for Windows (GraphPad Software, San Diego, California USA, www.graphpad.com). The specific statistical tests that were used for comparing the significance between experimental groups, as well as the experimental reproducibility, are all indicated in the relevant figure legends. Significance is indicated by asterisks as follows: \**p <* 0.05, \*\**p <* 0.01, \*\*\**p <* 0.001 and \*\*\*\**p <* 0.0001.

For all PM elimination kinetic assay experiments, 1-3 biological replicates were performed. Specific n numbers for each genotype are indicated in the figure legends. Note that for *rubicon*, several different mutant alleles and an shRNA transgene displayed similar phenotypes.

### Reporting summary

Further information on research design is available in the Nature Research Reporting Summary linked to this article.

## Data availability

The authors declare that the data supporting the findings of this study are available within the paper and its supplementary information files, and that all additional data are publicly available. MVB proteomics data are deposited into PRIDE repository, accession number [will be available before publication]. Costume codes for PME kinetics assay and Atg8a recruitment analysis are deposited into Github repository.

## Reagans and software

Reagents and software
| Reagent | Source | Identifier |
| --- | --- | --- |
| In-Fusion HD Cloning Kit | Takara-Clontech | 638909 |
| Quick-RNA Microprep Kit | Zymo Research | R1051 |
| High-Capacity cDNA Reverse Transcription Kit | Applied Biosystems™ | 4368814 |
| KAPA SYBR® FAST qPCR Master Mix Kit | KAPA Biosystems | KR0389_S-v2.17 |
| Halocarbon oil 27 | Sigma-Aldrich | H8773 |
| Halocarbon oil 700 | Sigma-Aldrich | H8898 |
| 2,7-Dichlorodihydrofluorescein diacetate (H2DCFDA) | Cayman chemical | 85155 |
| Protease inhibitor cocktail | Sigma-Aldrich | P8340 |
| OptiPrep™ Density Gradient Medium | Sigma-Aldrich | D1556 |
| MitoTracker™ Green FM | Thermo Fisher Scientific | M7514 |
| D(+) Limonene | Thermo Fisher Scientific | FL/1860/07 |
| Cocamide DEA | kind gift from Sano International<br>(https://www.sano-international.com/) |  |
| Ethoxylated alcohol (Bio-Soft N1-7) | kind gift from Sano International<br>(https://www.sano-international.com/) |  |
| Anti-GFP antibody | Abcam | Ab290 |
| Biotin Anti-GFP antibody | Abcam | Ab6658 |
| Anti-*Drosophila melanogaster* Atg8a polyclonal antibody | Creative-diagnostics | CABT-L1690 |
| Anti-pan polyglycylated Tubulin Antibody, clone AXO 49 | Sigma-Aldrich | MABS276 |
| Anti-Rubicon antibody | Biorbyt | orb186000 |
| Anti-Rubicon antibody | Abcam | Ab156052 |

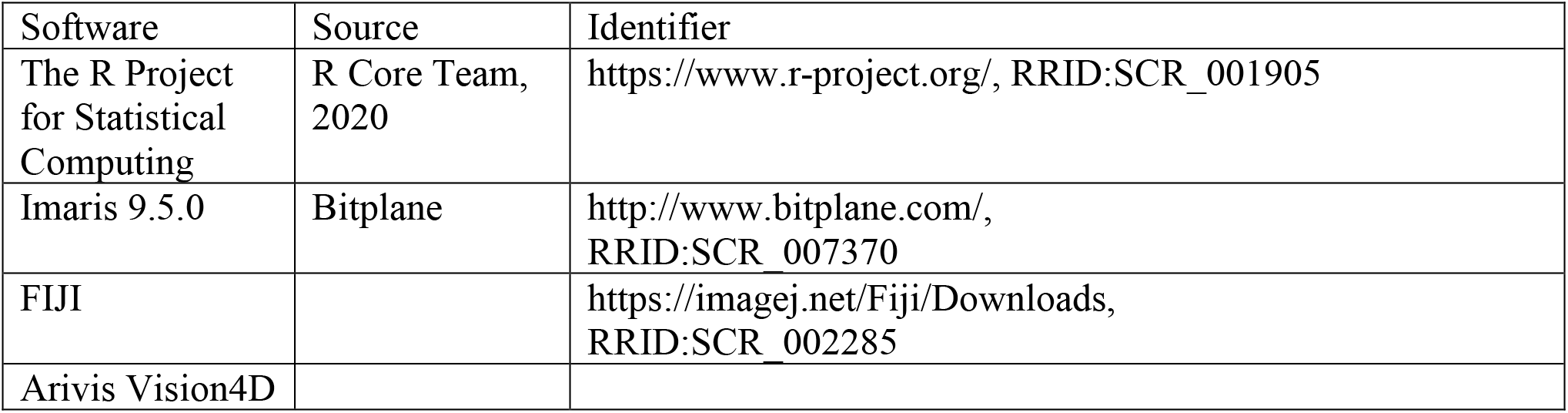
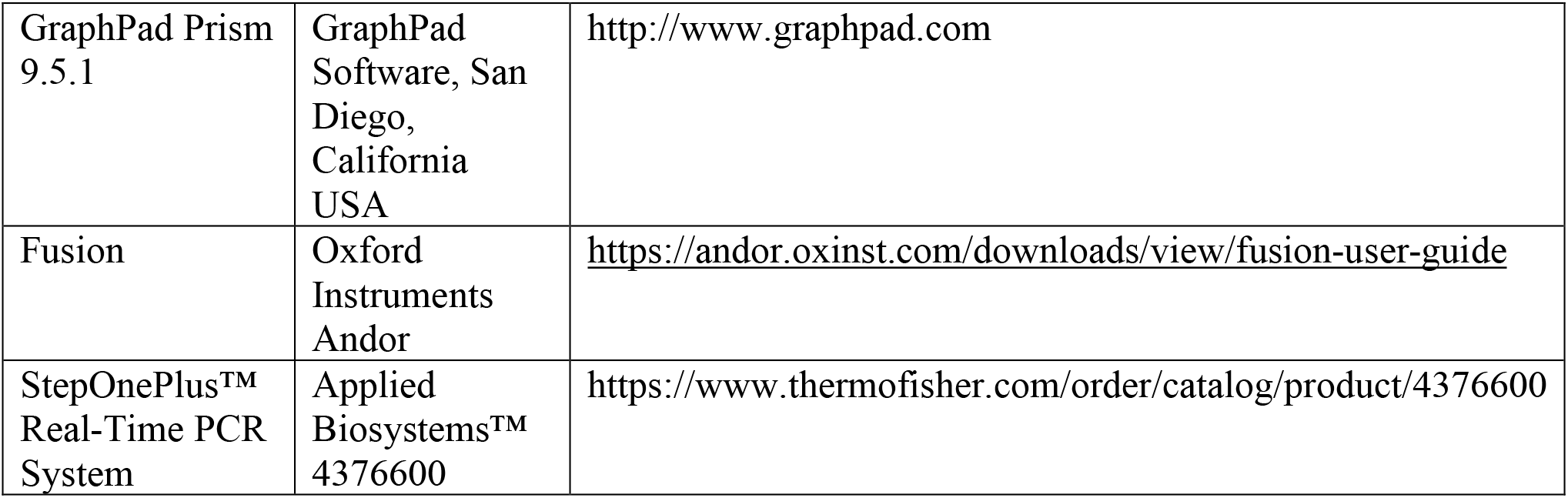

## Acknowledgments

We are grateful to Satomi Takeo, Graydon B. Gonsalvez, Robert Tsien, Suzanne Eaton, the *Drosophila* Genomics Resource Center (DGRC), the *Drosophila* RNAi Screening Center (DRSC) and Transgenic RNAi Project (TRiP), the Bloomington *Drosophila* Stock Center, and the Forchheimer Plasmid Bank of the Weizmann Institute of Science for providing additional stocks and reagents. We thank Johannes Bischof from FlyORF for help with generating endogenously tagged *rubicon*, Jordana Lindner-Ovadia for creating the graph in Extended Data Fig. 1c, Alina Kolpakova and Yoseph Addadi for help with microscopy imaging, Moshe Peretz for help with subcellular fractionation assay, Tsviya Olender for help with MS data analysis, and Ron Rotkopf for help with statistical analysis. We note Bar Lavi Lib, Doreen Padan, Ben Yashar, and Rebeca Gonzalez-Rolfe, for helping to carry out experiments during their rotation periods in the laboratory. We acknowledge the BioRender website which was used to create the illustrations presented in Fig. 1d, Extended Data Fig. 1a, and Extended Data Fig. 7. We warmly thank Genia Brodsky and Iryna Savych from the WIS Graphic Design Department for help with the graphic illustration in Fig. 1a and video editing and annotation, respectively. Electron microscopy studies were conducted at the Irving and Cherna Moskowitz Center for Nano and Bio-Nano Imaging at the Weizmann Institute of Science. We thank the Arama laboratory members for encouragement and advice. We warmly thank Eyal Schejter for support, advice, and excellent comments on the manuscript.

## Authors contributions

S.B-H. designed, performed, analyzed experiments, and performed statistical analyses. S.A. provided technical assistance with specific experiments. Y.P. and L.G. generated the Red-PM sperm producing transgenic flies (*dj-(MTS)tdTomato*) and the *UASp-eGFP-hCD63* fly line, respectively, and both were engaged in early stages of this study. O.G. wrote the data analysis code for the PM elimination kinetic assay and E.S. wrote the data analysis code for Atg8a recruitment assay. R.H.-K., E.M. and S.P. extracted and immunostained mice sperm. Z.R. and D.K. provided bovine sperm samples and related knowledge. N.D. performed the TEM experiments. D.M. performed the MS experiment and related initial analysis. S.P. helped with the pathway enrichment analysis of the proteomic data. K.Y.-S. helped with mice sperm experiments and was responsible for the supervision of some aspects of the study, E.A. led the project, designed experiments, interpreted results, was responsible for the general supervision of the study, wrote the manuscript, and procured funding.

