## Supplementary Material for "Egg MVBs elicit an antimicrobial pathway to degrade paternal mitochondria after fertilization"

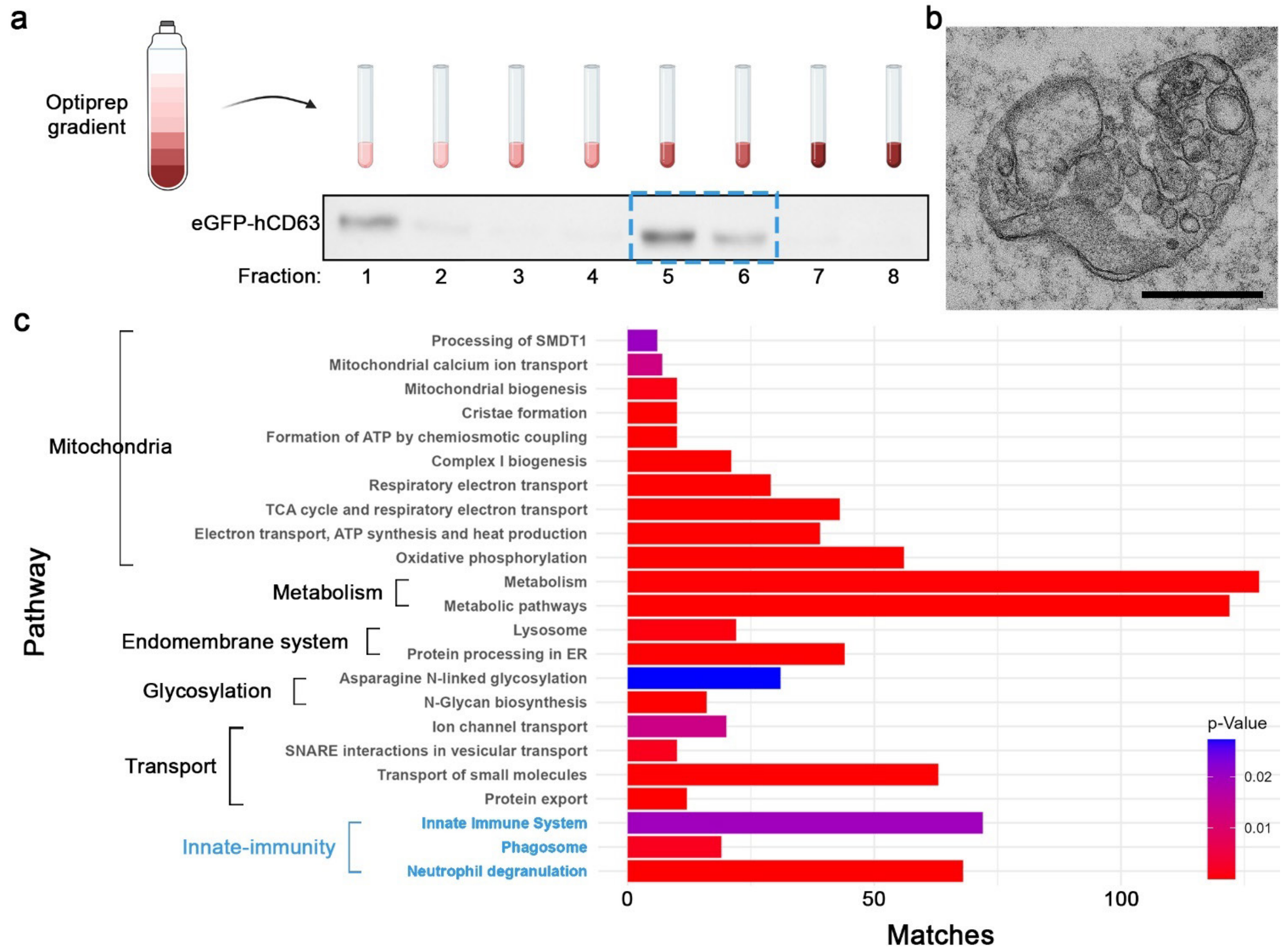


**Extended Data Fig. 1: Egg-derived MVBs contain components of innate immunity with elements of phagocytosis. a,** Western blot analysis of fractions obtained by OptiPrep$^{TM}$ density gradient isolation of MVBs from lysates prepared from eggs maternally expressing the MVB marker eGFP-hCD63. The peak of the eGFP signal appeared in fractions # 5 and 6 (band size, 52 kDa). Note that fraction #1 is the lowest density fraction, which likely contains the MVB unbound hCD63-eGFP. **b,** A TEM micrograph of a sample from combined fractions # 5 and 6, demonstrating the enrichment of egg MVBs in these fractions. Scale bar, 0.5 µm. **c,** Pathway enrichment analysis ($p < 0.05$) of the egg MVB proteins shows enrichment of innate-immunity ($p$ = 0.02) and phagosome ($p$ = 0.001) related factors. Enriched pathways are presented by the number of representative genes (matches), and the corresponding $p$ values (color scale). Pathway enrichment analysis was performed using the Flymine website.

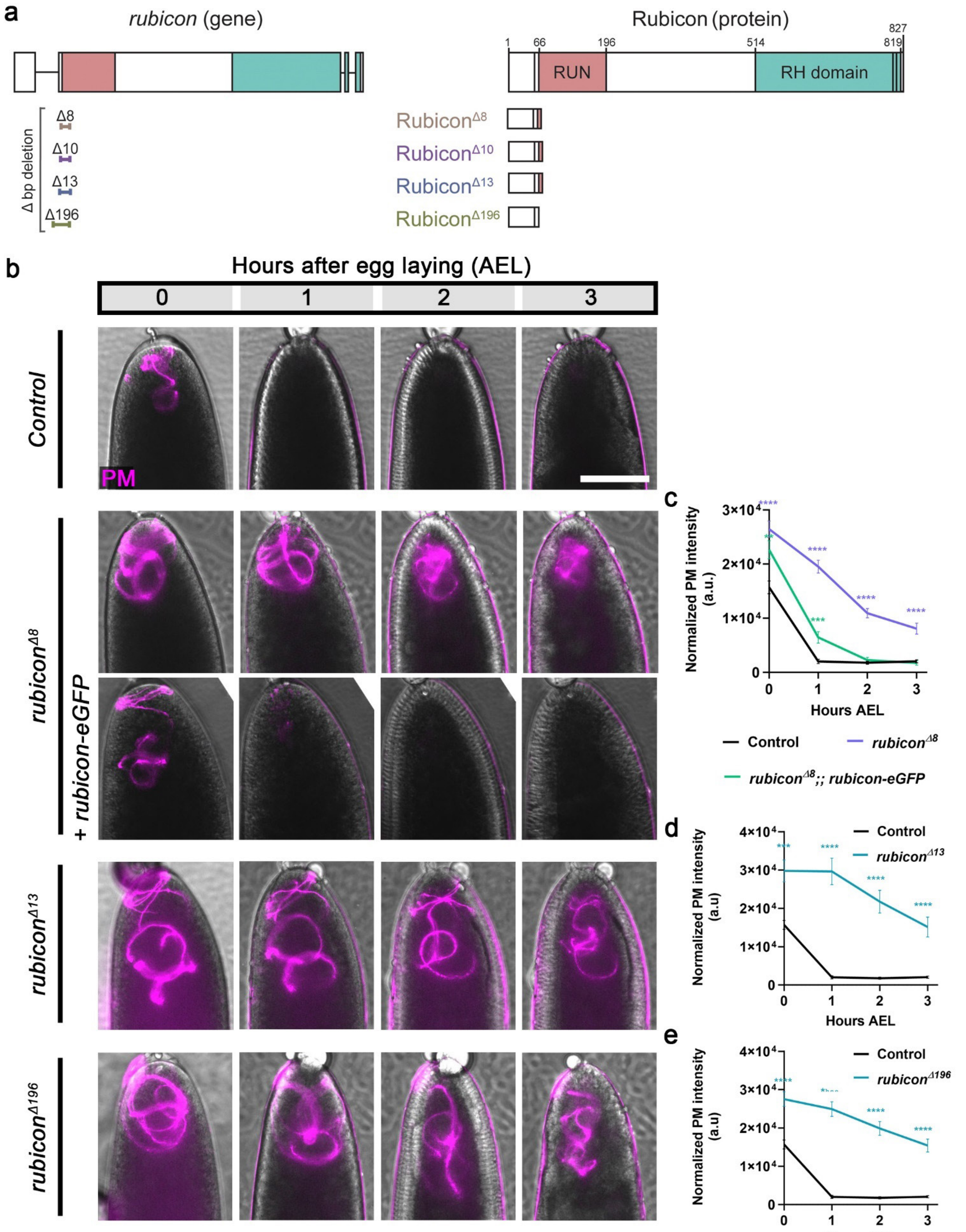


**Extended Data Fig. 2: PME is significantly attenuated in *rubicon* mutant embryos. a,** Schematic representations of the *rubicon* gene and mutant alleles (left), and the matching predicted proteins (right). Exons and introns are indicated by thick and thin bars of the DNA structure, respectively. The relative locations of the RUN (pink) and Rubicon homology (RH) (turquoise) domains are indicated. Different colored bars below the DNA structure indicate the locations of the deletions in the different *rubicon* mutant alleles, which result in frameshifts and consequent premature stop codons. **b,** PME is significantly attenuated in early fertilized eggs laid by females homozygous for the *rubicon* mutant alleles. A live imaging assay performed and presented as in Fig. 1e. Note the near complete restoration of PME kinetics in the $rubicon^{\Delta 8}$ mutant egg upon maternal expression of the *rubicon-eGFP* transgene. Scale bar, 100 μm. **c-e,** Quantifications of normalized red-PM fluorescence intensities (arbitrary units [a.u.]) in embryos represented in (**b**). Error bars indicate SEM. The respective numbers of scored embryos (n) laid by females of the control, $rubicon^{\Delta 8}$, $rubicon^{\Delta 8}$;; *rubicon-eGFP*, $rubicon^{\Delta 13}$, and $rubicon^{\Delta 196}$, are 61, 50, 47, 47, and 53. $^{**}p < 0.01$, $^{***}p < 0.001$, and $^{****}p < 0.0001$. Statistical tests for (**c**), two-way repeated measures ANOVA, followed by Dunnett's multiple comparisons test. Statistical tests for (**d,e**), two-way repeated measures ANOVA, followed by Šídák's multiple comparisons test.

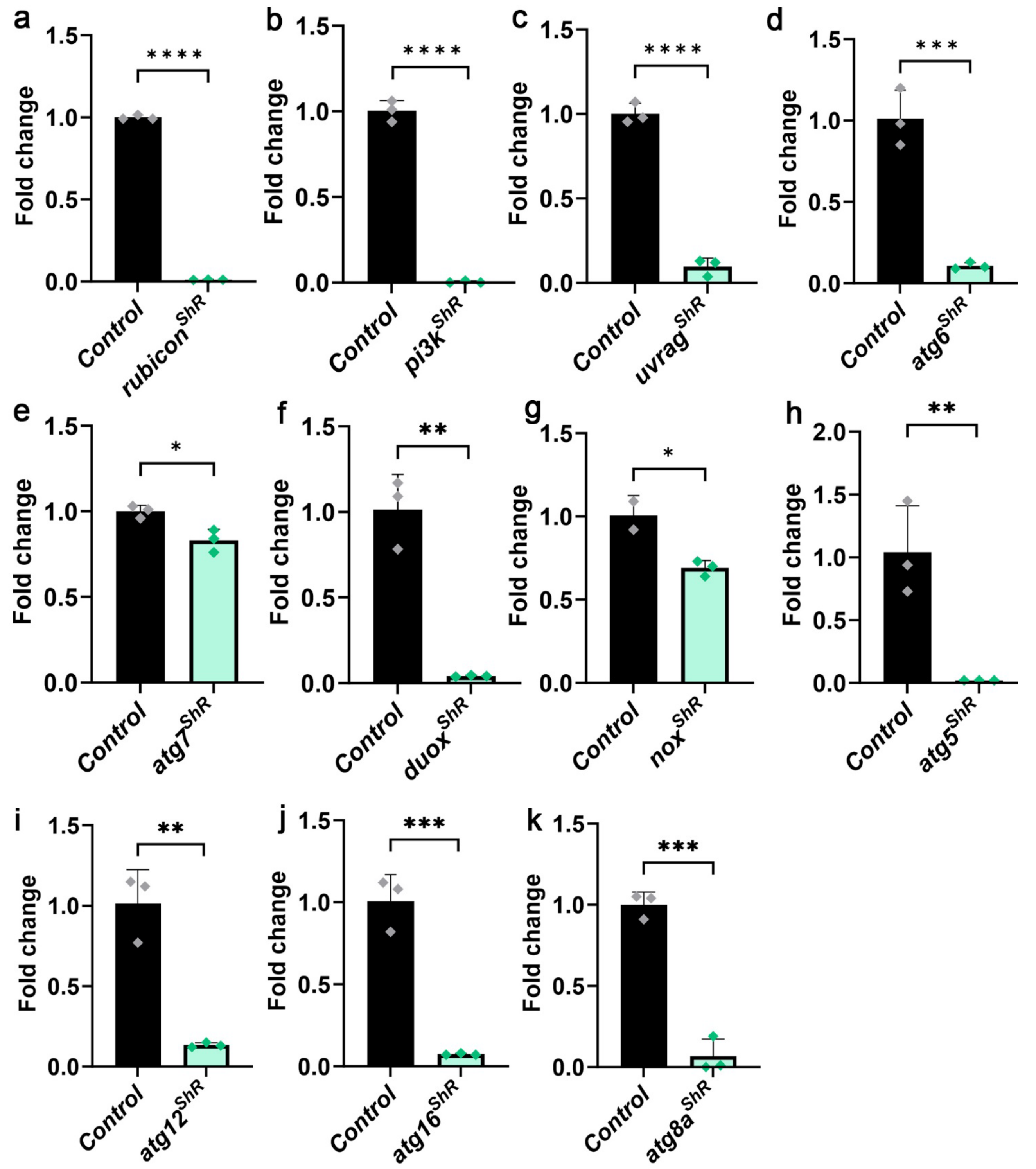


**Extended Data Fig. 3: Validation of the shRNA-mediated knockdowns used in this study. a-k,** Quantitative RT-qPCR analysis in early fertilized eggs with maternal knockdown of *rubicon*, $p < 0.0001$, n=3 (**a**); *pi3k*, $p < 0.0001$, n=3 (**b**); *uvrag*, $p < 0.0001$, n=3 (**c**); *atg6*, $p = 0.0009$, n=3 (**d**); *nox*, $p = 0.0221$, n=3 (**e**); *duox*, $p = 0.0012$, n=3 (**f**); *atg7*, $p = 0.0170$, n=3 (**g**); *atg5*, $p = 0.0088$, n=3 (**h**);

*atg12*, *p* = 0.0020, n=3 (**i**); *atg16*, *p* = 0.0006, n=3 (**j**); *atg8a*, *p* = 0.0003, n=3 (**k**). The maternal driver line (*matα-GAL4*/+) was used as control. Two-tailed unpaired student's t-test. Error bars indicate standard deviation (SD).

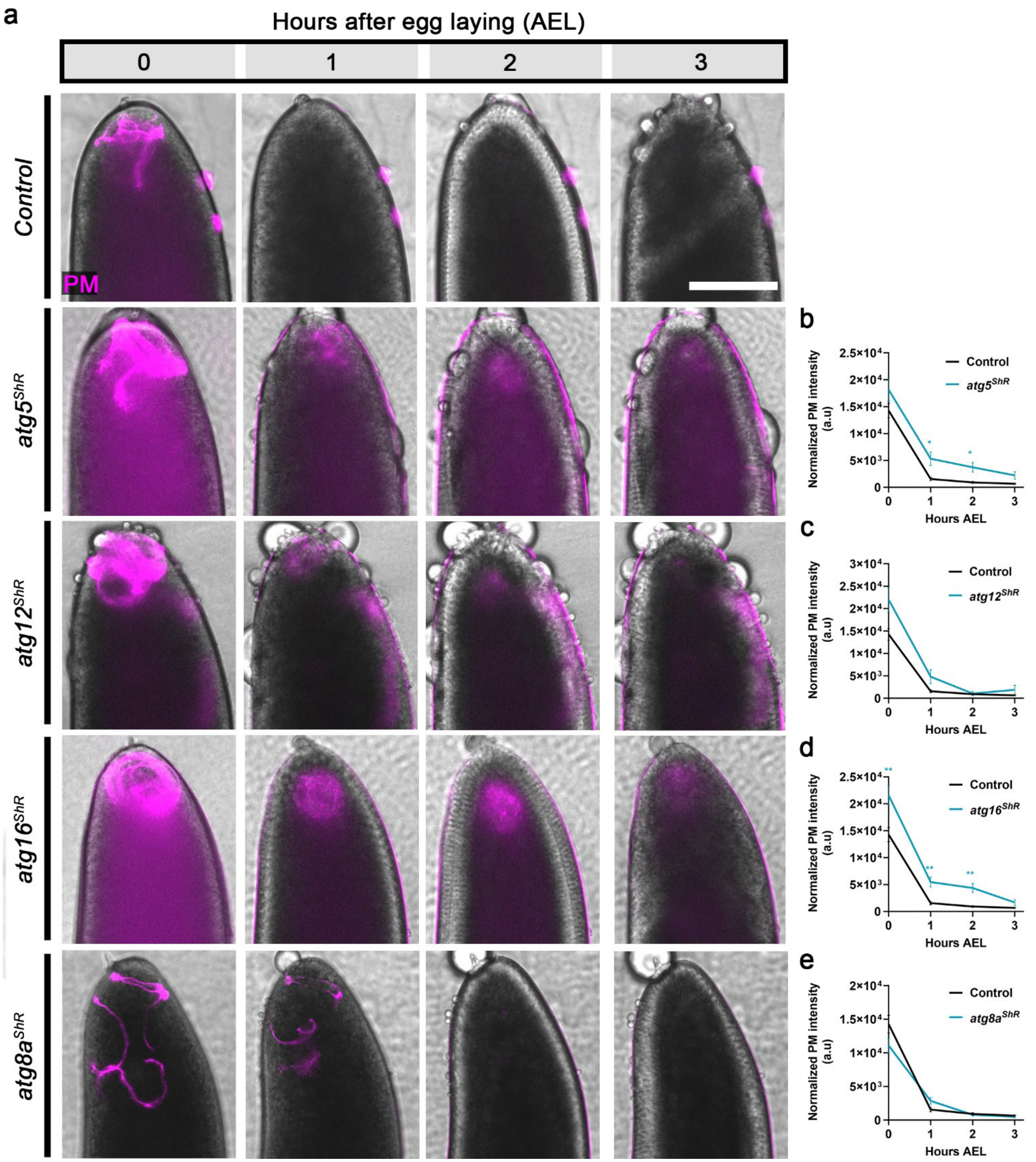


**Extended Data Fig. 4: Atg8a recruitment to the FVS occurs at an advanced decaying stage of the PM transgenic fluorescent signal. a,** A live imaging assay performed and presented as in Fig. 1e. Scale bar, 100 µm. **b-e,** Quantifications of normalized red-PM fluorescence intensities (arbitrary units [a.u.]) in embryos represented in (**a**). Error bars indicate SEM. The respective numbers of scored embryos (n) laid by females of the control, *atg5*$^{ShR}$, *atg12*$^{ShR}$; *atg16*$^{ShR}$, and *atg8a*$^{ShR}$, are 58, 28, 26, 34, and 51.

$*p < 0.05$ and $**p < 0.01$. Two-way repeated measures ANOVA, followed by Šídák's multiple comparisons test. Note the minor to no effect on the kinetics of the decaying PM transgenic fluorescent signal upon maternal shRNA-mediated knockdowns of major components of the Atg8/LC3 conjugation machinery, *atg5*, *atg12*, and *atg16*, as well as of *atg8a* itself.

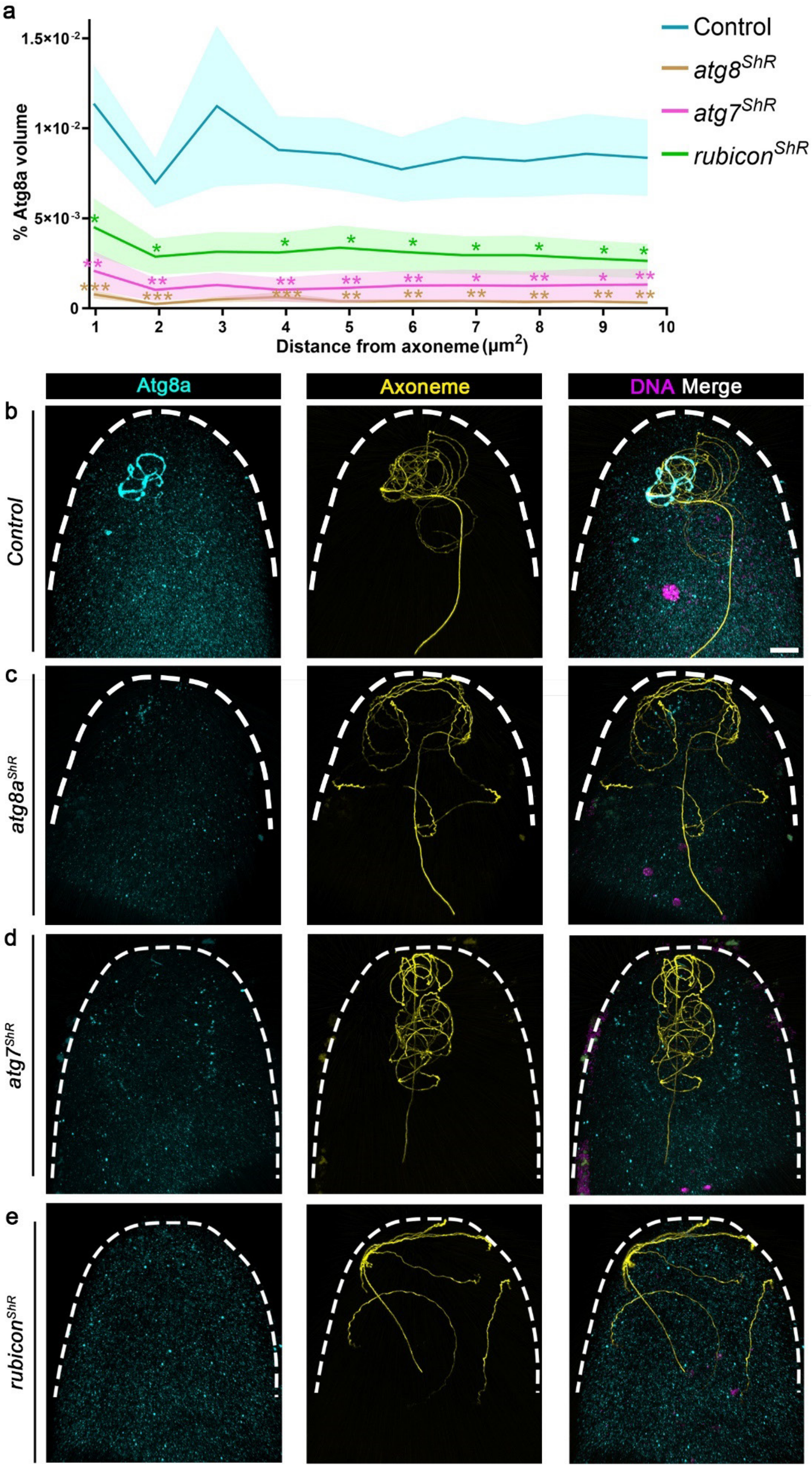


**Extended Data Fig. 5: Atg8a recruitment to the FVS requires Rubicon and Atg7. a,** The graphs depict staining volumes of Atg8a in 0–1-hour AEL fertilized eggs maternally expressing shRNA (*ShR*) transgenes against *atg8a*, *atg7* and *rubicon*. Corresponding WT early fertilized eggs served as control. Calculations were performed in intervals of 1 μm radius area around the axoneme for up to 10 μm. The respective numbers of scored embryos (n) laid by females of the control, $atg8a^{ShR}$, $atg7^{ShR}$, and $rubicon^{ShR}$, are 20, 12, 11, and 15. *$p < 0.05$, **$p < 0.01$, ***$p < 0.001$ and ****$p < 0.0001$. One-way ANOVA, followed by Holm-Šídák's multiple comparisons test. Error bars indicate SEM. Corresponding graphs depicting the Atg8a staining volumes in a limited 0-1 μm radius area around the axoneme are presented in Fig. 5c. **b,** Representative confocal images of the embryos used for the calculations in (**a**). Atg8a (cyan) and the axoneme (yellow) were detected by immunostaining. The 0–1-hour AEL embryos were further staged by the number of nuclei, as revealed by DAPI staining of the DNA (magenta).

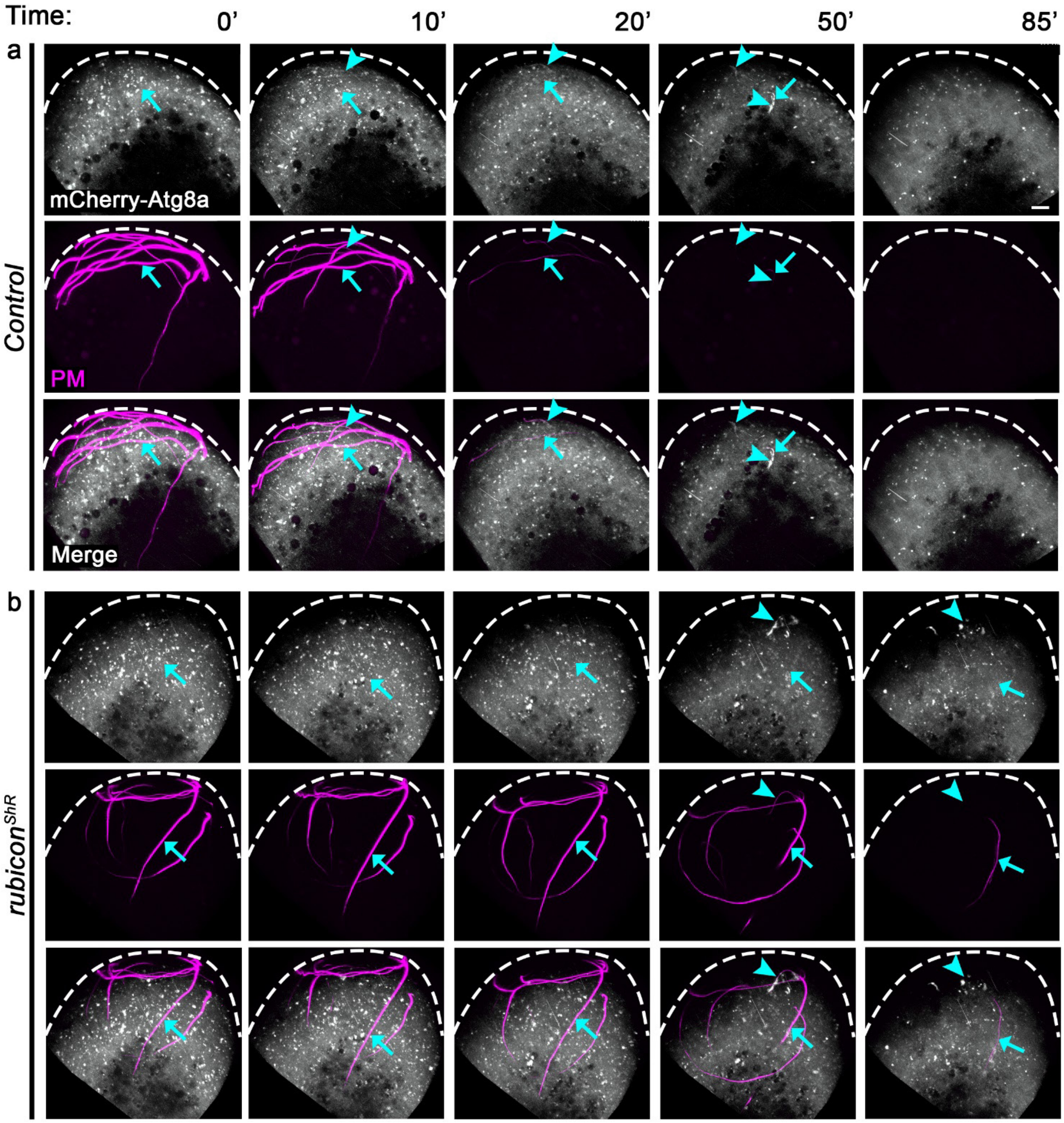


**Extended Data Fig. 6: Recruitment of Atg8a to the FVS is significantly delayed and inefficient in the absence of Rubicon. a,b,** Live imaging of the anterior regions of two early developing embryos fertilized by green-PM sperm (magenta) and maternally expressing an *mCherry-Atg8a* transgene (white) under the *matα-Gal4* driver (control, **a**) and, in addition, an shRNAi transgene against *rubicon* (**b**). Shown are representative images from Supplementary Video 5 (**a**) and Supplementary Video 6 (**b**) taken at the indicated min AEL. The arrows track the same regions on the PM as the embryos develop, while the arrowheads track mCherry-Atg8a recruitment to the same regions on the FVS. Whereas conjugation of mCherry-Atg8a to the FVS in an otherwise WT background is relatively fast (within 10-20 min AEL), with only a few short FVS segments displaying recruitment of mCherry-Atg8a at 40 min AEL (**a**; arrowheads), in the *rubicon* knockdown background, recruitment of the mCherry-Atg8a to the FVS is highly inefficient, such that it appears very late (50 min AEL) and only on a few short FVS segments (**b**; arrowheads). Scale bar, 10 μm.

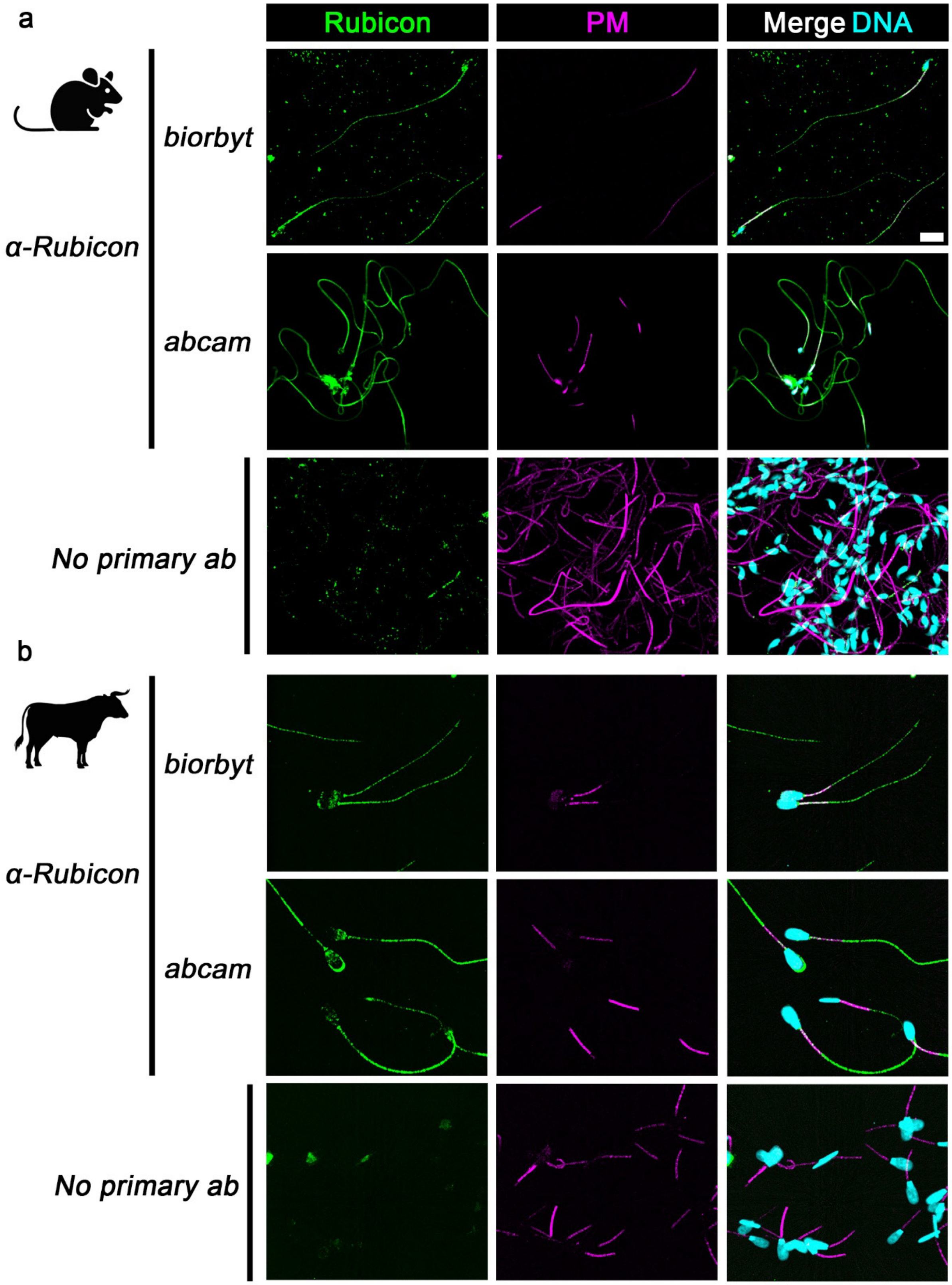


**Extended Data Fig. 7: Rubicon is expressed on the flagellum of mature mammalian sperm cells.**

Representative images of mature sperm cells from mice (**a**) and bulls (**b**) immunostained with two different anti-Rubicon antibodies (green; obtained from Biorbyt and Abcam). Secondary antibody only staining was performed to exclude possible non-specific staining. The sperm mitochondria are visualized with MitoTracker (magenta) and the sperm nuclei with DAPI (light blue). Scale bar, 10 μm.

**Description of Additional Supplementary Files**

File Name: Supplementary Video 1

Description:

**Live imaging of PME after fertilization in early *Drosophila* embryo.** A time-lapse confocal microscopy movie of the anterior region of a WT egg fertilized by red-PM sperm (magenta). The early fertilized egg was imaged for 43 min AEL. Note the PM pieces that bud off from the flagellum as PME advances (arrow). Scale bar, 10 μm.

File Name: Supplementary Video 2

Description:

**Live imaging of PME in an early *Drosophila* embryo maternally expressing the *rubicon-eGFP* transgene.** A time-lapse confocal microscopy movie of the anterior region of a WT egg maternally expressing Rubicon-eGFP (green) and fertilized by red-PM sperm (magenta). Rubicon positive MVBs engage and densely coat the sperm flagellum, forming FVSs that enwrap large flagellar segments withing which the PM is degraded. The early fertilized egg was imaged for 45 min AEL. Scale bar, 10 μm.

File Name: Supplementary Video 3

Description:

**Live imaging of PME in an early *Drosophila* embryo maternally expressing a PtdIns(3)P reporter.** A time-lapse confocal microscopy movie of the anterior region of a WT egg maternally expressing the PtdIns(3)P reporter TagRFPt-2xFYVE (magenta) and fertilized by green-PM sperm (green). The early fertilized egg was imaged for 15 min AEL. Note the association of PtdIns(3)P on the FVS (arrow), indicative of PI3KC3 activity. Scale bar, 10 μm.

File Name: Supplementary Video 4

Description:

**Live imaging of PME in an early *Drosophila* embryo maternally expressing both the *rubicon-eGFP* and an *mCherry-atg8a* transgenes.** A time-lapse confocal microscopy movie of the anterior region of a WT egg maternally expressing Rubicon-eGFP (green) and mCherry-Atg8a (magenta), and fertilized by WT sperm. The early fertilized egg was imaged for 60 min AEL. Note the 6-12 min lag between the formation of Rubicon-eGFP positive FVS and the consecutive recruitment of mCherry-Atg8a. Scale bar, 5 μm.

File Name: Supplementary Video 5

Description:

**Live imaging of PME in an early *Drosophila* embryo maternally expressing an *mCherry-atg8a* transgene.** A time-lapse confocal microscopy movie of the anterior region of a WT egg maternally expressing mCherry-Atg8a (white) and fertilized by green-PM sperm (magenta). The early fertilized egg was imaged for 90 min AEL. Note that mCherry-Atg8a is recruited to large PM segments within 10 minutes after egg laying (arrow). Scale bar, 10 μm.

File Name: Supplementary Video 6

Description:

**Live imaging of PME in an early *Drosophila* embryo compromised for *rubicon* and maternally expressing an *mCherry-atg8a* transgene.** A time-lapse confocal microscopy movie of the anterior region of an egg maternally expressing *rubicon*$^{ShR}$ and mCherry-Atg8a (white) and fertilized by green-PM sperm (magenta). The early fertilized egg was imaged for 3 hours AEL. Note that in the few events when mCherry-Atg8a is recruited to the FVS, it occurs only after 50 min AEL and on short FVS segments (arrow), while most of the flagellum is not associated with mCherry-Atg8a. Scale bar, 10 μm.

File Name: Supplementary Table 1

Description:

**Protein profiling of the egg MVBs.**

Sheet 1: Total proteins identified by the MS analysis in the MVB and lysate (control) fractions.

Sheet 2: MVB enriched proteins considered for the pathway enrichment analysis. The list contains all the proteins for which the fold change ratio (log2 of MVB PSM-log2 of lysate PSM) is above 1. In addition, proteins with less than 10 PSMs were discarded.
