## Supplementary Table 1 for "Egg MVBs elicit an antimicrobial pathway to degrade paternal mitochondria after fertilization"

| UniProt ID | Protein description | # PSMs MVB spectra | # of unique peptides MVB | # of PSMs lysate | # of unique peptides lysate | log2(PSM MVB) | log2(PSM lysate) | FC (log2 of MVB PSM-log2 of lysate PSM) |
|---|---|---|---|---|---|---|---|---|
| P98163 | YL_DROME Putative vitellogenin receptor OS=Drosophila melanogaster OX=7227 GN=yl PE=1 SV=2 | 388 | 97 | 131 | 45 | 8.599912842 | 7.033423002 | 1.566489841 |
| P22700 | ATC1_DROME Calcium-transporting ATPase sarcoplasmic/endoplasmic reticulum type OS=Drosophila melanogaster OX=7227 GN=Ca-P60A PE=1 SV=2 | 358 | 71 | 58 | 29 | 8.483815777 | 5.857980995 | 2.625834782 |
| Q94920 | VDAC_DROME Voltage-dependent anion-selective channel OS=Drosophila melanogaster OX=7227 GN=porin PE=1 SV=3 | 304 | 33 | 70 | 20 | 8.247927513 | 6.129283017 | 2.118644496 |
| Q9V496 | APLP_DROME Apolipophorins OS=Drosophila melanogaster OX=7227 GN=apolpp PE=1 SV=2 | 289 | 123 | 341 | 122 | 8.174925683 | 8.413627929 | -0.238702247 |
| P29844 | BIP_DROME Endoplasmic reticulum chaperone BiP OS=Drosophila melanogaster OX=7227 GN=Hsc70-3 PE=1 SV=2 | 279 | 51 | 94 | 25 | 8.124121312 | 6.554588852 | 1.56953246 |
| X2JB48 | X2JB48_DROME Stress-sensitive B, isoform E OS=Drosophila melanogaster OX=7227 GN=sesB PE=1 SV=1 | 274 | 38 | 78 | 23 | 8.098032083 | 6.285402219 | 1.812629864 |
| P13607 | ATNA_DROME Sodium/potassium-transporting ATPase subunit alpha OS=Drosophila melanogaster OX=7227 GN=Atpalpha PE=1 SV=3 | 204 | 58 | 22 | 14 | 7.672425342 | 4.459431619 | 3.212993723 |
| P02843 | VIT1_DROME Vitellogenin-1 OS=Drosophila melanogaster OX=7227 GN=Yp1 PE=1 SV=1 | 198 | 38 | 496 | 47 | 7.62935662 | 8.95419631 | -1.32483969 |
| P54399 | PDI_DROME Protein disulfide-isomerase OS=Drosophila melanogaster OX=7227 GN=Pdi PE=2 SV=1 | 190 | 41 | 86 | 30 | 7.569855608 | 6.426264755 | 1.143590854 |
| Q9V3U6 | Q9V3U6_DROME 26-29kD-proteinase OS=Drosophila melanogaster OX=7227 GN=26-29-p PE=1 SV=1 | 190 | 48 | 60 | 23 | 7.569855608 | 5.906890596 | 1.662965013 |
| A1Z6H7 | A1Z6H7_DROME Gp210 ortholog, isoform A OS=Drosophila melanogaster OX=7227 GN=Gp210 PE=1 SV=1 | 189 | 69 | 12 | 10 | 7.562242424 | 3.584962501 | 3.977279923 |
| Q9VEG6 | PERC_DROME Chorion peroxidase OS=Drosophila melanogaster OX=7227 GN=Pxt PE=2 SV=3 | 181 | 51 | 58 | 26 | 7.499845887 | 5.857980995 | 1.641864892 |
| Q95029 | CATL_DROME Cathepsin L OS=Drosophila melanogaster OX=7227 GN=Cp1 PE=2 SV=2 | 176 | 34 | 1 | 1 | 7.459431619 | 0 | 7.459431619 |
| Q3YMU0 | Q3YMU0_DROME Protein disulfide-isomerase (Fragment) OS=Drosophila melanogaster OX=7227 GN=ERp60 PE=1 SV=1 | 174 | 33 | 70 | 18 | 7.442943496 | 6.129283017 | 1.313660479 |
| A1Z843 | A1Z843_DROME CG1371 OS=Drosophila melanogaster OX=7227 GN=CG1371 PE=1 SV=1 | 172 | 53 | 19 | 12 | 7.426264755 | 4.247927513 | 3.178337241 |
| Q9VAY2 | Q9VAY2_DROME Glycoprotein 93 OS=Drosophila melanogaster OX=7227 GN=Gp93 PE=1 SV=1 | 162 | 48 | 75 | 24 | 7.339850003 | 6.22881869 | 1.111031312 |
| A0A0B4LFR4 | A0A0B4LFR4_DROME Oligosaccharide transferase delta subunit, isoform B OS=Drosophila melanogaster OX=7227 GN=OstDelta PE=1 SV=1 | 158 | 33 | 24 | 14 | 7.303780748 | 4.584962501 | 2.718818247 |
| L0MQ04 | L0MQ04_DROME ATP synthase subunit beta OS=Drosophila melanogaster OX=7227 GN=ATPsynbeta PE=1 SV=1 | 157 | 29 | 77 | 30 | 7.294620749 | 6.266786541 | 1.027834208 |
| Q9VKM7 | Q9VKM7_DROME AT01533p OS=Drosophila melanogaster OX=7227 GN=CG6087 PE=1 SV=2 | 157 | 31 | 24 | 10 | 7.294620749 | 4.584962501 | 2.709658248 |
| Q9VP27 | Q9VP27_DROME Golgi complex-localized glycoprotein 1, isoform A OS=Drosophila melanogaster OX=7227 GN=Glg1 PE=1 SV=2 | 147 | 41 | 1 | 1 | 7.199672345 | 0 | 7.199672345 |
| P11147 | HSP7D_DROME Heat shock 70 kDa protein cognate 4 OS=Drosophila melanogaster OX=7227 GN=Hsc70-4 PE=1 SV=3 | 141 | 39 | 311 | 57 | 7.139551352 | 8.28077077 | -1.141219418 |
| Q9VHY6 | Q9VHY6_DROME ER membrane protein complex 1, isoform A OS=Drosophila melanogaster OX=7227 GN=EMC1 PE=1 SV=1 | 137 | 46 | 14 | 10 | 7.098032083 | 3.807354922 | 3.290677161 |
| X2JEX8 | X2JEX8_DROME Yolk protein 3, isoform B OS=Drosophila melanogaster OX=7227 GN=Yp3 PE=1 SV=1 | 136 | 29 | 412 | 36 | 7.087462841 | 8.686500527 | -1.599037686 |
| TRYP_PIG | (Common contaminant protein) | 135 | 14 | 64 | 12 | 7.076815597 | 6 | 1.076815597 |
| O16043 | O16043_DROME Anon1A4 OS=Drosophila melanogaster OX=7227 GN=Df31 PE=1 SV=1 | 134 | 23 | 275 | 31 | 7.06608919 | 8.103287808 | -1.037198618 |
| P02844 | VIT2_DROME Vitellogenin-2 OS=Drosophila melanogaster OX=7227 GN=Yp2 PE=1 SV=2 | 124 | 31 | 336 | 43 | 6.95419631 | 8.392317423 | -1.438121112 |
| Q86BM5 | Q86BM5_DROME A kinase anchor protein 200, isoform D OS=Drosophila melanogaster OX=7227 GN=Akap200 PE=1 SV=1 | 124 | 23 | 2 | 2 | 6.95419631 | 1 | 5.95419631 |
| P35381 | ATPA_DROME ATP synthase subunit alpha, mitochondrial OS=Drosophila melanogaster OX=7227 GN=blw PE=1 SV=2 | 114 | 33 | 61 | 22 | 6.832890014 | 5.930737338 | 0.902152677 |
| Q7KMM4 | Q7KMM4_DROME BcDNA.GH04962 OS=Drosophila melanogaster OX=7227 GN=GCS2alpha PE=1 SV=1 | 112 | 44 | 34 | 18 | 6.807354922 | 5.087462841 | 1.719892081 |
| P10987 | ACT1_DROME Actin-5C OS=Drosophila melanogaster OX=7227 GN=Act5C PE=1 SV=4 | 111 | 26 | 262 | 31 | 6.794415866 | 8.033423002 | -1.239007135 |
| Q9VAL7 | Q9VAL7_DROME Calnexin 99A, isoform A OS=Drosophila melanogaster OX=7227 GN=Cnx99A PE=1 SV=2 | 109 | 31 | 1 | 1 | 6.768184325 | 0 | 6.768184325 |
| P08736 | EF1A1_DROME Elongation factor 1-alpha 1 OS=Drosophila melanogaster OX=7227 GN=eEF1alpha1 PE=1 SV=2 | 107 | 20 | 208 | 29 | 6.741466986 | 7.700439718 | -0.958972732 |
| P02828 | HSP83_DROME Heat shock protein 83 OS=Drosophila melanogaster OX=7227 GN=Hsp83 PE=1 SV=1 | 105 | 34 | 311 | 52 | 6.714245518 | 8.28077077 | -1.566525252 |
| Q7KRW4 | Q7KRW4_DROME Aminopeptidase OS=Drosophila melanogaster OX=7227 GN=CG14516 PE=1 SV=1 | 105 | 32 | 13 | 10 | 6.714245518 | 3.700439718 | 3.0138058 |
| Q9V491 | Q9V491_DROME Plexin A, isoform A OS=Drosophila melanogaster OX=7227 GN=PlexA PE=1 SV=1 | 105 | 58 | 8 | 8 | 6.714245518 | 3 | 3.714245518 |
| Q9VJD1 | Q9VJD1_DROME Glucosidase 2 beta subunit, isoform A OS=Drosophila melanogaster OX=7227 GN=GCS2beta PE=1 SV=1 | 104 | 29 | 32 | 14 | 6.700439718 | 5 | 1.700439718 |
| Q0E8E8 | Q0E8E8_DROME MIP08013p1 OS=Drosophila melanogaster OX=7227 GN=Mpcp2 PE=1 SV=1 | 103 | 27 | 27 | 9 | 6.686500527 | 4.754887502 | 1.931613025 |
| Q8T3L1 | Q8T3L1_DROME Acyl-CoA synthetase long-chain, isoform A OS=Drosophila melanogaster OX=7227 GN=Acsl PE=1 SV=1 | 102 | 38 | 16 | 12 | 6.672425342 | 4 | 2.672425342 |
| Q09332 | UGGG_DROME UDP-glucose:glycoprotein glucosyltransferase OS=Drosophila melanogaster OX=7227 GN=Ugt PE=1 SV=2 | 101 | 49 | 45 | 24 | 6.658211483 | 5.491853096 | 1.166358386 |
| P13060 | EF2_DROME Elongation factor 2 OS=Drosophila melanogaster OX=7227 GN=EF2 PE=1 SV=4 | 95 | 41 | 259 | 68 | 6.569855608 | 8.016808288 | -1.446952679 |
| Q9VMA7 | TGO1_DROME Transport and Golgi organization protein 1 OS=Drosophila melanogaster OX=7227 GN=Tango1 PE=1 SV=2 | 95 | 44 | 1 | 1 | 6.569855608 | 0 | 6.569855608 |
| M9PJN8 | M9PJN8_DROME Glyceraldehyde-3-phosphate dehydrogenase OS=Drosophila melanogaster OX=7227 GN=Gapdh2 PE=3 SV=1 | 94 | 23 | 173 | 27 | 6.554588852 | 7.434628228 | -0.880039376 |
| A0A0B4LFZ4 | A0A0B4LFZ4_DROME TER94, isoform E OS=Drosophila melanogaster OX=7227 GN=TER94 PE=1 SV=1 | 92 | 34 | 6 | 5 | 6.523561956 | 2.584962501 | 3.938599455 |
| Q24319 | OST48_DROME Dolichyl-diphosphooligosaccharide--protein glycosyltransferase 48 kDa subunit OS=Drosophila melanogaster OX=7227 GN=Ost48 PE=2 SV=2 | 90 | 24 | 17 | 11 | 6.491853096 | 4.087462841 | 2.404390255 |
| Q960Y8 | Q960Y8_DROME Aluminum tubes, isoform G OS=Drosophila melanogaster OX=7227 GN=alt PE=1 SV=1 | 88 | 40 | 1 | 1 | 6.459431619 | 0 | 6.459431619 |
| Q76NQ0 | Q76NQ0_DROME Dolichyl-diphosphooligosaccharide--protein glycosyltransferase subunit 1 OS=Drosophila melanogaster OX=7227 GN=CG5364 PE=1 SV=1 | 87 | 26 | 11 | 8 | 6.442943496 | 3.459431619 | 2.983511877 |
| Q9VE75 | Q9VE75_DROME V-type proton ATPase subunit a OS=Drosophila melanogaster OX=7227 GN=Vha100-2 PE=1 SV=2 | 87 | 37 | 13 | 10 | 6.442943496 | 3.700439718 | 2.742503778 |
| Q7K3D4 | Q7K3D4_DROME Peptidylprolyl isomerase OS=Drosophila melanogaster OX=7227 GN=zda PE=1 SV=1 | 86 | 21 | 13 | 9 | 6.426264755 | 3.700439718 | 2.725825037 |
| E1JHT6 | E1JHT6_DROME Reticulon-like protein OS=Drosophila melanogaster OX=7227 GN=Rtnl1 PE=1 SV=1 | 85 | 24 | 16 | 10 | 6.409390936 | 4 | 2.409390936 |
| K2C1_HUMAN | (Common contaminant protein) | 79 | 28 | 33 | 19 | 6.303780748 | 5.044394119 | 1.259386629 |
| P29413 | CALR_DROME Calreticulin OS=Drosophila melanogaster OX=7227 GN=Calr PE=1 SV=2 | 79 | 18 | 31 | 12 | 6.303780748 | 4.95419631 | 1.349584438 |
| Q9Y125 | Q9Y125_DROME BcDNA.GH08312 OS=Drosophila melanogaster OX=7227 GN=Sap-r PE=1 SV=1 | 78 | 28 | 17 | 11 | 6.285402219 | 4.087462841 | 2.197939378 |
| Q9VKZ7 | Q9VKZ7_DROME Nicalin OS=Drosophila melanogaster OX=7227 GN=CG4972-RA PE=1 SV=2 | 75 | 23 | 7 | 6 | 6.22881869 | 2.807354922 | 3.421463768 |
| Q9VV75 | Q9VV75_DROME AT02348p OS=Drosophila melanogaster OX=7227 GN=UQCR-C2 PE=1 SV=1 | 75 | 28 | 34 | 14 | 6.22881869 | 5.087462841 | 1.141355849 |
| Q27331 | VATA2_DROME V-type proton ATPase catalytic subunit A isoform 2 OS=Drosophila melanogaster OX=7227 GN=Vha68-2 PE=1 SV=2 | 72 | 29 | 59 | 24 | 6.169925001 | 5.882643049 | 0.287281952 |
| A0A023GQA5 | A0A023GQA5_DROME Lethal (2) 37Cc, isoform D OS=Drosophila melanogaster OX=7227 GN=l(2)37Cc PE=1 SV=1 | 72 | 20 | 31 | 14 | 6.169925001 | 4.95419631 | 1.215728691 |
| O46067 | O46067_DROME EG:25E8.1 protein OS=Drosophila melanogaster OX=7227 GN=CT9894 PE=1 SV=1 | 72 | 34 | 26 | 13 | 6.169925001 | 4.700439718 | 1.469485283 |
| A4V449 | A4V449_DROME NADH dehydrogenase (Ubiquinone) 75 kDa subunit, isoform B OS=Drosophila melanogaster OX=7227 GN=ND-75 PE=1 SV=1 | 69 | 30 | 26 | 16 | 6.108524457 | 4.700439718 | 1.408084739 |
| Q9V3K9 | Q9V3K9_DROME Carnitine palmitoyltransferase I OS=Drosophila melanogaster OX=7227 GN=whd PE=1 SV=1 | 69 | 27 | 7 | 6 | 6.108524457 | 2.807354922 | 3.301169535 |
| Q9VLN6 | Q9VLN6_DROME FI20146p1 OS=Drosophila melanogaster OX=7227 GN=Dme_CG7627 PE=1 SV=2 | 68 | 36 | 7 | 7 | 6.087462841 | 2.807354922 | 3.280107919 |
| K1C9_HUMAN | (Common contaminant protein) | 67 | 25 | 27 | 16 | 6.06608919 | 4.754887502 | 1.311201688 |
| M9PCF1 | M9PCF1_DROME Dual oxidase, isoform C OS=Drosophila melanogaster OX=7227 GN=Duox PE=4 SV=1 | 67 | 40 | 4 | 4 | 6.06608919 | 2 | 4.06608919 |
| Q7JWF1 | Q7JWF1_DROME Electron transfer flavoprotein-ubiquinone oxidoreductase, isoform A OS=Drosophila melanogaster OX=7227 GN=Etf-QO PE=1 SV=1 | 67 | 25 | 17 | 10 | 6.06608919 | 4.087462841 | 1.978626349 |
| Q9VB46 | Q9VB46_DROME FI18644p1 OS=Drosophila melanogaster OX=7227 GN=Hmu PE=1 SV=1 | 67 | 19 | 23 | 8 | 6.06608919 | 4.523561956 | 1.542527234 |
| O01666 | ATPG_DROME ATP synthase subunit gamma, mitochondrial OS=Drosophila melanogaster OX=7227 GN=ATPsyngamma PE=2 SV=2 | 66 | 18 | 20 | 10 | 6.044394119 | 4.321928095 | 1.722466024 |
| Q9VN14 | CONT_DROME Contactin OS=Drosophila melanogaster OX=7227 GN=Cont PE=1 SV=2 | 66 | 34 | 1 | 1 | 6.044394119 | 0 | 6.044394119 |
| A1Z713 | VPS13_DROME Vacuolar protein sorting-associated protein 13 OS=Drosophila melanogaster OX=7227 GN=Vps13 PE=1 SV=2 | 65 | 52 | 5 | 5 | 6.022367813 | 2.321928095 | 3.700439718 |
| A0A0B4KHW3 | A0A0B4KHW3_DROME Aralar1, isoform F OS=Drosophila melanogaster OX=7227 GN=aralar1 PE=1 SV=1 | 65 | 24 | 11 | 10 | 6.022367813 | 3.459431619 | 2.562936194 |
| Q7KTC4 | Q7KTC4_DROME Multidrug-Resistance like protein 1, isoform H OS=Drosophila melanogaster OX=7227 GN=MRP PE=1 SV=1 | 65 | 37 | 4 | 4 | 6.022367813 | 2 | 4.022367813 |
| Q95RA9 | GILT1_DROME GILT-like protein 1 OS=Drosophila melanogaster OX=7227 GN=GILT1 PE=1 SV=1 | 64 | 16 | 29 | 5 | 6 | 4.857980995 | 1.142019005 |
| M9MRM4 | M9MRM4_DROME Sac1 phosphatase, isoform B OS=Drosophila melanogaster OX=7227 GN=Sac1 PE=4 SV=1 | 64 | 25 | 8 | 6 | 6 | 3 | 3 |
| Q9VS57 | Q9VS57_DROME RE14391p OS=Drosophila melanogaster OX=7227 GN=Sec63 PE=1 SV=1 | 64 | 25 | 8 | 4 | 6 | 3 | 3 |
| Q8STG9 | Q8STG9_DROME DSec61alpha OS=Drosophila melanogaster OX=7227 GN=Sec61alpha PE=1 SV=1 | 63 | 13 | 11 | 5 | 5.977279923 | 3.459431619 | 2.517848305 |
| Q9W227 | Q9W227_DROME Peptidyl-prolyl cis-trans isomerase OS=Drosophila melanogaster OX=7227 GN=CG2852 PE=1 SV=1 | 63 | 18 | 41 | 15 | 5.977279923 | 5.357552005 | 0.619727919 |
| Q9VG31 | Q9VG31_DROME Malic enzyme OS=Drosophila melanogaster OX=7227 GN=Men PE=1 SV=1 | 63 | 26 | 1 | 1 | 5.977279923 | 0 | 5.977279923 |
| Q9V3P0 | PRDX1_DROME Peroxiredoxin 1 OS=Drosophila melanogaster OX=7227 GN=Jafrac1 PE=1 SV=1 | 62 | 16 | 88 | 21 | 5.95419631 | 6.459431619 | -0.505235308 |
| Q24560 | TBB1_DROME Tubulin beta-1 chain OS=Drosophila melanogaster OX=7227 GN=betaTub56D PE=1 SV=2 | 61 | 22 | 219 | 42 | 5.930737338 | 7.77478706 | -1.844049722 |
| Q9VFF0 | Q9VFF0_DROME GH01077p OS=Drosophila melanogaster OX=7227 GN=UQCR-C1 PE=1 SV=2 | 61 | 19 | 29 | 15 | 5.930737338 | 4.857980995 | 1.072756342 |
| Q9VXM4 | Q9VXM4_DROME LD12946p OS=Drosophila melanogaster OX=7227 GN=MSBP PE=1 SV=1 | 61 | 17 | 3 | 3 | 5.930737338 | 1.584962501 | 4.345774837 |
| Q9VVK7 | Q9VVK7_DROME NUCB1 OS=Drosophila melanogaster OX=7227 GN=NUCB1 PE=1 SV=3 | 60 | 17 | 5 | 5 | 5.906890596 | 2.321928095 | 3.584962501 |
| Q59DQ0 | Q59DQ0_DROME Calcium-transporting ATPase OS=Drosophila melanogaster OX=7227 GN=PMCA PE=1 SV=3 | 60 | 25 | 1 | 1 | 5.906890596 | 0 | 5.906890596 |

| UniProt ID | Protein description | # PSMs MVB spectra | # of unique peptides MVB | # of PSMs lysate | # of unique peptides lysate | log2(PSM MVB) | log2(PSM lysate) | FC (log2 of MVB PSM-log2 of lysate PSM) |
|---|---|---|---|---|---|---|---|---|
| O62619 | KPYK_DROME Pyruvate kinase OS=Drosophila melanogaster OX=7227 GN=PyK PE=2 SV=2 | 59 | 23 | 133 | 34 | 5.882643049 | 7.055282436 | -1.172639386 |
| A8DYI6 | A8DYI6_DROME Prohibitin 2, isoform E OS=Drosophila melanogaster OX=7227 GN=Phb2 PE=1 SV=1 | 59 | 18 | 26 | 12 | 5.882643049 | 4.700439718 | 1.182203331 |
| Q7KB18 | Q7KB18_DROME Epoxide hydrolase OS=Drosophila melanogaster OX=7227 GN=Jheh2 PE=1 SV=1 | 58 | 21 | 2 | 2 | 5.857980995 | 1 | 4.857980995 |
| P25007 | PPIA_DROME Peptidyl-prolyl cis-trans isomerase OS=Drosophila melanogaster OX=7227 GN=Cyp1 PE=1 SV=2 | 57 | 17 | 108 | 26 | 5.832890014 | 6.754887502 | -0.921997488 |
| Q9VUC1 | Q9VUC1_DROME Hsc70Cb, isoform A OS=Drosophila melanogaster OX=7227 GN=Hsc70Cb PE=1 SV=1 | 57 | 31 | 105 | 37 | 5.832890014 | 6.714245518 | -0.881355504 |
| P15357 | RS27A_DROME Ubiquitin-40S ribosomal protein S27a OS=Drosophila melanogaster OX=7227 GN=RpS27A PE=1 SV=2 | 57 | 11 | 77 | 14 | 5.832890014 | 6.266786541 | -0.433896527 |
| Q9VC57 | ATLAS_DROME Atlastin OS=Drosophila melanogaster OX=7227 GN=atl PE=1 SV=1 | 57 | 20 | 3 | 2 | 5.832890014 | 1.584962501 | 4.247927513 |
| O18332 | O18332_DROME FI01544p OS=Drosophila melanogaster OX=7227 GN=Rab1 PE=1 SV=1 | 57 | 10 | 15 | 7 | 5.832890014 | 3.906890596 | 1.925999419 |
| Q9VIF2 | Q9VIF2_DROME GM01519p OS=Drosophila melanogaster OX=7227 GN=Dmel\CG9248 PE=1 SV=1 | 57 | 18 | 9 | 6 | 5.832890014 | 3.169925001 | 2.662965013 |
| Q9VKJ6 | Q9VKJ6_DROME Cation-transporting ATPase OS=Drosophila melanogaster OX=7227 GN=BcDNA:GH06032 PE=1 SV=1 | 57 | 28 | 1 | 1 | 5.832890014 | 0 | 5.832890014 |
| K1C10_HUMAN | (Common contaminant protein) | 56 | 20 | 25 | 12 | 5.807354922 | 4.64385619 | 1.163498732 |
| Q961D3 | Q961D3_DROME LD17001p OS=Drosophila melanogaster OX=7227 GN=rdog PE=1 SV=1 | 56 | 35 | 8 | 6 | 5.807354922 | 3 | 2.807354922 |
| Q9W351 | Q9W351_DROME LD42637p OS=Drosophila melanogaster OX=7227 GN=Aladin PE=1 SV=1 | 56 | 20 | 6 | 5 | 5.807354922 | 2.584962501 | 3.222392421 |
| Q5U124 | Q5U124_DROME LP11544p OS=Drosophila melanogaster OX=7227 GN=Mal-A5 PE=1 SV=1 | 55 | 22 | 11 | 7 | 5.781359714 | 3.459431619 | 2.321928095 |
| Q9VZF1 | Q9VZF1_DROME GH07444p OS=Drosophila melanogaster OX=7227 GN=Dmel\CG1309 PE=1 SV=1 | 55 | 17 | 2 | 2 | 5.781359714 | 1 | 4.781359714 |
| A0A0B4KEL0 | A0A0B4KEL0_DROME Aldehyde dehydrogenase OS=Drosophila melanogaster OX=7227 GN=Aldh-III PE=1 SV=1 | 55 | 22 | 1 | 1 | 5.781359714 | 0 | 5.781359714 |
| A0A0B4JD57 | A0A0B4JD57_DROME Zipper, isoform F OS=Drosophila melanogaster OX=7227 GN=zip PE=1 SV=1 | 54 | 37 | 93 | 48 | 5.754887502 | 6.539158811 | -0.784271309 |
| O18335 | O18335_DROME Drab11 OS=Drosophila melanogaster OX=7227 GN=Rab11 PE=1 SV=1 | 54 | 13 | 27 | 10 | 5.754887502 | 4.754887502 | 1 |
| Q9VF20 | Q9VF20_DROME Heavy metal tolerance factor 1 OS=Drosophila melanogaster OX=7227 GN=Hmt-1 PE=1 SV=1 | 54 | 25 | 2 | 2 | 5.754887502 | 1 | 4.754887502 |
| Q24251 | ATP5H_DROME ATP synthase subunit d, mitochondrial OS=Drosophila melanogaster OX=7227 GN=ATPsynD PE=2 SV=2 | 53 | 15 | 25 | 6 | 5.727920455 | 4.64385619 | 1.084064265 |
| Q24439 | ATPO_DROME ATP synthase subunit O, mitochondrial OS=Drosophila melanogaster OX=7227 GN=ATPsynO PE=2 SV=2 | 53 | 20 | 25 | 9 | 5.727920455 | 4.64385619 | 1.084064265 |
| Q8INH6 | Q8INH6_DROME FI18373p1 OS=Drosophila melanogaster OX=7227 GN=CG11688 PE=1 SV=1 | 53 | 23 | 42 | 20 | 5.727920455 | 5.392317423 | 0.335603032 |
| Q9VWV6 | Q9VWV6_DROME Transferrin OS=Drosophila melanogaster OX=7227 GN=Tsf1 PE=1 SV=1 | 53 | 26 | 49 | 19 | 5.727920455 | 5.614709844 | 0.11321061 |
| X2J4W8 | X2J4W8_DROME Kruppel homolog 2, isoform C OS=Drosophila melanogaster OX=7227 GN=Kr-h2 PE=1 SV=1 | 53 | 11 | 6 | 4 | 5.727920455 | 2.584962501 | 3.142957954 |
| Q9V397 | Q9V397_DROME BcDNA.GH12558 OS=Drosophila melanogaster OX=7227 GN=Mtpalpha PE=1 SV=1 | 53 | 29 | 1 | 1 | 5.727920455 | 0 | 5.727920455 |
| Q9VK04 | Q9VK04_DROME Phospholipid-transporting ATPase OS=Drosophila melanogaster OX=7227 GN=CG16823 PE=1 SV=2 | 53 | 27 | 1 | 1 | 5.727920455 | 0 | 5.727920455 |
| A0A0B4KHF0 | A0A0B4KHF0_DROME Ferritin OS=Drosophila melanogaster OX=7227 GN=Fer2LCH PE=1 SV=1 | 52 | 14 | 13 | 9 | 5.700439718 | 3.700439718 | 2 |
| Q7K1Q6 | Q7K1Q6_DROME Jabba, isoform B OS=Drosophila melanogaster OX=7227 GN=Jabba PE=1 SV=1 | 52 | 15 | 55 | 14 | 5.700439718 | 5.781359714 | -0.080919995 |
| Q9V3A8 | Q9V3A8_DROME Ergic53, isoform A OS=Drosophila melanogaster OX=7227 GN=ergic53 PE=1 SV=1 | 52 | 14 | 5 | 4 | 5.700439718 | 2.321928095 | 3.378511623 |
| Q9VRE0 | Q9VRE0_DROME Catalytic subunit 3A of the oligosaccharyltransferase complex, isoform A OS=Drosophila melanogaster OX=7227 GN=Stt3A PE=1 SV=1 | 52 | 21 | 8 | 6 | 5.700439718 | 3 | 2.700439718 |
| P06605 | TBA3_DROME Tubulin alpha-3 chain OS=Drosophila melanogaster OX=7227 GN=alphaTub84D PE=2 SV=1 | 52 | 20 | 1 | 1 | 5.700439718 | 0 | 5.700439718 |
| Q23997 | C5210_DROME Imaginal disk growth factor 6 OS=Drosophila melanogaster OX=7227 GN=Idgf6 PE=1 SV=2 | 51 | 19 | 14 | 11 | 5.672425342 | 3.807354922 | 1.86507042 |
| Q7K3B7 | Q7K3B7_DROME LD40177p OS=Drosophila melanogaster OX=7227 GN=CG11208 PE=1 SV=1 | 51 | 18 | 22 | 13 | 5.672425342 | 4.459431619 | 1.212993723 |
| Q9V4E0 | Q9V4E0_DROME LD47962p OS=Drosophila melanogaster OX=7227 GN=ND-49 PE=1 SV=2 | 51 | 18 | 12 | 10 | 5.672425342 | 3.584962501 | 2.087462841 |
| Q9VS02 | Q9VS02_DROME Uncharacterized protein OS=Drosophila melanogaster OX=7227 GN=CG9953 PE=1 SV=1 | 51 | 20 | 14 | 7 | 5.672425342 | 3.807354922 | 1.86507042 |
| Q7JR71 | SODE_DROME Extracellular superoxide dismutase [Cu-Zn] OS=Drosophila melanogaster OX=7227 GN=Sod3 PE=1 SV=1 | 51 | 8 | 1 | 1 | 5.672425342 | 0 | 5.672425342 |
| O02649 | CH60A_DROME Heat shock protein 60A OS=Drosophila melanogaster OX=7227 GN=Hsp60A PE=1 SV=3 | 50 | 22 | 112 | 36 | 5.64385619 | 6.807354922 | -1.163498732 |
| A0A0B4LHX6 | A0A0B4LHX6_DROME Nucleoside diphosphate kinase OS=Drosophila melanogaster OX=7227 GN=awd PE=1 SV=1 | 50 | 12 | 74 | 11 | 5.64385619 | 6.209453366 | -0.565597176 |
| Q94516 | AT5F1_DROME ATP synthase subunit b, mitochondrial OS=Drosophila melanogaster OX=7227 GN=ATPsynB PE=2 SV=2 | 50 | 15 | 13 | 6 | 5.64385619 | 3.700439718 | 1.943416472 |
| Q9VWT3 | Q9VWT3_DROME Gamma-glutamyl transpeptidase, isoform A OS=Drosophila melanogaster OX=7227 GN=Ggt-1 PE=2 SV=2 | 50 | 19 | 9 | 9 | 5.64385619 | 3.169925001 | 2.473931188 |
| Q9VXI3 | Q9VXI3_DROME Uncharacterized protein, isoform C OS=Drosophila melanogaster OX=7227 GN=CG9911 PE=1 SV=1 | 50 | 23 | 17 | 10 | 5.64385619 | 4.087462841 | 1.556393349 |
| A0A0B4LGJ9 | A0A0B4LGJ9_DROME CG3209, isoform D OS=Drosophila melanogaster OX=7227 GN=CG3209 PE=1 SV=1 | 50 | 16 | 1 | 1 | 5.64385619 | 0 | 5.64385619 |
| E1JHR5 | E1JHR5_DROME Enolase, isoform F OS=Drosophila melanogaster OX=7227 GN=Eno PE=1 SV=1 | 49 | 18 | 107 | 25 | 5.614709844 | 6.741466986 | -1.126757142 |
| Q9W0M4 | Q9W0M4_DROME LP10861p OS=Drosophila melanogaster OX=7227 GN=Dmel\CG13887 PE=1 SV=1 | 49 | 12 | 10 | 5 | 5.614709844 | 3.321928095 | 2.292781749 |
| B7Z0L1 | B7Z0L1_DROME Fasciclin 1, isoform E OS=Drosophila melanogaster OX=7227 GN=Fas1 PE=1 SV=1 | 49 | 24 | 1 | 1 | 5.614709844 | 0 | 5.614709844 |
| Q9VLT3 | Q9VLT3_DROME LD23292p OS=Drosophila melanogaster OX=7227 GN=Mcr PE=1 SV=1 | 48 | 32 | 1 | 1 | 5.584962501 | 0 | 5.584962501 |
| A1Z7S0 | A1Z7S0_DROME Receptor mediated endocytosis 8, isoform A OS=Drosophila melanogaster OX=7227 GN=Rme-8 PE=1 SV=1 | 47 | 33 | 23 | 18 | 5.554588852 | 4.523561956 | 1.031026896 |
| Q7K485 | Q7K485_DROME CathD OS=Drosophila melanogaster OX=7227 GN=cathD PE=1 SV=1 | 47 | 18 | 20 | 8 | 5.554588852 | 4.321928095 | 1.232660757 |
| Q9VC63 | Q9VC63_DROME FI20190p1 OS=Drosophila melanogaster OX=7227 GN=Dme_CG5789 PE=1 SV=2 | 47 | 28 | 4 | 4 | 5.554588852 | 2 | 3.554588852 |
| Q9VJ45 | Q9VJ45_DROME UDP-glucuronosyltransferase OS=Drosophila melanogaster OX=7227 GN=BEST:LD25345 PE=2 SV=2 | 47 | 16 | 6 | 5 | 5.554588852 | 2.584962501 | 2.969626351 |
| O16797 | RL3_DROME 60S ribosomal protein L3 OS=Drosophila melanogaster OX=7227 GN=RpL3 PE=1 SV=3 | 46 | 20 | 105 | 24 | 5.523561956 | 6.714245518 | -1.190683562 |
| P31409 | VATB_DROME V-type proton ATPase subunit B OS=Drosophila melanogaster OX=7227 GN=Vha55 PE=2 SV=1 | 46 | 20 | 32 | 14 | 5.523561956 | 5 | 0.523561956 |
| P83094 | STIM_DROME Stromal interaction molecule homolog OS=Drosophila melanogaster OX=7227 GN=Stim PE=1 SV=1 | 46 | 18 | 1 | 1 | 5.523561956 | 0 | 5.523561956 |
| Q27597 | NCPR_DROME NADPH--cytochrome P450 reductase OS=Drosophila melanogaster OX=7227 GN=Cpr PE=2 SV=2 | 46 | 21 | 6 | 6 | 5.523561956 | 2.584962501 | 2.938599455 |
| Q9VCG4 | NDC1_DROME Nucleoporin Ndc1 OS=Drosophila melanogaster OX=7227 GN=Ndc1 PE=1 SV=1 | 46 | 15 | 4 | 3 | 5.523561956 | 2 | 3.523561956 |
| Q9VZ04 | Q9VZ04_DROME Glucosidase 1, isoform B OS=Drosophila melanogaster OX=7227 GN=GCS1 PE=1 SV=4 | 46 | 20 | 4 | 2 | 5.523561956 | 2 | 3.523561956 |
| Q9W2X6 | Q9W2X6_DROME ATP synthase, delta subunit, isoform A OS=Drosophila melanogaster OX=7227 GN=ATPsyndelta PE=1 SV=1 | 46 | 11 | 8 | 5 | 5.523561956 | 3 | 2.523561956 |
| M9PEC8 | M9PEC8_DROME Dynein heavy chain 64C, isoform G OS=Drosophila melanogaster OX=7227 GN=Dhc64C PE=1 SV=1 | 45 | 38 | 106 | 80 | 5.491853096 | 6.727920455 | -1.236067358 |
| P54611 | VATE_DROME V-type proton ATPase subunit E OS=Drosophila melanogaster OX=7227 GN=Vha26 PE=2 SV=1 | 45 | 18 | 39 | 13 | 5.491853096 | 5.285402219 | 0.206450877 |
| Q23983 | SNAP_DROME Alpha-soluble NSF attachment protein OS=Drosophila melanogaster OX=7227 GN=alphaSnap PE=1 SV=1 | 45 | 15 | 11 | 7 | 5.491853096 | 3.459431619 | 2.032421478 |
| Q06559 | RS3_DROME 40S ribosomal protein S3 OS=Drosophila melanogaster OX=7227 GN=RpS3 PE=1 SV=1 | 44 | 20 | 108 | 32 | 5.459431619 | 6.754887502 | -1.295455884 |
| P92177 | 1433E_DROME 14-3-3 protein epsilon OS=Drosophila melanogaster OX=7227 GN=14-3-3epsilon PE=1 SV=2 | 44 | 14 | 89 | 22 | 5.459431619 | 6.475733431 | -1.016301812 |
| Q9VN93 | CPR1_DROME Putative cysteine proteinase CG12163 OS=Drosophila melanogaster OX=7227 GN=CG12163 PE=2 SV=2 | 44 | 20 | 15 | 10 | 5.459431619 | 3.906890596 | 1.552541023 |
| A0A0B4LGL5 | A0A0B4LGL5_DROME CG8399, isoform B OS=Drosophila melanogaster OX=7227 GN=CG8399 PE=4 SV=1 | 44 | 17 | 1 | 1 | 5.459431619 | 0 | 5.459431619 |
| Q9V3J1 | VATH_DROME V-type proton ATPase subunit H OS=Drosophila melanogaster OX=7227 GN=VhaSFD PE=2 SV=2 | 43 | 19 | 20 | 12 | 5.426264755 | 4.321928095 | 1.10433666 |
| Q9VYV3 | TXND5_DROME Thioredoxin domain-containing protein 5 homolog OS=Drosophila melanogaster OX=7227 GN=prtp PE=1 SV=2 | 43 | 20 | 15 | 8 | 5.426264755 | 3.906890596 | 1.519374159 |
| Q9VQR9 | Q9VQR9_DROME FI04424p OS=Drosophila melanogaster OX=7227 GN=CG17593-RA PE=1 SV=1 | 43 | 16 | 1 | 1 | 5.426264755 | 0 | 5.426264755 |
| Q9V3F2 | Q9V3F2_DROME LD36009p OS=Drosophila melanogaster OX=7227 GN=lace PE=1 SV=1 | 43 | 21 | 1 | 1 | 5.426264755 | 0 | 5.426264755 |
| Q9XZ10 | Q9XZ10_DROME V-type proton ATPase subunit a OS=Drosophila melanogaster OX=7227 GN=Vha100-1 PE=1 SV=1 | 43 | 19 | 1 | 1 | 5.426264755 | 0 | 5.426264755 |
| Q9VHL2 | Q9VHL2_DROME T-complex protein 1 subunit eta OS=Drosophila melanogaster OX=7227 GN=CCT7 PE=1 SV=2 | 42 | 25 | 118 | 37 | 5.392317423 | 6.882643049 | -1.490325627 |
| Q7KKI0 | Q7KKI0_DROME T-complex protein 1 subunit epsilon OS=Drosophila melanogaster OX=7227 GN=CCT5 PE=1 SV=1 | 42 | 22 | 86 | 27 | 5.392317423 | 6.426264755 | -1.033947332 |
| Q9U5L1 | SRPRA_DROME Signal recognition particle receptor subunit alpha homolog OS=Drosophila melanogaster OX=7227 GN=Gtp-bp PE=1 SV=2 | 42 | 19 | 12 | 9 | 5.392317423 | 3.584962501 | 1.807354922 |
| Q9VYY2 | SPCS2_DROME Signal peptidase complex subunit 2 OS=Drosophila melanogaster OX=7227 GN=Spase25 PE=2 SV=1 | 42 | 11 | 2 | 2 | 5.392317423 | 1 | 4.392317423 |
| A1Z6V5 | A1Z6V5_DROME FI01422p OS=Drosophila melanogaster OX=7227 GN=Spn43Ab PE=1 SV=1 | 42 | 17 | 4 | 3 | 5.392317423 | 2 | 3.392317423 |
| Q9VPJ0 | Q9VPJ0_DROME RE58433p OS=Drosophila melanogaster OX=7227 GN=CT31762 PE=1 SV=1 | 42 | 14 | 1 | 1 | 5.392317423 | 0 | 5.392317423 |
| Q02748 | IF4A_DROME Eukaryotic initiation factor 4A OS=Drosophila melanogaster OX=7227 GN=eIF4A PE=1 SV=3 | 41 | 22 | 91 | 30 | 5.357552005 | 6.50779464 | -1.150242636 |
| Q9VHR8 | DPP3_DROME Dipeptidyl peptidase 3 OS=Drosophila melanogaster OX=7227 GN=DppIII PE=2 SV=2 | 41 | 22 | 73 | 32 | 5.357552005 | 6.189824559 | -0.832272554 |
| Q24323 | SEM2A_DROME Semaphorin-2A OS=Drosophila melanogaster OX=7227 GN=Sema2a PE=1 SV=2 | 41 | 22 | 4 | 4 | 5.357552005 | 2 | 3.357552005 |
| Q7JRE4 | MAN1_DROME Inner nuclear membrane protein Man1 OS=Drosophila melanogaster OX=7227 GN=MAN1 PE=1 SV=1 | 41 | 20 | 1 | 1 | 5.357552005 | 0 | 5.357552005 |
| Q9W303 | IDGF4_DROME Chitinase-like protein Idgf4 OS=Drosophila melanogaster OX=7227 GN=Idgf4 PE=2 SV=1 | 41 | 14 | 19 | 10 | 5.357552005 | 4.247927513 | 1.109624491 |
| A1Z9M2 | A1Z9M2_DROME Receptor expression-enhancing protein OS=Drosophila melanogaster OX=7227 GN=ReepB PE=3 SV=1 | 41 | 8 | 4 | 3 | 5.357552005 | 2 | 3.357552005 |

| UniProt ID | Protein description | # PSMs MVB spectra | # of unique peptides MVB | # of PSMs lysate | # of unique peptides lysate | log2(PSM MVB) | log2(PSM lysate) | FC (log2 of MVB PSM-log2 of lysate PSM) |
|---|---|---|---|---|---|---|---|---|
| Q9VQX6 | Q9VQX6_DROME LP05054p OS=Drosophila melanogaster OX=7227 GN=Dmel\CG3702 PE=1 SV=2 | 41 | 16 | 3 | 3 | 5.357552005 | 1.584962501 | 3.772589504 |
| Q9VRL0 | Q9VRL0_DROME Cytochrome c1, isoform A OS=Drosophila melanogaster OX=7227 GN=Cyt-c1 PE=1 SV=1 | 41 | 12 | 15 | 7 | 5.357552005 | 3.906890596 | 1.450661409 |
| Q7KSU6 | Q7KSU6_DROME Uncharacterized protein, isoform D OS=Drosophila melanogaster OX=7227 GN=CG8036 PE=1 SV=1 | 40 | 20 | 77 | 24 | 5.321928095 | 6.266786541 | -0.944858446 |
| Q9V4N3 | CYB5_DROME Cytochrome b5 OS=Drosophila melanogaster OX=7227 GN=Cyt-b5 PE=2 SV=1 | 40 | 7 | 6 | 2 | 5.321928095 | 2.584962501 | 2.736965594 |
| Q9V7Y2 | SGPL_DROME Sphingosine-1-phosphate lyase OS=Drosophila melanogaster OX=7227 GN=Sply PE=1 SV=1 | 40 | 20 | 4 | 3 | 5.321928095 | 2 | 3.321928095 |
| X2JGG6 | X2JGG6_DROME Heat shock protein 26, isoform B OS=Drosophila melanogaster OX=7227 GN=Hsp26 PE=3 SV=1 | 39 | 14 | 99 | 21 | 5.285402219 | 6.62935662 | -1.343954401 |
| P48605 | TCPG_DROME T-complex protein 1 subunit gamma OS=Drosophila melanogaster OX=7227 GN=CCT3 PE=2 SV=2 | 39 | 22 | 93 | 29 | 5.285402219 | 6.539158811 | -1.253756592 |
| Q9V438 | PDIA6_DROME Protein disulfide-isomerase A6 homolog OS=Drosophila melanogaster OX=7227 GN=CaBP1 PE=1 SV=1 | 39 | 12 | 16 | 9 | 5.285402219 | 4 | 1.285402219 |
| M9PCI6 | M9PCI6_DROME Echinoid, isoform C OS=Drosophila melanogaster OX=7227 GN=ed PE=4 SV=1 | 39 | 24 | 4 | 4 | 5.285402219 | 2 | 3.285402219 |
| Q9VGT3 | Q9VGT3_DROME GM04645p OS=Drosophila melanogaster OX=7227 GN=Ugt86Da PE=1 SV=1 | 39 | 15 | 2 | 1 | 5.285402219 | 1 | 4.285402219 |
| Q9VHX9 | Q9VHX9_DROME CD98 heavy chain, isoform A OS=Drosophila melanogaster OX=7227 GN=CD98hc PE=1 SV=1 | 39 | 24 | 6 | 5 | 5.285402219 | 2.584962501 | 2.700439718 |
| Q9VZL3 | Q9VZL3_DROME Sc2 OS=Drosophila melanogaster OX=7227 GN=Sc2 PE=1 SV=1 | 39 | 14 | 10 | 6 | 5.285402219 | 3.321928095 | 1.963474124 |
| Q9U969 | SWS_DROME Neuropathy target esterase sws OS=Drosophila melanogaster OX=7227 GN=sws PE=1 SV=2 | 39 | 29 | 1 | 1 | 5.285402219 | 0 | 5.285402219 |
| Q9W1G7 | Q9W1G7_DROME LD21576p OS=Drosophila melanogaster OX=7227 GN=Nap1 PE=1 SV=1 | 38 | 13 | 79 | 24 | 5.247927513 | 6.303780748 | -1.055853235 |
| P20240 | OTE_DROME Otefin OS=Drosophila melanogaster OX=7227 GN=Ote PE=1 SV=2 | 38 | 18 | 40 | 14 | 5.247927513 | 5.321928095 | -0.074000581 |
| Q9VEX6 | ATD3A_DROME ATPase family AAA domain-containing protein 3A homolog OS=Drosophila melanogaster OX=7227 GN=bor PE=1 SV=2 | 38 | 19 | 9 | 6 | 5.247927513 | 3.169925001 | 2.078002512 |
| M9NHG9 | M9NHG9_DROME Proline dehydrogenase OS=Drosophila melanogaster OX=7227 GN=slgA PE=1 SV=1 | 38 | 26 | 4 | 4 | 5.247927513 | 2 | 3.247927513 |
| M9PBQ4 | M9PBQ4_DROME Apoptosis inducing factor, isoform C OS=Drosophila melanogaster OX=7227 GN=AIF PE=1 SV=1 | 38 | 16 | 4 | 3 | 5.247927513 | 2 | 3.247927513 |
| Q9VC10 | Q9VC10_DROME Phosphatidylinositol glycan anchor biosynthesis class S OS=Drosophila melanogaster OX=7227 GN=PIG-S PE=1 SV=2 | 38 | 20 | 3 | 3 | 5.247927513 | 1.584962501 | 3.662965013 |
| Q9VEQ2 | Q9VEQ2_DROME GH23451p OS=Drosophila melanogaster OX=7227 GN=Dmel\CG3678 PE=1 SV=2 | 38 | 14 | 6 | 4 | 5.247927513 | 2.584962501 | 2.662965013 |
| Q9VN86 | Q9VN86_DROME AT14148p OS=Drosophila melanogaster OX=7227 GN=CT40966 PE=1 SV=1 | 38 | 17 | 3 | 3 | 5.247927513 | 1.584962501 | 3.662965013 |
| Q9VF91 | Q9VF91_DROME FI21235p1 OS=Drosophila melanogaster OX=7227 GN=Dmel\CG6236 PE=1 SV=1 | 38 | 22 | 1 | 1 | 5.247927513 | 0 | 5.247927513 |
| Q24298 | CADE_DROME DE-cadherin OS=Drosophila melanogaster OX=7227 GN=shg PE=1 SV=2 | 38 | 24 | 1 | 1 | 5.247927513 | 0 | 5.247927513 |
| A0A0B4LGF5 | A0A0B4LGF5_DROME Optic atrophy 1 ortholog, isoform C OS=Drosophila melanogaster OX=7227 GN=Opa1 PE=1 SV=1 | 38 | 29 | 1 | 1 | 5.247927513 | 0 | 5.247927513 |
| P09180 | RL4_DROME 60S ribosomal protein L4 OS=Drosophila melanogaster OX=7227 GN=RpL4 PE=1 SV=2 | 37 | 18 | 74 | 23 | 5.209453366 | 6.209453366 | -1 |
| Q9W1Y1 | EMC89_DROME ER membrane protein complex subunit 8/9 homolog OS=Drosophila melanogaster OX=7227 GN=EMC8-9 PE=1 SV=1 | 37 | 14 | 6 | 5 | 5.209453366 | 2.584962501 | 2.624490865 |
| A0A0B4K7R1 | A0A0B4K7R1_DROME Signal peptidase complex subunit 3 OS=Drosophila melanogaster OX=7227 GN=Spase22-23 PE=3 SV=1 | 37 | 9 | 3 | 1 | 5.209453366 | 1.584962501 | 3.624490865 |
| Q9V3N6 | Q9V3N6_DROME Transmembrane 9 superfamily member OS=Drosophila melanogaster OX=7227 GN=TM9SF4 PE=1 SV=1 | 37 | 16 | 4 | 3 | 5.209453366 | 2 | 3.209453366 |
| Q9VCC2 | Q9VCC2_DROME RE50040p OS=Drosophila melanogaster OX=7227 GN=Dmel\CG5510 PE=1 SV=1 | 37 | 13 | 4 | 4 | 5.209453366 | 2 | 3.209453366 |
| Q9W3G0 | Q9W3G0_DROME Phosphatidylinositol glycan anchor biosynthesis class T OS=Drosophila melanogaster OX=7227 GN=PIG-T PE=2 SV=1 | 37 | 16 | 3 | 3 | 5.209453366 | 1.584962501 | 3.624490865 |
| Q9W4N8 | Q9W4N8_DROME LD30122p OS=Drosophila melanogaster OX=7227 GN=Vap33 PE=1 SV=1 | 37 | 17 | 1 | 1 | 5.209453366 | 0 | 5.209453366 |
| X2J9A4 | X2J9A4_DROME Translocation protein 1, isoform D OS=Drosophila melanogaster OX=7227 GN=Trp1 PE=1 SV=1 | 37 | 7 | 1 | 1 | 5.209453366 | 0 | 5.209453366 |
| Q9VVA4 | Q9VVA4_DROME GH26789p OS=Drosophila melanogaster OX=7227 GN=Dmel\CG9674 PE=1 SV=2 | 36 | 28 | 89 | 47 | 5.169925001 | 6.475733431 | -1.30580843 |
| P29742 | CLH_DROME Clathrin heavy chain OS=Drosophila melanogaster OX=7227 GN=Chc PE=1 SV=1 | 36 | 28 | 85 | 46 | 5.169925001 | 6.409390936 | -1.239465935 |
| Q01604 | PGK_DROME Phosphoglycerate kinase OS=Drosophila melanogaster OX=7227 GN=Pgk PE=2 SV=2 | 36 | 20 | 76 | 24 | 5.169925001 | 6.247927513 | -1.078002512 |
| P08928 | LAM0_DROME Lamin Dm0 OS=Drosophila melanogaster OX=7227 GN=Lam PE=1 SV=4 | 36 | 20 | 32 | 19 | 5.169925001 | 5 | 0.169925001 |
| P91928 | MIC60_DROME MICOS complex subunit Mic60 OS=Drosophila melanogaster OX=7227 GN=Mitofilin PE=1 SV=4 | 36 | 23 | 18 | 12 | 5.169925001 | 4.169925001 | 1 |
| Q8MV48 | GALT7_DROME N-acetylgalactosaminyltransferase 7 OS=Drosophila melanogaster OX=7227 GN=GalNAc-T2 PE=2 SV=2 | 36 | 18 | 2 | 2 | 5.169925001 | 1 | 4.169925001 |
| Q9VTX8 | ODR4_DROME Protein odr-4 homolog OS=Drosophila melanogaster OX=7227 GN=CG10616 PE=3 SV=2 | 36 | 11 | 2 | 2 | 5.169925001 | 1 | 4.169925001 |
| O18333 | O18333_DROME GH01619p OS=Drosophila melanogaster OX=7227 GN=Rab2 PE=1 SV=1 | 36 | 17 | 11 | 8 | 5.169925001 | 3.459431619 | 1.710493383 |
| Q7KSQ0 | Q7KSQ0_DROME LD46175p OS=Drosophila melanogaster OX=7227 GN=sea PE=1 SV=1 | 36 | 17 | 4 | 3 | 5.169925001 | 2 | 3.169925001 |
| Q9VIH3 | Q9VIH3_DROME Microsomal triacylglycerol transfer protein OS=Drosophila melanogaster OX=7227 GN=Mtp PE=1 SV=2 | 36 | 22 | 9 | 8 | 5.169925001 | 3.169925001 | 2 |
| Q9VX36 | Q9VX36_DROME NADH dehydrogenase (Ubiquinone) 24 kDa subunit, isoform A OS=Drosophila melanogaster OX=7227 GN=ND-24 PE=1 SV=1 | 36 | 11 | 6 | 5 | 5.169925001 | 2.584962501 | 2.584962501 |
| Q7KUB1 | Q7KUB1_DROME Isocitrate dehydrogenase [NADP] OS=Drosophila melanogaster OX=7227 GN=Idh PE=1 SV=1 | 35 | 18 | 66 | 26 | 5.129283017 | 6.044394119 | -0.915111102 |
| M9PF61 | M9PF61_DROME Uncharacterized protein, isoform C OS=Drosophila melanogaster OX=7227 GN=CG6084 PE=1 SV=1 | 35 | 16 | 60 | 22 | 5.129283017 | 5.906890596 | -0.777607579 |
| A8JNP2 | A8JNP2_DROME Arginine kinase, isoform E OS=Drosophila melanogaster OX=7227 GN=Argk PE=1 SV=1 | 35 | 16 | 53 | 22 | 5.129283017 | 5.727920455 | -0.598637438 |
| P26308 | GBB1_DROME Guanine nucleotide-binding protein subunit beta-1 OS=Drosophila melanogaster OX=7227 GN=Gbeta13F PE=1 SV=1 | 35 | 9 | 6 | 3 | 5.129283017 | 2.584962501 | 2.544320516 |
| Q8T4E1 | GPI8_DROME Putative GPI-anchor transamidase OS=Drosophila melanogaster OX=7227 GN=PIG-K PE=2 SV=1 | 35 | 20 | 3 | 1 | 5.129283017 | 1.584962501 | 3.544320516 |
| Q7KLX3 | Q7KLX3_DROME GEO05407p1 OS=Drosophila melanogaster OX=7227 GN=Tapdelta PE=1 SV=1 | 35 | 6 | 6 | 3 | 5.129283017 | 2.584962501 | 2.544320516 |
| Q9VD29 | Q9VD29_DROME RE74312p OS=Drosophila melanogaster OX=7227 GN=Sar1 PE=1 SV=1 | 35 | 9 | 8 | 4 | 5.129283017 | 3 | 2.129283017 |
| Q9W464 | Q9W464_DROME Glycosylphosphatidylinositol anchor attachment 1 OS=Drosophila melanogaster OX=7227 GN=GAA1 PE=1 SV=1 | 35 | 15 | 6 | 4 | 5.129283017 | 2.584962501 | 2.544320516 |
| Q9V769 | C6A22_DROME Cytochrome P450 6a22 OS=Drosophila melanogaster OX=7227 GN=Cyp6a22 PE=2 SV=1 | 35 | 18 | 1 | 1 | 5.129283017 | 0 | 5.129283017 |
| Q8IQW2 | Q8IQW2_DROME Cytochrome c oxidase subunit OS=Drosophila melanogaster OX=7227 GN=COX6B PE=1 SV=1 | 35 | 8 | 1 | 1 | 5.129283017 | 0 | 5.129283017 |
| Q8T0L3 | Q8T0L3_DROME GH24511p OS=Drosophila melanogaster OX=7227 GN=Uba1 PE=1 SV=1 | 34 | 19 | 95 | 37 | 5.087462841 | 6.569855608 | -1.482392767 |
| P48148 | RHO1_DROME Ras-like GTP-binding protein Rho1 OS=Drosophila melanogaster OX=7227 GN=Rho1 PE=1 SV=1 | 34 | 10 | 13 | 6 | 5.087462841 | 3.700439718 | 1.387023123 |
| Q0E9F9 | Q0E9F9_DROME CG2915, isoform A OS=Drosophila melanogaster OX=7227 GN=CG2915-RB PE=1 SV=1 | 34 | 14 | 17 | 9 | 5.087462841 | 4.087462841 | 1 |
| Q9VY49 | Q9VY49_DROME LD24340p OS=Drosophila melanogaster OX=7227 GN=Dmel\CG11674 PE=2 SV=2 | 34 | 18 | 2 | 2 | 5.087462841 | 1 | 4.087462841 |
| Q9VZU4 | Q9VZU4_DROME LD25561p OS=Drosophila melanogaster OX=7227 GN=ND-30 PE=1 SV=1 | 34 | 10 | 10 | 5 | 5.087462841 | 3.321928095 | 1.765534746 |
| Q9W3X7 | Q9W3X7_DROME NADH dehydrogenase [ubiquinone] 1 beta subcomplex subunit 8, mitochondrial OS=Drosophila melanogaster OX=7227 GN=ND-ASHI PE=1 SV=1 | 34 | 8 | 2 | 2 | 5.087462841 | 1 | 4.087462841 |
| Q9VG81 | Q9VG81_DROME RH49330p OS=Drosophila melanogaster OX=7227 GN=Dmel\CG5167 PE=1 SV=1 | 34 | 15 | 1 | 1 | 5.087462841 | 0 | 5.087462841 |
| Q9VS47 | Q9VS47_DROME LD39967p OS=Drosophila melanogaster OX=7227 GN=Dmel\CG8602 PE=2 SV=1 | 34 | 7 | 1 | 1 | 5.087462841 | 0 | 5.087462841 |
| Q9VKC7 | Q9VKC7_DROME FI04402p OS=Drosophila melanogaster OX=7227 GN=CG6766-RA PE=1 SV=1 | 34 | 16 | 1 | 1 | 5.087462841 | 0 | 5.087462841 |
| P07764 | ALF_DROME Fructose-bisphosphate aldolase OS=Drosophila melanogaster OX=7227 GN=Ald PE=1 SV=5 | 33 | 17 | 62 | 21 | 5.044394119 | 5.95419631 | -0.909802191 |
| Q9VXY7 | LSD2_DROME Lipid storage droplets surface-binding protein 2 OS=Drosophila melanogaster OX=7227 GN=Lsd-2 PE=1 SV=1 | 33 | 16 | 48 | 18 | 5.044394119 | 5.584962501 | -0.540568381 |
| P32748 | PYRD_DROME Dihydroorotate dehydrogenase (quinone), mitochondrial OS=Drosophila melanogaster OX=7227 GN=Dhod PE=2 SV=2 | 33 | 17 | 9 | 8 | 5.044394119 | 3.169925001 | 1.874469118 |
| Q9V7N5 | VATC_DROME V-type proton ATPase subunit C OS=Drosophila melanogaster OX=7227 GN=Vha44 PE=2 SV=5 | 33 | 18 | 31 | 15 | 5.044394119 | 4.95419631 | 0.090197809 |
| A1Z803 | A1Z803_DROME CG1665 OS=Drosophila melanogaster OX=7227 GN=Marc PE=1 SV=1 | 33 | 13 | 2 | 2 | 5.044394119 | 1 | 4.044394119 |
| Q9VCM8 | Q9VCM8_DROME Uncharacterized protein, isoform A OS=Drosophila melanogaster OX=7227 GN=CG4408 PE=1 SV=2 | 33 | 13 | 12 | 7 | 5.044394119 | 3.584962501 | 1.459431619 |
| Q9VIQ8 | Q9VIQ8_DROME Cytochrome c oxidase subunit 4, isoform A OS=Drosophila melanogaster OX=7227 GN=COX4 PE=1 SV=1 | 33 | 9 | 11 | 5 | 5.044394119 | 3.459431619 | 1.584962501 |
| Q9W3U0 | Q9W3U0_DROME Acyl-coenzyme A oxidase OS=Drosophila melanogaster OX=7227 GN=Dmel\CG4586 PE=1 SV=1 | 33 | 21 | 17 | 12 | 5.044394119 | 4.087462841 | 0.956931278 |
| Q9W4K0 | Q9W4K0_DROME CHOp24, isoform A OS=Drosophila melanogaster OX=7227 GN=CHOp24 PE=1 SV=1 | 33 | 12 | 5 | 4 | 5.044394119 | 2.321928095 | 2.722466024 |
| Q9W5R5 | Q9W5R5_DROME LD47843p OS=Drosophila melanogaster OX=7227 GN=Slmap PE=1 SV=2 | 33 | 19 | 4 | 4 | 5.044394119 | 2 | 3.044394119 |
| Q9V415 | Q9V415_DROME HMG-coA reductase degradation 3, isoform A OS=Drosophila melanogaster OX=7227 GN=Hrd3 PE=1 SV=1 | 33 | 16 | 1 | 1 | 5.044394119 | 0 | 5.044394119 |
| Q8MVS5 | GLT35_DROME Polypeptide N-acetylgalactosaminyltransferase 35A OS=Drosophila melanogaster OX=7227 GN=Pgant35A PE=1 SV=2 | 33 | 20 | 1 | 1 | 5.044394119 | 0 | 5.044394119 |
| Q7K1W4 | Q7K1W4_DROME Epoxide hydrolase OS=Drosophila melanogaster OX=7227 GN=Jheh3 PE=2 SV=1 | 33 | 14 | 1 | 1 | 5.044394119 | 0 | 5.044394119 |
| Q8T4G5 | Q8T4G5_DROME SD01613p OS=Drosophila melanogaster OX=7227 GN=AFG3L2 PE=1 SV=1 | 33 | 20 | 1 | 1 | 5.044394119 | 0 | 5.044394119 |
| P41093 | RL18A_DROME 60S ribosomal protein L18a OS=Drosophila melanogaster OX=7227 GN=RpL18A PE=1 SV=1 | 32 | 15 | 72 | 17 | 5 | 6.169925001 | -1.169925001 |
| Q9VIE8 | Q9VIE8_DROME Aconitate hydratase, mitochondrial OS=Drosophila melanogaster OX=7227 GN=Acon PE=1 SV=2 | 32 | 18 | 69 | 32 | 5 | 6.108524457 | -1.108524457 |
| P23654 | NRT_DROME Neurotactin OS=Drosophila melanogaster OX=7227 GN=Nrt PE=1 SV=3 | 32 | 16 | 2 | 2 | 5 | 1 | 4 |
| P91929 | NDUAA_DROME NADH dehydrogenase [ubiquinone] 1 alpha subcomplex subunit 10, mitochondrial OS=Drosophila melanogaster OX=7227 GN=ND-42 PE=2 SV=2 | 32 | 12 | 7 | 5 | 5 | 2.807354922 | 2.192645078 |
| Q8SXY6 | TMEDA_DROME Transmembrane emp24 domain-containing protein bai OS=Drosophila melanogaster OX=7227 GN=bai PE=2 SV=1 | 32 | 11 | 5 | 3 | 5 | 2.321928095 | 2.678071905 |

| UniProt ID | Protein description | # PSMs MVB spectra | # of unique peptides MVB | # of PSMs lysate | # of unique peptides lysate | log2(PSM MVB) | log2(PSM lysate) | FC (log2 of MVB PSM-log2 of lysate PSM) |
|---|---|---|---|---|---|---|---|---|
| Q8SY19 | Q8SY19_DROME Microsomal glutathione S-transferase-like, isoform A OS=Drosophila melanogaster OX=7227 GN=Mgstl PE=1 SV=1 | 32 | 5 | 4 | 3 | 5 | 2 | 3 |
| Q9VJ58 | Q9VJ58_DROME Fas-associated factor 2, isoform A OS=Drosophila melanogaster OX=7227 GN=Faf2 PE=1 SV=1 | 32 | 17 | 2 | 2 | 5 | 1 | 4 |
| Q9VMI3 | Q9VMI3_DROME NADH dehydrogenase [ubiquinone] flavoprotein 1, mitochondrial OS=Drosophila melanogaster OX=7227 GN=ND-51 PE=1 SV=1 | 32 | 16 | 10 | 6 | 5 | 3.321928095 | 1.678071905 |
| Q9VUQ7 | Q9VUQ7_DROME RE36966p OS=Drosophila melanogaster OX=7227 GN=Dmel\CG7739 PE=1 SV=1 | 32 | 11 | 2 | 1 | 5 | 1 | 4 |
| Q9VZS7 | Q9VZS7_DROME Uncharacterized protein, isoform B OS=Drosophila melanogaster OX=7227 GN=Dmel\CG14967 PE=1 SV=2 | 32 | 28 | 3 | 2 | 5 | 1.584962501 | 3.415037499 |
| Q9VI57 | Q9VI57_DROME Dephospho-CoA kinase OS=Drosophila melanogaster OX=7227 GN=Dpck PE=2 SV=1 | 32 | 10 | 1 | 1 | 5 | 0 | 5 |
| Q7K3J0 | Q7K3J0_DROME CG8258 OS=Drosophila melanogaster OX=7227 GN=CCT8 PE=1 SV=1 | 31 | 21 | 88 | 34 | 4.95419631 | 6.459431619 | -1.505235308 |
| P06606 | TBA4_DROME Tubulin alpha-4 chain OS=Drosophila melanogaster OX=7227 GN=alphaTub67C PE=3 SV=1 | 31 | 16 | 66 | 19 | 4.95419631 | 6.044394119 | -1.090197809 |
| Q8MT58 | Q8MT58_DROME RE11562p OS=Drosophila melanogaster OX=7227 GN=CG17337 PE=1 SV=1 | 31 | 19 | 65 | 23 | 4.95419631 | 6.022367813 | -1.068171503 |
| Q9VQG4 | COLT_DROME Congested-like trachea protein OS=Drosophila melanogaster OX=7227 GN=colt PE=2 SV=1 | 31 | 11 | 3 | 2 | 4.95419631 | 1.584962501 | 3.36923381 |
| M9PGL7 | M9PGL7_DROME Translocase of outer membrane 40, isoform C OS=Drosophila melanogaster OX=7227 GN=Tom40 PE=4 SV=1 | 31 | 10 | 8 | 6 | 4.95419631 | 3 | 1.95419631 |
| Q7JQR3 | Q7JQR3_DROME Sulfhydryl oxidase OS=Drosophila melanogaster OX=7227 GN=Qsox1 PE=1 SV=1 | 31 | 20 | 5 | 5 | 4.95419631 | 2.321928095 | 2.632268215 |
| Q7K2N0 | Q7K2N0_DROME CG8735, isoform A OS=Drosophila melanogaster OX=7227 GN=Lnpk PE=1 SV=1 | 31 | 14 | 3 | 2 | 4.95419631 | 1.584962501 | 3.36923381 |
| Q9VUZ0 | Q9VUZ0_DROME Translocon-associated protein subunit beta OS=Drosophila melanogaster OX=7227 GN=SsRbeta PE=1 SV=2 | 31 | 5 | 2 | 2 | 4.95419631 | 1 | 3.95419631 |
| Q9W3N1 | Q9W3N1_DROME RH59310p OS=Drosophila melanogaster OX=7227 GN=spidey PE=1 SV=1 | 31 | 11 | 4 | 4 | 4.95419631 | 2 | 2.95419631 |
| E1JJF9 | E1JJF9_DROME Neuroglian, isoform D OS=Drosophila melanogaster OX=7227 GN=Nrg PE=1 SV=1 | 31 | 25 | 1 | 1 | 4.95419631 | 0 | 4.95419631 |
| Q9VTZ0 | Q9VTZ0_DROME GH08269p OS=Drosophila melanogaster OX=7227 GN=tral PE=1 SV=3 | 31 | 14 | 1 | 1 | 4.95419631 | 0 | 4.95419631 |
| Q9VXQ5 | Q9VXQ5_DROME Chaperonin containing TCP1 subunit 6 OS=Drosophila melanogaster OX=7227 GN=CCT6 PE=1 SV=1 | 30 | 18 | 77 | 28 | 4.906890596 | 6.266786541 | -1.359895945 |
| P32100 | RL7_DROME 60S ribosomal protein L7 OS=Drosophila melanogaster OX=7227 GN=RpL7 PE=1 SV=2 | 30 | 16 | 73 | 18 | 4.906890596 | 6.189824559 | -1.282933963 |
| P52029 | G6PI_DROME Glucose-6-phosphate isomerase OS=Drosophila melanogaster OX=7227 GN=Pgi PE=2 SV=2 | 30 | 18 | 73 | 20 | 4.906890596 | 6.189824559 | -1.282933963 |
| Q9VW68 | Q9VW68_DROME Gamma-aminobutyric acid transaminase, isoform A OS=Drosophila melanogaster OX=7227 GN=Gabat PE=1 SV=1 | 30 | 19 | 63 | 21 | 4.906890596 | 5.977279923 | -1.070389328 |
| Q9GU68 | IF5A_DROME Eukaryotic translation initiation factor 5A OS=Drosophila melanogaster OX=7227 GN=eEF5 PE=2 SV=2 | 30 | 11 | 55 | 17 | 4.906890596 | 5.781359714 | -0.874469118 |
| P29613 | TPIS_DROME Triosephosphate isomerase OS=Drosophila melanogaster OX=7227 GN=Tpi PE=1 SV=3 | 30 | 14 | 49 | 14 | 4.906890596 | 5.614709844 | -0.707819249 |
| O76742 | O76742_DROME CG5915 protein OS=Drosophila melanogaster OX=7227 GN=Rab7 PE=1 SV=1 | 30 | 14 | 9 | 6 | 4.906890596 | 3.169925001 | 1.736965594 |
| Q9VGJ9 | Q9VGJ9_DROME Heme oxygenase OS=Drosophila melanogaster OX=7227 GN=Ho PE=1 SV=1 | 30 | 12 | 6 | 5 | 4.906890596 | 2.584962501 | 2.321928095 |
| Q9VH64 | Q9VH64_DROME LD29322p OS=Drosophila melanogaster OX=7227 GN=Dmel\CG8507 PE=1 SV=1 | 30 | 16 | 14 | 9 | 4.906890596 | 3.807354922 | 1.099535674 |
| Q9VIK1 | Q9VIK1_DROME Transmembrane 9 superfamily member OS=Drosophila melanogaster OX=7227 GN=TM9SF2 PE=1 SV=1 | 30 | 14 | 3 | 3 | 4.906890596 | 1.584962501 | 3.321928095 |
| Q9VPE2 | Q9VPE2_DROME NADH dehydrogenase (Ubiquinone) 39 kDa subunit, isoform A OS=Drosophila melanogaster OX=7227 GN=ND-39 PE=1 SV=1 | 30 | 13 | 10 | 5 | 4.906890596 | 3.321928095 | 1.584962501 |
| Q9VQR2 | Q9VQR2_DROME CG8844 protein OS=Drosophila melanogaster OX=7227 GN=ND-PDSW PE=1 SV=1 | 30 | 10 | 4 | 4 | 4.906890596 | 2 | 2.906890596 |
| Q9VTZ5 | Q9VTZ5_DROME LD22449p OS=Drosophila melanogaster OX=7227 GN=Tsf2 PE=1 SV=1 | 30 | 22 | 3 | 2 | 4.906890596 | 1.584962501 | 3.321928095 |
| Q8T8R1 | Y3800_DROME CCHC-type zinc finger protein CG3800 OS=Drosophila melanogaster OX=7227 GN=CG3800 PE=1 SV=1 | 29 | 3 | 69 | 7 | 4.857980995 | 6.108524457 | -1.250543462 |
| P20432 | GSTD1_DROME Glutathione S-transferase D1 OS=Drosophila melanogaster OX=7227 GN=GstD1 PE=1 SV=1 | 29 | 10 | 59 | 14 | 4.857980995 | 5.882643049 | -1.024662054 |
| Q9VLB7 | Q9VLB7_DROME Rab GDP dissociation inhibitor OS=Drosophila melanogaster OX=7227 GN=Gdi PE=1 SV=1 | 29 | 16 | 55 | 23 | 4.857980995 | 5.781359714 | -0.923378718 |
| P91938 | TRXR1_DROME Thioredoxin reductase 1, mitochondrial OS=Drosophila melanogaster OX=7227 GN=Trxr-1 PE=1 SV=2 | 29 | 15 | 45 | 21 | 4.857980995 | 5.491853096 | -0.633872101 |
| O97067 | PTH2_DROME Probable peptidyl-tRNA hydrolase 2 OS=Drosophila melanogaster OX=7227 GN=CG1307 PE=2 SV=2 | 29 | 7 | 2 | 2 | 4.857980995 | 1 | 3.857980995 |
| P16378 | GNAO_DROME G protein alpha o subunit OS=Drosophila melanogaster OX=7227 GN=Galphao PE=1 SV=1 | 29 | 14 | 1 | 1 | 4.857980995 | 0 | 4.857980995 |
| Q24372 | LACH_DROME Lachesin OS=Drosophila melanogaster OX=7227 GN=Lac PE=1 SV=2 | 29 | 11 | 3 | 3 | 4.857980995 | 1.584962501 | 3.273018494 |
| Q9I7K5 | TMEDE_DROME Transmembrane emp24 domain-containing protein eca OS=Drosophila melanogaster OX=7227 GN=eca PE=2 SV=2 | 29 | 11 | 4 | 4 | 4.857980995 | 2 | 2.857980995 |
| Q9V3Q4 | Q9V3Q4_DROME GH25379p OS=Drosophila melanogaster OX=7227 GN=Jafrac2 PE=1 SV=1 | 29 | 9 | 8 | 6 | 4.857980995 | 3 | 1.857980995 |
| Q9VSN9 | Q9VSN9_DROME GM04779p OS=Drosophila melanogaster OX=7227 GN=SrpRbeta PE=1 SV=2 | 29 | 14 | 4 | 4 | 4.857980995 | 2 | 2.857980995 |
| Q7KRU8 | Q7KRU8_DROME Ferritin OS=Drosophila melanogaster OX=7227 GN=Fer1HCH PE=1 SV=1 | 29 | 11 | 1 | 1 | 4.857980995 | 0 | 4.857980995 |
| Q9W0S7 | SND1_DROME Staphylococcal nuclease domain-containing protein 1 OS=Drosophila melanogaster OX=7227 GN=Tudor-SN PE=1 SV=1 | 28 | 19 | 133 | 45 | 4.807354922 | 7.055282436 | -2.247927513 |
| Q9NJH0 | EF1G_DROME Elongation factor 1-gamma OS=Drosophila melanogaster OX=7227 GN=Ef1gamma PE=2 SV=2 | 28 | 14 | 74 | 21 | 4.807354922 | 6.209453366 | -1.402098444 |
| P46223 | RL7A_DROME 60S ribosomal protein L7a OS=Drosophila melanogaster OX=7227 GN=RpL7A PE=1 SV=2 | 28 | 14 | 72 | 18 | 4.807354922 | 6.169925001 | -1.362570079 |
| P29746 | BNB_DROME Protein bangles and beads OS=Drosophila melanogaster OX=7227 GN=bnb PE=1 SV=1 | 28 | 11 | 10 | 7 | 4.807354922 | 3.321928095 | 1.485426827 |
| P91679 | PEPT1_DROME Peptide transporter family 1 OS=Drosophila melanogaster OX=7227 GN=yin PE=1 SV=2 | 28 | 17 | 6 | 5 | 4.807354922 | 2.584962501 | 2.222392421 |
| P91927 | LETM1_DROME Mitochondrial proton/calcium exchanger protein OS=Drosophila melanogaster OX=7227 GN=Letm1 PE=2 SV=2 | 28 | 18 | 3 | 3 | 4.807354922 | 1.584962501 | 3.222392421 |
| A0A140SRF8 | A0A140SRF8_DROME Uncharacterized protein, isoform C OS=Drosophila melanogaster OX=7227 GN=CG11857 PE=1 SV=1 | 28 | 8 | 5 | 3 | 4.807354922 | 2.321928095 | 2.485426827 |
| Q0KI00 | Q0KI00_DROME Aminopeptidase OS=Drosophila melanogaster OX=7227 GN=SP1029 PE=1 SV=1 | 28 | 20 | 7 | 6 | 4.807354922 | 2.807354922 | 2 |
| Q7K172 | Q7K172_DROME LD04933p OS=Drosophila melanogaster OX=7227 GN=ste24a PE=1 SV=1 | 28 | 14 | 1 | 1 | 4.807354922 | 0 | 4.807354922 |
| Q8IPB7 | Q8IPB7_DROME Alpha-mannosidase OS=Drosophila melanogaster OX=7227 GN=LManII PE=1 SV=1 | 28 | 16 | 13 | 9 | 4.807354922 | 3.700439718 | 1.106915204 |
| Q9VL16 | Q9VL16_DROME RE45833p OS=Drosophila melanogaster OX=7227 GN=BEST:LD31953 PE=2 SV=1 | 28 | 10 | 5 | 4 | 4.807354922 | 2.321928095 | 2.485426827 |
| Q9VND3 | Q9VND3_DROME LD24833p OS=Drosophila melanogaster OX=7227 GN=Pi4KIIalpha PE=1 SV=3 | 28 | 17 | 2 | 2 | 4.807354922 | 1 | 3.807354922 |
| Q9W2V9 | Q9W2V9_DROME LD40707p OS=Drosophila melanogaster OX=7227 GN=l(1)G0289 PE=1 SV=1 | 28 | 13 | 7 | 6 | 4.807354922 | 2.807354922 | 2 |
| X2J9Z1 | X2J9Z1_DROME Silver, isoform N OS=Drosophila melanogaster OX=7227 GN=svr PE=1 SV=1 | 28 | 21 | 5 | 3 | 4.807354922 | 2.321928095 | 2.485426827 |
| Q9VHA8 | Q9VHA8_DROME LD25575p OS=Drosophila melanogaster OX=7227 GN=P58IPK PE=1 SV=1 | 28 | 14 | 1 | 1 | 4.807354922 | 0 | 4.807354922 |
| Q9VYG3 | Q9VYG3_DROME LD41462p OS=Drosophila melanogaster OX=7227 GN=Dmel\CG4332 PE=1 SV=1 | 28 | 12 | 1 | 1 | 4.807354922 | 0 | 4.807354922 |
| Q24046 | ATPB1_DROME Sodium/potassium-transporting ATPase subunit beta-1 OS=Drosophila melanogaster OX=7227 GN=nrv1 PE=1 SV=2 | 28 | 10 | 1 | 1 | 4.807354922 | 0 | 4.807354922 |
| Q9VQ61 | Q9VQ61_DROME Aspartate aminotransferase OS=Drosophila melanogaster OX=7227 GN=Got2 PE=1 SV=1 | 27 | 15 | 76 | 20 | 4.754887502 | 6.247927513 | -1.493040011 |
| Q9VN44 | Q9VN44_DROME FI07923p OS=Drosophila melanogaster OX=7227 GN=Karybeta3 PE=1 SV=1 | 27 | 16 | 63 | 27 | 4.754887502 | 5.977279923 | -1.222392421 |
| Q9VTP4 | R10AB_DROME 60S ribosomal protein L10a-2 OS=Drosophila melanogaster OX=7227 GN=RpL10Ab PE=1 SV=2 | 27 | 10 | 54 | 15 | 4.754887502 | 5.754887502 | -1 |
| X2JF40 | X2JF40_DROME Adenosylhomocysteinase OS=Drosophila melanogaster OX=7227 GN=Ahcy PE=3 SV=1 | 27 | 14 | 53 | 16 | 4.754887502 | 5.727920455 | -0.973032952 |
| Q9VKI8 | Q9VKI8_DROME GH03305p OS=Drosophila melanogaster OX=7227 GN=Dmel\CG6287 PE=1 SV=1 | 27 | 14 | 44 | 16 | 4.754887502 | 5.459431619 | -0.704544116 |
| M9PHG2 | M9PHG2_DROME Moesin, isoform M OS=Drosophila melanogaster OX=7227 GN=Moe PE=1 SV=1 | 27 | 16 | 42 | 22 | 4.754887502 | 5.392317423 | -0.637429921 |
| P61209 | ARF1_DROME ADP-ribosylation factor 1 OS=Drosophila melanogaster OX=7227 GN=Arf79F PE=1 SV=2 | 27 | 13 | 14 | 8 | 4.754887502 | 3.807354922 | 0.94753258 |
| Q8SY53 | Q8SY53_DROME GH11935p OS=Drosophila melanogaster OX=7227 GN=Ostgamma PE=1 SV=1 | 27 | 12 | 5 | 3 | 4.754887502 | 2.321928095 | 2.432959407 |
| Q9VJZ4 | Q9VJZ4_DROME AT12494p OS=Drosophila melanogaster OX=7227 GN=ND-B22 PE=1 SV=1 | 27 | 13 | 3 | 3 | 4.754887502 | 1.584962501 | 3.169925001 |
| Q9VMH4 | Q9VMH4_DROME FI09914p OS=Drosophila melanogaster OX=7227 GN=CG9175-RA PE=2 SV=1 | 27 | 13 | 2 | 2 | 4.754887502 | 1 | 3.754887502 |
| Q9XZ53 | Q9XZ53_DROME Catalytic subunit 3B of the oligosaccharyltransferase complex OS=Drosophila melanogaster OX=7227 GN=Stt3B PE=1 SV=1 | 27 | 11 | 2 | 2 | 4.754887502 | 1 | 3.754887502 |
| A0A0B4KHG2 | A0A0B4KHG2_DROME Nicastrin, isoform E OS=Drosophila melanogaster OX=7227 GN=Nct PE=1 SV=1 | 27 | 12 | 1 | 1 | 4.754887502 | 0 | 4.754887502 |
| M9MRV1 | M9MRV1_DROME Tweek, isoform G OS=Drosophila melanogaster OX=7227 GN=tweek PE=4 SV=1 | 27 | 25 | 1 | 1 | 4.754887502 | 0 | 4.754887502 |
| Q5U117 | Q5U117_DROME Enigma OS=Drosophila melanogaster OX=7227 GN=Egm PE=1 SV=1 | 27 | 17 | 1 | 1 | 4.754887502 | 0 | 4.754887502 |
| Q9VNR2 | Q9VNR2_DROME Calcium-transporting ATPase OS=Drosophila melanogaster OX=7227 GN=SPoCk PE=3 SV=1 | 27 | 22 | 1 | 1 | 4.754887502 | 0 | 4.754887502 |
| Q9VN21 | Q9VN21_DROME LD30155p OS=Drosophila melanogaster OX=7227 GN=lost PE=1 SV=1 | 26 | 17 | 113 | 29 | 4.700439718 | 6.820178962 | -2.119739244 |
| P41042 | RS4_DROME 40S ribosomal protein S4 OS=Drosophila melanogaster OX=7227 GN=RpS4 PE=1 SV=2 | 26 | 12 | 82 | 19 | 4.700439718 | 6.357552005 | -1.657112286 |
| Q0E993 | Q0E993_DROME Valyl-tRNA synthetase, isoform A OS=Drosophila melanogaster OX=7227 GN=ValRS PE=1 SV=1 | 26 | 18 | 49 | 25 | 4.700439718 | 5.614709844 | -0.914270126 |
| P13395 | SPTCA_DROME Spectrin alpha chain OS=Drosophila melanogaster OX=7227 GN=alpha-Spec PE=1 SV=2 | 26 | 22 | 26 | 23 | 4.700439718 | 4.700439718 | 0 |
| Q9VAM6 | CISD2_DROME CDGSH iron-sulfur domain-containing protein 2 homolog OS=Drosophila melanogaster OX=7227 GN=Cisd2 PE=2 SV=1 | 26 | 6 | 5 | 3 | 4.700439718 | 2.321928095 | 2.378511623 |
| Q9VLU5 | WWOX_DROME WW domain-containing oxidoreductase OS=Drosophila melanogaster OX=7227 GN=Wwox PE=2 SV=1 | 26 | 14 | 1 | 1 | 4.700439718 | 0 | 4.700439718 |
| Q7JZN0 | Q7JZN0_DROME Protein transport protein Sec61 subunit beta OS=Drosophila melanogaster OX=7227 GN=Sec61beta PE=1 SV=1 | 26 | 4 | 7 | 2 | 4.700439718 | 2.807354922 | 1.893084796 |
| Q8SXR7 | Q8SXR7_DROME CG3394, isoform A OS=Drosophila melanogaster OX=7227 GN=CG3394 PE=1 SV=1 | 26 | 16 | 2 | 2 | 4.700439718 | 1 | 3.700439718 |

| UniProt ID | Protein description | # PSMs MVB spectra | # of unique peptides MVB | # of PSMs lysate | # of unique peptides lysate | log2(PSM MVB) | log2(PSM lysate) | FC (log2 of MVB PSM-log2 of lysate PSM) |
|---|---|---|---|---|---|---|---|---|
| Q9VDT5 | Q9VDT5_DROME Carboxypeptidase OS=Drosophila melanogaster OX=7227 GN=CG4572-RC PE=1 SV=1 | 26 | 12 | 5 | 3 | 4.700439718 | 2.321928095 | 2.378511623 |
| Q9V6K1 | TPP2_DROME Tripeptidyl-peptidase 2 OS=Drosophila melanogaster OX=7227 GN=TppII PE=1 SV=2 | 26 | 17 | 1 | 1 | 4.700439718 | 0 | 4.700439718 |
| A0A0B4LEY0 | A0A0B4LEY0_DROME Rab32, isoform H OS=Drosophila melanogaster OX=7227 GN=Rab32 PE=1 SV=1 | 26 | 13 | 1 | 1 | 4.700439718 | 0 | 4.700439718 |
| A1ZA83 | A1ZA83_DROME CG8397 OS=Drosophila melanogaster OX=7227 GN=CG8397 PE=1 SV=1 | 26 | 9 | 1 | 1 | 4.700439718 | 0 | 4.700439718 |
| Q8IN49 | Q8IN49_DROME MIP05539p OS=Drosophila melanogaster OX=7227 GN=CG31195-RA PE=1 SV=2 | 26 | 17 | 1 | 1 | 4.700439718 | 0 | 4.700439718 |
| Q8IPG9 | Q8IPG9_DROME Basigin, isoform A OS=Drosophila melanogaster OX=7227 GN=Bsg PE=1 SV=1 | 26 | 11 | 1 | 1 | 4.700439718 | 0 | 4.700439718 |
| A4V391 | A4V391_DROME Chaperonin containing TCP1 subunit 1, isoform B OS=Drosophila melanogaster OX=7227 GN=CCT1 PE=1 SV=1 | 25 | 15 | 79 | 29 | 4.64385619 | 6.303780748 | -1.659924558 |
| Q9VZU7 | Q9VZU7_DROME Ubiquitin carboxyl-terminal hydrolase OS=Drosophila melanogaster OX=7227 GN=Usp5 PE=1 SV=1 | 25 | 19 | 78 | 29 | 4.64385619 | 6.285402219 | -1.641546029 |
| Q9VS34 | RL18_DROME 60S ribosomal protein L18 OS=Drosophila melanogaster OX=7227 GN=RpL18 PE=1 SV=1 | 25 | 5 | 48 | 6 | 4.64385619 | 5.584962501 | -0.941106311 |
| P00334 | ADH_DROME Alcohol dehydrogenase OS=Drosophila melanogaster OX=7227 GN=Adh PE=1 SV=2 | 25 | 12 | 45 | 18 | 4.64385619 | 5.491853096 | -0.847996907 |
| P61851 | SODC_DROME Superoxide dismutase [Cu-Zn] OS=Drosophila melanogaster OX=7227 GN=Sod1 PE=1 SV=2 | 25 | 8 | 40 | 10 | 4.64385619 | 5.321928095 | -0.678071905 |
| Q9W0B8 | Q9W0B8_DROME Coatomer subunit alpha OS=Drosophila melanogaster OX=7227 GN=alphaCOP PE=1 SV=1 | 25 | 20 | 36 | 28 | 4.64385619 | 5.169925001 | -0.526068812 |
| P08953 | TOLL_DROME Protein toll OS=Drosophila melanogaster OX=7227 GN=Tl PE=1 SV=1 | 25 | 14 | 2 | 2 | 4.64385619 | 1 | 3.64385619 |
| Q9U1M2 | TM120_DROME Transmembrane protein 120 homolog OS=Drosophila melanogaster OX=7227 GN=CG32795 PE=1 SV=1 | 25 | 9 | 2 | 1 | 4.64385619 | 1 | 3.64385619 |
| A0A0S0X8K7 | A0A0S0X8K7_DROME Tiny tim 50, isoform B OS=Drosophila melanogaster OX=7227 GN=ttm50 PE=4 SV=1 | 25 | 12 | 5 | 4 | 4.64385619 | 2.321928095 | 2.321928095 |
| Q8IQH6 | Q8IQH6_DROME Uncharacterized protein, isoform A OS=Drosophila melanogaster OX=7227 GN=CT30701 PE=1 SV=1 | 25 | 9 | 3 | 2 | 4.64385619 | 1.584962501 | 3.058893689 |
| Q8IRD4 | Q8IRD4_DROME Glutathione peroxidase OS=Drosophila melanogaster OX=7227 GN=PHGPx PE=1 SV=1 | 25 | 9 | 15 | 7 | 4.64385619 | 3.906890596 | 0.736965594 |
| Q9V3I2 | Q9V3I2_DROME Drab5 OS=Drosophila melanogaster OX=7227 GN=Rab5 PE=1 SV=1 | 25 | 10 | 8 | 6 | 4.64385619 | 3 | 1.64385619 |
| Q9VKC8 | Q9VKC8_DROME FI03495p OS=Drosophila melanogaster OX=7227 GN=Tom70 PE=1 SV=2 | 25 | 17 | 5 | 4 | 4.64385619 | 2.321928095 | 2.321928095 |
| Q9W1I7 | Q9W1I7_DROME CG5554, isoform A OS=Drosophila melanogaster OX=7227 GN=CG5554 PE=1 SV=1 | 25 | 10 | 5 | 2 | 4.64385619 | 2.321928095 | 2.321928095 |
| Q9W495 | PGAP1_DROME GPI inositol-deacylase OS=Drosophila melanogaster OX=7227 GN=PGAP1 PE=2 SV=3 | 25 | 14 | 1 | 1 | 4.64385619 | 0 | 4.64385619 |
| A0A0B4KI10 | A0A0B4KI10_DROME 26S proteasome non-ATPase regulatory subunit 1 OS=Drosophila melanogaster OX=7227 GN=Rpn2 PE=3 SV=1 | 24 | 18 | 80 | 33 | 4.584962501 | 6.321928095 | -1.736965594 |
| Q9VZ23 | RAN_DROME GTP-binding nuclear protein Ran OS=Drosophila melanogaster OX=7227 GN=Ran PE=1 SV=1 | 24 | 14 | 44 | 15 | 4.584962501 | 5.459431619 | -0.874469118 |
| Q961R8 | Q961R8_DROME GH09263p OS=Drosophila melanogaster OX=7227 GN=GlyRS PE=1 SV=1 | 24 | 13 | 40 | 15 | 4.584962501 | 5.321928095 | -0.736965594 |
| X2JEJ0 | X2JEJ0_DROME Uncharacterized protein, isoform D OS=Drosophila melanogaster OX=7227 GN=CG1440 PE=1 SV=1 | 24 | 14 | 29 | 15 | 4.584962501 | 4.857980995 | -0.273018494 |
| A0A0B4KG41 | A0A0B4KG41_DROME ADP-ribosylation factor-like 6 interacting protein 1, isoform B OS=Drosophila melanogaster OX=7227 GN=Arl6IP1 PE=4 SV=1 | 24 | 6 | 3 | 2 | 4.584962501 | 1.584962501 | 3 |
| Q9VBQ5 | Q9VBQ5_DROME LD38433p OS=Drosophila melanogaster OX=7227 GN=Dmel\CG5112 PE=1 SV=1 | 24 | 11 | 5 | 4 | 4.584962501 | 2.321928095 | 2.263034406 |
| Q9VDI5 | Q9VDI5_DROME Uncharacterized protein, isoform A OS=Drosophila melanogaster OX=7227 GN=CG17271 PE=1 SV=2 | 24 | 9 | 3 | 2 | 4.584962501 | 1.584962501 | 3 |
| Q9VJZ1 | Q9VJZ1_DROME FI21342p1 OS=Drosophila melanogaster OX=7227 GN=CG9302-RA PE=1 SV=1 | 24 | 14 | 11 | 8 | 4.584962501 | 3.459431619 | 1.125530882 |
| Q9VMB9 | Q9VMB9_DROME Cytochrome c oxidase subunit 5B, isoform A OS=Drosophila melanogaster OX=7227 GN=COX5B PE=1 SV=1 | 24 | 10 | 10 | 5 | 4.584962501 | 3.321928095 | 1.263034406 |
| Q9VTU2 | Q9VTU2_DROME NADH dehydrogenase (Ubiquinone) SGDH subunit, isoform A OS=Drosophila melanogaster OX=7227 GN=ND-SGDH PE=1 SV=1 | 24 | 6 | 7 | 3 | 4.584962501 | 2.807354922 | 1.777607579 |
| Q9VZ58 | Q9VZ58_DROME Purple acid phosphatase OS=Drosophila melanogaster OX=7227 GN=Dmel\CG1637 PE=1 SV=1 | 24 | 9 | 2 | 2 | 4.584962501 | 1 | 3.584962501 |
| Q9W335 | Q9W335_DROME LD26546p OS=Drosophila melanogaster OX=7227 GN=l(1)G0320 PE=1 SV=1 | 24 | 7 | 1 | 1 | 4.584962501 | 0 | 4.584962501 |
| B7YZF6 | B7YZF6_DROME Phospholipid-transporting ATPase OS=Drosophila melanogaster OX=7227 GN=CG42321 PE=3 SV=1 | 24 | 15 | 1 | 1 | 4.584962501 | 0 | 4.584962501 |
| M9PF16 | M9PF16_DROME Spectrin beta chain OS=Drosophila melanogaster OX=7227 GN=beta-Spec PE=3 SV=1 | 24 | 20 | 1 | 1 | 4.584962501 | 0 | 4.584962501 |
| Q8IMM8 | Q8IMM8_DROME MIP21506p OS=Drosophila melanogaster OX=7227 GN=mino PE=1 SV=1 | 24 | 17 | 1 | 1 | 4.584962501 | 0 | 4.584962501 |
| Q9VY87 | Q9VY87_DROME Cathepsin B1, isoform A OS=Drosophila melanogaster OX=7227 GN=CtsB1 PE=1 SV=1 | 24 | 7 | 1 | 1 | 4.584962501 | 0 | 4.584962501 |
| Q8MLY8 | RS8_DROME 40S ribosomal protein S8 OS=Drosophila melanogaster OX=7227 GN=RpS8 PE=1 SV=1 | 23 | 10 | 50 | 16 | 4.523561956 | 5.64385619 | -1.120294234 |
| O18404 | HCD2_DROME 3-hydroxyacyl-CoA dehydrogenase type-2 OS=Drosophila melanogaster OX=7227 GN=scu PE=1 SV=1 | 23 | 10 | 49 | 16 | 4.523561956 | 5.614709844 | -1.091147888 |
| Q9VKW5 | Q9VKW5_DROME LP07359p OS=Drosophila melanogaster OX=7227 GN=Dmel\CG5355 PE=1 SV=2 | 23 | 15 | 43 | 26 | 4.523561956 | 5.426264755 | -0.902702799 |
| A0A0B4KEH0 | A0A0B4KEH0_DROME 14-3-3zeta, isoform K OS=Drosophila melanogaster OX=7227 GN=14-3-3zeta PE=1 SV=1 | 23 | 7 | 33 | 7 | 4.523561956 | 5.044394119 | -0.520832163 |
| O77277 | TORS_DROME Torsin-like protein OS=Drosophila melanogaster OX=7227 GN=Torsin PE=2 SV=2 | 23 | 11 | 3 | 3 | 4.523561956 | 1.584962501 | 2.938599455 |
| Q7YU24 | MARF_DROME Transmembrane GTPase Marf OS=Drosophila melanogaster OX=7227 GN=Marf PE=1 SV=1 | 23 | 16 | 5 | 4 | 4.523561956 | 2.321928095 | 2.201633861 |
| Q9VAL0 | SPCS1_DROME Signal peptidase complex subunit 1 OS=Drosophila melanogaster OX=7227 GN=Spase12 PE=1 SV=2 | 23 | 4 | 1 | 1 | 4.523561956 | 0 | 4.523561956 |
| E1JHY0 | E1JHY0_DROME NADH-cytochrome b5 reductase OS=Drosophila melanogaster OX=7227 GN=CG5946 PE=1 SV=1 | 23 | 12 | 4 | 3 | 4.523561956 | 2 | 2.523561956 |
| Q6NR46 | Q6NR46_DROME RE58623p OS=Drosophila melanogaster OX=7227 GN=Spt-I PE=1 SV=1 | 23 | 9 | 3 | 3 | 4.523561956 | 1.584962501 | 2.938599455 |
| Q7K0F7 | Q7K0F7_DROME Carbonyl reductase, isoform A OS=Drosophila melanogaster OX=7227 GN=CG11200 PE=1 SV=1 | 23 | 11 | 2 | 2 | 4.523561956 | 1 | 3.523561956 |
| Q7K127 | Q7K127_DROME Alpha-galactosidase OS=Drosophila melanogaster OX=7227 GN=CG7997 PE=1 SV=1 | 23 | 10 | 9 | 5 | 4.523561956 | 3.169925001 | 1.353636955 |
| Q8IPT1 | Q8IPT1_DROME Uncharacterized protein, isoform D OS=Drosophila melanogaster OX=7227 GN=CG32441 PE=1 SV=2 | 23 | 7 | 4 | 3 | 4.523561956 | 2 | 2.523561956 |
| Q9V3Y4 | Q9V3Y4_DROME LD43650p OS=Drosophila melanogaster OX=7227 GN=Mtch PE=1 SV=1 | 23 | 14 | 4 | 3 | 4.523561956 | 2 | 2.523561956 |
| Q9VBX8 | Q9VBX8_DROME IP15857p OS=Drosophila melanogaster OX=7227 GN=CG11935 PE=1 SV=2 | 23 | 9 | 2 | 2 | 4.523561956 | 1 | 3.523561956 |
| Q9W2M4 | Q9W2M4_DROME Uncharacterized protein OS=Drosophila melanogaster OX=7227 GN=CG10527 PE=1 SV=1 | 23 | 8 | 10 | 4 | 4.523561956 | 3.321928095 | 1.201633861 |
| P16620 | PTP69_DROME Tyrosine-protein phosphatase 69D OS=Drosophila melanogaster OX=7227 GN=Ptp69D PE=1 SV=2 | 23 | 18 | 1 | 1 | 4.523561956 | 0 | 4.523561956 |
| P54385 | DHE3_DROME Glutamate dehydrogenase, mitochondrial OS=Drosophila melanogaster OX=7227 GN=Gdh PE=1 SV=2 | 22 | 14 | 74 | 28 | 4.459431619 | 6.209453366 | -1.750021747 |
| Q7KMQ0 | Q7KMQ0_DROME 26S proteasome regulatory complex subunit p48B OS=Drosophila melanogaster OX=7227 GN=Rpt1 PE=1 SV=1 | 22 | 12 | 64 | 22 | 4.459431619 | 6 | -1.540568381 |
| Q8MSW0 | Q8MSW0_DROME Isoleucyl-tRNA synthetase, isoform A OS=Drosophila melanogaster OX=7227 GN=IleRS PE=1 SV=1 | 22 | 19 | 57 | 31 | 4.459431619 | 5.832890014 | -1.373458396 |
| P41094 | RS18_DROME 40S ribosomal protein S18 OS=Drosophila melanogaster OX=7227 GN=RpS18 PE=1 SV=1 | 22 | 14 | 49 | 16 | 4.459431619 | 5.614709844 | -1.155278225 |
| Q9VBN5 | Q9VBN5_DROME 60S ribosomal protein L27 OS=Drosophila melanogaster OX=7227 GN=RpL27 PE=1 SV=1 | 22 | 8 | 34 | 10 | 4.459431619 | 5.087462841 | -0.628031223 |
| Q8IPU3 | Q8IPU3_DROME Tenzing norgay, isoform C OS=Drosophila melanogaster OX=7227 GN=tzn PE=1 SV=1 | 22 | 10 | 31 | 11 | 4.459431619 | 4.95419631 | -0.494764692 |
| Q9VE79 | Q9VE79_DROME Uncharacterized protein OS=Drosophila melanogaster OX=7227 GN=CG14309 PE=1 SV=2 | 22 | 12 | 31 | 16 | 4.459431619 | 4.95419631 | -0.494764692 |
| Q94523 | SDHA_DROME Succinate dehydrogenase [ubiquinone] flavoprotein subunit, mitochondrial OS=Drosophila melanogaster OX=7227 GN=SdhA PE=2 SV=3 | 22 | 13 | 27 | 13 | 4.459431619 | 4.754887502 | -0.295455884 |
| Q27367 | CRQ_DROME Protein croquemort OS=Drosophila melanogaster OX=7227 GN=crq PE=1 SV=3 | 22 | 13 | 2 | 1 | 4.459431619 | 1 | 3.459431619 |
| Q9V463 | NU154_DROME Nuclear pore complex protein Nup154 OS=Drosophila melanogaster OX=7227 GN=Nup154 PE=1 SV=1 | 22 | 13 | 16 | 10 | 4.459431619 | 4 | 0.459431619 |
| Q9VIU7 | DPM1_DROME Probable dolichol-phosphate mannosyltransferase OS=Drosophila melanogaster OX=7227 GN=CG10166 PE=3 SV=1 | 22 | 9 | 4 | 3 | 4.459431619 | 2 | 2.459431619 |
| Q9W0G1 | PTP61_DROME Tyrosine-protein phosphatase non-receptor type 61F OS=Drosophila melanogaster OX=7227 GN=Ptp61F PE=1 SV=1 | 22 | 12 | 5 | 3 | 4.459431619 | 2.321928095 | 2.137503524 |
| Q7K2E1 | Q7K2E1_DROME CG8839, isoform A OS=Drosophila melanogaster OX=7227 GN=CG8839 PE=1 SV=1 | 22 | 12 | 2 | 2 | 4.459431619 | 1 | 3.459431619 |
| Q9VCC6 | Q9VCC6_DROME GM05240p OS=Drosophila melanogaster OX=7227 GN=Dmel\CG6178 PE=1 SV=1 | 22 | 11 | 16 | 11 | 4.459431619 | 4 | 0.459431619 |
| Q9VDY8 | Q9VDY8_DROME MIP08680p OS=Drosophila melanogaster OX=7227 GN=vib PE=1 SV=1 | 22 | 9 | 17 | 8 | 4.459431619 | 4.087462841 | 0.371968777 |
| Q9VEY5 | Q9VEY5_DROME MICOS complex subunit OS=Drosophila melanogaster OX=7227 GN=Dmel\CG5903 PE=1 SV=1 | 22 | 10 | 5 | 3 | 4.459431619 | 2.321928095 | 2.137503524 |
| Q9VGK3 | Q9VGK3_DROME Peptidylprolyl isomerase OS=Drosophila melanogaster OX=7227 GN=BcDNA:GM09283 PE=1 SV=1 | 22 | 8 | 13 | 5 | 4.459431619 | 3.700439718 | 0.7589919 |
| Q9VNA3 | Q9VNA3_DROME GH21273p OS=Drosophila melanogaster OX=7227 GN=BEST:LD29996 PE=1 SV=1 | 22 | 10 | 2 | 2 | 4.459431619 | 1 | 3.459431619 |
| Q9VTY2 | Q9VTY2_DROME Uncharacterized protein, isoform A OS=Drosophila melanogaster OX=7227 GN=AKR PE=1 SV=1 | 22 | 10 | 23 | 10 | 4.459431619 | 4.523561956 | -0.064130337 |
| Q9VQ39 | Q9VQ39_DROME IP05441p OS=Drosophila melanogaster OX=7227 GN=CG17711 PE=1 SV=1 | 22 | 15 | 1 | 1 | 4.459431619 | 0 | 4.459431619 |
| Q9V3D4 | IDGF2_DROME Chitinase-like protein Idgf2 OS=Drosophila melanogaster OX=7227 GN=Idgf2 PE=1 SV=1 | 22 | 11 | 1 | 1 | 4.459431619 | 0 | 4.459431619 |
| Q9V4A7 | PLXB_DROME Plexin-B OS=Drosophila melanogaster OX=7227 GN=PlexB PE=1 SV=2 | 22 | 18 | 1 | 1 | 4.459431619 | 0 | 4.459431619 |
| E1JGQ9 | E1JGQ9_DROME Syndecan OS=Drosophila melanogaster OX=7227 GN=Sdc PE=1 SV=2 | 22 | 6 | 1 | 1 | 4.459431619 | 0 | 4.459431619 |
| Q7YU01 | Q7YU01_DROME LD11117p OS=Drosophila melanogaster OX=7227 GN=LpR2 PE=2 SV=1 | 22 | 14 | 1 | 1 | 4.459431619 | 0 | 4.459431619 |
| Q9VGF7 | Q9VGF7_DROME Glutamate carrier 1, isoform A OS=Drosophila melanogaster OX=7227 GN=GC1 PE=1 SV=1 | 22 | 8 | 1 | 1 | 4.459431619 | 0 | 4.459431619 |
| A0A0B4KGU4 | A0A0B4KGU4_DROME Belle, isoform B OS=Drosophila melanogaster OX=7227 GN=bel PE=4 SV=1 | 21 | 14 | 88 | 30 | 4.392317423 | 6.459431619 | -2.067114196 |
| P55830 | RS3A_DROME 40S ribosomal protein S3a OS=Drosophila melanogaster OX=7227 GN=RpS3A PE=1 SV=4 | 21 | 13 | 82 | 26 | 4.392317423 | 6.357552005 | -1.965234582 |
| Q9W392 | Q9W392_DROME Chaperonin containing TCP1 subunit 2 OS=Drosophila melanogaster OX=7227 GN=CCT2 PE=1 SV=2 | 21 | 14 | 74 | 32 | 4.392317423 | 6.209453366 | -1.817135943 |

| UniProt ID | Protein description | # PSMs MVB spectra | # of unique peptides MVB | # of PSMs lysate | # of unique peptides lysate | log2(PSM MVB) | log2(PSM lysate) | FC (log2 of MVB PSM-log2 of lysate PSM) |
|---|---|---|---|---|---|---|---|---|
| Q9VPN5 | Q9VPN5_DROME LD03220p OS=Drosophila melanogaster OX=7227 GN=Stip1 PE=1 SV=1 | 21 | 15 | 71 | 27 | 4.392317423 | 6.14974712 | -1.757429697 |
| A0A0B4K7J2 | RBP2_DROME E3 SUMO-protein ligase RanBP2 OS=Drosophila melanogaster OX=7227 GN=Nup358 PE=1 SV=1 | 21 | 13 | 66 | 43 | 4.392317423 | 6.044394119 | -1.652076697 |
| P31009 | RS2_DROME 40S ribosomal protein S2 OS=Drosophila melanogaster OX=7227 GN=RpS2 PE=1 SV=2 | 21 | 9 | 54 | 16 | 4.392317423 | 5.754887502 | -1.362570079 |
| P46415 | ADHX_DROME Alcohol dehydrogenase class-3 OS=Drosophila melanogaster OX=7227 GN=Fdh PE=1 SV=3 | 21 | 12 | 52 | 15 | 4.392317423 | 5.700439718 | -1.308122295 |
| Q9VNE9 | RL13A_DROME 60S ribosomal protein L13a OS=Drosophila melanogaster OX=7227 GN=RpL13A PE=1 SV=1 | 21 | 10 | 40 | 11 | 4.392317423 | 5.321928095 | -0.929610672 |
| P45594 | CADF_DROME Cofilin/actin-depolymerizing factor homolog OS=Drosophila melanogaster OX=7227 GN=tsr PE=1 SV=1 | 21 | 10 | 35 | 15 | 4.392317423 | 5.129283017 | -0.736965594 |
| Q9VFV9 | Q9VFV9_DROME DnaJ-like-2, isoform A OS=Drosophila melanogaster OX=7227 GN=Droj2 PE=1 SV=1 | 21 | 10 | 31 | 12 | 4.392317423 | 4.95419631 | -0.561878888 |
| K22E_HUMAN | (Common contaminant protein) | 21 | 16 | 8 | 5 | 4.392317423 | 3 | 1.392317423 |
| Q9VFC2 | SP88E_DROME Serine protease inhibitor 88Ea OS=Drosophila melanogaster OX=7227 GN=Spn88Ea PE=2 SV=1 | 21 | 15 | 8 | 5 | 4.392317423 | 3 | 1.392317423 |
| Q9W4P5 | VA0D1_DROME V-type proton ATPase subunit d 1 OS=Drosophila melanogaster OX=7227 GN=VhaAC39-1 PE=2 SV=1 | 21 | 8 | 3 | 2 | 4.392317423 | 1.584962501 | 2.807354922 |
| M9PF96 | M9PF96_DROME Peroxisomal membrane protein 70 kDa, isoform C OS=Drosophila melanogaster OX=7227 GN=Pmp70 PE=1 SV=1 | 21 | 14 | 1 | 1 | 4.392317423 | 0 | 4.392317423 |
| Q59E30 | Q59E30_DROME Failed axon connections, isoform C OS=Drosophila melanogaster OX=7227 GN=fax PE=1 SV=1 | 21 | 8 | 6 | 4 | 4.392317423 | 2.584962501 | 1.807354922 |
| Q9V3W2 | Q9V3W2_DROME GM23292p OS=Drosophila melanogaster OX=7227 GN=ND-B17 PE=1 SV=1 | 21 | 8 | 3 | 3 | 4.392317423 | 1.584962501 | 2.807354922 |
| Q9W125 | Q9W125_DROME NADH dehydrogenase [ubiquinone] 1 alpha subcomplex subunit 8 OS=Drosophila melanogaster OX=7227 GN=ND-19 PE=1 SV=1 | 21 | 8 | 3 | 2 | 4.392317423 | 1.584962501 | 2.807354922 |
| Q9W2G8 | Q9W2G8_DROME Acyl-coenzyme A oxidase OS=Drosophila melanogaster OX=7227 GN=Acox57D-d PE=1 SV=1 | 21 | 12 | 7 | 6 | 4.392317423 | 2.807354922 | 1.584962501 |
| Q9VKR0 | Q9VKR0_DROME ER membrane protein complex subunit 3 OS=Drosophila melanogaster OX=7227 GN=EMC3 PE=1 SV=1 | 21 | 12 | 1 | 1 | 4.392317423 | 0 | 4.392317423 |
| Q9W404 | Q9W404_DROME GH10714p OS=Drosophila melanogaster OX=7227 GN=Dmel\CG3842 PE=2 SV=1 | 21 | 12 | 1 | 1 | 4.392317423 | 0 | 4.392317423 |
| X2JD82 | X2JD82_DROME Open rectifier K[+] channel 1, isoform D OS=Drosophila melanogaster OX=7227 GN=Ork1 PE=3 SV=1 | 21 | 16 | 1 | 1 | 4.392317423 | 0 | 4.392317423 |
| Q9VLQ1 | ALG5_DROME Dolichyl-phosphate beta-glucosyltransferase OS=Drosophila melanogaster OX=7227 GN=wol PE=1 SV=1 | 21 | 9 | 1 | 1 | 4.392317423 | 0 | 4.392317423 |
| Q9VRB3 | CP6V1_DROME Probable cytochrome P450 6v1 OS=Drosophila melanogaster OX=7227 GN=Cyp6v1 PE=2 SV=1 | 21 | 15 | 1 | 1 | 4.392317423 | 0 | 4.392317423 |
| Q9W376 | NEUFC_DROME Neufericin homolog OS=Drosophila melanogaster OX=7227 GN=CG12056 PE=2 SV=1 | 21 | 6 | 1 | 1 | 4.392317423 | 0 | 4.392317423 |
| Q9NFV7 | Q9NFV7_DROME GH01829p OS=Drosophila melanogaster OX=7227 GN=Tep2 PE=1 SV=1 | 21 | 16 | 1 | 1 | 4.392317423 | 0 | 4.392317423 |
| O18640 | GBLP_DROME Guanine nucleotide-binding protein subunit beta-like protein OS=Drosophila melanogaster OX=7227 GN=Rack1 PE=1 SV=2 | 20 | 11 | 162 | 31 | 4.321928095 | 7.339850003 | -3.017921908 |
| A4UZZ4 | A4UZZ4_DROME Alpha-1,4 glucan phosphorylase OS=Drosophila melanogaster OX=7227 GN=GlyP PE=1 SV=1 | 20 | 17 | 113 | 46 | 4.321928095 | 6.820178962 | -2.498250868 |
| Q9VQL7 | Q9VQL7_DROME Fatty acid synthase 1, isoform A OS=Drosophila melanogaster OX=7227 GN=FASN1 PE=1 SV=1 | 20 | 17 | 81 | 49 | 4.321928095 | 6.339850003 | -2.017921908 |
| Q9VU68 | WDR1_DROME Actin-interacting protein 1 OS=Drosophila melanogaster OX=7227 GN=flr PE=2 SV=1 | 20 | 19 | 75 | 32 | 4.321928095 | 6.22881869 | -1.906890596 |
| Q9VW54 | Q9VW54_DROME 26S proteasome non-ATPase regulatory subunit 2 OS=Drosophila melanogaster OX=7227 GN=Rpn1 PE=1 SV=2 | 20 | 14 | 70 | 26 | 4.321928095 | 6.129283017 | -1.807354922 |
| P50887 | RL22_DROME 60S ribosomal protein L22 OS=Drosophila melanogaster OX=7227 GN=RpL22 PE=1 SV=2 | 20 | 8 | 61 | 10 | 4.321928095 | 5.930737338 | -1.608809243 |
| Q9VVU2 | Q9VVU2_DROME GEO07453p1 OS=Drosophila melanogaster OX=7227 GN=RpL26 PE=1 SV=1 | 20 | 8 | 55 | 12 | 4.321928095 | 5.781359714 | -1.459431619 |
| O18413 | PRS8_DROME 26S proteasome regulatory subunit 8 OS=Drosophila melanogaster OX=7227 GN=Rpt6 PE=1 SV=2 | 20 | 13 | 47 | 20 | 4.321928095 | 5.554588852 | -1.232660757 |
| P29845 | HSP7E_DROME Heat shock 70 kDa protein cognate 5 OS=Drosophila melanogaster OX=7227 GN=Hsc70-5 PE=1 SV=2 | 20 | 16 | 44 | 21 | 4.321928095 | 5.459431619 | -1.137503524 |
| Q9V3I1 | Q9V3I1_DROME BcDNA.LD08534 OS=Drosophila melanogaster OX=7227 GN=dUTPase PE=1 SV=1 | 20 | 8 | 41 | 11 | 4.321928095 | 5.357552005 | -1.03562391 |
| Q9V3Z4 | Q9V3Z4_DROME GH11341p OS=Drosophila melanogaster OX=7227 GN=Rpn5 PE=1 SV=1 | 20 | 10 | 41 | 17 | 4.321928095 | 5.357552005 | -1.03562391 |
| P08645 | RAP1_DROME Ras-related protein Rap1 OS=Drosophila melanogaster OX=7227 GN=Rap1 PE=2 SV=2 | 20 | 7 | 4 | 3 | 4.321928095 | 2 | 2.321928095 |
| Q9W1X9 | OCAD1_DROME OCIA domain-containing protein 1 OS=Drosophila melanogaster OX=7227 GN=asrij PE=1 SV=1 | 20 | 11 | 3 | 2 | 4.321928095 | 1.584962501 | 2.736965594 |
| Q7JRH2 | Q7JRH2_DROME Receptor expression-enhancing protein OS=Drosophila melanogaster OX=7227 GN=ReepA PE=1 SV=1 | 20 | 8 | 4 | 2 | 4.321928095 | 2 | 2.321928095 |
| Q7KJ73 | Q7KJ73_DROME Tetraspanin OS=Drosophila melanogaster OX=7227 GN=Tsp42Ee PE=1 SV=1 | 20 | 5 | 1 | 1 | 4.321928095 | 0 | 4.321928095 |
| Q8IQD4 | Q8IQD4_DROME Anion exchange protein OS=Drosophila melanogaster OX=7227 GN=CG8177 PE=1 SV=1 | 20 | 13 | 2 | 2 | 4.321928095 | 1 | 3.321928095 |
| Q8SYJ2 | Q8SYJ2_DROME GEO09626p1 OS=Drosophila melanogaster OX=7227 GN=ND-MLRQ PE=1 SV=1 | 20 | 6 | 7 | 4 | 4.321928095 | 2.807354922 | 1.514573173 |
| Q8T0V2 | Q8T0V2_DROME GH02495p OS=Drosophila melanogaster OX=7227 GN=MESK2 PE=1 SV=1 | 20 | 7 | 1 | 1 | 4.321928095 | 0 | 4.321928095 |
| Q9VJ43 | Q9VJ43_DROME LD10783p OS=Drosophila melanogaster OX=7227 GN=ScpX PE=1 SV=1 | 20 | 12 | 7 | 4 | 4.321928095 | 2.807354922 | 1.514573173 |
| Q9VJ75 | Q9VJ75_DROME GH08340p OS=Drosophila melanogaster OX=7227 GN=Dmel\CG10413 PE=1 SV=1 | 20 | 14 | 5 | 5 | 4.321928095 | 2.321928095 | 2 |
| Q9VVJ7 | Q9VVJ7_DROME CG7484 protein OS=Drosophila melanogaster OX=7227 GN=BcDNA:SD16138 PE=1 SV=2 | 20 | 8 | 4 | 2 | 4.321928095 | 2 | 2.321928095 |
| Q9VW40 | Q9VW40_DROME LD31235p OS=Drosophila melanogaster OX=7227 GN=TORIP PE=1 SV=2 | 20 | 7 | 2 | 2 | 4.321928095 | 1 | 3.321928095 |
| Q9VFP0 | Q9VFP0_DROME RH07106p OS=Drosophila melanogaster OX=7227 GN=Dmel\CG3061 PE=1 SV=1 | 20 | 10 | 1 | 1 | 4.321928095 | 0 | 4.321928095 |
| Q8IMZ9 | WFS1_DROME Wolframin OS=Drosophila melanogaster OX=7227 GN=wfs1 PE=1 SV=2 | 20 | 14 | 1 | 1 | 4.321928095 | 0 | 4.321928095 |
| Q9V427 | INX2_DROME Innexin inx2 OS=Drosophila melanogaster OX=7227 GN=Inx2 PE=1 SV=1 | 20 | 9 | 1 | 1 | 4.321928095 | 0 | 4.321928095 |
| Q9VCW1 | CP6D4_DROME Probable cytochrome P450 6d4 OS=Drosophila melanogaster OX=7227 GN=Cyp6d4 PE=2 SV=1 | 20 | 12 | 1 | 1 | 4.321928095 | 0 | 4.321928095 |
| Q9VLP9 | PORED_DROME Polyprenol reductase OS=Drosophila melanogaster OX=7227 GN=CG7840 PE=2 SV=1 | 20 | 9 | 1 | 1 | 4.321928095 | 0 | 4.321928095 |
| A0A0B4K7G9 | A0A0B4K7G9_DROME Nucleosome remodeling factor-38kD, isoform C OS=Drosophila melanogaster OX=7227 GN=Nurf-38 PE=1 SV=1 | 20 | 13 | 1 | 1 | 4.321928095 | 0 | 4.321928095 |
| A8DZ03 | A8DZ03_DROME Kuzbanian, isoform G OS=Drosophila melanogaster OX=7227 GN=kuz PE=1 SV=2 | 20 | 11 | 1 | 1 | 4.321928095 | 0 | 4.321928095 |
| O18338 | O18338_DROME LD44762p OS=Drosophila melanogaster OX=7227 GN=Rab8 PE=1 SV=1 | 20 | 9 | 1 | 1 | 4.321928095 | 0 | 4.321928095 |
| Q9VF33 | Q9VF33_DROME Alpha-mannosidase OS=Drosophila melanogaster OX=7227 GN=alpha-Man-IIb PE=2 SV=2 | 20 | 15 | 1 | 1 | 4.321928095 | 0 | 4.321928095 |
| Q9VXY9 | Q9VXY9_DROME Opossum, isoform A OS=Drosophila melanogaster OX=7227 GN=opm PE=1 SV=2 | 20 | 6 | 1 | 1 | 4.321928095 | 0 | 4.321928095 |
| X2J9B0 | X2J9B0_DROME Niemann-Pick type C-1a, isoform E OS=Drosophila melanogaster OX=7227 GN=Npc1a PE=1 SV=1 | 20 | 14 | 1 | 1 | 4.321928095 | 0 | 4.321928095 |
| X2JE30 | X2JE30_DROME Integrin beta OS=Drosophila melanogaster OX=7227 GN=mys PE=3 SV=1 | 20 | 12 | 1 | 1 | 4.321928095 | 0 | 4.321928095 |
| P52304 | POLO_DROME Serine/threonine-protein kinase polo OS=Drosophila melanogaster OX=7227 GN=polo PE=1 SV=2 | 19 | 11 | 48 | 18 | 4.247927513 | 5.584962501 | -1.337034987 |
| P12881 | PSA1_DROME Proteasome subunit alpha type-1 OS=Drosophila melanogaster OX=7227 GN=Prosalpha6 PE=1 SV=1 | 19 | 10 | 46 | 14 | 4.247927513 | 5.523561956 | -1.275634443 |
| P55935 | RS9_DROME 40S ribosomal protein S9 OS=Drosophila melanogaster OX=7227 GN=RpS9 PE=1 SV=2 | 19 | 12 | 46 | 17 | 4.247927513 | 5.523561956 | -1.275634443 |
| Q9W0A8 | Q9W0A8_DROME FI01658p OS=Drosophila melanogaster OX=7227 GN=RpL23A PE=1 SV=1 | 19 | 7 | 46 | 9 | 4.247927513 | 5.523561956 | -1.275634443 |
| Q9VEB1 | Q9VEB1_DROME IP09655p OS=Drosophila melanogaster OX=7227 GN=Mdh2 PE=1 SV=1 | 19 | 11 | 40 | 15 | 4.247927513 | 5.321928095 | -1.074000581 |
| P41126 | RL13_DROME 60S ribosomal protein L13 OS=Drosophila melanogaster OX=7227 GN=RpL13 PE=1 SV=1 | 19 | 8 | 36 | 14 | 4.247927513 | 5.169925001 | -0.921997488 |
| Q9VQL1 | Q9VQL1_DROME LP20978p OS=Drosophila melanogaster OX=7227 GN=SerRS PE=1 SV=1 | 19 | 11 | 31 | 17 | 4.247927513 | 4.95419631 | -0.706268797 |
| O18334 | RAB6_DROME Ras-related protein Rab6 OS=Drosophila melanogaster OX=7227 GN=Rab6 PE=1 SV=1 | 19 | 11 | 4 | 4 | 4.247927513 | 2 | 2.247927513 |
| P00408 | COX2_DROME Cytochrome c oxidase subunit 2 OS=Drosophila melanogaster OX=7227 GN=mt:CoII PE=3 SV=1 | 19 | 8 | 2 | 2 | 4.247927513 | 1 | 3.247927513 |
| P40792 | RAC1_DROME Ras-related protein Rac1 OS=Drosophila melanogaster OX=7227 GN=Rac1 PE=1 SV=2 | 19 | 7 | 11 | 6 | 4.247927513 | 3.459431619 | 0.788495895 |
| Q9VUR0 | UFSP2_DROME Probable Ufm1-specific protease 2 OS=Drosophila melanogaster OX=7227 GN=UFSP2 PE=2 SV=1 | 19 | 12 | 1 | 1 | 4.247927513 | 0 | 4.247927513 |
| A0A0B4KFY4 | A0A0B4KFY4_DROME CG3499, isoform D OS=Drosophila melanogaster OX=7227 GN=YME1L PE=1 SV=1 | 19 | 11 | 2 | 1 | 4.247927513 | 1 | 3.247927513 |
| A0A0B4LFF6 | A0A0B4LFF6_DROME CG8180, isoform B OS=Drosophila melanogaster OX=7227 GN=CG8180 PE=4 SV=1 | 19 | 10 | 2 | 2 | 4.247927513 | 1 | 3.247927513 |
| Q9VF27 | Q9VF27_DROME GM02062p OS=Drosophila melanogaster OX=7227 GN=ND-23 PE=1 SV=1 | 19 | 9 | 4 | 4 | 4.247927513 | 2 | 2.247927513 |
| Q9VN13 | Q9VN13_DROME Sideroflexin OS=Drosophila melanogaster OX=7227 GN=Sfxn1-3 PE=1 SV=1 | 19 | 10 | 1 | 1 | 4.247927513 | 0 | 4.247927513 |
| Q9VRN1 | Q9VRN1_DROME Transmembrane 9 superfamily member OS=Drosophila melanogaster OX=7227 GN=TM9SF3 PE=1 SV=2 | 19 | 8 | 1 | 1 | 4.247927513 | 0 | 4.247927513 |
| Q9W029 | Q9W029_DROME FI16876p1 OS=Drosophila melanogaster OX=7227 GN=yellow-g PE=1 SV=1 | 19 | 10 | 1 | 1 | 4.247927513 | 0 | 4.247927513 |
| Q960W6 | FACR3_DROME Putative fatty acyl-CoA reductase CG8306 OS=Drosophila melanogaster OX=7227 GN=CG8306 PE=2 SV=1 | 19 | 14 | 1 | 1 | 4.247927513 | 0 | 4.247927513 |
| O77434 | O77434_DROME EG:34F3.8 protein OS=Drosophila melanogaster OX=7227 GN=Sec22 PE=1 SV=2 | 19 | 7 | 1 | 1 | 4.247927513 | 0 | 4.247927513 |
| Q8INJ4 | Q8INJ4_DROME LD21727p OS=Drosophila melanogaster OX=7227 GN=3 PE=1 SV=2 | 19 | 14 | 1 | 1 | 4.247927513 | 0 | 4.247927513 |
| Q8SWX6 | Q8SWX6_DROME GH08893p OS=Drosophila melanogaster OX=7227 GN=Dmel\CG11790 PE=1 SV=1 | 19 | 9 | 1 | 1 | 4.247927513 | 0 | 4.247927513 |
| Q9V9W3 | Q9V9W3_DROME RE08669p OS=Drosophila melanogaster OX=7227 GN=RpL6 PE=1 SV=1 | 19 | 7 | 1 | 1 | 4.247927513 | 0 | 4.247927513 |
| Q9VT68 | Q9VT68_DROME Phosphatidylinositol-4,5-bisphosphate 4-phosphatase OS=Drosophila melanogaster OX=7227 GN=CG6707 PE=1 SV=2 | 19 | 11 | 1 | 1 | 4.247927513 | 0 | 4.247927513 |
| A8DZ29 | A8DZ29_DROME Eukaryotic translation initiation factor 4G1, isoform B OS=Drosophila melanogaster OX=7227 GN=eIF4G1 PE=1 SV=1 | 18 | 15 | 95 | 32 | 4.169925001 | 6.569855608 | -2.399930607 |
| Q9VRP2 | Q9VRP2_DROME Uncharacterized protein, isoform A OS=Drosophila melanogaster OX=7227 GN=BEST:CK00496 PE=1 SV=1 | 18 | 10 | 73 | 16 | 4.169925001 | 6.189824559 | -2.019899557 |

| UniProt ID | Protein description | # PSMs MVB spectra | # of unique peptides MVB | # of PSMs lysate | # of unique peptides lysate | log2(PSM MVB) | log2(PSM lysate) | FC (log2 of MVB PSM-log2 of lysate PSM) |
|---|---|---|---|---|---|---|---|---|
| Q9VK69 | Q9VK69_DROME T-complex protein 1 subunit delta OS=Drosophila melanogaster OX=7227 GN=CCT4 PE=1 SV=1 | 18 | 9 | 70 | 23 | 4.169925001 | 6.129283017 | -1.959358016 |
| Q8IP94 | Q8IP94_DROME Threonyl-tRNA synthetase, isoform B OS=Drosophila melanogaster OX=7227 GN=ThrRS PE=1 SV=1 | 18 | 14 | 65 | 27 | 4.169925001 | 6.022367813 | -1.852442812 |
| P52295 | IMA_DROME Importin subunit alpha OS=Drosophila melanogaster OX=7227 GN=Pen PE=1 SV=2 | 18 | 9 | 63 | 21 | 4.169925001 | 5.977279923 | -1.807354922 |
| M9NDC3 | M9NDC3_DROME Female sterile (2) ketel, isoform B OS=Drosophila melanogaster OX=7227 GN=Fs(2)Ket PE=4 SV=1 | 18 | 11 | 56 | 23 | 4.169925001 | 5.807354922 | -1.637429921 |
| P05389 | RLA2_DROME 60S acidic ribosomal protein P2 OS=Drosophila melanogaster OX=7227 GN=RpLP2 PE=1 SV=1 | 18 | 6 | 50 | 8 | 4.169925001 | 5.64385619 | -1.473931188 |
| Q9VSA3 | ACADM_DROME Probable medium-chain specific acyl-CoA dehydrogenase, mitochondrial OS=Drosophila melanogaster OX=7227 GN=CG12262 PE=2 SV=1 | 18 | 11 | 44 | 16 | 4.169925001 | 5.459431619 | -1.289506617 |
| Q0E9B6 | RS11_DROME 40S ribosomal protein S11 OS=Drosophila melanogaster OX=7227 GN=RpS11 PE=1 SV=1 | 18 | 5 | 43 | 8 | 4.169925001 | 5.426264755 | -1.256339753 |
| P55841 | RL14_DROME 60S ribosomal protein L14 OS=Drosophila melanogaster OX=7227 GN=RpL14 PE=1 SV=1 | 18 | 8 | 42 | 12 | 4.169925001 | 5.392317423 | -1.222392421 |
| Q8INP8 | Q8INP8_DROME Uncharacterized protein, isoform B OS=Drosophila melanogaster OX=7227 GN=CG11980 PE=1 SV=1 | 18 | 12 | 42 | 17 | 4.169925001 | 5.392317423 | -1.222392421 |
| Q7JR58 | Q7JR58_DROME CG6543, isoform A OS=Drosophila melanogaster OX=7227 GN=Echs1 PE=1 SV=1 | 18 | 10 | 35 | 12 | 4.169925001 | 5.129283017 | -0.959358016 |
| Q9VWH4 | IDH3A_DROME Probable isocitrate dehydrogenase [NAD] subunit alpha, mitochondrial OS=Drosophila melanogaster OX=7227 GN=l(1)G0156 PE=2 SV=1 | 18 | 13 | 34 | 18 | 4.169925001 | 5.087462841 | -0.91753784 |
| Q9VGZ3 | Q9VGZ3_DROME Iron regulatory protein 1B OS=Drosophila melanogaster OX=7227 GN=Irp-1B PE=1 SV=1 | 18 | 14 | 32 | 22 | 4.169925001 | 5 | -0.830074999 |
| O18405 | SURF4_DROME Surfeit locus protein 4 homolog OS=Drosophila melanogaster OX=7227 GN=Surf4 PE=2 SV=1 | 18 | 8 | 2 | 1 | 4.169925001 | 1 | 3.169925001 |
| Q94514 | COX5A_DROME Cytochrome c oxidase subunit 5A, mitochondrial OS=Drosophila melanogaster OX=7227 GN=COX5A PE=2 SV=2 | 18 | 7 | 6 | 5 | 4.169925001 | 2.584962501 | 1.584962501 |
| Q9VLM5 | DAD1_DROME Dolichyl-diphosphooligosaccharide--protein glycosyltransferase subunit DAD1 OS=Drosophila melanogaster OX=7227 GN=l(2)k12914 PE=3 SV=1 | 18 | 6 | 5 | 3 | 4.169925001 | 2.321928095 | 1.847996907 |
| Q9VS60 | SMSR1_DROME Sphingomyelin synthase-related 1 OS=Drosophila melanogaster OX=7227 GN=SMSr PE=1 SV=2 | 18 | 7 | 1 | 1 | 4.169925001 | 0 | 4.169925001 |
| Q9VXN2 | SUN_DROME Protein stunted OS=Drosophila melanogaster OX=7227 GN=sun PE=1 SV=1 | 18 | 5 | 2 | 2 | 4.169925001 | 1 | 3.169925001 |
| A8E6L6 | A8E6L6_DROME FI01906p OS=Drosophila melanogaster OX=7227 GN=O-fut1 PE=1 SV=1 | 18 | 8 | 5 | 4 | 4.169925001 | 2.321928095 | 1.847996907 |
| E1JJL4 | E1JJL4_DROME Furin 2, isoform I OS=Drosophila melanogaster OX=7227 GN=Fur2 PE=3 SV=1 | 18 | 13 | 1 | 1 | 4.169925001 | 0 | 4.169925001 |
| Q7JVK8 | Q7JVK8_DROME CG5323 OS=Drosophila melanogaster OX=7227 GN=CG5323 PE=1 SV=1 | 18 | 5 | 3 | 2 | 4.169925001 | 1.584962501 | 2.584962501 |
| Q9VL10 | Q9VL10_DROME AT10584p OS=Drosophila melanogaster OX=7227 GN=Mulk PE=1 SV=2 | 18 | 12 | 2 | 2 | 4.169925001 | 1 | 3.169925001 |
| Q9VXI6 | Q9VXI6_DROME Cytochrome b-c1 complex subunit 7 OS=Drosophila melanogaster OX=7227 GN=UQCR-14 PE=1 SV=1 | 18 | 9 | 3 | 2 | 4.169925001 | 1.584962501 | 2.584962501 |
| Q9W3C4 | Q9W3C4_DROME Beta-hexosaminidase OS=Drosophila melanogaster OX=7227 GN=Hexo2 PE=1 SV=1 | 18 | 12 | 2 | 2 | 4.169925001 | 1 | 3.169925001 |
| Q9VLK0 | Q9VLK0_DROME LD37974p OS=Drosophila melanogaster OX=7227 GN=PIG-U PE=2 SV=1 | 18 | 9 | 1 | 1 | 4.169925001 | 0 | 4.169925001 |
| Q9VMD5 | Q9VMD5_DROME Farjavit, isoform A OS=Drosophila melanogaster OX=7227 GN=frj PE=2 SV=1 | 18 | 7 | 1 | 1 | 4.169925001 | 0 | 4.169925001 |
| Q9VMU0 | Q9VMU0_DROME NADH dehydrogenase [ubiquinone] iron-sulfur protein 6, mitochondrial OS=Drosophila melanogaster OX=7227 GN=ND-13A PE=1 SV=1 | 18 | 7 | 1 | 1 | 4.169925001 | 0 | 4.169925001 |
| Q9VV49 | Q9VV49_DROME RE72803p OS=Drosophila melanogaster OX=7227 GN=AGPAT4 PE=2 SV=1 | 18 | 7 | 1 | 1 | 4.169925001 | 0 | 4.169925001 |
| Q9V3A4 | CSUP_DROME Protein catecholamines up OS=Drosophila melanogaster OX=7227 GN=Catsup PE=2 SV=1 | 18 | 1 | 1 | 1 | 4.169925001 | 0 | 4.169925001 |
| Q9VPB1 | ST7_DROME Protein ST7 homolog OS=Drosophila melanogaster OX=7227 GN=CG3634 PE=2 SV=1 | 18 | 12 | 1 | 1 | 4.169925001 | 0 | 4.169925001 |
| A4V4F2 | A4V4F2_DROME Flotillin 2, isoform F OS=Drosophila melanogaster OX=7227 GN=Flo2 PE=1 SV=1 | 18 | 12 | 1 | 1 | 4.169925001 | 0 | 4.169925001 |
| M9PF50 | M9PF50_DROME Piezo-type mechanosensitive ion channel component OS=Drosophila melanogaster OX=7227 GN=Piezo PE=1 SV=1 | 18 | 16 | 1 | 1 | 4.169925001 | 0 | 4.169925001 |
| O46342 | O46342_DROME Amiloride-sensitive Na+ channel OS=Drosophila melanogaster OX=7227 GN=rpk PE=2 SV=1 | 18 | 12 | 1 | 1 | 4.169925001 | 0 | 4.169925001 |
| Q7JRM5 | Q7JRM5_DROME GH12965p OS=Drosophila melanogaster OX=7227 GN=CG11127 PE=1 SV=1 | 18 | 12 | 1 | 1 | 4.169925001 | 0 | 4.169925001 |
| Q9VH65 | Q9VH65_DROME FI11941p OS=Drosophila melanogaster OX=7227 GN=CG12945-RB PE=2 SV=2 | 18 | 12 | 1 | 1 | 4.169925001 | 0 | 4.169925001 |
| Q9VIM0 | Q9VIM0_DROME GH14278p OS=Drosophila melanogaster OX=7227 GN=38C.55 PE=1 SV=1 | 18 | 9 | 1 | 1 | 4.169925001 | 0 | 4.169925001 |
| Q9VQ60 | Q9VQ60_DROME LD12115p OS=Drosophila melanogaster OX=7227 GN=Dmel\CG7289 PE=2 SV=2 | 18 | 12 | 1 | 1 | 4.169925001 | 0 | 4.169925001 |
| Q9VVA1 | Q9VVA1_DROME FI21241p1 OS=Drosophila melanogaster OX=7227 GN=CG9706-RA PE=1 SV=1 | 18 | 8 | 1 | 1 | 4.169925001 | 0 | 4.169925001 |
| Q9W2H8 | Q9W2H8_DROME CG9485, isoform B OS=Drosophila melanogaster OX=7227 GN=CG9485 PE=1 SV=3 | 18 | 14 | 1 | 1 | 4.169925001 | 0 | 4.169925001 |
| Q9W402 | Q9W402_DROME NADH dehydrogenase (Ubiquinone) B16.6 subunit, isoform A OS=Drosophila melanogaster OX=7227 GN=ND-B16.6 PE=1 SV=1 | 18 | 8 | 1 | 1 | 4.169925001 | 0 | 4.169925001 |
| Q9VFT4 | Q9VFT4_DROME AT27578p OS=Drosophila melanogaster OX=7227 GN=rin PE=1 SV=1 | 17 | 11 | 112 | 34 | 4.087462841 | 6.807354922 | -2.719892081 |
| E2QCF1 | E2QCF1_DROME ATP-citrate synthase OS=Drosophila melanogaster OX=7227 GN=ATPCL PE=1 SV=1 | 17 | 13 | 66 | 26 | 4.087462841 | 6.044394119 | -1.956931278 |
| P48591 | RIR1_DROME Ribonucleoside-diphosphate reductase large subunit OS=Drosophila melanogaster OX=7227 GN=RnrL PE=1 SV=2 | 17 | 9 | 55 | 23 | 4.087462841 | 5.781359714 | -1.693896872 |
| P17210 | KINH_DROME Kinesin heavy chain OS=Drosophila melanogaster OX=7227 GN=Khc PE=1 SV=2 | 17 | 14 | 52 | 28 | 4.087462841 | 5.700439718 | -1.612976877 |
| Q9V3G1 | RL8_DROME 60S ribosomal protein L8 OS=Drosophila melanogaster OX=7227 GN=RpL8 PE=1 SV=1 | 17 | 7 | 52 | 11 | 4.087462841 | 5.700439718 | -1.612976877 |
| Q9VLM8 | SYAC_DROME Alanine--tRNA ligase, cytoplasmic OS=Drosophila melanogaster OX=7227 GN=Aats-ala PE=2 SV=1 | 17 | 16 | 47 | 29 | 4.087462841 | 5.554588852 | -1.46712601 |
| Q9VLC5 | Q9VLC5_DROME Aldehyde dehydrogenase OS=Drosophila melanogaster OX=7227 GN=Aldh PE=1 SV=1 | 17 | 10 | 39 | 22 | 4.087462841 | 5.285402219 | -1.197939378 |
| Q9W414 | Q9W414_DROME Regulatory particle triple-A ATPase 4 OS=Drosophila melanogaster OX=7227 GN=Rpt4 PE=1 SV=3 | 17 | 10 | 38 | 16 | 4.087462841 | 5.247927513 | -1.160464672 |
| Q9VJ39 | Q9VJ39_DROME Leukotriene A(4) hydrolase OS=Drosophila melanogaster OX=7227 GN=Dmel\CG10602 PE=1 SV=4 | 17 | 11 | 28 | 14 | 4.087462841 | 4.807354922 | -0.719892081 |
| Q9VK60 | Q9VK60_DROME GH25425p OS=Drosophila melanogaster OX=7227 GN=Dmel\CG6180 PE=1 SV=2 | 17 | 8 | 19 | 9 | 4.087462841 | 4.247927513 | -0.160464672 |
| Q24179 | SLY1_DROME Protein sly1 homolog OS=Drosophila melanogaster OX=7227 GN=Slh PE=2 SV=4 | 17 | 11 | 3 | 3 | 4.087462841 | 1.584962501 | 2.502500341 |
| Q8MLV1 | LBR_DROME Lamin-B receptor OS=Drosophila melanogaster OX=7227 GN=LBR PE=1 SV=1 | 17 | 10 | 2 | 2 | 4.087462841 | 1 | 3.087462841 |
| Q95RU0 | CUE_DROME Protein cueball OS=Drosophila melanogaster OX=7227 GN=cue PE=1 SV=1 | 17 | 11 | 1 | 1 | 4.087462841 | 0 | 4.087462841 |
| Q9W141 | ATPK_DROME Putative ATP synthase subunit f, mitochondrial OS=Drosophila melanogaster OX=7227 GN=CG4692 PE=3 SV=1 | 17 | 5 | 5 | 2 | 4.087462841 | 2.321928095 | 1.765534746 |
| Q9XZ63 | ARMET_DROME Mesencephalic astrocyte-derived neurotrophic factor homolog OS=Drosophila melanogaster OX=7227 GN=Manf PE=2 SV=2 | 17 | 11 | 9 | 7 | 4.087462841 | 3.169925001 | 0.91753784 |
| A0A0B4KGF7 | A0A0B4KGF7_DROME Beta-Mannosidase, isoform H OS=Drosophila melanogaster OX=7227 GN=beta-Man PE=1 SV=1 | 17 | 11 | 4 | 3 | 4.087462841 | 2 | 2.087462841 |
| Q7JR49 | Q7JR49_DROME LD09158p OS=Drosophila melanogaster OX=7227 GN=VhaAC45 PE=1 SV=1 | 17 | 8 | 5 | 4 | 4.087462841 | 2.321928095 | 1.765534746 |
| Q7JZ25 | Q7JZ25_DROME Chloride channel protein OS=Drosophila melanogaster OX=7227 GN=ClC-b PE=1 SV=1 | 17 | 12 | 1 | 1 | 4.087462841 | 0 | 4.087462841 |
| Q9VLL5 | Q9VLL5_DROME LP03571p OS=Drosophila melanogaster OX=7227 GN=172F5T PE=2 SV=2 | 17 | 14 | 2 | 2 | 4.087462841 | 1 | 3.087462841 |
| Q9VTB4 | Q9VTB4_DROME NADH dehydrogenase (Ubiquinone) 13 kDa B subunit OS=Drosophila melanogaster OX=7227 GN=ND-13B PE=1 SV=2 | 17 | 6 | 3 | 2 | 4.087462841 | 1.584962501 | 2.502500341 |
| Q9VWI0 | Q9VWI0_DROME NADH dehydrogenase (Ubiquinone) 18 kDa subunit OS=Drosophila melanogaster OX=7227 GN=ND-18 PE=1 SV=2 | 17 | 7 | 6 | 4 | 4.087462841 | 2.584962501 | 1.502500341 |
| Q95ST2 | WLS_DROME Protein wntless OS=Drosophila melanogaster OX=7227 GN=wls PE=1 SV=1 | 17 | 7 | 1 | 1 | 4.087462841 | 0 | 4.087462841 |
| Q9NIV1 | E2AK3_DROME Eukaryotic translation initiation factor 2-alpha kinase OS=Drosophila melanogaster OX=7227 GN=PEK PE=1 SV=2 | 17 | 13 | 1 | 1 | 4.087462841 | 0 | 4.087462841 |
| Q9V7H4 | TM131_DROME Transmembrane protein 131 homolog OS=Drosophila melanogaster OX=7227 GN=CG8370 PE=1 SV=2 | 17 | 14 | 1 | 1 | 4.087462841 | 0 | 4.087462841 |
| Q7JRC3 | Q7JRC3_DROME Epoxide hydrolase OS=Drosophila melanogaster OX=7227 GN=Jheh1 PE=2 SV=1 | 17 | 9 | 1 | 1 | 4.087462841 | 0 | 4.087462841 |
| Q8IN25 | Q8IN25_DROME Aminopeptidase OS=Drosophila melanogaster OX=7227 GN=CG5845 PE=1 SV=1 | 17 | 12 | 1 | 1 | 4.087462841 | 0 | 4.087462841 |
| Q8INK3 | Q8INK3_DROME Fatty acid binding protein, isoform C OS=Drosophila melanogaster OX=7227 GN=fabp PE=1 SV=1 | 17 | 8 | 1 | 1 | 4.087462841 | 0 | 4.087462841 |
| Q8IQD8 | Q8IQD8_DROME Phospholipid scramblase OS=Drosophila melanogaster OX=7227 GN=scramb1 PE=3 SV=1 | 17 | 8 | 1 | 1 | 4.087462841 | 0 | 4.087462841 |
| Q8SXS0 | Q8SXS0_DROME RE40914p OS=Drosophila melanogaster OX=7227 GN=BcDNA:RE40914 PE=2 SV=1 | 17 | 10 | 1 | 1 | 4.087462841 | 0 | 4.087462841 |
| Q9VBC5 | Q9VBC5_DROME FI18833p1 OS=Drosophila melanogaster OX=7227 GN=CG5484-RA PE=1 SV=1 | 17 | 7 | 1 | 1 | 4.087462841 | 0 | 4.087462841 |
| Q9VK28 | Q9VK28_DROME LD10349p OS=Drosophila melanogaster OX=7227 GN=CT35580 PE=1 SV=1 | 17 | 15 | 1 | 1 | 4.087462841 | 0 | 4.087462841 |
| Q9VL02 | Q9VL02_DROME GH08677p OS=Drosophila melanogaster OX=7227 GN=nmd PE=2 SV=1 | 17 | 9 | 1 | 1 | 4.087462841 | 0 | 4.087462841 |
| Q9VN25 | EIF3A_DROME Eukaryotic translation initiation factor 3 subunit A OS=Drosophila melanogaster OX=7227 GN=eIF3a PE=1 SV=1 | 16 | 10 | 71 | 30 | 4 | 6.14974712 | -2.14974712 |
| Q9VL70 | Q9VL70_DROME HL08109p OS=Drosophila melanogaster OX=7227 GN=yip2 PE=1 SV=1 | 16 | 10 | 51 | 16 | 4 | 5.672425342 | -1.672425342 |
| P48375 | FKB12_DROME 12 kDa FK506-binding protein OS=Drosophila melanogaster OX=7227 GN=FK506-bp2 PE=3 SV=2 | 16 | 4 | 46 | 6 | 4 | 5.523561956 | -1.523561956 |
| Q9VFQ9 | Q9VFQ9_DROME Dipeptidase B, isoform A OS=Drosophila melanogaster OX=7227 GN=Dip-B PE=1 SV=2 | 16 | 10 | 44 | 18 | 4 | 5.459431619 | -1.459431619 |
| Q8IRH1 | Q8IRH1_DROME FI19310p1 OS=Drosophila melanogaster OX=7227 GN=Psa PE=1 SV=1 | 16 | 14 | 40 | 21 | 4 | 5.321928095 | -1.321928095 |
| Q9VB10 | Q9VB10_DROME GH01709p OS=Drosophila melanogaster OX=7227 GN=Dmel\CG5590 PE=1 SV=1 | 16 | 12 | 35 | 13 | 4 | 5.129283017 | -1.129283017 |
| M9MQH9 | M9MQH9_DROME Protein phosphatase 2A at 29B, isoform C OS=Drosophila melanogaster OX=7227 GN=Pp2A-29B PE=1 SV=1 | 16 | 11 | 33 | 16 | 4 | 5.044394119 | -1.044394119 |
| P17917 | PCNA_DROME Proliferating cell nuclear antigen OS=Drosophila melanogaster OX=7227 GN=PCNA PE=1 SV=2 | 16 | 8 | 32 | 12 | 4 | 5 | -1 |
| Q9Y112 | Q9Y112_DROME BcDNA.GH10614 OS=Drosophila melanogaster OX=7227 GN=AKR2E3 PE=1 SV=1 | 16 | 9 | 32 | 12 | 4 | 5 | -1 |
| Q9VJ19 | Q9VJ19_DROME RE25263p OS=Drosophila melanogaster OX=7227 GN=RpL30 PE=1 SV=1 | 16 | 5 | 31 | 9 | 4 | 4.95419631 | -0.95419631 |

| UniProt ID | Protein description | # PSMs MVB spectra | # of unique peptides MVB | # of PSMs lysate | # of unique peptides lysate | log2(PSM MVB) | log2(PSM lysate) | FC (log2 of MVB PSM-log2 of lysate PSM) |
|---|---|---|---|---|---|---|---|---|
| A1ZBJ2 | A1ZBJ2_DROME CG7461, isoform B OS=Drosophila melanogaster OX=7227 GN=CG7461 PE=1 SV=2 | 16 | 14 | 26 | 14 | 4 | 4.700439718 | -0.700439718 |
| Q9V784 | SAM50_DROME SAM50-like protein CG7639 OS=Drosophila melanogaster OX=7227 GN=CG7639 PE=2 SV=2 | 16 | 9 | 3 | 3 | 4 | 1.584962501 | 2.415037499 |
| Q9V7D2 | VATD1_DROME V-type proton ATPase subunit D 1 OS=Drosophila melanogaster OX=7227 GN=Vha36-1 PE=2 SV=1 | 16 | 8 | 8 | 5 | 4 | 3 | 1 |
| Q9VPQ2 | DJSHV_DROME DnaJ homolog shv OS=Drosophila melanogaster OX=7227 GN=shv PE=1 SV=1 | 16 | 6 | 4 | 3 | 4 | 2 | 2 |
| Q9W0H3 | LDAH_DROME Lipid droplet-associated hydrolase OS=Drosophila melanogaster OX=7227 GN=CG9186 PE=2 SV=1 | 16 | 11 | 1 | 1 | 4 | 0 | 4 |
| A0A0B4LGC8 | A0A0B4LGC8_DROME CG4585, isoform B OS=Drosophila melanogaster OX=7227 GN=CG4585 PE=4 SV=1 | 16 | 8 | 1 | 1 | 4 | 0 | 4 |
| Q6NN09 | Q6NN09_DROME ATP synthase, subunit C, isoform A OS=Drosophila melanogaster OX=7227 GN=ATPsynC PE=2 SV=1 | 16 | 1 | 13 | 1 | 4 | 3.700439718 | 0.299560282 |
| Q7K0X9 | Q7K0X9_DROME LD23667p OS=Drosophila melanogaster OX=7227 GN=Syx7 PE=1 SV=1 | 16 | 7 | 3 | membrane | 4 | 1.584962501 | 2.415037499 |
| Q7KTG2 | Q7KTG2_DROME Apolipoprotein lipid transfer particle OS=Drosophila melanogaster OX=7227 GN=Apoltp PE=1 SV=2 | 16 | 15 | 1 | 1 | 4 | 0 | 4 |
| Q95RF6 | Q95RF6_DROME LD34461p OS=Drosophila melanogaster OX=7227 GN=Tom20 PE=1 SV=1 | 16 | 6 | 7 | 3 | 4 | 2.807354922 | 1.192645078 |
| Q9VAJ9 | Q9VAJ9_DROME SD09259p OS=Drosophila melanogaster OX=7227 GN=CG 1907 PE=1 SV=1 | 16 | 8 | 3 | 2 | 4 | 1.584962501 | 2.415037499 |
| Q9VKM3 | Q9VKM3_DROME ATP synthase, subunit G, isoform A OS=Drosophila melanogaster OX=7227 GN=ATPsynG PE=1 SV=1 | 16 | 6 | 7 | 5 | 4 | 2.807354922 | 1.192645078 |
| Q9VZF6 | Q9VZF6_DROME GH04863p OS=Drosophila melanogaster OX=7227 GN=BcDNA:GH04863 PE=1 SV=1 | 16 | 14 | 2 | 2 | 4 | 1 | 3 |
| X2JGV8 | X2JGV8_DROME Uncharacterized protein, isoform B OS=Drosophila melanogaster OX=7227 GN=CG9231 PE=4 SV=1 | 16 | 6 | 2 | 2 | 4 | 1 | 3 |
| Q9VMM2 | Q9VMM2_DROME Uncharacterized protein, isoform B OS=Drosophila melanogaster OX=7227 GN=Dmel\CG11034 PE=3 SV=2 | 16 | 10 | 1 | 1 | 4 | 0 | 4 |
| Q9VYT6 | Q9VYT6_DROME RE49489p OS=Drosophila melanogaster OX=7227 GN=p24-1 PE=1 SV=1 | 16 | 6 | 1 | 1 | 4 | 0 | 4 |
| Q9Y114 | Q9Y114_DROME BcDNA.GH10229 OS=Drosophila melanogaster OX=7227 GN=BcDNA.GH10229 PE=1 SV=1 | 16 | 10 | 1 | 1 | 4 | 0 | 4 |
| Q7KRW1 | TRC8_DROME Protein TRC8 homolog OS=Drosophila melanogaster OX=7227 GN=Trc8 PE=1 SV=1 | 16 | 9 | 1 | membrane | 4 | 0 | 4 |
| Q9VGG5 | CAD87_DROME Cadherin-87A OS=Drosophila melanogaster OX=7227 GN=Cad87A PE=1 SV=4 | 16 | 13 | 1 | 1 | 4 | 0 | 4 |
| Q9VXT2 | DJC25_DROME DnaJ homolog subfamily C member 25 homolog OS=Drosophila melanogaster OX=7227 GN=CG7872 PE=2 SV=1 | 16 | 8 | 1 | 1 | 4 | 0 | 4 |
| A1Z9W2 | A1Z9W2_DROME RH47216p OS=Drosophila melanogaster OX=7227 GN=CG33506-RA PE=2 SV=1 | 16 | 6 | 1 | 1 | 4 | 0 | 4 |
| E1JHE4 | E1JHE4_DROME Fatty acid (Long chain) transport protein, isoform F OS=Drosophila melanogaster OX=7227 GN=Fatp PE=1 SV=3 | 16 | 12 | 1 | 1 | 4 | 0 | 4 |
| M9PDR3 | M9PDR3_DROME Leukocyte-antigen-related-like, isoform G OS=Drosophila melanogaster OX=7227 GN=Lar PE=4 SV=1 | 16 | 14 | 1 | 1 | 4 | 0 | 4 |
| Q4U2G8 | Q4U2G8_DROME LD02432p OS=Drosophila melanogaster OX=7227 GN=CG11961 PE=1 SV=1 | 16 | 13 | 1 | 1 | 4 | 0 | 4 |
| Q7JZN2 | Q7JZN2_DROME Multiple C2 domain and transmembrane region protein, isoform B OS=Drosophila melanogaster OX=7227 GN=Mctp PE=2 SV=1 | 16 | 9 | 1 | 1 | 4 | 0 | 4 |
| Q8IQH0 | Q8IQH0_DROME FI12817p OS=Drosophila melanogaster OX=7227 GN=Nrx-IV PE=1 SV=2 | 16 | 14 | 1 | 1 | 4 | 0 | 4 |
| Q8MLQ5 | Q8MLQ5_DROME Kazachoc, isoform D OS=Drosophila melanogaster OX=7227 GN=kcc PE=1 SV=1 | 16 | 11 | 1 | 1 | 4 | 0 | 4 |
| Q8WR08 | Q8WR08_DROME Dystroglycan type I OS=Drosophila melanogaster OX=7227 GN=Dg PE=1 SV=1 | 16 | 11 | 1 | 1 | 4 | 0 | 4 |
| Q9VG80 | Q9VG80_DROME Palmitoyltransferase OS=Drosophila melanogaster OX=7227 GN=CG5196-RA PE=2 SV=1 | 16 | 9 | 1 | 1 | 4 | 0 | 4 |
| Q9VKU2 | Q9VKU2_DROME LD46023p OS=Drosophila melanogaster OX=7227 GN=Dmel\CG7384 PE=2 SV=2 | 16 | 10 | 1 | 1 | 4 | 0 | 4 |
| Q9VNI9 | Q9VNI9_DROME IP18173p OS=Drosophila melanogaster OX=7227 GN=Cg2023 PE=1 SV=1 | 16 | 8 | 1 | 1 | 4 | 0 | 4 |
| Q9VPX6 | Q9VPX6_DROME Adenylyl cyclase-associated protein OS=Drosophila melanogaster OX=7227 GN=capt PE=1 SV=2 | 16 | 7 | 1 | 1 | 4 | 0 | 4 |
| Q9VQ29 | Q9VQ29_DROME Cytochrome b-c1 complex subunit Rieske, mitochondrial OS=Drosophila melanogaster OX=7227 GN=RFeSP PE=1 SV=3 | 16 | 7 | 1 | 1 | 4 | 0 | 4 |
| X2J9P9 | X2J9P9_DROME Cup, isoform C OS=Drosophila melanogaster OX=7227 GN=cup PE=4 SV=1 | 15 | 13 | 106 | 44 | 3.906890596 | 6.727920455 | -2.821029859 |
| P23128 | DDX6_DROME Putative ATP-dependent RNA helicase me31b OS=Drosophila melanogaster OX=7227 GN=me31B PE=1 SV=3 | 15 | 12 | 70 | 32 | 3.906890596 | 6.129283017 | -2.222392421 |
| Q7KN75 | Q7KN75_DROME Dodeca-satellite-binding protein 1, isoform A OS=Drosophila melanogaster OX=7227 GN=Dp1 PE=1 SV=1 | 15 | 15 | 66 | 32 | 3.906890596 | 6.044394119 | -2.137503524 |
| M9PG76 | M9PG76_DROME 60S acidic ribosomal protein P0 OS=Drosophila melanogaster OX=7227 GN=RpLP0 PE=3 SV=1 | 15 | 8 | 58 | 12 | 3.906890596 | 5.857980995 | -1.9510904 |
| Q9VL78 | FKB59_DROME FK506-binding protein 59 OS=Drosophila melanogaster OX=7227 GN=FKBP59 PE=1 SV=1 | 15 | 11 | 57 | 18 | 3.906890596 | 5.832890014 | -1.925999419 |
| A0A126GUT1 | A0A126GUT1_DROME Translationally controlled tumor protein, isoform B OS=Drosophila melanogaster OX=7227 GN=Tctp PE=3 SV=1 | 15 | 8 | 49 | 12 | 3.906890596 | 5.614709844 | -1.707819249 |
| P50882 | RL9_DROME 60S ribosomal protein L9 OS=Drosophila melanogaster OX=7227 GN=RpL9 PE=1 SV=2 | 15 | 10 | 41 | 14 | 3.906890596 | 5.357552005 | -1.450661409 |
| Q9VMI5 | Q9VMI5_DROME CG9135 protein OS=Drosophila melanogaster OX=7227 GN=Dmel\CG9135 PE=1 SV=1 | 15 | 9 | 30 | 14 | 3.906890596 | 4.906890596 | -1 |
| Q24572 | CAF1_DROME Probable histone-binding protein Caf1 OS=Drosophila melanogaster OX=7227 GN=Caf1 PE=1 SV=1 | 15 | 8 | 22 | 9 | 3.906890596 | 4.459431619 | -0.552541023 |
| P22465 | ANX10_DROME Annexin B10 OS=Drosophila melanogaster OX=7227 GN=AnxB10 PE=2 SV=3 | 15 | 10 | 20 | 11 | 3.906890596 | 4.321928095 | -0.415037499 |
| Q8T045 | RUMI_DROME O-glucosyltransferase rumi OS=Drosophila melanogaster OX=7227 GN=rumi PE=1 SV=1 | 15 | 9 | 9 | 7 | 3.906890596 | 3.169925001 | 0.736965594 |
| Q9U915 | KAD2_DROME Adenylate kinase OS=Drosophila melanogaster OX=7227 GN=Adk2 PE=1 SV=1 | 15 | 8 | 8 | 5 | 3.906890596 | 3 | 0.906890596 |
| Q9V3S9 | BGM_DROME Very long-chain-fatty-acid--CoA ligase bubblegum OS=Drosophila melanogaster OX=7227 GN=bgm PE=2 SV=1 | 15 | 12 | 8 | 8 | 3.906890596 | 3 | 0.906890596 |
| Q9VAF0 | S39AD_DROME Zinc transporter ZIP13 homolog OS=Drosophila melanogaster OX=7227 GN=Zip99C PE=2 SV=1 | 15 | 4 | 4 | 2 | 3.906890596 | 2 | 1.906890596 |
| Q9VRJ2 | FITM_DROME FIT family protein CG10671 OS=Drosophila melanogaster OX=7227 GN=CG10671 PE=1 SV=1 | 15 | 5 | 1 | 1 | 3.906890596 | 0 | 3.906890596 |
| A0A0B4KHH2 | A0A0B4KHH2_DROME Septin interacting protein 3, isoform B OS=Drosophila melanogaster OX=7227 GN=sip3 PE=1 SV=1 | 15 | 9 | 1 | 1 | 3.906890596 | 0 | 3.906890596 |
| O97066 | O97066_DROME Signal peptidase complex catalytic subunit SEC11 OS=Drosophila melanogaster OX=7227 GN=twr PE=1 SV=1 | 15 | 8 | 1 | 1 | 3.906890596 | 0 | 3.906890596 |
| Q8T0Q4 | Q8T0Q4_DROME GH13992p OS=Drosophila melanogaster OX=7227 GN=shrb PE=1 SV=1 | 15 | 7 | 13 | 7 | 3.906890596 | 3.700439718 | 0.206450877 |
| Q9VAP8 | Q9VAP8_DROME alpha-1,2-Mannosidase OS=Drosophila melanogaster OX=7227 GN=alpha-Man-Ib PE=1 SV=1 | 15 | 12 | 2 | 1 | 3.906890596 | 1 | 2.906890596 |
| Q9VNF9 | Q9VNF9_DROME Ethanol sensitive with low memory, isoform A OS=Drosophila melanogaster OX=7227 GN=elm PE=1 SV=1 | 15 | 7 | 3 | 3 | 3.906890596 | 1.584962501 | 2.321928095 |
| Q9VQD7 | Q9VQD7_DROME NADH dehydrogenase [ubiquinone] 1 alpha subcomplex subunit 12 OS=Drosophila melanogaster OX=7227 GN=ND-B17.2 PE=1 SV=1 | 15 | 6 | 5 | 4 | 3.906890596 | 2.321928095 | 1.584962501 |
| Q9VU45 | Q9VU45_DROME LD27581p OS=Drosophila melanogaster OX=7227 GN=Syx13 PE=1 SV=1 | 15 | 5 | 2 | 2 | 3.906890596 | 1 | 2.906890596 |
| Q9VV51 | Q9VV51_DROME RE10780p OS=Drosophila melanogaster OX=7227 GN=AGPAT3 PE=1 SV=1 | 15 | 8 | 2 | 2 | 3.906890596 | 1 | 2.906890596 |
| Q9VXK7 | Q9VXK7_DROME LD31474p OS=Drosophila melanogaster OX=7227 GN=ND-20 PE=2 SV=1 | 15 | 8 | 2 | 2 | 3.906890596 | 1 | 2.906890596 |
| Q9W2S1 | Q9W2S1_DROME FI17817p1 OS=Drosophila melanogaster OX=7227 GN=Tango5 PE=1 SV=1 | 15 | 7 | 4 | 3 | 3.906890596 | 2 | 1.906890596 |
| Q9VPQ7 | Q9VPQ7_DROME SD07518p OS=Drosophila melanogaster OX=7227 GN=Spp PE=1 SV=1 | 15 | 6 | 1 | 1 | 3.906890596 | 0 | 3.906890596 |
| Q6WV16 | GALT6_DROME N-acetylgalactosaminyltransferase 6 OS=Drosophila melanogaster OX=7227 GN=pgant6 PE=2 SV=2 | 15 | 9 | 1 | 1 | 3.906890596 | 0 | 3.906890596 |
| A0A0B4KHC7 | A0A0B4KHC7_DROME Uncharacterized protein, isoform B OS=Drosophila melanogaster OX=7227 GN=CG7685 PE=4 SV=1 | 15 | 6 | 1 | 1 | 3.906890596 | 0 | 3.906890596 |
| A0A0B4KHL4 | A0A0B4KHL4_DROME Gilgamesh, isoform L OS=Drosophila melanogaster OX=7227 GN=gish PE=1 SV=1 | 15 | 12 | 1 | 1 | 3.906890596 | 0 | 3.906890596 |
| O97132 | O97132_DROME Anoctamin OS=Drosophila melanogaster OX=7227 GN=Axs PE=3 SV=1 | 15 | 11 | 1 | 1 | 3.906890596 | 0 | 3.906890596 |
| Q0E8F2 | Q0E8F2_DROME FI05451p OS=Drosophila melanogaster OX=7227 GN=CG18022 PE=1 SV=1 | 15 | 7 | 1 | 1 | 3.906890596 | 0 | 3.906890596 |
| Q8SYL0 | Q8SYL0_DROME CDP-diacylglycerol--glycerol-3-phosphate 3-phosphatidyltransferase OS=Drosophila melanogaster OX=7227 GN=Dmel\CG7718 PE=2 SV=1 | 15 | 10 | 1 | 1 | 3.906890596 | 0 | 3.906890596 |
| Q9VGF5 | Q9VGF5_DROME LP09696p OS=Drosophila melanogaster OX=7227 GN=Tango9 PE=1 SV=1 | 15 | 4 | 1 | 1 | 3.906890596 | 0 | 3.906890596 |
| Q9VGH0 | Q9VGH0_DROME LD28817p OS=Drosophila melanogaster OX=7227 GN=Dmel\CG6962 PE=1 SV=1 | 15 | 11 | 1 | 1 | 3.906890596 | 0 | 3.906890596 |
| Q9VIU4 | Q9VIU4_DROME FI05338p OS=Drosophila melanogaster OX=7227 GN=CG10355 PE=1 SV=2 | 15 | 7 | 1 | 1 | 3.906890596 | 0 | 3.906890596 |
| Q9VVX6 | Q9VVX6_DROME Blocked early in transport 1, isoform A OS=Drosophila melanogaster OX=7227 GN=Bet1 PE=2 SV=1 | 15 | 6 | 1 | 1 | 3.906890596 | 0 | 3.906890596 |
| Q9VW73 | Q9VW73_DROME FI16105p1 OS=Drosophila melanogaster OX=7227 GN=CG14184-RB PE=1 SV=1 | 15 | 8 | 1 | 1 | 3.906890596 | 0 | 3.906890596 |
| X2JDR3 | X2JDR3_DROME Sidekick, isoform I OS=Drosophila melanogaster OX=7227 GN=sdk PE=4 SV=1 | 15 | 14 | 1 | 1 | 3.906890596 | 0 | 3.906890596 |
| Q9W5R8 | RL5_DROME 60S ribosomal protein L5 OS=Drosophila melanogaster OX=7227 GN=RpL5 PE=1 SV=1 | 14 | 7 | 106 | 15 | 3.807354922 | 6.727920455 | -2.920565533 |
| M9PG62 | M9PG62_DROME Regulatory particle non-ATPase 3, isoform B OS=Drosophila melanogaster OX=7227 GN=Rpn3 PE=1 SV=1 | 14 | 7 | 61 | 31 | 3.807354922 | 5.930737338 | -2.123382416 |
| A0A0B4K7Z5 | A0A0B4K7Z5_DROME Regulatory particle non-ATPase 6, isoform C OS=Drosophila melanogaster OX=7227 GN=Rpn6 PE=1 SV=1 | 14 | 12 | 57 | 23 | 3.807354922 | 5.832890014 | -2.025535092 |
| Q9TVM2 | XPO1_DROME Exportin-1 OS=Drosophila melanogaster OX=7227 GN=emb PE=1 SV=1 | 14 | 11 | 49 | 24 | 3.807354922 | 5.614709844 | -1.807354922 |
| O96827 | EF1B_DROME Probable elongation factor 1-beta OS=Drosophila melanogaster OX=7227 GN=eEF1beta PE=1 SV=3 | 14 | 9 | 46 | 16 | 3.807354922 | 5.523561956 | -1.716207034 |
| A0A126BEE6 | A0A126BEE6_DROME Uncharacterized protein, isoform B OS=Drosophila melanogaster OX=7227 GN=CG8223 PE=4 SV=1 | 14 | 8 | 46 | 15 | 3.807354922 | 5.523561956 | -1.716207034 |
| Q9VV60 | Q9VV60_DROME Tyrosine--tRNA ligase OS=Drosophila melanogaster OX=7227 GN=TyrRS PE=1 SV=1 | 14 | 11 | 39 | 23 | 3.807354922 | 5.285402219 | -1.478047297 |
| P17704 | RS17_DROME 40S ribosomal protein S17 OS=Drosophila melanogaster OX=7227 GN=RpS17 PE=1 SV=2 | 14 | 7 | 36 | 12 | 3.807354922 | 5.169925001 | -1.362570079 |
| Q9VB05 | Q9VB05_DROME ALG-2 interacting protein X OS=Drosophila melanogaster OX=7227 GN=ALiX PE=1 SV=1 | 14 | 12 | 33 | 21 | 3.807354922 | 5.044394119 | -1.237039197 |
| P48149 | RS15A_DROME 40S ribosomal protein S15Aa OS=Drosophila melanogaster OX=7227 GN=RpS15Aa PE=1 SV=2 | 14 | 8 | 32 | 11 | 3.807354922 | 5 | -1.192645078 |

| UniProt ID | Protein description | # PSMs MVB spectra | # of unique peptides MVB | # of PSMs lysate | # of unique peptides lysate | log2(PSM MVB) | log2(PSM lysate) | FC (log2 of MVB PSM-log2 of lysate PSM) |
|---|---|---|---|---|---|---|---|---|
| Q9W1H8 | Q9W1H8_DROME GH13256p OS=Drosophila melanogaster OX=7227 GN=Thiolase PE=1 SV=1 | 14 | 9 | 30 | 13 | 3.807354922 | 4.906890596 | -1.099535674 |
| Q9W457 | Q9W457_DROME Serine hydroxymethyltransferase OS=Drosophila melanogaster OX=7227 GN=Shmt PE=1 SV=1 | 14 | 10 | 29 | 15 | 3.807354922 | 4.857980995 | -1.050626073 |
| Q9VC18 | Q9VC18_DROME RE29555p OS=Drosophila melanogaster OX=7227 GN=153761_at PE=1 SV=1 | 14 | 13 | 28 | 17 | 3.807354922 | 4.807354922 | -1 |
| P36951 | HYI_DROME Putative hydroxypyruvate isomerase OS=Drosophila melanogaster OX=7227 GN=Gip PE=2 SV=1 | 14 | 9 | 26 | 8 | 3.807354922 | 4.700439718 | -0.893084796 |
| P36241 | RL19_DROME 60S ribosomal protein L19 OS=Drosophila melanogaster OX=7227 GN=RpL19 PE=1 SV=2 | 14 | 8 | 24 | 10 | 3.807354922 | 4.584962501 | -0.777607579 |
| F0JAI1 | F0JAI1_DROME Arginine methyltransferase 4, isoform B OS=Drosophila melanogaster OX=7227 GN=Art4 PE=1 SV=1 | 14 | 8 | 24 | 10 | 3.807354922 | 4.584962501 | -0.777607579 |
| Q9V9M7 | Q9V9M7_DROME RE62581p OS=Drosophila melanogaster OX=7227 GN=RpL21 PE=1 SV=1 | 14 | 6 | 24 | 9 | 3.807354922 | 4.584962501 | -0.777607579 |
| A0A0B4LGX1 | A0A0B4LGX1_DROME CG5174, isoform P OS=Drosophila melanogaster OX=7227 GN=CG5174 PE=1 SV=1 | 14 | 6 | 19 | 9 | 3.807354922 | 4.247927513 | -0.440572591 |
| O62621 | COPB2_DROME Coatomer subunit beta' OS=Drosophila melanogaster OX=7227 GN=beta'COP PE=2 SV=2 | 14 | 7 | 12 | 11 | 3.807354922 | 3.584962501 | 0.222392421 |
| Q8T9B6 | MESD_DROME LDLR chaperone boca OS=Drosophila melanogaster OX=7227 GN=boca PE=1 SV=2 | 14 | 8 | 8 | 5 | 3.807354922 | 3 | 0.807354922 |
| Q9VHV5 | ARL8_DROME ADP-ribosylation factor-like protein 8 OS=Drosophila melanogaster OX=7227 GN=Gie PE=1 SV=1 | 14 | 7 | 1 | 1 | 3.807354922 | 0 | 3.807354922 |
| Q9W5N0 | COA7_DROME Cytochrome c oxidase assembly factor 7 homolog OS=Drosophila melanogaster OX=7227 GN=CG13865 PE=2 SV=1 | 14 | 9 | 2 | 2 | 3.807354922 | 1 | 2.807354922 |
| Q9XZH6 | VATG_DROME V-type proton ATPase subunit G OS=Drosophila melanogaster OX=7227 GN=Vha13 PE=3 SV=1 | 14 | 3 | 2 | 1 | 3.807354922 | 1 | 2.807354922 |
| Q0E8H9 | Q0E8H9_DROME Beta-hexosaminidase OS=Drosophila melanogaster OX=7227 GN=Hexo1 PE=1 SV=1 | 14 | 9 | 3 | 2 | 3.807354922 | 1.584962501 | 2.222392421 |
| Q7JX87 | Q7JX87_DROME Peroxiredoxin 2540-2 OS=Drosophila melanogaster OX=7227 GN=Prx2540-2 PE=1 SV=1 | 14 | 6 | 13 | 8 | 3.807354922 | 3.700439718 | 0.106915204 |
| Q7JYX5 | Q7JYX5_DROME RH15778p OS=Drosophila melanogaster OX=7227 GN=Vamp7 PE=1 SV=1 | 14 | 8 | 2 | 1 | 3.807354922 | 1 | 2.807354922 |
| Q8IRR0 | Q8IRR0_DROME Uncharacterized protein, isoform B OS=Drosophila melanogaster OX=7227 GN=CG3566 PE=1 SV=3 | 14 | 6 | 2 | 2 | 3.807354922 | 1 | 2.807354922 |
| Q9VF14 | Q9VF14_DROME Endoribonuclease U-specific OS=Drosophila melanogaster OX=7227 GN=EndoU PE=2 SV=2 | 14 | 7 | 2 | 2 | 3.807354922 | 1 | 2.807354922 |
| Q9VVH5 | Q9VVH5_DROME AT13736p OS=Drosophila melanogaster OX=7227 GN=UQCR-Q PE=1 SV=1 | 14 | 5 | 3 | 2 | 3.807354922 | 1.584962501 | 2.222392421 |
| Q9VZS1 | Q9VZS1_DROME LD28127p OS=Drosophila melanogaster OX=7227 GN=Cg17746 PE=1 SV=1 | 14 | 9 | 7 | 6 | 3.807354922 | 2.807354922 | 1 |
| Q9W1F7 | Q9W1F7_DROME CG2970, isoform A OS=Drosophila melanogaster OX=7227 GN=CG2970 PE=1 SV=2 | 14 | 6 | 5 | 5 | 3.807354922 | 2.321928095 | 1.485426827 |
| S5WMX5 | S5WMX5_DROME FI22366p1 OS=Drosophila melanogaster OX=7227 GN=Ppt1 PE=1 SV=1 | 14 | 8 | 3 | 2 | 3.807354922 | 1.584962501 | 2.222392421 |
| Q9VCE0 | Q9VCE0_DROME Phosphatidylserine decarboxylase proenzyme, mitochondrial OS=Drosophila melanogaster OX=7227 GN=Pisd PE=1 SV=1 | 14 | 9 | 1 | 1 | 3.807354922 | 0 | 3.807354922 |
| P13706 | GPDA_DROME Glycerol-3-phosphate dehydrogenase [NAD(+)], cytoplasmic OS=Drosophila melanogaster OX=7227 GN=Gpdh PE=1 SV=3 | 14 | 11 | 1 | 1 | 3.807354922 | 0 | 3.807354922 |
| Q7K237 | C1GLT_DROME Glycoprotein-N-acetylgalactosamine 3-beta-galactosyltransferase 1 OS=Drosophila melanogaster OX=7227 GN=C1GalTA PE=2 SV=1 | 14 | 9 | 1 | 1 | 3.807354922 | 0 | 3.807354922 |
| Q9VRI9 | CP6T1_DROME Probable cytochrome P450 6t1 OS=Drosophila melanogaster OX=7227 GN=Cyp6t1 PE=2 SV=1 | 14 | 10 | 1 | 1 | 3.807354922 | 0 | 3.807354922 |
| Q9W1K0 | TM14_DROME Transmembrane protein 14 homolog OS=Drosophila melanogaster OX=7227 GN=CG5532 PE=3 SV=1 | 14 | 5 | 1 | 1 | 3.807354922 | 0 | 3.807354922 |
| A0A0B4JCZ4 | A0A0B4JCZ4_DROME Syntaxin 1A, isoform B OS=Drosophila melanogaster OX=7227 GN=Syx1A PE=3 SV=1 | 14 | 5 | 1 | 1 | 3.807354922 | 0 | 3.807354922 |
| A0A0B4KF86 | A0A0B4KF86_DROME Autophagy-related protein 9 OS=Drosophila melanogaster OX=7227 GN=Atg9 PE=1 SV=1 | 14 | 8 | 1 | 1 | 3.807354922 | 0 | 3.807354922 |
| Q0E9E2 | Q0E9E2_DROME Pyruvate carboxylase OS=Drosophila melanogaster OX=7227 GN=PCB PE=1 SV=1 | 14 | 11 | 1 | 1 | 3.807354922 | 0 | 3.807354922 |
| Q7KK54 | Q7KK54_DROME MIP07328p OS=Drosophila melanogaster OX=7227 GN=Sema1b PE=1 SV=1 | 14 | 13 | 1 | 1 | 3.807354922 | 0 | 3.807354922 |
| Q8IQN2 | Q8IQN2_DROME Chloride channel protein OS=Drosophila melanogaster OX=7227 GN=ClC-c PE=3 SV=1 | 14 | 9 | 1 | 1 | 3.807354922 | 0 | 3.807354922 |
| Q9V4C6 | Q9V4C6_DROME Zinc/iron regulated transporter-related protein 102B, isoform E OS=Drosophila melanogaster OX=7227 GN=Zip102B PE=1 SV=3 | 14 | 5 | 1 | 1 | 3.807354922 | 0 | 3.807354922 |
| Q9VB99 | Q9VB99_DROME LD03691p OS=Drosophila melanogaster OX=7227 GN=Tsp97E PE=2 SV=1 | 14 | 4 | 1 | 1 | 3.807354922 | 0 | 3.807354922 |
| Q9VH76 | Q9VH76_DROME Synaptosomal-associated protein OS=Drosophila melanogaster OX=7227 GN=Snap24 PE=1 SV=1 | 14 | 10 | 1 | 1 | 3.807354922 | 0 | 3.807354922 |
| Q9VHF6 | Q9VHF6_DROME Another drosophila Unc-51-like kinase, isoform A OS=Drosophila melanogaster OX=7227 GN=Aduk PE=4 SV=1 | 14 | 9 | 1 | 1 | 3.807354922 | 0 | 3.807354922 |
| Q9VV76 | Q9VV76_DROME Syntaxin 8 OS=Drosophila melanogaster OX=7227 GN=Syx8 PE=1 SV=1 | 14 | 6 | 1 | 1 | 3.807354922 | 0 | 3.807354922 |
| Q9VX95 | Q9VX95_DROME LD21896p OS=Drosophila melanogaster OX=7227 GN=wus PE=2 SV=1 | 14 | 6 | 1 | 1 | 3.807354922 | 0 | 3.807354922 |
| Q9W0L6 | Q9W0L6_DROME GH22266p OS=Drosophila melanogaster OX=7227 GN=Dmel\CG13907 PE=1 SV=1 | 14 | 10 | 1 | 1 | 3.807354922 | 0 | 3.807354922 |
| P21187 | PABP_DROME Polyadenylate-binding protein OS=Drosophila melanogaster OX=7227 GN=pAbp PE=1 SV=3 | 13 | 11 | 75 | 30 | 3.700439718 | 6.22881869 | -2.528378972 |
| C4NYP8 | F10A1_DROME Hsc70-interacting protein 1 OS=Drosophila melanogaster OX=7227 GN=HIP PE=1 SV=2 | 13 | 10 | 63 | 17 | 3.700439718 | 5.977279923 | -2.276840205 |
| Q9XZU1 | XPO2_DROME Exportin-2 OS=Drosophila melanogaster OX=7227 GN=Cse1 PE=2 SV=2 | 13 | 10 | 51 | 19 | 3.700439718 | 5.672425342 | -1.971985624 |
| A0A0B4LEY6 | A0A0B4LEY6_DROME Nascent polypeptide associated complex protein alpha subunit, isoform D OS=Drosophila melanogaster OX=7227 GN=Nacalpha PE=4 SV=1 | 13 | 7 | 46 | 9 | 3.700439718 | 5.523561956 | -1.823122238 |
| X2JC82 | X2JC82_DROME Heat shock protein 27, isoform B OS=Drosophila melanogaster OX=7227 GN=Hsp27 PE=3 SV=1 | 13 | 9 | 43 | 11 | 3.700439718 | 5.426264755 | -1.725825037 |
| Q9VUY9 | PGM_DROME Phosphoglucomutase OS=Drosophila melanogaster OX=7227 GN=Pgm PE=1 SV=1 | 13 | 10 | 35 | 21 | 3.700439718 | 5.129283017 | -1.428843299 |
| Q9VQE0 | Q9VQE0_DROME Dynamin related protein 1, isoform A OS=Drosophila melanogaster OX=7227 GN=Drp1 PE=1 SV=1 | 13 | 10 | 35 | 22 | 3.700439718 | 5.129283017 | -1.428843299 |
| O61231 | RL10_DROME 60S ribosomal protein L10 OS=Drosophila melanogaster OX=7227 GN=RpL10 PE=1 SV=1 | 13 | 6 | 29 | 9 | 3.700439718 | 4.857980995 | -1.157541277 |
| Q9VAN7 | Q9VAN7_DROME Phosphoglycerate mutase OS=Drosophila melanogaster OX=7227 GN=Pglym78 PE=1 SV=2 | 13 | 8 | 28 | 9 | 3.700439718 | 4.807354922 | -1.106915204 |
| M9MRD0 | M9MRD0_DROME Ubiquitin carboxyl-terminal hydrolase OS=Drosophila melanogaster OX=7227 GN=Uch PE=3 SV=1 | 13 | 9 | 25 | 11 | 3.700439718 | 4.64385619 | -0.943416472 |
| Q9VGW7 | Q9VGW7_DROME Arginine methyltransferase 1 OS=Drosophila melanogaster OX=7227 GN=Art1 PE=1 SV=1 | 13 | 10 | 24 | 13 | 3.700439718 | 4.584962501 | -0.884522783 |
| P54397 | FKB39_DROME 39 kDa FK506-binding nuclear protein OS=Drosophila melanogaster OX=7227 GN=FK506-bp1 PE=1 SV=2 | 13 | 9 | 23 | 11 | 3.700439718 | 4.523561956 | -0.823122238 |
| P62152 | CALM_DROME Calmodulin OS=Drosophila melanogaster OX=7227 GN=Cam PE=1 SV=2 | 13 | 8 | 23 | 10 | 3.700439718 | 4.523561956 | -0.823122238 |
| P25843 | PROF_DROME Profilin OS=Drosophila melanogaster OX=7227 GN=chic PE=1 SV=1 | 13 | 6 | 22 | 5 | 3.700439718 | 4.459431619 | -0.7589919 |
| Q9VKX2 | Q9VKX2_DROME Malate dehydrogenase OS=Drosophila melanogaster OX=7227 GN=Mdh1 PE=1 SV=2 | 13 | 8 | 21 | 8 | 3.700439718 | 4.392317423 | -0.691877705 |
| Q9VAC4 | Q9VAC4_DROME GEO09167p1 OS=Drosophila melanogaster OX=7227 GN=Nph PE=1 SV=1 | 13 | 7 | 18 | 7 | 3.700439718 | 4.169925001 | -0.469485283 |
| Q8T6I0 | Q8T6I0_DROME EH domain containing protein OS=Drosophila melanogaster OX=7227 GN=Past1 PE=1 SV=1 | 13 | 7 | 17 | 15 | 3.700439718 | 4.087462841 | -0.387023123 |
| Q8INW9 | Q8INW9_DROME Fondue, isoform A OS=Drosophila melanogaster OX=7227 GN=fon PE=1 SV=1 | 13 | 5 | 14 | 6 | 3.700439718 | 3.807354922 | -0.106915204 |
| P20353 | GNAI_DROME G protein alpha i subunit OS=Drosophila melanogaster OX=7227 GN=Galphai PE=1 SV=2 | 13 | 9 | 3 | 3 | 3.700439718 | 1.584962501 | 2.115477217 |
| Q24407 | ATP5J_DROME ATP synthase-coupling factor 6, mitochondrial OS=Drosophila melanogaster OX=7227 GN=ATPsynCf6 PE=3 SV=1 | 13 | 4 | 3 | 2 | 3.700439718 | 1.584962501 | 2.115477217 |
| Q9VZE7 | CTLH1_DROME CTL-like protein 1 OS=Drosophila melanogaster OX=7227 GN=CG1311 PE=2 SV=1 | 13 | 8 | 1 | 1 | 3.700439718 | 0 | 3.700439718 |
| A0A0B7P9G0 | A0A0B7P9G0_DROME Unextended, isoform E OS=Drosophila melanogaster OX=7227 GN=uex PE=1 SV=1 | 13 | 11 | 1 | 1 | 3.700439718 | 0 | 3.700439718 |
| Q7JZK1 | Q7JZK1_DROME NADH dehydrogenase (Ubiquinone) B14 subunit, isoform A OS=Drosophila melanogaster OX=7227 GN=ND-B14 PE=1 SV=1 | 13 | 6 | 3 | 1 | 3.700439718 | 1.584962501 | 2.115477217 |
| Q9VAD0 | Q9VAD0_DROME Acid phosphatase 1, isoform A OS=Drosophila melanogaster OX=7227 GN=Acph-1 PE=1 SV=1 | 13 | 7 | 4 | 3 | 3.700439718 | 2 | 1.700439718 |
| Q9VBU5 | Q9VBU5_DROME LD07939p OS=Drosophila melanogaster OX=7227 GN=Dmel\CG10425 PE=1 SV=1 | 13 | 7 | 2 | 2 | 3.700439718 | 1 | 2.700439718 |
| Q9VBV8 | Q9VBV8_DROME Mannosyltransferase OS=Drosophila melanogaster OX=7227 GN=Dmel\CG11851 PE=2 SV=1 | 13 | 8 | 3 | 3 | 3.700439718 | 1.584962501 | 2.115477217 |
| Q9VJ59 | Q9VJ59_DROME PRA1 family protein OS=Drosophila melanogaster OX=7227 GN=Jwa PE=1 SV=5 | 13 | 7 | 4 | 3 | 3.700439718 | 2 | 1.700439718 |
| Q9VMS1 | Q9VMS1_DROME Cyclope, isoform A OS=Drosophila melanogaster OX=7227 GN=cype PE=1 SV=3 | 13 | 3 | 5 | 2 | 3.700439718 | 2.321928095 | 1.378511623 |
| Q9VAA6 | Q9VAA6_DROME GEO12235p1 OS=Drosophila melanogaster OX=7227 GN=Dmel\CG7950 PE=2 SV=3 | 13 | 4 | 1 | 1 | 3.700439718 | 0 | 3.700439718 |
| Q9VD55 | Q9VD55_DROME Cytochrome c heme lyase OS=Drosophila melanogaster OX=7227 GN=Cchl PE=1 SV=1 | 13 | 7 | 1 | 1 | 3.700439718 | 0 | 3.700439718 |
| Q9VJX8 | Q9VJX8_DROME Very-long-chain (3R)-3-hydroxyacyl-CoA dehydratase OS=Drosophila melanogaster OX=7227 GN=Hacd2 PE=1 SV=1 | 13 | 5 | 1 | 1 | 3.700439718 | 0 | 3.700439718 |
| Q9W423 | Q9W423_DROME LD18904p OS=Drosophila melanogaster OX=7227 GN=schlank PE=1 SV=2 | 13 | 5 | 1 | 1 | 3.700439718 | 0 | 3.700439718 |
| P53624 | MA1A1_DROME Mannosyl-oligosaccharide alpha-1,2-mannosidase IA OS=Drosophila melanogaster OX=7227 GN=alpha-Man-Ia PE=2 SV=2 | 13 | 8 | 1 | 1 | 3.700439718 | 0 | 3.700439718 |
| Q7K4W1 | CDKAL_DROME Threonylcarbamoyladenosine tRNA methylthiotransferase OS=Drosophila melanogaster OX=7227 GN=CG6550 PE=2 SV=1 | 13 | 11 | 1 | 1 | 3.700439718 | 0 | 3.700439718 |
| Q9Y140 | DHRS7_DROME Dehydrogenase/reductase SDR family protein 7-like OS=Drosophila melanogaster OX=7227 GN=CG7601 PE=2 SV=1 | 13 | 6 | 1 | 1 | 3.700439718 | 0 | 3.700439718 |
| A0A0B4KHS1 | A0A0B4KHS1_DROME Crumbs, isoform D OS=Drosophila melanogaster OX=7227 GN=crb PE=4 SV=1 | 13 | 12 | 1 | 1 | 3.700439718 | 0 | 3.700439718 |
| M9PBY3 | M9PBY3_DROME Uncharacterized protein, isoform E OS=Drosophila melanogaster OX=7227 GN=CG43783 PE=1 SV=1 | 13 | 11 | 1 | 1 | 3.700439718 | 0 | 3.700439718 |
| M9PBY4 | M9PBY4_DROME NADH dehydrogenase (Ubiquinone) 15 kDa subunit, isoform H OS=Drosophila melanogaster OX=7227 GN=ND-15 PE=4 SV=1 | 13 | 5 | 1 | 1 | 3.700439718 | 0 | 3.700439718 |
| O76268 | O76268_DROME LD11641p OS=Drosophila melanogaster OX=7227 GN=NTPase PE=1 SV=1 | 13 | 7 | 1 | 1 | 3.700439718 | 0 | 3.700439718 |
| Q0E8Q7 | Q0E8Q7_DROME Uncharacterized protein, isoform B OS=Drosophila melanogaster OX=7227 GN=Eato PE=1 SV=1 | 13 | 12 | 1 | 1 | 3.700439718 | 0 | 3.700439718 |
| Q1RL12 | Q1RL12_DROME IP16413p OS=Drosophila melanogaster OX=7227 GN=nrv2 PE=1 SV=1 | 13 | 11 | 1 | 1 | 3.700439718 | 0 | 3.700439718 |
| Q8IMC2 | Q8IMC2_DROME Eph receptor tyrosine kinase, isoform B OS=Drosophila melanogaster OX=7227 GN=Eph PE=3 SV=2 | 13 | 10 | 1 | 1 | 3.700439718 | 0 | 3.700439718 |

| UniProt ID | Protein description | # PSMs MVB spectra | # of unique peptides MVB | # of PSMs lysate | # of unique peptides lysate | log2(PSM MVB) | log2(PSM lysate) | FC (log2 of MVB PSM-log2 of lysate PSM) |
|---|---|---|---|---|---|---|---|---|
| Q8MYW9 | Q8MYW9_DROME O-acyltransferase OS=Drosophila melanogaster OX=7227 GN=Dmel\CG8112 PE=2 SV=1 | 13 | 7 | 1 | 1 | 3.700439718 | 0 | 3.700439718 |
| Q95TS8 | Q95TS8_DROME LD25288p OS=Drosophila melanogaster OX=7227 GN=Dmel\CG10880 PE=2 SV=1 | 13 | 7 | 1 | 1 | 3.700439718 | 0 | 3.700439718 |
| Q9VDK9 | Q9VDK9_DROME GH12359p OS=Drosophila melanogaster OX=7227 GN=Dmel\CG16953 PE=1 SV=3 | 13 | 6 | 1 | 1 | 3.700439718 | 0 | 3.700439718 |
| Q9VKC9 | Q9VKC9_DROME Dietary and metabolic glutamate transporter OS=Drosophila melanogaster OX=7227 GN=dmGlut PE=2 SV=2 | 13 | 7 | 1 | 1 | 3.700439718 | 0 | 3.700439718 |
| Q9VUM3 | Q9VUM3_DROME GH25868p OS=Drosophila melanogaster OX=7227 GN=Dmel\CG7011 PE=1 SV=1 | 13 | 9 | 1 | 1 | 3.700439718 | 0 | 3.700439718 |
| Q9VZL1 | Q9VZL1_DROME LP07226p OS=Drosophila melanogaster OX=7227 GN=mge PE=1 SV=1 | 13 | 5 | 1 | 1 | 3.700439718 | 0 | 3.700439718 |
| Q9W3M6 | Q9W3M6_DROME Alkyl transferase OS=Drosophila melanogaster OX=7227 GN=Dmel\CG10778 PE=1 SV=2 | 13 | 8 | 1 | 1 | 3.700439718 | 0 | 3.700439718 |
| Q9W4G1 | Q9W4G1_DROME LD07278p OS=Drosophila melanogaster OX=7227 GN=CT21430 PE=2 SV=1 | 13 | 9 | 1 | 1 | 3.700439718 | 0 | 3.700439718 |
| Q9VEZ3 | MSPS_DROME Protein mini spindles OS=Drosophila melanogaster OX=7227 GN=msps PE=1 SV=3 | 12 | 12 | 50 | 33 | 3.584962501 | 5.64385619 | -2.058893689 |
| Q9W237 | RS16_DROME 40S ribosomal protein S16 OS=Drosophila melanogaster OX=7227 GN=RpS16 PE=1 SV=1 | 12 | 7 | 40 | 11 | 3.584962501 | 5.321928095 | -1.736965594 |
| X2JAI4 | X2JAI4_DROME DNA helicase OS=Drosophila melanogaster OX=7227 GN=Mcm3 PE=1 SV=1 | 12 | 8 | 38 | 17 | 3.584962501 | 5.247927513 | -1.662965013 |
| Q9XZJ4 | PSA6_DROME Proteasome subunit alpha type-6 OS=Drosophila melanogaster OX=7227 GN=Prosalpha1 PE=1 SV=2 | 12 | 8 | 34 | 13 | 3.584962501 | 5.087462841 | -1.502500341 |
| P35421 | PUR4_DROME Phosphoribosylformylglycinamidine synthase OS=Drosophila melanogaster OX=7227 GN=ade2 PE=2 SV=2 | 12 | 9 | 33 | 19 | 3.584962501 | 5.044394119 | -1.459431619 |
| Q9VZS5 | RL28_DROME 60S ribosomal protein L28 OS=Drosophila melanogaster OX=7227 GN=RpL28 PE=1 SV=1 | 12 | 10 | 33 | 11 | 3.584962501 | 5.044394119 | -1.459431619 |
| O97428 | O97428_DROME Ciboulot, isoform A OS=Drosophila melanogaster OX=7227 GN=cib PE=1 SV=1 | 12 | 5 | 27 | 8 | 3.584962501 | 4.754887502 | -1.169925001 |
| Q9W1B9 | Q9W1B9_DROME GEO07602p1 OS=Drosophila melanogaster OX=7227 GN=RpL12 PE=1 SV=1 | 12 | 4 | 27 | 6 | 3.584962501 | 4.754887502 | -1.169925001 |
| Q9VRL1 | Q9VRL1_DROME GEO06356p1 OS=Drosophila melanogaster OX=7227 GN=Uev1A PE=1 SV=1 | 12 | 5 | 26 | 7 | 3.584962501 | 4.700439718 | -1.115477217 |
| Q95083 | PSA5_DROME Proteasome subunit alpha type-5 OS=Drosophila melanogaster OX=7227 GN=Prosalpha5 PE=2 SV=2 | 12 | 8 | 25 | 13 | 3.584962501 | 4.64385619 | -1.058893689 |
| A4UZL5 | A4UZL5_DROME Glutathione S transferase S1, isoform C OS=Drosophila melanogaster OX=7227 GN=GstS1 PE=1 SV=1 | 12 | 8 | 24 | 9 | 3.584962501 | 4.584962501 | -1 |
| Q9VNE2 | PKRA_DROME Protein krasavietz OS=Drosophila melanogaster OX=7227 GN=kra PE=1 SV=1 | 12 | 9 | 23 | 12 | 3.584962501 | 4.523561956 | -0.938599455 |
| Q27889 | PP2B2_DROME Serine/threonine-protein phosphatase 2B catalytic subunit 2 OS=Drosophila melanogaster OX=7227 GN=Pp2B-14D PE=1 SV=2 | 12 | 9 | 20 | 11 | 3.584962501 | 4.321928095 | -0.736965594 |
| P04359 | RL32_DROME 60S ribosomal protein L32 OS=Drosophila melanogaster OX=7227 GN=RpL32 PE=1 SV=3 | 12 | 5 | 18 | 9 | 3.584962501 | 4.169925001 | -0.584962501 |
| Q9VD66 | Q9VD66_DROME Female-specific independent of transformer OS=Drosophila melanogaster OX=7227 GN=fit PE=1 SV=1 | 12 | 6 | 18 | 9 | 3.584962501 | 4.169925001 | -0.584962501 |
| Q9VW59 | Q9VW59_DROME LD16419p OS=Drosophila melanogaster OX=7227 GN=RhoGDI PE=1 SV=1 | 12 | 7 | 18 | 8 | 3.584962501 | 4.169925001 | -0.584962501 |
| O17445 | RL15_DROME 60S ribosomal protein L15 OS=Drosophila melanogaster OX=7227 GN=RpL15 PE=1 SV=1 | 12 | 7 | 17 | 7 | 3.584962501 | 4.087462841 | -0.502500341 |
| P29327 | RS6_DROME 40S ribosomal protein S6 OS=Drosophila melanogaster OX=7227 GN=RpS6 PE=1 SV=1 | 12 | 6 | 17 | 5 | 3.584962501 | 4.087462841 | -0.502500341 |
| Q9VBH8 | Q9VBH8_DROME FI02850p OS=Drosophila melanogaster OX=7227 GN=RpL34a PE=1 SV=2 | 12 | 4 | 17 | 4 | 3.584962501 | 4.087462841 | -0.502500341 |
| Q8MSC4 | Q8MSC4_DROME CG3509 protein OS=Drosophila melanogaster OX=7227 GN=BigH1 PE=1 SV=1 | 12 | 6 | 16 | 8 | 3.584962501 | 4 | -0.415037499 |
| P08646 | RAS1_DROME Ras-like protein 1 OS=Drosophila melanogaster OX=7227 GN=Ras85D PE=1 SV=2 | 12 | 5 | 2 | 2 | 3.584962501 | 1 | 2.584962501 |
| Q9VD26 | ZFPL1_DROME Zinc finger protein-like 1 homolog OS=Drosophila melanogaster OX=7227 GN=CG5382 PE=1 SV=1 | 12 | 8 | 1 | 1 | 3.584962501 | 0 | 3.584962501 |
| Q9VMV6 | SELT_DROME Thioredoxin reductase-like selenoprotein T homolog CG3887 OS=Drosophila melanogaster OX=7227 GN=CG3887 PE=2 SV=1 | 12 | 4 | 1 | 1 | 3.584962501 | 0 | 3.584962501 |
| Q9VSU7 | USE1_DROME Vesicle transport protein USE1 OS=Drosophila melanogaster OX=7227 GN=Use1 PE=2 SV=3 | 12 | 5 | 1 | 1 | 3.584962501 | 0 | 3.584962501 |
| Q7JVK6 | Q7JVK6_DROME GM27569p OS=Drosophila melanogaster OX=7227 GN=trsn PE=1 SV=1 | 12 | 5 | 11 | 3 | 3.584962501 | 3.459431619 | 0.125530882 |
| Q8IQA1 | Q8IQA1_DROME Bax Inhibitor-1, isoform B OS=Drosophila melanogaster OX=7227 GN=BI-1 PE=3 SV=1 | 12 | 7 | 1 | 1 | 3.584962501 | 0 | 3.584962501 |
| Q9I7L9 | Q9I7L9_DROME Uncharacterized protein, isoform A OS=Drosophila melanogaster OX=7227 GN=38C.53 PE=4 SV=2 | 12 | 8 | 1 | 1 | 3.584962501 | 0 | 3.584962501 |
| Q9VPT9 | Q9VPT9_DROME LD27268p OS=Drosophila melanogaster OX=7227 GN=Dmel\CG3662 PE=1 SV=1 | 12 | 7 | 1 | 1 | 3.584962501 | 0 | 3.584962501 |
| Q9VWL4 | Q9VWL4_DROME GH07340p OS=Drosophila melanogaster OX=7227 GN=Dmel\CG7556 PE=1 SV=1 | 12 | 7 | 1 | 1 | 3.584962501 | 0 | 3.584962501 |
| Q9W1N3 | Q9W1N3_DROME Levy, isoform A OS=Drosophila melanogaster OX=7227 GN=levy PE=1 SV=1 | 12 | 6 | 1 | 1 | 3.584962501 | 0 | 3.584962501 |
| O97172 | U729_DROME UPF0729 protein CG18508 OS=Drosophila melanogaster OX=7227 GN=CG18508 PE=1 SV=1 | 12 | 4 | 1 | 1 | 3.584962501 | 0 | 3.584962501 |
| Q6WV20 | GALT1_DROME Polypeptide N-acetylgalactosaminyltransferase 1 OS=Drosophila melanogaster OX=7227 GN=GalNAc-T1 PE=2 SV=2 | 12 | 6 | 1 | 1 | 3.584962501 | 0 | 3.584962501 |
| Q9VDD7 | S35B1_DROME Solute carrier family 35 member B1 homolog OS=Drosophila melanogaster OX=7227 GN=meigo PE=2 SV=1 | 12 | 6 | 1 | 1 | 3.584962501 | 0 | 3.584962501 |
| A0A0B4KFG0 | A0A0B4KFG0_DROME Alpha-mannosidase OS=Drosophila melanogaster OX=7227 GN=alpha-Man-IIa PE=3 SV=1 | 12 | 11 | 1 | 1 | 3.584962501 | 0 | 3.584962501 |
| A0A0B4KHV0 | A0A0B4KHV0_DROME Furin 1, isoform J OS=Drosophila melanogaster OX=7227 GN=Fur1 PE=3 SV=1 | 12 | 9 | 1 | 1 | 3.584962501 | 0 | 3.584962501 |
| A0A0B4LGL2 | A0A0B4LGL2_DROME Sphingomyelin phosphodiesterase OS=Drosophila melanogaster OX=7227 GN=CG3376 PE=1 SV=1 | 12 | 8 | 1 | 1 | 3.584962501 | 0 | 3.584962501 |
| E1JI19 | E1JI19_DROME I'm not dead yet, isoform D OS=Drosophila melanogaster OX=7227 GN=Indy PE=1 SV=1 | 12 | 6 | 1 | 1 | 3.584962501 | 0 | 3.584962501 |
| Q7JWI7 | Q7JWI7_DROME RE60337p OS=Drosophila melanogaster OX=7227 GN=CG11665 PE=1 SV=1 | 12 | 3 | 1 | 1 | 3.584962501 | 0 | 3.584962501 |
| Q8I935 | Q8I935_DROME Uncharacterized protein, isoform D OS=Drosophila melanogaster OX=7227 GN=CG33523 PE=1 SV=2 | 12 | 6 | 1 | 1 | 3.584962501 | 0 | 3.584962501 |
| Q8INX7 | Q8INX7_DROME Fasciclin 3, isoform C OS=Drosophila melanogaster OX=7227 GN=Fas3 PE=4 SV=2 | 12 | 9 | 1 | 1 | 3.584962501 | 0 | 3.584962501 |
| Q961R9 | Q961R9_DROME GH09241p OS=Drosophila melanogaster OX=7227 GN=CT19169 PE=1 SV=1 | 12 | 7 | 1 | 1 | 3.584962501 | 0 | 3.584962501 |
| Q9V3F9 | Q9V3F9_DROME BcDNA.LD28120 OS=Drosophila melanogaster OX=7227 GN=out PE=2 SV=1 | 12 | 7 | 1 | 1 | 3.584962501 | 0 | 3.584962501 |
| Q9W3F3 | Q9W3F3_DROME RH03540p OS=Drosophila melanogaster OX=7227 GN=Dmel\CG15347 PE=1 SV=2 | 12 | 7 | 1 | 1 | 3.584962501 | 0 | 3.584962501 |
| A0A0B4LFL2 | A0A0B4LFL2_DROME Eukaryotic translation initiation factor 3 subunit C OS=Drosophila melanogaster OX=7227 GN=eIF3c PE=3 SV=1 | 11 | 11 | 62 | 25 | 3.459431619 | 5.95419631 | -2.494764692 |
| Q9V461 | MCM6_DROME DNA replication licensing factor Mcm6 OS=Drosophila melanogaster OX=7227 GN=Mcm6 PE=1 SV=1 | 11 | 9 | 47 | 24 | 3.459431619 | 5.554588852 | -2.095157233 |
| P48598 | IF4E_DROME Eukaryotic translation initiation factor 4E1 OS=Drosophila melanogaster OX=7227 GN=eIF4E1 PE=1 SV=1 | 11 | 6 | 40 | 10 | 3.459431619 | 5.321928095 | -1.862496476 |
| Q8IMI7 | Q8IMI7_DROME 40S ribosomal protein S7 OS=Drosophila melanogaster OX=7227 GN=RpS7 PE=1 SV=2 | 11 | 7 | 39 | 14 | 3.459431619 | 5.285402219 | -1.8259706 |
| O97477 | INO1_DROME Inositol-3-phosphate synthase OS=Drosophila melanogaster OX=7227 GN=Inos PE=1 SV=1 | 11 | 7 | 37 | 16 | 3.459431619 | 5.209453366 | -1.750021747 |
| P41092 | RL27A_DROME 60S ribosomal protein L27a OS=Drosophila melanogaster OX=7227 GN=RpL27A PE=1 SV=2 | 11 | 7 | 35 | 9 | 3.459431619 | 5.129283017 | -1.669851398 |
| Q9VAN0 | SERC_DROME Probable phosphoserine aminotransferase OS=Drosophila melanogaster OX=7227 GN=CG11899 PE=2 SV=1 | 11 | 7 | 35 | 16 | 3.459431619 | 5.129283017 | -1.669851398 |
| A0A0B4LIJ0 | A0A0B4LIJ0_DROME Regulatory particle triple-A ATPase 2, isoform B OS=Drosophila melanogaster OX=7227 GN=Rpt2 PE=3 SV=1 | 11 | 8 | 35 | 16 | 3.459431619 | 5.129283017 | -1.669851398 |
| Q03334 | RS13_DROME 40S ribosomal protein S13 OS=Drosophila melanogaster OX=7227 GN=RpS13 PE=1 SV=3 | 11 | 6 | 30 | 7 | 3.459431619 | 4.906890596 | -1.447458977 |
| A0A126GUR6 | A0A126GUR6_DROME Succinate--CoA ligase [ADP-forming] subunit beta, mitochondrial OS=Drosophila melanogaster OX=7227 GN=ScsbetaA PE=1 SV=1 | 11 | 7 | 30 | 12 | 3.459431619 | 4.906890596 | -1.447458977 |
| Q9XZ61 | Q9XZ61_DROME Ubiquitin carboxyl-terminal hydrolase OS=Drosophila melanogaster OX=7227 GN=Uch-L5 PE=1 SV=1 | 11 | 9 | 30 | 14 | 3.459431619 | 4.906890596 | -1.447458977 |
| P07486 | G3P1_DROME Glyceraldehyde-3-phosphate dehydrogenase 1 OS=Drosophila melanogaster OX=7227 GN=Gapdh1 PE=2 SV=2 | 11 | 4 | 28 | 5 | 3.459431619 | 4.807354922 | -1.347923303 |
| Q9VNA5 | PSB4_DROME Proteasome subunit beta type-4 OS=Drosophila melanogaster OX=7227 GN=Prosbeta7 PE=2 SV=1 | 11 | 7 | 26 | 9 | 3.459431619 | 4.700439718 | -1.2410081 |
| A0A0C4FEI8 | A0A0C4FEI8_DROME Granny smith, isoform F OS=Drosophila melanogaster OX=7227 GN=grsm PE=1 SV=1 | 11 | 10 | 25 | 9 | 3.459431619 | 4.64385619 | -1.184424571 |
| Q9VBP9 | NPL4_DROME Nuclear protein localization protein 4 homolog OS=Drosophila melanogaster OX=7227 GN=Npl4 PE=1 SV=3 | 11 | 8 | 22 | 13 | 3.459431619 | 4.459431619 | -1 |
| Q0KHZ6 | Q0KHZ6_DROME Uncharacterized protein, isoform A OS=Drosophila melanogaster OX=7227 GN=CG 7834 PE=1 SV=1 | 11 | 9 | 21 | 11 | 3.459431619 | 4.392317423 | -0.932885804 |
| P53997 | SET_DROME Protein SET OS=Drosophila melanogaster OX=7227 GN=Set PE=1 SV=2 | 11 | 6 | 18 | 7 | 3.459431619 | 4.169925001 | -0.710493383 |
| Q7JXC4 | Q7JXC4_DROME CG6459 protein OS=Drosophila melanogaster OX=7227 GN=P32 PE=1 SV=1 | 11 | 6 | 16 | 6 | 3.459431619 | 4 | -0.540568381 |
| Q8IMK7 | Q8IMK7_DROME Alphabet, isoform B OS=Drosophila melanogaster OX=7227 GN=alph PE=1 SV=1 | 11 | 8 | 16 | 7 | 3.459431619 | 4 | -0.540568381 |
| Q9W1L1 | Q9W1L1_DROME Mitochondrial ribosomal protein L43 OS=Drosophila melanogaster OX=7227 GN=mRpL43 PE=1 SV=1 | 11 | 4 | 15 | 6 | 3.459431619 | 3.906890596 | -0.447458977 |
| M9PG40 | M9PG40_DROME Uncharacterized protein, isoform D OS=Drosophila melanogaster OX=7227 GN=CG40045 PE=1 SV=1 | 11 | 7 | 14 | 8 | 3.459431619 | 3.807354922 | -0.347923303 |
| P23380 | VATL_DROME V-type proton ATPase 16 kDa proteolipid subunit OS=Drosophila melanogaster OX=7227 GN=Vha16-1 PE=2 SV=1 | 11 | 6 | 1 | 1 | 3.459431619 | 0 | 3.459431619 |
| Q24157 | BRN_DROME Beta-1,3-galactosyltransferase brn OS=Drosophila melanogaster OX=7227 GN=brn PE=2 SV=2 | 11 | 6 | 1 | 1 | 3.459431619 | 0 | 3.459431619 |
| Q8IQ70 | MCU_DROME Calcium uniporter protein, mitochondrial OS=Drosophila melanogaster OX=7227 GN=CG18769 PE=2 SV=2 | 11 | 7 | 3 | 2 | 3.459431619 | 1.584962501 | 1.874469118 |
| Q95SK3 | CK5P3_DROME CDK5RAP3-like protein OS=Drosophila melanogaster OX=7227 GN=CG30291 PE=2 SV=1 | 11 | 8 | 4 | 3 | 3.459431619 | 2 | 1.459431619 |
| Q9V3W0 | UK114_DROME RutC family protein UK114 OS=Drosophila melanogaster OX=7227 GN=UK114 PE=2 SV=1 | 11 | 7 | 8 | 5 | 3.459431619 | 3 | 0.459431619 |
| Q9VMX0 | RM28_DROME 39S ribosomal protein L28, mitochondrial OS=Drosophila melanogaster OX=7227 GN=mRpL28 PE=2 SV=1 | 11 | 7 | 10 | 6 | 3.459431619 | 3.321928095 | 0.137503524 |
| Q9VVH3 | MIC13_DROME MICOS complex subunit MIC13 homolog QIL1 OS=Drosophila melanogaster OX=7227 GN=QIL1 PE=2 SV=1 | 11 | 6 | 1 | 1 | 3.459431619 | 0 | 3.459431619 |
| M9PFY2 | M9PFY2_DROME Uncharacterized protein, isoform C OS=Drosophila melanogaster OX=7227 GN=CG6479 PE=1 SV=1 | 11 | 7 | 2 | 2 | 3.459431619 | 1 | 2.459431619 |
| Q7PLT4 | Q7PLT4_DROME Translocase of the inner mitochondrial membrane 17b, isoform A OS=Drosophila melanogaster OX=7227 GN=Tim17b PE=1 SV=1 | 11 | 4 | 2 | 1 | 3.459431619 | 1 | 2.459431619 |

| UniProt ID | Protein description | # PSMs MVB spectra | # of unique peptides MVB | # of PSMs lysate | # of unique peptides lysate | log2(PSM MVB) | log2(PSM lysate) | FC (log2 of MVB PSM-log2 of lysate PSM) |
|---|---|---|---|---|---|---|---|---|
| Q9V9S0 | Q9V9S0_DROME FI04419p OS=Drosophila melanogaster OX=7227 GN=Lamp1 PE=1 SV=3 | 11 | 5 | 3 | 2 | 3.459431619 | 1.584962501 | 1.874469118 |
| Q9VX83 | Q9VX83_DROME Amino acid transmembrane transporter OS=Drosophila melanogaster OX=7227 GN=CT15971 PE=2 SV=2 | 11 | 6 | 2 | 2 | 3.459431619 | 1 | 2.459431619 |
| X2J9U5 | X2J9U5_DROME Non-lysosomal glucosylceramidase OS=Drosophila melanogaster OX=7227 GN=CG33090 PE=3 SV=1 | 11 | 9 | 2 | 2 | 3.459431619 | 1 | 2.459431619 |
| Q9W2U8 | Q9W2U8_DROME Neb-cGP, isoform A OS=Drosophila melanogaster OX=7227 GN=Neb-cGP PE=1 SV=1 | 11 | 4 | 1 | 1 | 3.459431619 | 0 | 3.459431619 |
| Q9Y171 | Q9Y171_DROME BcDNA.GH02220 OS=Drosophila melanogaster OX=7227 GN=BcDNA.GH02220 PE=2 SV=1 | 11 | 7 | 1 | 1 | 3.459431619 | 0 | 3.459431619 |
| P07207 | NOTCH_DROME Neurogenic locus Notch protein OS=Drosophila melanogaster OX=7227 GN=N PE=1 SV=3 | 11 | 9 | 1 | 1 | 3.459431619 | 0 | 3.459431619 |
| Q0KHY3 | MESH_DROME Protein mesh OS=Drosophila melanogaster OX=7227 GN=mesh PE=1 SV=2 | 11 | 7 | 1 | 1 | 3.459431619 | 0 | 3.459431619 |
| Q9VTK2 | POMT1_DROME Protein O-mannosyltransferase 1 OS=Drosophila melanogaster OX=7227 GN=rt PE=2 SV=2 | 11 | 9 | 1 | 1 | 3.459431619 | 0 | 3.459431619 |
| A1Z729 | A1Z729_DROME CG2064 OS=Drosophila melanogaster OX=7227 GN=CG2064 PE=1 SV=1 | 11 | 10 | 1 | 1 | 3.459431619 | 0 | 3.459431619 |
| H1UUB1 | H1UUB1_DROME GEO12465p1 OS=Drosophila melanogaster OX=7227 GN=Syb PE=1 SV=1 | 11 | 4 | 1 | 1 | 3.459431619 | 0 | 3.459431619 |
| M9NEV7 | M9NEV7_DROME Uncharacterized protein, isoform B OS=Drosophila melanogaster OX=7227 GN=CG9336 PE=1 SV=1 | 11 | 4 | 1 | 1 | 3.459431619 | 0 | 3.459431619 |
| M9PH44 | M9PH44_DROME Prominin-like, isoform E OS=Drosophila melanogaster OX=7227 GN=prominin-like PE=1 SV=1 | 11 | 9 | 1 | 1 | 3.459431619 | 0 | 3.459431619 |
| O97418 | O97418_DROME EG:152A3.7 protein OS=Drosophila melanogaster OX=7227 GN=ND-B14.5A PE=1 SV=1 | 11 | 4 | 1 | 1 | 3.459431619 | 0 | 3.459431619 |
| Q8I930 | Q8I930_DROME GH14147p OS=Drosophila melanogaster OX=7227 GN=hoe1 PE=1 SV=1 | 11 | 8 | 1 | 1 | 3.459431619 | 0 | 3.459431619 |
| Q8MRN5 | Q8MRN5_DROME Aminopeptidase OS=Drosophila melanogaster OX=7227 GN=CG5839 PE=1 SV=1 | 11 | 10 | 1 | 1 | 3.459431619 | 0 | 3.459431619 |
| Q960U8 | Q960U8_DROME O-acyltransferase OS=Drosophila melanogaster OX=7227 GN=mdy PE=2 SV=1 | 11 | 5 | 1 | 1 | 3.459431619 | 0 | 3.459431619 |
| Q962I2 | Q962I2_DROME RAL2 OS=Drosophila melanogaster OX=7227 GN=Rala PE=1 SV=1 | 11 | 6 | 1 | 1 | 3.459431619 | 0 | 3.459431619 |
| Q9VAQ8 | Q9VAQ8_DROME Beta4GalNAcTB OS=Drosophila melanogaster OX=7227 GN=beta4GalNAcTB PE=2 SV=1 | 11 | 6 | 1 | 1 | 3.459431619 | 0 | 3.459431619 |
| Q9VH02 | Q9VH02_DROME LD30822p OS=Drosophila melanogaster OX=7227 GN=Dmel\CG6293 PE=1 SV=1 | 11 | 5 | 1 | 1 | 3.459431619 | 0 | 3.459431619 |
| Q9VHG4 | Q9VHG4_DROME LD30634p OS=Drosophila melanogaster OX=7227 GN=ATP6AP2 PE=1 SV=1 | 11 | 4 | 1 | 1 | 3.459431619 | 0 | 3.459431619 |
| Q9VIG0 | Q9VIG0_DROME Uncharacterized protein OS=Drosophila melanogaster OX=7227 GN=CG9257 PE=1 SV=1 | 11 | 7 | 1 | 1 | 3.459431619 | 0 | 3.459431619 |
| Q9VNQ3 | Q9VNQ3_DROME ER membrane protein complex subunit 4 OS=Drosophila melanogaster OX=7227 GN=BcDNA:LD31024 PE=1 SV=1 | 11 | 6 | 1 | 1 | 3.459431619 | 0 | 3.459431619 |
| Q9VP06 | Q9VP06_DROME HL02815p OS=Drosophila melanogaster OX=7227 GN=Alg11 PE=1 SV=2 | 11 | 8 | 1 | 1 | 3.459431619 | 0 | 3.459431619 |
| Q9W4A0 | Q9W4A0_DROME GH11193p OS=Drosophila melanogaster OX=7227 GN=Rab18 PE=1 SV=1 | 11 | 8 | 1 | 1 | 3.459431619 | 0 | 3.459431619 |
| Q9W4P0 | Q9W4P0_DROME LD22456p OS=Drosophila melanogaster OX=7227 GN=Dmel\CG2938 PE=2 SV=1 | 11 | 8 | 1 | 1 | 3.459431619 | 0 | 3.459431619 |
| Q9W4W8 | Q9W4W8_DROME Uncharacterized protein, isoform A OS=Drosophila melanogaster OX=7227 GN=Dm100G10.aj PE=1 SV=1 | 11 | 8 | 1 | 1 | 3.459431619 | 0 | 3.459431619 |
| P20477 | GLNA1_DROME Glutamine synthetase 1, mitochondrial OS=Drosophila melanogaster OX=7227 GN=Gs1 PE=2 SV=3 | 10 | 6 | 45 | 17 | 3.321928095 | 5.491853096 | -2.169925001 |
| Q07152 | IMDH_DROME Inosine-5'-monophosphate dehydrogenase OS=Drosophila melanogaster OX=7227 GN=ras PE=1 SV=1 | 10 | 8 | 43 | 19 | 3.321928095 | 5.426264755 | -2.10433666 |
| M9PB90 | M9PB90_DROME Cullin-associated and neddylation-dissociated 1, isoform B OS=Drosophila melanogaster OX=7227 GN=Cand1 PE=1 SV=1 | 10 | 8 | 37 | 21 | 3.321928095 | 5.209453366 | -1.887525271 |
| Q9V5C6 | PSA3_DROME Proteasome subunit alpha type-3 OS=Drosophila melanogaster OX=7227 GN=Prosalpha7 PE=1 SV=1 | 10 | 7 | 33 | 12 | 3.321928095 | 5.044394119 | -1.722466024 |
| Q27268 | DX39B_DROME ATP-dependent RNA helicase WM6 OS=Drosophila melanogaster OX=7227 GN=Hel25E PE=1 SV=1 | 10 | 9 | 30 | 11 | 3.321928095 | 4.906890596 | -1.584962501 |
| Q9V429 | THIO2_DROME Thioredoxin-2 OS=Drosophila melanogaster OX=7227 GN=Trx-2 PE=1 SV=3 | 10 | 6 | 29 | 4 | 3.321928095 | 4.857980995 | -1.5360529 |
| A0A0B4KGP8 | A0A0B4KGP8_DROME Eukaryotic translation initiation factor 2 subunit gamma, isoform D OS=Drosophila melanogaster OX=7227 GN=eIF2gamma PE=4 SV=1 | 10 | 9 | 29 | 12 | 3.321928095 | 4.857980995 | -1.5360529 |
| M9PFR8 | M9PFR8_DROME Uncharacterized protein, isoform B OS=Drosophila melanogaster OX=7227 GN=CG32164 PE=4 SV=1 | 10 | 9 | 27 | 18 | 3.321928095 | 4.754887502 | -1.432959407 |
| P41572 | 6PGD_DROME 6-phosphogluconate dehydrogenase, decarboxylating OS=Drosophila melanogaster OX=7227 GN=Pgd PE=2 SV=1 | 10 | 8 | 24 | 9 | 3.321928095 | 4.584962501 | -1.263034406 |
| P98159 | NUDEL_DROME Serine protease nudel OS=Drosophila melanogaster OX=7227 GN=ndl PE=1 SV=2 | 10 | 10 | 22 | 14 | 3.321928095 | 4.459431619 | -1.137503524 |
| Q9V405 | Q9V405_DROME 26S proteasome regulatory complex subunit p48A OS=Drosophila melanogaster OX=7227 GN=Rpt3 PE=1 SV=1 | 10 | 8 | 22 | 12 | 3.321928095 | 4.459431619 | -1.137503524 |
| A0A0B4KFH4 | A0A0B4KFH4_DROME Regulatory particle non-ATPase 8, isoform B OS=Drosophila melanogaster OX=7227 GN=Rpn8 PE=4 SV=1 | 10 | 7 | 21 | 9 | 3.321928095 | 4.392317423 | -1.070389328 |
| A1ZB71 | A1ZB71_DROME Glutathione S transferase E6 OS=Drosophila melanogaster OX=7227 GN=GstE6 PE=1 SV=1 | 10 | 8 | 19 | 8 | 3.321928095 | 4.247927513 | -0.925999419 |
| P45437 | COPB_DROME Coatomer subunit beta OS=Drosophila melanogaster OX=7227 GN=betaCOP PE=2 SV=2 | 10 | 8 | 18 | 13 | 3.321928095 | 4.169925001 | -0.847996907 |
| Q05344 | SSRP1_DROME FACT complex subunit Ssrp1 OS=Drosophila melanogaster OX=7227 GN=Ssrp PE=1 SV=2 | 10 | 7 | 18 | 13 | 3.321928095 | 4.169925001 | -0.847996907 |
| Q9VJG0 | XPP_DROME Xaa-Pro aminopeptidase ApepP OS=Drosophila melanogaster OX=7227 GN=ApepP PE=1 SV=1 | 10 | 8 | 16 | 10 | 3.321928095 | 4 | -0.678071905 |
| Q9VXP3 | Q9VXP3_DROME GH05406p OS=Drosophila melanogaster OX=7227 GN=mRpS30 PE=1 SV=1 | 10 | 8 | 15 | 12 | 3.321928095 | 3.906890596 | -0.584962501 |
| P54622 | SSBP_DROME Single-stranded DNA-binding protein, mitochondrial OS=Drosophila melanogaster OX=7227 GN=mtSSB PE=1 SV=2 | 10 | 6 | 12 | 4 | 3.321928095 | 3.584962501 | -0.263034406 |
| A0A0B4KH25 | A0A0B4KH25_DROME Histone H2A OS=Drosophila melanogaster OX=7227 GN=His2Av PE=3 SV=1 | 10 | 4 | 12 | 6 | 3.321928095 | 3.584962501 | -0.263034406 |
| Q8I725 | Q8I725_DROME Uncharacterized protein OS=Drosophila melanogaster OX=7227 GN=CG1236 PE=1 SV=2 | 10 | 8 | 12 | 8 | 3.321928095 | 3.584962501 | -0.263034406 |
| Q8IQZ7 | Q8IQZ7_DROME FI19014p1 OS=Drosophila melanogaster OX=7227 GN=Fim PE=1 SV=1 | 10 | 9 | 12 | 8 | 3.321928095 | 3.584962501 | -0.263034406 |
| P22464 | ANXB9_DROME Annexin B9 OS=Drosophila melanogaster OX=7227 GN=AnxB9 PE=2 SV=2 | 10 | 9 | 11 | 9 | 3.321928095 | 3.459431619 | -0.137503524 |
| P17336 | CATA_DROME Catalase OS=Drosophila melanogaster OX=7227 GN=Cat PE=1 SV=2 | 10 | 7 | 5 | 4 | 3.321928095 | 2.321928095 | 1 |
| P84040 | H4_DROME Histone H4 OS=Drosophila melanogaster OX=7227 GN=His4 PE=1 SV=2 | 10 | 4 | 9 | 5 | 3.321928095 | 3.169925001 | 0.152003093 |
| Q95T12 | FLOWR_DROME Calcium channel flower OS=Drosophila melanogaster OX=7227 GN=fwe PE=1 SV=1 | 10 | 5 | 2 | 1 | 3.321928095 | 1 | 2.321928095 |
| A0A0B4K6K7 | A0A0B4K6K7_DROME UFM1 specific ligase 1, isoform C OS=Drosophila melanogaster OX=7227 GN=Ufl1 PE=4 SV=1 | 10 | 8 | 4 | 3 | 3.321928095 | 2 | 1.321928095 |
| O96692 | O96692_DROME RE63021p OS=Drosophila melanogaster OX=7227 GN=Rap2l PE=1 SV=1 | 10 | 5 | 2 | 2 | 3.321928095 | 1 | 2.321928095 |
| Q0E8X7 | Q0E8X7_DROME Uncharacterized protein, isoform A OS=Drosophila melanogaster OX=7227 GN=roh PE=1 SV=1 | 10 | 4 | 6 | 2 | 3.321928095 | 2.584962501 | 0.736965594 |
| Q500Y7 | Q500Y7_DROME GEO11443p1 OS=Drosophila melanogaster OX=7227 GN=UQCR-6.4 PE=2 SV=1 | 10 | 3 | 2 | 1 | 3.321928095 | 1 | 2.321928095 |
| Q7JZI2 | Q7JZI2_DROME Tetraspanin OS=Drosophila melanogaster OX=7227 GN=Tsp74F PE=2 SV=1 | 10 | 5 | 3 | 3 | 3.321928095 | 1.584962501 | 1.736965594 |
| Q8SZU0 | Q8SZU0_DROME GH13879p OS=Drosophila melanogaster OX=7227 GN=Dmel\CG9331 PE=1 SV=1 | 10 | 7 | 10 | 8 | 3.321928095 | 3.321928095 | 0 |
| Q7K519 | Q7K519_DROME GH16429p OS=Drosophila melanogaster OX=7227 GN=RagC-D PE=1 SV=1 | 10 | 8 | 4 | 4 | 3.321928095 | 2 | 1.321928095 |
| Q7KY04 | Q7KY04_DROME Rab4 protein OS=Drosophila melanogaster OX=7227 GN=Rab4 PE=1 SV=1 | 10 | 7 | 4 | 4 | 3.321928095 | 2 | 1.321928095 |
| Q9VM22 | Q9VM22_DROME LD15196p OS=Drosophila melanogaster OX=7227 GN=Dmel\CG5149 PE=1 SV=1 | 10 | 8 | 3 | 3 | 3.321928095 | 1.584962501 | 1.736965594 |
| Q9VPJ9 | Q9VPJ9_DROME Uncharacterized protein, isoform B OS=Drosophila melanogaster OX=7227 GN=BEST:CK02656 PE=1 SV=2 | 10 | 8 | 3 | 3 | 3.321928095 | 1.584962501 | 1.736965594 |
| Q9W0U6 | Q9W0U6_DROME Uncharacterized protein OS=Drosophila melanogaster OX=7227 GN=CG7049 PE=1 SV=1 | 10 | 8 | 3 | 3 | 3.321928095 | 1.584962501 | 1.736965594 |
| Q9W1H1 | Q9W1H1_DROME CG5339 OS=Drosophila melanogaster OX=7227 GN=CG5339 PE=2 SV=1 | 10 | 8 | 3 | 2 | 3.321928095 | 1.584962501 | 1.736965594 |
| X2JCC5 | X2JCC5_DROME Ankyrin repeat and LEM domain containing 2, isoform G OS=Drosophila melanogaster OX=7227 GN=Ankle2 PE=1 SV=1 | 10 | 8 | 3 | 3 | 3.321928095 | 1.584962501 | 1.736965594 |
| X2JFD6 | X2JFD6_DROME Peroxisomal multifunctional enzyme type 2, isoform B OS=Drosophila melanogaster OX=7227 GN=Mfe2 PE=4 SV=1 | 10 | 10 | 2 | 2 | 3.321928095 | 1 | 2.321928095 |
| Q9V3E5 | Q9V3E5_DROME Tetraspanin OS=Drosophila melanogaster OX=7227 GN=Tsp66E PE=2 SV=1 | 10 | 4 | 1 | 1 | 3.321928095 | 0 | 3.321928095 |
| Q9VBZ8 | Q9VBZ8_DROME Uncharacterized protein OS=Drosophila melanogaster OX=7227 GN=CG11781 PE=1 SV=1 | 10 | 4 | 1 | 1 | 3.321928095 | 0 | 3.321928095 |
| Q9V576 | WUN_DROME Putative phosphatidate phosphatase OS=Drosophila melanogaster OX=7227 GN=wun PE=1 SV=2 | 10 | 4 | 1 | 1 | 3.321928095 | 0 | 3.321928095 |
| Q9VAP3 | CTLH2_DROME CTL-like protein 2 OS=Drosophila melanogaster OX=7227 GN=CG11880 PE=3 SV=1 | 10 | 7 | 1 | 1 | 3.321928095 | 0 | 3.321928095 |
| Q9VED0 | TAPT1_DROME Protein TAPT1 homolog OS=Drosophila melanogaster OX=7227 GN=CG7218 PE=2 SV=2 | 10 | 6 | 1 | 1 | 3.321928095 | 0 | 3.321928095 |
| Q9VHB6 | MTX1_DROME Metaxin-1 homolog OS=Drosophila melanogaster OX=7227 GN=CG9393 PE=1 SV=1 | 10 | 6 | 1 | 1 | 3.321928095 | 0 | 3.321928095 |
| Q9VZS6 | NSMA_DROME Putative neutral sphingomyelinase OS=Drosophila melanogaster OX=7227 GN=CG12034 PE=2 SV=2 | 10 | 5 | 1 | 1 | 3.321928095 | 0 | 3.321928095 |
| A0A0B4LF14 | A0A0B4LF14_DROME Peptidyl-alpha-hydroxyglycine-alpha-amidating lyase 1, isoform C OS=Drosophila melanogaster OX=7227 GN=Pal1 PE=1 SV=1 | 10 | 7 | 1 | 1 | 3.321928095 | 0 | 3.321928095 |
| A1Z936 | A1Z936_DROME FI18626p1 OS=Drosophila melanogaster OX=7227 GN=ZnT49B PE=1 SV=1 | 10 | 8 | 1 | 1 | 3.321928095 | 0 | 3.321928095 |
| A1ZBH2 | A1ZBH2_DROME Uncharacterized protein, isoform A OS=Drosophila melanogaster OX=7227 GN=CG15111 PE=1 SV=1 | 10 | 8 | 1 | 1 | 3.321928095 | 0 | 3.321928095 |
| A4V0S1 | A4V0S1_DROME Syntaxin 5, isoform B OS=Drosophila melanogaster OX=7227 GN=Syx5 PE=1 SV=1 | 10 | 5 | 1 | 1 | 3.321928095 | 0 | 3.321928095 |
| A8JR46 | A8JR46_DROME Inositol-requiring enzyme-1 OS=Drosophila melanogaster OX=7227 GN=Ire1 PE=4 SV=1 | 10 | 8 | 1 | 1 | 3.321928095 | 0 | 3.321928095 |
| J7K3P9 | J7K3P9_DROME AQP7777 transcript variant D OS=Drosophila melanogaster OX=7227 GN=Prip PE=1 SV=1 | 10 | 4 | 1 | 1 | 3.321928095 | 0 | 3.321928095 |
| M9PEW1 | M9PEW1_DROME Uncharacterized protein, isoform E OS=Drosophila melanogaster OX=7227 GN=CG18815 PE=1 SV=1 | 10 | 6 | 1 | 1 | 3.321928095 | 0 | 3.321928095 |
| O15971 | O15971_DROME LD39986p OS=Drosophila melanogaster OX=7227 GN=Rab10 PE=1 SV=1 | 10 | 4 | 1 | 1 | 3.321928095 | 0 | 3.321928095 |
| Q7KK97 | Q7KK97_DROME LD44652p OS=Drosophila melanogaster OX=7227 GN=sut1 PE=1 SV=1 | 10 | 5 | 1 | 1 | 3.321928095 | 0 | 3.321928095 |

| UniProt ID | Protein description | # PSMs MVB spectra | # of unique peptides MVB | # of PSMs lysate | # of unique peptides lysate | log2(PSM MVB) | log2(PSM lysate) | FC (log2 of MVB PSM-log2 of lysate PSM) |
|---|---|---|---|---|---|---|---|---|
| Q8T481 | Q8T481_DROME AT11343p OS=Drosophila melanogaster OX=7227 GN=BcDNA:AT11343 PE=1 SV=1 | 10 | 6 | 1 | 1 | 3.321928095 | 0 | 3.321928095 |
| Q95T98 | Q95T98_DROME Elongation of very long chain fatty acids protein OS=Drosophila melanogaster OX=7227 GN=CG9798 PE=2 SV=1 | 10 | 5 | 1 | 1 | 3.321928095 | 0 | 3.321928095 |
| Q9VB96 | Q9VB96_DROME Uncharacterized protein, isoform A OS=Drosophila melanogaster OX=7227 GN=CG31075 PE=1 SV=2 | 10 | 7 | 1 | 1 | 3.321928095 | 0 | 3.321928095 |
| Q9VCI4 | Q9VCI4_DROME LD32918p OS=Drosophila melanogaster OX=7227 GN=Dmel\CG10217 PE=2 SV=2 | 10 | 8 | 1 | 1 | 3.321928095 | 0 | 3.321928095 |
| Q9VMQ6 | Q9VMQ6_DROME RE57459p OS=Drosophila melanogaster OX=7227 GN=CG6922 PE=2 SV=2 | 10 | 4 | 1 | 1 | 3.321928095 | 0 | 3.321928095 |
| Q9VNW8 | Q9VNW8_DROME FI24011p1 OS=Drosophila melanogaster OX=7227 GN=CG7458-RA PE=2 SV=3 | 10 | 8 | 1 | 1 | 3.321928095 | 0 | 3.321928095 |
| Q9VRU2 | Q9VRU2_DROME LD30746p OS=Drosophila melanogaster OX=7227 GN=loj PE=1 SV=2 | 10 | 5 | 1 | 1 | 3.321928095 | 0 | 3.321928095 |
| Q9VSW1 | Q9VSW1_DROME UTP--glucose-1-phosphate uridylyltransferase OS=Drosophila melanogaster OX=7227 GN=UGP PE=1 SV=2 | 10 | 9 | 1 | 1 | 3.321928095 | 0 | 3.321928095 |
| Q9VXD6 | Q9VXD6_DROME LD39612p OS=Drosophila melanogaster OX=7227 GN=Dmel\CG9723 PE=2 SV=1 | 10 | 3 | 1 | 1 | 3.321928095 | 0 | 3.321928095 |
| Q24492 | RFA1_DROME Replication protein A 70 kDa DNA-binding subunit OS=Drosophila melanogaster OX=7227 GN=RpA-70 PE=1 SV=1 | 9 | 8 | 50 | 23 | 3.169925001 | 5.64385619 | -2.473931188 |
| X2JFR6 | X2JFR6_DROME Eukaryotic translation initiation factor 2 subunit alpha, isoform B OS=Drosophila melanogaster OX=7227 GN=eIF2alpha PE=4 SV=1 | 9 | 8 | 45 | 17 | 3.169925001 | 5.491853096 | -2.321928095 |
| Q9VGP4 | Q9VGP4_DROME LP08082p OS=Drosophila melanogaster OX=7227 GN=Ranbp9 PE=1 SV=1 | 9 | 7 | 39 | 19 | 3.169925001 | 5.285402219 | -2.115477217 |
| Q9XYU0 | MCM7_DROME DNA replication licensing factor Mcm7 OS=Drosophila melanogaster OX=7227 GN=Mcm7 PE=1 SV=1 | 9 | 9 | 37 | 21 | 3.169925001 | 5.209453366 | -2.039528364 |
| X2JKH9 | X2JKH9_DROME Uncharacterized protein, isoform F OS=Drosophila melanogaster OX=7227 GN=CG8184 PE=1 SV=1 | 9 | 4 | 34 | 24 | 3.169925001 | 5.087462841 | -1.91753784 |
| A1Z992 | A1Z992_DROME 1,4-Alpha-Glucan branching enzyme OS=Drosophila melanogaster OX=7227 GN=AGBE PE=1 SV=1 | 9 | 7 | 33 | 20 | 3.169925001 | 5.044394119 | -1.874469118 |
| Q9V597 | RL31_DROME 60S ribosomal protein L31 OS=Drosophila melanogaster OX=7227 GN=RpL31 PE=1 SV=1 | 9 | 1 | 32 | 4 | 3.169925001 | 5 | -1.830074999 |
| Q9VVU1 | Q9VVU1_DROME FI09602p OS=Drosophila melanogaster OX=7227 GN=CG 3902 PE=1 SV=1 | 9 | 6 | 26 | 11 | 3.169925001 | 4.700439718 | -1.530514717 |
| X2JEX7 | X2JEX7_DROME Ubiquitin-specific protease 7, isoform C OS=Drosophila melanogaster OX=7227 GN=Usp7 PE=3 SV=1 | 9 | 8 | 26 | 22 | 3.169925001 | 4.700439718 | -1.530514717 |
| P48159 | RL23_DROME 60S ribosomal protein L23 OS=Drosophila melanogaster OX=7227 GN=RpL23 PE=1 SV=2 | 9 | 6 | 25 | 9 | 3.169925001 | 4.64385619 | -1.473931188 |
| Q9V677 | ECM29_DROME Proteasome-associated protein ECM29 homolog OS=Drosophila melanogaster OX=7227 GN=CG8858 PE=1 SV=1 | 9 | 7 | 25 | 19 | 3.169925001 | 4.64385619 | -1.473931188 |
| Q7KUA4 | Q7KUA4_DROME LD22577p OS=Drosophila melanogaster OX=7227 GN=Uba2 PE=1 SV=1 | 9 | 7 | 25 | 13 | 3.169925001 | 4.64385619 | -1.473931188 |
| Q9VAC1 | Q9VAC1_DROME GM14349p OS=Drosophila melanogaster OX=7227 GN=BEST:GH02390 PE=1 SV=1 | 9 | 7 | 25 | 12 | 3.169925001 | 4.64385619 | -1.473931188 |
| Q7KMP8 | Q7KMP8_DROME 26S proteasome regulatory complex subunit p39A OS=Drosophila melanogaster OX=7227 GN=Rpn9 PE=1 SV=1 | 9 | 7 | 23 | 13 | 3.169925001 | 4.523561956 | -1.353636955 |
| P08181 | CSK2A_DROME Casein kinase II subunit alpha OS=Drosophila melanogaster OX=7227 GN=CkIIalpha PE=1 SV=2 | 9 | 8 | 22 | 11 | 3.169925001 | 4.459431619 | -1.289506617 |
| Q9VF51 | Q9VF51_DROME Aldehyde oxidase 3, isoform A OS=Drosophila melanogaster OX=7227 GN=AOX3 PE=1 SV=1 | 9 | 9 | 21 | 17 | 3.169925001 | 4.392317423 | -1.222392421 |
| P52486 | UBCD4_DROME Ubiquitin-conjugating enzyme E2-22 kDa OS=Drosophila melanogaster OX=7227 GN=Ubc4 PE=1 SV=2 | 9 | 6 | 20 | 7 | 3.169925001 | 4.321928095 | -1.152003093 |
| P26686 | SRR55_DROME Serine-arginine protein 55 OS=Drosophila melanogaster OX=7227 GN=B52 PE=1 SV=4 | 9 | 8 | 17 | 8 | 3.169925001 | 4.087462841 | -0.91753784 |
| A0A0B4LH53 | A0A0B4LH53_DROME N-ethylmaleimide-sensitive factor 2, isoform B OS=Drosophila melanogaster OX=7227 GN=Nsf2 PE=4 SV=1 | 9 | 8 | 16 | 11 | 3.169925001 | 4 | -0.830074999 |
| O97102 | O97102_DROME Small ubiquitin-related modifier OS=Drosophila melanogaster OX=7227 GN=smt3 PE=1 SV=1 | 9 | 6 | 12 | 4 | 3.169925001 | 3.584962501 | -0.415037499 |
| Q9VBI2 | Q9VBI2_DROME UMP-CMP kinase OS=Drosophila melanogaster OX=7227 GN=Dak1 PE=1 SV=1 | 9 | 5 | 12 | 7 | 3.169925001 | 3.584962501 | -0.415037499 |
| Q9VH69 | RS29_DROME 40S ribosomal protein S29 OS=Drosophila melanogaster OX=7227 GN=RpS29 PE=1 SV=1 | 9 | 4 | 11 | 4 | 3.169925001 | 3.459431619 | -0.289506617 |
| A0A0B4KGT8 | A0A0B4KGT8_DROME Cheerio, isoform M OS=Drosophila melanogaster OX=7227 GN=cher PE=1 SV=1 | 9 | 9 | 11 | 10 | 3.169925001 | 3.459431619 | -0.289506617 |
| Q8SX57 | Q8SX57_DROME LD44221p OS=Drosophila melanogaster OX=7227 GN=CG11005 PE=1 SV=1 | 9 | 5 | 11 | 7 | 3.169925001 | 3.459431619 | -0.289506617 |
| Q27333 | NT56_DROME Lethal(2)neighbour of tid protein OS=Drosophila melanogaster OX=7227 GN=l(2)not PE=1 SV=1 | 9 | 6 | 4 | 3 | 3.169925001 | 2 | 1.169925001 |
| Q7K1V5 | JAGN_DROME Protein jagunal OS=Drosophila melanogaster OX=7227 GN=jagn PE=1 SV=1 | 9 | 3 | 1 | 1 | 3.169925001 | 0 | 3.169925001 |
| Q9V3D2 | HEM6_DROME Oxygen-dependent coproporphyrinogen-III oxidase OS=Drosophila melanogaster OX=7227 GN=Coprox PE=2 SV=1 | 9 | 8 | 3 | 3 | 3.169925001 | 1.584962501 | 1.584962501 |
| Q9V9J3 | SRC42_DROME Tyrosine-protein kinase Src42A OS=Drosophila melanogaster OX=7227 GN=Src42A PE=1 SV=1 | 9 | 9 | 1 | 1 | 3.169925001 | 0 | 3.169925001 |
| Q9VE50 | GOSR1_DROME Golgi SNAP receptor complex member 1 OS=Drosophila melanogaster OX=7227 GN=Gos28 PE=2 SV=1 | 9 | 5 | 1 | 1 | 3.169925001 | 0 | 3.169925001 |
| Q9VND8 | RHEB_DROME GTP-binding protein Rheb homolog OS=Drosophila melanogaster OX=7227 GN=Rheb PE=1 SV=1 | 9 | 5 | 5 | 4 | 3.169925001 | 2.321928095 | 0.847996907 |
| Q9XYN7 | PSB3_DROME Proteasome subunit beta type-3 OS=Drosophila melanogaster OX=7227 GN=Prosbeta3 PE=1 SV=1 | 9 | 4 | 6 | 4 | 3.169925001 | 2.584962501 | 0.584962501 |
| A1Z8W9 | A1Z8W9_DROME Gartenzwerg, isoform A OS=Drosophila melanogaster OX=7227 GN=garz PE=1 SV=1 | 9 | 6 | 3 | 3 | 3.169925001 | 1.584962501 | 1.584962501 |
| A8DQW8 | A8DQW8_DROME Closca OS=Drosophila melanogaster OX=7227 GN=clos PE=4 SV=3 | 9 | 9 | 7 | 7 | 3.169925001 | 2.807354922 | 0.362570079 |
| M9PEX7 | M9PEX7_DROME Uncharacterized protein, isoform B OS=Drosophila melanogaster OX=7227 GN=CG3408 PE=1 SV=1 | 9 | 6 | 1 | 1 | 3.169925001 | 0 | 3.169925001 |
| Q7K332 | Q7K332_DROME GH17623p OS=Drosophila melanogaster OX=7227 GN=CG3364 PE=2 SV=1 | 9 | 7 | 4 | 4 | 3.169925001 | 2 | 1.169925001 |
| Q8IPK9 | Q8IPK9_DROME Receptor protein serine/threonine kinase OS=Drosophila melanogaster OX=7227 GN=tkv PE=3 SV=1 | 9 | 6 | 2 | 2 | 3.169925001 | 1 | 2.169925001 |
| Q9VGW9 | Q9VGW9_DROME LD25118p OS=Drosophila melanogaster OX=7227 GN=mRpL37 PE=1 SV=3 | 9 | 7 | 7 | 6 | 3.169925001 | 2.807354922 | 0.362570079 |
| Q9VJD0 | Q9VJD0_DROME Mitochondrial inner membrane protease ATP23 OS=Drosophila melanogaster OX=7227 GN=Dmel\CG5131 PE=1 SV=2 | 9 | 8 | 3 | 2 | 3.169925001 | 1.584962501 | 1.584962501 |
| Q9VXZ0 | Q9VXZ0_DROME GEO07190p1 OS=Drosophila melanogaster OX=7227 GN=ND-B18 PE=1 SV=1 | 9 | 4 | 2 | 2 | 3.169925001 | 1 | 2.169925001 |
| Q9VZ34 | Q9VZ34_DROME RH72958p OS=Drosophila melanogaster OX=7227 GN=Dmel\CG2076 PE=1 SV=1 | 9 | 4 | 2 | 2 | 3.169925001 | 1 | 2.169925001 |
| Q9W028 | Q9W028_DROME IP19120p OS=Drosophila melanogaster OX=7227 GN=yellow-g2 PE=1 SV=1 | 9 | 8 | 2 | 2 | 3.169925001 | 1 | 2.169925001 |
| X2J7M7 | X2J7M7_DROME Uncharacterized protein, isoform E OS=Drosophila melanogaster OX=7227 GN=CG4658 PE=1 SV=1 | 9 | 7 | 2 | 1 | 3.169925001 | 1 | 2.169925001 |
| X2JB52 | X2JB52_DROME Purple acid phosphatase OS=Drosophila melanogaster OX=7227 GN=CG1637 PE=1 SV=1 | 9 | 6 | 2 | 2 | 3.169925001 | 1 | 2.169925001 |
| Q9VHJ4 | Q9VHJ4_DROME GH04846p OS=Drosophila melanogaster OX=7227 GN=RagA-B PE=1 SV=1 | 9 | 5 | 1 | 1 | 3.169925001 | 0 | 3.169925001 |
| Q9VMX4 | Q9VMX4_DROME AT19154p OS=Drosophila melanogaster OX=7227 GN=Scox PE=1 SV=1 | 9 | 6 | 1 | 1 | 3.169925001 | 0 | 3.169925001 |
| B7Z031 | PESTE_DROME Protein peste OS=Drosophila melanogaster OX=7227 GN=pes PE=2 SV=1 | 9 | 6 | 1 | 1 | 3.169925001 | 0 | 3.169925001 |
| O61491 | FLOT1_DROME Flotillin-1 OS=Drosophila melanogaster OX=7227 GN=Flo1 PE=2 SV=1 | 9 | 8 | 1 | 1 | 3.169925001 | 0 | 3.169925001 |
| O97422 | B3GI_DROME Galactosylgalactosylxylosylprotein 3-beta-glucuronosyltransferase I OS=Drosophila melanogaster OX=7227 GN=GlcAT-I PE=2 SV=2 | 9 | 6 | 1 | 1 | 3.169925001 | 0 | 3.169925001 |
| P10041 | DL_DROME Neurogenic locus protein delta OS=Drosophila melanogaster OX=7227 GN=Dl PE=1 SV=2 | 9 | 6 | 1 | 1 | 3.169925001 | 0 | 3.169925001 |
| P35992 | PTP10_DROME Tyrosine-protein phosphatase 10D OS=Drosophila melanogaster OX=7227 GN=Ptp10D PE=1 SV=4 | 9 | 7 | 1 | 1 | 3.169925001 | 0 | 3.169925001 |
| P49858 | CNI_DROME Protein cornichon OS=Drosophila melanogaster OX=7227 GN=cni PE=1 SV=1 | 9 | 3 | 1 | 1 | 3.169925001 | 0 | 3.169925001 |
| Q0KHU5 | PCAT_DROME Lysophosphatidylcholine acyltransferase OS=Drosophila melanogaster OX=7227 GN=LPCAT PE=2 SV=1 | 9 | 7 | 1 | 1 | 3.169925001 | 0 | 3.169925001 |
| Q7JW12 | TMX2_DROME Thioredoxin-related transmembrane protein 2 homolog OS=Drosophila melanogaster OX=7227 GN=CG11007 PE=2 SV=1 | 9 | 7 | 1 | 1 | 3.169925001 | 0 | 3.169925001 |
| Q8IQ56 | TMM11_DROME Transmembrane protein 11 homolog, mitochondrial OS=Drosophila melanogaster OX=7227 GN=Pmi PE=1 SV=3 | 9 | 6 | 1 | 1 | 3.169925001 | 0 | 3.169925001 |
| Q9U6M0 | ECSIT_DROME Evolutionarily conserved signaling intermediate in Toll pathway, mitochondrial OS=Drosophila melanogaster OX=7227 GN=ECSIT PE=1 SV=2 | 9 | 3 | 1 | 1 | 3.169925001 | 0 | 3.169925001 |
| Q9VLR9 | MPPE1_DROME Metallophosphoesterase 1 homolog OS=Drosophila melanogaster OX=7227 GN=PGAP5 PE=2 SV=4 | 9 | 6 | 1 | 1 | 3.169925001 | 0 | 3.169925001 |
| Q9VSB9 | TMM43_DROME Transmembrane protein 43 homolog OS=Drosophila melanogaster OX=7227 GN=CG8111 PE=2 SV=1 | 9 | 5 | 1 | 1 | 3.169925001 | 0 | 3.169925001 |
| Q9VV72 | MINP1_DROME Multiple inositol polyphosphate phosphatase 1 OS=Drosophila melanogaster OX=7227 GN=Mipp1 PE=1 SV=2 | 9 | 8 | 1 | 1 | 3.169925001 | 0 | 3.169925001 |
| Q9VWE0 | DOME_DROME Cytokine receptor OS=Drosophila melanogaster OX=7227 GN=dome PE=1 SV=1 | 9 | 8 | 1 | 1 | 3.169925001 | 0 | 3.169925001 |
| Q9VZM5 | PIGB_DROME GPI mannosyltransferase 3 OS=Drosophila melanogaster OX=7227 GN=PIG-B PE=2 SV=2 | 9 | 6 | 1 | 1 | 3.169925001 | 0 | 3.169925001 |
| A0A0B4LET2 | A0A0B4LET2_DROME Cyp6u1, isoform B OS=Drosophila melanogaster OX=7227 GN=Cyp6u1 PE=3 SV=1 | 9 | 6 | 1 | 1 | 3.169925001 | 0 | 3.169925001 |
| A0A0B4LF89 | A0A0B4LF89_DROME Beta-hexosaminidase OS=Drosophila melanogaster OX=7227 GN=fdl PE=1 SV=1 | 9 | 7 | 1 | 1 | 3.169925001 | 0 | 3.169925001 |
| A1Z7C4 | A1Z7C4_DROME LDL receptor protein 1, isoform G OS=Drosophila melanogaster OX=7227 GN=LRP1 PE=1 SV=2 | 9 | 8 | 1 | 1 | 3.169925001 | 0 | 3.169925001 |
| A1ZBL2 | A1ZBL2_DROME CG7744 OS=Drosophila melanogaster OX=7227 GN=CG7744-RA PE=1 SV=1 | 9 | 7 | 1 | 1 | 3.169925001 | 0 | 3.169925001 |
| A4V2I0 | A4V2I0_DROME Amalgam, isoform C OS=Drosophila melanogaster OX=7227 GN=Ama PE=1 SV=1 | 9 | 5 | 1 | 1 | 3.169925001 | 0 | 3.169925001 |
| A8DYJ3 | A8DYJ3_DROME Calpain-A, isoform C OS=Drosophila melanogaster OX=7227 GN=CalpA PE=1 SV=1 | 9 | 7 | 1 | 1 | 3.169925001 | 0 | 3.169925001 |
| B7YZM7 | B7YZM7_DROME Uncharacterized protein, isoform L OS=Drosophila melanogaster OX=7227 GN=CG42672 PE=1 SV=1 | 9 | 7 | 1 | 1 | 3.169925001 | 0 | 3.169925001 |
| E1JHC8 | E1JHC8_DROME Galactosylgalactosylxylosylprotein 3-beta-glucuronosyltransferase OS=Drosophila melanogaster OX=7227 GN=GlcAT-S PE=2 SV=1 | 9 | 5 | 1 | 1 | 3.169925001 | 0 | 3.169925001 |
| L0MLL1 | L0MLL1_DROME Cation-transporting ATPase OS=Drosophila melanogaster OX=7227 GN=anne PE=3 SV=1 | 9 | 8 | 1 | 1 | 3.169925001 | 0 | 3.169925001 |
| M9PB53 | M9PB53_DROME Sodium/hydrogen exchanger OS=Drosophila melanogaster OX=7227 GN=Nhe3 PE=3 SV=1 | 9 | 4 | 1 | 1 | 3.169925001 | 0 | 3.169925001 |
| M9PFD0 | M9PFD0_DROME Uncharacterized protein, isoform C OS=Drosophila melanogaster OX=7227 GN=ABCA PE=4 SV=1 | 9 | 8 | 1 | 1 | 3.169925001 | 0 | 3.169925001 |
| M9PFY9 | M9PFY9_DROME Nedd4, isoform M OS=Drosophila melanogaster OX=7227 GN=Nedd4 PE=1 SV=1 | 9 | 5 | 1 | 1 | 3.169925001 | 0 | 3.169925001 |

| UniProt ID | Protein description | # PSMs MVB spectra | # of unique peptides MVB | # of PSMs lysate | # of unique peptides lysate | log2(PSM MVB) | log2(PSM lysate) | FC (log2 of MVB PSM-log2 of lysate PSM) |
|---|---|---|---|---|---|---|---|---|
| Q6IDF5 | Q6IDF5_DROME NADH dehydrogenase (Ubiquinone) B15 subunit OS=Drosophila melanogaster OX=7227 GN=ND-B15 PE=1 SV=1 | 9 | 2 | 1 | 1 | 3.169925001 | 0 | 3.169925001 |
| Q6NN55 | Q6NN55_DROME Oysgedart, isoform A OS=Drosophila melanogaster OX=7227 GN=oys PE=2 SV=1 | 9 | 5 | 1 | 1 | 3.169925001 | 0 | 3.169925001 |
| Q7JZ56 | Q7JZ56_DROME Metallo endopeptidase OS=Drosophila melanogaster OX=7227 GN=S2P PE=2 SV=1 | 9 | 4 | 1 | 1 | 3.169925001 | 0 | 3.169925001 |
| Q7K0S1 | Q7K0S1_DROME LD39211p OS=Drosophila melanogaster OX=7227 GN=CG14764 PE=2 SV=1 | 9 | 5 | 1 | 1 | 3.169925001 | 0 | 3.169925001 |
| Q7K0W1 | Q7K0W1_DROME CG8531 OS=Drosophila melanogaster OX=7227 GN=CG8531 PE=1 SV=1 | 9 | 8 | 1 | 1 | 3.169925001 | 0 | 3.169925001 |
| Q95SH7 | Q95SH7_DROME GH26007p OS=Drosophila melanogaster OX=7227 GN=Dmel\CG14969 PE=1 SV=1 | 9 | 5 | 1 | 1 | 3.169925001 | 0 | 3.169925001 |
| Q95U32 | Q95U32_DROME GH11867p OS=Drosophila melanogaster OX=7227 GN=Rpb7 PE=2 SV=1 | 9 | 6 | 1 | 1 | 3.169925001 | 0 | 3.169925001 |
| Q9V3V2 | Q9V3V2_DROME Peptidylprolyl isomerase OS=Drosophila melanogaster OX=7227 GN=Fkbp14 PE=1 SV=1 | 9 | 5 | 1 | 1 | 3.169925001 | 0 | 3.169925001 |
| Q9V9T9 | Q9V9T9_DROME Beta-glucuronidase OS=Drosophila melanogaster OX=7227 GN=betaGlu PE=1 SV=1 | 9 | 8 | 1 | 1 | 3.169925001 | 0 | 3.169925001 |
| Q9VKZ9 | Q9VKZ9_DROME FI06463p OS=Drosophila melanogaster OX=7227 GN=l(2)SH0834 PE=1 SV=1 | 9 | 4 | 1 | 1 | 3.169925001 | 0 | 3.169925001 |
| Q9VQ34 | Q9VQ34_DROME GH17801p OS=Drosophila melanogaster OX=7227 GN=Dmel\CG17660 PE=1 SV=3 | 9 | 6 | 1 | 1 | 3.169925001 | 0 | 3.169925001 |
| Q9VR44 | Q9VR44_DROME GH05102p OS=Drosophila melanogaster OX=7227 GN=151989_at PE=1 SV=1 | 9 | 2 | 1 | 1 | 3.169925001 | 0 | 3.169925001 |
| Q9W2B3 | Q9W2B3_DROME CG9304, isoform A OS=Drosophila melanogaster OX=7227 GN=CG9304 PE=4 SV=3 | 9 | 3 | 1 | 1 | 3.169925001 | 0 | 3.169925001 |
| Q9W4J6 | Q9W4J6_DROME LD19190p OS=Drosophila melanogaster OX=7227 GN=Dmel\CG11436 PE=2 SV=1 | 9 | 8 | 1 | 1 | 3.169925001 | 0 | 3.169925001 |
| X2JC94 | X2JC94_DROME Phosphoglycerate mutase 5, isoform B OS=Drosophila melanogaster OX=7227 GN=Pgam5 PE=4 SV=1 | 9 | 8 | 1 | 1 | 3.169925001 | 0 | 3.169925001 |
| X2JFT2 | X2JFT2_DROME Gone early, isoform B OS=Drosophila melanogaster OX=7227 GN=goe PE=1 SV=1 | 9 | 9 | 1 | 1 | 3.169925001 | 0 | 3.169925001 |
| P38979 | RSSA_DROME 40S ribosomal protein SA OS=Drosophila melanogaster OX=7227 GN=sta PE=1 SV=3 | 8 | 6 | 62 | 16 | 3 | 5.95419631 | -2.95419631 |
| A0A0B4K618 | A0A0B4K618_DROME Fmr1, isoform G OS=Drosophila melanogaster OX=7227 GN=Fmr1 PE=1 SV=1 | 8 | 6 | 54 | 16 | 3 | 5.754887502 | -2.754887502 |
| A0A0B4LEV2 | A0A0B4LEV2_DROME DNA helicase OS=Drosophila melanogaster OX=7227 GN=dpa PE=1 SV=1 | 8 | 7 | 51 | 24 | 3 | 5.672425342 | -2.672425342 |
| Q9VBX3 | Q9VBX3_DROME Vig2, isoform B OS=Drosophila melanogaster OX=7227 GN=vig2 PE=1 SV=2 | 8 | 7 | 49 | 11 | 3 | 5.614709844 | -2.614709844 |
| Q9V3G7 | PSMD6_DROME 26S proteasome non-ATPase regulatory subunit 6 OS=Drosophila melanogaster OX=7227 GN=Rpn7 PE=2 SV=1 | 8 | 6 | 36 | 14 | 3 | 5.169925001 | -2.169925001 |
| Q7K3Z3 | Q7K3Z3_DROME GH01724p OS=Drosophila melanogaster OX=7227 GN=p47 PE=1 SV=1 | 8 | 6 | 36 | 13 | 3 | 5.169925001 | -2.169925001 |
| Q9VSD6 | Q9VSD6_DROME D-Importin 7/RanBP7 OS=Drosophila melanogaster OX=7227 GN=msk PE=1 SV=1 | 8 | 5 | 36 | 19 | 3 | 5.169925001 | -2.169925001 |
| Q7JZD3 | Q7JZD3_DROME Eb1, isoform A OS=Drosophila melanogaster OX=7227 GN=Eb1 PE=1 SV=1 | 8 | 6 | 33 | 13 | 3 | 5.044394119 | -2.044394119 |
| C7LAE4 | C7LAE4_DROME LP18603p OS=Drosophila melanogaster OX=7227 GN=Rm62 PE=1 SV=1 | 8 | 6 | 30 | 14 | 3 | 4.906890596 | -1.906890596 |
| Q7K2G1 | ADRM1_DROME Proteasomal ubiquitin receptor ADRM1 homolog OS=Drosophila melanogaster OX=7227 GN=Rpn13 PE=1 SV=1 | 8 | 4 | 29 | 12 | 3 | 4.857980995 | -1.857980995 |
| Q9VQF7 | BACC_DROME Bacchus OS=Drosophila melanogaster OX=7227 GN=Bacc PE=2 SV=1 | 8 | 4 | 26 | 7 | 3 | 4.700439718 | -1.700439718 |
| Q7KV27 | Q7KV27_DROME Uncharacterized protein, isoform A OS=Drosophila melanogaster OX=7227 GN=ALT PE=1 SV=1 | 8 | 6 | 26 | 12 | 3 | 4.700439718 | -1.700439718 |
| Q9VYV4 | Q9VYV4_DROME Amun, isoform A OS=Drosophila melanogaster OX=7227 GN=Amun PE=1 SV=1 | 8 | 6 | 26 | 12 | 3 | 4.700439718 | -1.700439718 |
| Q9VU35 | Q9VU35_DROME RH34413p OS=Drosophila melanogaster OX=7227 GN=Dmel\CG11267 PE=1 SV=1 | 8 | 5 | 25 | 9 | 3 | 4.64385619 | -1.64385619 |
| Q7K3W2 | Q7K3W2_DROME CG8728 OS=Drosophila melanogaster OX=7227 GN=CG8728-RA PE=1 SV=1 | 8 | 6 | 24 | 16 | 3 | 4.584962501 | -1.584962501 |
| Q9VPY2 | Q9VPY2_DROME Phospholipase A2 activator protein, isoform A OS=Drosophila melanogaster OX=7227 GN=Plap PE=1 SV=1 | 8 | 8 | 24 | 16 | 3 | 4.584962501 | -1.584962501 |
| Q9VMU4 | RL37A_DROME 60S ribosomal protein L37a OS=Drosophila melanogaster OX=7227 GN=RpL37A PE=1 SV=3 | 8 | 3 | 23 | 4 | 3 | 4.523561956 | -1.523561956 |
| P40796 | LA_DROME La protein homolog OS=Drosophila melanogaster OX=7227 GN=La PE=1 SV=2 | 8 | 7 | 22 | 11 | 3 | 4.459431619 | -1.459431619 |
| P41375 | IF2B_DROME Eukaryotic translation initiation factor 2 subunit 2 OS=Drosophila melanogaster OX=7227 GN=eIF2beta PE=1 SV=1 | 8 | 6 | 21 | 11 | 3 | 4.392317423 | -1.392317423 |
| Q9W3U9 | Q9W3U9_DROME CG14434-PA OS=Drosophila melanogaster OX=7227 GN=Dmel\CG14434 PE=1 SV=1 | 8 | 6 | 21 | 8 | 3 | 4.392317423 | -1.392317423 |
| Q9W3X6 | Q9W3X6_DROME FI05334p OS=Drosophila melanogaster OX=7227 GN=Fum1 PE=1 SV=3 | 8 | 6 | 21 | 9 | 3 | 4.392317423 | -1.392317423 |
| Q0KIB3 | Q0KIB3_DROME Histone acetyltransferase type B catalytic subunit OS=Drosophila melanogaster OX=7227 GN=Hat1 PE=1 SV=1 | 8 | 7 | 19 | 9 | 3 | 4.247927513 | -1.247927513 |
| Q9Y162 | Q9Y162_DROME BcDNA.GH02678 OS=Drosophila melanogaster OX=7227 GN=Vps4 PE=1 SV=1 | 8 | 8 | 19 | 12 | 3 | 4.247927513 | -1.247927513 |
| P53501 | ACT3_DROME Actin-57B OS=Drosophila melanogaster OX=7227 GN=Act57B PE=1 SV=1 | 8 | 4 | 18 | 5 | 3 | 4.169925001 | -1.169925001 |
| Q9VIE7 | Q9VIE7_DROME Burgundy, isoform B OS=Drosophila melanogaster OX=7227 GN=bur PE=1 SV=3 | 8 | 6 | 17 | 11 | 3 | 4.087462841 | -1.087462841 |
| Q9W0Y1 | TINA1_DROME Troponin C-akin-1 protein OS=Drosophila melanogaster OX=7227 GN=Tina-1 PE=2 SV=1 | 8 | 5 | 14 | 9 | 3 | 3.807354922 | -0.807354922 |
| Q9VIH1 | Q9VIH1_DROME CG9273 protein OS=Drosophila melanogaster OX=7227 GN=RPA2 PE=1 SV=2 | 8 | 6 | 14 | 7 | 3 | 3.807354922 | -0.807354922 |
| Q9VJZ6 | Q9VJZ6_DROME LD40224p OS=Drosophila melanogaster OX=7227 GN=Dmel\CG6523 PE=1 SV=1 | 8 | 6 | 13 | 6 | 3 | 3.700439718 | -0.700439718 |
| Q9VJZ5 | Q9VJZ5_DROME LD07294p OS=Drosophila melanogaster OX=7227 GN=Tap42 PE=1 SV=2 | 8 | 5 | 12 | 7 | 3 | 3.584962501 | -0.584962501 |
| Q9VZS3 | ETIF1_DROME Eukaryotic translation initiation factor eIF1 OS=Drosophila melanogaster OX=7227 GN=eIF1 PE=1 SV=1 | 8 | 4 | 11 | 5 | 3 | 3.459431619 | -0.459431619 |
| Q27237 | TID_DROME Protein tumorous imaginal discs, mitochondrial OS=Drosophila melanogaster OX=7227 GN=l(2)tid PE=1 SV=2 | 8 | 7 | 8 | 5 | 3 | 3 | 0 |
| P18475 | TORSO_DROME Tyrosine-protein kinase receptor torso OS=Drosophila melanogaster OX=7227 GN=tor PE=1 SV=2 | 8 | 7 | 1 | 1 | 3 | 0 | 3 |
| P21914 | SDHB_DROME Succinate dehydrogenase [ubiquinone] iron-sulfur subunit, mitochondrial OS=Drosophila melanogaster OX=7227 GN=SdhB PE=2 SV=2 | 8 | 6 | 7 | 6 | 3 | 2.807354922 | 0.192645078 |
| Q7KML2 | ACOX1_DROME Probable peroxisomal acyl-coenzyme A oxidase 1 OS=Drosophila melanogaster OX=7227 GN=CG5009 PE=1 SV=1 | 8 | 7 | 3 | 3 | 3 | 1.584962501 | 1.415037499 |
| Q9VNA4 | Y1161_DROME Uncharacterized protein CG1161 OS=Drosophila melanogaster OX=7227 GN=CG1161 PE=3 SV=1 | 8 | 2 | 2 | 1 | 3 | 1 | 2 |
| A0A0B4KG81 | A0A0B4KG81_DROME Uncharacterized protein, isoform B OS=Drosophila melanogaster OX=7227 GN=CG15099 PE=4 SV=1 | 8 | 7 | 2 | 2 | 3 | 1 | 2 |
| A0A0B4LGI9 | A0A0B4LGI9_DROME CG3760, isoform D OS=Drosophila melanogaster OX=7227 GN=CG3760 PE=1 SV=1 | 8 | 6 | 1 | 1 | 3 | 0 | 3 |
| O96824 | O96824_DROME EG:131F2.3 protein OS=Drosophila melanogaster OX=7227 GN=BcDNA:RE59545 PE=1 SV=1 | 8 | 5 | 1 | 1 | 3 | 0 | 3 |
| Q7JVK1 | Q7JVK1_DROME HL07956p OS=Drosophila melanogaster OX=7227 GN=mRpL18 PE=2 SV=1 | 8 | 5 | 7 | 5 | 3 | 2.807354922 | 0.192645078 |
| Q9U6R9 | Q9U6R9_DROME GH13039p OS=Drosophila melanogaster OX=7227 GN=gammaSnap1 PE=1 SV=1 | 8 | 6 | 4 | 4 | 3 | 2 | 1 |
| Q9VC05 | Q9VC05_DROME Oxysterol-binding protein OS=Drosophila melanogaster OX=7227 GN=Osbp PE=1 SV=1 | 8 | 6 | 5 | 5 | 3 | 2.321928095 | 0.678071905 |
| Q9VEV3 | Q9VEV3_DROME LP07287p OS=Drosophila melanogaster OX=7227 GN=Dmel\CG14894 PE=1 SV=2 | 8 | 4 | 5 | 3 | 3 | 2.321928095 | 0.678071905 |
| Q9VGS3 | Q9VGS3_DROME RH44771p OS=Drosophila melanogaster OX=7227 GN=SdhC PE=1 SV=2 | 8 | 5 | 4 | 2 | 3 | 2 | 1 |
| Q9VH37 | Q9VH37_DROME IP06524p OS=Drosophila melanogaster OX=7227 GN=Dmel\CG12811 PE=1 SV=1 | 8 | 4 | 3 | 2 | 3 | 1.584962501 | 1.415037499 |
| Q9VSR5 | Q9VSR5_DROME GM03767p OS=Drosophila melanogaster OX=7227 GN=mRpL12 PE=1 SV=1 | 8 | 5 | 4 | 3 | 3 | 2 | 1 |
| Q9VIX4 | Q9VIX4_DROME Acyl-coenzyme A oxidase OS=Drosophila melanogaster OX=7227 GN=CG17544-RA PE=1 SV=2 | 8 | 7 | 1 | 1 | 3 | 0 | 3 |
| Q9VQM2 | Q9VQM2_DROME NADH dehydrogenase [ubiquinone] 1 subunit C2 OS=Drosophila melanogaster OX=7227 GN=ND-B14.5B PE=1 SV=1 | 8 | 3 | 1 | 1 | 3 | 0 | 3 |
| Q9VV87 | Q9VV87_DROME Elongation of very long chain fatty acids protein OS=Drosophila melanogaster OX=7227 GN=Baldspot PE=2 SV=2 | 8 | 3 | 1 | 1 | 3 | 0 | 3 |
| P49283 | MVL_DROME Protein Malvolio OS=Drosophila melanogaster OX=7227 GN=Mvl PE=2 SV=2 | 8 | 4 | 1 | 1 | 3 | 0 | 3 |
| Q23979 | MY61F_DROME Myosin-IB OS=Drosophila melanogaster OX=7227 GN=Myo61F PE=1 SV=3 | 8 | 5 | 1 | 1 | 3 | 0 | 3 |
| Q6WV17 | GALT5_DROME Polypeptide N-acetylgalactosaminyltransferase 5 OS=Drosophila melanogaster OX=7227 GN=pgant5 PE=2 SV=2 | 8 | 6 | 1 | 1 | 3 | 0 | 3 |
| Q9V771 | C6A23_DROME Probable cytochrome P450 6a23 OS=Drosophila melanogaster OX=7227 GN=Cyp6a23 PE=2 SV=2 | 8 | 5 | 1 | 1 | 3 | 0 | 3 |
| Q9VWR8 | DPY19_DROME C-mannosyltransferase dpy-19 homolog OS=Drosophila melanogaster OX=7227 GN=CG6659 PE=2 SV=1 | 8 | 6 | 1 | 1 | 3 | 0 | 3 |
| A0A0B4K6E2 | A0A0B4K6E2_DROME Na/Ca-exchange protein, isoform G OS=Drosophila melanogaster OX=7227 GN=Calx PE=1 SV=1 | 8 | 7 | 1 | 1 | 3 | 0 | 3 |
| A0A0B4LFT2 | A0A0B4LFT2_DROME Cytochrome P450-4e2, isoform B OS=Drosophila melanogaster OX=7227 GN=Cyp4e2 PE=3 SV=1 | 8 | 5 | 1 | 1 | 3 | 0 | 3 |
| A8JUV7 | A8JUV7_DROME Brother of ihog, isoform G OS=Drosophila melanogaster OX=7227 GN=boi PE=1 SV=3 | 8 | 7 | 1 | 1 | 3 | 0 | 3 |
| M9NE97 | M9NE97_DROME Copper transporter 1A, isoform C OS=Drosophila melanogaster OX=7227 GN=Ctr1A PE=4 SV=1 | 8 | 4 | 1 | 1 | 3 | 0 | 3 |
| M9PDG9 | M9PDG9_DROME Uncharacterized protein, isoform B OS=Drosophila melanogaster OX=7227 GN=CG12773 PE=1 SV=1 | 8 | 6 | 1 | 1 | 3 | 0 | 3 |
| M9PGB8 | M9PGB8_DROME Uncharacterized protein, isoform E OS=Drosophila melanogaster OX=7227 GN=CG4293 PE=4 SV=1 | 8 | 7 | 1 | 1 | 3 | 0 | 3 |
| O18336 | O18336_DROME MIP20481p OS=Drosophila melanogaster OX=7227 GN=Rab14 PE=1 SV=1 | 8 | 7 | 1 | 1 | 3 | 0 | 3 |
| O77134 | O77134_DROME 9 kD basic protein OS=Drosophila melanogaster OX=7227 GN=ATPsynE PE=1 SV=1 | 8 | 1 | 1 | 1 | 3 | 0 | 3 |
| Q2PDR0 | Q2PDR0_DROME GEO11137p1 OS=Drosophila melanogaster OX=7227 GN=Dmel\CG33977 PE=1 SV=1 | 8 | 3 | 1 | 1 | 3 | 0 | 3 |
| Q7JXE1 | Q7JXE1_DROME CG8297 OS=Drosophila melanogaster OX=7227 GN=bug PE=2 SV=1 | 8 | 4 | 1 | 1 | 3 | 0 | 3 |
| Q7K4Y0 | Q7K4Y0_DROME Desaturase 1, isoform A OS=Drosophila melanogaster OX=7227 GN=Desat1 PE=1 SV=1 | 8 | 6 | 1 | 1 | 3 | 0 | 3 |
| Q7KUM8 | Q7KUM8_DROME Ecdysone-induced protein 28/29kD, isoform E OS=Drosophila melanogaster OX=7227 GN=Eip71CD PE=1 SV=2 | 8 | 6 | 1 | 1 | 3 | 0 | 3 |

| UniProt ID | Protein description | # PSMs MVB spectra | # of unique peptides MVB | # of PSMs lysate | # of unique peptides lysate | log2(PSM MVB) | log2(PSM lysate) | FC (log2 of MVB PSM-log2 of lysate PSM) |
|---|---|---|---|---|---|---|---|---|
| Q7PLG1 | Q7PLG1_DROME FI01132p OS=Drosophila melanogaster OX=7227 GN=CG41283 PE=2 SV=2 | 8 | 5 | 1 | 1 | 3 | 0 | 3 |
| Q7PLL1 | Q7PLL1_DROME Eukaryotic translation initiation factor 4B, isoform E OS=Drosophila melanogaster OX=7227 GN=eIF4B PE=1 SV=2 | 8 | 4 | 1 | 1 | 3 | 0 | 3 |
| Q960J9 | Q960J9_DROME LD47544p OS=Drosophila melanogaster OX=7227 GN=CG1733 PE=1 SV=1 | 8 | 5 | 1 | 1 | 3 | 0 | 3 |
| Q9VA18 | Q9VA18_DROME Coiled-coil-helix-coiled-coil-helix domain containing 3 OS=Drosophila melanogaster OX=7227 GN=Chchd3 PE=1 SV=1 | 8 | 5 | 1 | 1 | 3 | 0 | 3 |
| Q9VEU2 | Q9VEU2_DROME Derlin OS=Drosophila melanogaster OX=7227 GN=Der-2 PE=1 SV=1 | 8 | 5 | 1 | 1 | 3 | 0 | 3 |
| Q9VGW2 | Q9VGW2_DROME LD09343p OS=Drosophila melanogaster OX=7227 GN=TTLL15 PE=2 SV=1 | 8 | 8 | 1 | 1 | 3 | 0 | 3 |
| Q9VI53 | Q9VI53_DROME LD44267p OS=Drosophila melanogaster OX=7227 GN=Dmel\CG1105 PE=1 SV=1 | 8 | 6 | 1 | 1 | 3 | 0 | 3 |
| Q9VK30 | Q9VK30_DROME GH19029p OS=Drosophila melanogaster OX=7227 GN=Dmel\CG5287 PE=2 SV=2 | 8 | 4 | 1 | 1 | 3 | 0 | 3 |
| Q9VKC2 | Q9VKC2_DROME Amino acid transporter protein JHI-21 OS=Drosophila melanogaster OX=7227 GN=JhI-21 PE=1 SV=1 | 8 | 5 | 1 | 1 | 3 | 0 | 3 |
| Q9VND4 | Q9VND4_DROME GEO12035p1 OS=Drosophila melanogaster OX=7227 GN=Dmel\CG14671 PE=2 SV=1 | 8 | 4 | 1 | 1 | 3 | 0 | 3 |
| Q9VPB8 | Q9VPB8_DROME Peroxin 14 OS=Drosophila melanogaster OX=7227 GN=Pex14 PE=1 SV=1 | 8 | 6 | 1 | 1 | 3 | 0 | 3 |
| Q9VR90 | Q9VR90_DROME Syntaxin 16 OS=Drosophila melanogaster OX=7227 GN=Syx16 PE=1 SV=1 | 8 | 5 | 1 | 1 | 3 | 0 | 3 |
| Q9VSC3 | Q9VSC3_DROME Ribonuclease X25 OS=Drosophila melanogaster OX=7227 GN=RNaseX25 PE=1 SV=1 | 8 | 6 | 1 | 1 | 3 | 0 | 3 |
| Q9VX13 | Q9VX13_DROME FI05221p OS=Drosophila melanogaster OX=7227 GN=Dmel\CG15814 PE=2 SV=1 | 8 | 5 | 1 | 1 | 3 | 0 | 3 |
| Q9VZU8 | Q9VZU8_DROME RE67594p OS=Drosophila melanogaster OX=7227 GN=Alg2 PE=1 SV=1 | 8 | 5 | 1 | 1 | 3 | 0 | 3 |
| Q9VZW0 | Q9VZW0_DROME LD11247p OS=Drosophila melanogaster OX=7227 GN=Sk2 PE=2 SV=3 | 8 | 6 | 1 | 1 | 3 | 0 | 3 |
| Q9VZX8 | Q9VZX8_DROME AT02196p OS=Drosophila melanogaster OX=7227 GN=Dmel\CG12093 PE=1 SV=1 | 8 | 7 | 1 | 1 | 3 | 0 | 3 |
| Q9W366 | Q9W366_DROME RE59932p OS=Drosophila melanogaster OX=7227 GN=Dmel\CG12121 PE=1 SV=1 | 8 | 6 | 1 | 1 | 3 | 0 | 3 |
| Q9W4F7 | Q9W4F7_DROME LD22376p OS=Drosophila melanogaster OX=7227 GN=Dmel\CG6903 PE=1 SV=1 | 8 | 5 | 1 | 1 | 3 | 0 | 3 |
| Q9Y124 | Q9Y124_DROME BcDNA.GH08385 OS=Drosophila melanogaster OX=7227 GN=BcDNA.GH08385 PE=1 SV=1 | 8 | 6 | 1 | 1 | 3 | 0 | 3 |
| Q9I7D3 | CAPR1_DROME Caprin homolog OS=Drosophila melanogaster OX=7227 GN=Capr PE=1 SV=1 | 7 | 5 | 76 | 31 | 2.807354922 | 6.247927513 | -3.440572591 |
| A0A126GUM8 | A0A126GUM8_DROME Lingerer, isoform L OS=Drosophila melanogaster OX=7227 GN=lig PE=1 SV=1 | 7 | 6 | 76 | 17 | 2.807354922 | 6.247927513 | -3.440572591 |
| P15348 | TOP2_DROME DNA topoisomerase 2 OS=Drosophila melanogaster OX=7227 GN=Top2 PE=1 SV=1 | 7 | 5 | 43 | 26 | 2.807354922 | 5.426264755 | -2.618909833 |
| A0A0B4LFB3 | A0A0B4LFB3_DROME Eukaryotic translation initiation factor 3 subunit M OS=Drosophila melanogaster OX=7227 GN=eIF3m PE=3 SV=1 | 7 | 5 | 42 | 13 | 2.807354922 | 5.392317423 | -2.584962501 |
| P49735 | MCM2_DROME DNA replication licensing factor Mcm2 OS=Drosophila melanogaster OX=7227 GN=Mcm2 PE=1 SV=1 | 7 | 5 | 40 | 19 | 2.807354922 | 5.321928095 | -2.514573173 |
| Q9VJ86 | Q9VJ86_DROME Bicoid stability factor OS=Drosophila melanogaster OX=7227 GN=bsf PE=1 SV=2 | 7 | 7 | 40 | 26 | 2.807354922 | 5.321928095 | -2.514573173 |
| Q9VCA5 | SYFB_DROME Phenylalanine--tRNA ligase beta subunit OS=Drosophila melanogaster OX=7227 GN=beta-PheRS PE=1 SV=1 | 7 | 6 | 37 | 21 | 2.807354922 | 5.209453366 | -2.402098444 |
| Q9VK85 | Q9VK85_DROME Eukaryotic translation release factor 3, isoform A OS=Drosophila melanogaster OX=7227 GN=eRF3 PE=1 SV=1 | 7 | 5 | 37 | 18 | 2.807354922 | 5.209453366 | -2.402098444 |
| O76927 | RS21_DROME 40S ribosomal protein S21 OS=Drosophila melanogaster OX=7227 GN=RpS21 PE=1 SV=1 | 7 | 4 | 35 | 8 | 2.807354922 | 5.129283017 | -2.321928095 |
| Q9VSL8 | Q9VSL8_DROME Rhea, isoform B OS=Drosophila melanogaster OX=7227 GN=rhea PE=1 SV=1 | 7 | 7 | 35 | 27 | 2.807354922 | 5.129283017 | -2.321928095 |
| Q9VKZ8 | Q9VKZ8_DROME LP08774p OS=Drosophila melanogaster OX=7227 GN=Usp14 PE=1 SV=1 | 7 | 6 | 32 | 10 | 2.807354922 | 5 | -2.192645078 |
| C0HKA0 | RS14A_DROME 40S ribosomal protein S14a OS=Drosophila melanogaster OX=7227 GN=RpS14a PE=1 SV=1 | 7 | 2 | 31 | 9 | 2.807354922 | 4.95419631 | -2.146841388 |
| Q8IRG6 | SPT16_DROME FACT complex subunit spt16 OS=Drosophila melanogaster OX=7227 GN=dre4 PE=1 SV=2 | 7 | 6 | 31 | 18 | 2.807354922 | 4.95419631 | -2.146841388 |
| Q9VCK0 | EI3D1_DROME Eukaryotic translation initiation factor 3 subunit D-1 OS=Drosophila melanogaster OX=7227 GN=eIF3d1 PE=1 SV=1 | 7 | 6 | 31 | 13 | 2.807354922 | 4.95419631 | -2.146841388 |
| P09052 | VASA1_DROME ATP-dependent RNA helicase vasa OS=Drosophila melanogaster OX=7227 GN=vas PE=1 SV=3 | 7 | 6 | 27 | 18 | 2.807354922 | 4.754887502 | -1.94753258 |
| Q9V9Q4 | Q9V9Q4_DROME LD43819p OS=Drosophila melanogaster OX=7227 GN=Dmel\CG1416 PE=1 SV=2 | 7 | 4 | 27 | 13 | 2.807354922 | 4.754887502 | -1.94753258 |
| Q7KW39 | MMSA_DROME Probable methylmalonate-semialdehyde dehydrogenase [acylating], mitochondrial OS=Drosophila melanogaster OX=7227 GN=CG17896 PE=2 SV=1 | 7 | 6 | 26 | 11 | 2.807354922 | 4.700439718 | -1.893084796 |
| Q7K0E6 | Q7K0E6_DROME Aspartyl-tRNA synthetase, isoform A OS=Drosophila melanogaster OX=7227 GN=AspRS PE=1 SV=1 | 7 | 5 | 26 | 11 | 2.807354922 | 4.700439718 | -1.893084796 |
| A0A0B4K6D2 | A0A0B4K6D2_DROME Uncharacterized protein, isoform B OS=Drosophila melanogaster OX=7227 GN=p23 PE=1 SV=1 | 7 | 5 | 25 | 9 | 2.807354922 | 4.64385619 | -1.836501268 |
| F0JAG6 | F0JAG6_DROME Alpha actinin, isoform F OS=Drosophila melanogaster OX=7227 GN=Actn PE=1 SV=1 | 7 | 6 | 25 | 17 | 2.807354922 | 4.64385619 | -1.836501268 |
| Q0E940 | EIF3B_DROME Eukaryotic translation initiation factor 3 subunit B OS=Drosophila melanogaster OX=7227 GN=eIF3b PE=1 SV=1 | 7 | 7 | 23 | 15 | 2.807354922 | 4.523561956 | -1.716207034 |
| Q24276 | CDC37_DROME Hsp90 co-chaperone Cdc37 OS=Drosophila melanogaster OX=7227 GN=Cdc37 PE=1 SV=1 | 7 | 5 | 23 | 11 | 2.807354922 | 4.523561956 | -1.716207034 |
| Q9V3V6 | Q9V3V6_DROME 26S proteasome regulatory complex subunit p50 OS=Drosophila melanogaster OX=7227 GN=Rpt5 PE=1 SV=1 | 7 | 5 | 23 | 15 | 2.807354922 | 4.523561956 | -1.716207034 |
| Q9W1H4 | DNLI1_DROME DNA ligase 1 OS=Drosophila melanogaster OX=7227 GN=DNA-ligI PE=1 SV=2 | 7 | 5 | 22 | 13 | 2.807354922 | 4.459431619 | -1.652076697 |
| M9NG39 | M9NG39_DROME Protein on ecdysone puffs, isoform D OS=Drosophila melanogaster OX=7227 GN=Pep PE=1 SV=1 | 7 | 4 | 21 | 10 | 2.807354922 | 4.392317423 | -1.584962501 |
| Q9VDK7 | Q9VDK7_DROME Signal recognition particle subunit SRP72 OS=Drosophila melanogaster OX=7227 GN=Srp72 PE=1 SV=1 | 7 | 5 | 21 | 15 | 2.807354922 | 4.392317423 | -1.584962501 |
| P40301 | PSA2_DROME Proteasome subunit alpha type-2 OS=Drosophila melanogaster OX=7227 GN=Prosalpha2 PE=1 SV=1 | 7 | 6 | 20 | 8 | 2.807354922 | 4.321928095 | -1.514573173 |
| Q9VH07 | RUVB1_DROME RuvB-like helicase 1 OS=Drosophila melanogaster OX=7227 GN=pont PE=1 SV=1 | 7 | 6 | 20 | 11 | 2.807354922 | 4.321928095 | -1.514573173 |
| Q9VJD4 | Q9VJD4_DROME LD24721p OS=Drosophila melanogaster OX=7227 GN=Sgt PE=1 SV=1 | 7 | 5 | 20 | 10 | 2.807354922 | 4.321928095 | -1.514573173 |
| P48592 | RIR2_DROME Ribonucleoside-diphosphate reductase subunit M2 OS=Drosophila melanogaster OX=7227 GN=RnrS PE=1 SV=2 | 7 | 5 | 19 | 6 | 2.807354922 | 4.247927513 | -1.440572591 |
| Q9VXK6 | IF5_DROME Eukaryotic translation initiation factor 5 OS=Drosophila melanogaster OX=7227 GN=eIF5 PE=1 SV=1 | 7 | 5 | 19 | 12 | 2.807354922 | 4.247927513 | -1.440572591 |
| A0A0B4LES3 | A0A0B4LES3_DROME CG3107, isoform D OS=Drosophila melanogaster OX=7227 GN=CG3107 PE=4 SV=1 | 7 | 6 | 19 | 15 | 2.807354922 | 4.247927513 | -1.440572591 |
| A0A0B4KI24 | A0A0B4KI24_DROME Nucleoplasmin, isoform B OS=Drosophila melanogaster OX=7227 GN=Nlp PE=4 SV=1 | 7 | 5 | 18 | 6 | 2.807354922 | 4.169925001 | -1.362570079 |
| Q9VEJ0 | Q9VEJ0_DROME Peroxiredoxin 3 OS=Drosophila melanogaster OX=7227 GN=Prx3 PE=1 SV=2 | 7 | 6 | 18 | 10 | 2.807354922 | 4.169925001 | -1.362570079 |
| P35128 | UBE2N_DROME Ubiquitin-conjugating enzyme E2 N OS=Drosophila melanogaster OX=7227 GN=ben PE=1 SV=1 | 7 | 5 | 17 | 7 | 2.807354922 | 4.087462841 | -1.280107919 |
| Q9VFE4 | RS5B_DROME 40S ribosomal protein S5b OS=Drosophila melanogaster OX=7227 GN=RpS5b PE=2 SV=1 | 7 | 5 | 17 | 8 | 2.807354922 | 4.087462841 | -1.280107919 |
| A0A0B4KGX1 | A0A0B4KGX1_DROME Jaguar, isoform L OS=Drosophila melanogaster OX=7227 GN=jar PE=1 SV=1 | 7 | 6 | 17 | 14 | 2.807354922 | 4.087462841 | -1.280107919 |
| Q7JXZ2 | Q7JXZ2_DROME Eip55E, isoform A OS=Drosophila melanogaster OX=7227 GN=Eip55E PE=1 SV=1 | 7 | 5 | 17 | 9 | 2.807354922 | 4.087462841 | -1.280107919 |
| Q960M4 | Q960M4_DROME LD45324p OS=Drosophila melanogaster OX=7227 GN=Prx5 PE=1 SV=1 | 7 | 5 | 17 | 7 | 2.807354922 | 4.087462841 | -1.280107919 |
| Q960Z0 | KI10A_DROME Kinesin-like protein Klp10A OS=Drosophila melanogaster OX=7227 GN=Klp10A PE=1 SV=1 | 7 | 7 | 16 | 10 | 2.807354922 | 4 | -1.192645078 |
| Q9VL18 | EF1D_DROME Probable elongation factor 1-delta OS=Drosophila melanogaster OX=7227 GN=eEF1delta PE=1 SV=1 | 7 | 5 | 16 | 9 | 2.807354922 | 4 | -1.192645078 |
| O62602 | O62602_DROME Licorne OS=Drosophila melanogaster OX=7227 GN=lic PE=1 SV=1 | 7 | 4 | 16 | 10 | 2.807354922 | 4 | -1.192645078 |
| Q9VBU9 | Q9VBU9_DROME 40S ribosomal protein S27 OS=Drosophila melanogaster OX=7227 GN=RpS27 PE=1 SV=1 | 7 | 3 | 15 | 5 | 2.807354922 | 3.906890596 | -1.099535674 |
| Q7K221 | Q7K221_DROME Glutamate oxaloacetate transaminase 1, isoform A OS=Drosophila melanogaster OX=7227 GN=Got1 PE=1 SV=1 | 7 | 6 | 14 | 9 | 2.807354922 | 3.807354922 | -1 |
| X2JL49 | X2JL49_DROME Uncharacterized protein, isoform B OS=Drosophila melanogaster OX=7227 GN=CG7332 PE=4 SV=1 | 7 | 5 | 14 | 8 | 2.807354922 | 3.807354922 | -1 |
| Q9VUJ0 | RM39_DROME 39S ribosomal protein L39, mitochondrial OS=Drosophila melanogaster OX=7227 GN=mRpL39 PE=2 SV=2 | 7 | 5 | 13 | 6 | 2.807354922 | 3.700439718 | -0.893084796 |
| Q9VRD9 | Q9VRD9_DROME Cystathionine beta-synthase OS=Drosophila melanogaster OX=7227 GN=Cbs PE=1 SV=1 | 7 | 6 | 13 | 10 | 2.807354922 | 3.700439718 | -0.893084796 |
| P54353 | DOD_DROME Putative peptidyl-prolyl cis-trans isomerase dodo OS=Drosophila melanogaster OX=7227 GN=dod PE=2 SV=3 | 7 | 5 | 11 | 6 | 2.807354922 | 3.459431619 | -0.652076697 |
| P81900 | KAPR2_DROME cAMP-dependent protein kinase type II regulatory subunit OS=Drosophila melanogaster OX=7227 GN=Pka-R2 PE=1 SV=2 | 7 | 6 | 11 | 9 | 2.807354922 | 3.459431619 | -0.652076697 |
| Q9VAY9 | Q9VAY9_DROME GH07821p OS=Drosophila melanogaster OX=7227 GN=mRpS22 PE=1 SV=1 | 7 | 7 | 11 | 8 | 2.807354922 | 3.459431619 | -0.652076697 |
| Q9VZW7 | Q9VZW7_DROME Carnitine palmitoyltransferase 2 OS=Drosophila melanogaster OX=7227 GN=CPT2 PE=1 SV=1 | 7 | 6 | 11 | 9 | 2.807354922 | 3.459431619 | -0.652076697 |
| Q9V3Y2 | DRE2_DROME Anamorsin homolog OS=Drosophila melanogaster OX=7227 GN=CIAPIN1 PE=2 SV=1 | 7 | 5 | 9 | 7 | 2.807354922 | 3.169925001 | -0.362570079 |
| Q9V9U7 | Q9V9U7_DROME Tubulin-specific chaperone A OS=Drosophila melanogaster OX=7227 GN=Dmel\CG1890 PE=1 SV=1 | 7 | 4 | 9 | 5 | 2.807354922 | 3.169925001 | -0.362570079 |
| Q7K012 | Q7K012_DROME Brahma associated protein 55kD OS=Drosophila melanogaster OX=7227 GN=Bap55 PE=1 SV=1 | 7 | 4 | 8 | 5 | 2.807354922 | 3 | -0.192645078 |
| Q9VSU6 | Q9VSU6_DROME Dihydropteridine reductase OS=Drosophila melanogaster OX=7227 GN=Dhpr PE=1 SV=1 | 7 | 5 | 8 | 6 | 2.807354922 | 3 | -0.192645078 |
| O01404 | PHM_DROME Peptidylglycine alpha-hydroxylating monooxygenase OS=Drosophila melanogaster OX=7227 GN=Phm PE=1 SV=2 | 7 | 5 | 2 | 2 | 2.807354922 | 1 | 1.807354922 |
| P22058 | CPD1_DROME Chromosomal protein D1 OS=Drosophila melanogaster OX=7227 GN=D1 PE=1 SV=3 | 7 | 4 | 4 | 3 | 2.807354922 | 2 | 0.807354922 |
| P40945 | ARF2_DROME ADP-ribosylation factor 2 OS=Drosophila melanogaster OX=7227 GN=Arf102F PE=2 SV=2 | 7 | 4 | 3 | 3 | 2.807354922 | 1.584962501 | 1.222392421 |
| P42325 | NCAH_DROME Neurocalcin homolog OS=Drosophila melanogaster OX=7227 GN=Nca PE=1 SV=2 | 7 | 4 | 3 | 2 | 2.807354922 | 1.584962501 | 1.222392421 |
| Q24583 | VATF1_DROME V-type proton ATPase subunit F 1 OS=Drosophila melanogaster OX=7227 GN=Vha14-1 PE=1 SV=1 | 7 | 4 | 7 | 4 | 2.807354922 | 2.807354922 | 0 |
| Q7JWD3 | ASNA_DROME ATPase ASNA1 homolog OS=Drosophila melanogaster OX=7227 GN=CG1598 PE=2 SV=1 | 7 | 4 | 7 | 3 | 2.807354922 | 2.807354922 | 0 |

| UniProt ID | Protein description | # PSMs MVB spectra | # of unique peptides MVB | # of PSMs lysate | # of unique peptides lysate | log2(PSM MVB) | log2(PSM lysate) | FC (log2 of MVB PSM-log2 of lysate PSM) |
|---|---|---|---|---|---|---|---|---|
| Q9VHS2 | COX7A_DROME Cytochrome c oxidase subunit 7A, mitochondrial OS=Drosophila melanogaster OX=7227 GN=COX7A PE=3 SV=1 | 7 | 4 | 4 | 3 | 2.807354922 | 2 | 0.807354922 |
| Q9VXK0 | NIPSN_DROME Protein NipSnap OS=Drosophila melanogaster OX=7227 GN=Nipsnap PE=2 SV=2 | 7 | 5 | 5 | 4 | 2.807354922 | 2.321928095 | 0.485426827 |
| A0A0B4KH14 | A0A0B4KH14_DROME Uncharacterized protein, isoform B OS=Drosophila melanogaster OX=7227 GN=CG5447 PE=4 SV=1 | 7 | 5 | 1 | 1 | 2.807354922 | 0 | 2.807354922 |
| M9PC74 | M9PC74_DROME Uncharacterized protein, isoform G OS=Drosophila melanogaster OX=7227 GN=CG31650 PE=1 SV=1 | 7 | 7 | 2 | 2 | 2.807354922 | 1 | 1.807354922 |
| Q7JW46 | Q7JW46_DROME RE25483p OS=Drosophila melanogaster OX=7227 GN=CG11825 PE=2 SV=1 | 7 | 3 | 2 | 2 | 2.807354922 | 1 | 1.807354922 |
| Q9VB02 | Q9VB02_DROME SD09912p OS=Drosophila melanogaster OX=7227 GN=Dmel\CG12428 PE=1 SV=1 | 7 | 7 | 3 | 3 | 2.807354922 | 1.584962501 | 1.222392421 |
| Q9VVW7 | Q9VVW7_DROME Canopy b OS=Drosophila melanogaster OX=7227 GN=CNPYb PE=1 SV=2 | 7 | 3 | 7 | 2 | 2.807354922 | 2.807354922 | 0 |
| Q9VLS0 | Q9VLS0_DROME LD18607p OS=Drosophila melanogaster OX=7227 GN=Dmel\CG8460 PE=1 SV=1 | 7 | 4 | 3 | 3 | 2.807354922 | 1.584962501 | 1.222392421 |
| Q9VVG5 | Q9VVG5_DROME RH19679p OS=Drosophila melanogaster OX=7227 GN=BcDNA:RH19679 PE=1 SV=2 | 7 | 3 | 3 | 1 | 2.807354922 | 1.584962501 | 1.222392421 |
| Q9W3M8 | Q9W3M8_DROME LD34211p OS=Drosophila melanogaster OX=7227 GN=Ykt6 PE=1 SV=1 | 7 | 4 | 3 | 3 | 2.807354922 | 1.584962501 | 1.222392421 |
| Q9W5W8 | Q9W5W8_DROME RE22677p OS=Drosophila melanogaster OX=7227 GN=Dmel\CG9577 PE=1 SV=2 | 7 | 7 | 6 | 4 | 2.807354922 | 2.584962501 | 0.222392421 |
| Q9VGS9 | Q9VGS9_DROME LD21102p OS=Drosophila melanogaster OX=7227 GN=Ugt35a PE=1 SV=1 | 7 | 5 | 1 | 1 | 2.807354922 | 0 | 2.807354922 |
| Q9VHH1 | Q9VHH1_DROME LD42037p OS=Drosophila melanogaster OX=7227 GN=Dmel\CG8420 PE=1 SV=1 | 7 | 5 | 1 | 1 | 2.807354922 | 0 | 2.807354922 |
| P48607 | SPZ_DROME Protein spaetzle OS=Drosophila melanogaster OX=7227 GN=spz PE=1 SV=3 | 7 | 5 | 1 | 1 | 2.807354922 | 0 | 2.807354922 |
| Q8INQ7 | KTAP2_DROME Protein KRTCAP2 homolog OS=Drosophila melanogaster OX=7227 GN=CG31460 PE=2 SV=1 | 7 | 3 | 1 | 1 | 2.807354922 | 0 | 2.807354922 |
| Q8T0R7 | IDGF5_DROME Chitinase-like protein Idgf5 OS=Drosophila melanogaster OX=7227 GN=Idgf5 PE=2 SV=3 | 7 | 5 | 1 | 1 | 2.807354922 | 0 | 2.807354922 |
| Q9V4U9 | C6A13_DROME Probable cytochrome P450 6a13 OS=Drosophila melanogaster OX=7227 GN=Cyp6a13 PE=2 SV=1 | 7 | 4 | 1 | 1 | 2.807354922 | 0 | 2.807354922 |
| Q9VKH6 | PPT2_DROME Lysosomal thioesterase PPT2 homolog OS=Drosophila melanogaster OX=7227 GN=Ppt2 PE=2 SV=1 | 7 | 5 | 1 | 1 | 2.807354922 | 0 | 2.807354922 |
| Q9VT04 | PATH_DROME Proton-coupled amino acid transporter-like protein pathetic OS=Drosophila melanogaster OX=7227 GN=path PE=1 SV=1 | 7 | 4 | 1 | 1 | 2.807354922 | 0 | 2.807354922 |
| Q9VUW9 | HIP14_DROME Palmitoyltransferase Hip14 OS=Drosophila melanogaster OX=7227 GN=Hip14 PE=1 SV=1 | 7 | 7 | 1 | 1 | 2.807354922 | 0 | 2.807354922 |
| Q9VVD9 | S35B3_DROME Adenosine 3'-phospho 5'-phosphosulfate transporter 2 OS=Drosophila melanogaster OX=7227 GN=Papst2 PE=1 SV=2 | 7 | 3 | 1 | 1 | 2.807354922 | 0 | 2.807354922 |
| Q9VVX5 | MBOA5_DROME Lysophospholipid acyltransferase 5 OS=Drosophila melanogaster OX=7227 GN=nes PE=1 SV=1 | 7 | 2 | 1 | 1 | 2.807354922 | 0 | 2.807354922 |
| Q9VYF2 | IMPA3_DROME Putative inositol monophosphatase 3 OS=Drosophila melanogaster OX=7227 GN=CG15743 PE=2 SV=1 | 7 | 4 | 1 | 1 | 2.807354922 | 0 | 2.807354922 |
| Q9Y117 | GALT3_DROME Polypeptide N-acetylgalactosaminyltransferase 3 OS=Drosophila melanogaster OX=7227 GN=pgant3 PE=1 SV=1 | 7 | 6 | 1 | 1 | 2.807354922 | 0 | 2.807354922 |
| A0A0B4K7S0 | A0A0B4K7S0_DROME Fat facets, isoform D OS=Drosophila melanogaster OX=7227 GN=faf PE=3 SV=1 | 7 | 6 | 1 | 1 | 2.807354922 | 0 | 2.807354922 |
| A0A0B4KFZ0 | A0A0B4KFZ0_DROME Upstream of RpII140, isoform B OS=Drosophila melanogaster OX=7227 GN=140up PE=4 SV=1 | 7 | 3 | 1 | 1 | 2.807354922 | 0 | 2.807354922 |
| A0A0B4LF00 | A0A0B4LF00_DROME Palmitoyltransferase OS=Drosophila melanogaster OX=7227 GN=CG1407 PE=1 SV=1 | 7 | 3 | 1 | 1 | 2.807354922 | 0 | 2.807354922 |
| A0A0B4LGN1 | A0A0B4LGN1_DROME Inositol 1,4,5,-tris-phosphate receptor, isoform C OS=Drosophila melanogaster OX=7227 GN=Itp-r83A PE=4 SV=1 | 7 | 5 | 1 | 1 | 2.807354922 | 0 | 2.807354922 |
| A1Z9E0 | A1Z9E0_DROME Bb in a boxcar, isoform B OS=Drosophila melanogaster OX=7227 GN=bbc PE=1 SV=1 | 7 | 3 | 1 | 1 | 2.807354922 | 0 | 2.807354922 |
| A1ZB84 | A1ZB84_DROME FI22105p1 OS=Drosophila melanogaster OX=7227 GN=PIG-O PE=1 SV=1 | 7 | 7 | 1 | 1 | 2.807354922 | 0 | 2.807354922 |
| A8JUV9 | A8JUV9_DROME Shaggy, isoform M OS=Drosophila melanogaster OX=7227 GN=sgg PE=1 SV=1 | 7 | 4 | 1 | 1 | 2.807354922 | 0 | 2.807354922 |
| E2QCN3 | E2QCN3_DROME Alpha/beta hydrolase 1, isoform B OS=Drosophila melanogaster OX=7227 GN=Hydr1 PE=4 SV=1 | 7 | 6 | 1 | 1 | 2.807354922 | 0 | 2.807354922 |
| M9NFR5 | M9NFR5_DROME Ypsilon schachtel, isoform B OS=Drosophila melanogaster OX=7227 GN=yps PE=1 SV=1 | 7 | 3 | 1 | 1 | 2.807354922 | 0 | 2.807354922 |
| M9NG31 | M9NG31_DROME Defective chorion 1, isoform G OS=Drosophila melanogaster OX=7227 GN=dec-1 PE=4 SV=1 | 7 | 5 | 1 | 1 | 2.807354922 | 0 | 2.807354922 |
| M9PE16 | M9PE16_DROME Uncharacterized protein, isoform H OS=Drosophila melanogaster OX=7227 GN=CG4238 PE=4 SV=1 | 7 | 7 | 1 | 1 | 2.807354922 | 0 | 2.807354922 |
| M9PGK1 | M9PGK1_DROME Uncharacterized protein, isoform C OS=Drosophila melanogaster OX=7227 GN=CG3655 PE=4 SV=1 | 7 | 4 | 1 | 1 | 2.807354922 | 0 | 2.807354922 |
| M9PHV4 | M9PHV4_DROME Uncharacterized protein, isoform F OS=Drosophila melanogaster OX=7227 GN=CG42354 PE=1 SV=1 | 7 | 6 | 1 | 1 | 2.807354922 | 0 | 2.807354922 |
| Q0E8E1 | Q0E8E1_DROME GH08173p OS=Drosophila melanogaster OX=7227 GN=AC PE=2 SV=1 | 7 | 3 | 1 | 1 | 2.807354922 | 0 | 2.807354922 |
| Q6AWJ5 | Q6AWJ5_DROME LP12324p OS=Drosophila melanogaster OX=7227 GN=svr PE=1 SV=1 | 7 | 4 | 1 | 1 | 2.807354922 | 0 | 2.807354922 |
| Q7JQ36 | Q7JQ36_DROME Receptor protein serine/threonine kinase OS=Drosophila melanogaster OX=7227 GN=sax PE=2 SV=1 | 7 | 5 | 1 | 1 | 2.807354922 | 0 | 2.807354922 |
| Q7JUS1 | Q7JUS1_DROME AT09608p OS=Drosophila melanogaster OX=7227 GN=CG30491;CG30495 PE=1 SV=1 | 7 | 6 | 1 | 1 | 2.807354922 | 0 | 2.807354922 |
| Q7K4L9 | Q7K4L9_DROME CG8405, isoform A OS=Drosophila melanogaster OX=7227 GN=CG8405 PE=1 SV=1 | 7 | 6 | 1 | 1 | 2.807354922 | 0 | 2.807354922 |
| Q7KTP4 | Q7KTP4_DROME Reticulon-like protein OS=Drosophila melanogaster OX=7227 GN=Rtnl1 PE=1 SV=1 | 7 | 2 | 1 | 1 | 2.807354922 | 0 | 2.807354922 |
| Q7KUL1 | Q7KUL1_DROME Dally-like, isoform C OS=Drosophila melanogaster OX=7227 GN=dlp PE=1 SV=2 | 7 | 6 | 1 | 1 | 2.807354922 | 0 | 2.807354922 |
| Q8IRZ0 | Q8IRZ0_DROME Hexosyltransferase OS=Drosophila melanogaster OX=7227 GN=CG3038-RB PE=2 SV=1 | 7 | 7 | 1 | 1 | 2.807354922 | 0 | 2.807354922 |
| Q9VB20 | Q9VB20_DROME Distracted, isoform B OS=Drosophila melanogaster OX=7227 GN=dsd PE=4 SV=2 | 7 | 6 | 1 | 1 | 2.807354922 | 0 | 2.807354922 |
| Q9VBZ9 | Q9VBZ9_DROME Beta-4-galactosyltransferase 7 OS=Drosophila melanogaster OX=7227 GN=beta4GalT7 PE=1 SV=1 | 7 | 4 | 1 | 1 | 2.807354922 | 0 | 2.807354922 |
| Q9VC58 | Q9VC58_DROME LD37002p OS=Drosophila melanogaster OX=7227 GN=Syx18 PE=1 SV=1 | 7 | 3 | 1 | 1 | 2.807354922 | 0 | 2.807354922 |
| Q9VCF1 | Q9VCF1_DROME GH22341p OS=Drosophila melanogaster OX=7227 GN=Dmel\CG13603 PE=1 SV=1 | 7 | 3 | 1 | 1 | 2.807354922 | 0 | 2.807354922 |
| Q9VEE9 | Q9VEE9_DROME LD22559p OS=Drosophila melanogaster OX=7227 GN=Alg1 PE=1 SV=1 | 7 | 6 | 1 | 1 | 2.807354922 | 0 | 2.807354922 |
| Q9VHB1 | Q9VHB1_DROME Mitochondrial pyruvate carrier OS=Drosophila melanogaster OX=7227 GN=Dmel\CG9399 PE=1 SV=1 | 7 | 6 | 1 | 1 | 2.807354922 | 0 | 2.807354922 |
| Q9VNP7 | Q9VNP7_DROME LD37241p OS=Drosophila melanogaster OX=7227 GN=slif PE=2 SV=1 | 7 | 3 | 1 | 1 | 2.807354922 | 0 | 2.807354922 |
| Q9VTE1 | Q9VTE1_DROME GEO12001p1 OS=Drosophila melanogaster OX=7227 GN=CG6316 PE=2 SV=2 | 7 | 2 | 1 | 1 | 2.807354922 | 0 | 2.807354922 |
| Q9VTF5 | Q9VTF5_DROME RE42071p OS=Drosophila melanogaster OX=7227 GN=Dmel\CG7638 PE=2 SV=2 | 7 | 4 | 1 | 1 | 2.807354922 | 0 | 2.807354922 |
| Q9VUB9 | Q9VUB9_DROME LD24362p OS=Drosophila melanogaster OX=7227 GN=06 PE=1 SV=1 | 7 | 5 | 1 | 1 | 2.807354922 | 0 | 2.807354922 |
| Q9VXA9 | Q9VXA9_DROME RE04047p OS=Drosophila melanogaster OX=7227 GN=Dmel\CG4789 PE=2 SV=2 | 7 | 6 | 1 | 1 | 2.807354922 | 0 | 2.807354922 |
| Q9VYI1 | Q9VYI1_DROME Protein YIPF OS=Drosophila melanogaster OX=7227 GN=Dmel\CG4645 PE=1 SV=2 | 7 | 2 | 1 | 1 | 2.807354922 | 0 | 2.807354922 |
| Q9VYK6 | Q9VYK6_DROME Tomosyn, isoform C OS=Drosophila melanogaster OX=7227 GN=Tomosyn PE=1 SV=3 | 7 | 6 | 1 | 1 | 2.807354922 | 0 | 2.807354922 |
| Q9VYT4 | Q9VYT4_DROME ATP7, isoform B OS=Drosophila melanogaster OX=7227 GN=ATP7 PE=3 SV=3 | 7 | 7 | 1 | 1 | 2.807354922 | 0 | 2.807354922 |
| Q9VYW6 | Q9VYW6_DROME Rhomboid-like protein OS=Drosophila melanogaster OX=7227 GN=rho-4 PE=1 SV=2 | 7 | 5 | 1 | 1 | 2.807354922 | 0 | 2.807354922 |
| X2JCI0 | X2JCI0_DROME Fasciclin 2, isoform H OS=Drosophila melanogaster OX=7227 GN=Fas2 PE=1 SV=1 | 7 | 7 | 1 | 1 | 2.807354922 | 0 | 2.807354922 |
| X2JFG0 | X2JFG0_DROME Rudimentary, isoform D OS=Drosophila melanogaster OX=7227 GN=r PE=1 SV=1 | 6 | 6 | 60 | 36 | 2.584962501 | 5.906890596 | -3.321928095 |
| Q9VL89 | Q9VL89_DROME Methionine aminopeptidase 2 OS=Drosophila melanogaster OX=7227 GN=und PE=1 SV=1 | 6 | 4 | 46 | 12 | 2.584962501 | 5.523561956 | -2.938599455 |
| Q8SXM8 | Q8SXM8_DROME Lysine--tRNA ligase OS=Drosophila melanogaster OX=7227 GN=LysRS PE=1 SV=1 | 6 | 6 | 45 | 19 | 2.584962501 | 5.491853096 | -2.906890596 |
| Q9V434 | Q9V434_DROME Asparaginyl-tRNA synthetase, isoform A OS=Drosophila melanogaster OX=7227 GN=AsnRS PE=1 SV=1 | 6 | 6 | 36 | 17 | 2.584962501 | 5.169925001 | -2.584962501 |
| Q8INM3 | SLE_DROME Protein slender lobes OS=Drosophila melanogaster OX=7227 GN=sle PE=1 SV=1 | 6 | 5 | 29 | 16 | 2.584962501 | 4.857980995 | -2.273018494 |
| Q9VXR5 | Q9VXR5_DROME LD02975p OS=Drosophila melanogaster OX=7227 GN=BEST:LD38280 PE=1 SV=1 | 6 | 5 | 29 | 14 | 2.584962501 | 4.857980995 | -2.273018494 |
| Q7KN90 | SYCC_DROME Cysteine--tRNA ligase, cytoplasmic OS=Drosophila melanogaster OX=7227 GN=CysRS PE=1 SV=1 | 6 | 6 | 28 | 17 | 2.584962501 | 4.807354922 | -2.222392421 |
| Q7PLL6 | Q7PLL6_DROME Uncharacterized protein, isoform A OS=Drosophila melanogaster OX=7227 GN=CG17514 PE=1 SV=2 | 6 | 6 | 27 | 22 | 2.584962501 | 4.754887502 | -2.169925001 |
| Q9VZP5 | Q9VZP5_DROME Eukaryotic translation initiation factor 5B, isoform B OS=Drosophila melanogaster OX=7227 GN=eIF5B PE=1 SV=2 | 6 | 6 | 27 | 15 | 2.584962501 | 4.754887502 | -2.169925001 |
| Q9VGW6 | MCM5_DROME DNA replication licensing factor Mcm5 OS=Drosophila melanogaster OX=7227 GN=Mcm5 PE=1 SV=1 | 6 | 5 | 26 | 16 | 2.584962501 | 4.700439718 | -2.115477217 |
| E1JGL8 | E1JGL8_DROME Uncharacterized protein, isoform C OS=Drosophila melanogaster OX=7227 GN=CG30122 PE=1 SV=1 | 6 | 5 | 26 | 9 | 2.584962501 | 4.700439718 | -2.115477217 |
| A0A0B4LGZ5 | A0A0B4LGZ5_DROME Ribosomal protein L11, isoform B OS=Drosophila melanogaster OX=7227 GN=RpL11 PE=1 SV=1 | 6 | 3 | 24 | 7 | 2.584962501 | 4.584962501 | -2 |
| Q9VNB9 | Q9VNB9_DROME GEO07455p1 OS=Drosophila melanogaster OX=7227 GN=RpL35A PE=1 SV=1 | 6 | 4 | 22 | 8 | 2.584962501 | 4.459431619 | -1.874469118 |
| Q9VQB4 | Q9VQB4_DROME LD06553p OS=Drosophila melanogaster OX=7227 GN=CT12127 PE=1 SV=1 | 6 | 5 | 22 | 10 | 2.584962501 | 4.459431619 | -1.874469118 |
| Q9VTU4 | EIF3L_DROME Eukaryotic translation initiation factor 3 subunit L OS=Drosophila melanogaster OX=7227 GN=eIF3l PE=2 SV=1 | 6 | 6 | 21 | 10 | 2.584962501 | 4.392317423 | -1.807354922 |
| M9PHW7 | M9PHW7_DROME Uncharacterized protein, isoform D OS=Drosophila melanogaster OX=7227 GN=CG6767 PE=1 SV=1 | 6 | 5 | 21 | 10 | 2.584962501 | 4.392317423 | -1.807354922 |
| P54358 | DPOD1_DROME DNA polymerase delta catalytic subunit OS=Drosophila melanogaster OX=7227 GN=DNApol-delta PE=2 SV=2 | 6 | 6 | 20 | 15 | 2.584962501 | 4.321928095 | -1.736965594 |
| Q24524 | SING_DROME Protein singed OS=Drosophila melanogaster OX=7227 GN=sn PE=1 SV=1 | 6 | 4 | 20 | 11 | 2.584962501 | 4.321928095 | -1.736965594 |
| Q24253 | Q24253_DROME AP complex subunit beta OS=Drosophila melanogaster OX=7227 GN=AP-1-2beta PE=1 SV=1 | 6 | 6 | 19 | 13 | 2.584962501 | 4.247927513 | -1.662965013 |
| Q7KN94 | Q7KN94_DROME Walrus, isoform A OS=Drosophila melanogaster OX=7227 GN=wal PE=1 SV=1 | 6 | 5 | 19 | 12 | 2.584962501 | 4.247927513 | -1.662965013 |

| UniProt ID | Protein description | # PSMs MVB spectra | # of unique peptides MVB | # of PSMs lysate | # of unique peptides lysate | log2(PSM MVB) | log2(PSM lysate) | FC (log2 of MVB PSM-log2 of lysate PSM) |
|---|---|---|---|---|---|---|---|---|
| Q9W3J5 | SYFA_DROME Phenylalanine--tRNA ligase alpha subunit OS=Drosophila melanogaster OX=7227 GN=alpha-PheRS PE=1 SV=1 | 6 | 4 | 18 | 10 | 2.584962501 | 4.169925001 | -1.584962501 |
| M9PED6 | M9PED6_DROME Uncharacterized protein, isoform B OS=Drosophila melanogaster OX=7227 GN=CG4603 PE=1 SV=1 | 6 | 4 | 18 | 6 | 2.584962501 | 4.169925001 | -1.584962501 |
| Q9W1G0 | TALDO_DROME Probable transaldolase OS=Drosophila melanogaster OX=7227 GN=Taldo PE=2 SV=2 | 6 | 5 | 17 | 8 | 2.584962501 | 4.087462841 | -1.502500341 |
| Q9VLT7 | Q9VLT7_DROME IP17351p OS=Drosophila melanogaster OX=7227 GN=RpL36A PE=1 SV=2 | 6 | 4 | 17 | 3 | 2.584962501 | 4.087462841 | -1.502500341 |
| P23572 | CDK1_DROME Cyclin-dependent kinase 1 OS=Drosophila melanogaster OX=7227 GN=Cdk1 PE=1 SV=1 | 6 | 6 | 16 | 10 | 2.584962501 | 4 | -1.415037499 |
| Q8SWU7 | OLA1_DROME Obg-like ATPase 1 OS=Drosophila melanogaster OX=7227 GN=CG1354 PE=1 SV=1 | 6 | 6 | 16 | 9 | 2.584962501 | 4 | -1.415037499 |
| Q9VSS2 | SRP68_DROME Signal recognition particle subunit SRP68 OS=Drosophila melanogaster OX=7227 GN=Srp68 PE=2 SV=1 | 6 | 4 | 16 | 12 | 2.584962501 | 4 | -1.415037499 |
| P40304 | PSB1_DROME Proteasome subunit beta type-1 OS=Drosophila melanogaster OX=7227 GN=Prosbeta6 PE=2 SV=2 | 6 | 4 | 15 | 6 | 2.584962501 | 3.906890596 | -1.321928095 |
| A0A0B4KHL2 | A0A0B4KHL2_DROME Coatomer subunit gamma OS=Drosophila melanogaster OX=7227 GN=gammaCOP PE=1 SV=1 | 6 | 6 | 15 | 12 | 2.584962501 | 3.906890596 | -1.321928095 |
| Q9V466 | NU107_DROME Nuclear pore complex protein Nup107 OS=Drosophila melanogaster OX=7227 GN=Nup107 PE=1 SV=1 | 6 | 6 | 14 | 11 | 2.584962501 | 3.807354922 | -1.222392421 |
| Q9VRP3 | Q9VRP3_DROME AT08565p OS=Drosophila melanogaster OX=7227 GN=Txl PE=1 SV=1 | 6 | 4 | 14 | 7 | 2.584962501 | 3.807354922 | -1.222392421 |
| P91926 | AP2A_DROME AP-2 complex subunit alpha OS=Drosophila melanogaster OX=7227 GN=AP-2alpha PE=1 SV=1 | 6 | 6 | 13 | 11 | 2.584962501 | 3.700439718 | -1.115477217 |
| Q01637 | UMPS_DROME Uridine 5'-monophosphate synthase OS=Drosophila melanogaster OX=7227 GN=r-l PE=2 SV=2 | 6 | 4 | 13 | 8 | 2.584962501 | 3.700439718 | -1.115477217 |
| Q7KSM5 | Q7KSM5_DROME Activator of SUMO 1 OS=Drosophila melanogaster OX=7227 GN=Aos1 PE=1 SV=1 | 6 | 4 | 13 | 9 | 2.584962501 | 3.700439718 | -1.115477217 |
| X2J6D4 | X2J6D4_DROME Cytochrome c proximal, isoform B OS=Drosophila melanogaster OX=7227 GN=Cyt-c-p PE=2 SV=1 | 6 | 3 | 12 | 3 | 2.584962501 | 3.584962501 | -1 |
| P22817 | IDE_DROME Insulin-degrading enzyme OS=Drosophila melanogaster OX=7227 GN=Ide PE=1 SV=4 | 6 | 5 | 11 | 7 | 2.584962501 | 3.459431619 | -0.874469118 |
| E1JIR3 | E1JIR3_DROME Cyclin-dependent kinase 2, isoform C OS=Drosophila melanogaster OX=7227 GN=Cdk2 PE=1 SV=1 | 6 | 4 | 11 | 6 | 2.584962501 | 3.459431619 | -0.874469118 |
| Q8IPP8 | Q8IPP8_DROME Uncharacterized protein OS=Drosophila melanogaster OX=7227 GN=CG31548 PE=1 SV=1 | 6 | 4 | 11 | 6 | 2.584962501 | 3.459431619 | -0.874469118 |
| Q95SI7 | Q95SI7_DROME GH23390p OS=Drosophila melanogaster OX=7227 GN=Dmel\CG6028 PE=1 SV=1 | 6 | 5 | 10 | 6 | 2.584962501 | 3.321928095 | -0.736965594 |
| Q05856 | RSMB_DROME Small nuclear ribonucleoprotein-associated protein B OS=Drosophila melanogaster OX=7227 GN=SmB PE=1 SV=1 | 6 | 4 | 9 | 4 | 2.584962501 | 3.169925001 | -0.584962501 |
| Q7K1Y4 | CIAO1_DROME Probable cytosolic iron-sulfur protein assembly protein Ciao1 OS=Drosophila melanogaster OX=7227 GN=Ciao1 PE=1 SV=1 | 6 | 4 | 9 | 6 | 2.584962501 | 3.169925001 | -0.584962501 |
| Q9VI10 | SMD2_DROME Probable small nuclear ribonucleoprotein Sm D2 OS=Drosophila melanogaster OX=7227 GN=SmD2 PE=1 SV=1 | 6 | 4 | 9 | 5 | 2.584962501 | 3.169925001 | -0.584962501 |
| Q9VJY6 | RL24_DROME 60S ribosomal protein L24 OS=Drosophila melanogaster OX=7227 GN=RpL24 PE=1 SV=1 | 6 | 4 | 9 | 6 | 2.584962501 | 3.169925001 | -0.584962501 |
| A8JRC1 | A8JRC1_DROME Uncharacterized protein, isoform D OS=Drosophila melanogaster OX=7227 GN=CG5028 PE=1 SV=1 | 6 | 4 | 9 | 6 | 2.584962501 | 3.169925001 | -0.584962501 |
| Q9VQI7 | Q9VQI7_DROME Peroxiredoxin 6005 OS=Drosophila melanogaster OX=7227 GN=Prx6005 PE=1 SV=1 | 6 | 5 | 9 | 7 | 2.584962501 | 3.169925001 | -0.584962501 |
| Q9VRD4 | Q9VRD4_DROME Uncharacterized protein, isoform A OS=Drosophila melanogaster OX=7227 GN=Dmel\CG1532 PE=1 SV=1 | 6 | 5 | 9 | 6 | 2.584962501 | 3.169925001 | -0.584962501 |
| P07182 | CH36_DROME Chorion protein S36 OS=Drosophila melanogaster OX=7227 GN=Cp36 PE=3 SV=2 | 6 | 5 | 8 | 5 | 2.584962501 | 3 | -0.415037499 |
| Q9VRJ4 | Q9VRJ4_DROME SD02021p OS=Drosophila melanogaster OX=7227 GN=DHRS4 PE=1 SV=1 | 6 | 3 | 8 | 7 | 2.584962501 | 3 | -0.415037499 |
| P02283 | H2B_DROME Histone H2B OS=Drosophila melanogaster OX=7227 GN=His2B PE=1 SV=2 | 6 | 2 | 7 | 3 | 2.584962501 | 2.807354922 | -0.222392421 |
| M9NEW1 | M9NEW1_DROME Myosin light chain cytoplasmic, isoform B OS=Drosophila melanogaster OX=7227 GN=Mlc-c PE=4 SV=1 | 6 | 4 | 7 | 4 | 2.584962501 | 2.807354922 | -0.222392421 |
| Q8IQC1 | TVP23_DROME Uncharacterized Golgi apparatus membrane protein-like protein CG5021 OS=Drosophila melanogaster OX=7227 GN=CG5021 PE=2 SV=2 | 6 | 2 | 2 | 2 | 2.584962501 | 1 | 1.584962501 |
| Q9VH39 | NDUF4_DROME Protein NDUFAF4 homolog OS=Drosophila melanogaster OX=7227 GN=CG11722 PE=2 SV=1 | 6 | 5 | 2 | 2 | 2.584962501 | 1 | 1.584962501 |
| Q9VLP3 | MSS4_DROME Guanine nucleotide exchange factor MSS4 homolog OS=Drosophila melanogaster OX=7227 GN=strat PE=1 SV=2 | 6 | 4 | 4 | 3 | 2.584962501 | 2 | 0.584962501 |
| Q9VMY1 | RM24_DROME Probable 39S ribosomal protein L24, mitochondrial OS=Drosophila melanogaster OX=7227 GN=mRpL24 PE=2 SV=1 | 6 | 6 | 4 | 2 | 2.584962501 | 2 | 0.584962501 |
| Q9VQ62 | NPC2_DROME NPC intracellular cholesterol transporter 2 homolog a OS=Drosophila melanogaster OX=7227 GN=Npc2a PE=2 SV=1 | 6 | 2 | 2 | 1 | 2.584962501 | 1 | 1.584962501 |
| Q9VZL6 | LTOR4_DROME Ragulator complex protein LAMTOR4 homolog OS=Drosophila melanogaster OX=7227 GN=CG14977 PE=1 SV=2 | 6 | 2 | 1 | 1 | 2.584962501 | 0 | 2.584962501 |
| Q9W3V8 | ALG8_DROME Probable dolichyl pyrophosphate Glc1Man9GlcNAc2 alpha-1,3-glucosyltransferase OS=Drosophila melanogaster OX=7227 GN=xit PE=2 SV=1 | 6 | 5 | 1 | 1 | 2.584962501 | 0 | 2.584962501 |
| Q9Y1A3 | TIM8_DROME Mitochondrial import inner membrane translocase subunit Tim8 OS=Drosophila melanogaster OX=7227 GN=Tim8 PE=3 SV=1 | 6 | 4 | 1 | 1 | 2.584962501 | 0 | 2.584962501 |
| E1JIL1 | E1JIL1_DROME Uncharacterized protein, isoform D OS=Drosophila melanogaster OX=7227 GN=CG31303 PE=1 SV=1 | 6 | 3 | 1 | 1 | 2.584962501 | 0 | 2.584962501 |
| Q9V3L7 | Q9V3L7_DROME GEO04710p1 OS=Drosophila melanogaster OX=7227 GN=NP15.6 PE=1 SV=1 | 6 | 4 | 4 | 3 | 2.584962501 | 2 | 0.584962501 |
| Q9VAP1 | Q9VAP1_DROME Uncharacterized protein OS=Drosophila melanogaster OX=7227 GN=CG14512 PE=4 SV=1 | 6 | 5 | 2 | 1 | 2.584962501 | 1 | 1.584962501 |
| Q9VNC1 | Q9VNC1_DROME Mitochondrial ribosomal protein L44 OS=Drosophila melanogaster OX=7227 GN=mRpL44 PE=1 SV=1 | 6 | 6 | 5 | 5 | 2.584962501 | 2.321928095 | 0.263034406 |
| Q9VR96 | Q9VR96_DROME IP02009p OS=Drosophila melanogaster OX=7227 GN=Obp19c PE=2 SV=1 | 6 | 4 | 4 | 2 | 2.584962501 | 2 | 0.584962501 |
| Q9VXH7 | Q9VXH7_DROME FI17510p1 OS=Drosophila melanogaster OX=7227 GN=CG9917-RA PE=1 SV=1 | 6 | 4 | 2 | 2 | 2.584962501 | 1 | 1.584962501 |
| Q9W0M0 | Q9W0M0_DROME Uncharacterized protein OS=Drosophila melanogaster OX=7227 GN=CG13890 PE=1 SV=1 | 6 | 4 | 4 | 2 | 2.584962501 | 2 | 0.584962501 |
| Q9W2E8 | Q9W2E8_DROME NADH dehydrogenase (Ubiquinone) B12 subunit, isoform A OS=Drosophila melanogaster OX=7227 GN=ND-B12 PE=2 SV=1 | 6 | 3 | 3 | 2 | 2.584962501 | 1.584962501 | 1 |
| Q9VWL3 | Q9VWL3_DROME Uncharacterized protein OS=Drosophila melanogaster OX=7227 GN=CG7453 PE=1 SV=2 | 6 | 3 | 1 | 1 | 2.584962501 | 0 | 2.584962501 |
| P38040 | GBG1_DROME Guanine nucleotide-binding protein subunit gamma-1 OS=Drosophila melanogaster OX=7227 GN=Ggamma1 PE=2 SV=1 | 6 | 3 | 1 | 1 | 2.584962501 | 0 | 2.584962501 |
| Q7KVA1 | XYLT_DROME Xylosyltransferase oxt OS=Drosophila melanogaster OX=7227 GN=oxt PE=2 SV=1 | 6 | 5 | 1 | 1 | 2.584962501 | 0 | 2.584962501 |
| Q8SZ63 | GOGA5_DROME Golgin-84 OS=Drosophila melanogaster OX=7227 GN=Golgin84 PE=1 SV=2 | 6 | 4 | 1 | 1 | 2.584962501 | 0 | 2.584962501 |
| Q9V492 | MPV17_DROME Mpv17-like protein OS=Drosophila melanogaster OX=7227 GN=CG11077 PE=2 SV=1 | 6 | 4 | 1 | 1 | 2.584962501 | 0 | 2.584962501 |
| Q9VAS7 | INX3_DROME Innexin inx3 OS=Drosophila melanogaster OX=7227 GN=Inx3 PE=1 SV=1 | 6 | 5 | 1 | 1 | 2.584962501 | 0 | 2.584962501 |
| Q9VDD1 | DDRGK_DROME DDRGK domain-containing protein 1 OS=Drosophila melanogaster OX=7227 GN=CG5862 PE=2 SV=1 | 6 | 3 | 1 | 1 | 2.584962501 | 0 | 2.584962501 |
| Q9VRX6 | INX4_DROME Innexin inx4 OS=Drosophila melanogaster OX=7227 GN=zpg PE=2 SV=1 | 6 | 3 | 1 | 1 | 2.584962501 | 0 | 2.584962501 |
| Q9VSL7 | FOI_DROME Zinc transporter foi OS=Drosophila melanogaster OX=7227 GN=foi PE=1 SV=3 | 6 | 4 | 1 | 1 | 2.584962501 | 0 | 2.584962501 |
| Q9W589 | OFUT2_DROME GDP-fucose protein O-fucosyltransferase 2 OS=Drosophila melanogaster OX=7227 GN=O-fut2 PE=1 SV=1 | 6 | 5 | 1 | 1 | 2.584962501 | 0 | 2.584962501 |
| Q9XZ08 | EXT3_DROME Exostosin-3 OS=Drosophila melanogaster OX=7227 GN=botv PE=1 SV=1 | 6 | 6 | 1 | 1 | 2.584962501 | 0 | 2.584962501 |
| A0A0B4KG54 | A0A0B4KG54_DROME S-methyl-5'-thioadenosine phosphorylase OS=Drosophila melanogaster OX=7227 GN=Mtap PE=3 SV=1 | 6 | 5 | 1 | 1 | 2.584962501 | 0 | 2.584962501 |
| A0A0B4KGQ0 | A0A0B4KGQ0_DROME Ran binding protein 3, isoform B OS=Drosophila melanogaster OX=7227 GN=RanBP3 PE=1 SV=1 | 6 | 4 | 1 | 1 | 2.584962501 | 0 | 2.584962501 |
| A0A0B4KGQ4 | A0A0B4KGQ4_DROME Easter, isoform B OS=Drosophila melanogaster OX=7227 GN=ea PE=3 SV=1 | 6 | 5 | 1 | 1 | 2.584962501 | 0 | 2.584962501 |
| A0A0B4KGV7 | A0A0B4KGV7_DROME Uncharacterized protein, isoform B OS=Drosophila melanogaster OX=7227 GN=CG13601 PE=4 SV=1 | 6 | 4 | 1 | 1 | 2.584962501 | 0 | 2.584962501 |
| A0A0B4LGL6 | A0A0B4LGL6_DROME Unzipped, isoform B OS=Drosophila melanogaster OX=7227 GN=uzip PE=4 SV=1 | 6 | 4 | 1 | 1 | 2.584962501 | 0 | 2.584962501 |
| A0A0C4DHA6 | A0A0C4DHA6_DROME Curled, isoform G OS=Drosophila melanogaster OX=7227 GN=cu PE=4 SV=1 | 6 | 5 | 1 | 1 | 2.584962501 | 0 | 2.584962501 |
| A0A0S0X0Z7 | A0A0S0X0Z7_DROME Protein tweety homolog OS=Drosophila melanogaster OX=7227 GN=CG3638 PE=3 SV=1 | 6 | 4 | 1 | 1 | 2.584962501 | 0 | 2.584962501 |
| B7YZW3 | B7YZW3_DROME Uncharacterized protein, isoform E OS=Drosophila melanogaster OX=7227 GN=CG42389 PE=1 SV=1 | 6 | 5 | 1 | 1 | 2.584962501 | 0 | 2.584962501 |
| M9PCX4 | M9PCX4_DROME Uncharacterized protein, isoform E OS=Drosophila melanogaster OX=7227 GN=CG3961 PE=1 SV=1 | 6 | 4 | 1 | 1 | 2.584962501 | 0 | 2.584962501 |
| M9PG98 | M9PG98_DROME alpha-1,2-Mannosidase OS=Drosophila melanogaster OX=7227 GN=Edem1 PE=1 SV=1 | 6 | 6 | 1 | 1 | 2.584962501 | 0 | 2.584962501 |
| M9PGG4 | M9PGG4_DROME Armadillo, isoform F OS=Drosophila melanogaster OX=7227 GN=arm PE=4 SV=1 | 6 | 5 | 1 | 1 | 2.584962501 | 0 | 2.584962501 |
| Q3ZAN1 | Q3ZAN1_DROME Golgi pH regulator OS=Drosophila melanogaster OX=7227 GN=GPHR PE=2 SV=1 | 6 | 4 | 1 | 1 | 2.584962501 | 0 | 2.584962501 |
| Q6IGN6 | Q6IGN6_DROME HDC05827 OS=Drosophila melanogaster OX=7227 GN=CG34228 PE=1 SV=1 | 6 | 4 | 1 | 1 | 2.584962501 | 0 | 2.584962501 |
| Q6NP91 | Q6NP91_DROME RE44586p OS=Drosophila melanogaster OX=7227 GN=CG11210 PE=1 SV=1 | 6 | 5 | 1 | 1 | 2.584962501 | 0 | 2.584962501 |
| Q7JYH3 | Q7JYH3_DROME NADH dehydrogenase (Ubiquinone) B14.7 subunit OS=Drosophila melanogaster OX=7227 GN=ND-B14.7 PE=1 SV=1 | 6 | 2 | 1 | 1 | 2.584962501 | 0 | 2.584962501 |
| Q7JZZ3 | Q7JZZ3_DROME RE03883p OS=Drosophila melanogaster OX=7227 GN=CG13321 PE=1 SV=1 | 6 | 5 | 1 | 1 | 2.584962501 | 0 | 2.584962501 |
| Q7K579 | Q7K579_DROME Diacylglycerol kinase OS=Drosophila melanogaster OX=7227 GN=Dgkepsilon PE=1 SV=1 | 6 | 5 | 1 | 1 | 2.584962501 | 0 | 2.584962501 |
| Q7K8Y5 | Q7K8Y5_DROME Serpin 4 OS=Drosophila melanogaster OX=7227 GN=Spn42Da PE=1 SV=1 | 6 | 6 | 1 | 1 | 2.584962501 | 0 | 2.584962501 |
| Q8IP48 | Q8IP48_DROME Zinc transporter 35C, isoform B OS=Drosophila melanogaster OX=7227 GN=ZnT35C PE=4 SV=1 | 6 | 4 | 1 | 1 | 2.584962501 | 0 | 2.584962501 |
| Q8IQQ7 | Q8IQQ7_DROME Bloated tubules, isoform B OS=Drosophila melanogaster OX=7227 GN=blot PE=2 SV=1 | 6 | 4 | 1 | 1 | 2.584962501 | 0 | 2.584962501 |
| Q8SWX4 | Q8SWX4_DROME Aminopeptidase OS=Drosophila melanogaster OX=7227 GN=CG5839 PE=1 SV=1 | 6 | 6 | 1 | 1 | 2.584962501 | 0 | 2.584962501 |
| Q94515 | Q94515_DROME Des-1 protein OS=Drosophila melanogaster OX=7227 GN=ifc PE=1 SV=1 | 6 | 3 | 1 | 1 | 2.584962501 | 0 | 2.584962501 |
| Q960K3 | Q960K3_DROME LD47387p OS=Drosophila melanogaster OX=7227 GN=Dmel\CG1103 PE=2 SV=1 | 6 | 2 | 1 | 1 | 2.584962501 | 0 | 2.584962501 |
| Q961B9 | Q961B9_DROME LD24073p OS=Drosophila melanogaster OX=7227 GN=Dmel\CG5027 PE=2 SV=1 | 6 | 6 | 1 | 1 | 2.584962501 | 0 | 2.584962501 |

| UniProt ID | Protein description | # PSMs MVB spectra | # of unique peptides MVB | # of PSMs lysate | # of unique peptides lysate | log2(PSM MVB) | log2(PSM lysate) | FC (log2 of MVB PSM-log2 of lysate PSM) |
|---|---|---|---|---|---|---|---|---|
| Q9VEJ4 | Q9VEJ4_DROME HL05730p OS=Drosophila melanogaster OX=7227 GN=Dmel\CG5823 PE=1 SV=2 | 6 | 3 | 1 | 1 | 2.584962501 | 0 | 2.584962501 |
| Q9VG14 | Q9VG14_DROME Uncharacterized protein OS=Drosophila melanogaster OX=7227 GN=CG7091 PE=4 SV=3 | 6 | 3 | 1 | 1 | 2.584962501 | 0 | 2.584962501 |
| Q9VGF0 | Q9VGF0_DROME Midline fasciclin, isoform C OS=Drosophila melanogaster OX=7227 GN=mfas PE=1 SV=1 | 6 | 4 | 1 | 1 | 2.584962501 | 0 | 2.584962501 |
| Q9VIG6 | Q9VIG6_DROME FI23927p1 OS=Drosophila melanogaster OX=7227 GN=Dmel\CG9265 PE=2 SV=3 | 6 | 5 | 1 | 1 | 2.584962501 | 0 | 2.584962501 |
| Q9VIL1 | Q9VIL1_DROME GEO05273p1 OS=Drosophila melanogaster OX=7227 GN=38D.27 PE=2 SV=2 | 6 | 2 | 1 | 1 | 2.584962501 | 0 | 2.584962501 |
| Q9VLP5 | Q9VLP5_DROME LD42916p OS=Drosophila melanogaster OX=7227 GN=Dme_CG7806 PE=1 SV=1 | 6 | 5 | 1 | 1 | 2.584962501 | 0 | 2.584962501 |
| Q9VMR0 | Q9VMR0_DROME GM05057p OS=Drosophila melanogaster OX=7227 GN=Dmel\CG7382 PE=1 SV=1 | 6 | 4 | 1 | 1 | 2.584962501 | 0 | 2.584962501 |
| Q9VPR1 | Q9VPR1_DROME GH02075p OS=Drosophila melanogaster OX=7227 GN=Tspo PE=1 SV=1 | 6 | 3 | 1 | 1 | 2.584962501 | 0 | 2.584962501 |
| Q9VQZ1 | Q9VQZ1_DROME Lethal (2) k05819, isoform A OS=Drosophila melanogaster OX=7227 GN=l(2)k05819 PE=2 SV=3 | 6 | 3 | 1 | 1 | 2.584962501 | 0 | 2.584962501 |
| Q9VU37 | Q9VU37_DROME Protein-S-isoprenylcysteine O-methyltransferase OS=Drosophila melanogaster OX=7227 GN=ste14 PE=2 SV=1 | 6 | 4 | 1 | 1 | 2.584962501 | 0 | 2.584962501 |
| Q9VWJ1 | Q9VWJ1_DROME Uncharacterized protein, isoform A OS=Drosophila melanogaster OX=7227 GN=CG8034 PE=1 SV=2 | 6 | 4 | 1 | 1 | 2.584962501 | 0 | 2.584962501 |
| Q9VXU7 | Q9VXU7_DROME IP07261p OS=Drosophila melanogaster OX=7227 GN=Alg14 PE=2 SV=1 | 6 | 4 | 1 | 1 | 2.584962501 | 0 | 2.584962501 |
| Q9VZD9 | Q9VZD9_DROME Peroxisomal membrane protein 34, isoform A OS=Drosophila melanogaster OX=7227 GN=PMP34 PE=3 SV=2 | 6 | 3 | 1 | 1 | 2.584962501 | 0 | 2.584962501 |
| Q9W0H8 | Q9W0H8_DROME IP16409p OS=Drosophila melanogaster OX=7227 GN=scf PE=1 SV=2 | 6 | 5 | 1 | 1 | 2.584962501 | 0 | 2.584962501 |
| Q9W388 | Q9W388_DROME AP-1 complex subunit gamma OS=Drosophila melanogaster OX=7227 GN=AP-1gamma PE=1 SV=2 | 6 | 6 | 1 | 1 | 2.584962501 | 0 | 2.584962501 |
| Q9W3M4 | Q9W3M4_DROME LD24308p OS=Drosophila melanogaster OX=7227 GN=l(1)G0193 PE=1 SV=1 | 6 | 4 | 1 | 1 | 2.584962501 | 0 | 2.584962501 |
| Q9W5X0 | Q9W5X0_DROME LD21953p OS=Drosophila melanogaster OX=7227 GN=Rab35 PE=1 SV=1 | 6 | 6 | 1 | 1 | 2.584962501 | 0 | 2.584962501 |
| Q9XZ28 | Q9XZ28_DROME Uncharacterized protein, isoform A OS=Drosophila melanogaster OX=7227 GN=BcDNA:LD21794 PE=2 SV=1 | 6 | 6 | 1 | 1 | 2.584962501 | 0 | 2.584962501 |
| Q9VUJ1 | Q9VUJ1_DROME Proteasome subunit beta type OS=Drosophila melanogaster OX=7227 GN=Prosbeta2 PE=1 SV=1 | 5 | 3 | 38 | 12 | 2.321928095 | 5.247927513 | -2.925999419 |
| A1ZBE9 | A1ZBE9_DROME Methionyl-tRNA synthetase OS=Drosophila melanogaster OX=7227 GN=MetRS PE=1 SV=1 | 5 | 5 | 35 | 22 | 2.321928095 | 5.129283017 | -2.807354922 |
| A8DY81 | A8DY81_DROME Not1, isoform C OS=Drosophila melanogaster OX=7227 GN=Not1 PE=1 SV=1 | 5 | 5 | 34 | 27 | 2.321928095 | 5.087462841 | -2.765534746 |
| O02195 | EIF3I_DROME Eukaryotic translation initiation factor 3 subunit I OS=Drosophila melanogaster OX=7227 GN=eIF3i PE=1 SV=1 | 5 | 3 | 33 | 12 | 2.321928095 | 5.044394119 | -2.722466024 |
| P55035 | PSMD4_DROME 26S proteasome non-ATPase regulatory subunit 4 OS=Drosophila melanogaster OX=7227 GN=Rpn10 PE=1 SV=2 | 5 | 5 | 31 | 12 | 2.321928095 | 4.95419631 | -2.632268215 |
| Q9VUV9 | U520_DROME Putative U5 small nuclear ribonucleoprotein 200 kDa helicase OS=Drosophila melanogaster OX=7227 GN=l(3)72Ab PE=2 SV=4 | 5 | 4 | 30 | 20 | 2.321928095 | 4.906890596 | -2.584962501 |
| Q9GQR5 | Q9GQR5_DROME RH02809p OS=Drosophila melanogaster OX=7227 GN=stai PE=1 SV=1 | 5 | 3 | 29 | 7 | 2.321928095 | 4.857980995 | -2.5360529 |
| M9PD27 | M9PD27_DROME Eukaryotic translation release factor 1, isoform H OS=Drosophila melanogaster OX=7227 GN=eRF1 PE=1 SV=1 | 5 | 4 | 28 | 11 | 2.321928095 | 4.807354922 | -2.485426827 |
| M9PF24 | M9PF24_DROME Kinesin light chain, isoform B OS=Drosophila melanogaster OX=7227 GN=Klc PE=4 SV=1 | 5 | 5 | 27 | 16 | 2.321928095 | 4.754887502 | -2.432959407 |
| Q9VWG3 | RS10B_DROME 40S ribosomal protein S10b OS=Drosophila melanogaster OX=7227 GN=RpS10b PE=1 SV=2 | 5 | 4 | 24 | 7 | 2.321928095 | 4.584962501 | -2.263034406 |
| A8JNJ6 | A8JNJ6_DROME Karst, isoform E OS=Drosophila melanogaster OX=7227 GN=kst PE=1 SV=2 | 5 | 5 | 24 | 23 | 2.321928095 | 4.584962501 | -2.263034406 |
| M9MRQ9 | M9MRQ9_DROME Succinate--CoA ligase [ADP/GDP-forming] subunit alpha, mitochondrial OS=Drosophila melanogaster OX=7227 GN=Scsalpha1 PE=3 SV=1 | 5 | 4 | 24 | 8 | 2.321928095 | 4.584962501 | -2.263034406 |
| Q9VWD9 | Q9VWD9_DROME LD38919p OS=Drosophila melanogaster OX=7227 GN=Ubqn PE=1 SV=1 | 5 | 3 | 24 | 14 | 2.321928095 | 4.584962501 | -2.263034406 |
| O77410 | EIF3E_DROME Eukaryotic translation initiation factor 3 subunit E OS=Drosophila melanogaster OX=7227 GN=eIF3e PE=1 SV=1 | 5 | 4 | 23 | 9 | 2.321928095 | 4.523561956 | -2.201633861 |
| Q9VXN4 | SYRC_DROME Probable arginine--tRNA ligase, cytoplasmic OS=Drosophila melanogaster OX=7227 GN=ArgRS PE=2 SV=1 | 5 | 5 | 23 | 16 | 2.321928095 | 4.523561956 | -2.201633861 |
| A0A0B4KHJ5 | A0A0B4KHJ5_DROME Glycogen [starch] synthase OS=Drosophila melanogaster OX=7227 GN=GlyS PE=1 SV=1 | 5 | 3 | 23 | 13 | 2.321928095 | 4.523561956 | -2.201633861 |
| Q9VMQ9 | Q9VMQ9_DROME Clot protein OS=Drosophila melanogaster OX=7227 GN=cl PE=1 SV=1 | 5 | 5 | 20 | 9 | 2.321928095 | 4.321928095 | -2 |
| P34739 | TTF2_DROME Transcription termination factor 2 OS=Drosophila melanogaster OX=7227 GN=lds PE=1 SV=2 | 5 | 5 | 18 | 13 | 2.321928095 | 4.169925001 | -1.847996907 |
| A1Z9E3 | A1Z9E3_DROME Elongation factor Tu OS=Drosophila melanogaster OX=7227 GN=mEFTu1 PE=1 SV=1 | 5 | 5 | 18 | 12 | 2.321928095 | 4.169925001 | -1.847996907 |
| Q0E965 | Q0E965_DROME SR protein kinase, isoform A OS=Drosophila melanogaster OX=7227 GN=SRPK PE=1 SV=1 | 5 | 4 | 18 | 14 | 2.321928095 | 4.169925001 | -1.847996907 |
| Q9V436 | Q9V436_DROME 26S proteasome regulatory complex subunit p30 OS=Drosophila melanogaster OX=7227 GN=Rpn12 PE=1 SV=1 | 5 | 5 | 18 | 10 | 2.321928095 | 4.169925001 | -1.847996907 |
| Q9VCW3 | NU133_DROME Nuclear pore complex protein Nup133 OS=Drosophila melanogaster OX=7227 GN=Nup133 PE=2 SV=2 | 5 | 5 | 17 | 10 | 2.321928095 | 4.087462841 | -1.765534746 |
| Q7K5K3 | Q7K5K3_DROME Pyruvate dehydrogenase E1 component subunit beta OS=Drosophila melanogaster OX=7227 GN=Pdhb PE=1 SV=1 | 5 | 5 | 17 | 9 | 2.321928095 | 4.087462841 | -1.765534746 |
| Q9W229 | Q9W229_DROME 40S ribosomal protein S24 OS=Drosophila melanogaster OX=7227 GN=RpS24 PE=1 SV=1 | 5 | 3 | 17 | 6 | 2.321928095 | 4.087462841 | -1.765534746 |
| P18053 | PSA4_DROME Proteasome subunit alpha type-4 OS=Drosophila melanogaster OX=7227 GN=Prosalpha3 PE=1 SV=2 | 5 | 3 | 16 | 6 | 2.321928095 | 4 | -1.678071905 |
| H8F4T6 | H8F4T6_DROME FI19911p1 OS=Drosophila melanogaster OX=7227 GN=CG16935-RA PE=1 SV=1 | 5 | 5 | 16 | 10 | 2.321928095 | 4 | -1.678071905 |
| Q9V3V9 | Q9V3V9_DROME Endonuclease G inhibitor, isoform A OS=Drosophila melanogaster OX=7227 GN=EndoGI PE=1 SV=1 | 5 | 4 | 16 | 9 | 2.321928095 | 4 | -1.678071905 |
| Q9VD58 | Q9VD58_DROME GH26270p OS=Drosophila melanogaster OX=7227 GN=Idh3b PE=1 SV=1 | 5 | 5 | 16 | 10 | 2.321928095 | 4 | -1.678071905 |
| Q94899 | CSN2_DROME COP9 signalosome complex subunit 2 OS=Drosophila melanogaster OX=7227 GN=alien PE=1 SV=2 | 5 | 5 | 15 | 10 | 2.321928095 | 3.906890596 | -1.584962501 |
| Q9W401 | CISY_DROME Probable citrate synthase, mitochondrial OS=Drosophila melanogaster OX=7227 GN=kdn PE=2 SV=1 | 5 | 5 | 15 | 10 | 2.321928095 | 3.906890596 | -1.584962501 |
| Q8I8U7 | TRA1_DROME Transcription-associated protein 1 OS=Drosophila melanogaster OX=7227 GN=Nipped-A PE=1 SV=4 | 5 | 4 | 14 | 12 | 2.321928095 | 3.807354922 | -1.485426827 |
| Q8T3U2 | RS23_DROME 40S ribosomal protein S23 OS=Drosophila melanogaster OX=7227 GN=RpS23 PE=1 SV=1 | 5 | 3 | 14 | 6 | 2.321928095 | 3.807354922 | -1.485426827 |
| A1ZBF1 | A1ZBF1_DROME FI16628p1 OS=Drosophila melanogaster OX=7227 GN=CG18190-RB PE=1 SV=2 | 5 | 5 | 14 | 11 | 2.321928095 | 3.807354922 | -1.485426827 |
| O77266 | O77266_DROME Uncharacterized protein, isoform A OS=Drosophila melanogaster OX=7227 GN=CG4199 PE=1 SV=3 | 5 | 5 | 14 | 10 | 2.321928095 | 3.807354922 | -1.485426827 |
| Q7JUP3 | Q7JUP3_DROME Caspar, isoform A OS=Drosophila melanogaster OX=7227 GN=casp PE=1 SV=1 | 5 | 5 | 14 | 9 | 2.321928095 | 3.807354922 | -1.485426827 |
| Q7K0B6 | Q7K0B6_DROME Glutathione S transferase T1 OS=Drosophila melanogaster OX=7227 GN=GstT1 PE=1 SV=1 | 5 | 5 | 14 | 9 | 2.321928095 | 3.807354922 | -1.485426827 |
| Q9VYW3 | Q9VYW3_DROME GEO08380p1 OS=Drosophila melanogaster OX=7227 GN=RPA3 PE=1 SV=1 | 5 | 5 | 14 | 7 | 2.321928095 | 3.807354922 | -1.485426827 |
| Q9W3C3 | Q9W3C3_DROME LD10016p OS=Drosophila melanogaster OX=7227 GN=Dmel\CG2004 PE=1 SV=1 | 5 | 4 | 14 | 7 | 2.321928095 | 3.807354922 | -1.485426827 |
| P05205 | HP1_DROME Heterochromatin protein 1 OS=Drosophila melanogaster OX=7227 GN=Su(var)205 PE=1 SV=2 | 5 | 3 | 13 | 7 | 2.321928095 | 3.700439718 | -1.378511623 |
| P56538 | IF6_DROME Eukaryotic translation initiation factor 6 OS=Drosophila melanogaster OX=7227 GN=eIF6 PE=2 SV=3 | 5 | 3 | 13 | 7 | 2.321928095 | 3.700439718 | -1.378511623 |
| A1Z7J6 | A1Z7J6_DROME Sec31 ortholog, isoform B OS=Drosophila melanogaster OX=7227 GN=Sec31 PE=1 SV=1 | 5 | 4 | 13 | 9 | 2.321928095 | 3.700439718 | -1.378511623 |
| O76752 | O76752_DROME CG12117-PA OS=Drosophila melanogaster OX=7227 GN=Sptr PE=1 SV=1 | 5 | 3 | 13 | 7 | 2.321928095 | 3.700439718 | -1.378511623 |
| Q9VEP6 | Q9VEP6_DROME Adenylosuccinate lyase OS=Drosophila melanogaster OX=7227 GN=AdSL PE=1 SV=1 | 5 | 3 | 13 | 10 | 2.321928095 | 3.700439718 | -1.378511623 |
| P42271 | TBG2_DROME Tubulin gamma-2 chain OS=Drosophila melanogaster OX=7227 GN=gammaTub37C PE=1 SV=3 | 5 | 5 | 12 | 10 | 2.321928095 | 3.584962501 | -1.263034406 |
| Q9Y0Y2 | PURA_DROME Adenylosuccinate synthetase OS=Drosophila melanogaster OX=7227 GN=AdSS PE=2 SV=1 | 5 | 3 | 12 | 7 | 2.321928095 | 3.584962501 | -1.263034406 |
| A0A0C4DHG2 | A0A0C4DHG2_DROME Gelsolin, isoform J OS=Drosophila melanogaster OX=7227 GN=Gel PE=4 SV=1 | 5 | 3 | 12 | 9 | 2.321928095 | 3.584962501 | -1.263034406 |
| M9PBK5 | M9PBK5_DROME Ribosomal protein LP1, isoform B OS=Drosophila melanogaster OX=7227 GN=RpLP1 PE=3 SV=1 | 5 | 2 | 12 | 4 | 2.321928095 | 3.584962501 | -1.263034406 |
| Q9VQ78 | Q9VQ78_DROME LD16031p OS=Drosophila melanogaster OX=7227 GN=TBCD PE=1 SV=1 | 5 | 3 | 12 | 7 | 2.321928095 | 3.584962501 | -1.263034406 |
| P55828 | RS20_DROME 40S ribosomal protein S20 OS=Drosophila melanogaster OX=7227 GN=RpS20 PE=1 SV=1 | 5 | 3 | 11 | 3 | 2.321928095 | 3.459431619 | -1.137503524 |
| Q9V3H2 | PSDE_DROME 26S proteasome non-ATPase regulatory subunit 14 OS=Drosophila melanogaster OX=7227 GN=Rpn11 PE=1 SV=1 | 5 | 4 | 11 | 6 | 2.321928095 | 3.459431619 | -1.137503524 |
| Q9VWA1 | CLC_DROME Clathrin light chain OS=Drosophila melanogaster OX=7227 GN=Clc PE=2 SV=1 | 5 | 3 | 11 | 6 | 2.321928095 | 3.459431619 | -1.137503524 |
| Q9VXX8 | RL371_DROME Probable 60S ribosomal protein L37-A OS=Drosophila melanogaster OX=7227 GN=RpL37a PE=1 SV=1 | 5 | 2 | 11 | 2 | 2.321928095 | 3.459431619 | -1.137503524 |
| A0A126GUV0 | A0A126GUV0_DROME Uncharacterized protein, isoform C OS=Drosophila melanogaster OX=7227 GN=CG10407 PE=1 SV=1 | 5 | 4 | 11 | 8 | 2.321928095 | 3.459431619 | -1.137503524 |
| Q9VSL6 | Q9VSL6_DROME Glutathione S transferase O1 OS=Drosophila melanogaster OX=7227 GN=GstO1 PE=1 SV=1 | 5 | 4 | 10 | 7 | 2.321928095 | 3.321928095 | -1 |
| Q9VCX3 | RM45_DROME Probable 39S ribosomal protein L45, mitochondrial OS=Drosophila melanogaster OX=7227 GN=mRpL45 PE=2 SV=1 | 5 | 5 | 9 | 5 | 2.321928095 | 3.169925001 | -0.847996907 |
| E2QD64 | E2QD64_DROME Uncharacterized protein, isoform B OS=Drosophila melanogaster OX=7227 GN=NO66 PE=1 SV=1 | 5 | 4 | 9 | 8 | 2.321928095 | 3.169925001 | -0.847996907 |
| Q9VHA1 | Q9VHA1_DROME Spermidine synthase, isoform A OS=Drosophila melanogaster OX=7227 GN=SpdS PE=1 SV=1 | 5 | 3 | 9 | 4 | 2.321928095 | 3.169925001 | -0.847996907 |
| Q9W5E1 | RBX1A_DROME RING-box protein 1A OS=Drosophila melanogaster OX=7227 GN=Roc1a PE=1 SV=1 | 5 | 3 | 8 | 3 | 2.321928095 | 3 | -0.678071905 |
| Q7JVH6 | Q7JVH6_DROME CG9436 OS=Drosophila melanogaster OX=7227 GN=CG9436 PE=1 SV=1 | 5 | 4 | 8 | 6 | 2.321928095 | 3 | -0.678071905 |
| Q9VD02 | Q9VD02_DROME GH14779p OS=Drosophila melanogaster OX=7227 GN=Dmel\CG7054 PE=1 SV=1 | 5 | 2 | 8 | 3 | 2.321928095 | 3 | -0.678071905 |
| Q9VEK8 | Q9VEK8_DROME GH07711p OS=Drosophila melanogaster OX=7227 GN=sds22 PE=1 SV=1 | 5 | 4 | 8 | 5 | 2.321928095 | 3 | -0.678071905 |
| Q9VG76 | MYCBP_DROME c-Myc-binding protein homolog OS=Drosophila melanogaster OX=7227 GN=CG17202 PE=2 SV=2 | 5 | 3 | 7 | 3 | 2.321928095 | 2.807354922 | -0.485426827 |
| Q9VW56 | PFD6_DROME Probable prefoldin subunit 6 OS=Drosophila melanogaster OX=7227 GN=Pfdn6 PE=2 SV=1 | 5 | 3 | 7 | 3 | 2.321928095 | 2.807354922 | -0.485426827 |

| UniProt ID | Protein description | # PSMs MVB spectra | # of unique peptides MVB | # of PSMs lysate | # of unique peptides lysate | log2(PSM MVB) | log2(PSM lysate) | FC (log2 of MVB PSM-log2 of lysate PSM) |
|---|---|---|---|---|---|---|---|---|
| A0A0B4LEK8 | A0A0B4LEK8_DROME Sec24AB ortholog, isoform C OS=Drosophila melanogaster OX=7227 GN=Sec24AB PE=1 SV=1 | 5 | 5 | 7 | 4 | 2.321928095 | 2.807354922 | -0.485426827 |
| A4V3J5 | A4V3J5_DROME Heterogeneous nuclear ribonucleoprotein at 98DE, isoform C OS=Drosophila melanogaster OX=7227 GN=Hrb98DE PE=1 SV=1 | 5 | 3 | 7 | 3 | 2.321928095 | 2.807354922 | -0.485426827 |
| F0JAN9 | F0JAN9_DROME RE19853p OS=Drosophila melanogaster OX=7227 GN=RhoGAP68F PE=1 SV=1 | 5 | 3 | 7 | 6 | 2.321928095 | 2.807354922 | -0.485426827 |
| Q86BQ4 | Q86BQ4_DROME Uncharacterized protein, isoform B OS=Drosophila melanogaster OX=7227 GN=CG2862 PE=1 SV=1 | 5 | 3 | 7 | 4 | 2.321928095 | 2.807354922 | -0.485426827 |
| Q9VH72 | Q9VH72_DROME TA01656p1 OS=Drosophila melanogaster OX=7227 GN=BVR PE=1 SV=1 | 5 | 4 | 7 | 4 | 2.321928095 | 2.807354922 | -0.485426827 |
| Q9W5N2 | RL38_DROME 60S ribosomal protein L38 OS=Drosophila melanogaster OX=7227 GN=RpL38 PE=1 SV=1 | 5 | 2 | 6 | 3 | 2.321928095 | 2.584962501 | -0.263034406 |
| Q9VE85 | Q9VE85_DROME Nucleoporin 43kD OS=Drosophila melanogaster OX=7227 GN=Nup43 PE=1 SV=1 | 5 | 2 | 6 | 4 | 2.321928095 | 2.584962501 | -0.263034406 |
| Q9VLS4 | Q9VLS4_DROME GEO10176p1 OS=Drosophila melanogaster OX=7227 GN=Dmel\CG8498 PE=1 SV=1 | 5 | 4 | 6 | 4 | 2.321928095 | 2.584962501 | -0.263034406 |
| P23625 | GNAQ_DROME G protein alpha q subunit OS=Drosophila melanogaster OX=7227 GN=Galphaq PE=2 SV=2 | 5 | 4 | 2 | 2 | 2.321928095 | 1 | 1.321928095 |
| P40793 | CDC42_DROME Cdc42 homolog OS=Drosophila melanogaster OX=7227 GN=Cdc42 PE=1 SV=1 | 5 | 3 | 2 | 2 | 2.321928095 | 1 | 1.321928095 |
| Q9VBV5 | SIL1_DROME Nucleotide exchange factor SIL1 OS=Drosophila melanogaster OX=7227 GN=CG10420 PE=2 SV=1 | 5 | 4 | 2 | 2 | 2.321928095 | 1 | 1.321928095 |
| Q9VE34 | EPG5_DROME Ectopic P granules protein 5 homolog OS=Drosophila melanogaster OX=7227 GN=CG14299 PE=1 SV=2 | 5 | 4 | 3 | 3 | 2.321928095 | 1.584962501 | 0.736965594 |
| Q9VF89 | RM09_DROME 39S ribosomal protein L9, mitochondrial OS=Drosophila melanogaster OX=7227 GN=mRpL9 PE=2 SV=1 | 5 | 3 | 1 | 1 | 2.321928095 | 0 | 2.321928095 |
| Q9VJD2 | LTOR3_DROME Ragulator complex protein LAMTOR3 homolog OS=Drosophila melanogaster OX=7227 GN=CG5110 PE=2 SV=1 | 5 | 3 | 1 | 1 | 2.321928095 | 0 | 2.321928095 |
| Q9VKB9 | RBGPR_DROME Rab3 GTPase-activating protein regulatory subunit OS=Drosophila melanogaster OX=7227 GN=Rab3-GAP PE=1 SV=2 | 5 | 4 | 3 | 3 | 2.321928095 | 1.584962501 | 0.736965594 |
| Q9VQ93 | GOLP3_DROME Golgi phosphoprotein 3 homolog sauron OS=Drosophila melanogaster OX=7227 GN=sau PE=1 SV=1 | 5 | 3 | 1 | 1 | 2.321928095 | 0 | 2.321928095 |
| Q9VTH0 | PLOD_DROME Procollagen-lysine,2-oxoglutarate 5-dioxygenase OS=Drosophila melanogaster OX=7227 GN=Plod PE=1 SV=1 | 5 | 4 | 1 | 1 | 2.321928095 | 0 | 2.321928095 |
| Q9W021 | RM23_DROME 39S ribosomal protein L23, mitochondrial OS=Drosophila melanogaster OX=7227 GN=mRpL23 PE=2 SV=1 | 5 | 3 | 1 | 1 | 2.321928095 | 0 | 2.321928095 |
| Q9XY35 | QCR9_DROME Cytochrome b-c1 complex subunit 9 OS=Drosophila melanogaster OX=7227 GN=ox PE=3 SV=1 | 5 | 2 | 1 | 1 | 2.321928095 | 0 | 2.321928095 |
| M9PDU4 | M9PDU4_DROME Acyl carrier protein OS=Drosophila melanogaster OX=7227 GN=ND-ACP PE=1 SV=1 | 5 | 2 | 5 | 3 | 2.321928095 | 2.321928095 | 0 |
| M9PEZ2 | M9PEZ2_DROME Fibroblast growth factor receptor 1 oncogene partner 2, isoform B OS=Drosophila melanogaster OX=7227 GN=Fgop2 PE=4 SV=1 | 5 | 4 | 1 | 1 | 2.321928095 | 0 | 2.321928095 |
| Q86BR8 | Q86BR8_DROME Mitochondrial transcription factor A, isoform B OS=Drosophila melanogaster OX=7227 GN=TFAM PE=1 SV=1 | 5 | 5 | 5 | 3 | 2.321928095 | 2.321928095 | 0 |
| M9PHN1 | M9PHN1_DROME Discontinuous actin hexagon, isoform B OS=Drosophila melanogaster OX=7227 GN=dah PE=4 SV=1 | 5 | 4 | 2 | 2 | 2.321928095 | 1 | 1.321928095 |
| O61722 | O61722_DROME PRL-1 phosphatase, isoform A OS=Drosophila melanogaster OX=7227 GN=PRL-1 PE=1 SV=2 | 5 | 4 | 4 | 2 | 2.321928095 | 2 | 0.321928095 |
| Q9V3D0 | Q9V3D0_DROME LD33485p OS=Drosophila melanogaster OX=7227 GN=mRpL4 PE=1 SV=1 | 5 | 4 | 5 | 3 | 2.321928095 | 2.321928095 | 0 |
| O76904 | O76904_DROME EG:8D8.1 protein OS=Drosophila melanogaster OX=7227 GN=fs(1)N PE=2 SV=1 | 5 | 5 | 4 | 4 | 2.321928095 | 2 | 0.321928095 |
| Q0E9B5 | Q0E9B5_DROME CG8841, isoform A OS=Drosophila melanogaster OX=7227 GN=CG8841 PE=1 SV=1 | 5 | 4 | 4 | 3 | 2.321928095 | 2 | 0.321928095 |
| Q6NP72 | Q6NP72_DROME GEO09659p1 OS=Drosophila melanogaster OX=7227 GN=CG13220 PE=1 SV=1 | 5 | 3 | 1 | 1 | 2.321928095 | 0 | 2.321928095 |
| Q7K010 | Q7K010_DROME Tetraspanin OS=Drosophila melanogaster OX=7227 GN=Tsp42Ef PE=2 SV=1 | 5 | 3 | 2 | 1 | 2.321928095 | 1 | 1.321928095 |
| Q9VEA1 | Q9VEA1_DROME Eukaryotic translation initiation factor 1A, isoform A OS=Drosophila melanogaster OX=7227 GN=eIF1A PE=1 SV=1 | 5 | 2 | 5 | 3 | 2.321928095 | 2.321928095 | 0 |
| Q9VBI3 | Q9VBI3_DROME RH72336p OS=Drosophila melanogaster OX=7227 GN=Vps2 PE=1 SV=1 | 5 | 4 | 3 | 3 | 2.321928095 | 1.584962501 | 0.736965594 |
| Q9VDZ5 | Q9VDZ5_DROME GH27195p OS=Drosophila melanogaster OX=7227 GN=CG42359-RA PE=1 SV=3 | 5 | 1 | 3 | 1 | 2.321928095 | 1.584962501 | 0.736965594 |
| Q9VF15 | Q9VF15_DROME GEO09476p1 OS=Drosophila melanogaster OX=7227 GN=glob1 PE=1 SV=1 | 5 | 4 | 4 | 3 | 2.321928095 | 2 | 0.321928095 |
| Q9VRQ9 | Q9VRQ9_DROME GH11174p OS=Drosophila melanogaster OX=7227 GN=QC PE=1 SV=2 | 5 | 4 | 5 | 3 | 2.321928095 | 2.321928095 | 0 |
| Q9VVT6 | Q9VVT6_DROME GEO08966p1 OS=Drosophila melanogaster OX=7227 GN=Grx1 PE=2 SV=1 | 5 | 3 | 3 | 2 | 2.321928095 | 1.584962501 | 0.736965594 |
| Q9VXQ0 | Q9VXQ0_DROME Mitochondrial ribosomal protein L3 OS=Drosophila melanogaster OX=7227 GN=mRpL3 PE=1 SV=2 | 5 | 4 | 2 | 2 | 2.321928095 | 1 | 1.321928095 |
| Q9W3C8 | Q9W3C8_DROME LD36568p OS=Drosophila melanogaster OX=7227 GN=CG11285 PE=1 SV=1 | 5 | 4 | 3 | 3 | 2.321928095 | 1.584962501 | 0.736965594 |
| Q9VEC4 | Q9VEC4_DROME Uncharacterized protein, isoform A OS=Drosophila melanogaster OX=7227 GN=CG7208 PE=4 SV=3 | 5 | 5 | 1 | 1 | 2.321928095 | 0 | 2.321928095 |
| Q9VG01 | Q9VG01_DROME RE12073p OS=Drosophila melanogaster OX=7227 GN=Dmel\CG46280 PE=1 SV=1 | 5 | 5 | 1 | 1 | 2.321928095 | 0 | 2.321928095 |
| Q9VQ96 | Q9VQ96_DROME RH13166p OS=Drosophila melanogaster OX=7227 GN=CG31686 PE=2 SV=1 | 5 | 3 | 1 | 1 | 2.321928095 | 0 | 2.321928095 |
| Q9W2G9 | Q9W2G9_DROME Acyl-coenzyme A oxidase OS=Drosophila melanogaster OX=7227 GN=Acox57D-p PE=1 SV=1 | 5 | 5 | 1 | 1 | 2.321928095 | 0 | 2.321928095 |
| P09208 | INSR_DROME Insulin-like receptor OS=Drosophila melanogaster OX=7227 GN=InR PE=1 SV=3 | 5 | 4 | 1 | 1 | 2.321928095 | 0 | 2.321928095 |
| P33270 | CP6A2_DROME Cytochrome P450 6a2 OS=Drosophila melanogaster OX=7227 GN=Cyp6a2 PE=2 SV=2 | 5 | 4 | 1 | 1 | 2.321928095 | 0 | 2.321928095 |
| P35220 | CTNA_DROME Catenin alpha OS=Drosophila melanogaster OX=7227 GN=alpha-Cat PE=1 SV=2 | 5 | 5 | 1 | 1 | 2.321928095 | 0 | 2.321928095 |
| Q27593 | CP6A8_DROME Cytochrome P450 6a8 OS=Drosophila melanogaster OX=7227 GN=Cyp6a8 PE=2 SV=2 | 5 | 4 | 1 | 1 | 2.321928095 | 0 | 2.321928095 |
| Q7JQ32 | GC76C_DROME Receptor-type guanylate cyclase Gyc76C OS=Drosophila melanogaster OX=7227 GN=Gyc76C PE=1 SV=1 | 5 | 5 | 1 | 1 | 2.321928095 | 0 | 2.321928095 |
| Q8IMX7 | MIRO_DROME Mitochondrial Rho GTPase OS=Drosophila melanogaster OX=7227 GN=Miro PE=1 SV=1 | 5 | 3 | 1 | 1 | 2.321928095 | 0 | 2.321928095 |
| Q8IN78 | TIM22_DROME Mitochondrial import inner membrane translocase subunit Tim22 OS=Drosophila melanogaster OX=7227 GN=Tim22 PE=2 SV=1 | 5 | 4 | 1 | 1 | 2.321928095 | 0 | 2.321928095 |
| Q8IRI6 | GTR1_DROME Glucose transporter type 1 OS=Drosophila melanogaster OX=7227 GN=Glut1 PE=2 SV=4 | 5 | 4 | 1 | 1 | 2.321928095 | 0 | 2.321928095 |
| Q8T8L8 | ALG10_DROME Putative Dol-P-Glc:Glc(2)Man(9)GlcNAc(2)-PP-Dol alpha-1,2-glucosyltransferase OS=Drosophila melanogaster OX=7227 GN=Alg10 PE=2 SV=1 | 5 | 5 | 1 | 1 | 2.321928095 | 0 | 2.321928095 |
| Q9GQQ0 | SPIN_DROME Protein spinster OS=Drosophila melanogaster OX=7227 GN=spin PE=1 SV=1 | 5 | 3 | 1 | 1 | 2.321928095 | 0 | 2.321928095 |
| Q9NI63 | PMYT1_DROME Membrane-associated tyrosine- and threonine-specific cdc2-inhibitory kinase OS=Drosophila melanogaster OX=7227 GN=Myt1 PE=1 SV=2 | 5 | 5 | 1 | 1 | 2.321928095 | 0 | 2.321928095 |
| Q9VIW3 | RAGP1_DROME Ran GTPase-activating protein OS=Drosophila melanogaster OX=7227 GN=RanGAP PE=1 SV=1 | 5 | 4 | 1 | 1 | 2.321928095 | 0 | 2.321928095 |
| Q9VRG7 | CNEP1_DROME CTD nuclear envelope phosphatase 1 homolog OS=Drosophila melanogaster OX=7227 GN=Dd PE=2 SV=1 | 5 | 4 | 1 | 1 | 2.321928095 | 0 | 2.321928095 |
| Q9VYV5 | FUT8_DROME Alpha-(1,6)-fucosyltransferase OS=Drosophila melanogaster OX=7227 GN=FucT6 PE=1 SV=1 | 5 | 4 | 1 | 1 | 2.321928095 | 0 | 2.321928095 |
| Q9VZJ9 | MUL1_DROME Mitochondrial E3 ubiquitin protein ligase 1 OS=Drosophila melanogaster OX=7227 GN=Mul1 PE=1 SV=1 | 5 | 4 | 1 | 1 | 2.321928095 | 0 | 2.321928095 |
| Q9W2M2 | TREA_DROME Trehalase OS=Drosophila melanogaster OX=7227 GN=Treh PE=1 SV=1 | 5 | 5 | 1 | 1 | 2.321928095 | 0 | 2.321928095 |
| A0A0B4KEU5 | A0A0B4KEU5_DROME Coronin OS=Drosophila melanogaster OX=7227 GN=coro PE=1 SV=1 | 5 | 3 | 1 | 1 | 2.321928095 | 0 | 2.321928095 |
| A0A0B4KFQ0 | A0A0B4KFQ0_DROME BIR repeat containing ubiquitin-conjugating enzyme, isoform E OS=Drosophila melanogaster OX=7227 GN=Bruce PE=1 SV=1 | 5 | 4 | 1 | 1 | 2.321928095 | 0 | 2.321928095 |
| A0A0S0WGR0 | A0A0S0WGR0_DROME Uncharacterized protein, isoform C OS=Drosophila melanogaster OX=7227 GN=CG7970 PE=1 SV=1 | 5 | 2 | 1 | 1 | 2.321928095 | 0 | 2.321928095 |
| A0A0S0WIH4 | A0A0S0WIH4_DROME Myosin heavy chain, isoform V OS=Drosophila melanogaster OX=7227 GN=Mhc PE=1 SV=1 | 5 | 5 | 1 | 1 | 2.321928095 | 0 | 2.321928095 |
| A1Z7Z4 | A1Z7Z4_DROME CG1648, isoform B OS=Drosophila melanogaster OX=7227 GN=CG1648 PE=1 SV=1 | 5 | 2 | 1 | 1 | 2.321928095 | 0 | 2.321928095 |
| A1Z8R8 | A1Z8R8_DROME Rhomboid-7, isoform A OS=Drosophila melanogaster OX=7227 GN=rho-7 PE=4 SV=1 | 5 | 3 | 1 | 1 | 2.321928095 | 0 | 2.321928095 |
| A1Z8T3 | A1Z8T3_DROME GEO10360p1 OS=Drosophila melanogaster OX=7227 GN=CG18343-RA PE=2 SV=1 | 5 | 1 | 1 | 1 | 2.321928095 | 0 | 2.321928095 |
| A8DY62 | A8DY62_DROME Uncharacterized protein, isoform B OS=Drosophila melanogaster OX=7227 GN=PIG-X PE=4 SV=1 | 5 | 4 | 1 | 1 | 2.321928095 | 0 | 2.321928095 |
| B7YZZ0 | B7YZZ0_DROME GH19557p OS=Drosophila melanogaster OX=7227 GN=CG42371-RA PE=2 SV=1 | 5 | 2 | 1 | 1 | 2.321928095 | 0 | 2.321928095 |
| M9MSM5 | M9MSM5_DROME Dishevelled associated activator of morphogenesis, isoform D OS=Drosophila melanogaster OX=7227 GN=DAAM PE=4 SV=1 | 5 | 3 | 1 | 1 | 2.321928095 | 0 | 2.321928095 |
| M9NCX9 | M9NCX9_DROME Galectin OS=Drosophila melanogaster OX=7227 GN=galectin PE=1 SV=1 | 5 | 4 | 1 | 1 | 2.321928095 | 0 | 2.321928095 |
| M9PFT8 | M9PFT8_DROME Uncharacterized protein, isoform B OS=Drosophila melanogaster OX=7227 GN=CG5235 PE=4 SV=1 | 5 | 5 | 1 | 1 | 2.321928095 | 0 | 2.321928095 |
| M9PIC3 | M9PIC3_DROME Uncharacterized protein, isoform B OS=Drosophila melanogaster OX=7227 GN=CG4825 PE=1 SV=1 | 5 | 4 | 1 | 1 | 2.321928095 | 0 | 2.321928095 |
| O96067 | O96067_DROME DJTB-like protein OS=Drosophila melanogaster OX=7227 GN=JTBR PE=2 SV=1 | 5 | 2 | 1 | 1 | 2.321928095 | 0 | 2.321928095 |
| Q7JS69 | Q7JS69_DROME FI04632p OS=Drosophila melanogaster OX=7227 GN=nrv3 PE=1 SV=1 | 5 | 5 | 1 | 1 | 2.321928095 | 0 | 2.321928095 |
| Q7JZE1 | Q7JZE1_DROME Peroxin 11 OS=Drosophila melanogaster OX=7227 GN=Pex11 PE=1 SV=1 | 5 | 5 | 1 | 1 | 2.321928095 | 0 | 2.321928095 |
| Q7K155 | Q7K155_DROME LD07878p OS=Drosophila melanogaster OX=7227 GN=Dmel\CG6928 PE=2 SV=1 | 5 | 4 | 1 | 1 | 2.321928095 | 0 | 2.321928095 |
| Q7KN74 | Q7KN74_DROME Lethal (2) k14710, isoform A OS=Drosophila melanogaster OX=7227 GN=l(2)k14710 PE=2 SV=1 | 5 | 5 | 1 | 1 | 2.321928095 | 0 | 2.321928095 |
| Q7KUR6 | Q7KUR6_DROME Uncharacterized protein, isoform B OS=Drosophila melanogaster OX=7227 GN=CG5535 PE=1 SV=1 | 5 | 4 | 1 | 1 | 2.321928095 | 0 | 2.321928095 |
| Q8MRW1 | Q8MRW1_DROME SD19278p OS=Drosophila melanogaster OX=7227 GN=Tim23 PE=1 SV=1 | 5 | 3 | 1 | 1 | 2.321928095 | 0 | 2.321928095 |
| Q8T0J5 | Q8T0J5_DROME GH26851p OS=Drosophila melanogaster OX=7227 GN=Dmel\CG7675 PE=1 SV=1 | 5 | 4 | 1 | 1 | 2.321928095 | 0 | 2.321928095 |
| Q9V3I6 | Q9V3I6_DROME RE32624p OS=Drosophila melanogaster OX=7227 GN=BG:DS02740.11 PE=2 SV=1 | 5 | 2 | 1 | 1 | 2.321928095 | 0 | 2.321928095 |
| Q9V420 | Q9V420_DROME FI02878p OS=Drosophila melanogaster OX=7227 GN=Su(P) PE=1 SV=1 | 5 | 4 | 1 | 1 | 2.321928095 | 0 | 2.321928095 |
| Q9VA78 | Q9VA78_DROME Sphingomyelin phosphodiesterase OS=Drosophila melanogaster OX=7227 GN=CG15533 PE=1 SV=1 | 5 | 3 | 1 | 1 | 2.321928095 | 0 | 2.321928095 |

| UniProt ID | Protein description | # PSMs MVB spectra | # of unique peptides MVB | # of PSMs lysate | # of unique peptides lysate | log2(PSM MVB) | log2(PSM lysate) | FC (log2 of MVB PSM-log2 of lysate PSM) |
|---|---|---|---|---|---|---|---|---|
| Q9VG00 | Q9VG00_DROME GEO09174p1 OS=Drosophila melanogaster OX=7227 GN=l(3)87Df PE=2 SV=1 | 5 | 3 | 1 | 1 | 2.321928095 | 0 | 2.321928095 |
| Q9VHB5 | Q9VHB5_DROME SD10846p OS=Drosophila melanogaster OX=7227 GN=Vps45 PE=1 SV=1 | 5 | 4 | 1 | 1 | 2.321928095 | 0 | 2.321928095 |
| Q9VI65 | Q9VI65_DROME LD01482p OS=Drosophila melanogaster OX=7227 GN=Dmel\CG10068 PE=2 SV=1 | 5 | 4 | 1 | 1 | 2.321928095 | 0 | 2.321928095 |
| Q9VJQ7 | Q9VJQ7_DROME LD27791p OS=Drosophila melanogaster OX=7227 GN=mol PE=2 SV=2 | 5 | 4 | 1 | 1 | 2.321928095 | 0 | 2.321928095 |
| Q9VLR4 | Q9VLR4_DROME SD10877p OS=Drosophila melanogaster OX=7227 GN=Dmel\CG8372 PE=2 SV=1 | 5 | 3 | 1 | 1 | 2.321928095 | 0 | 2.321928095 |
| Q9VLW9 | Q9VLW9_DROME Homocysteine-induced endoplasmic reticulum protein, isoform A OS=Drosophila melanogaster OX=7227 GN=Herp PE=1 SV=1 | 5 | 4 | 1 | 1 | 2.321928095 | 0 | 2.321928095 |
| Q9VMC3 | Q9VMC3_DROME LD35051p OS=Drosophila melanogaster OX=7227 GN=CG13765 PE=1 SV=2 | 5 | 4 | 1 | 1 | 2.321928095 | 0 | 2.321928095 |
| Q9VNB2 | Q9VNB2_DROME RE42193p OS=Drosophila melanogaster OX=7227 GN=CC1172 PE=2 SV=1 | 5 | 5 | 1 | 1 | 2.321928095 | 0 | 2.321928095 |
| Q9VP18 | Q9VP18_DROME V-type proton ATPase subunit OS=Drosophila melanogaster OX=7227 GN=VhaM9.7-b PE=1 SV=2 | 5 | 3 | 1 | 1 | 2.321928095 | 0 | 2.321928095 |
| Q9VR55 | Q9VR55_DROME LD29159p OS=Drosophila melanogaster OX=7227 GN=ADP PE=1 SV=1 | 5 | 3 | 1 | 1 | 2.321928095 | 0 | 2.321928095 |
| Q9VR57 | Q9VR57_DROME RE18169p OS=Drosophila melanogaster OX=7227 GN=Dmel\CG12576 PE=1 SV=1 | 5 | 3 | 1 | 1 | 2.321928095 | 0 | 2.321928095 |
| Q9VTU8 | Q9VTU8_DROME Carbonic anhydrase 2, isoform A OS=Drosophila melanogaster OX=7227 GN=CAH2 PE=2 SV=1 | 5 | 5 | 1 | 1 | 2.321928095 | 0 | 2.321928095 |
| Q9VU64 | Q9VU64_DROME LD27069p OS=Drosophila melanogaster OX=7227 GN=Dmel\CG10171 PE=2 SV=1 | 5 | 4 | 1 | 1 | 2.321928095 | 0 | 2.321928095 |
| Q9VVM1 | Q9VVM1_DROME Uncharacterized protein, isoform B OS=Drosophila melanogaster OX=7227 GN=Dmel\CG7408 PE=1 SV=2 | 5 | 4 | 1 | 1 | 2.321928095 | 0 | 2.321928095 |
| Q9VW35 | Q9VW35_DROME SD02261p OS=Drosophila melanogaster OX=7227 GN=Trpml PE=1 SV=1 | 5 | 4 | 1 | 1 | 2.321928095 | 0 | 2.321928095 |
| Q9VWT1 | Q9VWT1_DROME GH22474p OS=Drosophila melanogaster OX=7227 GN=HisRS PE=1 SV=1 | 5 | 5 | 1 | 1 | 2.321928095 | 0 | 2.321928095 |
| Q9VXG0 | Q9VXG0_DROME FI04422p OS=Drosophila melanogaster OX=7227 GN=CG9947-RA PE=2 SV=1 | 5 | 3 | 1 | 1 | 2.321928095 | 0 | 2.321928095 |
| Q9VY98 | Q9VY98_DROME LP12254p OS=Drosophila melanogaster OX=7227 GN=A PE=2 SV=1 | 5 | 3 | 1 | 1 | 2.321928095 | 0 | 2.321928095 |
| Q9VZW1 | Q9VZW1_DROME Phospholipid scramblase OS=Drosophila melanogaster OX=7227 GN=scramb2 PE=1 SV=1 | 5 | 3 | 1 | 1 | 2.321928095 | 0 | 2.321928095 |
| Q9W023 | Q9W023_DROME FI03626p OS=Drosophila melanogaster OX=7227 GN=Dmel\CG1317 PE=1 SV=2 | 5 | 3 | 1 | 1 | 2.321928095 | 0 | 2.321928095 |
| Q9W1F5 | Q9W1F5_DROME CG3803 OS=Drosophila melanogaster OX=7227 GN=CG3803 PE=2 SV=1 | 5 | 3 | 1 | 1 | 2.321928095 | 0 | 2.321928095 |
| Q9W1W8 | Q9W1W8_DROME CG3831 protein OS=Drosophila melanogaster OX=7227 GN=CG3831 PE=2 SV=2 | 5 | 3 | 1 | 1 | 2.321928095 | 0 | 2.321928095 |
| Q9W3K9 | Q9W3K9_DROME FI02989p OS=Drosophila melanogaster OX=7227 GN=Ldsdh1 PE=1 SV=1 | 5 | 4 | 1 | 1 | 2.321928095 | 0 | 2.321928095 |
| Q9W3Q0 | Q9W3Q0_DROME FI07418p OS=Drosophila melanogaster OX=7227 GN=Rab39 PE=2 SV=1 | 5 | 4 | 1 | 1 | 2.321928095 | 0 | 2.321928095 |
| Q9W4F5 | Q9W4F5_DROME Protein tyrosine phosphatase 4E, isoform A OS=Drosophila melanogaster OX=7227 GN=Ptp4E PE=2 SV=1 | 5 | 4 | 1 | 1 | 2.321928095 | 0 | 2.321928095 |
| Q9W4P6 | Q9W4P6_DROME FI03654p OS=Drosophila melanogaster OX=7227 GN=BcDNA:GH06717 PE=1 SV=2 | 5 | 3 | 1 | 1 | 2.321928095 | 0 | 2.321928095 |
| Q9Y1A7 | Q9Y1A7_DROME LD25378p OS=Drosophila melanogaster OX=7227 GN=mnd PE=2 SV=1 | 5 | 3 | 1 | 1 | 2.321928095 | 0 | 2.321928095 |
| X2JCM1 | X2JCM1_DROME Ankyrin 2, isoform AA OS=Drosophila melanogaster OX=7227 GN=Ank2 PE=1 SV=1 | 5 | 4 | 1 | 1 | 2.321928095 | 0 | 2.321928095 |
| Q7KN62 | TERA_DROME Transitional endoplasmic reticulum ATPase TER94 OS=Drosophila melanogaster OX=7227 GN=TER94 PE=1 SV=1 | 4 | 3 | 119 | 54 | 2 | 6.894817763 | -4.894817763 |
| Q8IMF5 | Q8IMF5_DROME Microtubule-associated protein 205, isoform B OS=Drosophila melanogaster OX=7227 GN=Map205 PE=1 SV=1 | 4 | 4 | 48 | 18 | 2 | 5.584962501 | -3.584962501 |
| Q9VLT5 | POE_DROME Protein purity of essence OS=Drosophila melanogaster OX=7227 GN=poe PE=1 SV=1 | 4 | 4 | 47 | 37 | 2 | 5.554588852 | -3.554588852 |
| O61613 | NMT_DROME Glycylpeptide N-tetradecanoyltransferase OS=Drosophila melanogaster OX=7227 GN=Nmt PE=2 SV=2 | 4 | 4 | 41 | 16 | 2 | 5.357552005 | -3.357552005 |
| P39018 | RS19A_DROME 40S ribosomal protein S19a OS=Drosophila melanogaster OX=7227 GN=RpS19a PE=1 SV=3 | 4 | 3 | 35 | 8 | 2 | 5.129283017 | -3.129283017 |
| A0A0B4LGZ3 | A0A0B4LGZ3_DROME Serine/threonine-protein phosphatase 2A 55 kDa regulatory subunit B OS=Drosophila melanogaster OX=7227 GN=tws PE=1 SV=1 | 4 | 4 | 34 | 16 | 2 | 5.087462841 | -3.087462841 |
| Q9VWI2 | Q9VWI2_DROME N(Alpha)-acetyltransferase 15/16, isoform A OS=Drosophila melanogaster OX=7227 GN=Naa15-16 PE=1 SV=1 | 4 | 4 | 33 | 22 | 2 | 5.044394119 | -3.044394119 |
| P35600 | RFC1_DROME Replication factor C subunit 1 OS=Drosophila melanogaster OX=7227 GN=Gnf1 PE=1 SV=2 | 4 | 3 | 30 | 21 | 2 | 4.906890596 | -2.906890596 |
| Q7JZW2 | Q7JZW2_DROME GEO07301p1 OS=Drosophila melanogaster OX=7227 GN=RpS15 PE=1 SV=1 | 4 | 3 | 26 | 7 | 2 | 4.700439718 | -2.700439718 |
| Q9V3K3 | RUVB2_DROME RuvB-like helicase 2 OS=Drosophila melanogaster OX=7227 GN=rept PE=1 SV=1 | 4 | 3 | 25 | 11 | 2 | 4.64385619 | -2.64385619 |
| Q9VUM0 | MSH6_DROME Probable DNA mismatch repair protein Msh6 OS=Drosophila melanogaster OX=7227 GN=Msh6 PE=1 SV=2 | 4 | 3 | 25 | 19 | 2 | 4.64385619 | -2.64385619 |
| A1Z6L9 | A1Z6L9_DROME FI21274p1 OS=Drosophila melanogaster OX=7227 GN=Trap1 PE=1 SV=1 | 4 | 4 | 24 | 17 | 2 | 4.584962501 | -2.584962501 |
| A1Z8U0 | A1Z8U0_DROME Pre-mRNA processing factor 8 OS=Drosophila melanogaster OX=7227 GN=Prp8 PE=1 SV=1 | 4 | 4 | 24 | 21 | 2 | 4.584962501 | -2.584962501 |
| Q9W3N9 | Q9W3N9_DROME LD24105p OS=Drosophila melanogaster OX=7227 GN=154229_at PE=1 SV=1 | 4 | 4 | 24 | 11 | 2 | 4.584962501 | -2.584962501 |
| Q7KLW9 | Q7KLW9_DROME BcDNA.LD02793 OS=Drosophila melanogaster OX=7227 GN=Prp19 PE=1 SV=1 | 4 | 4 | 23 | 12 | 2 | 4.523561956 | -2.523561956 |
| Q9VSL4 | Q9VSL4_DROME Glutathione S transferase O2, isoform B OS=Drosophila melanogaster OX=7227 GN=GstO2 PE=1 SV=2 | 4 | 3 | 23 | 8 | 2 | 4.523561956 | -2.523561956 |
| Q7K581 | Q7K581_DROME CG8963, isoform A OS=Drosophila melanogaster OX=7227 GN=CG8963 PE=1 SV=1 | 4 | 3 | 20 | 8 | 2 | 4.321928095 | -2.321928095 |
| Q9I7S8 | PUR6_DROME Multifunctional protein ADE2 OS=Drosophila melanogaster OX=7227 GN=ade5 PE=2 SV=2 | 4 | 3 | 19 | 11 | 2 | 4.247927513 | -2.247927513 |
| Q7K7G0 | Q7K7G0_DROME Coatomer subunit delta OS=Drosophila melanogaster OX=7227 GN=deltaCOP PE=1 SV=1 | 4 | 4 | 19 | 10 | 2 | 4.247927513 | -2.247927513 |
| Q9VNX8 | EIF2A_DROME Eukaryotic translation initiation factor 2A OS=Drosophila melanogaster OX=7227 GN=CG7414 PE=1 SV=1 | 4 | 4 | 18 | 14 | 2 | 4.169925001 | -2.169925001 |
| Q7K148 | Q7K148_DROME Proteasome subunit beta type OS=Drosophila melanogaster OX=7227 GN=Prosbeta5 PE=1 SV=1 | 4 | 2 | 18 | 7 | 2 | 4.169925001 | -2.169925001 |
| Q7KM15 | Q7KM15_DROME Transcription factor BTF3 OS=Drosophila melanogaster OX=7227 GN=bic PE=1 SV=1 | 4 | 3 | 18 | 5 | 2 | 4.169925001 | -2.169925001 |
| Q8IN02 | Q8IN02_DROME BcDNA.LD12153 OS=Drosophila melanogaster OX=7227 GN=Dph5 PE=1 SV=2 | 4 | 4 | 18 | 6 | 2 | 4.169925001 | -2.169925001 |
| A1ZAK3 | A1ZAK3_DROME Dek, isoform C OS=Drosophila melanogaster OX=7227 GN=Dek PE=1 SV=1 | 4 | 4 | 17 | 8 | 2 | 4.087462841 | -2.087462841 |
| Q9VDT1 | Q9VDT1_DROME Arc42 OS=Drosophila melanogaster OX=7227 GN=Arc42 PE=1 SV=1 | 4 | 3 | 17 | 12 | 2 | 4.087462841 | -2.087462841 |
| Q9VL68 | Q9VL68_DROME RH73277p OS=Drosophila melanogaster OX=7227 GN=Dmel\CG4598 PE=1 SV=2 | 4 | 3 | 17 | 9 | 2 | 4.087462841 | -2.087462841 |
| P48588 | RS25_DROME 40S ribosomal protein S25 OS=Drosophila melanogaster OX=7227 GN=RpS25 PE=1 SV=3 | 4 | 2 | 16 | 6 | 2 | 4 | -2 |
| A1Z7K8 | A1Z7K8_DROME CG8235 OS=Drosophila melanogaster OX=7227 GN=AIMP1 PE=1 SV=1 | 4 | 4 | 16 | 9 | 2 | 4 | -2 |
| M9NDI3 | M9NDI3_DROME Sorting nexin 6, isoform B OS=Drosophila melanogaster OX=7227 GN=Snx6 PE=1 SV=1 | 4 | 4 | 16 | 10 | 2 | 4 | -2 |
| Q0KI98 | Q0KI98_DROME FI06027p OS=Drosophila melanogaster OX=7227 GN=TrpRS PE=1 SV=1 | 4 | 3 | 16 | 11 | 2 | 4 | -2 |
| Q9VKW3 | Q9VKW3_DROME Replication factor C subunit 3 OS=Drosophila melanogaster OX=7227 GN=RfC3 PE=1 SV=2 | 4 | 4 | 16 | 13 | 2 | 4 | -2 |
| O46037 | VINC_DROME Vinculin OS=Drosophila melanogaster OX=7227 GN=Vinc PE=1 SV=1 | 4 | 4 | 15 | 12 | 2 | 3.906890596 | -1.906890596 |
| X2J516 | X2J516_DROME Trifunctional purine biosynthetic protein adenosine-3 OS=Drosophila melanogaster OX=7227 GN=Gart PE=1 SV=1 | 4 | 3 | 15 | 13 | 2 | 3.906890596 | -1.906890596 |
| Q9V9X4 | MTNA_DROME Methylthioribose-1-phosphate isomerase OS=Drosophila melanogaster OX=7227 GN=CG11334 PE=2 SV=1 | 4 | 4 | 14 | 11 | 2 | 3.807354922 | -1.807354922 |
| A1Z6X6 | A1Z6X6_DROME Glyoxalase 1 OS=Drosophila melanogaster OX=7227 GN=Glo1 PE=1 SV=1 | 4 | 2 | 14 | 6 | 2 | 3.807354922 | -1.807354922 |
| Q7K3M5 | Q7K3M5_DROME DHX15 ortholog, isoform A OS=Drosophila melanogaster OX=7227 GN=Dhx15 PE=1 SV=1 | 4 | 4 | 14 | 11 | 2 | 3.807354922 | -1.807354922 |
| Q9U9Q1 | Q9U9Q1_DROME Replication factor C 38kD subunit, isoform A OS=Drosophila melanogaster OX=7227 GN=RfC38 PE=1 SV=1 | 4 | 4 | 14 | 10 | 2 | 3.807354922 | -1.807354922 |
| Q9VAJ2 | Q9VAJ2_DROME Bub3, isoform A OS=Drosophila melanogaster OX=7227 GN=Bub3 PE=1 SV=1 | 4 | 4 | 13 | 10 | 2 | 3.700439718 | -1.700439718 |
| Q9VKW1 | Q9VKW1_DROME LD41958p OS=Drosophila melanogaster OX=7227 GN=Wdfy2 PE=1 SV=1 | 4 | 3 | 13 | 8 | 2 | 3.700439718 | -1.700439718 |
| Q9XYZ9 | Q9XYZ9_DROME Glutathione S transferase E12, isoform A OS=Drosophila melanogaster OX=7227 GN=GstE12 PE=1 SV=1 | 4 | 2 | 13 | 6 | 2 | 3.700439718 | -1.700439718 |
| P02516 | HSP23_DROME Heat shock protein 23 OS=Drosophila melanogaster OX=7227 GN=Hsp23 PE=2 SV=2 | 4 | 2 | 12 | 6 | 2 | 3.584962501 | -1.584962501 |
| Q9VXP4 | PA1B2_DROME Platelet-activating factor acetylhydrolase IB subunit beta homolog OS=Drosophila melanogaster OX=7227 GN=Paf-AHalpha PE=1 SV=1 | 4 | 3 | 12 | 6 | 2 | 3.584962501 | -1.584962501 |
| Q9VGQ1 | Q9VGQ1_DROME GM01350p OS=Drosophila melanogaster OX=7227 GN=Dmel\CG5214 PE=1 SV=1 | 4 | 4 | 12 | 6 | 2 | 3.584962501 | -1.584962501 |
| X2JG40 | X2JG40_DROME Gamma-tubulin complex component OS=Drosophila melanogaster OX=7227 GN=Grip84 PE=1 SV=1 | 4 | 3 | 12 | 11 | 2 | 3.584962501 | -1.584962501 |
| Q05783 | HMGD_DROME High mobility group protein D OS=Drosophila melanogaster OX=7227 GN=HmgD PE=1 SV=1 | 4 | 1 | 11 | 4 | 2 | 3.459431619 | -1.459431619 |
| Q9V4S8 | CSN7_DROME COP9 signalosome complex subunit 7 OS=Drosophila melanogaster OX=7227 GN=CSN7 PE=1 SV=2 | 4 | 3 | 11 | 7 | 2 | 3.459431619 | -1.459431619 |
| Q9VIM5 | ARPC2_DROME Actin-related protein 2/3 complex subunit 2 OS=Drosophila melanogaster OX=7227 GN=Arpc2 PE=2 SV=2 | 4 | 3 | 11 | 6 | 2 | 3.459431619 | -1.459431619 |
| Q9VYA0 | NADE_DROME Probable glutamine-dependent NAD(+) synthetase OS=Drosophila melanogaster OX=7227 GN=CG9940 PE=1 SV=1 | 4 | 3 | 11 | 8 | 2 | 3.459431619 | -1.459431619 |
| E1JH01 | E1JH01_DROME COP9 signalosome subunit 4, isoform B OS=Drosophila melanogaster OX=7227 GN=CSN4 PE=1 SV=1 | 4 | 4 | 11 | 10 | 2 | 3.459431619 | -1.459431619 |
| Q95U54 | Q95U54_DROME GH06271p OS=Drosophila melanogaster OX=7227 GN=Mms19 PE=1 SV=1 | 4 | 3 | 11 | 9 | 2 | 3.459431619 | -1.459431619 |
| Q9VB23 | Q9VB23_DROME Serine/threonine protein phosphatase 2A regulatory subunit OS=Drosophila melanogaster OX=7227 GN=wdb PE=1 SV=2 | 4 | 3 | 11 | 9 | 2 | 3.459431619 | -1.459431619 |
| Q9VCR2 | Q9VCR2_DROME Uncharacterized protein, isoform A OS=Drosophila melanogaster OX=7227 GN=Dmel\CG6726 PE=1 SV=2 | 4 | 4 | 11 | 6 | 2 | 3.459431619 | -1.459431619 |
| Q9VTW1 | Q9VTW1_DROME RE15265p OS=Drosophila melanogaster OX=7227 GN=Sms PE=1 SV=1 | 4 | 3 | 11 | 6 | 2 | 3.459431619 | -1.459431619 |

| UniProt ID | Protein description | # PSMs MVB spectra | # of unique peptides MVB | # of PSMs lysate | # of unique peptides lysate | log2(PSM MVB) | log2(PSM lysate) | FC (log2 of MVB PSM-log2 of lysate PSM) |
|---|---|---|---|---|---|---|---|---|
| P22769 | PSA71_DROME Proteasome subunit alpha type-7-1 OS=Drosophila melanogaster OX=7227 GN=Prosalpha4 PE=1 SV=2 | 4 | 3 | 10 | 4 | 2 | 3.321928095 | -1.321928095 |
| P45888 | ARP2_DROME Actin-related protein 2 OS=Drosophila melanogaster OX=7227 GN=Arp2 PE=2 SV=3 | 4 | 3 | 10 | 8 | 2 | 3.321928095 | -1.321928095 |
| P46863 | KL61_DROME Kinesin-like protein Klp61F OS=Drosophila melanogaster OX=7227 GN=Klp61F PE=1 SV=2 | 4 | 4 | 10 | 9 | 2 | 3.321928095 | -1.321928095 |
| Q7JWD6 | Q7JWD6_DROME Elongin C, isoform A OS=Drosophila melanogaster OX=7227 GN=EloC PE=1 SV=1 | 4 | 2 | 10 | 4 | 2 | 3.321928095 | -1.321928095 |
| Q8IMX4 | Q8IMX4_DROME FI04408p OS=Drosophila melanogaster OX=7227 GN=Rox8 PE=1 SV=1 | 4 | 2 | 10 | 5 | 2 | 3.321928095 | -1.321928095 |
| Q9VFW4 | Q9VFW4_DROME MIP09035p OS=Drosophila melanogaster OX=7227 GN=poly PE=1 SV=1 | 4 | 3 | 10 | 6 | 2 | 3.321928095 | -1.321928095 |
| Q9V8M5 | 3HIDH_DROME Probable 3-hydroxyisobutyrate dehydrogenase, mitochondrial OS=Drosophila melanogaster OX=7227 GN=CG15093 PE=2 SV=2 | 4 | 4 | 9 | 5 | 2 | 3.169925001 | -1.169925001 |
| O62530 | O62530_DROME Adaptor protein complex 2, mu subunit, isoform A OS=Drosophila melanogaster OX=7227 GN=AP-2mu PE=1 SV=1 | 4 | 3 | 9 | 8 | 2 | 3.169925001 | -1.169925001 |
| Q8MLS2 | Q8MLS2_DROME Ribose-5-phosphate isomerase, isoform B OS=Drosophila melanogaster OX=7227 GN=Rpi PE=1 SV=3 | 4 | 4 | 9 | 8 | 2 | 3.169925001 | -1.169925001 |
| Q9VC06 | Q9VC06_DROME LD37516p OS=Drosophila melanogaster OX=7227 GN=Dmel\CG11771 PE=1 SV=1 | 4 | 4 | 9 | 8 | 2 | 3.169925001 | -1.169925001 |
| P49630 | RL36_DROME 60S ribosomal protein L36 OS=Drosophila melanogaster OX=7227 GN=RpL36 PE=1 SV=1 | 4 | 3 | 8 | 4 | 2 | 3 | -1 |
| A1Z7P1 | A1Z7P1_DROME Phosphomannomutase 45A OS=Drosophila melanogaster OX=7227 GN=Pmm45A PE=1 SV=1 | 4 | 3 | 8 | 5 | 2 | 3 | -1 |
| Q9VEC2 | Q9VEC2_DROME Proteasome assembly chaperone 2 OS=Drosophila melanogaster OX=7227 GN=CG12321-RA PE=1 SV=1 | 4 | 4 | 8 | 5 | 2 | 3 | -1 |
| P40423 | SQH_DROME Myosin regulatory light chain sqh OS=Drosophila melanogaster OX=7227 GN=sqh PE=1 SV=1 | 4 | 4 | 7 | 4 | 2 | 2.807354922 | -0.807354922 |
| Q9V9S8 | HEMH_DROME Ferrochelatase, mitochondrial OS=Drosophila melanogaster OX=7227 GN=FeCh PE=2 SV=1 | 4 | 3 | 7 | 5 | 2 | 2.807354922 | -0.807354922 |
| Q9XZ58 | CSN5_DROME COP9 signalosome complex subunit 5 OS=Drosophila melanogaster OX=7227 GN=CSN5 PE=1 SV=1 | 4 | 3 | 7 | 5 | 2 | 2.807354922 | -0.807354922 |
| M9PI82 | M9PI82_DROME Mo25, isoform B OS=Drosophila melanogaster OX=7227 GN=Mo25 PE=4 SV=1 | 4 | 3 | 7 | 6 | 2 | 2.807354922 | -0.807354922 |
| Q9VAG9 | Q9VAG9_DROME LD34542p OS=Drosophila melanogaster OX=7227 GN=Dmel\CG7789 PE=1 SV=1 | 4 | 4 | 7 | 6 | 2 | 2.807354922 | -0.807354922 |
| Q8SYG2 | CSN3_DROME COP9 signalosome complex subunit 3 OS=Drosophila melanogaster OX=7227 GN=CSN3 PE=2 SV=2 | 4 | 3 | 6 | 3 | 2 | 2.584962501 | -0.584962501 |
| Q9VTF9 | UFD1_DROME Ubiquitin fusion degradation protein 1 homolog OS=Drosophila melanogaster OX=7227 GN=Ufd1-like PE=2 SV=1 | 4 | 2 | 6 | 5 | 2 | 2.584962501 | -0.584962501 |
| M9NDM4 | M9NDM4_DROME Will die slowly, isoform B OS=Drosophila melanogaster OX=7227 GN=wds PE=4 SV=1 | 4 | 4 | 6 | 5 | 2 | 2.584962501 | -0.584962501 |
| Q9VL71 | Q9VL71_DROME LD29830p OS=Drosophila melanogaster OX=7227 GN=Srp54 PE=1 SV=1 | 4 | 3 | 6 | 6 | 2 | 2.584962501 | -0.584962501 |
| Q9VWG1 | Q9VWG1_DROME LD37169p OS=Drosophila melanogaster OX=7227 GN=Dmel21.3 PE=1 SV=1 | 4 | 3 | 6 | 4 | 2 | 2.584962501 | -0.584962501 |
| Q9W0S6 | Q9W0S6_DROME IP18235p OS=Drosophila melanogaster OX=7227 GN=mRpL17 PE=1 SV=2 | 4 | 3 | 6 | 3 | 2 | 2.584962501 | -0.584962501 |
| Q9NK57 | NIF3L_DROME NIF3-like protein 1 OS=Drosophila melanogaster OX=7227 GN=anon-35F/36A PE=1 SV=3 | 4 | 4 | 5 | 5 | 2 | 2.321928095 | -0.321928095 |
| Q9VMC6 | Q9VMC6_DROME GH06693p OS=Drosophila melanogaster OX=7227 GN=152078_at PE=1 SV=1 | 4 | 4 | 5 | 4 | 2 | 2.321928095 | -0.321928095 |
| A0A0B4K692 | NEP2_DROME Neprilysin-2 OS=Drosophila melanogaster OX=7227 GN=Nep2 PE=1 SV=1 | 4 | 2 | 4 | 2 | 2 | 2 | 0 |
| P00399 | COX1_DROME Cytochrome c oxidase subunit 1 OS=Drosophila melanogaster OX=7227 GN=mt:CoI PE=2 SV=2 | 4 | 3 | 2 | 2 | 2 | 1 | 1 |
| P14199 | REF2P_DROME Protein ref(2)P OS=Drosophila melanogaster OX=7227 GN=ref(2)P PE=1 SV=2 | 4 | 3 | 2 | 1 | 2 | 1 | 1 |
| P18935 | CYB_DROME Cytochrome b OS=Drosophila melanogaster OX=7227 GN=mt:Cyt-b PE=3 SV=2 | 4 | 3 | 1 | 1 | 2 | 0 | 2 |
| P40688 | SWA_DROME Protein swallow OS=Drosophila melanogaster OX=7227 GN=swa PE=1 SV=1 | 4 | 3 | 3 | 3 | 2 | 1.584962501 | 0.415037499 |
| P84345 | ATP8_DROME ATP synthase protein 8 OS=Drosophila melanogaster OX=7227 GN=mt:ATPase8 PE=3 SV=1 | 4 | 2 | 1 | 1 | 2 | 0 | 2 |
| Q709R6 | BOCKS_DROME LEM domain-containing protein Bocksbeutel OS=Drosophila melanogaster OX=7227 GN=bocks PE=1 SV=1 | 4 | 4 | 3 | 2 | 2 | 1.584962501 | 0.415037499 |
| Q9U9P7 | PITC1_DROME Cytoplasmic phosphatidylinositol transfer protein 1 OS=Drosophila melanogaster OX=7227 GN=rdgBbeta PE=2 SV=1 | 4 | 4 | 1 | 1 | 2 | 0 | 2 |
| Q9UAS6 | CMTR2_DROME Cap-specific mRNA (nucleoside-2'-O-)-methyltransferase 2 OS=Drosophila melanogaster OX=7227 GN=cmtr2 PE=2 SV=2 | 4 | 4 | 3 | 2 | 2 | 1.584962501 | 0.415037499 |
| Q9VQ26 | RB3GP_DROME Rab3 GTPase-activating protein catalytic subunit OS=Drosophila melanogaster OX=7227 GN=CG31935 PE=1 SV=2 | 4 | 3 | 4 | 3 | 2 | 2 | 0 |
| Q9VVI9 | CHMP5_DROME Charged multivesicular body protein 5 OS=Drosophila melanogaster OX=7227 GN=Vps60 PE=1 SV=2 | 4 | 3 | 3 | 3 | 2 | 1.584962501 | 0.415037499 |
| Q9VZ49 | ENDOU_DROME Poly(U)-specific endoribonuclease homolog OS=Drosophila melanogaster OX=7227 GN=CG2145 PE=2 SV=1 | 4 | 4 | 1 | 1 | 2 | 0 | 2 |
| A1Z9I0 | A1Z9I0_DROME CG6357 OS=Drosophila melanogaster OX=7227 GN=CG6357 PE=1 SV=1 | 4 | 4 | 1 | 1 | 2 | 0 | 2 |
| A8JUT4 | A8JUT4_DROME Nuclear transport factor-2, isoform B OS=Drosophila melanogaster OX=7227 GN=Ntf-2 PE=1 SV=1 | 4 | 2 | 3 | 2 | 2 | 1.584962501 | 0.415037499 |
| M9PC89 | M9PC89_DROME Small ribonucleoprotein particle U1 subunit 70K, isoform C OS=Drosophila melanogaster OX=7227 GN=snRNP-U1-70K PE=4 SV=1 | 4 | 3 | 3 | 3 | 2 | 1.584962501 | 0.415037499 |
| M9PCE8 | M9PCE8_DROME Angiotensin-converting enzyme OS=Drosophila melanogaster OX=7227 GN=Acer PE=3 SV=1 | 4 | 4 | 1 | 1 | 2 | 0 | 2 |
| M9PEL1 | M9PEL1_DROME Ras opposite, isoform B OS=Drosophila melanogaster OX=7227 GN=Rop PE=3 SV=1 | 4 | 3 | 2 | 2 | 2 | 1 | 1 |
| M9PF42 | M9PF42_DROME Connector of kinase to AP-1, isoform G OS=Drosophila melanogaster OX=7227 GN=Cka PE=1 SV=1 | 4 | 4 | 3 | 3 | 2 | 1.584962501 | 0.415037499 |
| M9PG83 | M9PG83_DROME Cysteine string protein, isoform D OS=Drosophila melanogaster OX=7227 GN=Csp PE=1 SV=1 | 4 | 3 | 2 | 2 | 2 | 1 | 1 |
| Q7K084 | Q7K084_DROME Odorant-binding protein 44a, isoform A OS=Drosophila melanogaster OX=7227 GN=Obp44a PE=1 SV=1 | 4 | 3 | 2 | 1 | 2 | 1 | 1 |
| Q95TP0 | Q95TP0_DROME LD34582p OS=Drosophila melanogaster OX=7227 GN=TBCC PE=1 SV=1 | 4 | 3 | 3 | 3 | 2 | 1.584962501 | 0.415037499 |
| Q9V3Y7 | Q9V3Y7_DROME Uncharacterized protein, isoform A OS=Drosophila melanogaster OX=7227 GN=BG:DS01759.2 PE=1 SV=1 | 4 | 3 | 2 | 2 | 2 | 1 | 1 |
| Q9VJ25 | Q9VJ25_DROME GEO07313p1 OS=Drosophila melanogaster OX=7227 GN=BcDNA:RE09053 PE=2 SV=1 | 4 | 1 | 2 | 1 | 2 | 1 | 1 |
| Q9VJ44 | Q9VJ44_DROME SD06839p OS=Drosophila melanogaster OX=7227 GN=Dmel\CG17597 PE=1 SV=2 | 4 | 3 | 2 | 2 | 2 | 1 | 1 |
| Q9VPF6 | Q9VPF6_DROME FI06804p OS=Drosophila melanogaster OX=7227 GN=mRpL15 PE=1 SV=1 | 4 | 3 | 4 | 4 | 2 | 2 | 0 |
| Q9VRY6 | Q9VRY6_DROME LD14015p OS=Drosophila melanogaster OX=7227 GN=Dmel\CG10103 PE=1 SV=1 | 4 | 2 | 3 | 3 | 2 | 1.584962501 | 0.415037499 |
| Q9VU36 | Q9VU36_DROME LP04985p OS=Drosophila melanogaster OX=7227 GN=mRpL20 PE=1 SV=1 | 4 | 3 | 3 | 2 | 2 | 1.584962501 | 0.415037499 |
| Q9W1I8 | Q9W1I8_DROME LD03592p OS=Drosophila melanogaster OX=7227 GN=Snap29 PE=1 SV=1 | 4 | 3 | 3 | 2 | 2 | 1.584962501 | 0.415037499 |
| Q9W4I3 | Q9W4I3_DROME FI01736p OS=Drosophila melanogaster OX=7227 GN=mRpL30 PE=1 SV=1 | 4 | 2 | 4 | 2 | 2 | 2 | 0 |
| Q9VEH2 | Q9VEH2_DROME Uncharacterized protein OS=Drosophila melanogaster OX=7227 GN=CG7523 PE=1 SV=1 | 4 | 2 | 1 | 1 | 2 | 0 | 2 |
| Q9VGE7 | Q9VGE7_DROME Beta-galactosidase OS=Drosophila melanogaster OX=7227 GN=Ect3 PE=1 SV=1 | 4 | 4 | 1 | 1 | 2 | 0 | 2 |
| Q9VTB3 | Q9VTB3_DROME GH06691p OS=Drosophila melanogaster OX=7227 GN=Dmel\CG11811 PE=1 SV=1 | 4 | 3 | 1 | 1 | 2 | 0 | 2 |
| Q9VUM2 | Q9VUM2_DROME RH56103p OS=Drosophila melanogaster OX=7227 GN=Dmel\CG6878 PE=1 SV=1 | 4 | 2 | 1 | 1 | 2 | 0 | 2 |
| O61307 | TENM_DROME Teneurin-m OS=Drosophila melanogaster OX=7227 GN=Ten-m PE=1 SV=2 | 4 | 4 | 1 | 1 | 2 | 0 | 2 |
| P56079 | CDSA_DROME Phosphatidate cytidylyltransferase, photoreceptor-specific OS=Drosophila melanogaster OX=7227 GN=CdsA PE=1 SV=2 | 4 | 2 | 1 | 1 | 2 | 0 | 2 |
| Q95SX8 | NAA60_DROME N-alpha-acetyltransferase 60 OS=Drosophila melanogaster OX=7227 GN=Naa60 PE=1 SV=1 | 4 | 3 | 1 | 1 | 2 | 0 | 2 |
| Q9U4F3 | SURF1_DROME SURF1-like protein OS=Drosophila melanogaster OX=7227 GN=Surf1 PE=2 SV=1 | 4 | 2 | 1 | 1 | 2 | 0 | 2 |
| Q9V3X4 | BSCL2_DROME Seipin OS=Drosophila melanogaster OX=7227 GN=Seipin PE=2 SV=1 | 4 | 3 | 1 | 1 | 2 | 0 | 2 |
| Q9VQP9 | OSTC_DROME Putative oligosaccharyltransferase complex subunit CG9662 OS=Drosophila melanogaster OX=7227 GN=CG9662 PE=2 SV=3 | 4 | 2 | 1 | 1 | 2 | 0 | 2 |
| Q9VRL2 | GOSR2_DROME Probable Golgi SNAP receptor complex member 2 OS=Drosophila melanogaster OX=7227 GN=Membrin PE=2 SV=1 | 4 | 4 | 1 | 1 | 2 | 0 | 2 |
| Q9VVA8 | TM258_DROME Transmembrane protein 258 homolog OS=Drosophila melanogaster OX=7227 GN=CG9669 PE=3 SV=1 | 4 | 2 | 1 | 1 | 2 | 0 | 2 |
| Q9VW71 | FAT2_DROME Fat-like cadherin-related tumor suppressor homolog OS=Drosophila melanogaster OX=7227 GN=kug PE=2 SV=3 | 4 | 4 | 1 | 1 | 2 | 0 | 2 |
| Q9VXG4 | ANX11_DROME Annexin B11 OS=Drosophila melanogaster OX=7227 GN=AnxB11 PE=2 SV=2 | 4 | 4 | 1 | 1 | 2 | 0 | 2 |
| A0A0B4K6T3 | A0A0B4K6T3_DROME Calcium-independent receptor for alpha-latrotoxin, isoform H OS=Drosophila melanogaster OX=7227 GN=Cirl PE=3 SV=1 | 4 | 3 | 1 | 1 | 2 | 0 | 2 |
| A0A0B4K7I3 | A0A0B4K7I3_DROME Down syndrome cell adhesion molecule 1, isoform BN OS=Drosophila melanogaster OX=7227 GN=Dscam1 PE=1 SV=1 | 4 | 4 | 1 | 1 | 2 | 0 | 2 |
| A0A0B4KEH8 | A0A0B4KEH8_DROME Uncharacterized protein, isoform D OS=Drosophila melanogaster OX=7227 GN=CG17883 PE=4 SV=1 | 4 | 4 | 1 | 1 | 2 | 0 | 2 |
| A0A0B4KH03 | A0A0B4KH03_DROME Tetraspanin OS=Drosophila melanogaster OX=7227 GN=Tsp96F PE=1 SV=1 | 4 | 2 | 1 | 1 | 2 | 0 | 2 |
| A0A0B4KH68 | A0A0B4KH68_DROME Lysosomal enzyme receptor protein, isoform E OS=Drosophila melanogaster OX=7227 GN=Lerp PE=1 SV=1 | 4 | 4 | 1 | 1 | 2 | 0 | 2 |
| A0A0B4LEX9 | A0A0B4LEX9_DROME Uncharacterized protein, isoform E OS=Drosophila melanogaster OX=7227 GN=CG42326 PE=4 SV=1 | 4 | 3 | 1 | 1 | 2 | 0 | 2 |
| A0A0B4LFK0 | A0A0B4LFK0_DROME Picot, isoform C OS=Drosophila melanogaster OX=7227 GN=Picot PE=4 SV=1 | 4 | 4 | 1 | 1 | 2 | 0 | 2 |
| A0A0B4LFU5 | A0A0B4LFU5_DROME Stambha A, isoform D OS=Drosophila melanogaster OX=7227 GN=stmA PE=4 SV=1 | 4 | 4 | 1 | 1 | 2 | 0 | 2 |
| A1Z6J2 | A1Z6J2_DROME SREBP cleavage activating protein OS=Drosophila melanogaster OX=7227 GN=SCAP PE=4 SV=1 | 4 | 3 | 1 | 1 | 2 | 0 | 2 |
| A1Z763 | A1Z763_DROME MIP13269p OS=Drosophila melanogaster OX=7227 GN=CG30377-RA PE=1 SV=1 | 4 | 3 | 1 | 1 | 2 | 0 | 2 |
| A1Z7G9 | A1Z7G9_DROME Hexosyltransferase OS=Drosophila melanogaster OX=7227 GN=beta3GalTII PE=2 SV=1 | 4 | 3 | 1 | 1 | 2 | 0 | 2 |
| A1Z920 | A1Z920_DROME FI24025p1 OS=Drosophila melanogaster OX=7227 GN=fra PE=1 SV=1 | 4 | 4 | 1 | 1 | 2 | 0 | 2 |

| UniProt ID | Protein description | # PSMs MVB spectra | # of unique peptides MVB | # of PSMs lysate | # of unique peptides lysate | log2(PSM MVB) | log2(PSM lysate) | FC (log2 of MVB PSM-log2 of lysate PSM) |
|---|---|---|---|---|---|---|---|---|
| A4V392 | A4V392_DROME Adiponectin receptor, isoform C OS=Drosophila melanogaster OX=7227 GN=AdipoR PE=1 SV=1 | 4 | 3 | 1 | 1 | 2 | 0 | 2 |
| A8DY83 | A8DY83_DROME Uncharacterized protein, isoform B OS=Drosophila melanogaster OX=7227 GN=CG12929 PE=4 SV=1 | 4 | 1 | 1 | 1 | 2 | 0 | 2 |
| A8DYC5 | A8DYC5_DROME GEO13345p1 OS=Drosophila melanogaster OX=7227 GN=ND-MWFE PE=2 SV=1 | 4 | 3 | 1 | 1 | 2 | 0 | 2 |
| A8Y535 | A8Y535_DROME Cytochrome b-c1 complex subunit 6 OS=Drosophila melanogaster OX=7227 GN=UQCR-11 PE=1 SV=2 | 4 | 1 | 1 | 1 | 2 | 0 | 2 |
| B7Z090 | B7Z090_DROME Presenilin OS=Drosophila melanogaster OX=7227 GN=Psn PE=1 SV=1 | 4 | 1 | 1 | 1 | 2 | 0 | 2 |
| B7Z0M9 | B7Z0M9_DROME Oxysterol-binding protein OS=Drosophila melanogaster OX=7227 GN=CG42668 PE=1 SV=1 | 4 | 4 | 1 | 1 | 2 | 0 | 2 |
| B7Z0V2 | B7Z0V2_DROME Uncharacterized protein, isoform I OS=Drosophila melanogaster OX=7227 GN=CG2698 PE=4 SV=2 | 4 | 4 | 1 | 1 | 2 | 0 | 2 |
| E1JJ71 | E1JJ71_DROME Dementin, isoform H OS=Drosophila melanogaster OX=7227 GN=Dmtn PE=4 SV=1 | 4 | 4 | 1 | 1 | 2 | 0 | 2 |
| F3YD52 | F3YD52_DROME FI14544p OS=Drosophila melanogaster OX=7227 GN=CG12910-RA PE=2 SV=1 | 4 | 4 | 1 | 1 | 2 | 0 | 2 |
| L0MLP2 | L0MLP2_DROME Calcium/calmodulin-dependent protein kinase II, isoform M OS=Drosophila melanogaster OX=7227 GN=CaMKII PE=1 SV=1 | 4 | 4 | 1 | 1 | 2 | 0 | 2 |
| L7EEU0 | L7EEU0_DROME Dead box protein 80, isoform E OS=Drosophila melanogaster OX=7227 GN=Dbp80 PE=1 SV=1 | 4 | 4 | 1 | 1 | 2 | 0 | 2 |
| M9PEK6 | M9PEK6_DROME Mushroom body defect, isoform L OS=Drosophila melanogaster OX=7227 GN=mud PE=1 SV=1 | 4 | 4 | 1 | 1 | 2 | 0 | 2 |
| M9PF10 | M9PF10_DROME Uncharacterized protein, isoform C OS=Drosophila melanogaster OX=7227 GN=CG5989 PE=4 SV=1 | 4 | 4 | 1 | 1 | 2 | 0 | 2 |
| M9PH36 | M9PH36_DROME Uncharacterized protein, isoform E OS=Drosophila melanogaster OX=7227 GN=CG34449 PE=1 SV=1 | 4 | 2 | 1 | 1 | 2 | 0 | 2 |
| M9PHD6 | M9PHD6_DROME Uncharacterized protein, isoform B OS=Drosophila melanogaster OX=7227 GN=CG4880 PE=4 SV=1 | 4 | 3 | 1 | 1 | 2 | 0 | 2 |
| M9PHI3 | M9PHI3_DROME Protein-tyrosine sulfotransferase OS=Drosophila melanogaster OX=7227 GN=Tango13 PE=3 SV=1 | 4 | 4 | 1 | 1 | 2 | 0 | 2 |
| M9PJN9 | M9PJN9_DROME Shibire, isoform O OS=Drosophila melanogaster OX=7227 GN=shi PE=1 SV=1 | 4 | 4 | 1 | 1 | 2 | 0 | 2 |
| Q2PDM2 | Q2PDM2_DROME Rho GTPase activating protein at 102A, isoform C OS=Drosophila melanogaster OX=7227 GN=RhoGAP102A PE=4 SV=2 | 4 | 1 | 1 | 1 | 2 | 0 | 2 |
| Q4QPU3 | Q4QPU3_DROME GEO12027p1 OS=Drosophila melanogaster OX=7227 GN=CG15908 PE=2 SV=1 | 4 | 4 | 1 | 1 | 2 | 0 | 2 |
| Q6NN28 | Q6NN28_DROME RE21974p OS=Drosophila melanogaster OX=7227 GN=Tm1 PE=1 SV=1 | 4 | 4 | 1 | 1 | 2 | 0 | 2 |
| Q7JR96 | Q7JR96_DROME SD19533p OS=Drosophila melanogaster OX=7227 GN=Snx16 PE=1 SV=1 | 4 | 4 | 1 | 1 | 2 | 0 | 2 |
| Q7JWS8 | Q7JWS8_DROME AT17867p OS=Drosophila melanogaster OX=7227 GN=Mtl PE=1 SV=1 | 4 | 4 | 1 | 1 | 2 | 0 | 2 |
| Q7K0L5 | Q7K0L5_DROME LP01562p OS=Drosophila melanogaster OX=7227 GN=veil PE=1 SV=1 | 4 | 4 | 1 | 1 | 2 | 0 | 2 |
| Q7K3T4 | Q7K3T4_DROME Palmitoyltransferase OS=Drosophila melanogaster OX=7227 GN=CG8314 PE=2 SV=1 | 4 | 3 | 1 | 1 | 2 | 0 | 2 |
| Q7K5K0 | Q7K5K0_DROME GASZ ortholog OS=Drosophila melanogaster OX=7227 GN=Gasz PE=2 SV=1 | 4 | 3 | 1 | 1 | 2 | 0 | 2 |
| Q7KTG4 | Q7KTG4_DROME Organic anion transporting polypeptide 30B, isoform D OS=Drosophila melanogaster OX=7227 GN=Oatp30B PE=4 SV=1 | 4 | 4 | 1 | 1 | 2 | 0 | 2 |
| Q7KTG6 | Q7KTG6_DROME Phospholipid-transporting ATPase OS=Drosophila melanogaster OX=7227 GN=CG33298 PE=3 SV=1 | 4 | 4 | 1 | 1 | 2 | 0 | 2 |
| Q7KTS3 | Q7KTS3_DROME Uncharacterized protein, isoform B OS=Drosophila melanogaster OX=7227 GN=Dmel\CG2519 PE=4 SV=1 | 4 | 2 | 1 | 1 | 2 | 0 | 2 |
| Q7KTY3 | Q7KTY3_DROME CG5976 OS=Drosophila melanogaster OX=7227 GN=isoQC PE=2 SV=1 | 4 | 3 | 1 | 1 | 2 | 0 | 2 |
| Q7KV99 | Q7KV99_DROME Uncharacterized protein, isoform D OS=Drosophila melanogaster OX=7227 GN=CG1275 PE=1 SV=1 | 4 | 2 | 1 | 1 | 2 | 0 | 2 |
| Q86P97 | Q86P97_DROME GM06145p OS=Drosophila melanogaster OX=7227 GN=SppL PE=2 SV=1 | 4 | 3 | 1 | 1 | 2 | 0 | 2 |
| Q8IMG1 | Q8IMG1_DROME Uncharacterized protein, isoform C OS=Drosophila melanogaster OX=7227 GN=CG1910 PE=1 SV=1 | 4 | 4 | 1 | 1 | 2 | 0 | 2 |
| Q8IR26 | Q8IR26_DROME RE35552p OS=Drosophila melanogaster OX=7227 GN=CG32581-RA PE=2 SV=2 | 4 | 3 | 1 | 1 | 2 | 0 | 2 |
| Q8IR72 | Q8IR72_DROME FI19011p1 OS=Drosophila melanogaster OX=7227 GN=CG32638-RB PE=1 SV=2 | 4 | 2 | 1 | 1 | 2 | 0 | 2 |
| Q95R73 | Q95R73_DROME SD03412p OS=Drosophila melanogaster OX=7227 GN=CG18278 PE=1 SV=1 | 4 | 2 | 1 | 1 | 2 | 0 | 2 |
| Q960F6 | Q960F6_DROME SD03655p OS=Drosophila melanogaster OX=7227 GN=Dmel\CG12004 PE=2 SV=1 | 4 | 3 | 1 | 1 | 2 | 0 | 2 |
| Q9I7K4 | Q9I7K4_DROME LD18367p OS=Drosophila melanogaster OX=7227 GN=Dmel\CG18542 PE=1 SV=1 | 4 | 2 | 1 | 1 | 2 | 0 | 2 |
| Q9V395 | Q9V395_DROME BcDNA.LD28419 OS=Drosophila melanogaster OX=7227 GN=rtet PE=2 SV=1 | 4 | 2 | 1 | 1 | 2 | 0 | 2 |
| Q9V3S7 | Q9V3S7_DROME BcDNA.GH12326 OS=Drosophila melanogaster OX=7227 GN=BcDNA.GH12326 PE=1 SV=1 | 4 | 4 | 1 | 1 | 2 | 0 | 2 |
| Q9VBW0 | Q9VBW0_DROME Uncharacterized protein OS=Drosophila melanogaster OX=7227 GN=CG13663 PE=4 SV=1 | 4 | 2 | 1 | 1 | 2 | 0 | 2 |
| Q9VDC8 | Q9VDC8_DROME FI06469p OS=Drosophila melanogaster OX=7227 GN=Dmel\CG3337 PE=2 SV=1 | 4 | 2 | 1 | 1 | 2 | 0 | 2 |
| Q9VDT4 | Q9VDT4_DROME Uncharacterized protein, isoform A OS=Drosophila melanogaster OX=7227 GN=BcDNA:AT09463 PE=4 SV=1 | 4 | 2 | 1 | 1 | 2 | 0 | 2 |
| Q9VEU9 | Q9VEU9_DROME FI21838p1 OS=Drosophila melanogaster OX=7227 GN=CG14883-RA PE=2 SV=2 | 4 | 3 | 1 | 1 | 2 | 0 | 2 |
| Q9VG64 | Q9VG64_DROME RE35942p OS=Drosophila melanogaster OX=7227 GN=Dmel\CG18549 PE=1 SV=2 | 4 | 3 | 1 | 1 | 2 | 0 | 2 |
| Q9VGN8 | Q9VGN8_DROME CG5276 protein OS=Drosophila melanogaster OX=7227 GN=Dmel\CG5276 PE=2 SV=1 | 4 | 4 | 1 | 1 | 2 | 0 | 2 |
| Q9VGQ2 | Q9VGQ2_DROME RE71331p OS=Drosophila melanogaster OX=7227 GN=BcDNA:RH54117 PE=1 SV=1 | 4 | 2 | 1 | 1 | 2 | 0 | 2 |
| Q9VIC3 | Q9VIC3_DROME Alpha-Esterase-1 OS=Drosophila melanogaster OX=7227 GN=alpha-Est1 PE=2 SV=1 | 4 | 4 | 1 | 1 | 2 | 0 | 2 |
| Q9VIU8 | Q9VIU8_DROME GH25853p OS=Drosophila melanogaster OX=7227 GN=Dmel\CG10132 PE=1 SV=2 | 4 | 2 | 1 | 1 | 2 | 0 | 2 |
| Q9VIW9 | Q9VIW9_DROME Uncharacterized protein, isoform A OS=Drosophila melanogaster OX=7227 GN=Dmel\CG16771 PE=1 SV=1 | 4 | 4 | 1 | 1 | 2 | 0 | 2 |
| Q9VJ04 | Q9VJ04_DROME LD35811p OS=Drosophila melanogaster OX=7227 GN=Dmel\CG10702 PE=2 SV=1 | 4 | 4 | 1 | 1 | 2 | 0 | 2 |
| Q9VJK9 | Q9VJK9_DROME GEO06505p1 OS=Drosophila melanogaster OX=7227 GN=BEST:LD27171 PE=2 SV=2 | 4 | 2 | 1 | 1 | 2 | 0 | 2 |
| Q9VK99 | Q9VK99_DROME Atilla, isoform B OS=Drosophila melanogaster OX=7227 GN=atilla PE=4 SV=3 | 4 | 2 | 1 | 1 | 2 | 0 | 2 |
| Q9VKA1 | Q9VKA1_DROME FI02817p OS=Drosophila melanogaster OX=7227 GN=Dmel\CG6583 PE=2 SV=1 | 4 | 2 | 1 | 1 | 2 | 0 | 2 |
| Q9VKJ8 | Q9VKJ8_DROME LD15174p OS=Drosophila melanogaster OX=7227 GN=Dmel\CG6201 PE=2 SV=2 | 4 | 3 | 1 | 1 | 2 | 0 | 2 |
| Q9VLM7 | Q9VLM7_DROME UT01133p OS=Drosophila melanogaster OX=7227 GN=Dmel\CG13392 PE=1 SV=1 | 4 | 3 | 1 | 1 | 2 | 0 | 2 |
| Q9VPX8 | Q9VPX8_DROME RH48586p OS=Drosophila melanogaster OX=7227 GN=Tango14 PE=2 SV=1 | 4 | 3 | 1 | 1 | 2 | 0 | 2 |
| Q9VQK4 | Q9VQK4_DROME Alpha4GT1 OS=Drosophila melanogaster OX=7227 GN=alpha4GT1 PE=2 SV=1 | 4 | 2 | 1 | 1 | 2 | 0 | 2 |
| Q9VRZ7 | Q9VRZ7_DROME RE68357p OS=Drosophila melanogaster OX=7227 GN=Dmel\CG10075 PE=2 SV=1 | 4 | 4 | 1 | 1 | 2 | 0 | 2 |
| Q9VUL8 | Q9VUL8_DROME LD41491p OS=Drosophila melanogaster OX=7227 GN=Pex3 PE=1 SV=1 | 4 | 4 | 1 | 1 | 2 | 0 | 2 |
| Q9VVJ5 | Q9VVJ5_DROME Nedd4 family interacting protein OS=Drosophila melanogaster OX=7227 GN=Ndfip PE=2 SV=1 | 4 | 3 | 1 | 1 | 2 | 0 | 2 |
| Q9VW21 | Q9VW21_DROME Phosphatidylinositol glycan anchor biosynthesis class F OS=Drosophila melanogaster OX=7227 GN=PIG-F PE=4 SV=2 | 4 | 2 | 1 | 1 | 2 | 0 | 2 |
| Q9VWZ2 | Q9VWZ2_DROME LD30826p OS=Drosophila melanogaster OX=7227 GN=Dmel\CG7536 PE=1 SV=2 | 4 | 4 | 1 | 1 | 2 | 0 | 2 |
| Q9VXE8 | Q9VXE8_DROME RE34144p OS=Drosophila melanogaster OX=7227 GN=Ubc7 PE=1 SV=1 | 4 | 2 | 1 | 1 | 2 | 0 | 2 |
| Q9VXZ2 | Q9VXZ2_DROME RE23381p OS=Drosophila melanogaster OX=7227 GN=5541 PE=1 SV=1 | 4 | 4 | 1 | 1 | 2 | 0 | 2 |
| Q9VYJ4 | Q9VYJ4_DROME 1-acyl-sn-glycerol-3-phosphate acyltransferase OS=Drosophila melanogaster OX=7227 GN=AGPAT1 PE=2 SV=1 | 4 | 3 | 1 | 1 | 2 | 0 | 2 |
| Q9VYX8 | Q9VYX8_DROME LD04844p OS=Drosophila melanogaster OX=7227 GN=Dmel\CG1572 PE=1 SV=1 | 4 | 3 | 1 | 1 | 2 | 0 | 2 |
| Q9VZF2 | Q9VZF2_DROME LD05244p OS=Drosophila melanogaster OX=7227 GN=Dmel\CG15011 PE=2 SV=2 | 4 | 4 | 1 | 1 | 2 | 0 | 2 |
| Q9W0C5 | Q9W0C5_DROME ATP binding cassette subfamily B member 7, isoform A OS=Drosophila melanogaster OX=7227 GN=ABCB7 PE=1 SV=2 | 4 | 3 | 1 | 1 | 2 | 0 | 2 |
| Q9W133 | Q9W133_DROME AarF domain containing kinase, isoform A OS=Drosophila melanogaster OX=7227 GN=Adck PE=1 SV=2 | 4 | 4 | 1 | 1 | 2 | 0 | 2 |
| Q9W236 | Q9W236_DROME Vacuolar protein sorting 20, isoform A OS=Drosophila melanogaster OX=7227 GN=Vps20 PE=1 SV=1 | 4 | 3 | 1 | 1 | 2 | 0 | 2 |
| Q9W4A6 | Q9W4A6_DROME RE56564p OS=Drosophila melanogaster OX=7227 GN=OtopLa PE=1 SV=1 | 4 | 3 | 1 | 1 | 2 | 0 | 2 |
| Q9W4X6 | Q9W4X6_DROME Tetraspanin OS=Drosophila melanogaster OX=7227 GN=Tsp3A PE=2 SV=1 | 4 | 3 | 1 | 1 | 2 | 0 | 2 |
| Q9W569 | Q9W569_DROME GEO08311p1 OS=Drosophila melanogaster OX=7227 GN=Dmel\CG3740 PE=1 SV=1 | 4 | 2 | 1 | 1 | 2 | 0 | 2 |
| Q9Y166 | Q9Y166_DROME BcDNA.GH02431 OS=Drosophila melanogaster OX=7227 GN=Dic1 PE=1 SV=1 | 4 | 3 | 1 | 1 | 2 | 0 | 2 |
| X2J5D8 | X2J5D8_DROME Alpha-(1,3)-fucosyltransferase OS=Drosophila melanogaster OX=7227 GN=FucTB PE=1 SV=1 | 4 | 4 | 1 | 1 | 2 | 0 | 2 |
| X2J6Q6 | X2J6Q6_DROME Palmitoyltransferase OS=Drosophila melanogaster OX=7227 GN=GABPI PE=3 SV=1 | 4 | 3 | 1 | 1 | 2 | 0 | 2 |
| X2JBS2 | X2JBS2_DROME Inactivation no afterpotential E, isoform G OS=Drosophila melanogaster OX=7227 GN=inaE PE=4 SV=1 | 4 | 2 | 1 | 1 | 2 | 0 | 2 |
| P28668 | SYEP_DROME Bifunctional glutamate/proline--tRNA ligase OS=Drosophila melanogaster OX=7227 GN=GluProRS PE=1 SV=2 | 3 | 3 | 69 | 37 | 1.584962501 | 6.108524457 | -4.523561956 |
| Q8IPF5 | Q8IPF5_DROME A kinase anchor protein 200, isoform B OS=Drosophila melanogaster OX=7227 GN=Akap200 PE=1 SV=1 | 3 | 1 | 53 | 15 | 1.584962501 | 5.727920455 | -4.142957954 |
| A8DY67 | A8DY67_DROME Acetyl-CoA carboxylase, isoform E OS=Drosophila melanogaster OX=7227 GN=ACC PE=1 SV=2 | 3 | 3 | 37 | 28 | 1.584962501 | 5.209453366 | -3.624490865 |
| A1Z968 | A1Z968_DROME NAT1, isoform D OS=Drosophila melanogaster OX=7227 GN=NAT1 PE=1 SV=1 | 3 | 3 | 35 | 22 | 1.584962501 | 5.129283017 | -3.544320516 |

| UniProt ID | Protein description | # PSMs MVB spectra | # of unique peptides MVB | # of PSMs lysate | # of unique peptides lysate | log2(PSM MVB) | log2(PSM lysate) | FC (log2 of MVB PSM-log2 of lysate PSM) |
|---|---|---|---|---|---|---|---|---|
| Q9Y105 | SYQ_DROME Probable glutamine--tRNA ligase OS=Drosophila melanogaster OX=7227 GN=Aats-gln PE=2 SV=1 | 3 | 3 | 30 | 18 | 1.584962501 | 4.906890596 | -3.321928095 |
| Q9VQR8 | Q9VQR8_DROME Leucyl-tRNA synthetase, isoform A OS=Drosophila melanogaster OX=7227 GN=LeuRS PE=1 SV=2 | 3 | 3 | 29 | 16 | 1.584962501 | 4.857980995 | -3.273018494 |
| A0A0B4JD97 | A0A0B4JD97_DROME Transforming acidic coiled-coil protein, isoform K OS=Drosophila melanogaster OX=7227 GN=tacc PE=1 SV=1 | 3 | 3 | 27 | 18 | 1.584962501 | 4.754887502 | -3.169925001 |
| P80455 | RS12_DROME 40S ribosomal protein S12 OS=Drosophila melanogaster OX=7227 GN=RpS12 PE=1 SV=2 | 3 | 3 | 26 | 8 | 1.584962501 | 4.700439718 | -3.115477217 |
| Q9VVA6 | Q9VVA6_DROME Nuclear migration protein NudC OS=Drosophila melanogaster OX=7227 GN=nudC PE=1 SV=2 | 3 | 3 | 26 | 11 | 1.584962501 | 4.700439718 | -3.115477217 |
| A1Z8P9 | TPR_DROME Nucleoprotein TPR OS=Drosophila melanogaster OX=7227 GN=Mtor PE=1 SV=1 | 3 | 3 | 25 | 21 | 1.584962501 | 4.64385619 | -3.058893689 |
| Q9VVL7 | Q9VVL7_DROME Dihydrolipoyl dehydrogenase OS=Drosophila melanogaster OX=7227 GN=Dmel\CG7430 PE=1 SV=1 | 3 | 3 | 25 | 12 | 1.584962501 | 4.64385619 | -3.058893689 |
| M9PEE1 | M9PEE1_DROME Smallminded, isoform H OS=Drosophila melanogaster OX=7227 GN=smid PE=1 SV=1 | 3 | 3 | 24 | 17 | 1.584962501 | 4.584962501 | -3 |
| Q9XZ06 | NU931_DROME Nuclear pore complex protein Nup93-1 OS=Drosophila melanogaster OX=7227 GN=Nup93-1 PE=1 SV=1 | 3 | 3 | 23 | 13 | 1.584962501 | 4.523561956 | -2.938599455 |
| Q9VSS1 | Q9VSS1_DROME Pixie, isoform A OS=Drosophila melanogaster OX=7227 GN=pix PE=1 SV=1 | 3 | 3 | 23 | 16 | 1.584962501 | 4.523561956 | -2.938599455 |
| A0A0B4K7L1 | A0A0B4K7L1_DROME ATP-dependent 6-phosphofructokinase OS=Drosophila melanogaster OX=7227 GN=Pfk PE=1 SV=1 | 3 | 3 | 22 | 13 | 1.584962501 | 4.459431619 | -2.874469118 |
| Q9VA37 | DJ1B_DROME Protein dj-1beta OS=Drosophila melanogaster OX=7227 GN=dj-1beta PE=1 SV=3 | 3 | 3 | 20 | 11 | 1.584962501 | 4.321928095 | -2.736965594 |
| Q9W4H6 | Q9W4H6_DROME Pyruvate dehydrogenase E1 component subunit alpha OS=Drosophila melanogaster OX=7227 GN=Pdha PE=1 SV=2 | 3 | 2 | 20 | 8 | 1.584962501 | 4.321928095 | -2.736965594 |
| Q9VJJ0 | Q9VJJ0_DROME Proteasome subunit beta type OS=Drosophila melanogaster OX=7227 GN=Prosbeta4 PE=1 SV=1 | 3 | 2 | 19 | 11 | 1.584962501 | 4.247927513 | -2.662965013 |
| Q9XZ29 | Q9XZ29_DROME EG:BACR25B3.9 protein OS=Drosophila melanogaster OX=7227 GN=Klp3A PE=1 SV=1 | 3 | 2 | 19 | 15 | 1.584962501 | 4.247927513 | -2.662965013 |
| P43248 | MSH2_DROME DNA mismatch repair protein spellchecker 1 OS=Drosophila melanogaster OX=7227 GN=spel1 PE=3 SV=4 | 3 | 2 | 18 | 14 | 1.584962501 | 4.169925001 | -2.584962501 |
| Q9VCH5 | NUP98_DROME Nuclear pore complex protein Nup98-Nup96 OS=Drosophila melanogaster OX=7227 GN=Nup98-96 PE=1 SV=3 | 3 | 3 | 17 | 15 | 1.584962501 | 4.087462841 | -2.502500341 |
| Q9VXF9 | Q9VXF9_DROME AT13091p OS=Drosophila melanogaster OX=7227 GN=rngo PE=1 SV=1 | 3 | 3 | 17 | 10 | 1.584962501 | 4.087462841 | -2.502500341 |
| Q9W0H6 | Q9W0H6_DROME Uncharacterized protein OS=Drosophila melanogaster OX=7227 GN=CG9149 PE=1 SV=2 | 3 | 3 | 17 | 9 | 1.584962501 | 4.087462841 | -2.502500341 |
| O18373 | SPS1_DROME Selenide, water dikinase OS=Drosophila melanogaster OX=7227 GN=SelD PE=2 SV=1 | 3 | 3 | 16 | 7 | 1.584962501 | 4 | -2.415037499 |
| P23696 | PP2A_DROME Serine/threonine-protein phosphatase PP2A OS=Drosophila melanogaster OX=7227 GN=mts PE=2 SV=1 | 3 | 3 | 16 | 10 | 1.584962501 | 4 | -2.415037499 |
| A0A0B4K812 | A0A0B4K812_DROME Lissencephaly-1 homolog OS=Drosophila melanogaster OX=7227 GN=Lis-1 PE=3 SV=1 | 3 | 3 | 16 | 7 | 1.584962501 | 4 | -2.415037499 |
| Q8IR13 | Q8IR13_DROME Eukaryotic translation initiation factor 4H1, isoform B OS=Drosophila melanogaster OX=7227 GN=eIF4H1 PE=1 SV=1 | 3 | 3 | 16 | 8 | 1.584962501 | 4 | -2.415037499 |
| Q9VBP6 | Q9VBP6_DROME GH21316p OS=Drosophila melanogaster OX=7227 GN=Ssadh PE=1 SV=1 | 3 | 3 | 16 | 10 | 1.584962501 | 4 | -2.415037499 |
| Q86BI3 | Q86BI3_DROME Zinc-finger protein at 72D, isoform B OS=Drosophila melanogaster OX=7227 GN=Zn72D PE=1 SV=1 | 3 | 2 | 15 | 10 | 1.584962501 | 3.906890596 | -2.321928095 |
| C0HL66 | H33A_DROME Histone H3.3A OS=Drosophila melanogaster OX=7227 GN=His3.3A PE=1 SV=1 | 3 | 2 | 14 | 5 | 1.584962501 | 3.807354922 | -2.222392421 |
| Q9VN50 | EI3F1_DROME Eukaryotic translation initiation factor 3 subunit F-1 OS=Drosophila melanogaster OX=7227 GN=eIF3f1 PE=2 SV=1 | 3 | 3 | 14 | 8 | 1.584962501 | 3.807354922 | -2.222392421 |
| Q9VNI3 | HPF1_DROME Histone PARylation factor 1-like OS=Drosophila melanogaster OX=7227 GN=CG1218 PE=1 SV=2 | 3 | 3 | 14 | 6 | 1.584962501 | 3.807354922 | -2.222392421 |
| O77430 | O77430_DROME GEO01111p1 OS=Drosophila melanogaster OX=7227 GN=SkpA PE=1 SV=1 | 3 | 2 | 14 | 4 | 1.584962501 | 3.807354922 | -2.222392421 |
| Q7KK96 | Q7KK96_DROME Structural maintenance of chromosomes protein OS=Drosophila melanogaster OX=7227 GN=SMC2 PE=1 SV=1 | 3 | 3 | 14 | 11 | 1.584962501 | 3.807354922 | -2.222392421 |
| Q9VM69 | Q9VM69_DROME CG10206-PA OS=Drosophila melanogaster OX=7227 GN=nop5 PE=1 SV=2 | 3 | 2 | 14 | 9 | 1.584962501 | 3.807354922 | -2.222392421 |
| O44081 | DKC1_DROME H/ACA ribonucleoprotein complex subunit 4 OS=Drosophila melanogaster OX=7227 GN=Nop60B PE=1 SV=1 | 3 | 3 | 13 | 10 | 1.584962501 | 3.700439718 | -2.115477217 |
| Q9W2U4 | PP4R2_DROME Serine/threonine-protein phosphatase 4 regulatory subunit 2 OS=Drosophila melanogaster OX=7227 GN=PPP4R2r PE=1 SV=2 | 3 | 3 | 13 | 7 | 1.584962501 | 3.700439718 | -2.115477217 |
| A0A0B4KFW0 | A0A0B4KFW0_DROME Hippo, isoform B OS=Drosophila melanogaster OX=7227 GN=hpo PE=4 SV=1 | 3 | 3 | 13 | 10 | 1.584962501 | 3.700439718 | -2.115477217 |
| Q9V3D9 | Q9V3D9_DROME Signal recognition particle 54 kDa protein OS=Drosophila melanogaster OX=7227 GN=Srp54k PE=1 SV=1 | 3 | 3 | 13 | 8 | 1.584962501 | 3.700439718 | -2.115477217 |
| X2JIQ5 | X2JIQ5_DROME Ribosomal protein L17, isoform E OS=Drosophila melanogaster OX=7227 GN=RpL17 PE=3 SV=1 | 3 | 2 | 13 | 6 | 1.584962501 | 3.700439718 | -2.115477217 |
| Q7K550 | EIF3J_DROME Eukaryotic translation initiation factor 3 subunit J OS=Drosophila melanogaster OX=7227 GN=eIF3j PE=1 SV=1 | 3 | 3 | 12 | 7 | 1.584962501 | 3.584962501 | -2 |
| Q7KRY6 | NHK1_DROME Nucleosomal histone kinase 1 OS=Drosophila melanogaster OX=7227 GN=ball PE=1 SV=1 | 3 | 1 | 12 | 8 | 1.584962501 | 3.584962501 | -2 |
| A1Z934 | A1Z934_DROME CG8778 OS=Drosophila melanogaster OX=7227 GN=CG8778 PE=1 SV=1 | 3 | 3 | 12 | 6 | 1.584962501 | 3.584962501 | -2 |
| Q9VG69 | Q9VG69_DROME LP03547p OS=Drosophila melanogaster OX=7227 GN=Dmel\CG5844 PE=1 SV=1 | 3 | 3 | 12 | 8 | 1.584962501 | 3.584962501 | -2 |
| Q8IQV9 | NU205_DROME Nuclear pore complex protein Nup205 OS=Drosophila melanogaster OX=7227 GN=Nup205 PE=2 SV=1 | 3 | 3 | 11 | 10 | 1.584962501 | 3.459431619 | -1.874469118 |
| A0A0B4K6U6 | A0A0B4K6U6_DROME Squid, isoform E OS=Drosophila melanogaster OX=7227 GN=sqd PE=4 SV=1 | 3 | 3 | 11 | 4 | 1.584962501 | 3.459431619 | -1.874469118 |
| A4V448 | A4V448_DROME Glutamate-cysteine ligase catalytic subunit, isoform B OS=Drosophila melanogaster OX=7227 GN=Gclc PE=1 SV=1 | 3 | 3 | 11 | 9 | 1.584962501 | 3.459431619 | -1.874469118 |
| Q53ZT0 | Q53ZT0_DROME DNAJ-1 OS=Drosophila melanogaster OX=7227 GN=DnaJ-1 PE=1 SV=1 | 3 | 2 | 11 | 8 | 1.584962501 | 3.459431619 | -1.874469118 |
| Q7JY94 | Q7JY94_DROME Autophagy-related 7, isoform A OS=Drosophila melanogaster OX=7227 GN=Atg7 PE=1 SV=1 | 3 | 2 | 11 | 8 | 1.584962501 | 3.459431619 | -1.874469118 |
| Q9VPY7 | Q9VPY7_DROME LD42035p OS=Drosophila melanogaster OX=7227 GN=Dmel\CG5126 PE=1 SV=1 | 3 | 2 | 11 | 10 | 1.584962501 | 3.459431619 | -1.874469118 |
| Q9VUB8 | Q9VUB8_DROME Endosulfine, isoform A OS=Drosophila melanogaster OX=7227 GN=endos PE=1 SV=1 | 3 | 2 | 11 | 4 | 1.584962501 | 3.459431619 | -1.874469118 |
| P20232 | TFS2_DROME Transcription elongation factor S-II OS=Drosophila melanogaster OX=7227 GN=TfIIS PE=2 SV=1 | 3 | 2 | 10 | 5 | 1.584962501 | 3.321928095 | -1.736965594 |
| P48604 | GRPE_DROME GrpE protein homolog, mitochondrial OS=Drosophila melanogaster OX=7227 GN=Roe1 PE=2 SV=2 | 3 | 2 | 10 | 6 | 1.584962501 | 3.321928095 | -1.736965594 |
| Q9U3Z7 | NH2L1_DROME NHP2-like protein 1 homolog OS=Drosophila melanogaster OX=7227 GN=hoip PE=2 SV=1 | 3 | 3 | 10 | 4 | 1.584962501 | 3.321928095 | -1.736965594 |
| Q9VMY9 | GUAD_DROME Guanine deaminase OS=Drosophila melanogaster OX=7227 GN=DhpD PE=1 SV=1 | 3 | 3 | 10 | 8 | 1.584962501 | 3.321928095 | -1.736965594 |
| Q9W2D9 | EIF3K_DROME Eukaryotic translation initiation factor 3 subunit K OS=Drosophila melanogaster OX=7227 GN=eIF3k PE=2 SV=1 | 3 | 3 | 10 | 5 | 1.584962501 | 3.321928095 | -1.736965594 |
| Q9VNX4 | Q9VNX4_DROME Delta-1-Pyrroline-5-carboxylate dehydrogenase 1, isoform A OS=Drosophila melanogaster OX=7227 GN=P5CDh1 PE=1 SV=1 | 3 | 3 | 10 | 7 | 1.584962501 | 3.321928095 | -1.736965594 |
| P45889 | ACTZ_DROME Actin-related protein 1 OS=Drosophila melanogaster OX=7227 GN=Arp1 PE=2 SV=2 | 3 | 3 | 9 | 6 | 1.584962501 | 3.169925001 | -1.584962501 |
| Q9VC45 | ASP_DROME Protein abnormal spindle OS=Drosophila melanogaster OX=7227 GN=asp PE=1 SV=3 | 3 | 3 | 9 | 8 | 1.584962501 | 3.169925001 | -1.584962501 |
| Q9VCZ8 | PFD5_DROME Probable prefoldin subunit 5 OS=Drosophila melanogaster OX=7227 GN=Pfdn5 PE=2 SV=1 | 3 | 3 | 9 | 6 | 1.584962501 | 3.169925001 | -1.584962501 |
| Q9VQM4 | TYDP1_DROME Probable tyrosyl-DNA phosphodiesterase OS=Drosophila melanogaster OX=7227 GN=gkt PE=2 SV=1 | 3 | 2 | 9 | 6 | 1.584962501 | 3.169925001 | -1.584962501 |
| O44226 | O44226_DROME Elongin B OS=Drosophila melanogaster OX=7227 GN=EloB PE=1 SV=1 | 3 | 3 | 9 | 4 | 1.584962501 | 3.169925001 | -1.584962501 |
| Q8INE8 | Q8INE8_DROME Arginine methyltransferase 3, isoform B OS=Drosophila melanogaster OX=7227 GN=Art3 PE=1 SV=1 | 3 | 2 | 9 | 6 | 1.584962501 | 3.169925001 | -1.584962501 |
| Q9V9W7 | Q9V9W7_DROME Rough deal OS=Drosophila melanogaster OX=7227 GN=rod PE=1 SV=2 | 3 | 3 | 9 | 8 | 1.584962501 | 3.169925001 | -1.584962501 |
| Q9VWV2 | Q9VWV2_DROME Uncharacterized protein, isoform D OS=Drosophila melanogaster OX=7227 GN=CG32549 PE=1 SV=2 | 3 | 3 | 9 | 6 | 1.584962501 | 3.169925001 | -1.584962501 |
| Q9W2I2 | Q9W2I2_DROME Queuosine salvage protein OS=Drosophila melanogaster OX=7227 GN=CG9752 PE=1 SV=1 | 3 | 2 | 9 | 5 | 1.584962501 | 3.169925001 | -1.584962501 |
| Q9XZ16 | Q9XZ16_DROME Ubiquitin-like domain-containing C-terminal domain phosphatase 1 OS=Drosophila melanogaster OX=7227 GN=Ublcp1 PE=1 SV=1 | 3 | 2 | 9 | 5 | 1.584962501 | 3.169925001 | -1.584962501 |
| Q960T2 | SAP47_DROME Synapse-associated protein of 47 kDa OS=Drosophila melanogaster OX=7227 GN=Sap47 PE=1 SV=1 | 3 | 3 | 8 | 7 | 1.584962501 | 3 | -1.415037499 |
| Q9V3J4 | SEC13_DROME Protein SEC13 homolog OS=Drosophila melanogaster OX=7227 GN=Sec13 PE=1 SV=1 | 3 | 3 | 8 | 5 | 1.584962501 | 3 | -1.415037499 |
| Q9VX98 | DENR_DROME Density-regulated protein homolog OS=Drosophila melanogaster OX=7227 GN=DENR PE=1 SV=3 | 3 | 2 | 8 | 5 | 1.584962501 | 3 | -1.415037499 |
| A0A0B4K859 | A0A0B4K859_DROME CG8771, isoform C OS=Drosophila melanogaster OX=7227 GN=Nup188 PE=1 SV=1 | 3 | 2 | 8 | 7 | 1.584962501 | 3 | -1.415037499 |
| A0A0B4KH09 | A0A0B4KH09_DROME Groucho, isoform I OS=Drosophila melanogaster OX=7227 GN=gro PE=1 SV=1 | 3 | 3 | 8 | 5 | 1.584962501 | 3 | -1.415037499 |
| M9PF40 | M9PF40_DROME Prefoldin 2, isoform B OS=Drosophila melanogaster OX=7227 GN=Pfdn2 PE=4 SV=1 | 3 | 2 | 8 | 4 | 1.584962501 | 3 | -1.415037499 |
| Q541C9 | Q541C9_DROME RE18955p OS=Drosophila melanogaster OX=7227 GN=Ts PE=1 SV=1 | 3 | 3 | 8 | 7 | 1.584962501 | 3 | -1.415037499 |
| Q5BI50 | Q5BI50_DROME Cullin 4, isoform A OS=Drosophila melanogaster OX=7227 GN=Cul4 PE=1 SV=1 | 3 | 3 | 8 | 7 | 1.584962501 | 3 | -1.415037499 |
| Q7PLP4 | Q7PLP4_DROME Uncharacterized protein, isoform B OS=Drosophila melanogaster OX=7227 GN=BEST:GM10035 PE=1 SV=1 | 3 | 3 | 8 | 6 | 1.584962501 | 3 | -1.415037499 |
| Q9VB64 | Q9VB64_DROME Uncharacterized protein, isoform A OS=Drosophila melanogaster OX=7227 GN=CG31063 PE=1 SV=3 | 3 | 3 | 8 | 7 | 1.584962501 | 3 | -1.415037499 |
| Q9VQT8 | Q9VQT8_DROME Immune induced molecule 33, isoform A OS=Drosophila melanogaster OX=7227 GN=IM33 PE=1 SV=1 | 3 | 2 | 8 | 2 | 1.584962501 | 3 | -1.415037499 |
| Q9VSK9 | Q9VSK9_DROME GH24787p OS=Drosophila melanogaster OX=7227 GN=CT20937 PE=1 SV=1 | 3 | 3 | 8 | 7 | 1.584962501 | 3 | -1.415037499 |
| Q9VW13 | Q9VW13_DROME FI01412p OS=Drosophila melanogaster OX=7227 GN=CG9330-RA PE=1 SV=1 | 3 | 3 | 8 | 7 | 1.584962501 | 3 | -1.415037499 |
| O01382 | DRICE_DROME Caspase OS=Drosophila melanogaster OX=7227 GN=Drice PE=1 SV=2 | 3 | 3 | 7 | 5 | 1.584962501 | 2.807354922 | -1.222392421 |
| Q7JXF5 | NUP62_DROME Nuclear pore glycoprotein p62 OS=Drosophila melanogaster OX=7227 GN=Nup62 PE=2 SV=1 | 3 | 3 | 7 | 5 | 1.584962501 | 2.807354922 | -1.222392421 |
| D0IQK3 | D0IQK3_DROME AMP deaminase OS=Drosophila melanogaster OX=7227 GN=AMPdeam PE=1 SV=2 | 3 | 1 | 7 | 6 | 1.584962501 | 2.807354922 | -1.222392421 |
| M9PFS0 | M9PFS0_DROME Target of rapamycin, isoform B OS=Drosophila melanogaster OX=7227 GN=Tor PE=1 SV=1 | 3 | 3 | 7 | 4 | 1.584962501 | 2.807354922 | -1.222392421 |
| Q9VAI9 | Q9VAI9_DROME GEO07291p1 OS=Drosophila melanogaster OX=7227 GN=Obp99c PE=1 SV=1 | 3 | 3 | 7 | 3 | 1.584962501 | 2.807354922 | -1.222392421 |

| UniProt ID | Protein description | # PSMs MVB spectra | # of unique peptides MVB | # of PSMs lysate | # of unique peptides lysate | log2(PSM MVB) | log2(PSM lysate) | FC (log2 of MVB PSM-log2 of lysate PSM) |
|---|---|---|---|---|---|---|---|---|
| Q9VHT3 | Q9VHT3_DROME CG9617 protein OS=Drosophila melanogaster OX=7227 GN=Sgt1 PE=1 SV=1 | 3 | 2 | 7 | 5 | 1.584962501 | 2.807354922 | -1.222392421 |
| X2JCY2 | X2JCY2_DROME CTP synthase OS=Drosophila melanogaster OX=7227 GN=CTPsyn PE=3 SV=1 | 3 | 3 | 7 | 6 | 1.584962501 | 2.807354922 | -1.222392421 |
| P12370 | KAPC_DROME cAMP-dependent protein kinase catalytic subunit OS=Drosophila melanogaster OX=7227 GN=Pka-C1 PE=1 SV=3 | 3 | 3 | 6 | 5 | 1.584962501 | 2.584962501 | -1 |
| P40320 | METK_DROME S-adenosylmethionine synthase OS=Drosophila melanogaster OX=7227 GN=Sam-S PE=2 SV=2 | 3 | 3 | 6 | 5 | 1.584962501 | 2.584962501 | -1 |
| P42207 | SEPT1_DROME Septin-1 OS=Drosophila melanogaster OX=7227 GN=Sep1 PE=1 SV=1 | 3 | 3 | 6 | 3 | 1.584962501 | 2.584962501 | -1 |
| Q7K7A9 | FEN1_DROME Flap endonuclease 1 OS=Drosophila melanogaster OX=7227 GN=Fen1 PE=2 SV=1 | 3 | 3 | 6 | 5 | 1.584962501 | 2.584962501 | -1 |
| Q95RN0 | F172A_DROME FAM172 family protein homolog CG10038 OS=Drosophila melanogaster OX=7227 GN=CG10038 PE=2 SV=1 | 3 | 3 | 6 | 5 | 1.584962501 | 2.584962501 | -1 |
| M9NDU6 | M9NDU6_DROME Casein kinase II beta subunit, isoform I OS=Drosophila melanogaster OX=7227 GN=CkIIbeta PE=1 SV=1 | 3 | 2 | 6 | 4 | 1.584962501 | 2.584962501 | -1 |
| Q9VD00 | Q9VD00_DROME FI07666p OS=Drosophila melanogaster OX=7227 GN=Dmel\CG5377 PE=1 SV=1 | 3 | 2 | 6 | 5 | 1.584962501 | 2.584962501 | -1 |
| Q9VHE4 | Q9VHE4_DROME LD31229p OS=Drosophila melanogaster OX=7227 GN=Dmel\CG8132 PE=1 SV=1 | 3 | 3 | 6 | 5 | 1.584962501 | 2.584962501 | -1 |
| Q9W4J4 | Q9W4J4_DROME GM14292p OS=Drosophila melanogaster OX=7227 GN=Dmel\CG11444 PE=1 SV=1 | 3 | 3 | 6 | 3 | 1.584962501 | 2.584962501 | -1 |
| X2JER0 | X2JER0_DROME Uncharacterized protein, isoform B OS=Drosophila melanogaster OX=7227 GN=CG2200 PE=4 SV=1 | 3 | 3 | 6 | 5 | 1.584962501 | 2.584962501 | -1 |
| P25160 | ARL1_DROME ADP-ribosylation factor-like protein 1 OS=Drosophila melanogaster OX=7227 GN=Arl1 PE=2 SV=5 | 3 | 3 | 5 | 3 | 1.584962501 | 2.321928095 | -0.736965594 |
| A1Z6S6 | A1Z6S6_DROME RE54322p OS=Drosophila melanogaster OX=7227 GN=vimar PE=1 SV=1 | 3 | 3 | 5 | 3 | 1.584962501 | 2.321928095 | -0.736965594 |
| Q5U191 | Q5U191_DROME CG1882, isoform A OS=Drosophila melanogaster OX=7227 GN=CG1882 PE=1 SV=1 | 3 | 3 | 5 | 2 | 1.584962501 | 2.321928095 | -0.736965594 |
| Q7KN61 | Q7KN61_DROME Farnesyl pyrophosphate synthase OS=Drosophila melanogaster OX=7227 GN=Fpps PE=1 SV=1 | 3 | 3 | 5 | 4 | 1.584962501 | 2.321928095 | -0.736965594 |
| Q8IML4 | Q8IML4_DROME V-type proton ATPase subunit a OS=Drosophila melanogaster OX=7227 GN=Vha100-1 PE=1 SV=1 | 3 | 2 | 5 | 4 | 1.584962501 | 2.321928095 | -0.736965594 |
| Q9VBK9 | Q9VBK9_DROME Uncharacterized protein OS=Drosophila melanogaster OX=7227 GN=CG5913 PE=1 SV=1 | 3 | 3 | 5 | 4 | 1.584962501 | 2.321928095 | -0.736965594 |
| Q9VDH8 | Q9VDH8_DROME 40S ribosomal protein S30 OS=Drosophila melanogaster OX=7227 GN=RpS30 PE=1 SV=1 | 3 | 2 | 5 | 2 | 1.584962501 | 2.321928095 | -0.736965594 |
| Q9VY48 | Q9VY48_DROME Mitochondrial ribosomal protein L38 OS=Drosophila melanogaster OX=7227 GN=mRpL38 PE=1 SV=2 | 3 | 3 | 5 | 5 | 1.584962501 | 2.321928095 | -0.736965594 |
| P35415 | MYSP1_DROME Paramyosin, long form OS=Drosophila melanogaster OX=7227 GN=Prm PE=1 SV=1 | 3 | 2 | 4 | 1 | 1.584962501 | 2 | -0.415037499 |
| Q7KNF2 | PABP2_DROME Polyadenylate-binding protein 2 OS=Drosophila melanogaster OX=7227 GN=Pabp2 PE=1 SV=1 | 3 | 3 | 4 | 3 | 1.584962501 | 2 | -0.415037499 |
| Q9VHN6 | RM19_DROME 39S ribosomal protein L19, mitochondrial OS=Drosophila melanogaster OX=7227 GN=mRpL19 PE=2 SV=1 | 3 | 3 | 4 | 3 | 1.584962501 | 2 | -0.415037499 |
| Q9VN95 | ENOPH_DROME Enolase-phosphatase E1 OS=Drosophila melanogaster OX=7227 GN=CG12173 PE=2 SV=2 | 3 | 3 | 4 | 3 | 1.584962501 | 2 | -0.415037499 |
| Q7KVL7 | Q7KVL7_DROME Vacuolar protein sorting-associated protein 35 OS=Drosophila melanogaster OX=7227 GN=Vps35 PE=1 SV=2 | 3 | 3 | 4 | 4 | 1.584962501 | 2 | -0.415037499 |
| Q8SY69 | Q8SY69_DROME MICOS complex subunit MIC10 OS=Drosophila melanogaster OX=7227 GN=BcDNA:RH61753 PE=2 SV=1 | 3 | 2 | 4 | 2 | 1.584962501 | 2 | -0.415037499 |
| Q9V3E3 | Q9V3E3_DROME GM06185p OS=Drosophila melanogaster OX=7227 GN=mRpL50 PE=1 SV=1 | 3 | 2 | 4 | 3 | 1.584962501 | 2 | -0.415037499 |
| Q9VN55 | Q9VN55_DROME Antimeros, isoform A OS=Drosophila melanogaster OX=7227 GN=atms PE=1 SV=1 | 3 | 2 | 4 | 3 | 1.584962501 | 2 | -0.415037499 |
| Q27884 | PPP6_DROME Serine/threonine-protein phosphatase 6 catalytic subunit OS=Drosophila melanogaster OX=7227 GN=PpV PE=2 SV=1 | 3 | 2 | 3 | 3 | 1.584962501 | 1.584962501 | 0 |
| Q7JXF7 | SEELE_DROME Protein seele OS=Drosophila melanogaster OX=7227 GN=sel PE=1 SV=1 | 3 | 1 | 2 | 2 | 1.584962501 | 1 | 0.584962501 |
| Q7K566 | BMCP_DROME Mitochondrial uncoupling protein Bmcp OS=Drosophila melanogaster OX=7227 GN=Bmcp PE=2 SV=1 | 3 | 3 | 2 | 2 | 1.584962501 | 1 | 0.584962501 |
| Q8SX37 | CDIPT_DROME CDP-diacylglycerol--inositol 3-phosphatidyltransferase OS=Drosophila melanogaster OX=7227 GN=Pis PE=1 SV=1 | 3 | 2 | 1 | 1 | 1.584962501 | 0 | 1.584962501 |
| Q9V3N1 | SP27A_DROME Serine protease inhibitor 27A OS=Drosophila melanogaster OX=7227 GN=Spn27A PE=1 SV=1 | 3 | 3 | 2 | 2 | 1.584962501 | 1 | 0.584962501 |
| Q9VLS9 | LACB2_DROME Beta-lactamase-like protein 2 homolog OS=Drosophila melanogaster OX=7227 GN=CG12375 PE=2 SV=1 | 3 | 3 | 3 | 2 | 1.584962501 | 1.584962501 | 0 |
| Q9VLV5 | RUXE_DROME Probable small nuclear ribonucleoprotein E OS=Drosophila melanogaster OX=7227 GN=SmE PE=1 SV=1 | 3 | 3 | 2 | 2 | 1.584962501 | 1 | 0.584962501 |
| Q9VTZ6 | PMM_DROME Probable phosphomannomutase OS=Drosophila melanogaster OX=7227 GN=CG10688 PE=2 SV=1 | 3 | 3 | 3 | 3 | 1.584962501 | 1.584962501 | 0 |
| Q9W462 | UBP30_DROME Ubiquitin carboxyl-terminal hydrolase 30 homolog OS=Drosophila melanogaster OX=7227 GN=Usp30 PE=2 SV=1 | 3 | 3 | 2 | 2 | 1.584962501 | 1 | 0.584962501 |
| A1Z765 | A1Z765_DROME Uncharacterized protein, isoform A OS=Drosophila melanogaster OX=7227 GN=CG12159 PE=1 SV=1 | 3 | 1 | 2 | 1 | 1.584962501 | 1 | 0.584962501 |
| E1JJS1 | E1JJS1_DROME Glutathione S transferase T3, isoform B OS=Drosophila melanogaster OX=7227 GN=GstT3 PE=3 SV=1 | 3 | 3 | 2 | 2 | 1.584962501 | 1 | 0.584962501 |
| E2QCK0 | E2QCK0_DROME Transport and golgi organization 11, isoform B OS=Drosophila melanogaster OX=7227 GN=Tango11 PE=1 SV=1 | 3 | 3 | 1 | 1 | 1.584962501 | 0 | 1.584962501 |
| M9MSK3 | M9MSK3_DROME Tyrosine-protein kinase OS=Drosophila melanogaster OX=7227 GN=Src64B PE=3 SV=1 | 3 | 3 | 1 | 1 | 1.584962501 | 0 | 1.584962501 |
| M9PFQ2 | M9PFQ2_DROME Virus-induced RNA 1, isoform G OS=Drosophila melanogaster OX=7227 GN=vir-1 PE=1 SV=1 | 3 | 2 | 3 | 2 | 1.584962501 | 1.584962501 | 0 |
| O46308 | O46308_DROME EG:8D8.5 protein OS=Drosophila melanogaster OX=7227 GN=png PE=2 SV=1 | 3 | 3 | 1 | 1 | 1.584962501 | 0 | 1.584962501 |
| Q0KHS7 | Q0KHS7_DROME Lipid storage droplet-2, isoform B OS=Drosophila melanogaster OX=7227 GN=Lsd-2 PE=1 SV=1 | 3 | 1 | 3 | 1 | 1.584962501 | 1.584962501 | 0 |
| Q7KVF6 | Q7KVF6_DROME IP20706p OS=Drosophila melanogaster OX=7227 GN=CG33228-RA PE=2 SV=2 | 3 | 2 | 2 | 2 | 1.584962501 | 1 | 0.584962501 |
| Q8IP97 | Q8IP97_DROME Peroxin 19, isoform A OS=Drosophila melanogaster OX=7227 GN=Pex19 PE=1 SV=2 | 3 | 2 | 2 | 1 | 1.584962501 | 1 | 0.584962501 |
| Q9V3R3 | Q9V3R3_DROME CG3704-PA OS=Drosophila melanogaster OX=7227 GN=Dmel\CG3704 PE=1 SV=1 | 3 | 3 | 2 | 2 | 1.584962501 | 1 | 0.584962501 |
| Q9VGA0 | Q9VGA0_DROME Glutathione S transferase D9, isoform A OS=Drosophila melanogaster OX=7227 GN=GstD9 PE=1 SV=1 | 3 | 3 | 2 | 2 | 1.584962501 | 1 | 0.584962501 |
| Q9VI16 | Q9VI16_DROME Carnitine O-Acetyl-Transferase, isoform A OS=Drosophila melanogaster OX=7227 GN=CRAT PE=1 SV=2 | 3 | 3 | 3 | 2 | 1.584962501 | 1.584962501 | 0 |
| Q9VJ11 | Q9VJ11_DROME Calcium load-activated calcium channel OS=Drosophila melanogaster OX=7227 GN=B2 PE=1 SV=1 | 3 | 2 | 2 | 2 | 1.584962501 | 1 | 0.584962501 |
| Q9VKU3 | Q9VKU3_DROME CG6094 protein OS=Drosophila melanogaster OX=7227 GN=Dmel\CG6094 PE=1 SV=1 | 3 | 3 | 3 | 3 | 1.584962501 | 1.584962501 | 0 |
| Q9VL69 | Q9VL69_DROME LD05267p OS=Drosophila melanogaster OX=7227 GN=AI945207 PE=1 SV=1 | 3 | 1 | 2 | 1 | 1.584962501 | 1 | 0.584962501 |
| Q9VP13 | Q9VP13_DROME CR6-interacting factor OS=Drosophila melanogaster OX=7227 GN=CRIF PE=1 SV=1 | 3 | 2 | 3 | 2 | 1.584962501 | 1.584962501 | 0 |
| Q9VW57 | Q9VW57_DROME Grasp65 OS=Drosophila melanogaster OX=7227 GN=Grasp65 PE=1 SV=1 | 3 | 2 | 2 | 2 | 1.584962501 | 1 | 0.584962501 |
| Q9W127 | Q9W127_DROME CG3663 OS=Drosophila melanogaster OX=7227 GN=CG3663 PE=1 SV=1 | 3 | 2 | 2 | 2 | 1.584962501 | 1 | 0.584962501 |
| Q9W323 | Q9W323_DROME Tyrosine-protein phosphatase OS=Drosophila melanogaster OX=7227 GN=Ptpmeg2 PE=1 SV=2 | 3 | 3 | 3 | 2 | 1.584962501 | 1.584962501 | 0 |
| Q9W3T9 | Q9W3T9_DROME GM01152p OS=Drosophila melanogaster OX=7227 GN=CT9706 PE=1 SV=2 | 3 | 3 | 3 | 3 | 1.584962501 | 1.584962501 | 0 |
| Q9VBG7 | Q9VBG7_DROME Uncharacterized protein, isoform A OS=Drosophila melanogaster OX=7227 GN=CG14558 PE=1 SV=2 | 3 | 2 | 1 | 1 | 1.584962501 | 0 | 1.584962501 |
| Q9VIK0 | Q9VIK0_DROME FI06578p OS=Drosophila melanogaster OX=7227 GN=38E.13 PE=2 SV=1 | 3 | 3 | 1 | 1 | 1.584962501 | 0 | 1.584962501 |
| Q9W0K9 | Q9W0K9_DROME LD10220p OS=Drosophila melanogaster OX=7227 GN=Dmel\CG9205 PE=1 SV=1 | 3 | 3 | 1 | 1 | 1.584962501 | 0 | 1.584962501 |
| Q9W3H4 | Q9W3H4_DROME LD36273p OS=Drosophila melanogaster OX=7227 GN=sni PE=1 SV=1 | 3 | 3 | 1 | 1 | 1.584962501 | 0 | 1.584962501 |
| A0A126GUP6 | MP1_DROME Melanization protease 1 OS=Drosophila melanogaster OX=7227 GN=MP1 PE=1 SV=1 | 3 | 3 | 1 | 1 | 1.584962501 | 0 | 1.584962501 |
| P11046 | LAMB1_DROME Laminin subunit beta-1 OS=Drosophila melanogaster OX=7227 GN=LanB1 PE=1 SV=4 | 3 | 3 | 1 | 1 | 1.584962501 | 0 | 1.584962501 |
| P36975 | SNP25_DROME Synaptosomal-associated protein 25 OS=Drosophila melanogaster OX=7227 GN=Snap25 PE=2 SV=1 | 3 | 3 | 1 | 1 | 1.584962501 | 0 | 1.584962501 |
| Q00637 | SODM_DROME Superoxide dismutase [Mn], mitochondrial OS=Drosophila melanogaster OX=7227 GN=Sod2 PE=2 SV=3 | 3 | 2 | 1 | 1 | 1.584962501 | 0 | 1.584962501 |
| Q05547 | PP13_DROME Serine/threonine-protein phosphatase alpha-3 isoform OS=Drosophila melanogaster OX=7227 GN=Pp1-13C PE=1 SV=1 | 3 | 3 | 1 | 1 | 1.584962501 | 0 | 1.584962501 |
| Q86BN8 | PTPM1_DROME Phosphatidylglycerophosphatase and protein-tyrosine phosphatase 1 OS=Drosophila melanogaster OX=7227 GN=PTPMT1 PE=2 SV=1 | 3 | 3 | 1 | 1 | 1.584962501 | 0 | 1.584962501 |
| Q8MRC9 | GALT9_DROME Putative polypeptide N-acetylgalactosaminyltransferase 9 OS=Drosophila melanogaster OX=7227 GN=pgant9 PE=2 SV=2 | 3 | 3 | 1 | 1 | 1.584962501 | 0 | 1.584962501 |
| Q8T0B1 | NEPR1_DROME Nuclear envelope phosphatase-regulatory subunit 1 homolog OS=Drosophila melanogaster OX=7227 GN=CG8009 PE=2 SV=1 | 3 | 2 | 1 | 1 | 1.584962501 | 0 | 1.584962501 |
| Q95SS8 | TMM70_DROME Transmembrane protein 70 homolog, mitochondrial OS=Drosophila melanogaster OX=7227 GN=CG7506 PE=2 SV=1 | 3 | 2 | 1 | 1 | 1.584962501 | 0 | 1.584962501 |
| Q9VAY3 | MFRN_DROME Mitoferrin OS=Drosophila melanogaster OX=7227 GN=mfrn PE=2 SV=1 | 3 | 3 | 1 | 1 | 1.584962501 | 0 | 1.584962501 |
| Q9VER6 | MODSP_DROME Modular serine protease OS=Drosophila melanogaster OX=7227 GN=modSP PE=1 SV=2 | 3 | 3 | 1 | 1 | 1.584962501 | 0 | 1.584962501 |
| Q9VHT4 | FUCT1_DROME GDP-fucose transporter 1 OS=Drosophila melanogaster OX=7227 GN=nac PE=1 SV=1 | 3 | 2 | 1 | 1 | 1.584962501 | 0 | 1.584962501 |
| Q9VMW8 | MPU1_DROME Mannose-P-dolichol utilization defect 1 protein homolog OS=Drosophila melanogaster OX=7227 GN=CG3792 PE=2 SV=2 | 3 | 2 | 1 | 1 | 1.584962501 | 0 | 1.584962501 |
| Q9VP04 | ORMDL_DROME ORM1-like protein OS=Drosophila melanogaster OX=7227 GN=ORMDL PE=2 SV=1 | 3 | 3 | 1 | 1 | 1.584962501 | 0 | 1.584962501 |
| Q9W223 | CP6D2_DROME Probable cytochrome P450 6d2 OS=Drosophila melanogaster OX=7227 GN=Cyp6d2 PE=2 SV=1 | 3 | 3 | 1 | 1 | 1.584962501 | 0 | 1.584962501 |
| Q9W2D6 | TIM10_DROME Mitochondrial import inner membrane translocase subunit Tim10 OS=Drosophila melanogaster OX=7227 GN=Tim10 PE=3 SV=1 | 3 | 3 | 1 | 1 | 1.584962501 | 0 | 1.584962501 |
| Q9W3F7 | MIGA_DROME Mitoguardin OS=Drosophila melanogaster OX=7227 GN=Miga PE=1 SV=2 | 3 | 3 | 1 | 1 | 1.584962501 | 0 | 1.584962501 |
| Q9W568 | HFW1_DROME Protein halfway OS=Drosophila melanogaster OX=7227 GN=hfw PE=1 SV=3 | 3 | 2 | 1 | 1 | 1.584962501 | 0 | 1.584962501 |
| A0A0B4K6K9 | A0A0B4K6K9_DROME La related protein, isoform F OS=Drosophila melanogaster OX=7227 GN=larp PE=1 SV=1 | 3 | 3 | 1 | 1 | 1.584962501 | 0 | 1.584962501 |
| A0A0B4KER9 | A0A0B4KER9_DROME Tout-velu, isoform B OS=Drosophila melanogaster OX=7227 GN=ttv PE=4 SV=1 | 3 | 2 | 1 | 1 | 1.584962501 | 0 | 1.584962501 |

| UniProt ID | Protein description | # PSMs MVB spectra | # of unique peptides MVB | # of PSMs lysate | # of unique peptides lysate | log2(PSM MVB) | log2(PSM lysate) | FC (log2 of MVB PSM-log2 of lysate PSM) |
|---|---|---|---|---|---|---|---|---|
| A0A0B4KGI5 | A0A0B4KGI5_DROME Carboxylic ester hydrolase OS=Drosophila melanogaster OX=7227 GN=Ace PE=1 SV=1 | 3 | 3 | 1 | 1 | 1.584962501 | 0 | 1.584962501 |
| A0A0B4KHB2 | A0A0B4KHB2_DROME C-terminal binding protein, isoform G OS=Drosophila melanogaster OX=7227 GN=CtBP PE=1 SV=1 | 3 | 3 | 1 | 1 | 1.584962501 | 0 | 1.584962501 |
| A0A0B4KHK1 | A0A0B4KHK1_DROME Supernumerary limbs, isoform B OS=Drosophila melanogaster OX=7227 GN=slmb PE=4 SV=1 | 3 | 3 | 1 | 1 | 1.584962501 | 0 | 1.584962501 |
| A0A0B4LEJ6 | A0A0B4LEJ6_DROME Wunen-2, isoform B OS=Drosophila melanogaster OX=7227 GN=wun2 PE=4 SV=1 | 3 | 2 | 1 | 1 | 1.584962501 | 0 | 1.584962501 |
| A1Z705 | A1Z705_DROME CG2144 OS=Drosophila melanogaster OX=7227 GN=PIG-G PE=2 SV=1 | 3 | 2 | 1 | 1 | 1.584962501 | 0 | 1.584962501 |
| A1Z838 | A1Z838_DROME CG2292 OS=Drosophila melanogaster OX=7227 GN=PIG-N PE=4 SV=1 | 3 | 3 | 1 | 1 | 1.584962501 | 0 | 1.584962501 |
| A1Z8R6 | A1Z8R6_DROME Hexosyltransferase OS=Drosophila melanogaster OX=7227 GN=GalT1 PE=3 SV=1 | 3 | 3 | 1 | 1 | 1.584962501 | 0 | 1.584962501 |
| A1ZA77 | A1ZA77_DROME CG8320 OS=Drosophila melanogaster OX=7227 GN=CG8320-RA PE=1 SV=1 | 3 | 2 | 1 | 1 | 1.584962501 | 0 | 1.584962501 |
| A1ZBB1 | A1ZBB1_DROME Galectin OS=Drosophila melanogaster OX=7227 GN=CG5335-RA PE=2 SV=1 | 3 | 3 | 1 | 1 | 1.584962501 | 0 | 1.584962501 |
| A4UZ58 | A4UZ58_DROME Klaroid, isoform E OS=Drosophila melanogaster OX=7227 GN=koi PE=1 SV=1 | 3 | 3 | 1 | 1 | 1.584962501 | 0 | 1.584962501 |
| B7YZH3 | B7YZH3_DROME Transient receptor potential cation channel, subfamily M ortholog, isoform G OS=Drosophila melanogaster OX=7227 GN=Trpm PE=4 SV=1 | 3 | 2 | 1 | 1 | 1.584962501 | 0 | 1.584962501 |
| B7YZJ3 | B7YZJ3_DROME Uncharacterized protein OS=Drosophila melanogaster OX=7227 GN=CG42239 PE=1 SV=1 | 3 | 1 | 1 | 1 | 1.584962501 | 0 | 1.584962501 |
| B7YZR7 | B7YZR7_DROME Mapmodulin, isoform B OS=Drosophila melanogaster OX=7227 GN=Mapmodulin PE=1 SV=1 | 3 | 2 | 1 | 1 | 1.584962501 | 0 | 1.584962501 |
| B7Z0M0 | B7Z0M0_DROME FI17308p1 OS=Drosophila melanogaster OX=7227 GN=Vti1b PE=1 SV=1 | 3 | 3 | 1 | 1 | 1.584962501 | 0 | 1.584962501 |
| E1JGQ3 | E1JGQ3_DROME Glycogenin, isoform I OS=Drosophila melanogaster OX=7227 GN=Gyg PE=1 SV=1 | 3 | 2 | 1 | 1 | 1.584962501 | 0 | 1.584962501 |
| E1JHB7 | E1JHB7_DROME PDGF-and VEGF-receptor related, isoform J OS=Drosophila melanogaster OX=7227 GN=Pvr PE=1 SV=1 | 3 | 3 | 1 | 1 | 1.584962501 | 0 | 1.584962501 |
| E1JIU2 | E1JIU2_DROME Inwardly rectifying potassium channel 1, isoform C OS=Drosophila melanogaster OX=7227 GN=Irk1 PE=1 SV=1 | 3 | 3 | 1 | 1 | 1.584962501 | 0 | 1.584962501 |
| E2QCJ5 | E2QCJ5_DROME CG4050, isoform B OS=Drosophila melanogaster OX=7227 GN=Tmtc3 PE=4 SV=1 | 3 | 2 | 1 | 1 | 1.584962501 | 0 | 1.584962501 |
| M9NF14 | M9NF14_DROME IGF-II mRNA-binding protein, isoform L OS=Drosophila melanogaster OX=7227 GN=Imp PE=1 SV=1 | 3 | 3 | 1 | 1 | 1.584962501 | 0 | 1.584962501 |
| M9PB91 | M9PB91_DROME Alpha-1,3-glucosyltransferase OS=Drosophila melanogaster OX=7227 GN=gny PE=3 SV=1 | 3 | 2 | 1 | 1 | 1.584962501 | 0 | 1.584962501 |
| M9PBP5 | M9PBP5_DROME Derlin OS=Drosophila melanogaster OX=7227 GN=Der-1 PE=3 SV=1 | 3 | 1 | 1 | 1 | 1.584962501 | 0 | 1.584962501 |
| M9PBX3 | M9PBX3_DROME Uncharacterized protein, isoform H OS=Drosophila melanogaster OX=7227 GN=CG31689 PE=1 SV=1 | 3 | 2 | 1 | 1 | 1.584962501 | 0 | 1.584962501 |
| M9PCE7 | M9PCE7_DROME Uncharacterized protein, isoform E OS=Drosophila melanogaster OX=7227 GN=CG13784 PE=4 SV=1 | 3 | 2 | 1 | 1 | 1.584962501 | 0 | 1.584962501 |
| M9PCV1 | M9PCV1_DROME alpha-1,2-Mannosidase OS=Drosophila melanogaster OX=7227 GN=Edem2 PE=1 SV=1 | 3 | 3 | 1 | 1 | 1.584962501 | 0 | 1.584962501 |
| M9PCZ1 | M9PCZ1_DROME Uncharacterized protein, isoform K OS=Drosophila melanogaster OX=7227 GN=CG9449 PE=3 SV=1 | 3 | 1 | 1 | 1 | 1.584962501 | 0 | 1.584962501 |
| M9PFK7 | M9PFK7_DROME Argonaute 2, isoform E OS=Drosophila melanogaster OX=7227 GN=AGO2 PE=1 SV=1 | 3 | 3 | 1 | 1 | 1.584962501 | 0 | 1.584962501 |
| M9PHS8 | M9PHS8_DROME Uncharacterized protein, isoform C OS=Drosophila melanogaster OX=7227 GN=CG9784 PE=1 SV=1 | 3 | 3 | 1 | 1 | 1.584962501 | 0 | 1.584962501 |
| Q0E8E5 | Q0E8E5_DROME RE19996p1 OS=Drosophila melanogaster OX=7227 GN=CG7255-RE PE=2 SV=1 | 3 | 2 | 1 | 1 | 1.584962501 | 0 | 1.584962501 |
| Q6II06 | Q6II06_DROME Vesicle transport protein OS=Drosophila melanogaster OX=7227 GN=BP1075 PE=3 SV=1 | 3 | 2 | 1 | 1 | 1.584962501 | 0 | 1.584962501 |
| Q7JW07 | Q7JW07_DROME LD08826p OS=Drosophila melanogaster OX=7227 GN=CG12935 PE=1 SV=1 | 3 | 3 | 1 | 1 | 1.584962501 | 0 | 1.584962501 |
| Q7JW61 | Q7JW61_DROME LD43649p OS=Drosophila melanogaster OX=7227 GN=Usp20-33 PE=2 SV=1 | 3 | 3 | 1 | 1 | 1.584962501 | 0 | 1.584962501 |
| Q7JYK2 | Q7JYK2_DROME CG1941, isoform A OS=Drosophila melanogaster OX=7227 GN=CG1941 PE=1 SV=1 | 3 | 2 | 1 | 1 | 1.584962501 | 0 | 1.584962501 |
| Q7JYX3 | Q7JYX3_DROME GEO08273p1 OS=Drosophila melanogaster OX=7227 GN=CG15083 PE=2 SV=1 | 3 | 2 | 1 | 1 | 1.584962501 | 0 | 1.584962501 |
| Q7K2B1 | Q7K2B1_DROME CG7222, isoform A OS=Drosophila melanogaster OX=7227 GN=CG7222 PE=1 SV=1 | 3 | 3 | 1 | 1 | 1.584962501 | 0 | 1.584962501 |
| Q7K3T3 | Q7K3T3_DROME Fatty acyl-CoA reductase OS=Drosophila melanogaster OX=7227 GN=CG1441 PE=1 SV=1 | 3 | 2 | 1 | 1 | 1.584962501 | 0 | 1.584962501 |
| Q7KSP6 | Q7KSP6_DROME SET domain binding factor, isoform B OS=Drosophila melanogaster OX=7227 GN=Sbf PE=1 SV=1 | 3 | 3 | 1 | 1 | 1.584962501 | 0 | 1.584962501 |
| Q7KTX7 | Q7KTX7_DROME E3 ubiquitin-protein ligase parkin OS=Drosophila melanogaster OX=7227 GN=park PE=1 SV=1 | 3 | 3 | 1 | 1 | 1.584962501 | 0 | 1.584962501 |
| Q7KV24 | Q7KV24_DROME Uncharacterized protein OS=Drosophila melanogaster OX=7227 GN=CG15744 PE=3 SV=1 | 3 | 2 | 1 | 1 | 1.584962501 | 0 | 1.584962501 |
| Q8INW5 | Q8INW5_DROME MIP27352p OS=Drosophila melanogaster OX=7227 GN=Dmel\CG10237 PE=1 SV=1 | 3 | 3 | 1 | 1 | 1.584962501 | 0 | 1.584962501 |
| Q8MLU9 | Q8MLU9_DROME Uncharacterized protein OS=Drosophila melanogaster OX=7227 GN=CG13492 PE=4 SV=3 | 3 | 3 | 1 | 1 | 1.584962501 | 0 | 1.584962501 |
| Q9V9S4 | Q9V9S4_DROME FI21844p1 OS=Drosophila melanogaster OX=7227 GN=Dmel\CG2003 PE=2 SV=1 | 3 | 3 | 1 | 1 | 1.584962501 | 0 | 1.584962501 |
| Q9VA77 | Q9VA77_DROME Sphingomyelin phosphodiesterase OS=Drosophila melanogaster OX=7227 GN=CG15534 PE=3 SV=2 | 3 | 3 | 1 | 1 | 1.584962501 | 0 | 1.584962501 |
| Q9VAH7 | Q9VAH7_DROME RE62393p OS=Drosophila melanogaster OX=7227 GN=yata PE=1 SV=1 | 3 | 3 | 1 | 1 | 1.584962501 | 0 | 1.584962501 |
| Q9VAL5 | Q9VAL5_DROME Protein tyrosine phosphatase 99A, isoform F OS=Drosophila melanogaster OX=7227 GN=Ptp99A PE=4 SV=3 | 3 | 3 | 1 | 1 | 1.584962501 | 0 | 1.584962501 |
| Q9VB73 | Q9VB73_DROME Palmitoyltransferase OS=Drosophila melanogaster OX=7227 GN=CG5880-RA PE=2 SV=2 | 3 | 2 | 1 | 1 | 1.584962501 | 0 | 1.584962501 |
| Q9VED8 | Q9VED8_DROME Deoxyribonuclease II OS=Drosophila melanogaster OX=7227 GN=DNaseII PE=1 SV=1 | 3 | 3 | 1 | 1 | 1.584962501 | 0 | 1.584962501 |
| Q9VF06 | Q9VF06_DROME GH08991p OS=Drosophila melanogaster OX=7227 GN=Dmel\CG4287 PE=2 SV=1 | 3 | 3 | 1 | 1 | 1.584962501 | 0 | 1.584962501 |
| Q9VFE2 | Q9VFE2_DROME FI18055p1 OS=Drosophila melanogaster OX=7227 GN=Sdr PE=2 SV=1 | 3 | 3 | 1 | 1 | 1.584962501 | 0 | 1.584962501 |
| Q9VHQ5 | Q9VHQ5_DROME RE12402p OS=Drosophila melanogaster OX=7227 GN=Dmel\CG11737 PE=2 SV=1 | 3 | 1 | 1 | 1 | 1.584962501 | 0 | 1.584962501 |
| Q9VI20 | Q9VI20_DROME FI17865p1 OS=Drosophila melanogaster OX=7227 GN=godzilla PE=1 SV=1 | 3 | 2 | 1 | 1 | 1.584962501 | 0 | 1.584962501 |
| Q9VIH9 | Q9VIH9_DROME GEO04151p1 OS=Drosophila melanogaster OX=7227 GN=BcDNA:GH07967 PE=2 SV=1 | 3 | 2 | 1 | 1 | 1.584962501 | 0 | 1.584962501 |
| Q9VKD2 | Q9VKD2_DROME Very-long-chain (3R)-3-hydroxyacyl-CoA dehydratase OS=Drosophila melanogaster OX=7227 GN=Hacd1 PE=1 SV=1 | 3 | 2 | 1 | 1 | 1.584962501 | 0 | 1.584962501 |
| Q9VKQ0 | Q9VKQ0_DROME Uncharacterized protein, isoform A OS=Drosophila melanogaster OX=7227 GN=CG17130 PE=4 SV=2 | 3 | 1 | 1 | 1 | 1.584962501 | 0 | 1.584962501 |
| Q9VLH9 | Q9VLH9_DROME Alpha-mannosidase OS=Drosophila melanogaster OX=7227 GN=LManVI PE=1 SV=2 | 3 | 3 | 1 | 1 | 1.584962501 | 0 | 1.584962501 |
| Q9VMX7 | Q9VMX7_DROME Jonah 25Bi OS=Drosophila melanogaster OX=7227 GN=Jon25Bi PE=1 SV=1 | 3 | 3 | 1 | 1 | 1.584962501 | 0 | 1.584962501 |
| Q9VN28 | Q9VN28_DROME LD26442p OS=Drosophila melanogaster OX=7227 GN=Dmel\CG14650 PE=2 SV=1 | 3 | 3 | 1 | 1 | 1.584962501 | 0 | 1.584962501 |
| Q9VNA6 | Q9VNA6_DROME Ceramide kinase, isoform A OS=Drosophila melanogaster OX=7227 GN=Cerk PE=2 SV=2 | 3 | 2 | 1 | 1 | 1.584962501 | 0 | 1.584962501 |
| Q9VNL0 | Q9VNL0_DROME Gasp, isoform A OS=Drosophila melanogaster OX=7227 GN=Gasp PE=1 SV=1 | 3 | 3 | 1 | 1 | 1.584962501 | 0 | 1.584962501 |
| Q9VPA9 | Q9VPA9_DROME LD24894p OS=Drosophila melanogaster OX=7227 GN=Dmel\CG3618 PE=2 SV=1 | 3 | 3 | 1 | 1 | 1.584962501 | 0 | 1.584962501 |
| Q9VPM9 | Q9VPM9_DROME AT02829p OS=Drosophila melanogaster OX=7227 GN=Dmel\CG11601 PE=2 SV=1 | 3 | 1 | 1 | 1 | 1.584962501 | 0 | 1.584962501 |
| Q9VPR7 | Q9VPR7_DROME Coiled, isoform A OS=Drosophila melanogaster OX=7227 GN=cold PE=2 SV=1 | 3 | 2 | 1 | 1 | 1.584962501 | 0 | 1.584962501 |
| Q9VQI2 | Q9VQI2_DROME LD08641p OS=Drosophila melanogaster OX=7227 GN=CT10107 PE=2 SV=1 | 3 | 3 | 1 | 1 | 1.584962501 | 0 | 1.584962501 |
| Q9VSE8 | Q9VSE8_DROME LD39959p OS=Drosophila melanogaster OX=7227 GN=Uxs PE=2 SV=1 | 3 | 2 | 1 | 1 | 1.584962501 | 0 | 1.584962501 |
| Q9VSG9 | Q9VSG9_DROME GH24192p OS=Drosophila melanogaster OX=7227 GN=Dmel\CG13671 PE=1 SV=2 | 3 | 2 | 1 | 1 | 1.584962501 | 0 | 1.584962501 |
| Q9VSV2 | Q9VSV2_DROME Transporter OS=Drosophila melanogaster OX=7227 GN=CG4476-RB PE=2 SV=2 | 3 | 2 | 1 | 1 | 1.584962501 | 0 | 1.584962501 |
| Q9VTG0 | Q9VTG0_DROME Phosphate transporter OS=Drosophila melanogaster OX=7227 GN=NaPi-III PE=2 SV=2 | 3 | 3 | 1 | 1 | 1.584962501 | 0 | 1.584962501 |
| Q9VUV6 | Q9VUV6_DROME GXIVsPLA2, isoform A OS=Drosophila melanogaster OX=7227 GN=GXIVsPLA2 PE=2 SV=2 | 3 | 3 | 1 | 1 | 1.584962501 | 0 | 1.584962501 |
| Q9VV83 | Q9VV83_DROME Uncharacterized protein OS=Drosophila melanogaster OX=7227 GN=CG4098 PE=1 SV=1 | 3 | 2 | 1 | 1 | 1.584962501 | 0 | 1.584962501 |
| Q9VVK5 | Q9VVK5_DROME Adenosine deaminase-related growth factor A OS=Drosophila melanogaster OX=7227 GN=Adgf-A PE=1 SV=1 | 3 | 3 | 1 | 1 | 1.584962501 | 0 | 1.584962501 |
| Q9VW58 | Q9VW58_DROME LD33138p OS=Drosophila melanogaster OX=7227 GN=Dmel\CG8004 PE=1 SV=1 | 3 | 3 | 1 | 1 | 1.584962501 | 0 | 1.584962501 |
| Q9VW66 | Q9VW66_DROME Uncharacterized protein, isoform A OS=Drosophila melanogaster OX=7227 GN=BcDNA:RH69723 PE=1 SV=1 | 3 | 3 | 1 | 1 | 1.584962501 | 0 | 1.584962501 |
| Q9VX77 | Q9VX77_DROME Coiled-coil-helix-coiled-coil-helix domain containing 2, isoform A OS=Drosophila melanogaster OX=7227 GN=Chchd2 PE=1 SV=1 | 3 | 1 | 1 | 1 | 1.584962501 | 0 | 1.584962501 |
| Q9VXH4 | Q9VXH4_DROME RE16337p OS=Drosophila melanogaster OX=7227 GN=Dmel\CG9921 PE=1 SV=1 | 3 | 1 | 1 | 1 | 1.584962501 | 0 | 1.584962501 |
| Q9VXQ8 | Q9VXQ8_DROME GM02445p OS=Drosophila melanogaster OX=7227 GN=Dmel\CG11679 PE=2 SV=1 | 3 | 2 | 1 | 1 | 1.584962501 | 0 | 1.584962501 |
| Q9VXV4 | Q9VXV4_DROME RH04535p OS=Drosophila melanogaster OX=7227 GN=Dmel\CG11655 PE=1 SV=1 | 3 | 2 | 1 | 1 | 1.584962501 | 0 | 1.584962501 |
| Q9VXV6 | Q9VXV6_DROME Secretory carrier-associated membrane protein OS=Drosophila melanogaster OX=7227 GN=Scamp PE=1 SV=1 | 3 | 2 | 1 | 1 | 1.584962501 | 0 | 1.584962501 |
| Q9VYL5 | Q9VYL5_DROME GH19726p OS=Drosophila melanogaster OX=7227 GN=Dmel\CG1824 PE=1 SV=1 | 3 | 2 | 1 | 1 | 1.584962501 | 0 | 1.584962501 |
| Q9VYR2 | Q9VYR2_DROME RE13973p OS=Drosophila melanogaster OX=7227 GN=Dmel\CG1492 PE=2 SV=2 | 3 | 3 | 1 | 1 | 1.584962501 | 0 | 1.584962501 |
| Q9VYX5 | Q9VYX5_DROME CG11699-RA protein OS=Drosophila melanogaster OX=7227 GN=Dmel\CG11699 PE=2 SV=1 | 3 | 2 | 1 | 1 | 1.584962501 | 0 | 1.584962501 |
| Q9VZA6 | Q9VZA6_DROME Hexosyltransferase OS=Drosophila melanogaster OX=7227 GN=BcDNA:GM04393 PE=2 SV=1 | 3 | 2 | 1 | 1 | 1.584962501 | 0 | 1.584962501 |
| Q9W1H9 | Q9W1H9_DROME CG5569 OS=Drosophila melanogaster OX=7227 GN=CG5569 PE=2 SV=1 | 3 | 3 | 1 | 1 | 1.584962501 | 0 | 1.584962501 |

| UniProt ID | Protein description | # PSMs MVB spectra | # of unique peptides MVB | # of PSMs lysate | # of unique peptides lysate | log2(PSM MVB) | log2(PSM lysate) | FC (log2 of MVB PSM-log2 of lysate PSM) |
|---|---|---|---|---|---|---|---|---|
| Q9W1R8 | Q9W1R8_DROME CG3500, isoform A OS=Drosophila melanogaster OX=7227 GN=CG3500-RA PE=2 SV=1 | 3 | 2 | 1 | 1 | 1.584962501 | 0 | 1.584962501 |
| Q9W258 | Q9W258_DROME Babos, isoform A OS=Drosophila melanogaster OX=7227 GN=babos PE=2 SV=2 | 3 | 2 | 1 | 1 | 1.584962501 | 0 | 1.584962501 |
| Q9W280 | Q9W280_DROME GH11358p OS=Drosophila melanogaster OX=7227 GN=CG11170 PE=2 SV=1 | 3 | 3 | 1 | 1 | 1.584962501 | 0 | 1.584962501 |
| Q9W380 | Q9W380_DROME RH45008p OS=Drosophila melanogaster OX=7227 GN=BcDNA:RH45008 PE=2 SV=1 | 3 | 2 | 1 | 1 | 1.584962501 | 0 | 1.584962501 |
| Q9W513 | Q9W513_DROME GEO10196p1 OS=Drosophila melanogaster OX=7227 GN=Dmel\CG17776 PE=2 SV=1 | 3 | 1 | 1 | 1 | 1.584962501 | 0 | 1.584962501 |
| E1JJD6 | E1JJD6_DROME Twenty-four, isoform F OS=Drosophila melanogaster OX=7227 GN=tyf PE=1 SV=2 | 2 | 2 | 35 | 16 | 1 | 5.129283017 | -4.129283017 |
| P48809 | RB27C_DROME Heterogeneous nuclear ribonucleoprotein 27C OS=Drosophila melanogaster OX=7227 GN=Hrb27C PE=1 SV=2 | 2 | 2 | 29 | 12 | 1 | 4.857980995 | -3.857980995 |
| Q9XYZ5 | DDB1_DROME DNA damage-binding protein 1 OS=Drosophila melanogaster OX=7227 GN=pic PE=1 SV=1 | 2 | 2 | 26 | 15 | 1 | 4.700439718 | -3.700439718 |
| Q8MZI3 | Q8MZI3_DROME GH10652p OS=Drosophila melanogaster OX=7227 GN=BcDNA:HL01868 PE=1 SV=1 | 2 | 2 | 26 | 12 | 1 | 4.700439718 | -3.700439718 |
| Q9V3W7 | Q9V3W7_DROME LD40489p OS=Drosophila melanogaster OX=7227 GN=SF2 PE=1 SV=1 | 2 | 2 | 24 | 9 | 1 | 4.584962501 | -3.584962501 |
| O76922 | AUB_DROME Protein aubergine OS=Drosophila melanogaster OX=7227 GN=aub PE=1 SV=1 | 2 | 2 | 23 | 16 | 1 | 4.523561956 | -3.523561956 |
| Q9V3E9 | Q9V3E9_DROME FI17138p1 OS=Drosophila melanogaster OX=7227 GN=spag PE=1 SV=1 | 2 | 2 | 23 | 14 | 1 | 4.523561956 | -3.523561956 |
| Q9VYT3 | Q9VYT3_DROME FI23714p1 OS=Drosophila melanogaster OX=7227 GN=Nrd1 PE=1 SV=2 | 2 | 2 | 23 | 19 | 1 | 4.523561956 | -3.523561956 |
| Q23972 | SMG_DROME Protein Smaug OS=Drosophila melanogaster OX=7227 GN=smg PE=1 SV=2 | 2 | 2 | 22 | 11 | 1 | 4.459431619 | -3.459431619 |
| Q9VKD3 | NFS1_DROME Probable cysteine desulfurase, mitochondrial OS=Drosophila melanogaster OX=7227 GN=CG12264 PE=2 SV=1 | 2 | 1 | 22 | 12 | 1 | 4.459431619 | -3.459431619 |
| A1ZBW0 | A1ZBW0_DROME Heterogeneous nuclear ribonucleoprotein K, isoform C OS=Drosophila melanogaster OX=7227 GN=HnRNP-K PE=1 SV=1 | 2 | 2 | 22 | 11 | 1 | 4.459431619 | -3.459431619 |
| Q24154 | RL29_DROME 60S ribosomal protein L29 OS=Drosophila melanogaster OX=7227 GN=RpL29 PE=1 SV=1 | 2 | 1 | 21 | 4 | 1 | 4.392317423 | -3.392317423 |
| Q9VH81 | Q9VH81_DROME GH12714p OS=Drosophila melanogaster OX=7227 GN=PpD3 PE=1 SV=1 | 2 | 2 | 21 | 12 | 1 | 4.392317423 | -3.392317423 |
| Q9VQQ6 | Q9VQQ6_DROME FI18122p1 OS=Drosophila melanogaster OX=7227 GN=Snx1 PE=1 SV=1 | 2 | 2 | 20 | 13 | 1 | 4.321928095 | -3.321928095 |
| M9PCG1 | M9PCG1_DROME Dihydrolipoamide acetyltransferase component of pyruvate dehydrogenase complex OS=Drosophila melanogaster OX=7227 GN=muc PE=1 SV=1 | 2 | 2 | 19 | 10 | 1 | 4.247927513 | -3.247927513 |
| P04052 | RPB1_DROME DNA-directed RNA polymerase II subunit RPB1 OS=Drosophila melanogaster OX=7227 GN=RpII215 PE=3 SV=4 | 2 | 2 | 18 | 14 | 1 | 4.169925001 | -3.169925001 |
| Q24368 | ISWI_DROME Chromatin-remodeling complex ATPase chain Iswi OS=Drosophila melanogaster OX=7227 GN=Iswi PE=1 SV=1 | 2 | 2 | 17 | 14 | 1 | 4.087462841 | -3.087462841 |
| Q9U9Q4 | EIF3H_DROME Eukaryotic translation initiation factor 3 subunit H OS=Drosophila melanogaster OX=7227 GN=eIF3h PE=1 SV=2 | 2 | 2 | 17 | 9 | 1 | 4.087462841 | -3.087462841 |
| Q6NL44 | Q6NL44_DROME GH28815p OS=Drosophila melanogaster OX=7227 GN=CG18238 PE=1 SV=1 | 2 | 2 | 17 | 11 | 1 | 4.087462841 | -3.087462841 |
| Q7JRH5 | Q7JRH5_DROME CG5726 OS=Drosophila melanogaster OX=7227 GN=CG5726 PE=1 SV=1 | 2 | 2 | 17 | 12 | 1 | 4.087462841 | -3.087462841 |
| Q9VHR5 | Q9VHR5_DROME Uncharacterized protein OS=Drosophila melanogaster OX=7227 GN=CG9684 PE=1 SV=2 | 2 | 2 | 17 | 10 | 1 | 4.087462841 | -3.087462841 |
| Q9VRV8 | Q9VRV8_DROME FI21453p1 OS=Drosophila melanogaster OX=7227 GN=Tnpo PE=1 SV=1 | 2 | 2 | 17 | 7 | 1 | 4.087462841 | -3.087462841 |
| P32392 | ARP3_DROME Actin-related protein 3 OS=Drosophila melanogaster OX=7227 GN=Arp3 PE=2 SV=3 | 2 | 2 | 16 | 10 | 1 | 4 | -3 |
| Q9VGR7 | Y6693_DROME J domain-containing protein CG6693 OS=Drosophila melanogaster OX=7227 GN=CG6693 PE=1 SV=1 | 2 | 2 | 16 | 6 | 1 | 4 | -3 |
| Q9VZ20 | Q9VZ20_DROME Dynein light intermediate chain, isoform A OS=Drosophila melanogaster OX=7227 GN=Dlic PE=1 SV=2 | 2 | 2 | 16 | 9 | 1 | 4 | -3 |
| Q24311 | CUL1_DROME Cullin homolog 1 OS=Drosophila melanogaster OX=7227 GN=Cul1 PE=1 SV=2 | 2 | 2 | 15 | 11 | 1 | 3.906890596 | -2.906890596 |
| Q9W1V3 | FBRL_DROME rRNA 2'-O-methyltransferase fibrillarin OS=Drosophila melanogaster OX=7227 GN=Fib PE=2 SV=1 | 2 | 2 | 15 | 9 | 1 | 3.906890596 | -2.906890596 |
| Q9W4M9 | NSUN2_DROME tRNA (cytosine(34)-C(5))-methyltransferase OS=Drosophila melanogaster OX=7227 GN=Nsun2 PE=2 SV=1 | 2 | 2 | 15 | 10 | 1 | 3.906890596 | -2.906890596 |
| Q9W1N4 | Q9W1N4_DROME HL01517p OS=Drosophila melanogaster OX=7227 GN=wmd PE=1 SV=1 | 2 | 2 | 15 | 11 | 1 | 3.906890596 | -2.906890596 |
| Q9W330 | Q9W330_DROME Phosphotransferase OS=Drosophila melanogaster OX=7227 GN=Hex-A PE=1 SV=1 | 2 | 2 | 15 | 10 | 1 | 3.906890596 | -2.906890596 |
| P32234 | 128UP_DROME GTP-binding protein 128up OS=Drosophila melanogaster OX=7227 GN=128up PE=2 SV=2 | 2 | 2 | 14 | 10 | 1 | 3.807354922 | -2.807354922 |
| Q9VDQ7 | NAA25_DROME Phagocyte signaling-impaired protein OS=Drosophila melanogaster OX=7227 GN=psidin PE=2 SV=1 | 2 | 2 | 14 | 12 | 1 | 3.807354922 | -2.807354922 |
| O97182 | O97182_DROME Actin-related protein 2/3 complex subunit OS=Drosophila melanogaster OX=7227 GN=Arpc1 PE=1 SV=1 | 2 | 2 | 14 | 12 | 1 | 3.807354922 | -2.807354922 |
| Q9VDV2 | Q9VDV2_DROME AT06125p OS=Drosophila melanogaster OX=7227 GN=BEST:GH15838 PE=1 SV=1 | 2 | 2 | 14 | 11 | 1 | 3.807354922 | -2.807354922 |
| Q7K2D2 | DCTN2_DROME Dynactin subunit 2 OS=Drosophila melanogaster OX=7227 GN=DCTN2-p50 PE=1 SV=1 | 2 | 2 | 13 | 8 | 1 | 3.700439718 | -2.700439718 |
| A0A0B4K7G4 | A0A0B4K7G4_DROME Modulo, isoform C OS=Drosophila melanogaster OX=7227 GN=mod PE=1 SV=1 | 2 | 2 | 13 | 8 | 1 | 3.700439718 | -2.700439718 |
| Q8IN05 | Q8IN05_DROME Liquid facets-Related, isoform D OS=Drosophila melanogaster OX=7227 GN=lqfR PE=1 SV=2 | 2 | 2 | 13 | 8 | 1 | 3.700439718 | -2.700439718 |
| Q9VC48 | Q9VC48_DROME Methionine aminopeptidase OS=Drosophila melanogaster OX=7227 GN=Dmel\CG13630 PE=1 SV=1 | 2 | 2 | 13 | 9 | 1 | 3.700439718 | -2.700439718 |
| A0A0B4LH64 | A0A0B4LH64_DROME Ataxin-2, isoform D OS=Drosophila melanogaster OX=7227 GN=Atx2 PE=4 SV=1 | 2 | 1 | 12 | 8 | 1 | 3.584962501 | -2.584962501 |
| A1ZBM2 | A1ZBM2_DROME Tubulin-binding cofactor B OS=Drosophila melanogaster OX=7227 GN=TBCB PE=1 SV=1 | 2 | 2 | 12 | 7 | 1 | 3.584962501 | -2.584962501 |
| E2QC56 | E2QC56_DROME Uncharacterized protein, isoform B OS=Drosophila melanogaster OX=7227 GN=CG10417 PE=1 SV=1 | 2 | 2 | 12 | 8 | 1 | 3.584962501 | -2.584962501 |
| M9PC99 | M9PC99_DROME Secretory 24CD, isoform C OS=Drosophila melanogaster OX=7227 GN=Sec24CD PE=1 SV=1 | 2 | 2 | 12 | 11 | 1 | 3.584962501 | -2.584962501 |
| Q8SXC2 | Q8SXC2_DROME FI04487p OS=Drosophila melanogaster OX=7227 GN=Dmel\CG6950 PE=1 SV=1 | 2 | 2 | 12 | 11 | 1 | 3.584962501 | -2.584962501 |
| Q9VV39 | Q9VV39_DROME Mitochondrial ribosomal protein S34 OS=Drosophila melanogaster OX=7227 GN=mRpS34 PE=1 SV=2 | 2 | 2 | 12 | 7 | 1 | 3.584962501 | -2.584962501 |
| Q9VYG8 | Q9VYG8_DROME CG15717-PA OS=Drosophila melanogaster OX=7227 GN=Dmel\CG15717 PE=1 SV=1 | 2 | 2 | 12 | 6 | 1 | 3.584962501 | -2.584962501 |
| Q9W2E7 | Q9W2E7_DROME Rae1 OS=Drosophila melanogaster OX=7227 GN=Rae1 PE=1 SV=1 | 2 | 2 | 12 | 9 | 1 | 3.584962501 | -2.584962501 |
| O61443 | MK38B_DROME Mitogen-activated protein kinase p38b OS=Drosophila melanogaster OX=7227 GN=p38b PE=1 SV=1 | 2 | 2 | 11 | 7 | 1 | 3.459431619 | -2.459431619 |
| Q8SY96 | NFU1_DROME NFU1 iron-sulfur cluster scaffold homolog, mitochondrial OS=Drosophila melanogaster OX=7227 GN=CG32500 PE=2 SV=1 | 2 | 2 | 10 | 6 | 1 | 3.321928095 | -2.321928095 |
| Q9GYU8 | NUP88_DROME Nuclear pore complex protein Nup88 OS=Drosophila melanogaster OX=7227 GN=mbo PE=1 SV=2 | 2 | 2 | 10 | 9 | 1 | 3.321928095 | -2.321928095 |
| Q9I7K0 | JUPIT_DROME Microtubule-associated protein Jupiter OS=Drosophila melanogaster OX=7227 GN=Jupiter PE=1 SV=2 | 2 | 2 | 10 | 4 | 1 | 3.321928095 | -2.321928095 |
| Q9VBX1 | NEMF_DROME Nuclear export mediator factor NEMF homolog OS=Drosophila melanogaster OX=7227 GN=Clbn PE=1 SV=2 | 2 | 2 | 10 | 6 | 1 | 3.321928095 | -2.321928095 |
| Q9VLR5 | TCP4_DROME RNA polymerase II transcriptional coactivator OS=Drosophila melanogaster OX=7227 GN=Ssb-c31a PE=2 SV=1 | 2 | 2 | 10 | 6 | 1 | 3.321928095 | -2.321928095 |
| M9PEU5 | M9PEU5_DROME NEDD8-activating enzyme E1 regulatory subunit OS=Drosophila melanogaster OX=7227 GN=APP-BP1 PE=3 SV=1 | 2 | 2 | 10 | 6 | 1 | 3.321928095 | -2.321928095 |
| Q9V464 | ASF1_DROME Histone chaperone asf1 OS=Drosophila melanogaster OX=7227 GN=asf1 PE=1 SV=1 | 2 | 2 | 9 | 6 | 1 | 3.169925001 | -2.169925001 |
| Q9VNV3 | DDX1_DROME ATP-dependent RNA helicase Ddx1 OS=Drosophila melanogaster OX=7227 GN=Ddx1 PE=2 SV=1 | 2 | 1 | 9 | 8 | 1 | 3.169925001 | -2.169925001 |
| Q9VVI2 | EDC3_DROME Enhancer of mRNA-decapping protein 3 OS=Drosophila melanogaster OX=7227 GN=Edc3 PE=1 SV=1 | 2 | 2 | 9 | 8 | 1 | 3.169925001 | -2.169925001 |
| Q9W2N0 | CAPZA_DROME F-actin-capping protein subunit alpha OS=Drosophila melanogaster OX=7227 GN=cpa PE=2 SV=1 | 2 | 2 | 9 | 5 | 1 | 3.169925001 | -2.169925001 |
| B7YZZ9 | B7YZZ9_DROME DNA-(apurinic or apyrimidinic site) lyase OS=Drosophila melanogaster OX=7227 GN=Rrp1 PE=1 SV=1 | 2 | 2 | 9 | 7 | 1 | 3.169925001 | -2.169925001 |
| Q7K1H0 | Q7K1H0_DROME GH09096p OS=Drosophila melanogaster OX=7227 GN=CG6491 PE=1 SV=1 | 2 | 2 | 9 | 7 | 1 | 3.169925001 | -2.169925001 |
| Q8MMD3 | Q8MMD3_DROME Epidermal growth factor receptor pathway substrate clone 15, isoform C OS=Drosophila melanogaster OX=7227 GN=Eps-15 PE=1 SV=1 | 2 | 2 | 9 | 7 | 1 | 3.169925001 | -2.169925001 |
| Q8SYL1 | Q8SYL1_DROME Lethal (2) 09851, isoform A OS=Drosophila melanogaster OX=7227 GN=l(2)09851 PE=1 SV=1 | 2 | 2 | 9 | 6 | 1 | 3.169925001 | -2.169925001 |
| Q9VYR1 | Q9VYR1_DROME Regucalcin homologue OS=Drosophila melanogaster OX=7227 GN=regucalcin PE=1 SV=1 | 2 | 2 | 9 | 7 | 1 | 3.169925001 | -2.169925001 |
| Q9W2Z4 | Q9W2Z4_DROME Uncharacterized protein, isoform C OS=Drosophila melanogaster OX=7227 GN=CG2990 PE=4 SV=2 | 2 | 2 | 9 | 7 | 1 | 3.169925001 | -2.169925001 |
| Q9Y0Y5 | Q9Y0Y5_DROME Coatomer subunit epsilon OS=Drosophila melanogaster OX=7227 GN=epsilonCOP PE=1 SV=1 | 2 | 2 | 9 | 7 | 1 | 3.169925001 | -2.169925001 |
| A1ZAW5 | ICLN_DROME Methylosome subunit pICln OS=Drosophila melanogaster OX=7227 GN=icln PE=1 SV=1 | 2 | 2 | 8 | 6 | 1 | 3 | -2 |
| Q960X8 | HRS_DROME Hepatocyte growth factor-regulated tyrosine kinase substrate OS=Drosophila melanogaster OX=7227 GN=Hrs PE=1 SV=1 | 2 | 2 | 8 | 5 | 1 | 3 | -2 |
| Q9VJC7 | EFTS_DROME Elongation factor Ts, mitochondrial OS=Drosophila melanogaster OX=7227 GN=mEFTs PE=2 SV=1 | 2 | 2 | 8 | 5 | 1 | 3 | -2 |
| Q7K2X8 | Q7K2X8_DROME Nucleoporin at 44A, isoform A OS=Drosophila melanogaster OX=7227 GN=Nup44A PE=1 SV=1 | 2 | 2 | 8 | 7 | 1 | 3 | -2 |
| Q9U1K7 | Q9U1K7_DROME EG:140G11.5 protein OS=Drosophila melanogaster OX=7227 GN=EG:140G11.5 PE=1 SV=1 | 2 | 2 | 8 | 4 | 1 | 3 | -2 |
| Q9VSL9 | Q9VSL9_DROME RE62833p OS=Drosophila melanogaster OX=7227 GN=Dmel\CG6638 PE=1 SV=1 | 2 | 2 | 8 | 6 | 1 | 3 | -2 |
| Q9W074 | Q9W074_DROME FI04779p OS=Drosophila melanogaster OX=7227 GN=HBS1 PE=1 SV=2 | 2 | 2 | 8 | 6 | 1 | 3 | -2 |
| P07183 | CH38_DROME Chorion protein S38 OS=Drosophila melanogaster OX=7227 GN=Cp38 PE=2 SV=1 | 2 | 2 | 7 | 5 | 1 | 2.807354922 | -1.807354922 |
| P49028 | MGN_DROME Protein mago nashi OS=Drosophila melanogaster OX=7227 GN=mago PE=1 SV=1 | 2 | 2 | 7 | 4 | 1 | 2.807354922 | -1.807354922 |
| Q9NHD5 | SAN_DROME Probable N-acetyltransferase san OS=Drosophila melanogaster OX=7227 GN=san PE=1 SV=1 | 2 | 2 | 7 | 6 | 1 | 2.807354922 | -1.807354922 |
| Q9VAH9 | INT11_DROME Integrator complex subunit 11 OS=Drosophila melanogaster OX=7227 GN=IntS11 PE=1 SV=1 | 2 | 2 | 7 | 7 | 1 | 2.807354922 | -1.807354922 |
| Q9VB62 | DPOA2_DROME DNA polymerase alpha subunit B OS=Drosophila melanogaster OX=7227 GN=DNApol-alpha73 PE=1 SV=2 | 2 | 1 | 7 | 6 | 1 | 2.807354922 | -1.807354922 |

| UniProt ID | Protein description | # PSMs MVB spectra | # of unique peptides MVB | # of PSMs lysate | # of unique peptides lysate | log2(PSM MVB) | log2(PSM lysate) | FC (log2 of MVB PSM-log2 of lysate PSM) |
|---|---|---|---|---|---|---|---|---|
| M9NEH9 | M9NEH9_DROME Mushroom-body expressed, isoform K OS=Drosophila melanogaster OX=7227 GN=mub PE=1 SV=1 | 2 | 2 | 7 | 3 | 1 | 2.807354922 | -1.807354922 |
| O62531 | O62531_DROME Adaptor protein complex 1, mu subunit OS=Drosophila melanogaster OX=7227 GN=AP-1mu PE=1 SV=1 | 2 | 2 | 7 | 7 | 1 | 2.807354922 | -1.807354922 |
| Q7JVZ8 | Q7JVZ8_DROME Glutathione S transferase E11, isoform A OS=Drosophila melanogaster OX=7227 GN=GstE11 PE=1 SV=1 | 2 | 2 | 7 | 4 | 1 | 2.807354922 | -1.807354922 |
| Q8IMF8 | Q8IMF8_DROME Hephaestus, isoform C OS=Drosophila melanogaster OX=7227 GN=heph PE=1 SV=1 | 2 | 2 | 7 | 4 | 1 | 2.807354922 | -1.807354922 |
| Q9VG79 | Q9VG79_DROME Dipeptidase C, isoform A OS=Drosophila melanogaster OX=7227 GN=Dip-C PE=1 SV=2 | 2 | 2 | 7 | 6 | 1 | 2.807354922 | -1.807354922 |
| Q94517 | HDAC1_DROME Histone deacetylase Rpd3 OS=Drosophila melanogaster OX=7227 GN=Rpd3 PE=1 SV=2 | 2 | 2 | 6 | 4 | 1 | 2.584962501 | -1.584962501 |
| Q9VHD3 | MAAI1_DROME Probable maleylacetoacetate isomerase 1 OS=Drosophila melanogaster OX=7227 GN=GstZ1 PE=1 SV=1 | 2 | 1 | 6 | 4 | 1 | 2.584962501 | -1.584962501 |
| Q9VLU0 | BAF_DROME Barrier-to-autointegration factor OS=Drosophila melanogaster OX=7227 GN=baf PE=1 SV=1 | 2 | 2 | 6 | 3 | 1 | 2.584962501 | -1.584962501 |
| Q9VWS2 | NUP35_DROME Nucleoporin Nup35 OS=Drosophila melanogaster OX=7227 GN=Nup35 PE=1 SV=1 | 2 | 2 | 6 | 5 | 1 | 2.584962501 | -1.584962501 |
| Q7JW03 | Q7JW03_DROME CG2257 OS=Drosophila melanogaster OX=7227 GN=UbcE2H PE=1 SV=1 | 2 | 2 | 6 | 4 | 1 | 2.584962501 | -1.584962501 |
| Q7K738 | Q7K738_DROME EG:52C10.4 protein OS=Drosophila melanogaster OX=7227 GN=Ubc10 PE=1 SV=1 | 2 | 2 | 6 | 4 | 1 | 2.584962501 | -1.584962501 |
| Q7KNM2 | Q7KNM2_DROME Dorsal interacting protein 4 OS=Drosophila melanogaster OX=7227 GN=lwr PE=1 SV=1 | 2 | 2 | 6 | 2 | 1 | 2.584962501 | -1.584962501 |
| Q9VCK6 | Q9VCK6_DROME LD34157p OS=Drosophila melanogaster OX=7227 GN=Dmel\CG10184 PE=1 SV=1 | 2 | 2 | 6 | 4 | 1 | 2.584962501 | -1.584962501 |
| Q9VCV4 | Q9VCV4_DROME Iron regulatory protein 1A OS=Drosophila melanogaster OX=7227 GN=Irp-1A PE=1 SV=1 | 2 | 2 | 6 | 5 | 1 | 2.584962501 | -1.584962501 |
| Q9VDI1 | Q9VDI1_DROME LD46328p OS=Drosophila melanogaster OX=7227 GN=Synd PE=1 SV=2 | 2 | 2 | 6 | 6 | 1 | 2.584962501 | -1.584962501 |
| Q9VFE7 | Q9VFE7_DROME Nuclear pore protein OS=Drosophila melanogaster OX=7227 GN=Nup93-2 PE=1 SV=1 | 2 | 2 | 6 | 6 | 1 | 2.584962501 | -1.584962501 |
| Q9VG33 | Q9VG33_DROME Sulfurtransferase OS=Drosophila melanogaster OX=7227 GN=BcDNA:RH54517 PE=1 SV=2 | 2 | 2 | 6 | 4 | 1 | 2.584962501 | -1.584962501 |
| Q9VMH2 | Q9VMH2_DROME Actin-related protein 2/3 complex subunit 4 OS=Drosophila melanogaster OX=7227 GN=Arpc4 PE=1 SV=1 | 2 | 2 | 6 | 5 | 1 | 2.584962501 | -1.584962501 |
| Q9VQQ0 | Q9VQQ0_DROME Serine/threonine-protein phosphatase 2A activator OS=Drosophila melanogaster OX=7227 GN=Ptpa PE=1 SV=1 | 2 | 2 | 6 | 4 | 1 | 2.584962501 | -1.584962501 |
| P25171 | RCC1_DROME Regulator of chromosome condensation OS=Drosophila melanogaster OX=7227 GN=Rcc1 PE=1 SV=2 | 2 | 2 | 5 | 5 | 1 | 2.321928095 | -1.321928095 |
| P54359 | SEPT2_DROME Septin-2 OS=Drosophila melanogaster OX=7227 GN=Sep2 PE=2 SV=2 | 2 | 2 | 5 | 5 | 1 | 2.321928095 | -1.321928095 |
| Q9V431 | API5_DROME Apoptosis inhibitor 5 homolog OS=Drosophila melanogaster OX=7227 GN=Aac11 PE=2 SV=1 | 2 | 2 | 5 | 5 | 1 | 2.321928095 | -1.321928095 |
| Q9V877 | SUB_DROME Kinesin-like protein subito OS=Drosophila melanogaster OX=7227 GN=sub PE=1 SV=1 | 2 | 2 | 5 | 5 | 1 | 2.321928095 | -1.321928095 |
| Q9VVA0 | Y9705_DROME Cold shock domain-containing protein CG9705 OS=Drosophila melanogaster OX=7227 GN=CG9705 PE=1 SV=1 | 2 | 2 | 5 | 3 | 1 | 2.321928095 | -1.321928095 |
| Q9VZ64 | 6PGL_DROME Probable 6-phosphogluconolactonase OS=Drosophila melanogaster OX=7227 GN=CG17333 PE=2 SV=1 | 2 | 2 | 5 | 2 | 1 | 2.321928095 | -1.321928095 |
| Q9W1X8 | FCL_DROME Probable GDP-L-fucose synthase OS=Drosophila melanogaster OX=7227 GN=Gmer PE=2 SV=1 | 2 | 1 | 5 | 3 | 1 | 2.321928095 | -1.321928095 |
| Q9W328 | LST8_DROME Protein LST8 homolog OS=Drosophila melanogaster OX=7227 GN=Lst8 PE=1 SV=2 | 2 | 2 | 5 | 3 | 1 | 2.321928095 | -1.321928095 |
| M9PG47 | M9PG47_DROME 40S ribosomal protein S26 OS=Drosophila melanogaster OX=7227 GN=RpS26 PE=3 SV=1 | 2 | 1 | 5 | 1 | 1 | 2.321928095 | -1.321928095 |
| Q0E8X8 | Q0E8X8_DROME FI01416p OS=Drosophila melanogaster OX=7227 GN=CG13551 PE=1 SV=1 | 2 | 1 | 5 | 2 | 1 | 2.321928095 | -1.321928095 |
| Q7K1T1 | Q7K1T1_DROME LD44824p OS=Drosophila melanogaster OX=7227 GN=vlc PE=1 SV=1 | 2 | 2 | 5 | 5 | 1 | 2.321928095 | -1.321928095 |
| Q7K561 | Q7K561_DROME CG8503 OS=Drosophila melanogaster OX=7227 GN=SmydA-1 PE=2 SV=1 | 2 | 2 | 5 | 5 | 1 | 2.321928095 | -1.321928095 |
| Q9VEN3 | Q9VEN3_DROME FI20177p1 OS=Drosophila melanogaster OX=7227 GN=GckIII PE=1 SV=1 | 2 | 1 | 5 | 5 | 1 | 2.321928095 | -1.321928095 |
| Q9VHC8 | Q9VHC8_DROME LD31448p OS=Drosophila melanogaster OX=7227 GN=Dmel\CG8149 PE=1 SV=1 | 2 | 2 | 5 | 2 | 1 | 2.321928095 | -1.321928095 |
| Q9VNF3 | Q9VNF3_DROME RE01652p OS=Drosophila melanogaster OX=7227 GN=Dmel\CG12171 PE=1 SV=1 | 2 | 2 | 5 | 3 | 1 | 2.321928095 | -1.321928095 |
| Q9VNH5 | Q9VNH5_DROME GH04919p OS=Drosophila melanogaster OX=7227 GN=Dmel\CG2091 PE=1 SV=1 | 2 | 2 | 5 | 3 | 1 | 2.321928095 | -1.321928095 |
| Q9VRQ4 | Q9VRQ4_DROME LD47944p OS=Drosophila melanogaster OX=7227 GN=Dmel\CG5568 PE=2 SV=3 | 2 | 2 | 5 | 4 | 1 | 2.321928095 | -1.321928095 |
| Q9XYZ4 | Q9XYZ4_DROME Double-strand break repair protein OS=Drosophila melanogaster OX=7227 GN=mre11 PE=1 SV=1 | 2 | 1 | 5 | 5 | 1 | 2.321928095 | -1.321928095 |
| P25867 | UBCD1_DROME Ubiquitin-conjugating enzyme E2-17 kDa OS=Drosophila melanogaster OX=7227 GN=eff PE=2 SV=1 | 2 | 2 | 4 | 2 | 1 | 2 | -1 |
| P40417 | ERKA_DROME Mitogen-activated protein kinase ERK-A OS=Drosophila melanogaster OX=7227 GN=rl PE=1 SV=3 | 2 | 1 | 4 | 2 | 1 | 2 | -1 |
| P46461 | NSF1_DROME Vesicle-fusing ATPase 1 OS=Drosophila melanogaster OX=7227 GN=comt PE=2 SV=1 | 2 | 2 | 4 | 4 | 1 | 2 | -1 |
| Q9V4M2 | WECH_DROME Protein wech OS=Drosophila melanogaster OX=7227 GN=wech PE=1 SV=2 | 2 | 2 | 4 | 4 | 1 | 2 | -1 |
| Q9V535 | RBM8A_DROME RNA-binding protein 8A OS=Drosophila melanogaster OX=7227 GN=tsu PE=1 SV=1 | 2 | 1 | 4 | 3 | 1 | 2 | -1 |
| Q9VJ33 | NEDD8_DROME NEDD8 OS=Drosophila melanogaster OX=7227 GN=Nedd8 PE=1 SV=1 | 2 | 2 | 4 | 2 | 1 | 2 | -1 |
| Q9VL00 | OTUBL_DROME Ubiquitin thioesterase otubain-like OS=Drosophila melanogaster OX=7227 GN=CG4968 PE=2 SV=1 | 2 | 2 | 4 | 3 | 1 | 2 | -1 |
| Q9VMW7 | ITPA_DROME Inosine triphosphate pyrophosphatase OS=Drosophila melanogaster OX=7227 GN=CG8891 PE=2 SV=1 | 2 | 2 | 4 | 2 | 1 | 2 | -1 |
| Q9VU84 | DBNL_DROME Drebrin-like protein OS=Drosophila melanogaster OX=7227 GN=Abp1 PE=1 SV=1 | 2 | 2 | 4 | 4 | 1 | 2 | -1 |
| Q9VXB5 | RM22_DROME 39S ribosomal protein L22, mitochondrial OS=Drosophila melanogaster OX=7227 GN=mRpL22 PE=2 SV=2 | 2 | 2 | 4 | 3 | 1 | 2 | -1 |
| Q9VYG1 | PSMD5_DROME 26S proteasome non-ATPase regulatory subunit 5 OS=Drosophila melanogaster OX=7227 GN=CG12096 PE=1 SV=3 | 2 | 2 | 4 | 4 | 1 | 2 | -1 |
| Q9VZD5 | RT06_DROME Probable 28S ribosomal protein S6, mitochondrial OS=Drosophila melanogaster OX=7227 GN=mRpS6 PE=2 SV=1 | 2 | 2 | 4 | 3 | 1 | 2 | -1 |
| A0A0B4K699 | A0A0B4K699_DROME Mustard, isoform Y OS=Drosophila melanogaster OX=7227 GN=mtd PE=1 SV=1 | 2 | 2 | 4 | 3 | 1 | 2 | -1 |
| A1Z7Q8 | A1Z7Q8_DROME Alicorn, isoform A OS=Drosophila melanogaster OX=7227 GN=alc PE=1 SV=1 | 2 | 1 | 4 | 2 | 1 | 2 | -1 |
| O76863 | O76863_DROME EG:100G10.3 protein OS=Drosophila melanogaster OX=7227 GN=eIF2Bbeta PE=1 SV=1 | 2 | 2 | 4 | 4 | 1 | 2 | -1 |
| Q7K274 | Q7K274_DROME LD15586p OS=Drosophila melanogaster OX=7227 GN=l(2)k09913 PE=1 SV=1 | 2 | 1 | 4 | 2 | 1 | 2 | -1 |
| Q8I077 | Q8I077_DROME BcDNA.LD22910 OS=Drosophila melanogaster OX=7227 GN=Usp1 PE=1 SV=1 | 2 | 1 | 4 | 4 | 1 | 2 | -1 |
| Q8IPX7 | Q8IPX7_DROME RE44908p OS=Drosophila melanogaster OX=7227 GN=Rrp40 PE=1 SV=1 | 2 | 2 | 4 | 4 | 1 | 2 | -1 |
| Q8IQM5 | Q8IQM5_DROME Uncharacterized protein, isoform A OS=Drosophila melanogaster OX=7227 GN=Dmel\CG7656 PE=1 SV=1 | 2 | 1 | 4 | 3 | 1 | 2 | -1 |
| Q9VG38 | Q9VG38_DROME Suppressor of fused homolog OS=Drosophila melanogaster OX=7227 GN=Su(fu) PE=1 SV=1 | 2 | 2 | 4 | 3 | 1 | 2 | -1 |
| Q9VHQ0 | Q9VHQ0_DROME FI06460p OS=Drosophila melanogaster OX=7227 GN=mRpS18A PE=1 SV=3 | 2 | 2 | 4 | 3 | 1 | 2 | -1 |
| Q9VK44 | Q9VK44_DROME LD24832p OS=Drosophila melanogaster OX=7227 GN=CG 9934 PE=1 SV=1 | 2 | 2 | 4 | 4 | 1 | 2 | -1 |
| Q9VU38 | Q9VU38_DROME Adenosine kinase, isoform B OS=Drosophila melanogaster OX=7227 GN=AdenoK PE=1 SV=1 | 2 | 2 | 4 | 3 | 1 | 2 | -1 |
| Q9VUH6 | Q9VUH6_DROME Myopic OS=Drosophila melanogaster OX=7227 GN=mop PE=1 SV=2 | 2 | 2 | 4 | 3 | 1 | 2 | -1 |
| Q9VX69 | Q9VX69_DROME FI01450p OS=Drosophila melanogaster OX=7227 GN=Dmel\CG5162 PE=1 SV=1 | 2 | 2 | 4 | 4 | 1 | 2 | -1 |
| Q9VYT5 | Q9VYT5_DROME LD06533p OS=Drosophila melanogaster OX=7227 GN=Dmel\CG10347 PE=1 SV=1 | 2 | 1 | 4 | 3 | 1 | 2 | -1 |
| Q9W425 | Q9W425_DROME Rabconnectin-3A OS=Drosophila melanogaster OX=7227 GN=Rbcn-3A PE=1 SV=3 | 2 | 2 | 4 | 4 | 1 | 2 | -1 |
| Q08012 | DRK_DROME Protein enhancer of sevenless 2B OS=Drosophila melanogaster OX=7227 GN=drk PE=1 SV=1 | 2 | 2 | 3 | 3 | 1 | 1.584962501 | -0.584962501 |
| Q24324 | DSOR1_DROME Dual specificity mitogen-activated protein kinase kinase dSOR1 OS=Drosophila melanogaster OX=7227 GN=Dsor1 PE=1 SV=2 | 2 | 2 | 3 | 3 | 1 | 1.584962501 | -0.584962501 |
| Q94535 | U2AF1_DROME Splicing factor U2af 38 kDa subunit OS=Drosophila melanogaster OX=7227 GN=U2af38 PE=1 SV=2 | 2 | 2 | 3 | 3 | 1 | 1.584962501 | -0.584962501 |
| Q9I7H9 | OGFD1_DROME Prolyl 3-hydroxylase sudestada1 OS=Drosophila melanogaster OX=7227 GN=sud1 PE=2 SV=1 | 2 | 2 | 3 | 3 | 1 | 1.584962501 | -0.584962501 |
| Q9VSF4 | DHYS_DROME Probable deoxyhypusine synthase OS=Drosophila melanogaster OX=7227 GN=CG8005 PE=2 SV=2 | 2 | 2 | 3 | 3 | 1 | 1.584962501 | -0.584962501 |
| Q9W197 | 5NT3B_DROME 7-methylguanosine phosphate-specific 5'-nucleotidase OS=Drosophila melanogaster OX=7227 GN=cN-IIIB PE=1 SV=1 | 2 | 2 | 3 | 3 | 1 | 1.584962501 | -0.584962501 |
| A1Z7K2 | A1Z7K2_DROME FI03455p OS=Drosophila melanogaster OX=7227 GN=CG30349-RA PE=1 SV=1 | 2 | 1 | 3 | 3 | 1 | 1.584962501 | -0.584962501 |
| A8Y589 | A8Y589_DROME Apoptosis-linked gene-2 OS=Drosophila melanogaster OX=7227 GN=Alg-2 PE=1 SV=2 | 2 | 2 | 3 | 2 | 1 | 1.584962501 | -0.584962501 |
| Q7K0X3 | Q7K0X3_DROME Ctr9, isoform A OS=Drosophila melanogaster OX=7227 GN=Ctr9 PE=1 SV=1 | 2 | 2 | 3 | 3 | 1 | 1.584962501 | -0.584962501 |
| Q8T3V6 | Q8T3V6_DROME AT29239p OS=Drosophila melanogaster OX=7227 GN=mRpL47 PE=1 SV=1 | 2 | 2 | 3 | 3 | 1 | 1.584962501 | -0.584962501 |
| Q9VG51 | Q9VG51_DROME RE52028p OS=Drosophila melanogaster OX=7227 GN=Snx3 PE=1 SV=1 | 2 | 2 | 3 | 3 | 1 | 1.584962501 | -0.584962501 |
| Q9VHT5 | Q9VHT5_DROME LD31571p OS=Drosophila melanogaster OX=7227 GN=mRpL1 PE=1 SV=2 | 2 | 2 | 3 | 3 | 1 | 1.584962501 | -0.584962501 |
| Q9VHX4 | Q9VHX4_DROME LD24679p OS=Drosophila melanogaster OX=7227 GN=Dmel\CG2767 PE=1 SV=1 | 2 | 2 | 3 | 2 | 1 | 1.584962501 | -0.584962501 |
| Q9VUZ8 | Q9VUZ8_DROME GEO07442p1 OS=Drosophila melanogaster OX=7227 GN=PDCD-5 PE=1 SV=1 | 2 | 2 | 3 | 2 | 1 | 1.584962501 | -0.584962501 |
| P18930 | NU3M_DROME NADH-ubiquinone oxidoreductase chain 3 OS=Drosophila melanogaster OX=7227 GN=mt:ND3 PE=3 SV=1 | 2 | 1 | 1 | 1 | 1 | 0 | 1 |
| P40797 | PNUT_DROME Protein peanut OS=Drosophila melanogaster OX=7227 GN=pnut PE=1 SV=2 | 2 | 2 | 1 | 1 | 1 | 0 | 1 |
| P48609 | CDK5_DROME Cyclin-dependent kinase 5 homolog OS=Drosophila melanogaster OX=7227 GN=Cdk5 PE=1 SV=2 | 2 | 2 | 2 | 2 | 1 | 1 | 0 |

| UniProt ID | Protein description | # PSMs MVB spectra | # of unique peptides MVB | # of PSMs lysate | # of unique peptides lysate | log2(PSM MVB) | log2(PSM lysate) | FC (log2 of MVB PSM-log2 of lysate PSM) |
|---|---|---|---|---|---|---|---|---|
| Q7JZM8 | RM41_DROME 39S ribosomal protein L41, mitochondrial OS=Drosophila melanogaster OX=7227 GN=mRpL41 PE=2 SV=1 | 2 | 2 | 2 | 2 | 1 | 1 | 0 |
| Q9W451 | Q9W451_DROME Female sterile (1) M3 OS=Drosophila melanogaster OX=7227 GN=fs(1)M3 PE=4 SV=2 | 2 | 2 | 3 | 3 | 1 | 1.584962501 | -0.584962501 |
| Q9VAJ4 | OB99A_DROME General odorant-binding protein 99a OS=Drosophila melanogaster OX=7227 GN=Obp99a PE=2 SV=2 | 2 | 2 | 2 | 2 | 1 | 1 | 0 |
| Q9VJ38 | RM13_DROME 39S ribosomal protein L13, mitochondrial OS=Drosophila melanogaster OX=7227 GN=mRpL13 PE=2 SV=2 | 2 | 2 | 2 | 2 | 1 | 1 | 0 |
| Q9VLJ9 | RM51_DROME 39S ribosomal protein L51, mitochondrial OS=Drosophila melanogaster OX=7227 GN=mRpL51 PE=2 SV=2 | 2 | 2 | 2 | 2 | 1 | 1 | 0 |
| A0A0C4DHE7 | A0A0C4DHE7_DROME Glutamate dehydrogenase OS=Drosophila melanogaster OX=7227 GN=Gdh PE=1 SV=1 | 2 | 1 | 2 | 1 | 1 | 1 | 0 |
| A1Z7B8 | A1Z7B8_DROME GEO08456p1 OS=Drosophila melanogaster OX=7227 GN=CG30373 PE=2 SV=1 | 2 | 2 | 2 | 2 | 1 | 1 | 0 |
| A1Z7H7 | A1Z7H7_DROME CG8586, isoform A OS=Drosophila melanogaster OX=7227 GN=CG8586-RA PE=1 SV=1 | 2 | 2 | 1 | 1 | 1 | 0 | 1 |
| A1Z814 | A1Z814_DROME Oxysterol-binding protein OS=Drosophila melanogaster OX=7227 GN=CG1513-RA PE=1 SV=2 | 2 | 2 | 1 | 1 | 1 | 0 | 1 |
| A1ZB29 | A1ZB29_DROME CG5757, isoform A OS=Drosophila melanogaster OX=7227 GN=CG5757 PE=1 SV=1 | 2 | 2 | 1 | 1 | 1 | 0 | 1 |
| A8JV07 | A8JV07_DROME Uncharacterized protein, isoform B OS=Drosophila melanogaster OX=7227 GN=Dmel\CG9915 PE=1 SV=1 | 2 | 2 | 1 | 1 | 1 | 0 | 1 |
| E2QCZ3 | E2QCZ3_DROME LP22035p OS=Drosophila melanogaster OX=7227 GN=CG7705-RA PE=2 SV=1 | 2 | 2 | 1 | 1 | 1 | 0 | 1 |
| F0JAQ9 | F0JAQ9_DROME MIP27169p OS=Drosophila melanogaster OX=7227 GN=Wbp2 PE=1 SV=1 | 2 | 1 | 2 | 1 | 1 | 1 | 0 |
| M9PBU2 | M9PBU2_DROME Rac2, isoform B OS=Drosophila melanogaster OX=7227 GN=Rac2 PE=4 SV=1 | 2 | 2 | 1 | 1 | 1 | 0 | 1 |
| M9PEI9 | M9PEI9_DROME Probable methylthioribulose-1-phosphate dehydratase OS=Drosophila melanogaster OX=7227 GN=CG11134 PE=3 SV=1 | 2 | 2 | 2 | 2 | 1 | 1 | 0 |
| M9PEJ5 | M9PEJ5_DROME Retinal degeneration B, isoform J OS=Drosophila melanogaster OX=7227 GN=rdgB PE=1 SV=1 | 2 | 1 | 1 | 1 | 1 | 0 | 1 |
| Q0E8V7 | Q0E8V7_DROME GEO11073p1 OS=Drosophila melanogaster OX=7227 GN=Dmel\CG34132 PE=1 SV=1 | 2 | 2 | 1 | 1 | 1 | 0 | 1 |
| Q7K4M9 | Q7K4M9_DROME LD31969p OS=Drosophila melanogaster OX=7227 GN=CG15118 PE=1 SV=1 | 2 | 2 | 2 | 2 | 1 | 1 | 0 |
| Q7KUT5 | Q7KUT5_DROME HL01254p OS=Drosophila melanogaster OX=7227 GN=Ccdc58 PE=1 SV=1 | 2 | 1 | 2 | 1 | 1 | 1 | 0 |
| Q8IQF8 | Q8IQF8_DROME AT18092p OS=Drosophila melanogaster OX=7227 GN=34F4T PE=1 SV=1 | 2 | 1 | 2 | 1 | 1 | 1 | 0 |
| Q8IRJ9 | Q8IRJ9_DROME Glycerol kinase 1, isoform B OS=Drosophila melanogaster OX=7227 GN=Gk1 PE=1 SV=1 | 2 | 2 | 1 | 1 | 1 | 0 | 1 |
| Q9VJJ1 | Q9VJJ1_DROME COX assembly mitochondrial protein OS=Drosophila melanogaster OX=7227 GN=CG17996 PE=3 SV=1 | 2 | 1 | 2 | 1 | 1 | 1 | 0 |
| Q9VK46 | Q9VK46_DROME Protein YIPF OS=Drosophila melanogaster OX=7227 GN=Dmel\CG12404 PE=1 SV=1 | 2 | 1 | 2 | 1 | 1 | 1 | 0 |
| Q9VSW4 | Q9VSW4_DROME FI06490p OS=Drosophila melanogaster OX=7227 GN=mRRF1 PE=1 SV=1 | 2 | 2 | 2 | 2 | 1 | 1 | 0 |
| Q9VV89 | Q9VV89_DROME Coat protein (Coatomer) zeta, isoform A OS=Drosophila melanogaster OX=7227 GN=zetaCOP PE=1 SV=2 | 2 | 1 | 2 | 2 | 1 | 1 | 0 |
| Q9VIQ5 | Q9VIQ5_DROME RH02620p OS=Drosophila melanogaster OX=7227 GN=Dmel\CG10680 PE=2 SV=1 | 2 | 1 | 1 | 1 | 1 | 0 | 1 |
| Q9VJ10 | Q9VJ10_DROME Lethal (2) 37Bb OS=Drosophila melanogaster OX=7227 GN=l(2)37Bb PE=1 SV=1 | 2 | 2 | 1 | 1 | 1 | 0 | 1 |
| Q9VVA7 | Q9VVA7_DROME Tumor suppressor protein 101 OS=Drosophila melanogaster OX=7227 GN=TSG101 PE=1 SV=2 | 2 | 2 | 1 | 1 | 1 | 0 | 1 |
| Q9VWD5 | Q9VWD5_DROME LD35087p OS=Drosophila melanogaster OX=7227 GN=Dmel\CG14232 PE=1 SV=1 | 2 | 2 | 1 | 1 | 1 | 0 | 1 |
| Q9W350 | Q9W350_DROME C11.1, isoform A OS=Drosophila melanogaster OX=7227 GN=c11.1 PE=1 SV=2 | 2 | 2 | 1 | 1 | 1 | 0 | 1 |
| Q9W389 | Q9W389_DROME ADP-ribosylation factor related protein 1 OS=Drosophila melanogaster OX=7227 GN=Arfrp1 PE=1 SV=2 | 2 | 2 | 1 | 1 | 1 | 0 | 1 |
| Q9W5C2 | Q9W5C2_DROME Translocating chain-associated membrane protein OS=Drosophila melanogaster OX=7227 GN=TRAM PE=1 SV=1 | 2 | 1 | 1 | 1 | 1 | 0 | 1 |
| ALBU_BOVIN | (Common contaminant protein) | 2 | 2 | 1 | 1 | 1 | 0 | 1 |
| B7YZT2 | FIS1_DROME Mitochondrial fission 1 protein OS=Drosophila melanogaster OX=7227 GN=Fis1 PE=2 SV=1 | 2 | 2 | 1 | 1 | 1 | 0 | 1 |
| M9PD06 | UBP32_DROME Ubiquitin carboxyl-terminal hydrolase 32 OS=Drosophila melanogaster OX=7227 GN=Usp32 PE=3 SV=1 | 2 | 2 | 1 | 1 | 1 | 0 | 1 |
| O44252 | ROST_DROME Protein rolling stone OS=Drosophila melanogaster OX=7227 GN=rost PE=2 SV=2 | 2 | 2 | 1 | 1 | 1 | 0 | 1 |
| O61267 | LOK_DROME Ovarian-specific serine/threonine-protein kinase Lok OS=Drosophila melanogaster OX=7227 GN=lok PE=2 SV=1 | 2 | 2 | 1 | 1 | 1 | 0 | 1 |
| P0DKM0 | COA3_DROME Cytochrome c oxidase assembly factor 3, mitochondrial OS=Drosophila melanogaster OX=7227 GN=Ccdc56 PE=3 SV=1 | 2 | 2 | 1 | 1 | 1 | 0 | 1 |
| P18537 | FRIZ_DROME Frizzled OS=Drosophila melanogaster OX=7227 GN=fz PE=1 SV=1 | 2 | 2 | 1 | 1 | 1 | 0 | 1 |
| P25722 | HS2ST_DROME Heparin sulfate O-sulfotransferase OS=Drosophila melanogaster OX=7227 GN=Hs2st PE=2 SV=2 | 2 | 2 | 1 | 1 | 1 | 0 | 1 |
| P27716 | INX1_DROME Innexin inx1 OS=Drosophila melanogaster OX=7227 GN=ogre PE=1 SV=1 | 2 | 2 | 1 | 1 | 1 | 0 | 1 |
| P28466 | WNT5_DROME Protein Wnt-5 OS=Drosophila melanogaster OX=7227 GN=Wnt5 PE=1 SV=2 | 2 | 2 | 1 | 1 | 1 | 0 | 1 |
| P47938 | THIO1_DROME Thioredoxin-1 OS=Drosophila melanogaster OX=7227 GN=dhd PE=1 SV=1 | 2 | 1 | 1 | 1 | 1 | 0 | 1 |
| P82711 | C6A19_DROME Probable cytochrome P450 6a19 OS=Drosophila melanogaster OX=7227 GN=Cyp6a19 PE=3 SV=1 | 2 | 2 | 1 | 1 | 1 | 0 | 1 |
| P83097 | WSCK_DROME Putative tyrosine-protein kinase Wsck OS=Drosophila melanogaster OX=7227 GN=Wsck PE=2 SV=2 | 2 | 2 | 1 | 1 | 1 | 0 | 1 |
| Q00449 | MDR49_DROME Multidrug resistance protein homolog 49 OS=Drosophila melanogaster OX=7227 GN=Mdr49 PE=2 SV=2 | 2 | 2 | 1 | 1 | 1 | 0 | 1 |
| Q5BI42 | CAHD1_DROME VWFA and cache domain-containing protein CG16868 OS=Drosophila melanogaster OX=7227 GN=CG16868 PE=2 SV=1 | 2 | 2 | 1 | 1 | 1 | 0 | 1 |
| Q7KNA0 | DYM_DROME Dymeclin OS=Drosophila melanogaster OX=7227 GN=CG8230 PE=1 SV=1 | 2 | 2 | 1 | 1 | 1 | 0 | 1 |
| Q7KSC4 | MPC1_DROME Mitochondrial pyruvate carrier 1 OS=Drosophila melanogaster OX=7227 GN=Mpc1 PE=3 SV=1 | 2 | 1 | 1 | 1 | 1 | 0 | 1 |
| Q8MLZ7 | IDGF3_DROME Chitinase-like protein Idgf3 OS=Drosophila melanogaster OX=7227 GN=Idgf3 PE=1 SV=3 | 2 | 2 | 1 | 1 | 1 | 0 | 1 |
| Q8MM24 | IDGF1_DROME Chitinase-like protein Idgf1 OS=Drosophila melanogaster OX=7227 GN=Idgf1 PE=1 SV=2 | 2 | 2 | 1 | 1 | 1 | 0 | 1 |
| Q9U4H5 | AMX_DROME TM2 domain-containing protein almondex OS=Drosophila melanogaster OX=7227 GN=amx PE=2 SV=1 | 2 | 2 | 1 | 1 | 1 | 0 | 1 |
| Q9V3Z2 | SER7_DROME Serine protease 7 OS=Drosophila melanogaster OX=7227 GN=Sp7 PE=1 SV=1 | 2 | 1 | 1 | 1 | 1 | 0 | 1 |
| Q9V4I1 | CP9B2_DROME Cytochrome P450 9b2 OS=Drosophila melanogaster OX=7227 GN=Cyp9b2 PE=2 SV=1 | 2 | 2 | 1 | 1 | 1 | 0 | 1 |
| Q9V4T3 | C4AD1_DROME Probable cytochrome P450 4ad1 OS=Drosophila melanogaster OX=7227 GN=Cyp4ad1 PE=2 SV=1 | 2 | 2 | 1 | 1 | 1 | 0 | 1 |
| Q9V774 | C6A21_DROME Probable cytochrome P450 6a21 OS=Drosophila melanogaster OX=7227 GN=Cyp6a21 PE=2 SV=1 | 2 | 2 | 1 | 1 | 1 | 0 | 1 |
| Q9V7W1 | PIGV_DROME GPI mannosyltransferase 2 OS=Drosophila melanogaster OX=7227 GN=PIG-V PE=1 SV=1 | 2 | 1 | 1 | 1 | 1 | 0 | 1 |
| Q9V9A8 | PHTF_DROME Putative homeodomain transcription factor OS=Drosophila melanogaster OX=7227 GN=phtf PE=1 SV=2 | 2 | 2 | 1 | 1 | 1 | 0 | 1 |
| Q9VCK1 | GILT2_DROME GILT-like protein 2 OS=Drosophila melanogaster OX=7227 GN=GILT2 PE=2 SV=1 | 2 | 1 | 1 | 1 | 1 | 0 | 1 |
| Q9VF08 | TIM16_DROME Mitochondrial import inner membrane translocase subunit Tim16 OS=Drosophila melanogaster OX=7227 GN=blp PE=2 SV=1 | 2 | 2 | 1 | 1 | 1 | 0 | 1 |
| Q9VFP1 | CP6D5_DROME Probable cytochrome P450 6d5 OS=Drosophila melanogaster OX=7227 GN=Cyp6d5 PE=2 SV=1 | 2 | 1 | 1 | 1 | 1 | 0 | 1 |
| Q9VH19 | LMLN_DROME Leishmanolysin-like peptidase OS=Drosophila melanogaster OX=7227 GN=Invadolysin PE=2 SV=2 | 2 | 2 | 1 | 1 | 1 | 0 | 1 |
| Q9VH78 | ALG12_DROME Probable Dol-P-Man:Man(7)GlcNAc(2)-PP-Dol alpha-1,6-mannosyltransferase OS=Drosophila melanogaster OX=7227 GN=CG8412 PE=2 SV=1 | 2 | 2 | 1 | 1 | 1 | 0 | 1 |
| Q9VMT6 | C28D2_DROME Probable cytochrome P450 28d2 OS=Drosophila melanogaster OX=7227 GN=Cyp28d2 PE=3 SV=1 | 2 | 2 | 1 | 1 | 1 | 0 | 1 |
| Q9VR50 | S35E1_DROME Solute carrier family 35 member E1 homolog OS=Drosophila melanogaster OX=7227 GN=CG14621 PE=2 SV=1 | 2 | 1 | 1 | 1 | 1 | 0 | 1 |
| Q9VRJ8 | ASTER_DROME Protein Asterix OS=Drosophila melanogaster OX=7227 GN=CG10674 PE=3 SV=1 | 2 | 1 | 1 | 1 | 1 | 0 | 1 |
| Q9VWE9 | S61G2_DROME Protein transport protein Sec61 gamma-2 subunit OS=Drosophila melanogaster OX=7227 GN=Sec61gamma PE=3 SV=1 | 2 | 1 | 1 | 1 | 1 | 0 | 1 |
| Q9VWH8 | KISH_DROME Protein kish OS=Drosophila melanogaster OX=7227 GN=ksh PE=3 SV=2 | 2 | 1 | 1 | 1 | 1 | 0 | 1 |
| Q9VYD7 | TIM9_DROME Mitochondrial import inner membrane translocase subunit Tim9 OS=Drosophila melanogaster OX=7227 GN=Tim9a PE=3 SV=1 | 2 | 2 | 1 | 1 | 1 | 0 | 1 |
| Q9VYI3 | RM49_DROME Probable 39S ribosomal protein L49, mitochondrial OS=Drosophila melanogaster OX=7227 GN=mRpL49 PE=3 SV=1 | 2 | 2 | 1 | 1 | 1 | 0 | 1 |
| Q9VZU2 | HHAT_DROME Protein-cysteine N-palmitoyltransferase Rasp OS=Drosophila melanogaster OX=7227 GN=rasp PE=2 SV=1 | 2 | 2 | 1 | 1 | 1 | 0 | 1 |
| Q9W092 | CHIT2_DROME Probable chitinase 2 OS=Drosophila melanogaster OX=7227 GN=Cht2 PE=1 SV=1 | 2 | 2 | 1 | 1 | 1 | 0 | 1 |
| Q9Y0V3 | T10B_DROME Mitochondrial import inner membrane translocase subunit Tim10B OS=Drosophila melanogaster OX=7227 GN=Tim9b PE=3 SV=1 | 2 | 2 | 1 | 1 | 1 | 0 | 1 |
| A0A0B4JD45 | A0A0B4JD45_DROME Tafazzin, isoform D OS=Drosophila melanogaster OX=7227 GN=Taz PE=4 SV=1 | 2 | 2 | 1 | 1 | 1 | 0 | 1 |
| A0A0B4K6Q5 | A0A0B4K6Q5_DROME Phosphatidylinositol glycan anchor biosynthesis class L, isoform C OS=Drosophila melanogaster OX=7227 GN=PIG-L PE=4 SV=1 | 2 | 1 | 1 | 1 | 1 | 0 | 1 |
| A0A0B4K7D8 | A0A0B4K7D8_DROME CG6805, isoform B OS=Drosophila melanogaster OX=7227 GN=CG6805 PE=1 SV=1 | 2 | 2 | 1 | 1 | 1 | 0 | 1 |
| A0A0B4K833 | A0A0B4K833_DROME Calcineurin B2, isoform B OS=Drosophila melanogaster OX=7227 GN=CanB2 PE=4 SV=1 | 2 | 2 | 1 | 1 | 1 | 0 | 1 |
| A0A0B4KEI5 | A0A0B4KEI5_DROME Ars2, isoform E OS=Drosophila melanogaster OX=7227 GN=Ars2 PE=1 SV=1 | 2 | 2 | 1 | 1 | 1 | 0 | 1 |
| A0A0B4KEX0 | A0A0B4KEX0_DROME Protein clueless OS=Drosophila melanogaster OX=7227 GN=clu PE=1 SV=1 | 2 | 2 | 1 | 1 | 1 | 0 | 1 |
| A0A0B4KG59 | A0A0B4KG59_DROME Serine/threonine-protein kinase receptor OS=Drosophila melanogaster OX=7227 GN=put PE=1 SV=1 | 2 | 2 | 1 | 1 | 1 | 0 | 1 |
| A0A0B4KHF5 | A0A0B4KHF5_DROME Uncharacterized protein, isoform O OS=Drosophila melanogaster OX=7227 GN=CG2246 PE=1 SV=1 | 2 | 2 | 1 | 1 | 1 | 0 | 1 |

| UniProt ID | Protein description | # PSMs MVB spectra | # of unique peptides MVB | # of PSMs lysate | # of unique peptides lysate | log2(PSM MVB) | log2(PSM lysate) | FC (log2 of MVB PSM-log2 of lysate PSM) |
|---|---|---|---|---|---|---|---|---|
| A0A0B4KHK0 | A0A0B4KHK0_DROME Dihydroxyacetone phosphate acyltransferase, isoform C OS=Drosophila melanogaster OX=7227 GN=Dhap-at PE=3 SV=1 | 2 | 2 | 1 | 1 | 1 | 0 | 1 |
| A0A0B4LF97 | A0A0B4LF97_DROME Bedraggled, isoform D OS=Drosophila melanogaster OX=7227 GN=bdg PE=4 SV=1 | 2 | 2 | 1 | 1 | 1 | 0 | 1 |
| A0A0B4LG64 | A0A0B4LG64_DROME Metallophosphoesterase, isoform D OS=Drosophila melanogaster OX=7227 GN=Mppe PE=4 SV=1 | 2 | 2 | 1 | 1 | 1 | 0 | 1 |
| A0A0B4LGP3 | A0A0B4LGP3_DROME Epithelial membrane protein, isoform E OS=Drosophila melanogaster OX=7227 GN=emp PE=3 SV=1 | 2 | 2 | 1 | 1 | 1 | 0 | 1 |
| A0A0B4LGV1 | A0A0B4LGV1_DROME ER membrane protein complex 1, isoform C OS=Drosophila melanogaster OX=7227 GN=EMC1 PE=1 SV=1 | 2 | 1 | 1 | 1 | 1 | 0 | 1 |
| A0A0C4DHN2 | A0A0C4DHN2_DROME Uncharacterized protein, isoform B OS=Drosophila melanogaster OX=7227 GN=CG41378 PE=4 SV=1 | 2 | 2 | 1 | 1 | 1 | 0 | 1 |
| A0A0S0WGS4 | A0A0S0WGS4_DROME Uncharacterized protein, isoform E OS=Drosophila melanogaster OX=7227 GN=CG34310 PE=4 SV=1 | 2 | 2 | 1 | 1 | 1 | 0 | 1 |
| A1Z741 | A1Z741_DROME Uncharacterized protein, isoform B OS=Drosophila melanogaster OX=7227 GN=CG18812 PE=4 SV=1 | 2 | 2 | 1 | 1 | 1 | 0 | 1 |
| A1Z7S4 | A1Z7S4_DROME CG8026, isoform D OS=Drosophila melanogaster OX=7227 GN=CG8026 PE=3 SV=2 | 2 | 1 | 1 | 1 | 1 | 0 | 1 |
| A1Z823 | A1Z823_DROME PRA1 family protein OS=Drosophila melanogaster OX=7227 GN=CG1418-RA PE=2 SV=1 | 2 | 2 | 1 | 1 | 1 | 0 | 1 |
| A1Z9L5 | A1Z9L5_DROME FI21814p1 OS=Drosophila melanogaster OX=7227 GN=CG13016-RA PE=2 SV=1 | 2 | 2 | 1 | 1 | 1 | 0 | 1 |
| B7Z0C9 | B7Z0C9_DROME GH15104p1 OS=Drosophila melanogaster OX=7227 GN=Dmel\CG42307 PE=2 SV=1 | 2 | 2 | 1 | 1 | 1 | 0 | 1 |
| B7Z0U6 | B7Z0U6_DROME Bestrophin 1, isoform B OS=Drosophila melanogaster OX=7227 GN=Best1 PE=1 SV=1 | 2 | 2 | 1 | 1 | 1 | 0 | 1 |
| B7Z107 | B7Z107_DROME GH09380p1 OS=Drosophila melanogaster OX=7227 GN=Dmel\CG42376 PE=2 SV=1 | 2 | 2 | 1 | 1 | 1 | 0 | 1 |
| C6SV36 | C6SV36_DROME IP19903p OS=Drosophila melanogaster OX=7227 GN=CG3884-RB PE=1 SV=1 | 2 | 2 | 1 | 1 | 1 | 0 | 1 |
| E1JGN0 | E1JGN0_DROME Non-specific serine/threonine protein kinase OS=Drosophila melanogaster OX=7227 GN=par-1 PE=1 SV=1 | 2 | 2 | 1 | 1 | 1 | 0 | 1 |
| E1JI98 | E1JI98_DROME GEO02452p1 OS=Drosophila melanogaster OX=7227 GN=Dmel\CG42455 PE=2 SV=1 | 2 | 1 | 1 | 1 | 1 | 0 | 1 |
| M9ND00 | M9ND00_DROME Dynamin associated protein 160, isoform D OS=Drosophila melanogaster OX=7227 GN=Dap160 PE=1 SV=1 | 2 | 2 | 1 | 1 | 1 | 0 | 1 |
| M9NE76 | M9NE76_DROME Galactosylgalactosylxylosylprotein 3-beta-glucuronosyltransferase OS=Drosophila melanogaster OX=7227 GN=GlcAT-P PE=3 SV=1 | 2 | 2 | 1 | 1 | 1 | 0 | 1 |
| M9PB77 | M9PB77_DROME Semaphorin 1a, isoform I OS=Drosophila melanogaster OX=7227 GN=Sema1a PE=1 SV=1 | 2 | 2 | 1 | 1 | 1 | 0 | 1 |
| M9PBN2 | M9PBN2_DROME Neural lazarillo, isoform B OS=Drosophila melanogaster OX=7227 GN=NLaz PE=1 SV=1 | 2 | 2 | 1 | 1 | 1 | 0 | 1 |
| M9PCH6 | M9PCH6_DROME Omega, isoform H OS=Drosophila melanogaster OX=7227 GN=ome PE=1 SV=1 | 2 | 2 | 1 | 1 | 1 | 0 | 1 |
| M9PD26 | M9PD26_DROME Cytochrome b5-related, isoform B OS=Drosophila melanogaster OX=7227 GN=Cyt-b5-r PE=4 SV=1 | 2 | 2 | 1 | 1 | 1 | 0 | 1 |
| M9PD30 | M9PD30_DROME Uncharacterized protein, isoform G OS=Drosophila melanogaster OX=7227 GN=CG5850 PE=1 SV=1 | 2 | 1 | 1 | 1 | 1 | 0 | 1 |
| M9PDW0 | M9PDW0_DROME Neuronal synaptobrevin, isoform K OS=Drosophila melanogaster OX=7227 GN=nSyb PE=1 SV=1 | 2 | 2 | 1 | 1 | 1 | 0 | 1 |
| M9PE35 | M9PE35_DROME RabX6, isoform B OS=Drosophila melanogaster OX=7227 GN=RabX6 PE=4 SV=1 | 2 | 1 | 1 | 1 | 1 | 0 | 1 |
| M9PEG6 | M9PEG6_DROME Zinc transporter 63C, isoform F OS=Drosophila melanogaster OX=7227 GN=ZnT63C PE=1 SV=1 | 2 | 2 | 1 | 1 | 1 | 0 | 1 |
| O76899 | O76899_DROME EG:80H7.2 protein OS=Drosophila melanogaster OX=7227 GN=pck PE=4 SV=1 | 2 | 1 | 1 | 1 | 1 | 0 | 1 |
| O97479 | O97479_DROME FI05212p OS=Drosophila melanogaster OX=7227 GN=Sodh-1 PE=1 SV=1 | 2 | 2 | 1 | 1 | 1 | 0 | 1 |
| Q059A4 | Q059A4_DROME CG4627 OS=Drosophila melanogaster OX=7227 GN=CG4627 PE=2 SV=1 | 2 | 2 | 1 | 1 | 1 | 0 | 1 |
| Q0E8W1 | Q0E8W1_DROME Fatty acyl-CoA reductase OS=Drosophila melanogaster OX=7227 GN=CG30427 PE=3 SV=1 | 2 | 1 | 1 | 1 | 1 | 0 | 1 |
| Q4ABE8 | Q4ABE8_DROME Cadherin-N, isoform K OS=Drosophila melanogaster OX=7227 GN=CadN PE=1 SV=1 | 2 | 2 | 1 | 1 | 1 | 0 | 1 |
| Q4V429 | Q4V429_DROME GEO10184p1 OS=Drosophila melanogaster OX=7227 GN=CG13018 PE=2 SV=1 | 2 | 2 | 1 | 1 | 1 | 0 | 1 |
| Q5LJX8 | Q5LJX8_DROME Zydeco, isoform F OS=Drosophila melanogaster OX=7227 GN=zyd PE=3 SV=3 | 2 | 2 | 1 | 1 | 1 | 0 | 1 |
| Q6IGW6 | Q6IGW6_DROME GEO13367p1 OS=Drosophila melanogaster OX=7227 GN=CG34200 PE=2 SV=1 | 2 | 1 | 1 | 1 | 1 | 0 | 1 |
| Q7JQL2 | Q7JQL2_DROME LD20463p OS=Drosophila melanogaster OX=7227 GN=Sans PE=2 SV=1 | 2 | 2 | 1 | 1 | 1 | 0 | 1 |
| Q7K0A0 | Q7K0A0_DROME Mitochondrial ribosomal protein L42, isoform A OS=Drosophila melanogaster OX=7227 GN=mRpL42 PE=1 SV=1 | 2 | 2 | 1 | 1 | 1 | 0 | 1 |
| Q7K1I4 | Q7K1I4_DROME Tetraspanin OS=Drosophila melanogaster OX=7227 GN=Tsp42Ea PE=1 SV=1 | 2 | 2 | 1 | 1 | 1 | 0 | 1 |
| Q7K2X5 | Q7K2X5_DROME GH01839p OS=Drosophila melanogaster OX=7227 GN=CG14762 PE=1 SV=1 | 2 | 2 | 1 | 1 | 1 | 0 | 1 |
| Q7KSB7 | Q7KSB7_DROME Apoptotic signal-regulating kinase 1, isoform B OS=Drosophila melanogaster OX=7227 GN=Ask1 PE=1 SV=1 | 2 | 2 | 1 | 1 | 1 | 0 | 1 |
| Q7KTC7 | Q7KTC7_DROME Multidrug-Resistance like protein 1, isoform C OS=Drosophila melanogaster OX=7227 GN=MRP PE=1 SV=1 | 2 | 2 | 1 | 1 | 1 | 0 | 1 |
| Q7KUF9 | Q7KUF9_DROME Sugar baby, isoform F OS=Drosophila melanogaster OX=7227 GN=Sugb PE=1 SV=3 | 2 | 2 | 1 | 1 | 1 | 0 | 1 |
| Q7KUK8 | Q7KUK8_DROME GEO05070p1 OS=Drosophila melanogaster OX=7227 GN=Dmel\CG9628 PE=1 SV=1 | 2 | 1 | 1 | 1 | 1 | 0 | 1 |
| Q7PLE8 | Q7PLE8_DROME Rab21, isoform A OS=Drosophila melanogaster OX=7227 GN=Rab21 PE=4 SV=1 | 2 | 2 | 1 | 1 | 1 | 0 | 1 |
| Q8IGQ3 | Q8IGQ3_DROME RE48687p OS=Drosophila melanogaster OX=7227 GN=Dmel\CG13284 PE=2 SV=1 | 2 | 2 | 1 | 1 | 1 | 0 | 1 |
| Q8IGV3 | Q8IGV3_DROME RE23632p OS=Drosophila melanogaster OX=7227 GN=Dmel\CG12746 PE=2 SV=1 | 2 | 2 | 1 | 1 | 1 | 0 | 1 |
| Q8IMC1 | Q8IMC1_DROME Transporter OS=Drosophila melanogaster OX=7227 GN=Gat PE=1 SV=2 | 2 | 2 | 1 | 1 | 1 | 0 | 1 |
| Q8IQQ0 | Q8IQQ0_DROME Neural conserved at 73EF, isoform F OS=Drosophila melanogaster OX=7227 GN=Nc73EF PE=1 SV=1 | 2 | 2 | 1 | 1 | 1 | 0 | 1 |
| Q8IR19 | Q8IR19_DROME Uncharacterized protein, isoform A OS=Drosophila melanogaster OX=7227 GN=CG32576 PE=1 SV=1 | 2 | 1 | 1 | 1 | 1 | 0 | 1 |
| Q8IRD0 | Q8IRD0_DROME RH17559p OS=Drosophila melanogaster OX=7227 GN=Dmel\CG32267 PE=1 SV=1 | 2 | 1 | 1 | 1 | 1 | 0 | 1 |
| Q8MLN4 | Q8MLN4_DROME CG9380, isoform D OS=Drosophila melanogaster OX=7227 GN=CG9380 PE=1 SV=2 | 2 | 2 | 1 | 1 | 1 | 0 | 1 |
| Q8MT77 | Q8MT77_DROME LD28489p OS=Drosophila melanogaster OX=7227 GN=CG17338 PE=2 SV=1 | 2 | 2 | 1 | 1 | 1 | 0 | 1 |
| Q8SWW9 | Q8SWW9_DROME LD18126p OS=Drosophila melanogaster OX=7227 GN=EG:171D11.2 PE=1 SV=1 | 2 | 1 | 1 | 1 | 1 | 0 | 1 |
| Q8T8T9 | Q8T8T9_DROME AT30094p OS=Drosophila melanogaster OX=7227 GN=BcDNA:AT30094 PE=2 SV=1 | 2 | 1 | 1 | 1 | 1 | 0 | 1 |
| Q95SH2 | Q95SH2_DROME Charged multivesicular body protein 1, isoform A OS=Drosophila melanogaster OX=7227 GN=Chmp1 PE=1 SV=1 | 2 | 1 | 1 | 1 | 1 | 0 | 1 |
| Q9I7V5 | Q9I7V5_DROME LD43412p OS=Drosophila melanogaster OX=7227 GN=CG18870 PE=2 SV=1 | 2 | 2 | 1 | 1 | 1 | 0 | 1 |
| Q9V392 | Q9V392_DROME FI01909p OS=Drosophila melanogaster OX=7227 GN=NimC4 PE=2 SV=1 | 2 | 2 | 1 | 1 | 1 | 0 | 1 |
| Q9V3W9 | Q9V3W9_DROME DNA repair protein Rad23 OS=Drosophila melanogaster OX=7227 GN=Rad23 PE=1 SV=1 | 2 | 2 | 1 | 1 | 1 | 0 | 1 |
| Q9VDI6 | Q9VDI6_DROME GM10285p OS=Drosophila melanogaster OX=7227 GN=Dmel\CG17270 PE=2 SV=2 | 2 | 2 | 1 | 1 | 1 | 0 | 1 |
| Q9VE49 | Q9VE49_DROME LP06937p OS=Drosophila melanogaster OX=7227 GN=7702 PE=1 SV=1 | 2 | 2 | 1 | 1 | 1 | 0 | 1 |
| Q9VED7 | Q9VED7_DROME RE37682p OS=Drosophila melanogaster OX=7227 GN=CLLD6 PE=1 SV=1 | 2 | 1 | 1 | 1 | 1 | 0 | 1 |
| Q9VF46 | Q9VF46_DROME GH25158p OS=Drosophila melanogaster OX=7227 GN=Dmel\CG5399 PE=2 SV=1 | 2 | 1 | 1 | 1 | 1 | 0 | 1 |
| Q9VFX0 | Q9VFX0_DROME Aminopeptidase OS=Drosophila melanogaster OX=7227 GN=CG8773-RB PE=1 SV=2 | 2 | 2 | 1 | 1 | 1 | 0 | 1 |
| Q9VG91 | Q9VG91_DROME Uncharacterized protein, isoform A OS=Drosophila melanogaster OX=7227 GN=Dmel\CG10035 PE=4 SV=1 | 2 | 2 | 1 | 1 | 1 | 0 | 1 |
| Q9VGS1 | Q9VGS1_DROME LD23375p OS=Drosophila melanogaster OX=7227 GN=ZnT86D PE=2 SV=1 | 2 | 1 | 1 | 1 | 1 | 0 | 1 |
| Q9VHX6 | Q9VHX6_DROME IP07787p OS=Drosophila melanogaster OX=7227 GN=PIG-H PE=2 SV=1 | 2 | 2 | 1 | 1 | 1 | 0 | 1 |
| Q9VIK7 | Q9VIK7_DROME LD18032p OS=Drosophila melanogaster OX=7227 GN=38D.30 PE=1 SV=4 | 2 | 2 | 1 | 1 | 1 | 0 | 1 |
| Q9VIW6 | Q9VIW6_DROME RE17845p OS=Drosophila melanogaster OX=7227 GN=Rab9 PE=2 SV=1 | 2 | 2 | 1 | 1 | 1 | 0 | 1 |
| Q9VJH8 | Q9VJH8_DROME FI03416p OS=Drosophila melanogaster OX=7227 GN=Ugt36Bc PE=1 SV=1 | 2 | 2 | 1 | 1 | 1 | 0 | 1 |
| Q9VK58 | Q9VK58_DROME Pih1D1, isoform D OS=Drosophila melanogaster OX=7227 GN=Pih1D1 PE=1 SV=5 | 2 | 1 | 1 | 1 | 1 | 0 | 1 |
| Q9VKA0 | Q9VKA0_DROME Crooked OS=Drosophila melanogaster OX=7227 GN=crok PE=2 SV=1 | 2 | 2 | 1 | 1 | 1 | 0 | 1 |
| Q9VL22 | Q9VL22_DROME RH68195p OS=Drosophila melanogaster OX=7227 GN=Dmel\CG4908 PE=2 SV=1 | 2 | 1 | 1 | 1 | 1 | 0 | 1 |
| Q9VL61 | Q9VL61_DROME FI22810p1 OS=Drosophila melanogaster OX=7227 GN=Dmel\CG5853 PE=2 SV=1 | 2 | 2 | 1 | 1 | 1 | 0 | 1 |
| Q9VLR8 | Q9VLR8_DROME GH19970p OS=Drosophila melanogaster OX=7227 GN=SLC5A11 PE=2 SV=2 | 2 | 2 | 1 | 1 | 1 | 0 | 1 |
| Q9VLS8 | Q9VLS8_DROME LD29807p OS=Drosophila melanogaster OX=7227 GN=Dmel\CG8668 PE=1 SV=2 | 2 | 2 | 1 | 1 | 1 | 0 | 1 |
| Q9VLU9 | Q9VLU9_DROME Uncharacterized protein, isoform E OS=Drosophila melanogaster OX=7227 GN=CG7231 PE=4 SV=2 | 2 | 2 | 1 | 1 | 1 | 0 | 1 |
| Q9VMD2 | Q9VMD2_DROME LD47671p OS=Drosophila melanogaster OX=7227 GN=Dmel\CG9536 PE=1 SV=1 | 2 | 1 | 1 | 1 | 1 | 0 | 1 |
| Q9VMX8 | Q9VMX8_DROME Jonah 25Bii, isoform A OS=Drosophila melanogaster OX=7227 GN=Jon25Bii PE=1 SV=1 | 2 | 2 | 1 | 1 | 1 | 0 | 1 |
| Q9VN02 | Q9VN02_DROME GH14561p OS=Drosophila melanogaster OX=7227 GN=Vps24 PE=1 SV=1 | 2 | 2 | 1 | 1 | 1 | 0 | 1 |
| Q9VN16 | Q9VN16_DROME RE35255p OS=Drosophila melanogaster OX=7227 GN=Dmel\CG14646 PE=2 SV=1 | 2 | 2 | 1 | 1 | 1 | 0 | 1 |
| Q9VNR8 | Q9VNR8_DROME LD47277p OS=Drosophila melanogaster OX=7227 GN=Dmel\CG11367 PE=2 SV=3 | 2 | 2 | 1 | 1 | 1 | 0 | 1 |

| UniProt ID | Protein description | # PSMs MVB spectra | # of unique peptides MVB | # of PSMs lysate | # of unique peptides lysate | log2(PSM MVB) | log2(PSM lysate) | FC (log2 of MVB PSM-log2 of lysate PSM) |
|---|---|---|---|---|---|---|---|---|
| Q9VNW1 | Q9VNW1_DROME LD30543p OS=Drosophila melanogaster OX=7227 GN=Dmel\CG7133 PE=2 SV=2 | 2 | 2 | 1 | 1 | 1 | 0 | 1 |
| Q9VPB9 | Q9VPB9_DROME Peroxisomal membrane protein PEX16 OS=Drosophila melanogaster OX=7227 GN=Pex16 PE=1 SV=1 | 2 | 2 | 1 | 1 | 1 | 0 | 1 |
| Q9VPN0 | Q9VPN0_DROME GH04039p OS=Drosophila melanogaster OX=7227 GN=Dmel\CG3625 PE=2 SV=1 | 2 | 1 | 1 | 1 | 1 | 0 | 1 |
| Q9VPP0 | Q9VPP0_DROME Equilibrative nucleoside transporter 1, isoform A OS=Drosophila melanogaster OX=7227 GN=Ent1 PE=2 SV=2 | 2 | 2 | 1 | 1 | 1 | 0 | 1 |
| Q9VQ91 | Q9VQ91_DROME GH18329p OS=Drosophila melanogaster OX=7227 GN=papi PE=1 SV=1 | 2 | 2 | 1 | 1 | 1 | 0 | 1 |
| Q9VQB8 | Q9VQB8_DROME Phosphatidylinositol-glycan biosynthesis class W protein OS=Drosophila melanogaster OX=7227 GN=PIG-Wa PE=1 SV=2 | 2 | 2 | 1 | 1 | 1 | 0 | 1 |
| Q9VRF7 | Q9VRF7_DROME GH09530p OS=Drosophila melanogaster OX=7227 GN=CG1726 PE=1 SV=2 | 2 | 2 | 1 | 1 | 1 | 0 | 1 |
| Q9VSJ2 | Q9VSJ2_DROME Jonah 66Cii OS=Drosophila melanogaster OX=7227 GN=Jon66Cii PE=3 SV=1 | 2 | 2 | 1 | 1 | 1 | 0 | 1 |
| Q9VSX7 | Q9VSX7_DROME LD02310p OS=Drosophila melanogaster OX=7227 GN=4080 PE=2 SV=1 | 2 | 1 | 1 | 1 | 1 | 0 | 1 |
| Q9VT84 | Q9VT84_DROME IP10354p OS=Drosophila melanogaster OX=7227 GN=Dmel\CG6628 PE=2 SV=1 | 2 | 1 | 1 | 1 | 1 | 0 | 1 |
| Q9VTM6 | Q9VTM6_DROME Crimpled OS=Drosophila melanogaster OX=7227 GN=crim PE=2 SV=1 | 2 | 1 | 1 | 1 | 1 | 0 | 1 |
| Q9VW37 | Q9VW37_DROME LP12374p OS=Drosophila melanogaster OX=7227 GN=SREBP PE=1 SV=1 | 2 | 2 | 1 | 1 | 1 | 0 | 1 |
| Q9VWJ3 | Q9VWJ3_DROME LD05272p OS=Drosophila melanogaster OX=7227 GN=CG8020 PE=1 SV=2 | 2 | 2 | 1 | 1 | 1 | 0 | 1 |
| Q9VXY3 | Q9VXY3_DROME Dihydrolipoamide acetyltransferase component of pyruvate dehydrogenase complex OS=Drosophila melanogaster OX=7227 GN=Dmel\CG5599 PE=1 SV=1 | 2 | 2 | 1 | 1 | 1 | 0 | 1 |
| Q9VZ71 | Q9VZ71_DROME RE01453p OS=Drosophila melanogaster OX=7227 GN=Dmel\CG15211 PE=2 SV=1 | 2 | 1 | 1 | 1 | 1 | 0 | 1 |
| Q9VZZ0 | Q9VZZ0_DROME RE02644p OS=Drosophila melanogaster OX=7227 GN=Dmel\CG1246 PE=2 SV=1 | 2 | 1 | 1 | 1 | 1 | 0 | 1 |
| Q9W073 | Q9W073_DROME RH22148p OS=Drosophila melanogaster OX=7227 GN=Dmel\CG12025 PE=2 SV=1 | 2 | 1 | 1 | 1 | 1 | 0 | 1 |
| Q9W095 | Q9W095_DROME FI03690p OS=Drosophila melanogaster OX=7227 GN=Gk2 PE=2 SV=2 | 2 | 1 | 1 | 1 | 1 | 0 | 1 |
| Q9W228 | Q9W228_DROME RE07815p OS=Drosophila melanogaster OX=7227 GN=Ugt58Fa PE=1 SV=3 | 2 | 2 | 1 | 1 | 1 | 0 | 1 |
| Q9W297 | Q9W297_DROME Ceramide glucosyltransferase OS=Drosophila melanogaster OX=7227 GN=GlcT PE=2 SV=1 | 2 | 2 | 1 | 1 | 1 | 0 | 1 |
| Q9W2E4 | Q9W2E4_DROME CG9865, isoform A OS=Drosophila melanogaster OX=7227 GN=PIG-M PE=2 SV=1 | 2 | 2 | 1 | 1 | 1 | 0 | 1 |
| Q9W2J4 | Q9W2J4_DROME UDP-glucuronosyltransferase OS=Drosophila melanogaster OX=7227 GN=BEST:GH09393 PE=1 SV=2 | 2 | 2 | 1 | 1 | 1 | 0 | 1 |
| Q9W2M9 | Q9W2M9_DROME FI16623p1 OS=Drosophila melanogaster OX=7227 GN=CG15653-RA PE=1 SV=1 | 2 | 2 | 1 | 1 | 1 | 0 | 1 |
| Q9W3L4 | Q9W3L4_DROME GH20802p OS=Drosophila melanogaster OX=7227 GN=Dmel\CG2233 PE=1 SV=1 | 2 | 2 | 1 | 1 | 1 | 0 | 1 |
| Q9W4C4 | Q9W4C4_DROME GM14014p OS=Drosophila melanogaster OX=7227 GN=spoon PE=1 SV=2 | 2 | 1 | 1 | 1 | 1 | 0 | 1 |
| Q9W4M3 | Q9W4M3_DROME Femcoat OS=Drosophila melanogaster OX=7227 GN=Femcoat PE=4 SV=2 | 2 | 2 | 1 | 1 | 1 | 0 | 1 |
| Q9W4W6 | Q9W4W6_DROME CMP-sialic acid/UDP-galactose transporter OS=Drosophila melanogaster OX=7227 GN=Ugalt PE=1 SV=1 | 2 | 2 | 1 | 1 | 1 | 0 | 1 |
| Q9W4X4 | Q9W4X4_DROME Phosphatidylinositol 4-kinase III alpha, isoform C OS=Drosophila melanogaster OX=7227 GN=PI4KIIIalpha PE=1 SV=3 | 2 | 2 | 1 | 1 | 1 | 0 | 1 |
| Q9W509 | Q9W509_DROME EG:103B4.3 protein OS=Drosophila melanogaster OX=7227 GN=Mct1 PE=1 SV=2 | 2 | 2 | 1 | 1 | 1 | 0 | 1 |
| Q9W551 | Q9W551_DROME GEO02601p1 OS=Drosophila melanogaster OX=7227 GN=Dmel\CG14817 PE=2 SV=1 | 2 | 2 | 1 | 1 | 1 | 0 | 1 |
| X2J9E8 | X2J9E8_DROME Calsyntenin-1, isoform D OS=Drosophila melanogaster OX=7227 GN=Cals PE=4 SV=1 | 2 | 2 | 1 | 1 | 1 | 0 | 1 |
| P13496 | DCTN1_DROME Dynactin subunit 1 OS=Drosophila melanogaster OX=7227 GN=DCTN1-p150 PE=1 SV=2 | 1 | 1 | 23 | 15 | 0 | 4.523561956 | -4.523561956 |
| Q9VYW4 | Q9VYW4_DROME LD04461p OS=Drosophila melanogaster OX=7227 GN=Dmel\CG1703 PE=1 SV=1 | 1 | 1 | 23 | 15 | 0 | 4.523561956 | -4.523561956 |
| Q9VKM1 | PIWI_DROME Protein piwi OS=Drosophila melanogaster OX=7227 GN=piwi PE=1 SV=1 | 1 | 1 | 21 | 12 | 0 | 4.392317423 | -4.392317423 |
| Q86BY9 | RIG_DROME Protein rigor mortis OS=Drosophila melanogaster OX=7227 GN=rig PE=1 SV=1 | 1 | 1 | 19 | 14 | 0 | 4.247927513 | -4.247927513 |
| X2J9F3 | X2J9F3_DROME Nucleoporin 160kD, isoform B OS=Drosophila melanogaster OX=7227 GN=Nup160 PE=4 SV=1 | 1 | 1 | 19 | 14 | 0 | 4.247927513 | -4.247927513 |
| Q9VNF8 | Q9VNF8_DROME Protein transport protein SEC23 OS=Drosophila melanogaster OX=7227 GN=Sec23 PE=1 SV=2 | 1 | 1 | 18 | 10 | 0 | 4.169925001 | -4.169925001 |
| Q7K0D8 | NUP50_DROME Nuclear pore complex protein Nup50 OS=Drosophila melanogaster OX=7227 GN=Nup50 PE=2 SV=1 | 1 | 1 | 17 | 11 | 0 | 4.087462841 | -4.087462841 |
| Q9VCC0 | CHRD1_DROME Cysteine and histidine-rich domain-containing protein OS=Drosophila melanogaster OX=7227 GN=CHORD PE=1 SV=1 | 1 | 1 | 15 | 9 | 0 | 3.906890596 | -3.906890596 |
| Q9VKK1 | EDC4_DROME Enhancer of mRNA-decapping protein 4 homolog OS=Drosophila melanogaster OX=7227 GN=Ge-1 PE=1 SV=2 | 1 | 1 | 15 | 11 | 0 | 3.906890596 | -3.906890596 |
| A0A0B4LEV3 | A0A0B4LEV3_DROME CG8635, isoform B OS=Drosophila melanogaster OX=7227 GN=CG8635 PE=4 SV=1 | 1 | 1 | 15 | 8 | 0 | 3.906890596 | -3.906890596 |
| B7Z0E2 | B7Z0E2_DROME FI18812p1 OS=Drosophila melanogaster OX=7227 GN=Unr PE=1 SV=1 | 1 | 1 | 15 | 11 | 0 | 3.906890596 | -3.906890596 |
| Q9VY91 | Q9VY91_DROME LD21074p OS=Drosophila melanogaster OX=7227 GN=Pdcd4 PE=1 SV=2 | 1 | 1 | 15 | 9 | 0 | 3.906890596 | -3.906890596 |
| P53034 | RFC2_DROME Replication factor C subunit 2 OS=Drosophila melanogaster OX=7227 GN=RfC4 PE=1 SV=1 | 1 | 1 | 14 | 7 | 0 | 3.807354922 | -3.807354922 |
| A0A0B4LHQ4 | A0A0B4LHQ4_DROME Circadian trip, isoform M OS=Drosophila melanogaster OX=7227 GN=ctrip PE=1 SV=1 | 1 | 1 | 14 | 12 | 0 | 3.807354922 | -3.807354922 |
| B7Z0D3 | B7Z0D3_DROME Uncharacterized protein, isoform D OS=Drosophila melanogaster OX=7227 GN=CG7546 PE=1 SV=1 | 1 | 1 | 14 | 9 | 0 | 3.807354922 | -3.807354922 |
| Q95WY3 | Q95WY3_DROME FI04781p OS=Drosophila melanogaster OX=7227 GN=Nop56 PE=1 SV=1 | 1 | 1 | 14 | 8 | 0 | 3.807354922 | -3.807354922 |
| P39205 | CIN_DROME Molybdenum cofactor synthesis protein cinnamon OS=Drosophila melanogaster OX=7227 GN=cin PE=1 SV=3 | 1 | 1 | 13 | 8 | 0 | 3.700439718 | -3.700439718 |
| Q9VUR3 | AIMP2_DROME Probable aminoacyl tRNA synthase complex-interacting multifunctional protein 2 OS=Drosophila melanogaster OX=7227 GN=AIMP2 PE=3 SV=1 | 1 | 1 | 13 | 10 | 0 | 3.700439718 | -3.700439718 |
| Q9W4X7 | EI3G1_DROME Eukaryotic translation initiation factor 3 subunit G-1 OS=Drosophila melanogaster OX=7227 GN=eIF3g1 PE=1 SV=1 | 1 | 1 | 13 | 4 | 0 | 3.700439718 | -3.700439718 |
| Q7K533 | Q7K533_DROME GH14572p OS=Drosophila melanogaster OX=7227 GN=Gbp2 PE=1 SV=1 | 1 | 1 | 13 | 7 | 0 | 3.700439718 | -3.700439718 |
| Q9W088 | DPOD2_DROME DNA polymerase delta small subunit OS=Drosophila melanogaster OX=7227 GN=CG12018 PE=2 SV=1 | 1 | 1 | 12 | 7 | 0 | 3.584962501 | -3.584962501 |
| Q9VVM8 | Q9VVM8_DROME FI02004p OS=Drosophila melanogaster OX=7227 GN=CG5290-RA PE=1 SV=1 | 1 | 1 | 12 | 5 | 0 | 3.584962501 | -3.584962501 |
| Q8T6B9 | PUF68_DROME Poly(U)-binding-splicing factor half pint OS=Drosophila melanogaster OX=7227 GN=hfp PE=1 SV=2 | 1 | 1 | 11 | 6 | 0 | 3.459431619 | -3.459431619 |
| Q9W0M7 | SF3B3_DROME Splicing factor 3B subunit 3 OS=Drosophila melanogaster OX=7227 GN=Sf3b3 PE=1 SV=2 | 1 | 1 | 11 | 10 | 0 | 3.459431619 | -3.459431619 |
| Q24050 | Q24050_DROME Anon-i1 protein OS=Drosophila melanogaster OX=7227 GN=Dmel\CG2034 PE=1 SV=1 | 1 | 1 | 11 | 6 | 0 | 3.459431619 | -3.459431619 |
| Q9V3A7 | Q9V3A7_DROME Structural maintenance of chromosomes protein OS=Drosophila melanogaster OX=7227 GN=glu PE=1 SV=1 | 1 | 1 | 11 | 9 | 0 | 3.459431619 | -3.459431619 |
| Q9VAX8 | Q9VAX8_DROME LD28793p OS=Drosophila melanogaster OX=7227 GN=Dmel\CG4849 PE=1 SV=1 | 1 | 1 | 11 | 8 | 0 | 3.459431619 | -3.459431619 |
| Q9VI56 | Q9VI56_DROME GEO05313p1 OS=Drosophila melanogaster OX=7227 GN=Dmel\CG1943 PE=1 SV=1 | 1 | 1 | 11 | 4 | 0 | 3.459431619 | -3.459431619 |
| Q9VTB0 | Q9VTB0_DROME LD35289p OS=Drosophila melanogaster OX=7227 GN=Dmel\CG8003 PE=1 SV=1 | 1 | 1 | 11 | 6 | 0 | 3.459431619 | -3.459431619 |
| Q9W1H5 | Q9W1H5_DROME Decapping protein 1, isoform A OS=Drosophila melanogaster OX=7227 GN=DCP1 PE=1 SV=1 | 1 | 1 | 11 | 7 | 0 | 3.459431619 | -3.459431619 |
| Q9W1Q8 | Q9W1Q8_DROME LD05345p OS=Drosophila melanogaster OX=7227 GN=eIF2Bdelta PE=1 SV=1 | 1 | 1 | 11 | 6 | 0 | 3.459431619 | -3.459431619 |
| X2JJP5 | X2JJP5_DROME Secretory 16, isoform H OS=Drosophila melanogaster OX=7227 GN=Sec16 PE=1 SV=1 | 1 | 1 | 11 | 9 | 0 | 3.459431619 | -3.459431619 |
| Q94901 | LARK_DROME RNA-binding protein lark OS=Drosophila melanogaster OX=7227 GN=lark PE=1 SV=1 | 1 | 1 | 10 | 9 | 0 | 3.321928095 | -3.321928095 |
| Q9VHC7 | Q9VHC7_DROME FI21236p1 OS=Drosophila melanogaster OX=7227 GN=rump PE=1 SV=1 | 1 | 1 | 10 | 6 | 0 | 3.321928095 | -3.321928095 |
| Q9VP77 | Q9VP77_DROME LD23875p OS=Drosophila melanogaster OX=7227 GN=Dmel\CG10565 PE=1 SV=1 | 1 | 1 | 10 | 9 | 0 | 3.321928095 | -3.321928095 |
| Q9VWE7 | Q9VWE7_DROME LD26789p OS=Drosophila melanogaster OX=7227 GN=Ranbp21 PE=1 SV=1 | 1 | 1 | 10 | 8 | 0 | 3.321928095 | -3.321928095 |
| Q7K486 | ARMC6_DROME Armadillo repeat-containing protein 6 homolog OS=Drosophila melanogaster OX=7227 GN=CG5721 PE=1 SV=1 | 1 | 1 | 9 | 5 | 0 | 3.169925001 | -3.169925001 |
| Q7KUT2 | LONM_DROME Lon protease homolog, mitochondrial OS=Drosophila melanogaster OX=7227 GN=Lon PE=1 SV=1 | 1 | 1 | 9 | 8 | 0 | 3.169925001 | -3.169925001 |
| Q8MR62 | VIAF1_DROME Viral IAP-associated factor homolog OS=Drosophila melanogaster OX=7227 GN=viaf PE=1 SV=1 | 1 | 1 | 9 | 5 | 0 | 3.169925001 | -3.169925001 |
| Q94524 | DYLT_DROME Dynein light chain Tctex-type OS=Drosophila melanogaster OX=7227 GN=Dlc90F PE=1 SV=1 | 1 | 1 | 9 | 5 | 0 | 3.169925001 | -3.169925001 |
| Q9W445 | MCTS1_DROME Malignant T-cell-amplified sequence 1 homolog OS=Drosophila melanogaster OX=7227 GN=MCTS1 PE=1 SV=1 | 1 | 1 | 9 | 4 | 0 | 3.169925001 | -3.169925001 |
| A0A0B4K6L4 | A0A0B4K6L4_DROME Amidophosphoribosyltransferase OS=Drosophila melanogaster OX=7227 GN=Prat PE=1 SV=1 | 1 | 1 | 9 | 6 | 0 | 3.169925001 | -3.169925001 |
| Q9VGF9 | Q9VGF9_DROME Aurora A OS=Drosophila melanogaster OX=7227 GN=aurA PE=1 SV=1 | 1 | 1 | 9 | 6 | 0 | 3.169925001 | -3.169925001 |
| Q9W3Y3 | Q9W3Y3_DROME LD13807p OS=Drosophila melanogaster OX=7227 GN=Dmel\CG3226 PE=1 SV=1 | 1 | 1 | 9 | 5 | 0 | 3.169925001 | -3.169925001 |
| Q9XZ19 | Q9XZ19_DROME Uncharacterized protein OS=Drosophila melanogaster OX=7227 GN=BcDNA:LD21537 PE=1 SV=1 | 1 | 1 | 9 | 6 | 0 | 3.169925001 | -3.169925001 |
| A1Z9X0 | APKC_DROME Atypical protein kinase C OS=Drosophila melanogaster OX=7227 GN=aPKC PE=1 SV=1 | 1 | 1 | 8 | 3 | 0 | 3 | -3 |
| Q7KQM6 | GGYF1_DROME GIGYF family protein CG11148 OS=Drosophila melanogaster OX=7227 GN=CG11148 PE=1 SV=2 | 1 | 1 | 8 | 7 | 0 | 3 | -3 |
| Q9VU02 | SMD1_DROME Probable small nuclear ribonucleoprotein Sm D1 OS=Drosophila melanogaster OX=7227 GN=SmD1 PE=1 SV=1 | 1 | 1 | 8 | 4 | 0 | 3 | -3 |
| Q9VVU5 | CSN1_DROME COP9 signalosome complex subunit 1b OS=Drosophila melanogaster OX=7227 GN=CSN1b PE=2 SV=1 | 1 | 1 | 8 | 5 | 0 | 3 | -3 |
| M9PDJ9 | M9PDJ9_DROME Tetratricopeptide repeat protein 2, isoform C OS=Drosophila melanogaster OX=7227 GN=Tpr2 PE=1 SV=1 | 1 | 1 | 8 | 6 | 0 | 3 | -3 |

| UniProt ID | Protein description | # PSMs MVB spectra | # of unique peptides MVB | # of PSMs lysate | # of unique peptides lysate | log2(PSM MVB) | log2(PSM lysate) | FC (log2 of MVB PSM-log2 of lysate PSM) |
|---|---|---|---|---|---|---|---|---|
| M9PIL3 | M9PIL3_DROME ADP-ribosylation factor GTPase activating protein 3, isoform G OS=Drosophila melanogaster OX=7227 GN=ArfGAP3 PE=1 SV=1 | 1 | 1 | 8 | 7 | 0 | 3 | -3 |
| Q7JX41 | Q7JX41_DROME Geminin OS=Drosophila melanogaster OX=7227 GN=geminin PE=1 SV=1 | 1 | 1 | 8 | 3 | 0 | 3 | -3 |
| Q7KMH5 | Q7KMH5_DROME BcDNA.LD29892 OS=Drosophila melanogaster OX=7227 GN=Smyd4-4 PE=1 SV=1 | 1 | 1 | 8 | 7 | 0 | 3 | -3 |
| Q8IPA2 | Q8IPA2_DROME FI17312p1 OS=Drosophila melanogaster OX=7227 GN=Mal-B2 PE=1 SV=1 | 1 | 1 | 8 | 6 | 0 | 3 | -3 |
| Q8MKK1 | Q8MKK1_DROME RH09246p OS=Drosophila melanogaster OX=7227 GN=AIMP3 PE=1 SV=1 | 1 | 1 | 8 | 4 | 0 | 3 | -3 |
| Q8SXF0 | Q8SXF0_DROME Mitochondrial ribosomal protein S9 OS=Drosophila melanogaster OX=7227 GN=mRpS9 PE=1 SV=1 | 1 | 1 | 8 | 6 | 0 | 3 | -3 |
| Q9VP57 | Q9VP57_DROME LD15904p OS=Drosophila melanogaster OX=7227 GN=pzg PE=1 SV=1 | 1 | 1 | 8 | 4 | 0 | 3 | -3 |
| Q9VT75 | Q9VT75_DROME LD19812p OS=Drosophila melanogaster OX=7227 GN=vnc PE=1 SV=1 | 1 | 1 | 8 | 4 | 0 | 3 | -3 |
| Q9VVC8 | Q9VVC8_DROME LD23434p OS=Drosophila melanogaster OX=7227 GN=Dmel\CG6664 PE=1 SV=1 | 1 | 1 | 8 | 6 | 0 | 3 | -3 |
| Q9VVW6 | Q9VVW6_DROME LD22716p OS=Drosophila melanogaster OX=7227 GN=MESR6 PE=1 SV=1 | 1 | 1 | 8 | 7 | 0 | 3 | -3 |
| Q6AWN0 | MTND_DROME 1,2-dihydroxy-3-keto-5-methylthiopentene dioxygenase OS=Drosophila melanogaster OX=7227 GN=Adi1 PE=2 SV=2 | 1 | 1 | 7 | 5 | 0 | 2.807354922 | -2.807354922 |
| Q9V3D6 | CPSF2_DROME Probable cleavage and polyadenylation specificity factor subunit 2 OS=Drosophila melanogaster OX=7227 GN=Cpsf100 PE=1 SV=1 | 1 | 1 | 7 | 6 | 0 | 2.807354922 | -2.807354922 |
| Q9VFR0 | BCCIP_DROME Protein BCCIP homolog OS=Drosophila melanogaster OX=7227 GN=CG9286 PE=2 SV=2 | 1 | 1 | 7 | 4 | 0 | 2.807354922 | -2.807354922 |
| Q9VL31 | RIP_DROME RIP-like protein OS=Drosophila melanogaster OX=7227 GN=Ripalpha PE=2 SV=1 | 1 | 1 | 7 | 4 | 0 | 2.807354922 | -2.807354922 |
| Q9VYR0 | LS12A_DROME Protein LSM12 homolog A OS=Drosophila melanogaster OX=7227 GN=CG15735 PE=2 SV=2 | 1 | 1 | 7 | 2 | 0 | 2.807354922 | -2.807354922 |
| Q9W252 | RAD50_DROME DNA repair protein RAD50 OS=Drosophila melanogaster OX=7227 GN=rad50 PE=2 SV=4 | 1 | 1 | 7 | 6 | 0 | 2.807354922 | -2.807354922 |
| A1Z9I5 | A1Z9I5_DROME FI24033p1 OS=Drosophila melanogaster OX=7227 GN=tum PE=1 SV=1 | 1 | 1 | 7 | 7 | 0 | 2.807354922 | -2.807354922 |
| M9PCL5 | M9PCL5_DROME Mitochondrial ribosomal protein S7, isoform B OS=Drosophila melanogaster OX=7227 GN=mRpS7 PE=4 SV=1 | 1 | 1 | 7 | 5 | 0 | 2.807354922 | -2.807354922 |
| M9PGG8 | M9PGG8_DROME Uncharacterized protein, isoform B OS=Drosophila melanogaster OX=7227 GN=CG40016 PE=1 SV=1 | 1 | 1 | 7 | 4 | 0 | 2.807354922 | -2.807354922 |
| Q7KTW9 | Q7KTW9_DROME Asparagine synthetase OS=Drosophila melanogaster OX=7227 GN=AsnS PE=2 SV=1 | 1 | 1 | 7 | 5 | 0 | 2.807354922 | -2.807354922 |
| Q8IP62 | Q8IP62_DROME Mitochondrial ribosomal protein S23 OS=Drosophila melanogaster OX=7227 GN=mRpS23 PE=1 SV=1 | 1 | 1 | 7 | 5 | 0 | 2.807354922 | -2.807354922 |
| Q9VAJ1 | Q9VAJ1_DROME Condensin complex subunit 1 OS=Drosophila melanogaster OX=7227 GN=Cap-D2 PE=1 SV=1 | 1 | 1 | 7 | 5 | 0 | 2.807354922 | -2.807354922 |
| Q9VGH5 | Q9VGH5_DROME AT27789p OS=Drosophila melanogaster OX=7227 GN=glo PE=1 SV=1 | 1 | 1 | 7 | 6 | 0 | 2.807354922 | -2.807354922 |
| Q9VNI4 | Q9VNI4_DROME GH13924p OS=Drosophila melanogaster OX=7227 GN=Dmel\CG2046 PE=1 SV=1 | 1 | 1 | 7 | 5 | 0 | 2.807354922 | -2.807354922 |
| Q9VZX6 | Q9VZX6_DROME LD06441p OS=Drosophila melanogaster OX=7227 GN=mRpS35 PE=1 SV=1 | 1 | 1 | 7 | 4 | 0 | 2.807354922 | -2.807354922 |
| Q9Y118 | Q9Y118_DROME BG:DS07295.3 OS=Drosophila melanogaster OX=7227 GN=Pol32 PE=1 SV=1 | 1 | 1 | 7 | 6 | 0 | 2.807354922 | -2.807354922 |
| X2JEM4 | X2JEM4_DROME GEO07185p1 OS=Drosophila melanogaster OX=7227 GN=RpS28b PE=2 SV=1 | 1 | 1 | 7 | 3 | 0 | 2.807354922 | -2.807354922 |
| A1YK02 | NUP75_DROME Nuclear pore complex protein Nup75 OS=Drosophila melanogaster OX=7227 GN=Nup75 PE=2 SV=1 | 1 | 1 | 6 | 5 | 0 | 2.584962501 | -2.584962501 |
| Q9V637 | PSMF1_DROME Proteasome inhibitor PI31 subunit OS=Drosophila melanogaster OX=7227 GN=PI31 PE=1 SV=1 | 1 | 1 | 6 | 5 | 0 | 2.584962501 | -2.584962501 |
| Q9VYS3 | RENT1_DROME Regulator of nonsense transcripts 1 homolog OS=Drosophila melanogaster OX=7227 GN=Upf1 PE=1 SV=2 | 1 | 1 | 6 | 6 | 0 | 2.584962501 | -2.584962501 |
| Q9VYY3 | UBA5_DROME Ubiquitin-like modifier-activating enzyme 5 OS=Drosophila melanogaster OX=7227 GN=Uba5 PE=2 SV=1 | 1 | 1 | 6 | 4 | 0 | 2.584962501 | -2.584962501 |
| Q9W4X9 | ZW10_DROME Centromere/kinetochore protein zw10 OS=Drosophila melanogaster OX=7227 GN=Zw10 PE=1 SV=2 | 1 | 1 | 6 | 4 | 0 | 2.584962501 | -2.584962501 |
| Q5U0V7 | Q5U0V7_DROME GH06496p OS=Drosophila melanogaster OX=7227 GN=Synj PE=1 SV=1 | 1 | 1 | 6 | 6 | 0 | 2.584962501 | -2.584962501 |
| Q7KSW3 | Q7KSW3_DROME Sugarlethal, isoform B OS=Drosophila melanogaster OX=7227 GN=sgll PE=1 SV=1 | 1 | 1 | 6 | 4 | 0 | 2.584962501 | -2.584962501 |
| Q9VG04 | Q9VG04_DROME LP04475p OS=Drosophila melanogaster OX=7227 GN=Dmel\CG8031 PE=1 SV=1 | 1 | 1 | 6 | 4 | 0 | 2.584962501 | -2.584962501 |
| Q9VT60 | Q9VT60_DROME Calcium-independent phospholipase A2 VIA, isoform A OS=Drosophila melanogaster OX=7227 GN=iPLA2-VIA PE=1 SV=3 | 1 | 1 | 6 | 4 | 0 | 2.584962501 | -2.584962501 |
| X2JFX0 | X2JFX0_DROME Uncharacterized protein, isoform C OS=Drosophila melanogaster OX=7227 GN=Dmel\CG8188 PE=3 SV=1 | 1 | 1 | 6 | 5 | 0 | 2.584962501 | -2.584962501 |
| A1Z9A8 | PTCD3_DROME Protein PTCD3 homolog, mitochondrial OS=Drosophila melanogaster OX=7227 GN=CG4679 PE=2 SV=1 | 1 | 1 | 5 | 5 | 0 | 2.321928095 | -2.321928095 |
| O76932 | PP4C_DROME Serine/threonine-protein phosphatase 4 catalytic subunit OS=Drosophila melanogaster OX=7227 GN=Pp4-19C PE=2 SV=1 | 1 | 1 | 5 | 2 | 0 | 2.321928095 | -2.321928095 |
| Q95RA8 | MOB1_DROME MOB kinase activator-like 1 OS=Drosophila melanogaster OX=7227 GN=mats PE=1 SV=1 | 1 | 1 | 5 | 4 | 0 | 2.321928095 | -2.321928095 |
| Q9V3G3 | PPIE_DROME Peptidyl-prolyl cis-trans isomerase E OS=Drosophila melanogaster OX=7227 GN=cyp33 PE=1 SV=1 | 1 | 1 | 5 | 4 | 0 | 2.321928095 | -2.321928095 |
| Q9VKD6 | DGT2_DROME Augmin complex subunit dgt2 OS=Drosophila melanogaster OX=7227 GN=dgt2 PE=1 SV=1 | 1 | 1 | 5 | 4 | 0 | 2.321928095 | -2.321928095 |
| Q9VLD6 | BOREA_DROME Borealin OS=Drosophila melanogaster OX=7227 GN=borr PE=1 SV=1 | 1 | 1 | 5 | 5 | 0 | 2.321928095 | -2.321928095 |
| Q9VPP5 | RNH2A_DROME Ribonuclease H2 subunit A OS=Drosophila melanogaster OX=7227 GN=CG13690 PE=2 SV=1 | 1 | 1 | 5 | 3 | 0 | 2.321928095 | -2.321928095 |
| A0A0B4LF92 | A0A0B4LF92_DROME CG4646, isoform B OS=Drosophila melanogaster OX=7227 GN=CG4646 PE=4 SV=1 | 1 | 1 | 5 | 4 | 0 | 2.321928095 | -2.321928095 |
| H9XVM6 | H9XVM6_DROME Crk oncogene, isoform E OS=Drosophila melanogaster OX=7227 GN=Crk PE=1 SV=1 | 1 | 1 | 5 | 4 | 0 | 2.321928095 | -2.321928095 |
| Q7JW66 | Q7JW66_DROME LD21545p OS=Drosophila melanogaster OX=7227 GN=CG12129 PE=1 SV=1 | 1 | 1 | 5 | 3 | 0 | 2.321928095 | -2.321928095 |
| Q9V3F8 | Q9V3F8_DROME Pyrroline-5-carboxylate reductase OS=Drosophila melanogaster OX=7227 GN=P5cr PE=1 SV=1 | 1 | 1 | 5 | 3 | 0 | 2.321928095 | -2.321928095 |
| Q9V470 | Q9V470_DROME Succinate--CoA ligase [GDP-forming] subunit beta, mitochondrial OS=Drosophila melanogaster OX=7227 GN=ScsbetaG PE=1 SV=1 | 1 | 1 | 5 | 5 | 0 | 2.321928095 | -2.321928095 |
| Q9VF28 | Q9VF28_DROME Actin-related protein 2/3 complex subunit 3 OS=Drosophila melanogaster OX=7227 GN=Arpc3A PE=1 SV=4 | 1 | 1 | 5 | 4 | 0 | 2.321928095 | -2.321928095 |
| Q9VF77 | Q9VF77_DROME FI16517p1 OS=Drosophila melanogaster OX=7227 GN=Trax PE=1 SV=3 | 1 | 1 | 5 | 4 | 0 | 2.321928095 | -2.321928095 |
| Q9VGV8 | Q9VGV8_DROME GM03430p OS=Drosophila melanogaster OX=7227 GN=Dmel\CG4511 PE=2 SV=1 | 1 | 1 | 5 | 5 | 0 | 2.321928095 | -2.321928095 |
| Q9VI64 | Q9VI64_DROME LD30995p OS=Drosophila melanogaster OX=7227 GN=Dmel\CG10098 PE=1 SV=1 | 1 | 1 | 5 | 3 | 0 | 2.321928095 | -2.321928095 |
| Q9VKW8 | Q9VKW8_DROME U6 snRNA-associated Sm-like protein LSm4 OS=Drosophila melanogaster OX=7227 GN=BcDNA:RE35747 PE=1 SV=3 | 1 | 1 | 5 | 3 | 0 | 2.321928095 | -2.321928095 |
| Q9VMU7 | Q9VMU7_DROME FI16614p1 OS=Drosophila melanogaster OX=7227 GN=CG14036-RA PE=2 SV=1 | 1 | 1 | 5 | 4 | 0 | 2.321928095 | -2.321928095 |
| Q9VP29 | Q9VP29_DROME RE39629p OS=Drosophila melanogaster OX=7227 GN=Dmel\CG7519 PE=1 SV=1 | 1 | 1 | 5 | 4 | 0 | 2.321928095 | -2.321928095 |
| Q9W4K1 | Q9W4K1_DROME AP-3 complex subunit beta OS=Drosophila melanogaster OX=7227 GN=rb PE=1 SV=1 | 1 | 1 | 5 | 4 | 0 | 2.321928095 | -2.321928095 |
| O76454 | PHS_DROME Pterin-4-alpha-carbinolamine dehydratase OS=Drosophila melanogaster OX=7227 GN=Pcd PE=1 SV=2 | 1 | 1 | 4 | 2 | 0 | 2 | -2 |
| P25991 | SUF_DROME Protein suppressor of forked OS=Drosophila melanogaster OX=7227 GN=su(f) PE=1 SV=2 | 1 | 1 | 4 | 4 | 0 | 2 | -2 |
| P52485 | UBCD2_DROME Ubiquitin-conjugating enzyme E2-24 kDa OS=Drosophila melanogaster OX=7227 GN=Ubc2 PE=2 SV=1 | 1 | 1 | 4 | 3 | 0 | 2 | -2 |
| Q23976 | KU70_DROME ATP-dependent DNA helicase 2 subunit 1 OS=Drosophila melanogaster OX=7227 GN=Irbp PE=2 SV=3 | 1 | 1 | 4 | 3 | 0 | 2 | -2 |
| Q9V4Q8 | RU2A_DROME Probable U2 small nuclear ribonucleoprotein A' OS=Drosophila melanogaster OX=7227 GN=U2A PE=1 SV=1 | 1 | 1 | 4 | 3 | 0 | 2 | -2 |
| Q9XZS3 | TMA7_DROME Translation machinery-associated protein 7 homolog OS=Drosophila melanogaster OX=7227 GN=CG13364 PE=4 SV=1 | 1 | 1 | 4 | 3 | 0 | 2 | -2 |
| M9NF89 | M9NF89_DROME U2 snRNP auxiliary factor large subunit OS=Drosophila melanogaster OX=7227 GN=U2af50 PE=3 SV=1 | 1 | 1 | 4 | 4 | 0 | 2 | -2 |
| M9PBN7 | M9PBN7_DROME Capping protein beta, isoform B OS=Drosophila melanogaster OX=7227 GN=cpb PE=4 SV=1 | 1 | 1 | 4 | 3 | 0 | 2 | -2 |
| M9PGD7 | M9PGD7_DROME Uncharacterized protein, isoform B OS=Drosophila melanogaster OX=7227 GN=CG10802 PE=1 SV=1 | 1 | 1 | 4 | 3 | 0 | 2 | -2 |
| O97111 | O97111_DROME LD29223p OS=Drosophila melanogaster OX=7227 GN=par-6 PE=1 SV=1 | 1 | 1 | 4 | 3 | 0 | 2 | -2 |
| Q7KSD3 | Q7KSD3_DROME Frayed, isoform A OS=Drosophila melanogaster OX=7227 GN=fray PE=1 SV=1 | 1 | 1 | 4 | 3 | 0 | 2 | -2 |
| Q8IML6 | Q8IML6_DROME IP14839p OS=Drosophila melanogaster OX=7227 GN=Pdhb PE=1 SV=1 | 1 | 1 | 4 | 3 | 0 | 2 | -2 |
| Q8SY58 | Q8SY58_DROME GH02315p OS=Drosophila melanogaster OX=7227 GN=Dmel\CG4872 PE=1 SV=1 | 1 | 1 | 4 | 3 | 0 | 2 | -2 |
| Q9VC52 | Q9VC52_DROME Protoporphyrinogen oxidase OS=Drosophila melanogaster OX=7227 GN=Ppox PE=1 SV=1 | 1 | 1 | 4 | 2 | 0 | 2 | -2 |
| Q9VWS1 | Q9VWS1_DROME LD25271p OS=Drosophila melanogaster OX=7227 GN=Dmel\CG6617 PE=1 SV=1 | 1 | 1 | 4 | 4 | 0 | 2 | -2 |
| Q9VXI1 | Q9VXI1_DROME RH68464p OS=Drosophila melanogaster OX=7227 GN=Dmel\CG9914 PE=1 SV=2 | 1 | 1 | 4 | 4 | 0 | 2 | -2 |
| Q9W0B3 | Q9W0B3_DROME GH19706p OS=Drosophila melanogaster OX=7227 GN=Vta1 PE=1 SV=1 | 1 | 1 | 4 | 3 | 0 | 2 | -2 |
| Q9W1E5 | Q9W1E5_DROME Anaphase-promoting complex subunit 2 OS=Drosophila melanogaster OX=7227 GN=mr PE=1 SV=1 | 1 | 1 | 4 | 3 | 0 | 2 | -2 |
| O01939 | MST_DROME Protein misato OS=Drosophila melanogaster OX=7227 GN=mst PE=2 SV=1 | 1 | 1 | 3 | 3 | 0 | 1.584962501 | -1.584962501 |
| P43332 | SNRPA_DROME U1 small nuclear ribonucleoprotein A OS=Drosophila melanogaster OX=7227 GN=snf PE=1 SV=1 | 1 | 1 | 3 | 2 | 0 | 1.584962501 | -1.584962501 |
| P49963 | SRP19_DROME Signal recognition particle 19 kDa protein OS=Drosophila melanogaster OX=7227 GN=Srp19 PE=2 SV=2 | 1 | 1 | 3 | 3 | 0 | 1.584962501 | -1.584962501 |
| Q70PY2 | PGSB1_DROME Peptidoglycan-recognition protein SB1 OS=Drosophila melanogaster OX=7227 GN=PGRP-SB1 PE=2 SV=2 | 1 | 1 | 3 | 2 | 0 | 1.584962501 | -1.584962501 |
| Q99323 | MYSN_DROME Myosin heavy chain, non-muscle OS=Drosophila melanogaster OX=7227 GN=zip PE=1 SV=2 | 1 | 1 | 3 | 1 | 0 | 1.584962501 | -1.584962501 |
| Q9V359 | VPS28_DROME Vacuolar protein sorting-associated protein 28 homolog OS=Drosophila melanogaster OX=7227 GN=Vps28 PE=1 SV=1 | 1 | 1 | 3 | 3 | 0 | 1.584962501 | -1.584962501 |

| UniProt ID | Protein description | # PSMs MVB spectra | # of unique peptides MVB | # of PSMs lysate | # of unique peptides lysate | log2(PSM MVB) | log2(PSM lysate) | FC (log2 of MVB PSM-log2 of lysate PSM) |
|---|---|---|---|---|---|---|---|---|
| Q9V521 | PPO2_DROME Phenoloxidase 2 OS=Drosophila melanogaster OX=7227 GN=PPO2 PE=1 SV=1 | 1 | 1 | 3 | 3 | 0 | 1.584962501 | -1.584962501 |
| Q9VAM9 | MESH1_DROME Guanosine-3',5'-bis(diphosphate) 3'-pyrophosphohydrolase MESH1 OS=Drosophila melanogaster OX=7227 GN=Mesh1 PE=1 SV=1 | 1 | 1 | 3 | 2 | 0 | 1.584962501 | -1.584962501 |
| Q9VLU6 | SDHF4_DROME Succinate dehydrogenase assembly factor 4, mitochondrial OS=Drosophila melanogaster OX=7227 GN=Sirup PE=3 SV=2 | 1 | 1 | 3 | 2 | 0 | 1.584962501 | -1.584962501 |
| Q9VNT5 | TRXR2_DROME Thioredoxin reductase 2, mitochondrial OS=Drosophila melanogaster OX=7227 GN=Trxr-2 PE=2 SV=1 | 1 | 1 | 3 | 1 | 0 | 1.584962501 | -1.584962501 |
| Q9VU58 | NPLP2_DROME Neuropeptide-like 2 OS=Drosophila melanogaster OX=7227 GN=Nplp2 PE=1 SV=1 | 1 | 1 | 3 | 2 | 0 | 1.584962501 | -1.584962501 |
| Q9VVN2 | RT26_DROME Probable 28S ribosomal protein S26, mitochondrial OS=Drosophila melanogaster OX=7227 GN=mRpS26 PE=2 SV=1 | 1 | 1 | 3 | 1 | 0 | 1.584962501 | -1.584962501 |
| Q9VWP4 | SUOX_DROME Probable sulfite oxidase, mitochondrial OS=Drosophila melanogaster OX=7227 GN=CG7280 PE=2 SV=1 | 1 | 1 | 3 | 3 | 0 | 1.584962501 | -1.584962501 |
| Q9W552 | VPS26_DROME Vacuolar protein sorting-associated protein 26 OS=Drosophila melanogaster OX=7227 GN=Vps26 PE=2 SV=1 | 1 | 1 | 3 | 3 | 0 | 1.584962501 | -1.584962501 |
| A0A0B4KFE5 | A0A0B4KFE5_DROME NAD-dependent methylenetetrahydrofolate dehydrogenase, isoform C OS=Drosophila melanogaster OX=7227 GN=Nmdmc PE=3 SV=1 | 1 | 1 | 3 | 3 | 0 | 1.584962501 | -1.584962501 |
| A1Z933 | A1Z933_DROME GEO02273p1 OS=Drosophila melanogaster OX=7227 GN=CG33672-RA PE=1 SV=1 | 1 | 1 | 3 | 2 | 0 | 1.584962501 | -1.584962501 |
| B7YZX9 | B7YZX9_DROME Uncharacterized protein, isoform B OS=Drosophila melanogaster OX=7227 GN=38C.49 PE=1 SV=1 | 1 | 1 | 3 | 2 | 0 | 1.584962501 | -1.584962501 |
| H0RNC3 | H0RNC3_DROME FI17124p1 OS=Drosophila melanogaster OX=7227 GN=Rad60 PE=2 SV=1 | 1 | 1 | 3 | 3 | 0 | 1.584962501 | -1.584962501 |
| O77477 | O77477_DROME CG4882, isoform A OS=Drosophila melanogaster OX=7227 GN=CG4882 PE=1 SV=1 | 1 | 1 | 3 | 3 | 0 | 1.584962501 | -1.584962501 |
| Q7JWR4 | Q7JWR4_DROME GH19585p OS=Drosophila melanogaster OX=7227 GN=CG13319 PE=1 SV=1 | 1 | 1 | 3 | 3 | 0 | 1.584962501 | -1.584962501 |
| Q7K1C3 | Q7K1C3_DROME CG6984 OS=Drosophila melanogaster OX=7227 GN=CG6984 PE=1 SV=1 | 1 | 1 | 3 | 3 | 0 | 1.584962501 | -1.584962501 |
| Q7PLE6 | Q7PLE6_DROME Uncharacterized protein, isoform C OS=Drosophila melanogaster OX=7227 GN=CG17691 PE=1 SV=3 | 1 | 1 | 3 | 2 | 0 | 1.584962501 | -1.584962501 |
| Q8MSS7 | Q8MSS7_DROME Mitochondrial ribosomal protein S2 OS=Drosophila melanogaster OX=7227 GN=mRpS2 PE=2 SV=1 | 1 | 1 | 3 | 2 | 0 | 1.584962501 | -1.584962501 |
| Q95RV5 | Q95RV5_DROME Ribulose-phosphate 3-epimerase OS=Drosophila melanogaster OX=7227 GN=CG1364 PE=1 SV=1 | 1 | 1 | 3 | 2 | 0 | 1.584962501 | -1.584962501 |
| Q9V3E7 | Q9V3E7_DROME LD24793p OS=Drosophila melanogaster OX=7227 GN=Ref1 PE=1 SV=1 | 1 | 1 | 3 | 3 | 0 | 1.584962501 | -1.584962501 |
| Q9VA42 | Q9VA42_DROME Niemann-Pick type C-2g, isoform A OS=Drosophila melanogaster OX=7227 GN=Npc2g PE=1 SV=1 | 1 | 1 | 3 | 2 | 0 | 1.584962501 | -1.584962501 |
| Q9VA97 | Q9VA97_DROME Pyridoxal phosphate homeostasis protein OS=Drosophila melanogaster OX=7227 GN=Dmel\CG1983 PE=1 SV=2 | 1 | 1 | 3 | 3 | 0 | 1.584962501 | -1.584962501 |
| Q9VEN6 | Q9VEN6_DROME Mitochondrial ribosomal protein S11 OS=Drosophila melanogaster OX=7227 GN=mRpS11 PE=1 SV=1 | 1 | 1 | 3 | 3 | 0 | 1.584962501 | -1.584962501 |
| Q9VPE1 | Q9VPE1_DROME Uncharacterized protein, isoform C OS=Drosophila melanogaster OX=7227 GN=CG32425 PE=1 SV=2 | 1 | 1 | 3 | 2 | 0 | 1.584962501 | -1.584962501 |
| Q9VWS7 | Q9VWS7_DROME MIP23504p OS=Drosophila melanogaster OX=7227 GN=CG15047-RA PE=2 SV=1 | 1 | 1 | 3 | 2 | 0 | 1.584962501 | -1.584962501 |
| Q9VZZ6 | Q9VZZ6_DROME CG16985 protein OS=Drosophila melanogaster OX=7227 GN=BcDNA:RE30174 PE=1 SV=1 | 1 | 1 | 3 | 2 | 0 | 1.584962501 | -1.584962501 |
| Q9W022 | Q9W022_DROME GEO01508p1 OS=Drosophila melanogaster OX=7227 GN=8993 PE=1 SV=1 | 1 | 1 | 3 | 2 | 0 | 1.584962501 | -1.584962501 |
| Q9W4C3 | Q9W4C3_DROME Ubiquitinyl hydrolase 1 OS=Drosophila melanogaster OX=7227 GN=Usp16-45 PE=1 SV=1 | 1 | 1 | 3 | 3 | 0 | 1.584962501 | -1.584962501 |
| Q9W4Z2 | Q9W4Z2_DROME EG:BACH48C10.6 protein OS=Drosophila melanogaster OX=7227 GN=mRpL14 PE=2 SV=1 | 1 | 1 | 3 | 1 | 0 | 1.584962501 | -1.584962501 |
| Q9W547 | Q9W547_DROME LD21404p OS=Drosophila melanogaster OX=7227 GN=mRpL16 PE=1 SV=1 | 1 | 1 | 3 | 2 | 0 | 1.584962501 | -1.584962501 |
| X2J9A5 | X2J9A5_DROME Uncharacterized protein, isoform B OS=Drosophila melanogaster OX=7227 GN=CG31673 PE=1 SV=1 | 1 | 1 | 3 | 2 | 0 | 1.584962501 | -1.584962501 |
| A1A708 | Y4951_DROME Uncharacterized protein CG4951 OS=Drosophila melanogaster OX=7227 GN=CG4951 PE=1 SV=1 | 1 | 1 | 2 | 2 | 0 | 1 | -1 |
| O77263 | TRMB_DROME tRNA (guanine-N(7)-)-methyltransferase OS=Drosophila melanogaster OX=7227 GN=CG4045 PE=2 SV=1 | 1 | 1 | 2 | 2 | 0 | 1 | -1 |
| Q7K1Z5 | UFC1_DROME Ubiquitin-fold modifier-conjugating enzyme 1 OS=Drosophila melanogaster OX=7227 GN=Ufc1 PE=2 SV=1 | 1 | 1 | 2 | 2 | 0 | 1 | -1 |
| Q7K4B3 | ELP2_DROME Probable elongator complex protein 2 OS=Drosophila melanogaster OX=7227 GN=Elp2 PE=1 SV=1 | 1 | 1 | 2 | 2 | 0 | 1 | -1 |
| Q9V9Z1 | RM32_DROME 39S ribosomal protein L32, mitochondrial OS=Drosophila melanogaster OX=7227 GN=mRpL32 PE=2 SV=1 | 1 | 1 | 2 | 2 | 0 | 1 | -1 |
| Q9VCY3 | CSN6_DROME COP9 signalosome complex subunit 6 OS=Drosophila melanogaster OX=7227 GN=CSN6 PE=1 SV=1 | 1 | 1 | 2 | 2 | 0 | 1 | -1 |
| Q9VXE0 | RUXG_DROME Probable small nuclear ribonucleoprotein G OS=Drosophila melanogaster OX=7227 GN=SNRPG PE=1 SV=1 | 1 | 1 | 2 | 2 | 0 | 1 | -1 |
| Q9W0P5 | GALE_DROME UDP-glucose 4-epimerase OS=Drosophila melanogaster OX=7227 GN=Gale PE=1 SV=1 | 1 | 1 | 2 | 1 | 0 | 1 | -1 |
| Q9W2Y3 | NNRE_DROME NAD(P)H-hydrate epimerase OS=Drosophila melanogaster OX=7227 GN=CG2974 PE=2 SV=2 | 1 | 1 | 2 | 2 | 0 | 1 | -1 |
| A0A0B4KHB7 | A0A0B4KHB7_DROME AP complex subunit sigma OS=Drosophila melanogaster OX=7227 GN=AP-1sigma PE=3 SV=1 | 1 | 1 | 2 | 2 | 0 | 1 | -1 |
| A1Z7X7 | A1Z7X7_DROME Glutathione S transferase T2 OS=Drosophila melanogaster OX=7227 GN=GstT2 PE=1 SV=2 | 1 | 1 | 2 | 2 | 0 | 1 | -1 |
| A1ZAS8 | A1ZAS8_DROME Egg-derived tyrosine phosphatase, isoform C OS=Drosophila melanogaster OX=7227 GN=EDTP PE=4 SV=2 | 1 | 1 | 2 | 2 | 0 | 1 | -1 |
| E1JGK7 | E1JGK7_DROME GEO01604p1 OS=Drosophila melanogaster OX=7227 GN=lolal PE=1 SV=1 | 1 | 1 | 2 | 1 | 0 | 1 | -1 |
| M9PBX0 | M9PBX0_DROME Mothers against decapentaplegic homolog OS=Drosophila melanogaster OX=7227 GN=Mad PE=3 SV=1 | 1 | 1 | 2 | 1 | 0 | 1 | -1 |
| M9PE30 | M9PE30_DROME Transgelin OS=Drosophila melanogaster OX=7227 GN=Chd64 PE=1 SV=1 | 1 | 1 | 2 | 1 | 0 | 1 | -1 |
| O18645 | O18645_DROME Non-specific serine/threonine protein kinase OS=Drosophila melanogaster OX=7227 GN=AMPKalpha PE=1 SV=1 | 1 | 1 | 2 | 1 | 0 | 1 | -1 |
| Q7JWX3 | Q7JWX3_DROME GH10112p OS=Drosophila melanogaster OX=7227 GN=nec PE=1 SV=1 | 1 | 1 | 2 | 2 | 0 | 1 | -1 |
| Q7KN85 | Q7KN85_DROME ATP-citrate synthase OS=Drosophila melanogaster OX=7227 GN=ATPCL PE=1 SV=1 | 1 | 1 | 2 | 1 | 0 | 1 | -1 |
| Q8I0J3 | Q8I0J3_DROME Mitochondrial ribosomal protein S21, isoform A OS=Drosophila melanogaster OX=7227 GN=mRpS21 PE=1 SV=1 | 1 | 1 | 2 | 2 | 0 | 1 | -1 |
| Q8SXX1 | Q8SXX1_DROME Phosphatidylinositol 5-phosphate 4-kinase, isoform A OS=Drosophila melanogaster OX=7227 GN=PIP4K PE=1 SV=1 | 1 | 1 | 2 | 1 | 0 | 1 | -1 |
| Q9VBU4 | Q9VBU4_DROME Peptidyl-prolyl cis-trans isomerase OS=Drosophila melanogaster OX=7227 GN=BcDNA:RH66629 PE=1 SV=1 | 1 | 1 | 2 | 2 | 0 | 1 | -1 |
| Q9VCY1 | Q9VCY1_DROME Uncharacterized protein OS=Drosophila melanogaster OX=7227 GN=CG6937 PE=1 SV=1 | 1 | 1 | 2 | 2 | 0 | 1 | -1 |
| Q9VD14 | Q9VD14_DROME GH07286p OS=Drosophila melanogaster OX=7227 GN=Dmel\CG13850 PE=1 SV=3 | 1 | 1 | 2 | 2 | 0 | 1 | -1 |
| Q9VFN9 | Q9VFN9_DROME GEO07882p1 OS=Drosophila melanogaster OX=7227 GN=BcDNA:RE55111 PE=1 SV=1 | 1 | 1 | 2 | 1 | 0 | 1 | -1 |
| Q9VG59 | Q9VG59_DROME FI07645p OS=Drosophila melanogaster OX=7227 GN=CG5608-RA PE=1 SV=1 | 1 | 1 | 2 | 2 | 0 | 1 | -1 |
| Q9VGV9 | Q9VGV9_DROME GH11067p OS=Drosophila melanogaster OX=7227 GN=Dmel\CG6567 PE=1 SV=1 | 1 | 1 | 2 | 1 | 0 | 1 | -1 |
| Q9VHC0 | Q9VHC0_DROME LD23870p OS=Drosophila melanogaster OX=7227 GN=RnpS1 PE=1 SV=2 | 1 | 1 | 2 | 1 | 0 | 1 | -1 |
| Q9VHF4 | Q9VHF4_DROME RE70963p OS=Drosophila melanogaster OX=7227 GN=BcDNA:RE70963 PE=2 SV=2 | 1 | 1 | 2 | 2 | 0 | 1 | -1 |
| P04388 | RAS2_DROME Ras-like protein 2 OS=Drosophila melanogaster OX=7227 GN=Ras64B PE=1 SV=2 | 1 | 1 | 1 | 1 | 0 | 0 | 0 |
| Q9VHG1 | Q9VHG1_DROME RE66051p OS=Drosophila melanogaster OX=7227 GN=Vps16A PE=1 SV=1 | 1 | 1 | 2 | 2 | 0 | 1 | -1 |
| P07192 | MAL3_DROME Maltase A3 OS=Drosophila melanogaster OX=7227 GN=Mal-A3 PE=2 SV=2 | 1 | 1 | 1 | 1 | 0 | 0 | 0 |
| Q9VJE3 | Q9VJE3_DROME LD24839p OS=Drosophila melanogaster OX=7227 GN=BEST:LD24839 PE=1 SV=1 | 1 | 1 | 2 | 2 | 0 | 1 | -1 |
| Q9VKC5 | Q9VKC5_DROME RH17958p OS=Drosophila melanogaster OX=7227 GN=CG6770-RA PE=1 SV=1 | 1 | 1 | 2 | 1 | 0 | 1 | -1 |
| Q27297 | RAD51_DROME DNA repair protein Rad51 homolog OS=Drosophila melanogaster OX=7227 GN=spn-A PE=2 SV=1 | 1 | 1 | 1 | 1 | 0 | 0 | 0 |
| Q9VPX5 | Q9VPX5_DROME Vacuolar protein sorting-associated protein 29 OS=Drosophila melanogaster OX=7227 GN=Vps29 PE=1 SV=1 | 1 | 1 | 2 | 2 | 0 | 1 | -1 |
| Q9NBK5 | TRC_DROME Serine/threonine-protein kinase tricorner OS=Drosophila melanogaster OX=7227 GN=trc PE=1 SV=1 | 1 | 1 | 1 | 1 | 0 | 0 | 0 |
| Q9V8I2 | LTOR2_DROME Ragulator complex protein LAMTOR2 homolog OS=Drosophila melanogaster OX=7227 GN=CG5189 PE=2 SV=1 | 1 | 1 | 1 | 1 | 0 | 0 | 0 |
| Q9VAU9 | ZN330_DROME Zinc finger protein 330 homolog OS=Drosophila melanogaster OX=7227 GN=Noa36 PE=2 SV=1 | 1 | 1 | 1 | 1 | 0 | 0 | 0 |
| Q9VVX4 | Q9VVX4_DROME Mitochondrial ribosomal protein L21, isoform A OS=Drosophila melanogaster OX=7227 GN=mRpL21 PE=1 SV=2 | 1 | 1 | 2 | 2 | 0 | 1 | -1 |
| Q9VW70 | Q9VW70_DROME FI21813p1 OS=Drosophila melanogaster OX=7227 GN=AAF49079 PE=1 SV=2 | 1 | 1 | 2 | 2 | 0 | 1 | -1 |
| Q9VK68 | PITH1_DROME PITH domain-containing protein CG6153 OS=Drosophila melanogaster OX=7227 GN=CG6153 PE=2 SV=1 | 1 | 1 | 1 | 1 | 0 | 0 | 0 |
| Q9VKN7 | AURKB_DROME Aurora kinase B OS=Drosophila melanogaster OX=7227 GN=ial PE=1 SV=1 | 1 | 1 | 1 | 1 | 0 | 0 | 0 |
| Q9VX15 | Q9VX15_DROME LD35209p OS=Drosophila melanogaster OX=7227 GN=DmRFC2 PE=1 SV=2 | 1 | 1 | 2 | 2 | 0 | 1 | -1 |
| Q9VMQ5 | COQ6_DROME Ubiquinone biosynthesis monooxygenase COQ6, mitochondrial OS=Drosophila melanogaster OX=7227 GN=Coq6 PE=2 SV=1 | 1 | 1 | 1 | 1 | 0 | 0 | 0 |
| Q6NLJ9 | Q6NLJ9_DROME AT19138p OS=Drosophila melanogaster OX=7227 GN=CG30109 PE=1 SV=1 | 1 | 1 | 1 | 1 | 0 | 0 | 0 |
| Q7JVI6 | Q7JVI6_DROME Glutathione S transferase E13, isoform A OS=Drosophila melanogaster OX=7227 GN=GstE13 PE=1 SV=1 | 1 | 1 | 1 | 1 | 0 | 0 | 0 |
| Q7K4C7 | Q7K4C7_DROME LD46058p OS=Drosophila melanogaster OX=7227 GN=Pcyt1 PE=1 SV=1 | 1 | 1 | 1 | 1 | 0 | 0 | 0 |
| Q7K4J7 | Q7K4J7_DROME LD36653p OS=Drosophila melanogaster OX=7227 GN=CG17985 PE=2 SV=1 | 1 | 1 | 1 | 1 | 0 | 0 | 0 |
| Q8IMX8 | Q8IMX8_DROME U6 snRNA-associated Sm-like protein LSm3 OS=Drosophila melanogaster OX=7227 GN=LSm3 PE=1 SV=1 | 1 | 1 | 1 | 1 | 0 | 0 | 0 |
| Q94533 | Q94533_DROME Ribosomal protein S6 kinase OS=Drosophila melanogaster OX=7227 GN=S6k PE=1 SV=2 | 1 | 1 | 1 | 1 | 0 | 0 | 0 |
| Q9VCF8 | Q9VCF8_DROME LD23561p OS=Drosophila melanogaster OX=7227 GN=Dmel\CG5854 PE=1 SV=1 | 1 | 1 | 1 | 1 | 0 | 0 | 0 |

| UniProt ID | Protein description | # PSMs MVB spectra | # of unique peptides MVB | # of PSMs lysate | # of unique peptides lysate | log2(PSM MVB) | log2(PSM lysate) | FC (log2 of MVB PSM-log2 of lysate PSM) |
|---|---|---|---|---|---|---|---|---|
| Q9VES8 | Q9VES8_DROME RE28913p OS=Drosophila melanogaster OX=7227 GN=Dmel\CG10340 PE=1 SV=3 | 1 | 1 | 1 | 1 | 0 | 0 | 0 |
| Q9VI15 | Q9VI15_DROME GH12023p OS=Drosophila melanogaster OX=7227 GN=Dmel\CG10286 PE=1 SV=1 | 1 | 1 | 1 | 1 | 0 | 0 | 0 |
| Q9VJ28 | Q9VJ28_DROME FI05204p OS=Drosophila melanogaster OX=7227 GN=L2HGDH PE=1 SV=1 | 1 | 1 | 1 | 1 | 0 | 0 | 0 |
| Q9VRA0 | Q9VRA0_DROME Uncharacterized protein, isoform A OS=Drosophila melanogaster OX=7227 GN=Dmel\CG15456 PE=4 SV=1 | 1 | 1 | 1 | 1 | 0 | 0 | 0 |
| Q9VZJ4 | Q9VZJ4_DROME Uncharacterized protein, isoform B OS=Drosophila melanogaster OX=7227 GN=Dmel\CG11593 PE=4 SV=3 | 1 | 1 | 1 | 1 | 0 | 0 | 0 |
| Q9VZQ4 | Q9VZQ4_DROME Dro6 protein OS=Drosophila melanogaster OX=7227 GN=Drsl6 PE=2 SV=1 | 1 | 1 | 1 | 1 | 0 | 0 | 0 |
| Q9W0P4 | Q9W0P4_DROME GEO05053p1 OS=Drosophila melanogaster OX=7227 GN=BcDNA:LD06317 PE=1 SV=1 | 1 | 1 | 1 | 1 | 0 | 0 | 0 |
| Q9W5E7 | Q9W5E7_DROME LP07417p OS=Drosophila melanogaster OX=7227 GN=BcDNA:LP07417 PE=2 SV=1 | 1 | 1 | 1 | 1 | 0 | 0 | 0 |
| O44386 | ITA3_DROME Integrin alpha-PS3 OS=Drosophila melanogaster OX=7227 GN=scb PE=1 SV=2 | 1 | 1 | 1 | 1 | 0 | 0 | 0 |
| O76902 | PKHF1_DROME Pleckstrin homology domain-containing family F member 1 homolog OS=Drosophila melanogaster OX=7227 GN=rush PE=1 SV=1 | 1 | 1 | 1 | 1 | 0 | 0 | 0 |
| P04412 | EGFR_DROME Epidermal growth factor receptor OS=Drosophila melanogaster OX=7227 GN=Egfr PE=1 SV=3 | 1 | 1 | 1 | 1 | 0 | 0 | 0 |
| P07190 | MAL1_DROME Maltase A1 OS=Drosophila melanogaster OX=7227 GN=Mal-A1 PE=2 SV=2 | 1 | 1 | 1 | 1 | 0 | 0 | 0 |
| P09491 | TPM2_DROME Tropomyosin-2 OS=Drosophila melanogaster OX=7227 GN=Tm2 PE=2 SV=1 | 1 | 1 | 1 | 1 | 0 | 0 | 0 |
| P12080 | ITA2_DROME Integrin alpha-PS2 OS=Drosophila melanogaster OX=7227 GN=if PE=1 SV=2 | 1 | 1 | 1 | 1 | 0 | 0 | 0 |
| P18432 | MLR_DROME Myosin regulatory light chain 2 OS=Drosophila melanogaster OX=7227 GN=Mlc2 PE=1 SV=2 | 1 | 1 | 1 | 1 | 0 | 0 | 0 |
| P18929 | NU1M_DROME NADH-ubiquinone oxidoreductase chain 1 OS=Drosophila melanogaster OX=7227 GN=mt:ND1 PE=3 SV=3 | 1 | 1 | 1 | 1 | 0 | 0 | 0 |
| P18932 | NU5M_DROME NADH-ubiquinone oxidoreductase chain 5 OS=Drosophila melanogaster OX=7227 GN=mt:ND5 PE=3 SV=4 | 1 | 1 | 1 | 1 | 0 | 0 | 0 |
| P35004 | TRYB_DROME Trypsin beta OS=Drosophila melanogaster OX=7227 GN=betaTry PE=2 SV=2 | 1 | 1 | 1 | 1 | 0 | 0 | 0 |
| P35005 | TRYE_DROME Trypsin epsilon OS=Drosophila melanogaster OX=7227 GN=epsilonTry PE=2 SV=1 | 1 | 1 | 1 | 1 | 0 | 0 | 0 |
| P42519 | STAR_DROME Protein Star OS=Drosophila melanogaster OX=7227 GN=S PE=2 SV=1 | 1 | 1 | 1 | 1 | 0 | 0 | 0 |
| P83971 | LYSA_DROME Lysozyme A/C OS=Drosophila melanogaster OX=7227 GN=LysA PE=2 SV=1 | 1 | 1 | 1 | 1 | 0 | 0 | 0 |
| Q00174 | LAMA_DROME Laminin subunit alpha OS=Drosophila melanogaster OX=7227 GN=LanA PE=1 SV=2 | 1 | 1 | 1 | 1 | 0 | 0 | 0 |
| Q0KHQ5 | TAO_DROME Serine/threonine-protein kinase Tao OS=Drosophila melanogaster OX=7227 GN=Tao PE=1 SV=1 | 1 | 1 | 1 | 1 | 0 | 0 | 0 |
| Q24155 | TRUNK_DROME Protein trunk OS=Drosophila melanogaster OX=7227 GN=trk PE=3 SV=1 | 1 | 1 | 1 | 1 | 0 | 0 | 0 |
| Q24192 | RHOL_DROME Ras-like GTP-binding protein RhoL OS=Drosophila melanogaster OX=7227 GN=RhoL PE=2 SV=1 | 1 | 1 | 1 | 1 | 0 | 0 | 0 |
| Q7JWG9 | RM52_DROME 39S ribosomal protein L52, mitochondrial OS=Drosophila melanogaster OX=7227 GN=mRpL52 PE=2 SV=1 | 1 | 1 | 1 | 1 | 0 | 0 | 0 |
| Q8MR45 | ASPG1_DROME Putative N(4)-(beta-N-acetylglucosaminyl)-L-asparaginase CG1827 OS=Drosophila melanogaster OX=7227 GN=CG1827 PE=2 SV=1 | 1 | 1 | 1 | 1 | 0 | 0 | 0 |
| Q8MRQ4 | LMBD2_DROME LMBR1 domain-containing protein 2 homolog OS=Drosophila melanogaster OX=7227 GN=CG8135 PE=2 SV=2 | 1 | 1 | 1 | 1 | 0 | 0 | 0 |
| Q8SZ16 | SDHF3_DROME Succinate dehydrogenase assembly factor 3, mitochondrial OS=Drosophila melanogaster OX=7227 GN=Sdhaf3 PE=2 SV=1 | 1 | 1 | 1 | 1 | 0 | 0 | 0 |
| Q95T10 | XYLK_DROME Glycosaminoglycan xylosylkinase homolog OS=Drosophila melanogaster OX=7227 GN=CG3631 PE=2 SV=1 | 1 | 1 | 1 | 1 | 0 | 0 | 0 |
| Q9U6B8 | CRCM1_DROME Calcium release-activated calcium channel protein 1 OS=Drosophila melanogaster OX=7227 GN=olf186-F PE=1 SV=1 | 1 | 1 | 1 | 1 | 0 | 0 | 0 |
| Q9V3R8 | UBIA1_DROME UbiA prenyltransferase domain-containing protein 1 homolog OS=Drosophila melanogaster OX=7227 GN=heix PE=2 SV=1 | 1 | 1 | 1 | 1 | 0 | 0 | 0 |
| Q9VA73 | CMC_DROME Calcium-binding mitochondrial carrier protein Aralar1 OS=Drosophila melanogaster OX=7227 GN=aralar1 PE=2 SV=1 | 1 | 1 | 1 | 1 | 0 | 0 | 0 |
| Q9VAI1 | CIA30_DROME Complex I intermediate-associated protein 30, mitochondrial OS=Drosophila melanogaster OX=7227 GN=CIA30 PE=2 SV=1 | 1 | 1 | 1 | 1 | 0 | 0 | 0 |
| Q9VCI5 | DHSD_DROME Succinate dehydrogenase [ubiquinone] cytochrome b small subunit, mitochondrial OS=Drosophila melanogaster OX=7227 GN=SdhD PE=2 SV=1 | 1 | 1 | 1 | 1 | 0 | 0 | 0 |
| Q9VEB4 | MTEF5_DROME Transcription termination factor 5, mitochondrial OS=Drosophila melanogaster OX=7227 GN=mTerf5 PE=1 SV=1 | 1 | 1 | 1 | 1 | 0 | 0 | 0 |
| Q9VEG9 | NRM_DROME Nurim homolog OS=Drosophila melanogaster OX=7227 GN=nrm PE=2 SV=1 | 1 | 1 | 1 | 1 | 0 | 0 | 0 |
| Q9VEX9 | SAP18_DROME Histone deacetylase complex subunit SAP18 OS=Drosophila melanogaster OX=7227 GN=Bin1 PE=1 SV=1 | 1 | 1 | 1 | 1 | 0 | 0 | 0 |
| Q9VM51 | MME1_DROME Mitochondrial magnesium exporter 1 OS=Drosophila melanogaster OX=7227 GN=MME1 PE=2 SV=1 | 1 | 1 | 1 | 1 | 0 | 0 | 0 |
| Q9VM64 | IHOG_DROME Interference hedgehog OS=Drosophila melanogaster OX=7227 GN=ihog PE=1 SV=1 | 1 | 1 | 1 | 1 | 0 | 0 | 0 |
| Q9VMC8 | PHF5A_DROME PHD finger-like domain-containing protein 5A OS=Drosophila melanogaster OX=7227 GN=Phf5a PE=3 SV=2 | 1 | 1 | 1 | 1 | 0 | 0 | 0 |
| Q9VQG2 | APH1_DROME Gamma-secretase subunit Aph-1 OS=Drosophila melanogaster OX=7227 GN=aph-1 PE=1 SV=1 | 1 | 1 | 1 | 1 | 0 | 0 | 0 |
| Q9VS46 | RINT1_DROME RINT1-like protein OS=Drosophila melanogaster OX=7227 GN=Rint1 PE=2 SV=2 | 1 | 1 | 1 | 1 | 0 | 0 | 0 |
| Q9VTJ8 | TIM14_DROME Mitochondrial import inner membrane translocase subunit TIM14 OS=Drosophila melanogaster OX=7227 GN=Tim14 PE=3 SV=2 | 1 | 1 | 1 | 1 | 0 | 0 | 0 |
| Q9W0G7 | MSD1_DROME Augmin complex subunit msd1 OS=Drosophila melanogaster OX=7227 GN=msd1 PE=1 SV=2 | 1 | 1 | 1 | 1 | 0 | 0 | 0 |
| Q9W145 | STRT1_DROME Steroidogenic acute regulatory protein-like OS=Drosophila melanogaster OX=7227 GN=Start1 PE=1 SV=2 | 1 | 1 | 1 | 1 | 0 | 0 | 0 |
| Q9W422 | JOSL_DROME Josephin-like protein OS=Drosophila melanogaster OX=7227 GN=CG3781 PE=2 SV=3 | 1 | 1 | 1 | 1 | 0 | 0 | 0 |
| Q9W429 | S35B4_DROME UDP-xylose and UDP-N-acetylglucosamine transporter OS=Drosophila melanogaster OX=7227 GN=Efr PE=2 SV=1 | 1 | 1 | 1 | 1 | 0 | 0 | 0 |
| A0A0B4KEN4 | A0A0B4KEN4_DROME Vang-like protein OS=Drosophila melanogaster OX=7227 GN=Vang PE=3 SV=1 | 1 | 1 | 1 | 1 | 0 | 0 | 0 |
| A0A0B4KEW2 | A0A0B4KEW2_DROME Uncharacterized protein, isoform E OS=Drosophila melanogaster OX=7227 GN=CG30389 PE=4 SV=1 | 1 | 1 | 1 | 1 | 0 | 0 | 0 |
| A0A0B4KEW6 | A0A0B4KEW6_DROME Amphiphysin, isoform B OS=Drosophila melanogaster OX=7227 GN=Amph PE=1 SV=1 | 1 | 1 | 1 | 1 | 0 | 0 | 0 |
| A0A0B4KFH1 | A0A0B4KFH1_DROME Muskelin, isoform C OS=Drosophila melanogaster OX=7227 GN=muskelin PE=4 SV=1 | 1 | 1 | 1 | 1 | 0 | 0 | 0 |
| A0A0B4KFH6 | A0A0B4KFH6_DROME Protein SYS1 homolog OS=Drosophila melanogaster OX=7227 GN=Dmel\CG11753 PE=3 SV=1 | 1 | 1 | 1 | 1 | 0 | 0 | 0 |
| A0A0B4LEV0 | A0A0B4LEV0_DROME CG8321, isoform B OS=Drosophila melanogaster OX=7227 GN=CG8321 PE=4 SV=1 | 1 | 1 | 1 | 1 | 0 | 0 | 0 |
| A0A0B4LEW9 | A0A0B4LEW9_DROME Cyp6a14, isoform E (Fragment) OS=Drosophila melanogaster OX=7227 GN=Cyp6a14 PE=1 SV=1 | 1 | 1 | 1 | 1 | 0 | 0 | 0 |
| A0A0B4LF75 | A0A0B4LF75_DROME Off-track, isoform B OS=Drosophila melanogaster OX=7227 GN=otk PE=3 SV=1 | 1 | 1 | 1 | 1 | 0 | 0 | 0 |
| A0A0B4LFX4 | A0A0B4LFX4_DROME Coracle, isoform F OS=Drosophila melanogaster OX=7227 GN=cora PE=1 SV=1 | 1 | 1 | 1 | 1 | 0 | 0 | 0 |
| A0A0B4LGS0 | A0A0B4LGS0_DROME Alpha-amylase OS=Drosophila melanogaster OX=7227 GN=Amy-p PE=3 SV=1 | 1 | 1 | 1 | 1 | 0 | 0 | 0 |
| A0A0C4DHC4 | A0A0C4DHC4_DROME Uncharacterized protein, isoform G OS=Drosophila melanogaster OX=7227 GN=SLO2 PE=4 SV=1 | 1 | 1 | 1 | 1 | 0 | 0 | 0 |
| A0A126GUM4 | A0A126GUM4_DROME Ran-binding protein M, isoform G OS=Drosophila melanogaster OX=7227 GN=RanBPM PE=1 SV=1 | 1 | 1 | 1 | 1 | 0 | 0 | 0 |
| A1Z6G4 | A1Z6G4_DROME RE41154p1 OS=Drosophila melanogaster OX=7227 GN=CG17508-RA PE=1 SV=1 | 1 | 1 | 1 | 1 | 0 | 0 | 0 |
| A1Z8I8 | A1Z8I8_DROME Uncharacterized protein, isoform E OS=Drosophila melanogaster OX=7227 GN=CG42336 PE=1 SV=2 | 1 | 1 | 1 | 1 | 0 | 0 | 0 |
| A1Z9M0 | A1Z9M0_DROME Uncharacterized protein, isoform B OS=Drosophila melanogaster OX=7227 GN=CG18324 PE=3 SV=1 | 1 | 1 | 1 | 1 | 0 | 0 | 0 |
| A1ZAB1 | A1ZAB1_DROME Lambik OS=Drosophila melanogaster OX=7227 GN=lbk PE=1 SV=2 | 1 | 1 | 1 | 1 | 0 | 0 | 0 |
| A1ZAG1 | A1ZAG1_DROME CG4398 OS=Drosophila melanogaster OX=7227 GN=jtb PE=4 SV=1 | 1 | 1 | 1 | 1 | 0 | 0 | 0 |
| A1ZB72 | A1ZB72_DROME Glutathione S transferase E7 OS=Drosophila melanogaster OX=7227 GN=GstE7 PE=1 SV=1 | 1 | 1 | 1 | 1 | 0 | 0 | 0 |
| A1ZBK1 | A1ZBK1_DROME FI23922p1 OS=Drosophila melanogaster OX=7227 GN=CG10051-RA PE=2 SV=1 | 1 | 1 | 1 | 1 | 0 | 0 | 0 |
| A2VER2 | A2VER2_DROME IP18228p OS=Drosophila melanogaster OX=7227 GN=Dmel\CG42495 PE=2 SV=1 | 1 | 1 | 1 | 1 | 0 | 0 | 0 |
| A8DY79 | A8DY79_DROME Bruchpilot, isoform G OS=Drosophila melanogaster OX=7227 GN=brp PE=1 SV=2 | 1 | 1 | 1 | 1 | 0 | 0 | 0 |
| A8JQU1 | A8JQU1_DROME Uncharacterized protein, isoform C OS=Drosophila melanogaster OX=7227 GN=CG1208 PE=3 SV=1 | 1 | 1 | 1 | 1 | 0 | 0 | 0 |
| A8JRD2 | A8JRD2_DROME Lipophorin receptor 1, isoform S OS=Drosophila melanogaster OX=7227 GN=LpR1 PE=4 SV=3 | 1 | 1 | 1 | 1 | 0 | 0 | 0 |
| A8JTM7 | A8JTM7_DROME Megalin, isoform A OS=Drosophila melanogaster OX=7227 GN=mgl PE=1 SV=2 | 1 | 1 | 1 | 1 | 0 | 0 | 0 |
| C9QP21 | C9QP21_DROME Carboxylic ester hydrolase OS=Drosophila melanogaster OX=7227 GN=Est-6 PE=1 SV=1 | 1 | 1 | 1 | 1 | 0 | 0 | 0 |
| E1JGK8 | E1JGK8_DROME Uncharacterized protein, isoform A OS=Drosophila melanogaster OX=7227 GN=CG42518 PE=4 SV=1 | 1 | 1 | 1 | 1 | 0 | 0 | 0 |
| E1JIX4 | E1JIX4_DROME Uncharacterized protein OS=Drosophila melanogaster OX=7227 GN=CG42488 PE=4 SV=1 | 1 | 1 | 1 | 1 | 0 | 0 | 0 |
| E1JJF2 | E1JJF2_DROME Uncharacterized protein, isoform B OS=Drosophila melanogaster OX=7227 GN=CG4558 PE=4 SV=1 | 1 | 1 | 1 | 1 | 0 | 0 | 0 |
| F9VMG5 | F9VMG5_DROME GEO02462p1 OS=Drosophila melanogaster OX=7227 GN=CG31808-RC PE=2 SV=1 | 1 | 1 | 1 | 1 | 0 | 0 | 0 |
| L0MLM5 | L0MLM5_DROME Uncharacterized protein, isoform B OS=Drosophila melanogaster OX=7227 GN=CG32850 PE=4 SV=1 | 1 | 1 | 1 | 1 | 0 | 0 | 0 |
| M9NDD5 | M9NDD5_DROME Alpha/beta hydrolase2, isoform B OS=Drosophila melanogaster OX=7227 GN=Hydr2 PE=4 SV=1 | 1 | 1 | 1 | 1 | 0 | 0 | 0 |
| M9NDV5 | M9NDV5_DROME Uncharacterized protein, isoform G OS=Drosophila melanogaster OX=7227 GN=CG17715 PE=1 SV=1 | 1 | 1 | 1 | 1 | 0 | 0 | 0 |
| M9PB37 | M9PB37_DROME Uncharacterized protein, isoform F OS=Drosophila melanogaster OX=7227 GN=CG34126 PE=4 SV=1 | 1 | 1 | 1 | 1 | 0 | 0 | 0 |

| UniProt ID | Protein description | # PSMs MVB spectra | # of unique peptides MVB | # of PSMs lysate | # of unique peptides lysate | log2(PSM MVB) | log2(PSM lysate) | FC (log2 of MVB PSM-log2 of lysate PSM) |
|---|---|---|---|---|---|---|---|---|
| M9PC46 | M9PC46_DROME Roundabout 2, isoform B OS=Drosophila melanogaster OX=7227 GN=robo2 PE=4 SV=1 | 1 | 1 | 1 | 1 | 0 | 0 | 0 |
| M9PD12 | M9PD12_DROME Spichthyin, isoform B OS=Drosophila melanogaster OX=7227 GN=spict PE=4 SV=1 | 1 | 1 | 1 | 1 | 0 | 0 | 0 |
| M9PD31 | M9PD31_DROME Uncharacterized protein, isoform B OS=Drosophila melanogaster OX=7227 GN=CG5262 PE=4 SV=1 | 1 | 1 | 1 | 1 | 0 | 0 | 0 |
| M9PEA1 | M9PEA1_DROME Uncharacterized protein, isoform C OS=Drosophila melanogaster OX=7227 GN=CG18586 PE=4 SV=1 | 1 | 1 | 1 | 1 | 0 | 0 | 0 |
| M9PEM7 | M9PEM7_DROME Liquid facets, isoform G OS=Drosophila melanogaster OX=7227 GN=lqf PE=1 SV=1 | 1 | 1 | 1 | 1 | 0 | 0 | 0 |
| M9PES4 | M9PES4_DROME Defective proboscis extension response 6, isoform D OS=Drosophila melanogaster OX=7227 GN=dpr6 PE=4 SV=1 | 1 | 1 | 1 | 1 | 0 | 0 | 0 |
| M9PFF1 | M9PFF1_DROME ADP-ribosylation factor GTPase activating protein 1, isoform B OS=Drosophila melanogaster OX=7227 GN=ArfGAP1 PE=1 SV=1 | 1 | 1 | 1 | 1 | 0 | 0 | 0 |
| M9PFK5 | M9PFK5_DROME Uncharacterized protein, isoform B OS=Drosophila melanogaster OX=7227 GN=CG7510 PE=4 SV=1 | 1 | 1 | 1 | 1 | 0 | 0 | 0 |
| M9PHV9 | M9PHV9_DROME XK-related protein OS=Drosophila melanogaster OX=7227 GN=CG32579 PE=3 SV=1 | 1 | 1 | 1 | 1 | 0 | 0 | 0 |
| O76462 | O76462_DROME Bem46 OS=Drosophila melanogaster OX=7227 GN=Bem46 PE=2 SV=1 | 1 | 1 | 1 | 1 | 0 | 0 | 0 |
| O77262 | O77262_DROME EG:22E5.9 protein OS=Drosophila melanogaster OX=7227 GN=Actn PE=2 SV=1 | 1 | 1 | 1 | 1 | 0 | 0 | 0 |
| Q0E9A7 | Q0E9A7_DROME Uncharacterized protein, isoform E OS=Drosophila melanogaster OX=7227 GN=CG30054 PE=4 SV=3 | 1 | 1 | 1 | 1 | 0 | 0 | 0 |
| Q400M8 | Q400M8_DROME Meltrin, isoform D OS=Drosophila melanogaster OX=7227 GN=Meltrin PE=1 SV=2 | 1 | 1 | 1 | 1 | 0 | 0 | 0 |
| Q4QQ12 | Q4QQ12_DROME IP06605p OS=Drosophila melanogaster OX=7227 GN=CG11110 PE=2 SV=1 | 1 | 1 | 1 | 1 | 0 | 0 | 0 |
| Q60GL7 | Q60GL7_DROME Alpha1,3-mannose beta1,2-N-acetylglucosaminyltransferase 1 OS=Drosophila melanogaster OX=7227 GN=Mgat1 PE=2 SV=1 | 1 | 1 | 1 | 1 | 0 | 0 | 0 |
| Q6NLL3 | Q6NLL3_DROME GH21943p OS=Drosophila melanogaster OX=7227 GN=Dmel\CG11537 PE=2 SV=1 | 1 | 1 | 1 | 1 | 0 | 0 | 0 |
| Q6NR72 | Q6NR72_DROME CG4019, isoform A OS=Drosophila melanogaster OX=7227 GN=Eglp4 PE=1 SV=1 | 1 | 1 | 1 | 1 | 0 | 0 | 0 |
| Q7JUM3 | Q7JUM3_DROME CG6401, isoform A OS=Drosophila melanogaster OX=7227 GN=PIG-A PE=2 SV=1 | 1 | 1 | 1 | 1 | 0 | 0 | 0 |
| Q7JUN3 | Q7JUN3_DROME CG2316, isoform A OS=Drosophila melanogaster OX=7227 GN=Dmel\CG2316 PE=2 SV=1 | 1 | 1 | 1 | 1 | 0 | 0 | 0 |
| Q7JV69 | Q7JV69_DROME SD11922p OS=Drosophila melanogaster OX=7227 GN=Spn55B PE=1 SV=1 | 1 | 1 | 1 | 1 | 0 | 0 | 0 |
| Q7JXB9 | Q7JXB9_DROME Endonuclease G OS=Drosophila melanogaster OX=7227 GN=EndoG PE=1 SV=1 | 1 | 1 | 1 | 1 | 0 | 0 | 0 |
| Q7JXC6 | Q7JXC6_DROME CG8237 OS=Drosophila melanogaster OX=7227 GN=CG8237 PE=2 SV=1 | 1 | 1 | 1 | 1 | 0 | 0 | 0 |
| Q7JY99 | Q7JY99_DROME CG8298, isoform A OS=Drosophila melanogaster OX=7227 GN=CG8298 PE=2 SV=1 | 1 | 1 | 1 | 1 | 0 | 0 | 0 |
| Q7JZY1 | Q7JZY1_DROME RE06389p OS=Drosophila melanogaster OX=7227 GN=Dmel\CG2310 PE=1 SV=1 | 1 | 1 | 1 | 1 | 0 | 0 | 0 |
| Q7K036 | Q7K036_DROME AT23266p OS=Drosophila melanogaster OX=7227 GN=Tom7 PE=2 SV=1 | 1 | 1 | 1 | 1 | 0 | 0 | 0 |
| Q7K0L7 | Q7K0L7_DROME LP01207p OS=Drosophila melanogaster OX=7227 GN=Tpc1 PE=1 SV=1 | 1 | 1 | 1 | 1 | 0 | 0 | 0 |
| Q7K188 | Q7K188_DROME CG6329, isoform A OS=Drosophila melanogaster OX=7227 GN=CG6329 PE=2 SV=1 | 1 | 1 | 1 | 1 | 0 | 0 | 0 |
| Q7K275 | Q7K275_DROME LD15563p OS=Drosophila melanogaster OX=7227 GN=link PE=2 SV=1 | 1 | 1 | 1 | 1 | 0 | 0 | 0 |
| Q7K3N4 | Q7K3N4_DROME CG8888 OS=Drosophila melanogaster OX=7227 GN=CG8888 PE=1 SV=1 | 1 | 1 | 1 | 1 | 0 | 0 | 0 |
| Q7K483 | Q7K483_DROME SD08549p OS=Drosophila melanogaster OX=7227 GN=Dmel\CG7943 PE=2 SV=1 | 1 | 1 | 1 | 1 | 0 | 0 | 0 |
| Q7K5M6 | Q7K5M6_DROME GH04176p OS=Drosophila melanogaster OX=7227 GN=Sip1 PE=1 SV=1 | 1 | 1 | 1 | 1 | 0 | 0 | 0 |
| Q7KSB5 | Q7KSB5_DROME S-formylglutathione hydrolase OS=Drosophila melanogaster OX=7227 GN=CG4390 PE=1 SV=1 | 1 | 1 | 1 | 1 | 0 | 0 | 0 |
| Q7KT70 | Q7KT70_DROME FI18641p1 OS=Drosophila melanogaster OX=7227 GN=Gli PE=2 SV=1 | 1 | 1 | 1 | 1 | 0 | 0 | 0 |
| Q7KTD2 | Q7KTD2_DROME SR family splicing factor SC35, isoform A OS=Drosophila melanogaster OX=7227 GN=SC35 PE=1 SV=1 | 1 | 1 | 1 | 1 | 0 | 0 | 0 |
| Q7PL91 | Q7PL91_DROME GEO11417p1 OS=Drosophila melanogaster OX=7227 GN=ND-AGGG PE=2 SV=1 | 1 | 1 | 1 | 1 | 0 | 0 | 0 |
| Q7YTZ4 | Q7YTZ4_DROME GH22187p OS=Drosophila melanogaster OX=7227 GN=CG10737 PE=1 SV=1 | 1 | 1 | 1 | 1 | 0 | 0 | 0 |
| Q86P36 | Q86P36_DROME Patched-related, isoform A OS=Drosophila melanogaster OX=7227 GN=Ptr PE=2 SV=1 | 1 | 1 | 1 | 1 | 0 | 0 | 0 |
| Q8I0D4 | Q8I0D4_DROME RE20510p OS=Drosophila melanogaster OX=7227 GN=Dmel\CG9297 PE=1 SV=1 | 1 | 1 | 1 | 1 | 0 | 0 | 0 |
| Q8IP68 | Q8IP68_DROME Uncharacterized protein, isoform A OS=Drosophila melanogaster OX=7227 GN=BG:DS08249.4 PE=4 SV=1 | 1 | 1 | 1 | 1 | 0 | 0 | 0 |
| Q8IPS9 | Q8IPS9_DROME Uncharacterized protein, isoform B OS=Drosophila melanogaster OX=7227 GN=Dmel\CG32448 PE=4 SV=2 | 1 | 1 | 1 | 1 | 0 | 0 | 0 |
| Q8IQB7 | Q8IQB7_DROME MIP21654p OS=Drosophila melanogaster OX=7227 GN=BEST:GH25183 PE=1 SV=1 | 1 | 1 | 1 | 1 | 0 | 0 | 0 |
| Q8IRZ3 | Q8IRZ3_DROME GH16657p OS=Drosophila melanogaster OX=7227 GN=Sdic3 PE=2 SV=3 | 1 | 1 | 1 | 1 | 0 | 0 | 0 |
| Q8MMD0 | Q8MMD0_DROME RH04491p OS=Drosophila melanogaster OX=7227 GN=CG30423 PE=2 SV=3 | 1 | 1 | 1 | 1 | 0 | 0 | 0 |
| Q8MSC8 | Q8MSC8_DROME LD14533p OS=Drosophila melanogaster OX=7227 GN=Dmel\CG2211 PE=2 SV=1 | 1 | 1 | 1 | 1 | 0 | 0 | 0 |
| Q8MSE4 | Q8MSE4_DROME GM30016p2 OS=Drosophila melanogaster OX=7227 GN=PIG-P PE=2 SV=1 | 1 | 1 | 1 | 1 | 0 | 0 | 0 |
| Q8MZ03 | Q8MZ03_DROME RE30473p OS=Drosophila melanogaster OX=7227 GN=BcDNA:RE30473 PE=2 SV=1 | 1 | 1 | 1 | 1 | 0 | 0 | 0 |
| Q8SWV5 | Q8SWV5_DROME LD47995p OS=Drosophila melanogaster OX=7227 GN=CG10444 PE=1 SV=1 | 1 | 1 | 1 | 1 | 0 | 0 | 0 |
| Q8SYC3 | Q8SYC3_DROME RE68569p OS=Drosophila melanogaster OX=7227 GN=BcDNA:RE68569 PE=2 SV=1 | 1 | 1 | 1 | 1 | 0 | 0 | 0 |
| Q8SYD9 | Q8SYD9_DROME Endophilin B, isoform A OS=Drosophila melanogaster OX=7227 GN=EndoB PE=1 SV=1 | 1 | 1 | 1 | 1 | 0 | 0 | 0 |
| Q8SZ06 | Q8SZ06_DROME RE24541p OS=Drosophila melanogaster OX=7227 GN=Dmel\CG12567 PE=1 SV=1 | 1 | 1 | 1 | 1 | 0 | 0 | 0 |
| Q8T043 | Q8T043_DROME CG3409, isoform A OS=Drosophila melanogaster OX=7227 GN=chk PE=2 SV=1 | 1 | 1 | 1 | 1 | 0 | 0 | 0 |
| Q8T4C9 | Q8T4C9_DROME AT03632p OS=Drosophila melanogaster OX=7227 GN=BcDNA:AT03632 PE=1 SV=1 | 1 | 1 | 1 | 1 | 0 | 0 | 0 |
| Q95RS6 | Q95RS6_DROME IP12536p OS=Drosophila melanogaster OX=7227 GN=BcDNA:LD13269 PE=2 SV=1 | 1 | 1 | 1 | 1 | 0 | 0 | 0 |
| Q95RV3 | Q95RV3_DROME LD09786p OS=Drosophila melanogaster OX=7227 GN=Lhr PE=2 SV=1 | 1 | 1 | 1 | 1 | 0 | 0 | 0 |
| Q95T37 | Q95T37_DROME GH26622p OS=Drosophila melanogaster OX=7227 GN=Nmda1 PE=2 SV=1 | 1 | 1 | 1 | 1 | 0 | 0 | 0 |
| Q961J8 | Q961J8_DROME GH20109p OS=Drosophila melanogaster OX=7227 GN=CT14378 PE=1 SV=1 | 1 | 1 | 1 | 1 | 0 | 0 | 0 |
| Q9V9A9 | Q9V9A9_DROME L(2)01289 long form OS=Drosophila melanogaster OX=7227 GN=l(2)01289 PE=1 SV=2 | 1 | 1 | 1 | 1 | 0 | 0 | 0 |
| Q9VA81 | Q9VA81_DROME GEO10172p1 OS=Drosophila melanogaster OX=7227 GN=mRpS18C PE=2 SV=2 | 1 | 1 | 1 | 1 | 0 | 0 | 0 |
| Q9VB75 | Q9VB75_DROME GH08870p OS=Drosophila melanogaster OX=7227 GN=gb PE=1 SV=1 | 1 | 1 | 1 | 1 | 0 | 0 | 0 |
| Q9VBY7 | Q9VBY7_DROME Protein lin-7 homolog OS=Drosophila melanogaster OX=7227 GN=veli PE=1 SV=1 | 1 | 1 | 1 | 1 | 0 | 0 | 0 |
| Q9VCZ0 | Q9VCZ0_DROME Elongation of very long chain fatty acids protein OS=Drosophila melanogaster OX=7227 GN=Dmel\CG5326 PE=2 SV=1 | 1 | 1 | 1 | 1 | 0 | 0 | 0 |
| Q9VDV5 | Q9VDV5_DROME FI18650p1 OS=Drosophila melanogaster OX=7227 GN=CG6231-RD PE=2 SV=2 | 1 | 1 | 1 | 1 | 0 | 0 | 0 |
| Q9VDX6 | Q9VDX6_DROME Uncharacterized protein OS=Drosophila melanogaster OX=7227 GN=CG18493 PE=4 SV=2 | 1 | 1 | 1 | 1 | 0 | 0 | 0 |
| Q9VE24 | Q9VE24_DROME RE63280p OS=Drosophila melanogaster OX=7227 GN=Dmel\CG14291 PE=1 SV=1 | 1 | 1 | 1 | 1 | 0 | 0 | 0 |
| Q9VEX2 | Q9VEX2_DROME Gilgamesh, isoform A OS=Drosophila melanogaster OX=7227 GN=gish PE=1 SV=1 | 1 | 1 | 1 | 1 | 0 | 0 | 0 |
| Q9VFI3 | Q9VFI3_DROME GEO08281p1 OS=Drosophila melanogaster OX=7227 GN=Dmel\CG8066 PE=1 SV=1 | 1 | 1 | 1 | 1 | 0 | 0 | 0 |
| Q9VFW8 | Q9VFW8_DROME Aminopeptidase OS=Drosophila melanogaster OX=7227 GN=CG8774 PE=1 SV=2 | 1 | 1 | 1 | 1 | 0 | 0 | 0 |
| Q9VG45 | Q9VG45_DROME Dpp target protein OS=Drosophila melanogaster OX=7227 GN=Dtg PE=2 SV=1 | 1 | 1 | 1 | 1 | 0 | 0 | 0 |
| Q9VGN7 | Q9VGN7_DROME AT25352p OS=Drosophila melanogaster OX=7227 GN=Exd2 PE=2 SV=2 | 1 | 1 | 1 | 1 | 0 | 0 | 0 |
| Q9VGQ4 | Q9VGQ4_DROME GH25284p OS=Drosophila melanogaster OX=7227 GN=scpr-B PE=2 SV=2 | 1 | 1 | 1 | 1 | 0 | 0 | 0 |
| Q9VH62 | Q9VH62_DROME LD44003p OS=Drosophila melanogaster OX=7227 GN=Dmel\CG9467 PE=2 SV=1 | 1 | 1 | 1 | 1 | 0 | 0 | 0 |
| Q9VHN0 | Q9VHN0_DROME GEO07735p1 OS=Drosophila melanogaster OX=7227 GN=TMEM216 PE=2 SV=1 | 1 | 1 | 1 | 1 | 0 | 0 | 0 |
| Q9VHQ4 | Q9VHQ4_DROME RE08932p OS=Drosophila melanogaster OX=7227 GN=CG11740 PE=2 SV=2 | 1 | 1 | 1 | 1 | 0 | 0 | 0 |
| Q9VHV3 | Q9VHV3_DROME FI02011p OS=Drosophila melanogaster OX=7227 GN=CG7800-RA PE=1 SV=2 | 1 | 1 | 1 | 1 | 0 | 0 | 0 |
| Q9VHV9 | Q9VHV9_DROME Uncharacterized protein OS=Drosophila melanogaster OX=7227 GN=CG7900 PE=4 SV=3 | 1 | 1 | 1 | 1 | 0 | 0 | 0 |
| Q9VI84 | Q9VI84_DROME LD17053p OS=Drosophila melanogaster OX=7227 GN=Dmel\CG1227 PE=2 SV=1 | 1 | 1 | 1 | 1 | 0 | 0 | 0 |
| Q9VID1 | Q9VID1_DROME Tetraspanin OS=Drosophila melanogaster OX=7227 GN=Tsp39D PE=2 SV=3 | 1 | 1 | 1 | 1 | 0 | 0 | 0 |
| Q9VJ63 | Q9VJ63_DROME FI20169p1 OS=Drosophila melanogaster OX=7227 GN=CG10338-RA PE=2 SV=2 | 1 | 1 | 1 | 1 | 0 | 0 | 0 |
| Q9VK84 | Q9VK84_DROME Solute carrier organic anion transporter family member OS=Drosophila melanogaster OX=7227 GN=Oatp33Ea PE=2 SV=3 | 1 | 1 | 1 | 1 | 0 | 0 | 0 |
| Q9VKN8 | Q9VKN8_DROME Palmitoyltransferase OS=Drosophila melanogaster OX=7227 GN=Dnz1 PE=2 SV=1 | 1 | 1 | 1 | 1 | 0 | 0 | 0 |
| Q9VKP2 | Q9VKP2_DROME GH26960p OS=Drosophila melanogaster OX=7227 GN=Porin2 PE=2 SV=1 | 1 | 1 | 1 | 1 | 0 | 0 | 0 |

| UniProt ID | Protein description | # PSMs MVB spectra | # of unique peptides MVB | # of PSMs lysate | # of unique peptides lysate | log2(PSM MVB) | log2(PSM lysate) | FC (log2 of MVB PSM-log2 of lysate PSM) |
|---|---|---|---|---|---|---|---|---|
| Q9VLI0 | Q9VLI0_DROME Alpha-mannosidase OS=Drosophila melanogaster OX=7227 GN=LManV PE=1 SV=1 | 1 | 1 | 1 | 1 | 0 | 0 | 0 |
| Q9VMW1 | Q9VMW1_DROME Reticulon-like protein OS=Drosophila melanogaster OX=7227 GN=Rtnl1 PE=1 SV=2 | 1 | 1 | 1 | 1 | 0 | 0 | 0 |
| Q9VPC1 | Q9VPC1_DROME FI07223p OS=Drosophila melanogaster OX=7227 GN=CG4042-RA PE=1 SV=1 | 1 | 1 | 1 | 1 | 0 | 0 | 0 |
| Q9VQ88 | Q9VQ88_DROME Uncharacterized protein OS=Drosophila melanogaster OX=7227 GN=CG15386 PE=4 SV=1 | 1 | 1 | 1 | 1 | 0 | 0 | 0 |
| Q9VQI6 | Q9VQI6_DROME LD25952p OS=Drosophila melanogaster OX=7227 GN=Dmel\CG8814 PE=2 SV=1 | 1 | 1 | 1 | 1 | 0 | 0 | 0 |
| Q9VQL3 | Q9VQL3_DROME Cornichon related OS=Drosophila melanogaster OX=7227 GN=cnir PE=4 SV=1 | 1 | 1 | 1 | 1 | 0 | 0 | 0 |
| Q9VQR0 | Q9VQR0_DROME FI04421p OS=Drosophila melanogaster OX=7227 GN=CG3246-RA PE=1 SV=1 | 1 | 1 | 1 | 1 | 0 | 0 | 0 |
| Q9VQT7 | Q9VQT7_DROME RH15675p OS=Drosophila melanogaster OX=7227 GN=CG16713-RA PE=1 SV=1 | 1 | 1 | 1 | 1 | 0 | 0 | 0 |
| Q9VRM3 | Q9VRM3_DROME Sinuous OS=Drosophila melanogaster OX=7227 GN=sinu PE=1 SV=3 | 1 | 1 | 1 | 1 | 0 | 0 | 0 |
| Q9VRS6 | Q9VRS6_DROME Jonah 65Aiv OS=Drosophila melanogaster OX=7227 GN=Jon65Aiv PE=1 SV=1 | 1 | 1 | 1 | 1 | 0 | 0 | 0 |
| Q9VS80 | Q9VS80_DROME FI17864p1 OS=Drosophila melanogaster OX=7227 GN=CG8539-RA PE=2 SV=1 | 1 | 1 | 1 | 1 | 0 | 0 | 0 |
| Q9VSF0 | Q9VSF0_DROME Autophagy-related 18a, isoform A OS=Drosophila melanogaster OX=7227 GN=Atg18a PE=1 SV=2 | 1 | 1 | 1 | 1 | 0 | 0 | 0 |
| Q9VST8 | Q9VST8_DROME LD38503p OS=Drosophila melanogaster OX=7227 GN=Dmel\CG4942 PE=2 SV=1 | 1 | 1 | 1 | 1 | 0 | 0 | 0 |
| Q9VTT0 | Q9VTT0_DROME Semaphorin 5c, isoform A OS=Drosophila melanogaster OX=7227 GN=Sema5c PE=4 SV=3 | 1 | 1 | 1 | 1 | 0 | 0 | 0 |
| Q9VTZ7 | Q9VTZ7_DROME LD47970p OS=Drosophila melanogaster OX=7227 GN=Dmel\CG4069 PE=1 SV=1 | 1 | 1 | 1 | 1 | 0 | 0 | 0 |
| Q9VUH4 | Q9VUH4_DROME FI02046p OS=Drosophila melanogaster OX=7227 GN=Dmel\CG9384 PE=2 SV=2 | 1 | 1 | 1 | 1 | 0 | 0 | 0 |
| Q9VWG0 | Q9VWG0_DROME GH21048p OS=Drosophila melanogaster OX=7227 GN=Shawn PE=2 SV=2 | 1 | 1 | 1 | 1 | 0 | 0 | 0 |
| Q9VX14 | Q9VX14_DROME RH64870p OS=Drosophila melanogaster OX=7227 GN=Ucp4A PE=1 SV=1 | 1 | 1 | 1 | 1 | 0 | 0 | 0 |
| Q9VX17 | Q9VX17_DROME LD36024p OS=Drosophila melanogaster OX=7227 GN=Dmel\CG6398 PE=2 SV=1 | 1 | 1 | 1 | 1 | 0 | 0 | 0 |
| Q9VXY5 | Q9VXY5_DROME LP17418p OS=Drosophila melanogaster OX=7227 GN=CG9065-RA PE=1 SV=1 | 1 | 1 | 1 | 1 | 0 | 0 | 0 |
| Q9VY05 | Q9VY05_DROME GH11762p OS=Drosophila melanogaster OX=7227 GN=CT26924 PE=1 SV=1 | 1 | 1 | 1 | 1 | 0 | 0 | 0 |
| Q9VY92 | Q9VY92_DROME GEO07753p1 OS=Drosophila melanogaster OX=7227 GN=2 PE=1 SV=1 | 1 | 1 | 1 | 1 | 0 | 0 | 0 |
| Q9VZ01 | Q9VZ01_DROME RH44935p OS=Drosophila melanogaster OX=7227 GN=BcDNA:RH44935 PE=1 SV=1 | 1 | 1 | 1 | 1 | 0 | 0 | 0 |
| Q9VZ57 | Q9VZ57_DROME Purple acid phosphatase OS=Drosophila melanogaster OX=7227 GN=CG1637-RA PE=2 SV=1 | 1 | 1 | 1 | 1 | 0 | 0 | 0 |
| Q9VZE2 | Q9VZE2_DROME GEO04055p1 OS=Drosophila melanogaster OX=7227 GN=Dmel\CG15012 PE=1 SV=2 | 1 | 1 | 1 | 1 | 0 | 0 | 0 |
| Q9VZF3 | Q9VZF3_DROME RE09466p OS=Drosophila melanogaster OX=7227 GN=BcDNA:RE09466 PE=1 SV=2 | 1 | 1 | 1 | 1 | 0 | 0 | 0 |
| Q9VZP9 | Q9VZP9_DROME IP05517p OS=Drosophila melanogaster OX=7227 GN=Dmel\CG12012 PE=1 SV=2 | 1 | 1 | 1 | 1 | 0 | 0 | 0 |
| Q9W045 | Q9W045_DROME FI21255p1 OS=Drosophila melanogaster OX=7227 GN=BcDNA:SD08776 PE=2 SV=2 | 1 | 1 | 1 | 1 | 0 | 0 | 0 |
| Q9W060 | Q9W060_DROME Mauve OS=Drosophila melanogaster OX=7227 GN=mv PE=4 SV=2 | 1 | 1 | 1 | 1 | 0 | 0 | 0 |
| Q9W068 | Q9W068_DROME LP10272p OS=Drosophila melanogaster OX=7227 GN=Tmhs PE=2 SV=1 | 1 | 1 | 1 | 1 | 0 | 0 | 0 |
| Q9W0E9 | Q9W0E9_DROME Rhomboid-like protein OS=Drosophila melanogaster OX=7227 GN=stet PE=3 SV=2 | 1 | 1 | 1 | 1 | 0 | 0 | 0 |
| Q9W106 | Q9W106_DROME CG3589 OS=Drosophila melanogaster OX=7227 GN=CG3589 PE=2 SV=2 | 1 | 1 | 1 | 1 | 0 | 0 | 0 |
| Q9W251 | Q9W251_DROME Palmitoyltransferase OS=Drosophila melanogaster OX=7227 GN=CG10344 PE=3 SV=2 | 1 | 1 | 1 | 1 | 0 | 0 | 0 |
| Q9W259 | Q9W259_DROME FI23916p1 OS=Drosophila melanogaster OX=7227 GN=CG13506-RA PE=1 SV=2 | 1 | 1 | 1 | 1 | 0 | 0 | 0 |
| Q9XYM5 | Q9XYM5_DROME LD27861p OS=Drosophila melanogaster OX=7227 GN=Su(var)2-10 PE=1 SV=1 | 1 | 1 | 1 | 1 | 0 | 0 | 0 |
| Q9XZ15 | Q9XZ15_DROME LD17368p OS=Drosophila melanogaster OX=7227 GN=Osi6 PE=1 SV=2 | 1 | 1 | 1 | 1 | 0 | 0 | 0 |
| Q9Y128 | Q9Y128_DROME BcDNA.GH07688 OS=Drosophila melanogaster OX=7227 GN=cert PE=1 SV=1 | 1 | 1 | 1 | 1 | 0 | 0 | 0 |
| X2JCX4 | X2JCX4_DROME Uncharacterized protein, isoform A OS=Drosophila melanogaster OX=7227 GN=Dmel\CG45089 PE=4 SV=1 | 1 | 1 | 1 | 1 | 0 | 0 | 0 |
| X2JDV7 | X2JDV7_DROME Uncharacterized protein, isoform B OS=Drosophila melanogaster OX=7227 GN=CG5966 PE=3 SV=1 | 1 | 1 | 1 | 1 | 0 | 0 | 0 |
| X2JG15 | X2JG15_DROME Uncharacterized protein, isoform E OS=Drosophila melanogaster OX=7227 GN=CG34120 PE=4 SV=1 | 1 | 1 | 1 | 1 | 0 | 0 | 0 |
| X2JJK8 | X2JJK8_DROME Severe impairment of CI with lengthened youth, isoform B OS=Drosophila melanogaster OX=7227 GN=sicily PE=4 SV=1 | 1 | 1 | 1 | 1 | 0 | 0 | 0 |
